# CompCorona: A Web Portal for Comparative Analysis of the Host Transcriptome of PBMC and Lung SARS-CoV-2, SARS-CoV, and MERS-CoV

**DOI:** 10.1101/2023.01.21.524927

**Authors:** Rana Salihoğlu, Fatih Saraçoğlu, Mustafa Sibai, Talip Zengin, Başak Abak Masud, Onur Karasoy, Tuğba Önal-Süzek

**Affiliations:** Department of Bioinformatics, University of Würzburg, Würzburg, Germany; Department of Bioinformatics, Muğla Sıtkı Koçman University,Turkey; Josep Carreras Leukaemia Research Institute, Spain; Department of Molecular Biology and Genetics, Muğla Sıtkı Koçman University, Turkey

## Abstract

**Motivation:** Understanding the host response to SARS-CoV-2 infection is crucial for deciding on the correct treatment of this epidemic disease. Although several recent studies reported the comparative transcriptome analyses of the three coronaviridae (CoV) members; namely SARS-CoV, MERS-CoV, and SARS-CoV-2, there is yet to exist a web-tool to compare increasing number of host transcriptome response datasets against the pre-processed CoV member datasets. Therefore, we developed a web application called CompCorona, which allows users to compare their own transcriptome data of infected host cells with our pre-built datasets of the three epidemic CoVs, as well as perform functional enrichment and principal component analyses (PCA).

**Results:** Comparative analyses of the transcriptome profiles of the three CoVs revealed that numerous differentially regulated genes directly or indirectly related to several diseases (e.g., hypertension, male fertility, ALS, and epithelial dysfunction) are altered in response to CoV infections. Transcriptome similarities and differences between the host PBMC and lung tissue infected by SARS-CoV-2 are presented. Most of our findings are congruent with the clinical cases recorded in the literature. Hence, we anticipate that our results will significantly contribute to ongoing studies investigating the pre-and/or post-implications of SARS-CoV-2 infection. In addition, we implemented a user-friendly public website, CompCorona for biomedical researchers to compare users own CoV-infected host transcriptome data against the built-in CoV datasets and visualize their results via interactive PCA, UpSet and Pathway plots.

**Availability:** CompCorona is freely available on the web at http://compcorona.mu.edu.tr

## 1 Introduction

The Coronavirus disease (COVID-19) caused by the severe acute respiratory syndrome coronavirus 2 (SARS-CoV-2) attacking the respiratory system caused severe immune and inflammatory response similar to other coronaviruses like the Middle East respiratory syndrome coronavirus (MERS-CoV) and the severe acute respiratory syndrome coronavirus (SARS-CoV). Unlike the other seasonal endemic human coronaviruses (HCoVs) with mild respiratory responses COVID-19, MERS-CoV, and SARS-CoV spread to multiple countries. In 2003, SARS-CoV caused an outbreak in a dozen countries resulting in the deaths of around 800 people. In 2012, MERS-CoV outbreak has caused the deaths of more than 900 people. The last coronavirus outbreak in 2019, has resulted in more than 6.5 million death worldwide by the time this manuscript was written.

Although the number of cases and deaths caused by the recent pandemic has recently decreased compared to the beginning of 2022 due to the increasing amount of vaccine applications (∼ 13 billion doses; COVID-19 Dashboard, 2022), the risk of infection remains with the ever-increasing number of variants of SARS-CoV-2 (e.g., Delta, Omicron; Kupferschmidt and Wadman, 2021; He et al., 2021). In order to more effectively understand the underlying reasons of aberrant host response to SARS-CoV-2, particularly in light of continually evolving variants, the transcriptomes of three main types of human coronaviruses have been comparatively studied (i.e., SARS-CoV, MERS-CoV, and SARS-CoV-2, Daamen et al., 2021; Jha et al., 2021; Krishnamoorthy et al., 2021; Alsamman and Zayed, 2022). The differential gene expression (DEG) analyses of these studies elucidate the severe lung damage on the molecular level, as well as the more frequent cardiovascular complications reflected by changes in several genes of SARS-CoV-2 (e.g., TNF, IL32, CXCL1-3, FOXO1, and TFPI2) compared to the seasonal endemic HCoVs (Jha et al., 2021). In addition, the numerous common pathways among the three HCoVs have been found using both transcriptome and enrichment (KEGG and GO) analyses and linked with the implications (e.g., neurological, mitochondrial dysfunction, and anosmia and olfactory dysfunction) and drug repurposing (e.g., Deferoxamine, Verapamil, and Colchicine) for SARS-CoV-2 infection (Krishnamoorthy et al., 2021). However, further studies are still needed for comparative analyses of host response to HCoV infection to decise faster treatment options to treat the severely affected patients. To that end, several efforts have been made, such as establishing some recent COVID-19 databases (e.g., COVID-19db, Zhang et al., 2022; CoV-RDB, Tzou et al., 2022).

In this study, we compared the transcriptome profiles of host cells infected with SARS-CoV, MERS-CoV, and SARS-CoV-2 to understand the host response to SARS-CoV-2 infection. We performed enrichment analyses using KEGG and GO to identify the most critical pathways for each coronavirus. Our analysis of the up-and down-regulated genes revealed that SARS-CoV-2 infection might be linked to various diseases. We also created a web page to share the results of our comparative analysis of the differentially expressed genes across the SARS-CoV, MERS-CoV, and lung and blood SARS-CoV-2 datasets. Users can upload their own data and compare it to these datasets using a Venn diagram and interactively visualize the relevant KEGG pathways or plot the Principal Component Analysis (PCA) of all datasets through a user-friendly web interface.

## 2 Methods

### 2.1 Gene Expression Omnibus Datasets for RNA Sequencing Analysis

The human transcriptome data infected by SARS-CoV-2, MERS-CoV, and SARS-CoV were downloaded from NCBI-GEO (https://www.ncbi.nlm.nih.gov/geo/). We selected GSE147507 (postmortem lung tissue, A549 lung cell line) for SARS-CoV-2, GSE139516 (lung adenocarcinoma, Calu-3), and GSE56192 for MERS-CoV (MRC5 cells), GSE56192 for SARS-CoV (MRC5 cells) for re-analysis. Accessions of CRR119890, CRR125445, CRR125446, CRR119891, CRR119892, and CRR119893 were downloaded from Genome Sequence Archive (https://bigd.big.ac.cn/gsa) for the SARS-CoV-2 peripheral blood mononuclear cells (PBMC) (Supplementary 1). The collected RNA sequencing datasets were analyzed using the methods following the pipeline of the updated version of Griffith et al. (2015).

### 2.2 RNA-Seq Datasets Differential Gene Expression Analysis

The raw data quality control (QC) was carried out using FastQC and multiple QC. The low-quality reads and adapters were trimmed using the FLEXBAR software (Figure S1) that ensures accurate adapter trimming of sequence labels based on exact overlap sequence alignment on some data (Dodt et al., 2012). The GRCh38/hg38, a compilation of the human genome published in December 2013, was selected as a reference genome for indexing in RNA-Seq analysis. HISAT2 was used to map next-generation sequencing reads with a population of human genomes and a single reference genome (Kim et al., 2015). The StringTie tool was used to analyze SAM/BAM files generated by HISAT2 to produce accurate gene reconstructions and estimate expression levels. These data were then mapped to the human reference genome GRCh38 using HISAT2 (Figure S1). The Ballgown R package was used to perform statistical analysis on the assembled transcriptomes, including analysis of differential gene expression (with a threshold of p-value < 0.05 and logFC > 1), visualization of transcript structures, and matching of assembled transcripts to annotation.

### 2.3 The Gene Ontology (GO) and Kyoto Encyclopedia of Genes and Genome (KEGG) Enrichment Analysis of DEGs

Transcriptomic analysis was conducted on SARS-CoV-2, MERS-CoV, and SARS-CoV infected human cells to compare the host gene response to infection. The clusterProfiler package (version 4.2.1) was used to perform functional analysis and determine the impact of differential expression of the human genes to each of these three similar infection types. The enrichGO and enrichKEGG functions in clusterProfiler were employed for GO and KEGG pathway enrichment analysis, respectively, with a p-value of less than 0.05. The results were then visualized using the pathview R package, and the most significant pathways for each coronavirus were identified and comparative tables and graphs were created.

To perform Gene Set Enrichment Analysis (GSEA), we used the clusterProfiler R package to annotate a list of gene names and their corresponding log fold change values. The genes were then sorted based on their log fold change. For the GSEA analysis, we set the number of permutations to 10000 and the minimum and maximum gene set sizes to 3 and 600, respectively. We also set the p-value cut-off at 0.05 to identify significant pathways. The GSEA analysis identified either activated or suppressed pathways, which was determined by the normalized enrichment score for each pathway.

### 2.4 Protein Network Based on Hub Gene Analysis

The interaction between DEGs was compiled from the STRING database (http://string.embl.de/) and visualized using Cytoscape. We used the NetworkAnalyzer plug-in to determine the characteristics of a small-world network by calculating the distribution of the network node degree, the distribution of the shortest path, average aggregation coefficient, and the proximity to the center. ClusterViz app with Molecular complex detection (MCODE), a plug-in in Cytoscape, was used to analyze PPI network modules, and MCODE scores > 2 were set as cut-off criteria with the default parameters (degree cut-off = 2, node score cut-off = 0.2, K-core = 3 and max depth = 100). CytoHubba, a plug-in in Cytoscape, was used to identify hub genes in the PPI network module with the MCC method (Figure S2 and Table S3).

### 2.5 Creating the Web platform

A web application was developed to present gene sets to users in an interactive way. Using this interface, users can upload their gene sets to the application as CSV or TSV formatted files and visualize their own and our pre-analyzed datasets on an interactive Venn diagram and PCA plot. React, a JavaScript library, is utilized to increase the interactivity of the interface. Uploaded files are streamed and not saved on the server side in any way, and only the file name and content are received and processed by the user.

For KEGG pathway display, firstly, the pathways of all Homo sapiens (hsa) genes were collected with the help of the BioConductor package in R. This collected data was transferred to the MySQL database. Secondly, a REST service was developed using the Python-based Flask framework to provide a connection to the database. The data obtained from the client request sent to this service (the gene list and p-value value are sent as parameters) are converted into a network graph by the Highchart.js library. Users can interactively click on the diagram’s genes or pathway information by connecting to their NCBI info pages

## 3 Results

In this study, we focused on both the genes that are up-regulated and downregulated during SARS-CoV-2, MERS-CoV and SARS-CoV infections. Secondly, we investigated the similarities and differences in human peripheral blood mononuclear cells and lung tissue transcriptome of SARS-CoV-2 infection. We used the MCC method of the cytoHubba plug-in in Cytoscape to identify 20 hub genes for SARS-CoV-2 disease, including ISG15, IFIT1, STAT1, MX1, IRF7, IRF9, IFIT3, OAS1, OAS3, and IFI35 (as shown in Figure S2. C). Combining the results from the Molecular Complex Detection (MCODE) plug-in with CytoHubba, we identified 20 hub genes (as shown in Figures S2. B and C).

### 3.1 Changes in transcriptional features and functionalities of virus-host interaction in lung SARS-CoV-2 infection

After the transcriptome analysis of the lung SARS-CoV-2 infection (GSE147507) dataset, we identified 615 differentially expressed genes (DEGs) with a p-value of less than 0.05 and a logFC of greater than 0. Of these DEGs, 335 were up-regulated, and 280 were down-regulated. Some of the most highly up-regulated genes included ISG15, MX1, IFI6, IFI27, IFIT1, IRF7, and PARP9, which are involved in the immune response to viral infections and the inhibition of viral replication. Some of the most highly down-regulated genes included ANG, NIPA1, METTL21A, GNB3, SURF4, SENP2, PGPEP1, ARL6, CRTC1, YTHDC2, and VEGFA, which are involved in vascular permeability, endothelian dysfunction, magnesium transport, methylation, and protein synthesis.

The up-regulated genes among SARS-CoV-2 infected DEGs were analyzed using GO and KEGG enrichment analysis, with a p-value of less than 0.05. A dot-plot was used to illustrate the relationships and differences between the pathways, with the dot size and color representing the gene ratio of the particular pathway and the significance level, respectively. The GO enrichment analysis identified 150 GO terms, including 115 biological processes, 8 cellular components, and 27 molecular function terms. The most important GO terms were related to the regulation of IP-10 production, defense response to symbionts and viruses, response to type I interferons and viruses, regulation of viral processes and life cycles, cellular response to type I interferons, type I interferons, cytokine-mediated and MDA-5 signaling pathways, ISGF3 complex, CXCR chemokine receptor binding, and ISG15-protein conjugation (as shown in Figure S6 A).

The down-regulated genes in the lung SARS-CoV-2 infection dataset were found to be associated with several GO terms, including the structural constituent of Ribosome, cytosolic Ribosome, cytoplasmic translation, angiogenin-PRI complex, vacuolar and lysosomal lumen, and mitochondrial and inner membrane. These associations were identified based on the p-value threshold during the Gene Ontology enrichment analysis.

In the host response to SARS-CoV-2 infection, the up-regulated DEGs (differentially expressed genes) were mainly enriched in pathways related to inflammation, such as the type I interferon signaling and Coronavirus disease pathway. Other pathways enriched in SARS-CoV-2 infection include IL-17, TNF, and NF-kappa B signaling pathways related to immunity, NOD-like receptors, Measles, Influenza A, viral carcinogenesis, Hepatitis C, Kaposi sarcoma-associated herpesvirus, and Epstein-Barr virus infection pathways. The GO (Gene Ontology) enrichment analysis of these DEGs emphasized the GO terms of the interferon response, response to the virus, viral process, and cytokine-mediated. Other GO terms associated with SARS-CoV-2 included catalytic activity, acting on RNA, cytokine and biotic stimulus regulation, cytoplasmic MDA-5 pattern, interferon-alpha, and antiviral response.

An analysis of the SARS-CoV-2 virus and its effects on the body identified 24 significantly affected pathways. Of these pathways, 19 were found to be activated and 5 were suppressed. The suppressed pathways included the ribosome, lysosome, alcoholism, focal adhesion, and Ras signaling pathways. In contrast, the pathways that were activated had Legionellosis, chemokine, viral protein interaction with cytokine and cytokine receptor, NOD and RIG-I-like receptors, Kaposi sarcoma-associated herpesvirus infection, herpes simplex virus 1, human papillomavirus and Epstein-Barr virus infections, coronavirus disease (COVID-19), influenza A, measles, and hepatitis C.

In comparing lung SARS-CoV-2 with blood SARS-CoV-2, MERS-CoV, and SARS-CoV, 345 genes were identified as being differentially expressed in lung SARS-CoV-2. The genes IRF7, FXN, HERC6, TNK2, and RTEL1 were highly expressed, while GNB3, SURF4, SENP2, ANG, PIK3R4, and MFSD8 were found to be down-regulated.

### 3.2 Comparing Gene Expression Analysis of blood and lung of SARS-CoV-2

A total of 1703 different DEGs expressed genes were identified between blood samples from healthy controls and SARS-CoV-2 patients. It was observed that 903 genes were expressed at higher levels in blood SARS-CoV-2 patients (Figure S5), while the expression of 800 genes was decreased (Figure S3 and Supplementary Table 1). Gene expression similarities and differences between the PBMC and lung tissue of SARS-CoV-2 were determined, and the 83 DEGs were shared between lung tissue and PBMC (Figure S4. B). The heatmap of common genes revealed that the most frequently downregulated genes in PBMC were AHNAK, which are associated with peripheral NK cell maturity, and the neurotransmission-associated genes, such as VAMP2 and CXCL8 including chemokine activity and interleukin-8 receptor binding.

IFI27 and OAS1 genes, among the most critical genes up-regulated in blood and lung SARS-CoV-2 infection, are the defense response against the other organism, including the cellular protein metabolic process; and are involved in processes such as the apoptotic signaling pathway. The LAP3 gene contains peptidase activity and aminopeptidase, and the SULT1A1 gene includes sulfotransferase and flavonol 3-sulfotransferase activity. Moreover, another critical up-regulated FN1 gene is involved in embryogenesis, wound healing, blood coagulation, host defense, and cell adhesion and migration processes, including metastasis and fibrosis.

### 3.3 Inflammatory mediators in SARS-CoV-2

The up-regulation of several chemokines and cytokines in SARS-CoV-2 infection has been observed, including CXCL1, CXCL2, CXCL3, CXCL5, CXCL8, TNFSF12, CCL20, and IL4. These proteins play roles in the inflammatory response and hematopoietic regulation, and IL18, a member of the IL-1 family, is also up-regulated. The expression of these genes has been observed in activated monocytes and neutrophils and at sites of inflammation. In vitro, CXCL2 has been shown to suppress the proliferation of hematopoietic progenitor cells.

Immunoglobulin transcripts (IGHG1, IGKC, IGHA1, VSIG4, IGHG3, and IGHG4), leukocyte immunoglobulin-like receptor (LILRB4), fibronectin (FN1), complement factors (C1QA, C1QB, and C1QC), RAB13, IFI27, CES1, WDR13, inflammation and especially the upstream of the immune response (PBCLEC12A) regulation was observed in blood SARS-CoV-2 samples.

The inflammatory cytokines (CXCL8) and CLC involved in the apoptotic process, such as carbohydrate binding and cysteine-type endopeptidase activity, are among the most down-regulated genes (HLA-F, HLA-E, and HLA-DQA1). AHNAK, critical in the calcium entry and exit required during the immune response, is a downregulated gene. Interestingly, the expression of DEFA1B (Defensin Alpha 1B) and DEFA3 genes, which play a role in pathways related to defensins and the innate immune system, is relatively low.

### 3.4 Comparing the GO and KEGG enrichment analysis results of PBMC and lung SARS-CoV-2 infection

Up/down-regulated DEGs were used to detect lung and blood SARS-CoV-2 pathogenesis by GO assays. DEGs obtained from lung and blood SARS-CoV-2 infected samples were classified as up-regulated genes with log2FC values >0 after p-value <0.05 filtering, and genes with log2FC <0 were classified as down-regulated, and GO enrichment analysis was applied separately. Genes up-regulated in blood and lung SARS-CoV-2 are related to biological processes, mainly interferon response, immune response, and cytokine mediation. Down-regulated genes are primarily associated with the cellular component lysosomal and vacuolar lumen, mitochondrial, and organelle inner membrane. It was observed in the GO enrichment analysis that the function of natural killer cells, which play an essential role in controlling viral infections, is down-regulated in blood SARS-CoV-2 infection.

In a study of blood samples from patients infected with SARS-CoV-2, the Gene Ontology (GO) enrichment analysis identified 346 terms divided into molecular function, biological processes, and cellular components. Of these, 30 were related to molecular function, 184 to biological processes, and 132 to cellular components. The GO enrichment analysis was also applied to the up-regulated and down-regulated genes identified in the study.

Up-regulated genes in blood SARS-CoV-2, which are most significant as a result of GO enrichment analysis, mitochondrial protein-containing complex, mitochondrial inner membrane, inner organelle membrane, aerobic respiration, generation of precursor metabolites and energy, inner mitochondrial membrane protein complex, ATP metabolic process, cellular respiration, immunoglobulin complex, mitochondrial matrix, oxidative phosphorylation, and mitochondrial respiratory were highlighted.

The down-regulated genes in blood samples from people infected with SARS-CoV-2 were involved in various processes related to histone modification, chromatin organization, mRNA processing, and protein acetylation. These processes included histone acetylation, internal peptidyl-lysine acetylation, internal protein amino acid acetylation, regulation of histone modification, chromatin binding, transcription coregulator activity, and regulation of the mRNA metabolic process.

The natural killer cell-mediated cytotoxicity, Th1, Th2, and Th17 cell differentiation, graft-versus-host, inflammatory bowel disease, antigen processing, presentation, RNA degradation, and FoxO signaling pathway are located in the KEGG analysis performed with genes down-regulated in blood SARS-CoV-2 infection (pathways with a p-value of < 0.001). In addition, the most significant paths expressed by up-regulated genes are oxidative phosphorylation, Alzheimer’s disease, diabetic cardiomyopathy, Proteasome, chemical carcinogenesis, pathways of neurodegeneration, thermogenesis, and carbon metabolism.

In summary, the analysis of gene expression in SARS-CoV-2 infection in the lung and blood showed several pathways and gene functions that were either up-regulated or down-regulated. These included pathways related to the immune response, inflammation, and recognition of pathogen-associated molecular modes in the lung, as well as processes involved in histone modification, chromatin organization, mRNA processing, and protein modification in the blood. The analysis also identified specific genes, such as IRF7 and PPP2R2A, that may play essential roles in SARS-CoV-2 infection and could be potential targets for therapeutic intervention. Additionally, a web page was developed where the transcriptome analysis results can be compared with other data on SARS-CoV, MERS-CoV, and SARS-CoV-2 infection, and relevant pathways can be visualized in an interactive interface.

KEGG enrichment pathway analysis was performed to determine metabolic shifts and differentiated metabolic mechanisms in SARS-CoV-2 infected patients. Two different pathway analyses were performed for up-regulated and down-regulated genes according to DEGs obtained as a result of transcriptome analysis. There are differences in the pathway analysis results in lung and blood SARS-CoV-2 infected samples. Lung SARS-CoV-2 down-regulated genes cause changes in the lysosome and ribosome pathways. Up-regulated genes are Hepatitis C, COVID-19, NF-Kappa B, IL-17, TNF, RIG-I-like receptor, and AGE-RAGE signaling pathway are closely related. As a result of the enrichment analysis, increased activation of NF-KB was observed in the lung SARS-CoV-2 infection (Figure S7. A).

### 3.5 Other viral infections transcriptome analysis and GO/KEGG enrichment results

#### 3.5.1 SARS-CoV

We identified 3468 DEGs (Supplementary Table 1) by applying p value <0.05 and logFC >0 threshold in the SARS-CoV dataset (GSE56192), in which RNA-seq transcriptome analysis was applied. While 2090 genes were downregulated, 1378 genes were up-regulated. The top downregulated genes were IFI6, MX1, CXCL8, IFIT1, IFITM1, IFI27, SNORD3A, NPIPP1, ISG15, and AREG, whereas the top up-regulated genes were ACAD10, DMXL1, MIR22, TRIM13, DGLUCY, MATR3, SNORA18, PDIA3P2, CHFR, and SNORA8.

An analysis of the genes that are up-regulated in response to SARS-CoV infection found that they were associated with several processes, including the transition between different phases of the cell cycle, cell division, and the movement of substances between the nucleus and cytoplasm of a cell. In addition, the GO terms of down-regulated genes such as cytoplasmic translation, ribonucleoprotein complex biogenesis, apoptotic signaling pathway, and autophagy were expressed at high levels. In addition, the results of KEGG enrichment analysis based on the up-regulated DEGs were closely associated with nucleocytoplasmic transport, Coronavirus disease − COVID−19, ECM−receptor interaction, oxytocin, PI3K−Akt, and MAPK signaling pathway. Some immune-related pathways were enriched in SARS-CoV (Figure S6. C).

Based on the KEGG enrichment analysis results, the following pathways were activated during human cytomegalovirus infection: Endocytosis, spliceosome, human T-cell leukemia virus 1 infection, human immunodeficiency virus 1 infection, MAPK signaling, pathogenic Escherichia coli infection, and chemokine signaling pathway. In addition, the ErbB oncogene was up-regulated (as shown in Figure S7. C). The GSEA KEGG analysis on SARS-CoV infection identified 60 enriched pathways, with 16 pathways activated and 44 pathways suppressed after viral infection. Pathways related to the innate immune system, such as TNF, RIG-I-like receptor, and influenza A, were suppressed in SARS-CoV disease. In addition to these immune pathways, the following pathways were up-regulated: nucleocytoplasmic transport, dilated cardiomyopathy, cell cycle, calcium, taste transduction, Fanconi anemia, neuroactive ligand-receptor interaction, focal adhesion, Renin secretion, and ECM-receptor interaction.

#### 3.5.2 MERS-CoV

Transcriptome analysis of the MERS-CoV dataset (GSE56192) was performed and a total of 1073 DEGs was determined (Supplementary Table 1) by applying a p-value <0.05 and logFC >0 thresholds. As a result of the analysis, 641 genes were downregulated, while 432 genes were up-regulated (Figure S3). Top 15 up-regulated genes; ATF3, ZBTB38, AREG, PDE4B, ATF3, MEDAG, MMP10, SPIN1, RAPGEF6, SNORD14C, IL33, CSF3, OSBPL6, CXCL1, and NRG1. Top 10 downregulated genes; HCG20, TOMM6, ATG10, OAZ3, VDR, SNORA6, ASS1, EBPL, APOLD1, BBS1, and IFT22.

Tissue morphogenesis, cell−cell signaling by wnt, cholesterol biosynthetic process, and lung morphogenesis biological processes were the leading GO terms associated with up-regulated DEGs. That also pointed to the steroid biosynthesis KEGG pathway. In addition, down-regulated genes predominantly expressed the GO terms of cytoplasmic translation, rRNA metabolic process, respiratory electron transport chain, and Arp2/3 complex−mediated actin nucleation (Figure S6. D). The KEGG analysis resulted in the ribosome, Coronavirus disease, Salmonella infection, pathogenic Escherichia coli infection, and apoptosis pathways (Figure S7. D).

In the case of MERS-CoV, we found 32 enriched pathways (26 activated and 6 suppressed after viral infection). While ribosome, Coronavirus disease - COVID-19, drug metabolism - other enzymes, protein processing in the endoplasmic reticulum, spliceosome were activated, JAK-STAT, Longevity regulating, PI3K-Akt, ErbB, and Cytokine-cytokine receptor interaction were suppressed.

### 3.6 Transcriptional features shared between SARS-CoV-2 and SARS-CoV, and MERS-CoV infections

When the gene expression analysis results of MERS-CoV, SARS-CoV, and SARS-CoV-2 (lung and blood) virus infection datasets were compared, the shared genes were RPLP0P6 and CXCL1.

There were 60 common DEGs shared between MERS-CoV and SARS-CoV-2. Of these, 12 genes with LogFC >1, AREG, CXCL1, and TCIM are highly associated with cytokine activity, inflammatory response, apoptotic process, and NF-kappa B signaling pathway.

The 134 unique DEGs were detected that are common between SARS-CoV and SARS-CoV-2 but not found in MERS-CoV. While 80 were up-regulated in SARS-CoV-2, it was noteworthy that they were downregulated in SARS (Supplementary Table 1). Genes with LogFC value > 4, STIM1, PARP9, MRPL53, IFI27, IFIT1, ISG15, MX1, and IFI6 are closely related to apoptosis, cellular response to the virus, host-virus interaction, immunity, host-virus interaction/defense, mitochondrial translation, and type I interferon regulation/response pathways.

### 3.7 CompCorona webpage where CoVs datasets can be compared

A website has been created where the results of this study’s RNA-seq transcriptome analysis can be compared with user data (http://compcorona.mu.edu.tr and Fig. 1). CompCorona contains DEG data for MERS-CoV, SARS-CoV, SARS-CoV-2 infections in the lung and blood, and these can be compared to each other in a Venn diagram using the D3.js library (Fig. 1). CompCorona allows users to download the gene set of the intersection and provides an interactive interface to visualize relevant pathways. The website also uses principle component analysis to interactively identify common and different genes among SARS-CoV, MERS-CoV, SARS-CoV-2, and blood SARS-CoV-2 datasets and presents this information in an interactive 3D PCA plot using the Plotly library.

**Fig. 1.**
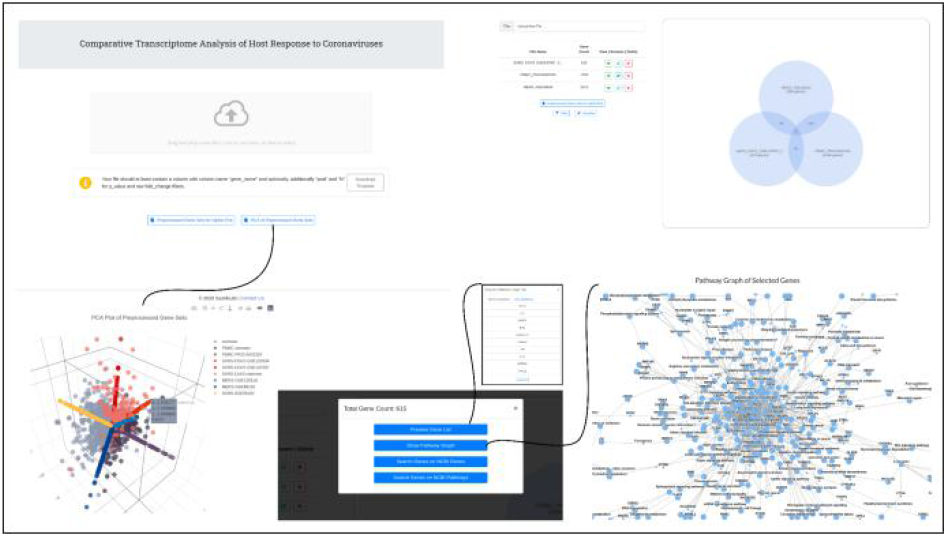
CompCorona website structure.

## 4 Discussion

### 4.1 The common pathways between SARS-CoV-2 and other HcoVs

The comparison of GSEA KEGG enrichment analysis of SARS-CoV and SARS-CoV-2 infection revealed 12 common pathways (Fig. 1). The Ribosome was suppressed for MERS-CoV, SARS-CoV, and SARS-CoV-2 (Krishnamoorthy et al., 2021, Zhang et al., 2022). Focal adhesion was active for SARS-CoV but suppressed for SARS-CoV-2 infection. Hence, these pathways may be essential in disease progression and virus replication. The remaining Coronavirus disease (Gugliandolo et al., 2021), Rheumatoid arthritis (Choy and Panayi., 2001; Alsamman and Zayed, 2020), TNF (Heller and Krönke, 1994), Epstein-Barr virus infection (Ascherio, and Munger, 2010), Influenza A (Ehrhardt, and Ludwig, 2009), Legionellosis (Newton et al., 2010), Hepatitis B (Caballero et al., 2018), Measles (Gerlier and Valentin, 2009), RIG-I-like receptor (Kawai, and Akira, 2008), Kaposi sarcoma-associated herpesvirus infection pathways (Lee et al., 2012) are suppressed in SARS-CoV, while SARS-CoV-2 was activated. According to a study, certain drugs currently used to treat COVID-19 can potentially induce the re-activation of Kaposi sarcoma-associated herpes virus (KSHV), a significant human oncogenic virus, through the manipulation of intracellular signaling pathways (Chen et al., 2021). It also shows that KSHV+ patients exposed or treated for SARS-CoV-2 infection may have an increased risk of developing virus-associated cancer even after fully recovering (Chen et al., 2021).

In our results, TNF signaling pathway was also suppressed in blood SARS-CoV-2. Complement and coagulation cascades and pertussis pathways were active in lung and blood SARS-CoV-2 infection datasets. It was observed that SARS-CoV and MERS-CoV infections shared Parkinson’s disease (Olanow and McNaught, 2006), Osteoclast differentiation (Takayanagi, 2010), and Spliceosome pathways. GSEA KEGG analysis showed that the osteoclast differentiation pathway was active in MERS-CoV but suppressed in SARS-CoV. Another finding of our research was the activation of the NOD-like receptor (NLR) signaling pathway in both SARS-CoV-2 and multiple sclerosis (MS). This suggests that the NLR signaling pathway may play a critical role in developing SARS-CoV-2-related MS (Qiu et al., 2022).

Epstein-Barr virus (EBV) and SARS-CoV-2 coinfection has been linked to increased fever and inflammation in infected patients (Chen et al., 2021). EBV can induce ACE2 expression when it enters the lytic replicative cycle in epithelial cells. A study using vesicular stomatitis virus (VSV) particles pseudotyped with the SARS-CoV-2 spike protein found that lytic EBV replication increases ACE2-dependent SARS-CoV-2 pseudovirus entry (Verma et al., 2021).

A high incidence of HSV-1 re-activation has been virologically and clinically observed in patients with severe SARS-CoV-2 pneumonia, especially in those treated with steroids (Franceschini et al., 2021).

Through our analysis of the GSEA/KEGG pathways, we identified pathways related to the immune response, inflammatory response, and recognition of pathogen-associated molecular modes that may be affected by SARS-CoV-2. These pathways included the IL-17 signaling pathway, NF-kappa B signaling pathway, NOD-like receptor signaling pathway, and TNF signaling pathway (as shown in Figure S8 A). The IL-17 signaling pathway plays a crucial role in the clearance of extracellular pathogens and has been linked to acute respiratory distress syndrome. IL-17 is a potent inflammatory cytokine that can stimulate the expression of chemokines and attract cells to parenchymal tissue (Khader and Cooper, 2008). Therefore, the activation of the IL-17 pathway may increase cytokine expression in patients with SARS-CoV-2.

The IL-17 signaling pathway plays an essential role in maintaining the homeostasis of tissues (Moseley et al., 2003). However, dysregulation of IL-17 production can lead to chronic inflammation, autoimmune disorders, and potentially even cancer (Sahu et al., 2021). In mice, reducing the levels of IL-17 significantly diminished the severity of chronic graft-versus-host disease induced by IL-1 (Part et al., 2018). Given these findings, targeting IL-17 could be an effective strategy for preventing acute respiratory distress syndrome (ARDS) in SARS-CoV-2 infection. Therefore, we believe a clinical trial of a drug targeting IL-17 could be a promising addition to the search for effective therapies for SARS-CoV-2 disease (Pacha et al., 2020).

### 4.2 Down- or up-regulated genes in SARS-CoV-2 and their possible implications

The comparative transcriptome analysis reveals down- and up-regulated genes that are notably altered after SARS-CoV-2 infection (Supplementary Table 1). Among these genes in the supplementary table 1, we selected those in the top 10 of both down- and up-regulated gene lists and investigated their possible implications for the disorders, which are the known consequences of SARS-CoV-2 infection.

One of our down-regulated genes, the GNB3 (guanine nucleotide-binding protein), includes three SNPs (single nucleotide polymorphisms) that significantly affect the blood pressure response to ACE inhibitors (Chaudhary et al., 2015; and references therein). This gene is well associated with hypertension (Benjafield et al., 1998; Yamamoto et al., 2004; Danoviz et al., 2006; Grove et al., 2007) and is congruent with the rise in blood pressure during the SARS-CoV-2 infection (e.g., Ran et al., 2020; Angeli et al., 2022; Laffin et al., 2022). The deficiency of SENP2 (sentrin/sumo-specific protease 2), which is another down-regulated gene in our analysis, has been related to early mortality in mice (Qi et al., 2014), mitochondrial dysfunction (Nan et al., 2022), embryonic heart and brain development in placenta (Maruyama et al., 2016), and SMA-like phenotype (Zhang et al., 2021). Despite the little relation between newborn health and SARS-CoV-2 infection (Kyle et al., 2020), we here would like to propound this anomaly that appeared in our analysis. The potential role of YTHDC2 (YTH domain-containing 2) as a mediator of m6A function in SARS-CoV-2 infection (Kamel et al., 2021) has been known in the literature. A deficiency in this gene can also be associated with spermatogenesis (Hsu et al., 2017) and lung cancer (Wang et al., 2021). Therefore, we here speculate that the down-regulation of this gene in our analysis can be linked with the male fertility problems that have been discussed as one of the SARS-CoV-2 implications (Vishvkarma and Rajender, 2020; Dutta and Sengupta, 2021; Haghpanah et al., 2021). We also found the METTL21A gene (a non-histone methyltransferase) as down-regulated, which is generally responsible for the protein-lysine N-methyltransferase activity (Nishizawa et al., 2014; Hamamoto et al., 2015; Jakobsson et al., 2016).

Angiogenin (ANG) and vascular endothelial growth factor A (VEGFA) genes, which are down-regulated in our analysis are both involved in vascular endothelial cell (EC) dysfunction have a relation with the Amyotrophic lateral sclerosis (ALS) neurodegenerative disease (e.g., Lambrechts et al., 2006; Galleria et al., 2008; Silva-Hucha et al., 2021), which is also evident in relation with the SARS-CoV-2 infection (e.g., Shi et al., 2021; Li et al., 2022; Lingor et al., 2022). In addition, the latter are proangiogenic cytokines that maintain tumor angiogenesis and preclude antitumor immunity (Schmittnaegel et al., 2017). The decrease in the surfeit locus protein 4 (SURF4) gene, known as a cargo receptor homolog, causes a decreased insulin secretion due to proinsulin retention in the endoplasmic reticulum and reduction of mature insulin in secretory granules (Saegusa et al., 2022). This role of the SURF4 gene is also suitable for studies investigating the relationship between insulin resistance and SARS-CoV-2 (Affinati et al., 2021; Merikangas et al., 2022). The NIPA1 (nonimprinted in Prader-Willi/Angelman syndrome 1) causes a neurodegenerative disease (hereditary spastic paraplegia, HSP type 6), including some phenotypic similarities with ALS (Amyotrophic lateral sclerosis; Rainier et al., 2003) that also has a link with SARS-CoV-2 infection (Li and Bedlack, 2021).

Among the genes that are unique in lung SARS-CoV-2 compared to SARS-CoV, MERS-CoV, and HCoV-2259E, and blood SARS-CoV-2, the top up-regulated ones are IRF7, HERC6, EXOG, ZNF566, POLR2J3, MMP17, MED17, GATAD2A, RTEL1, TRIM34, PPP2R2A, and SPIRE2.

HERC6 (Probable E3 ubiquitin-protein ligase 6) is a member of the HERC ubiquitin ligase family and exhibits antiviral activity induced by interferon (Paparisto et al., 2018). When HERC6 is overexpressed, it promotes the inflammatory response (Cao et al., 2022; Scott et al., 2022). It has been mentioned that this gene is overexpressed in SARS-CoV-2 infection patients (Li et al., 2021). Human EXOG (hEXOG) is a 5′-exonuclease crucial for mitochondrial DNA repair; The enzyme belongs to a family of nonspecific nucleases that includes the apoptotic endonuclease EndoG, and the accumulation of certain types of mitochondrial damage in its absence can directly trigger cell death (Van

Houten et al., 2016; Szymanski et al., 2017). The over-expression of EXOG in our study suggests that it may be necessary to treat COVID-19. The ubiquitous expression of the FAM228B gene, whose function has not yet been determined, in most tissues, including the brain, shows its functional importance. Changes in the expression of FAM228B are mentioned in depressed individuals who are treated with antidepressants and die from suicide (Fiori et al. 2021). Also, overlapping relationships have been identified between the severity of inflammation, brain structure, and function during acute COVID-19, with the severity of depression and post-traumatic distress in survivors (Benedetti et al., 2021; Yang et al., 2021). We believe that the over-expression of FAM228B in SARS-CoV-2 patients, as identified in our analysis, may be related to brain function.

In our analysis, the ZNF566 gene was found to be up-regulated. This gene plays a central role in regulating gene expression and in the process of cardiac regeneration, including the transition of endocardial and epicardial epithelial cells to mesenchymal cells. Atrial fibrillation (AF) and cardioembolic stroke are linked, and the ZNF566 gene is significantly correlated with novel biomarkers of AF-related stroke (Zou et al., 2019; Ekkert et al., 2021). There have been reports of recurrent cardioembolic stroke occurring after vaccination with the BNT162b2 (Pfizer) COVID-19 mRNA vaccine (Yoshida et al., 2022). COVID-19-associated cardiomyopathy increases the risk of cardioembolic stroke (Venketasubramanian et al., 2021). Another up-regulated gene, SPIRE2 (Spire Type Actin Nucleation Factor 2), has been linked to cardiovascular diseases (Wong and Shapiro, 2019; Tkacz et al., 2022). In addition, SPIRE2 expression is significantly reduced in the brain tissues of individuals with epilepsy (Hao et al., 2022). Over-expression of SPIRE2 has been observed in patients with SARS-CoV-2 infection. Therefore, it is interesting to study the potential effects of SARS-CoV-2 on the seizure rate in epilepsy patients (Kuroda, 2021). Further investigation of the impact of COVID-19 on epilepsy patients may also be important.

One of the essential findings in our study was the up-regulation of the POLR2J3 gene. This gene plays a role in RNA synthesis as a component of RNA polymerase II. The function of POLR2J3 has been linked to infertility (Taylor et al., 2017), which has been observed in patients with recurrent miscarriage (RM) (Ran et al., 2022). Another up-regulated gene in our analysis, MMP17 (Matrix Metallopeptidase 17), is responsible for the shedding of ACE2 (Kumar et al., 2021). MMP17, expressed by smooth muscle cells, is also necessary for repairing injuries in the intestinal epithelium caused by inflammation (Martin-Alonso et al., 2021). Over-expression of MMP17 has been associated with various types of cancer, including breast, ovarian, prostate, lung, liver, adrenal, cervix, gastric, colon, and oral mucosa, as well as with cardiovascular disease and arthritis (Grant et al., 1999; Wang et al., 2015; Yip et al., 2019; Xiao et al., 2022). Additionally, MMP17 may indirectly contribute to cartilage degradation in normal and pathological processes through its activation in ADAMTS-4 (Sohail et al., 2008).

The MED17 subunit is involved in the regulation of non-coding RNAs. A mutation that causes a frameshift in MED17 has been found to reduce the accumulation of miRNAs, indicating the importance of MED17 in miRNA biogenesis. The up-regulation of MED17 in our analysis suggests its potential role in the modulation of miRNA expression in viroid pathogenesis (Nozawa et al., 2017). MED17 has not yet been studied in the context of SARS-CoV-2, but it could be a valuable research topic for future studies. Over-expression of GATAD2A (GATA zinc finger domain containing 2A) has been shown to impair autophagy and promote H1N1 replication (Mo et al., 2021). Circ-GATAD2A has overexpressed in IAV H1N1 infection (Sajjad et al., 2021). GATAD2A has also been identified as a risk gene for schizophrenia in various tissues (Hoffmann and Spengler, 2019; Morris et al., 2019; Liu et al., 2020). Both COVID-19 and schizophrenia involve neuroinflammation, which may worsen the outcomes of both diseases when they occur together (Tendilla-Beltrán and Flores, 2021). GATAD2A has also been linked to cardiometabolic disease and type-2 diabetes due to the increased risk of metabolic syndrome associated with the use of atypical antipsychotic drugs (Ma et al., 2018; Merikangas et al., 2022). It is also related to total cholesterol, triglycerides (Liu et al., 2020), and tumorigenesis. Our study showed a close relationship between GATAD2A and thyroid cancer progression (Wang et al., 2017). Our literature review suggests that the thyroid gland and the entire hypothalamic-pituitary-thyroid axis may be vulnerable to damage from SARS-CoV-2 infection (Scappaticcio et al., 2021).

Pulmonary fibrosis is a fatal disease that leads to a progressive loss of respiratory function. RTEL1 is a DNA helicase that plays a role in DNA replication, genome stability, DNA repair, and telomere maintenance (Jenkins, 2020). RTEL1 has been identified as a gene associated with pulmonary fibrosis (Kropski and Loyd, 2015; Matto and Pillai, 2021). Higher expression of RTEL1 has been linked to important oncogenic pathways and metabolic changes in adrenocortical carcinoma (Yuan et al., 2022). The unresolved DNA damage response has been suggested to accelerate pulmonary aging and increase the risk of pulmonary fibrosis. It has recently been reported that SARS-CoV-2 can cause acute pulmonary epithelial senescence and subsequent fibrosis, although the exact mechanism is not yet understood (Hong et al., 2022). TRIM34, up-regulated in our study, has been shown to play a role in the restriction of HIV-1 infection and programmed cell death activated by IAV (Ohainle et al., 2019; Wang et al., 2022). HIV-1-induced immune dysfunction has also been linked to early SARS-CoV-2 clearance (Zhao et al., 2020). Further research on the role of TRIM34 in these processes could be beneficial.

PPP2R2A is a regulatory subunit that enhances Th1 and Th17 differentiation towards PP2A and may be a potential therapeutic target for conditions associated with Th1 and Th17 cell expansion (Pan et al., 2021). up-regulation of PPP2R2A expression, which can activate the phosphorylation of Akt, has also been observed in pancreatic adenocarcinoma (Wang et al., 2016). Further research on the relationship between PPP2R2A and SARS-CoV-2 could be beneficial. IRF7 is another gene that is highly expressed in SARS-CoV-2 infected patients, as observed through transcriptome analysis. Type I/III interferons, interferon regulatory factor 7 (IRF7), and interferon-stimulated genes (ISGs) are highly expressed in oropharyngeal cells of SARS-CoV-2 infected patients (Scagnolari et al., 2021; Hasan et al., 2021).

## 5. Conclusion

Although several comparative host transcriptome analysis studies of SARS, MERS and SARS-CoV2 have been published and several web portals facilitating analysis of the sequence data of the SARS,MERS and SARS-Cov2 viral sequences exist, there is no web portal enabling comparative analysis of host transcriptome data of these three CoVs. CompCorona enables visualization of the pre-processed CoV infected host transcriptome datasets by interactive UpSet, pathway and 3-dimensional PCA plots along with enrichment analysis. In the near future, we are planning to extend our web portal by adding more enrichment options and few more pre-processed (such as seasonal human coronavirus transcriptome) datasets to enhance the user experience of CompCorona web portal.

