## Supplementary material for "CompCorona: A Web Portal for Comparative Analysis of the Host Transcriptome of PBMC and Lung SARS-CoV-2, SARS-CoV, and MERS-CoV": Figures_1_9_Supplementary

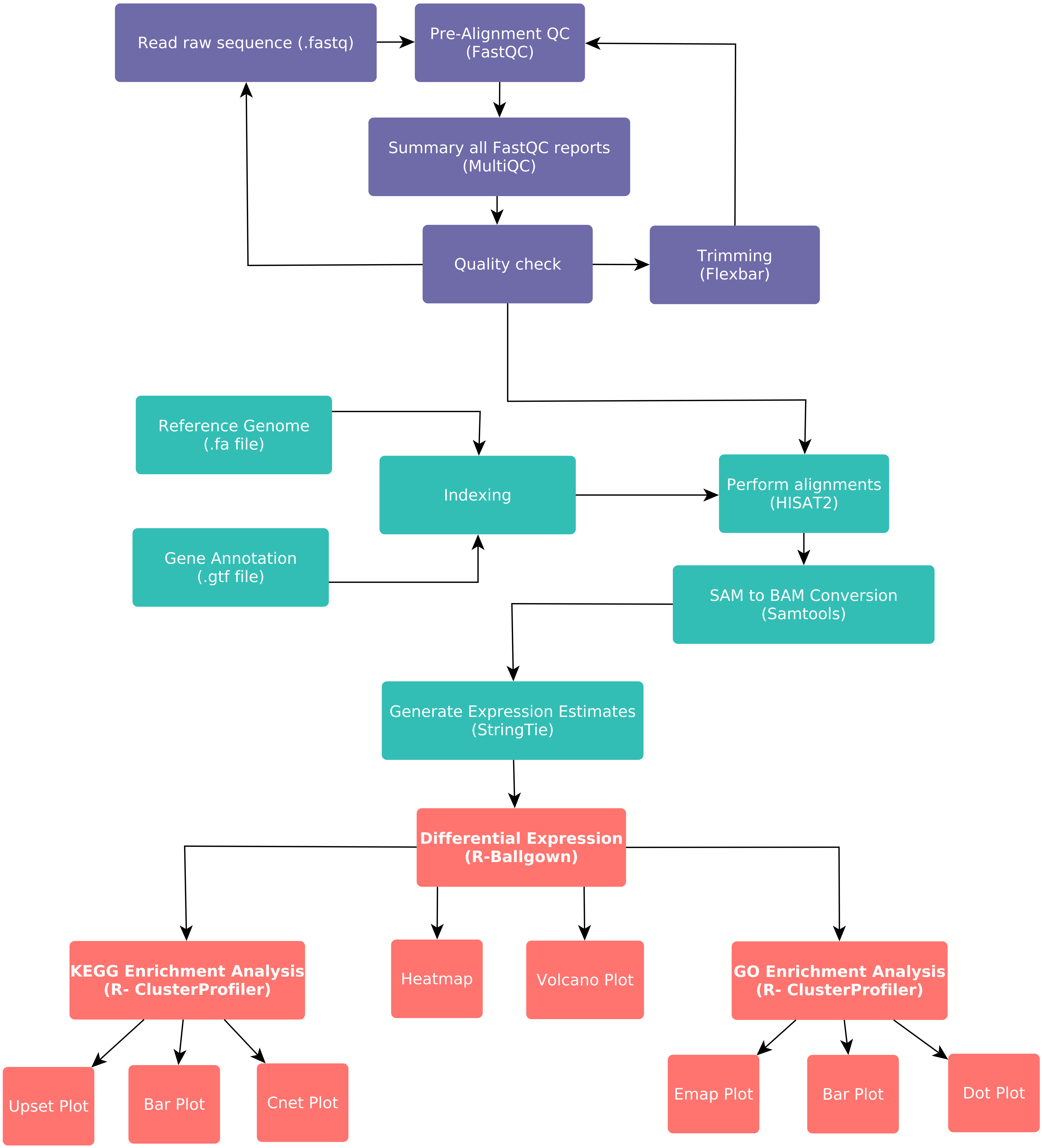

Read raw sequence (.fastq)
Pre-Alignment QC (FastQC)
Summary all FastQC reports (MultiQC)
Quality check
Trimming (Flexbar)
Reference Genome (.fa file)
Indexing
Perform alignments (HISAT2)
Gene Annotation (.gtf file)
SAM to BAM Conversion (Samtools)
Generate Expression Estimates (StringTie)
Differential Expression (R-Ballgown)
KEGG Enrichment Analysis (R- ClusterProfiler)
Heatmap
Volcano Plot
GO Enrichment Analysis (R- ClusterProfiler)
Upset Plot
Bar Plot
Cnet Plot
Emap Plot
Bar Plot
Dot Plot

Figure2

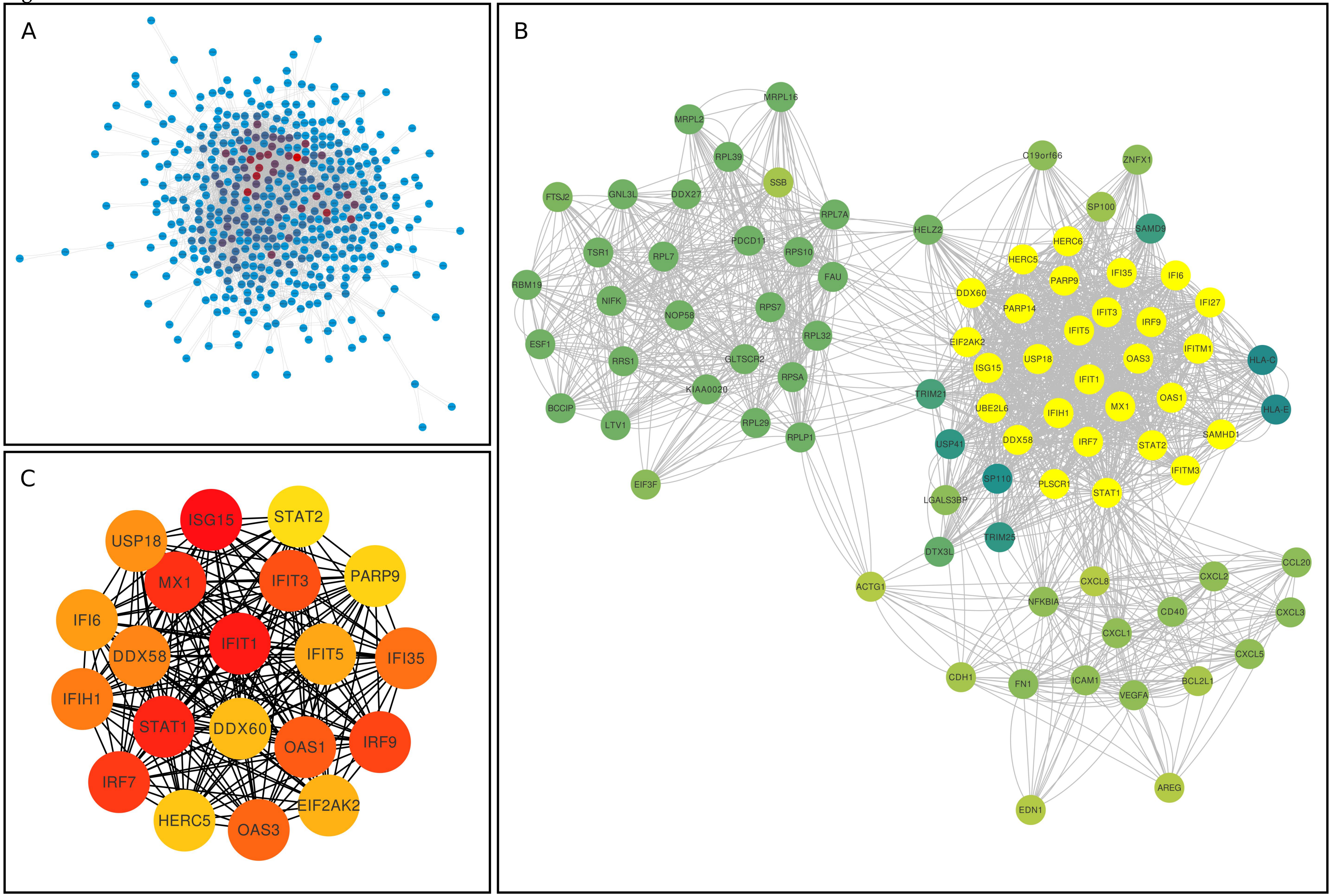

A
B
MRPL16
MRPL2
RPL39
SSB
FTSJ2
GNL3L
DDX27
RPL7A
TSR1
PDCD11
RPS10
RPL7
FAU
RBM19
NIFK
NOP58
RPS7
RPL32
ESF1
RRS1
GLTSCR2
RPSA
KIAA0020
BCCIP
LTV1
RPL29
RPLP1
EIF3F
C19orf66
ZNFX1
SP100
SAMD9
HELZ2
HERC6
HERC5
IFI35
IFI6
PARP9
DDX60
IFI27
PARP14
IFIT3
IRF9
IFIT5
EIF2AK2
IFITM1
HLA-C
USP18
OAS3
ISG15
IFIT1
TRIM21
OAS1
HLA-E
MX1
UBE2L6
IFIH1
USP41
DDX58
IRF7
STAT2
SAMHD1
IFITM3
SP110
PLSCR1
STAT1
LGALS3BP
TRIM25
DTX3L
CCL20
CXCL8
CXCL2
ACTG1
NFKBIA
CD40
CXCL3
CXCL1
CXCL5
CDH1
FN1
ICAM1
BCL2L1
VEGFA
AREG
EDN1
C
ISG15
STAT2
USP18
MX1
IFIT3
PARP9
IFI6
DDX58
IFIT1
IFIT5
IFI35
IFIH1
STAT1
DDX60
OAS1
IRF9
IRF7
HERC5
OAS3
EIF2AK2

Figure3

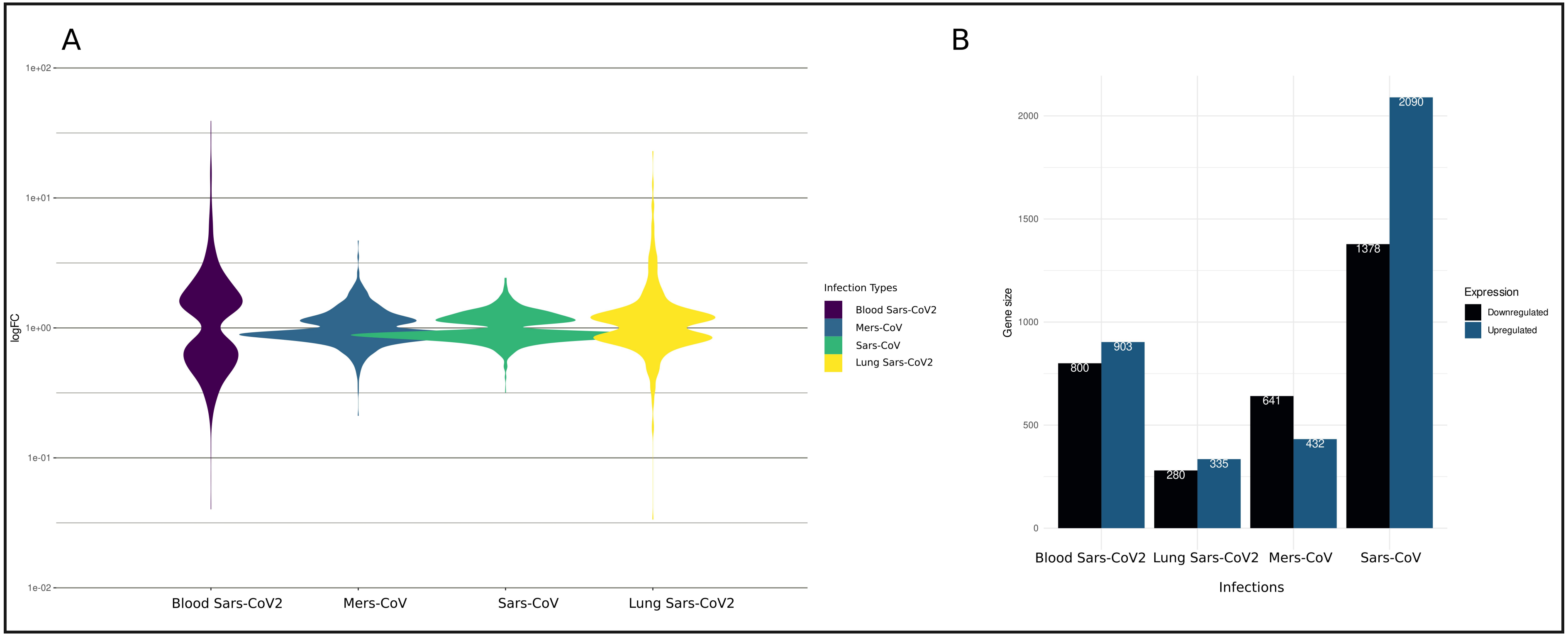
A
1e+02
1e+01
1e+00
1e-01
1e-02
logFC
Blood Sars-CoV2
Mers-CoV
Sars-CoV
Lung Sars-CoV2
Infection Types
Blood Sars-CoV2
Mers-CoV
Sars-CoV
Lung Sars-CoV2
B
Gene size
2000
1500
1000
500
0
2090
1378
903
800
641
432
335
280
Blood Sars-CoV2
Lung Sars-CoV2
Mers-CoV
Sars-CoV
Infections
Expression
Downregulated
Upregulated

Figure4

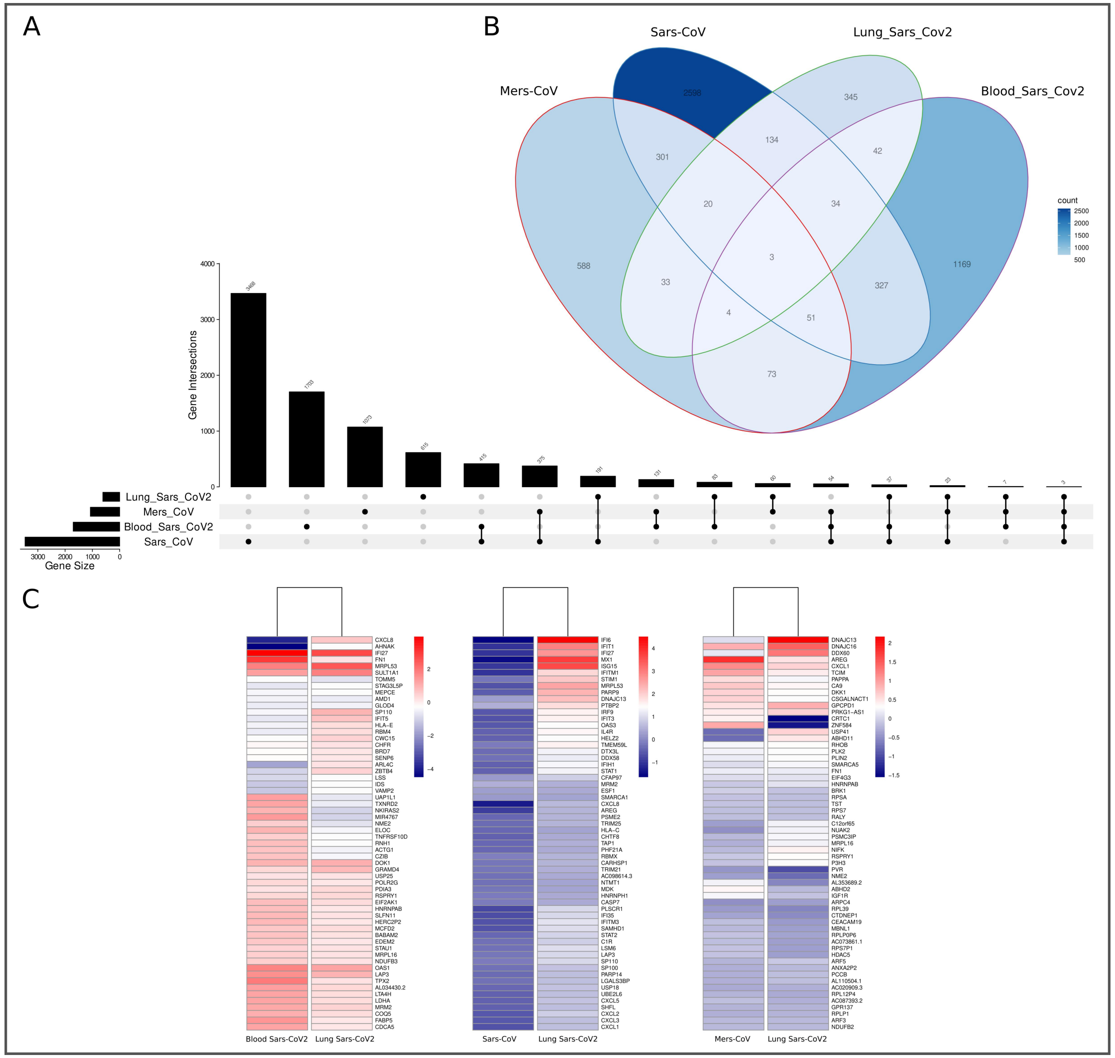
A
B
Sars-CoV
Lung_Sars_Cov2
Mers-CoV
Blood_Sars_Cov2
2598
345
301
134
42
20
34
588
3
1169
33
327
4
51
73
count
2500
2000
1500
1000
500
Gene Intersections
4000
3000
2000
1000
0
Lung_Sars_CoV2
Mers_CoV
Blood_Sars_CoV2
Sars_CoV
3000 2000 1000 0
Gene Size
C
Blood Sars-CoV2
Lung Sars-CoV2
Sars-CoV
Lung Sars-CoV2
Mers-CoV
Lung Sars-CoV2

Figure 5

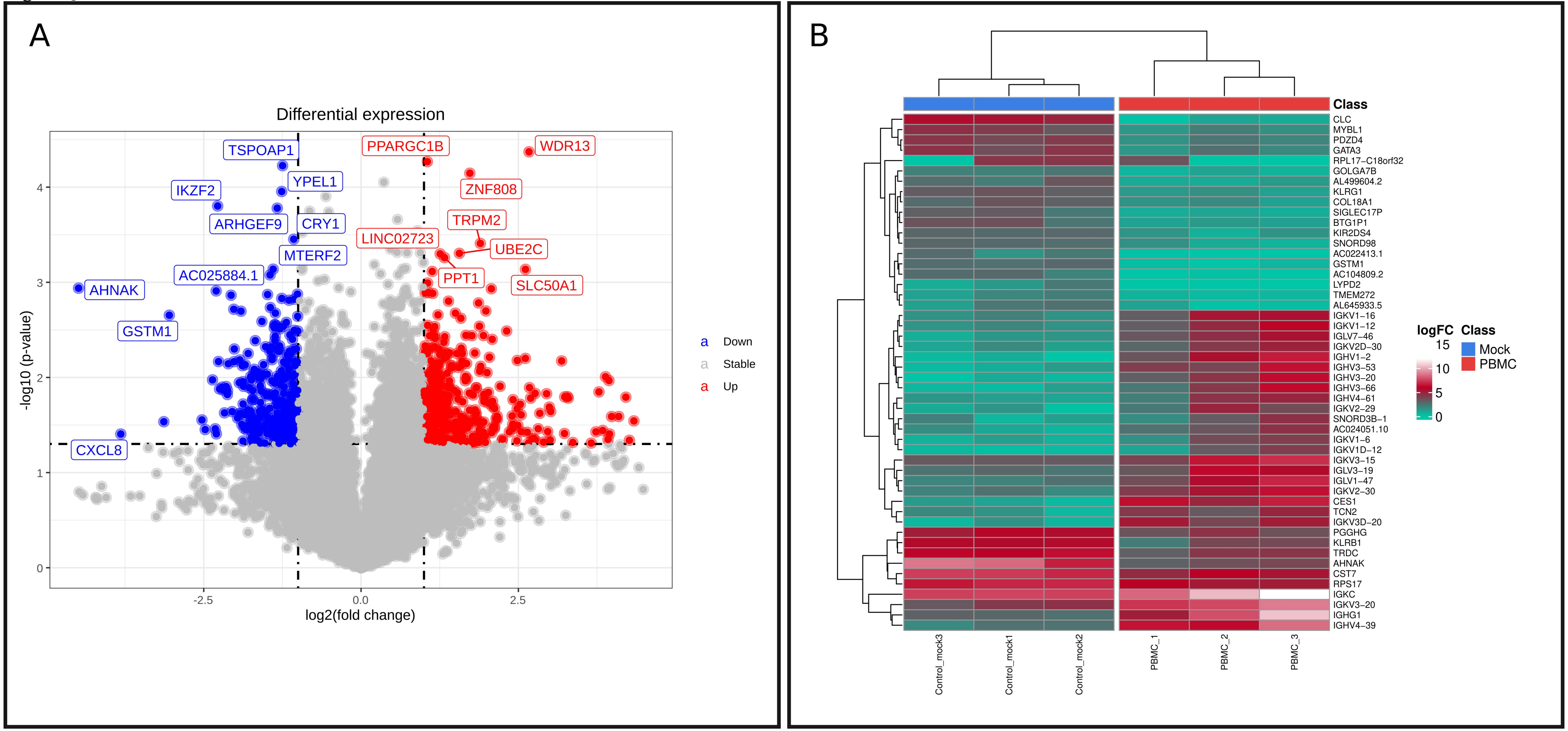

A
Differential expression
TSPOAP1
YPEL1
IKZF2
ARHGEF9
CRY1
MTERF2
AC025884.1
AHNAK
GSTM1
CXCL8
PPARGC1B
WDR13
ZNF808
TRPM2
LINC02723
UBE2C
PPT1
SLC50A1
-log10 (p-value)
log2(fold change)
a Down
a Stable
a Up
B
Class
logFC
Mock
PBMC
Control_mock3
Control_mock1
Control_mock2
PBMC_1
PBMC_2
PBMC_3
CLC
MYBL1
PDZD4
GATA3
RPL17−C18orf32
GOLGA7B
AL499604.2
KLRG1
COL18A1
SIGLEC17P
BTG1P1
KIR2DS4
SNORD98
AC022413.1
GSTM1
AC104809.2
LYPD2
TMEM272
AL645933.5
IGKV1−16
IGKV1−12
IGLV7−46
IGKV2D−30
IGHV1−2
IGHV3−53
IGHV3−20
IGHV3−66
IGHV4−61
IGKV2−29
SNORD3B−1
AC024051.10
IGKV1−6
IGKV1D−12
IGKV3−15
IGLV3−19
IGLV1−47
IGKV2−30
CES1
TCN2
IGKV3D−20
PGGHG
KLRB1
TRDC
AHNAK
CST7
RPS17
IGKC
IGKV3−20
IGHG1
IGHV4−39

### Figure 6

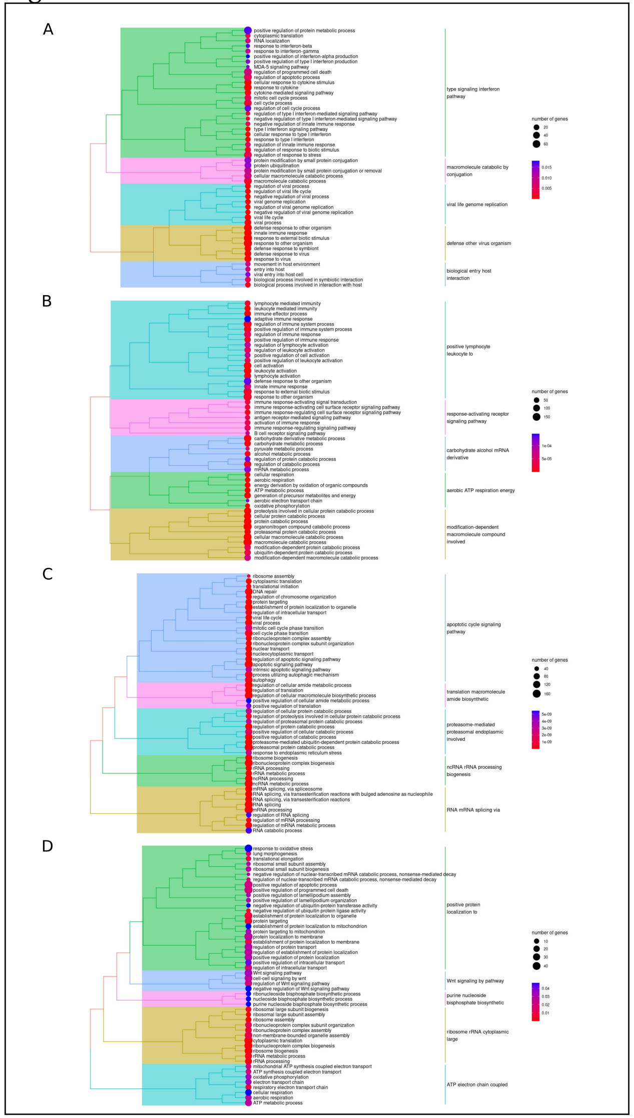

Figure 7

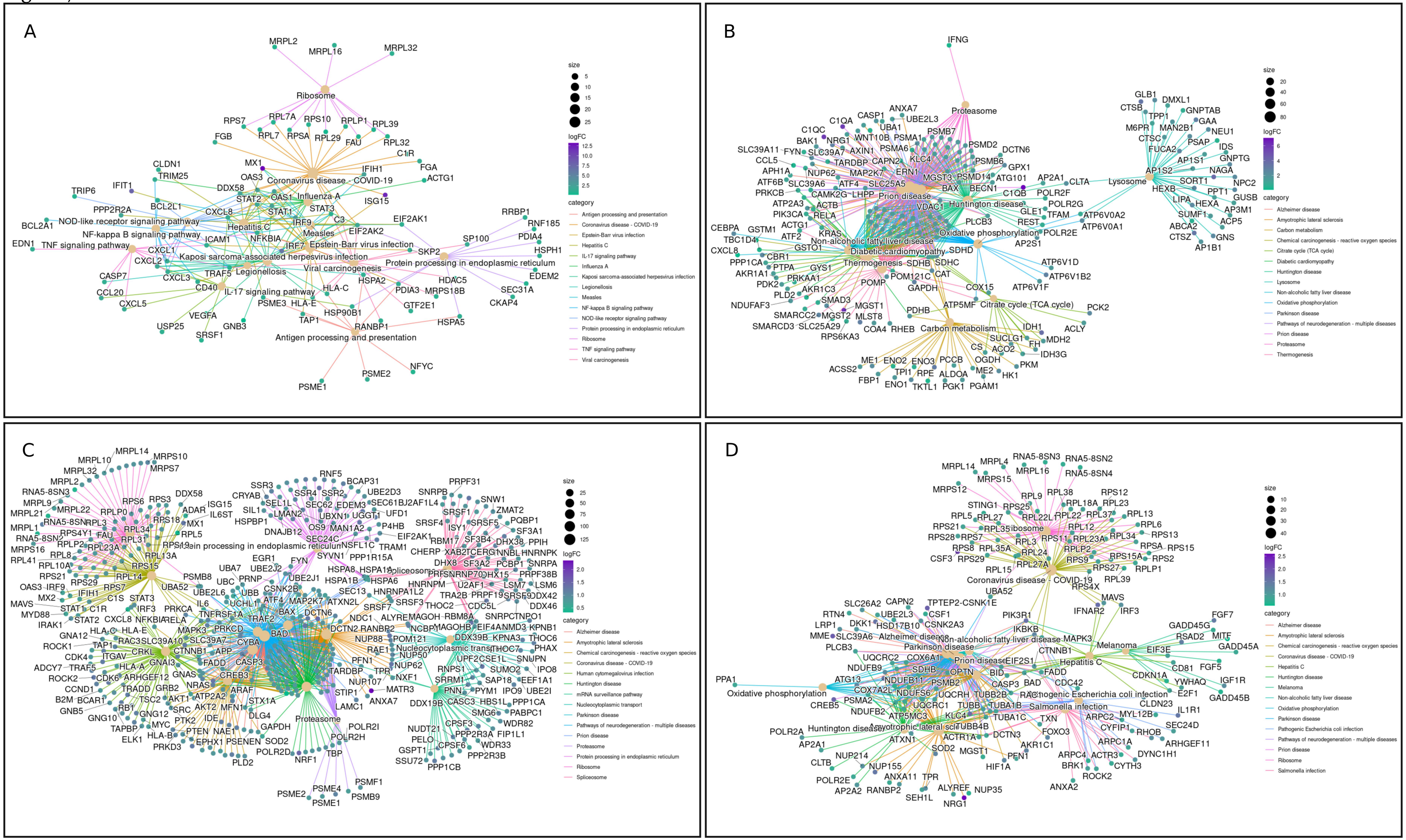

A
Ribosome
Coronavirus disease - COVID-19
Influenza A
NOD-like receptor signaling pathway
Hepatitis C
NF-kappa B signaling pathway
TNF signaling pathway
Measles
Epstein-Barr virus infection
Kaposi sarcoma-associated herpesvirus infection
Viral carcinogenesis
Protein processing in endoplasmic reticulum
Legionellosis
IL-17 signaling pathway
Antigen processing and presentation
size
logFC
category
B
Proteasome
Lysosome
Prion disease
Huntington disease
Non-alcoholic fatty liver disease
Oxidative phosphorylation
Diabetic cardiomyopathy
Thermogenesis
Citrate cycle (TCA cycle)
Carbon metabolism
C
Protein processing in endoplasmic reticulum
Spliceosome
Nucleocytoplasmic transport
Proteasome
D
Ribosome
Coronavirus disease - COVID-19
Alzheimer disease
Non-alcoholic fatty liver disease
Prion disease
Melanoma
Hepatitis C
Oxidative phosphorylation
Pathogenic Escherichia coli infection
Salmonella infection
Huntington disease
Amyotrophic lateral sclerosis

### Figure 8

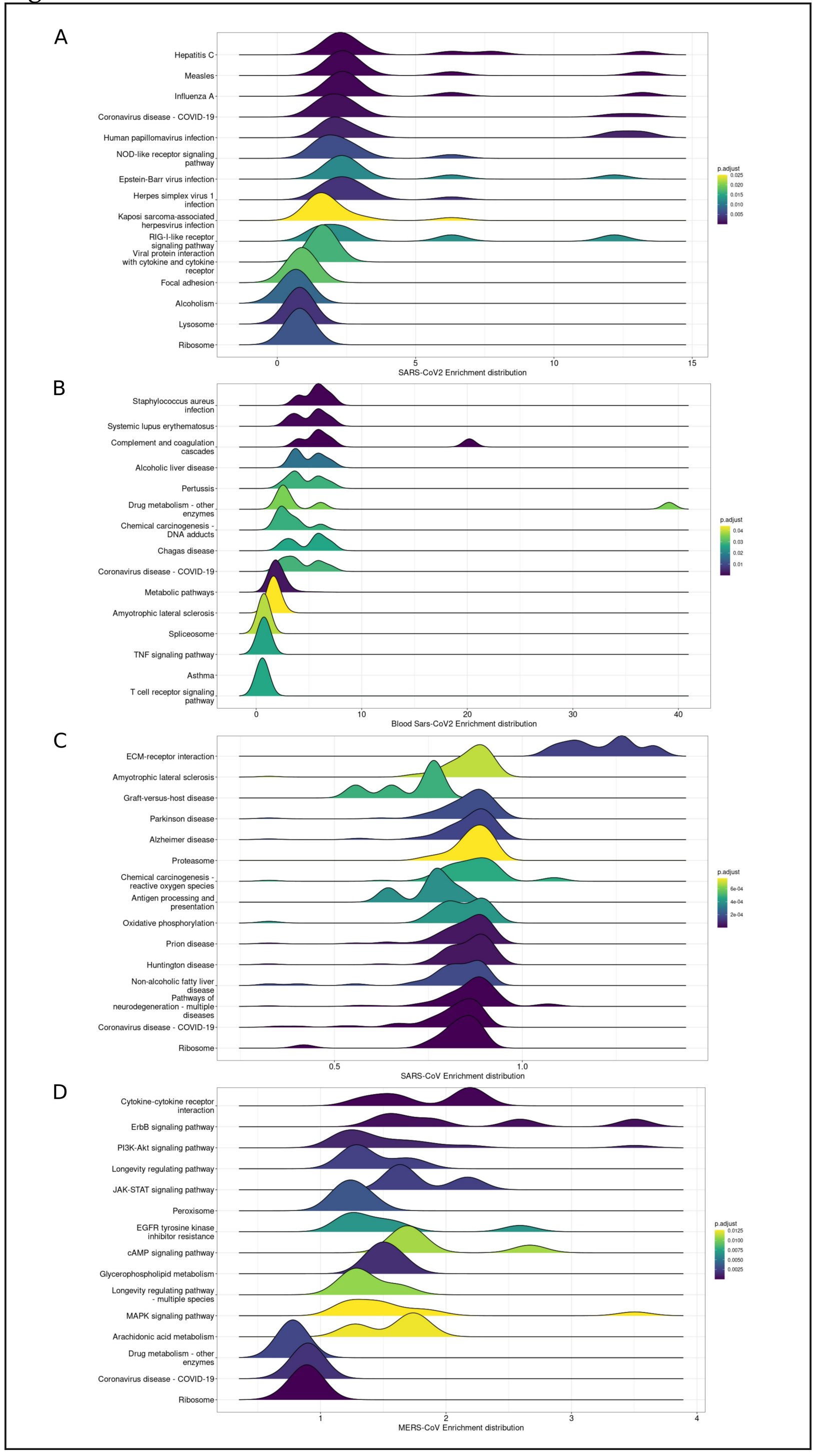
A
Hepatitis C
Measles
Influenza A
Coronavirus disease - COVID-19
Human papillomavirus infection
NOD-like receptor signaling pathway
Epstein-Barr virus infection
Herpes simplex virus 1 infection
Kaposi sarcoma-associated herpesvirus infection
RIG-I-like receptor signaling pathway
Viral protein interaction with cytokine and cytokine receptor
Focal adhesion
Alcoholism
Lysosome
Ribosome
p.adjust
0 5 10 15
SARS-CoV2 Enrichment distribution
B
Staphylococcus aureus infection
Systemic lupus erythematosus
Complement and coagulation cascades
Alcoholic liver disease
Pertussis
Drug metabolism - other enzymes
Chemical carcinogenesis - DNA adducts
Chagas disease
Coronavirus disease - COVID-19
Metabolic pathways
Amyotrophic lateral sclerosis
Spliceosome
TNF signaling pathway
Asthma
T cell receptor signaling pathway
p.adjust
0 10 20 30 40
Blood Sars-CoV2 Enrichment distribution
C
ECM-receptor interaction
Amyotrophic lateral sclerosis
Graft-versus-host disease
Parkinson disease
Alzheimer disease
Proteasome
Chemical carcinogenesis - reactive oxygen species
Antigen processing and presentation
Oxidative phosphorylation
Prion disease
Huntington disease
Non-alcoholic fatty liver disease
Pathways of neurodegeneration - multiple diseases
Coronavirus disease - COVID-19
Ribosome
p.adjust
0.5 1.0
SARS-CoV Enrichment distribution
D
Cytokine-cytokine receptor interaction
ErbB signaling pathway
PI3K-Akt signaling pathway
Longevity regulating pathway
JAK-STAT signaling pathway
Peroxisome
EGFR tyrosine kinase inhibitor resistance
cAMP signaling pathway
Glycerophospholipid metabolism
Longevity regulating pathway - multiple species
MAPK signaling pathway
Arachidonic acid metabolism
Drug metabolism - other enzymes
Coronavirus disease - COVID-19
Ribosome
p.adjust
1 2 3 4
MERS-CoV Enrichment distribution

**SARS-Cov2_DEGs**

| id | feature | fc | pval | qval | gene_name | de |
|---|---|---|---|---|---|---|
| ENSG0000022463 | gene | 0,033606837 | 0,04003026 | 0,765722912 | RPS27AP16 | -4,895101408 |
| ENSG0000027750 | gene | 0,164101059 | 0,003671247 | 0,493619508 | AC007383.4 | -2,607343549 |
| ENSG0000014440 | gene | 0,169918461 | 0,028136883 | 0,727417021 | METTL21A | -2,557085492 |
| ENSG0000011166 | gene | 0,18802193 | 0,006432253 | 0,560473839 | GNB3 | -2,411027152 |
| ENSG0000014824 | gene | 0,254017884 | 2,10045E-05 | 0,067991613 | SURF4 | -1,976998022 |
| ENSG0000016890 | gene | 0,278264188 | 0,0452138 | 0,776242869 | LRRC28 | -1,845472848 |
| ENSG0000016390 | gene | 0,284212508 | 0,018535967 | 0,669155317 | SENP2 | -1,814958047 |
| ENSG0000013051 | gene | 0,325010802 | 0,000282542 | 0,17148505 | PGPEP1 | -1,621440427 |
| ENSG0000011396 | gene | 0,340469676 | 0,010885541 | 0,575745644 | ARL6 | -1,554401785 |
| ENSG0000010566 | gene | 0,341004629 | 0,040792661 | 0,765722912 | CRTC1 | -1,55213677 |
| ENSG0000021427 | gene | 0,342405416 | 0,000892769 | 0,298954595 | ANG | -1,546222573 |
| ENSG0000017011 | gene | 0,345682561 | 0,034856571 | 0,761794317 | NIPA1 | -1,532480272 |
| ENSG0000004718 | gene | 0,361423715 | 0,012856328 | 0,605697652 | YTHDC2 | -1,468236923 |
| ENSG0000017157 | gene | 0,387903287 | 0,037689775 | 0,765722912 | ZNF584 | -1,366231093 |
| ENSG0000013011 | gene | 0,404750714 | 0,006610906 | 0,560473839 | GNL3L | -1,30489447 |
| ENSG0000009214 | gene | 0,420373162 | 0,012172536 | 0,60034321 | G2E3 | -1,250257527 |
| ENSG0000025525 | gene | 0,422682744 | 0,027982555 | 0,727417021 | AL078612.2 | -1,242352879 |
| ENSG0000016224 | gene | 0,426391216 | 0,0453228 | 0,776242869 | RPL29 | -1,229750377 |
| ENSG0000010104 | gene | 0,440564497 | 0,002459398 | 0,493619508 | SGK2 | -1,182574854 |
| ENSG0000016407 | gene | 0,451875231 | 0,000548567 | 0,253673251 | MFSD8 | -1,146003614 |
| ENSG0000017853 | gene | 0,455426867 | 0,036498789 | 0,765722912 | SLC25A20 | -1,134708692 |
| ENSG0000019645 | gene | 0,458482338 | 0,029867873 | 0,745621897 | PIK3R4 | -1,125061936 |
| ENSG0000027724 | gene | 0,462073106 | 0,029952549 | 0,745818463 | MIR6784 | -1,113806972 |
| ENSG0000016539 | gene | 0,46798039 | 0,032344129 | 0,751763165 | WRN | -1,095480017 |
| ENSG0000025838 | gene | 0,480723547 | 0,029019106 | 0,735782087 | PPT2-EGFL8 | -1,056720624 |
| ENSG0000017225 | gene | 0,482675233 | 0,015975282 | 0,65735579 | SERHL | -1,050875293 |
| ENSG0000011761 | gene | 0,489823127 | 0,009420846 | 0,57333908 | RSRP1 | -1,029667203 |
| ENSG0000015760 | gene | 0,489895041 | 0,035129134 | 0,761794317 | TMEM164 | -1,029455406 |
| ENSG0000012354 | gene | 0,494640006 | 0,010845106 | 0,575745644 | NDUFAF4 | -1,015549165 |
| ENSG0000007300 | gene | 0,504388383 | 0,006494328 | 0,560473839 | PVR | -0,987393046 |
| ENSG0000011271 | gene | 0,507545057 | 0,034762362 | 0,761794317 | VEGFA | -0,978392193 |
| ENSG0000016825 | gene | 0,512823339 | 0,038779659 | 0,765722912 | NKIRAS2 | -0,963466174 |
| ENSG0000017533 | gene | 0,512963291 | 0,001388158 | 0,421262513 | BANF1 | -0,963072509 |
| ENSG0000026426 | gene | 0,518690098 | 0,00861686 | 0,566479818 | MIR4767 | -0,947055267 |
| ENSG0000015751 | gene | 0,533656844 | 0,023108 | 0,700392548 | TSC22D3 | -0,906015749 |
| ENSG0000026290 | gene | 0,54781914 | 0,029778635 | 0,745621897 | AC027796.4 | -0,868228423 |
| ENSG0000023124 | gene | 0,549782549 | 0,031322669 | 0,749219164 | C1DP1 | -0,863066981 |
| ENSG0000006722 | gene | 0,558273145 | 0,01526966 | 0,636410574 | STOML1 | -0,840956936 |
| ENSG0000016971 | gene | 0,568421867 | 0,003193115 | 0,493619508 | CNBP | -0,814966039 |
| ENSG0000017539 | gene | 0,578132286 | 0,046775661 | 0,784665672 | EIF3F | -0,790528451 |
| ENSG0000012478 | gene | 0,582712827 | 0,040537296 | 0,765722912 | RREB1 | -0,779143026 |
| ENSG0000017576 | gene | 0,585967636 | 0,036430135 | 0,765722912 | TOMM5 | -0,77110711 |
| ENSG0000023696 | gene | 0,589520314 | 0,023633523 | 0,700392548 | GGT8P | -0,762386569 |
| ENSG0000023679 | gene | 0,59030577 | 0,046367675 | 0,783814537 | AL513175.2 | -0,76046565 |
| ENSG0000024367 | gene | 0,595957293 | 0,03761916 | 0,765722912 | NME2 | -0,746719145 |
| ENSG0000000581 | gene | 0,601435744 | 0,034666964 | 0,761794317 | MYCBP2 | -0,733517483 |
| ENSG0000013997 | gene | 0,610976183 | 0,023953087 | 0,700392548 | SLC38A6 | -0,710811952 |
| ENSG0000014096 | gene | 0,614131213 | 0,016378547 | 0,6599671 | OSGIN1 | -0,703381165 |
| ENSG0000027518 | gene | 0,614761005 | 0,008114167 | 0,560473839 | AC048341.2 | -0,701902438 |
| ENSG0000019735 | gene | 0,619697503 | 0,044904264 | 0,776242869 | UAP1L1 | -0,690363939 |
| ENSG0000013422 | gene | 0,62020154 | 0,032367476 | 0,751763165 | PSRC1 | -0,689190988 |
| ENSG0000013883 | gene | 0,630451694 | 0,033383333 | 0,754653699 | MAPK8IP3 | -0,665542261 |
| ENSG0000022695 | gene | 0,631052698 | 0,006832487 | 0,560473839 | DANCR | -0,664167608 |
| ENSG0000007686 | gene | 0,637704426 | 0,020084804 | 0,682146839 | RAP1GAP | -0,649040201 |
| ENSG0000025847 | gene | 0,639221808 | 0,049918014 | 0,788217613 | LINC02313 | -0,645611467 |

| | | | | | | |
|---|---|---|---|---|---|---|
| ENSG0000008599 | gene | 0,644852533 | 0,043370404 | 0,767591009 | RAD54L | -0,632958817 |
| ENSG0000025528 | gene | 0,650167628 | 0,006667442 | 0,560473839 | AP006621.3 | -0,621116369 |
| ENSG0000014341 | gene | 0,650803889 | 0,005127671 | 0,524190612 | CERS2 | -0,619705222 |
| ENSG0000010147 | gene | 0,651588763 | 0,008814582 | 0,566479818 | ACOT8 | -0,617966371 |
| ENSG0000017047 | gene | 0,657524406 | 0,009136845 | 0,570481883 | KRT4 | -0,60488365 |
| ENSG0000016241 | gene | 0,658016943 | 0,035579373 | 0,765722912 | KLHL21 | -0,603803364 |
| ENSG0000010908 | gene | 0,667933669 | 9,0942E-05 | 0,092192886 | TMEM97 | -0,582223257 |
| ENSG0000014377 | gene | 0,67459005 | 0,00049806 | 0,2418332 | CDC42BPA | -0,567917055 |
| ENSG0000019891 | gene | 0,680643636 | 0,031946129 | 0,750135666 | RPL39 | -0,55502845 |
| ENSG0000023370 | gene | 0,68372293 | 0,010700114 | 0,575745644 | AL353689.2 | -0,548516286 |
| ENSG0000018447 | gene | 0,688197336 | 0,019996664 | 0,682146839 | TXNRD2 | -0,539105787 |
| ENSG0000016594 | gene | 0,689533614 | 0,009877171 | 0,57333908 | IFI27L1 | -0,536307213 |
| ENSG0000022660 | gene | 0,690578195 | 0,015550455 | 0,642597757 | FTLP3 | -0,534123313 |
| ENSG0000017582 | gene | 0,691277559 | 0,008144475 | 0,560473839 | CTDNEP1 | -0,532663002 |
| ENSG0000012350 | gene | 0,693308145 | 0,005064424 | 0,524190612 | AMD1 | -0,528431385 |
| ENSG0000021018 | gene | 0,69358461 | 0,045013036 | 0,776242869 | MT-TS2 | -0,527856207 |
| ENSG0000011535 | gene | 0,694194546 | 0,023294364 | 0,700392548 | POLE4 | -0,526588065 |
| ENSG0000026279 | gene | 0,696187329 | 0,043025625 | 0,767591009 | AC130343.1 | -0,522452539 |
| ENSG0000017317 | gene | 0,702108567 | 0,040795944 | 0,765722912 | MTX1 | -0,510233963 |
| ENSG0000016203 | gene | 0,704288758 | 0,038595919 | 0,765722912 | SPSB3 | -0,50576104 |
| ENSG0000016769 | gene | 0,707640247 | 0,020674633 | 0,697122783 | GLOD4 | -0,498911991 |
| ENSG0000004060 | gene | 0,715363866 | 0,018703191 | 0,669657331 | RTN4R | -0,483250847 |
| ENSG0000026596 | gene | 0,716144366 | 0,024566171 | 0,702826025 | GACAT2 | -0,481677649 |
| ENSG0000027994 | gene | 0,716187028 | 0,016179861 | 0,658924168 | AC013489.3 | -0,481591706 |
| ENSG0000017200 | gene | 0,71894428 | 0,005222027 | 0,524190612 | THOP1 | -0,476048132 |
| ENSG0000020493 | gene | 0,723775759 | 0,029316716 | 0,739466574 | ATP6V0E2-AS1 | -0,466385306 |
| ENSG0000017165 | gene | 0,724434734 | 0,025583031 | 0,702826025 | NMRAL2P | -0,465072375 |
| ENSG0000026728 | gene | 0,724724263 | 0,022872991 | 0,700392548 | AC005306.1 | -0,464495899 |
| ENSG0000025866 | gene | 0,726895182 | 0,016890181 | 0,663077244 | AL117190.2 | -0,460180751 |
| ENSG0000016097 | gene | 0,729934311 | 0,042027395 | 0,767591009 | PPP1R16A | -0,454161457 |
| ENSG0000016069 | gene | 0,731292495 | 0,001533041 | 0,425554113 | SHC1 | -0,451479538 |
| ENSG0000013800 | gene | 0,732088874 | 0,00736261 | 0,560473839 | IFT172 | -0,449909295 |
| ENSG0000011265 | gene | 0,733859397 | 0,010802685 | 0,575745644 | MRPL2 | -0,446424416 |
| ENSG0000018400 | gene | 0,735442764 | 0,037303398 | 0,765722912 | ACTG1 | -0,443315027 |
| ENSG0000014471 | gene | 0,737940456 | 0,039154888 | 0,765722912 | RPL32 | -0,438423685 |
| ENSG0000007768 | gene | 0,74041954 | 0,047325219 | 0,785311848 | JADE1 | -0,433585126 |
| ENSG0000018656 | gene | 0,741802168 | 0,012126822 | 0,60034321 | CEACAM19 | -0,43089361 |
| ENSG0000009991 | gene | 0,743468076 | 0,023599552 | 0,700392548 | MED15 | -0,427657299 |
| ENSG0000014980 | gene | 0,745287181 | 0,000449495 | 0,229739128 | FAU | -0,42413165 |
| ENSG0000026859 | gene | 0,748444155 | 0,008205971 | 0,560473839 | AL355312.3 | -0,418033421 |
| ENSG0000016020 | gene | 0,755250701 | 0,034186264 | 0,761428461 | CBS | -0,404972476 |
| ENSG0000016489 | gene | 0,758906545 | 0,02323448 | 0,700392548 | FMC1 | -0,398005858 |
| ENSG0000015458 | gene | 0,759469529 | 0,019145241 | 0,674862552 | ELOC | -0,396936012 |
| ENSG0000016238 | gene | 0,76072435 | 0,033540386 | 0,754653699 | CZIB | -0,39455431 |
| ENSG0000015260 | gene | 0,762514078 | 0,019250018 | 0,674862552 | MBNL1 | -0,391164121 |
| ENSG0000026326 | gene | 0,763600907 | 0,048520967 | 0,785311848 | RPS7P1 | -0,389109279 |
| ENSG0000010884 | gene | 0,763913716 | 0,049181568 | 0,788217613 | HDAC5 | -0,3885184 |
| ENSG0000017527 | gene | 0,768429438 | 0,003892752 | 0,497401554 | TP53I11 | -0,380015306 |
| ENSG0000020443 | gene | 0,769484792 | 0,021697767 | 0,700392548 | GPANK1 | -0,378035281 |
| ENSG0000024155 | gene | 0,770290323 | 0,008311012 | 0,560473839 | ARPC4 | -0,376525794 |
| ENSG0000016710 | gene | 0,771009192 | 0,018031878 | 0,665808244 | FAM102A | -0,375180034 |
| ENSG0000017994 | gene | 0,772974861 | 0,043053547 | 0,767591009 | FIZ1 | -0,3715066 |
| ENSG0000016888 | gene | 0,775182606 | 0,048779491 | 0,787085197 | TNIP2 | -0,367391896 |
| ENSG0000006613 | gene | 0,775610049 | 0,024190623 | 0,702826025 | NFYC | -0,366596599 |
| ENSG0000024229 | gene | 0,775823773 | 0,013432319 | 0,618536995 | AC073861.1 | -0,366199112 |
| ENSG0000019883 | gene | 0,778155516 | 0,003481897 | 0,493619508 | OPA1 | -0,361869586 |
| ENSG0000016918 | gene | 0,778165775 | 0,025823029 | 0,702826025 | NSMCE1 | -0,361850566 |

| | | | | | | |
|---|---|---|---|---|---|---|
| ENSG000001696 | gene | 0,779077354 | 0,007090774 | 0,560473839 | HEXD | -0,360161515 |
| ENSG000001575 | gene | 0,780940354 | 0,042230149 | 0,767591009 | ETS2 | -0,356715732 |
| ENSG000001230 | gene | 0,782139514 | 0,010755251 | 0,575745644 | CDKN2C | -0,354502125 |
| ENSG000001658 | gene | 0,783722537 | 0,047026611 | 0,784665672 | NSMF | -0,351585111 |
| ENSG000001030 | gene | 0,784086378 | 0,016276452 | 0,658924168 | FA2H | -0,350915499 |
| ENSG000000104 | gene | 0,785077934 | 0,02631536 | 0,702826025 | IDS | -0,349092219 |
| ENSG000001229 | gene | 0,786743818 | 0,025155263 | 0,702826025 | RBM19 | -0,346034157 |
| ENSG000001735 | gene | 0,787478915 | 0,036374022 | 0,765722912 | TNFRSF10D | -0,3446868 |
| ENSG000001468 | gene | 0,788221818 | 0,01394982 | 0,621406878 | MEPCE | -0,34332641 |
| ENSG000002404 | gene | 0,78830373 | 0,042157357 | 0,767591009 | AC005586.1 | -0,343176493 |
| ENSG000001833 | gene | 0,791426563 | 0,034316718 | 0,761794317 | C19orf71 | -0,337472608 |
| ENSG000002422 | gene | 0,792096059 | 0,036966128 | 0,765722912 | STAG3L5P | -0,336252695 |
| ENSG000001177 | gene | 0,79524243 | 0,012475324 | 0,605697652 | PPP1R8 | -0,330533362 |
| ENSG000001386 | gene | 0,795455041 | 0,009659896 | 0,57333908 | SEC31A | -0,330147703 |
| ENSG000001826 | gene | 0,796868328 | 0,007488691 | 0,560473839 | SNORC | -0,327586738 |
| ENSG000001048 | gene | 0,797756623 | 0,009995316 | 0,57333908 | AMH | -0,325979414 |
| ENSG000002135 | gene | 0,79792613 | 0,006471293 | 0,560473839 | RPLP0P6 | -0,325672903 |
| ENSG000002355 | gene | 0,798593629 | 0,002761979 | 0,493619508 | AC010978.1 | -0,324466532 |
| ENSG000001610 | gene | 0,800277186 | 0,028546068 | 0,729502277 | SQSTM1 | -0,321428313 |
| ENSG000000231 | gene | 0,800862226 | 0,006866123 | 0,560473839 | RNH1 | -0,32037402 |
| ENSG000001249 | gene | 0,801208239 | 0,032436285 | 0,751763165 | AHNAK | -0,319750838 |
| ENSG000001602 | gene | 0,801363512 | 0,023763074 | 0,700392548 | LSS | -0,319471273 |
| ENSG000001782 | gene | 0,802437554 | 0,045482656 | 0,777609275 | GALNT11 | -0,31753897 |
| ENSG000002436 | gene | 0,802701602 | 0,008270893 | 0,560473839 | AC018638.5 | -0,317064318 |
| ENSG000002040 | gene | 0,802871166 | 0,022192992 | 0,700392548 | ARMCX7P | -0,316759592 |
| ENSG000001571 | gene | 0,805270411 | 0,045189923 | 0,776242869 | TIMP4 | -0,312454772 |
| ENSG000002202 | gene | 0,807391658 | 0,020941023 | 0,700392548 | VAMP2 | -0,308659413 |
| ENSG000001050 | gene | 0,809637028 | 0,008755149 | 0,566479818 | VRK3 | -0,304652824 |
| ENSG000002285 | gene | 0,80964405 | 0,039246429 | 0,765722912 | EEF1A1P11 | -0,30464031 |
| ENSG000001634 | gene | 0,810246533 | 0,016615791 | 0,663077244 | TMEM79 | -0,303567152 |
| ENSG000001784 | gene | 0,810924443 | 0,033400227 | 0,754653699 | RPL10P16 | -0,302360595 |
| ENSG000001062 | gene | 0,812611462 | 0,024914053 | 0,702826025 | SNX8 | -0,29936238 |
| ENSG000001162 | gene | 0,813466392 | 0,024848811 | 0,702826025 | QSOX1 | -0,297845353 |
| ENSG000002583 | gene | 0,81369951 | 0,017715632 | 0,663077244 | AC034102.6 | -0,297431973 |
| ENSG000000883 | gene | 0,815785665 | 0,012911071 | 0,605697652 | PDRG1 | -0,293737939 |
| ENSG000002501 | gene | 0,816581702 | 0,019225677 | 0,674862552 | EEF1A1P13 | -0,292330854 |
| ENSG000001858 | gene | 0,81659494 | 0,033416777 | 0,754653699 | RPL12P4 | -0,292307467 |
| ENSG000001615 | gene | 0,817304749 | 0,008764492 | 0,566479818 | FDXR | -0,291053978 |
| ENSG000001243 | gene | 0,817463781 | 0,038129013 | 0,765722912 | NAGK | -0,290773285 |
| ENSG000001671 | gene | 0,817539892 | 0,01865203 | 0,669657331 | COQ4 | -0,290638965 |
| ENSG000001731 | gene | 0,817888645 | 0,002257158 | 0,487094782 | ADCK5 | -0,29002366 |
| ENSG000001648 | gene | 0,819947283 | 0,02632791 | 0,702826025 | CDK5 | -0,286396937 |
| ENSG000002639 | gene | 0,820196159 | 0,008925076 | 0,566479818 | AC087393.2 | -0,285959107 |
| ENSG000001343 | gene | 0,824156927 | 0,046017319 | 0,782616789 | LPIN1 | -0,279009029 |
| ENSG000001769 | gene | 0,824316633 | 0,040724219 | 0,765722912 | DPP7 | -0,27872949 |
| ENSG000000547 | gene | 0,824907198 | 0,048494986 | 0,785311848 | ATP9A | -0,277696269 |
| ENSG000002594 | gene | 0,825346715 | 0,044539593 | 0,776242869 | AC023355.2 | -0,276927795 |
| ENSG000000811 | gene | 0,826242075 | 0,003245229 | 0,493619508 | EXD2 | -0,275363565 |
| ENSG000002325 | gene | 0,827896718 | 0,013800713 | 0,620457054 | RPL3P4 | -0,272477295 |
| ENSG000002306 | gene | 0,828712064 | 0,036754672 | 0,765722912 | RPS23P8 | -0,271057171 |
| ENSG000001723 | gene | 0,829500249 | 0,014929858 | 0,633117253 | COPRS | -0,269685681 |
| ENSG000001581 | gene | 0,829690242 | 0,003287956 | 0,493619508 | TPRG1L | -0,269355275 |
| ENSG000002688 | gene | 0,829720023 | 0,037811835 | 0,765722912 | AC020909.3 | -0,269303494 |
| ENSG000001712 | gene | 0,830530914 | 0,001660085 | 0,444131997 | NPTX1 | -0,267894226 |
| ENSG000000825 | gene | 0,830800585 | 0,043963012 | 0,771136021 | TRAF5 | -0,267425863 |
| ENSG000001004 | gene | 0,830820655 | 0,022158578 | 0,700392548 | CERK | -0,267391011 |
| ENSG000001853 | gene | 0,831100037 | 0,030403104 | 0,749158036 | TCN2 | -0,266905955 |

| | | | | | | |
|---|---|---|---|---|---|---|
| ENSG0000017362 | gene | 0,831416489 | 0,008039437 | 0,560473839 | LRFN4 | -0,266356735 |
| ENSG0000011566 | gene | 0,834968831 | 0,043194253 | 0,767591009 | STK16 | -0,260205752 |
| ENSG0000016067 | gene | 0,837468472 | 0,005549264 | 0,544332311 | CHTOP | -0,255893216 |
| ENSG0000014760 | gene | 0,838384786 | 0,003611306 | 0,493619508 | RPL7 | -0,254315558 |
| ENSG0000015525 | gene | 0,839045904 | 0,029307717 | 0,739466574 | MARVELD1 | -0,253178353 |
| ENSG0000010493 | gene | 0,840236951 | 0,011042664 | 0,579650325 | DMPK | -0,251131862 |
| ENSG0000010973 | gene | 0,840327692 | 0,011249792 | 0,583167084 | MFSD10 | -0,250976068 |
| ENSG0000009932 | gene | 0,841473284 | 0,048038479 | 0,785311848 | MZF1 | -0,249010627 |
| ENSG0000011965 | gene | 0,842790107 | 0,023894805 | 0,700392548 | NPC2 | -0,246754716 |
| ENSG0000014830 | gene | 0,842916756 | 0,030982302 | 0,749158036 | RPL7A | -0,246537933 |
| ENSG0000018504 | gene | 0,843286923 | 0,013439533 | 0,618536995 | CIB1 | -0,245904512 |
| ENSG0000022519 | gene | 0,844425722 | 0,047874711 | 0,785311848 | PLEKHM1 | -0,243957569 |
| ENSG0000014068 | gene | 0,845071888 | 0,018479841 | 0,669155317 | C16orf58 | -0,242854023 |
| ENSG0000013149 | gene | 0,845737364 | 0,011289817 | 0,583167084 | NDUFA2 | -0,241718377 |
| ENSG0000012608 | gene | 0,846221698 | 0,016149656 | 0,658924168 | UROD | -0,240892416 |
| ENSG0000018127 | gene | 0,846372437 | 0,007933193 | 0,560473839 | FRAT2 | -0,240635449 |
| ENSG0000000466 | gene | 0,847464014 | 0,003803245 | 0,497082126 | CAMKK1 | -0,238775987 |
| ENSG0000013886 | gene | 0,849069592 | 0,038677124 | 0,765722912 | GUCD1 | -0,236045289 |
| ENSG0000024431 | gene | 0,849077865 | 0,026162185 | 0,702826025 | AC024293.1 | -0,236031231 |
| ENSG0000021367 | gene | 0,849214554 | 0,022819958 | 0,700392548 | NCKIPSD | -0,235798998 |
| ENSG0000024430 | gene | 0,849743515 | 0,009485725 | 0,57333908 | AL589743.1 | -0,234900648 |
| ENSG0000013894 | gene | 0,850439348 | 0,00797639 | 0,560473839 | RNF185 | -0,233719747 |
| ENSG0000024670 | gene | 0,851463312 | 0,021909991 | 0,700392548 | H2AJ | -0,231983728 |
| ENSG0000010572 | gene | 0,853108553 | 0,033267972 | 0,754653699 | ATP13A1 | -0,229198768 |
| ENSG0000017326 | gene | 0,854634929 | 0,031716588 | 0,750135666 | GPR137 | -0,226619814 |
| ENSG0000009067 | gene | 0,85510689 | 0,027148516 | 0,722299273 | MCOLN1 | -0,225823323 |
| ENSG0000012873 | gene | 0,855160132 | 0,015105987 | 0,636410574 | HERC2 | -0,2257335 |
| ENSG0000021438 | gene | 0,855589287 | 0,035767818 | 0,765722912 | RPS3AP26 | -0,225009677 |
| ENSG0000013781 | gene | 0,858991731 | 0,048307461 | 0,785311848 | RPLP1 | -0,219283851 |
| ENSG0000000885 | gene | 0,85899502 | 0,001165677 | 0,365157756 | RHOBTB2 | -0,219278328 |
| ENSG0000013015 | gene | 0,859640183 | 0,042392814 | 0,767591009 | ECSIT | -0,218195173 |
| ENSG0000013645 | gene | 0,859934837 | 0,008666966 | 0,566479818 | SRSF1 | -0,217700754 |
| ENSG0000005112 | gene | 0,860478799 | 0,043186703 | 0,767591009 | HOMER3 | -0,216788449 |
| ENSG0000012286 | gene | 0,860648099 | 0,043992313 | 0,771136021 | CHST3 | -0,216504625 |
| ENSG0000008707 | gene | 0,861121663 | 0,013155379 | 0,61125306 | TRIP6 | -0,215711012 |
| ENSG0000005100 | gene | 0,862060566 | 0,038918803 | 0,765722912 | FAM160A2 | -0,214138862 |
| ENSG0000014068 | gene | 0,862700606 | 0,02601525 | 0,702826025 | TGFB1I1 | -0,213068126 |
| ENSG0000011512 | gene | 0,863286508 | 0,023350337 | 0,700392548 | TP53I3 | -0,212088654 |
| ENSG0000012296 | gene | 0,864250254 | 0,007966404 | 0,560473839 | CIT | -0,210478973 |
| ENSG0000023675 | gene | 0,864618881 | 0,021499124 | 0,700392548 | DNAJC9-AS1 | -0,209863753 |
| ENSG0000014841 | gene | 0,86494707 | 0,021041128 | 0,700392548 | NACC2 | -0,209316244 |
| ENSG0000014892 | gene | 0,865369389 | 0,024677598 | 0,702826025 | BTBD10 | -0,208612007 |
| ENSG0000025988 | gene | 0,865492307 | 0,039312832 | 0,765722912 | NR4A1AS | -0,208407098 |
| ENSG0000017273 | gene | 0,866246482 | 0,000170382 | 0,137881717 | LRRC20 | -0,207150507 |
| ENSG0000017686 | gene | 0,870448709 | 0,033907738 | 0,756961016 | AL358781.1 | -0,200168806 |
| ENSG0000013428 | gene | 0,871158291 | 0,042256698 | 0,767591009 | ARF3 | -0,198993212 |
| ENSG0000014053 | gene | 0,872087677 | 0,038206643 | 0,765722912 | NTRK3 | -0,197454909 |
| ENSG0000002284 | gene | 0,872432566 | 0,040821744 | 0,765722912 | RNF10 | -0,196884471 |
| ENSG0000018255 | gene | 0,876232853 | 0,028877699 | 0,735782087 | ADI1 | -0,190613788 |
| ENSG0000014838 | gene | 0,87635289 | 0,028103864 | 0,727417021 | INPP5E | -0,190416163 |
| ENSG0000000983 | gene | 0,878322918 | 0,034398615 | 0,761794317 | POMT2 | -0,187176646 |
| ENSG0000015294 | gene | 0,878589737 | 0,00630263 | 0,560473839 | RAD17 | -0,186738448 |
| ENSG0000019715 | gene | 0,87981523 | 0,04119009 | 0,765859559 | ABCB8 | -0,18472752 |
| ENSG0000016764 | gene | 0,879996647 | 0,017310537 | 0,663077244 | C19orf33 | -0,184430069 |
| ENSG0000023199 | gene | 0,880545124 | 0,045665087 | 0,779356164 | ANXA2P2 | -0,183531157 |
| ENSG0000009026 | gene | 0,881116564 | 0,023944557 | 0,700392548 | NDUFB2 | -0,182595208 |
| ENSG0000021361 | gene | 0,882018325 | 0,001936086 | 0,47003333 | HEXA | -0,181119465 |

| | | | | | | |
|---|---|---|---|---|---|---|
| ENSG0000023302 | gene | 0,882857451 | 0,014435641 | 0,628093865 | NPIPA9 | -0,179747581 |
| ENSG0000021362 | gene | 0,883281402 | 0,046010577 | 0,782616789 | LBH | -0,17905496 |
| ENSG0000016316 | gene | 0,883441856 | 0,008916446 | 0,566479818 | RNF149 | -0,178792908 |
| ENSG0000028497 | gene | 0,883721878 | 0,049758096 | 0,788217613 | BX255925.3 | -0,178335694 |
| ENSG0000014052 | gene | 0,883736255 | 0,016696006 | 0,663077244 | ABHD2 | -0,178312223 |
| ENSG0000017332 | gene | 0,885751132 | 0,015234301 | 0,636410574 | MAP3K11 | -0,175026691 |
| ENSG0000016637 | gene | 0,886564704 | 0,012660848 | 0,605697652 | ATP9B | -0,173702167 |
| ENSG0000016802 | gene | 0,886780616 | 0,018294034 | 0,667092139 | RPSA | -0,173350861 |
| ENSG0000011405 | gene | 0,887694349 | 0,022552942 | 0,700392548 | PCCB | -0,171865081 |
| ENSG0000017844 | gene | 0,887899615 | 0,03957076 | 0,765722912 | COX14 | -0,171531519 |
| ENSG0000010661 | gene | 0,88986864 | 0,012752362 | 0,605697652 | RHEB | -0,16833571 |
| ENSG0000016591 | gene | 0,891304401 | 0,042199912 | 0,767591009 | PACSIN3 | -0,166009865 |
| ENSG0000010012 | gene | 0,891600191 | 0,029694438 | 0,745621897 | EIF3L | -0,16553117 |
| ENSG0000006460 | gene | 0,892284232 | 0,035017532 | 0,761794317 | CTSA | -0,16442475 |
| ENSG0000012831 | gene | 0,893522708 | 0,027765239 | 0,727417021 | TST | -0,1624237 |
| ENSG0000010434 | gene | 0,894232367 | 0,011976032 | 0,599480655 | LAPTM4B | -0,161278329 |
| ENSG0000025874 | gene | 0,894599598 | 0,026130051 | 0,702826025 | AL110504.1 | -0,160685985 |
| ENSG0000012456 | gene | 0,895596403 | 0,016975283 | 0,663077244 | SNRPC | -0,15907936 |
| ENSG0000027854 | gene | 0,896176855 | 0,047122513 | 0,784917185 | ACACA | -0,158144627 |
| ENSG0000013468 | gene | 0,897286131 | 0,049291837 | 0,788217613 | YARS1 | -0,156359983 |
| ENSG0000013590 | gene | 0,89832196 | 0,007536909 | 0,560473839 | DOCK10 | -0,154695492 |
| ENSG0000017314 | gene | 0,898750564 | 0,009145454 | 0,570481883 | MRPL57 | -0,154007323 |
| ENSG0000016580 | gene | 0,899198667 | 0,049768101 | 0,788217613 | ZNF219 | -0,153288198 |
| ENSG0000013174 | gene | 0,89999883 | 0,00984516 | 0,57333908 | STARD3 | -0,152004969 |
| ENSG0000022204 | gene | 0,901264646 | 0,014758826 | 0,630703367 | CYTOR | -0,149977296 |
| ENSG0000026109 | gene | 0,902634571 | 0,030056659 | 0,746496713 | AC141586.3 | -0,147786059 |
| ENSG0000017186 | gene | 0,902651296 | 0,042603214 | 0,767591009 | RPS7 | -0,147759329 |
| ENSG0000007475 | gene | 0,902678678 | 0,024722804 | 0,702826025 | ZZEF1 | -0,147715565 |
| ENSG0000012597 | gene | 0,90341159 | 0,031886184 | 0,750135666 | RALY | -0,146544672 |
| ENSG0000019875 | gene | 0,909989863 | 0,008136244 | 0,560473839 | CDC42BPB | -0,136077621 |
| ENSG0000010537 | gene | 0,912563801 | 0,03271175 | 0,754653699 | NOP53 | -0,132002668 |
| ENSG0000012461 | gene | 0,912873177 | 0,019877673 | 0,682146839 | RPS10 | -0,131513651 |
| ENSG0000016879 | gene | 0,915699791 | 0,011155642 | 0,582432479 | ABHD15 | -0,127053401 |
| ENSG0000014884 | gene | 0,916438114 | 0,009897985 | 0,57333908 | PDCD11 | -0,125890635 |
| ENSG0000010032 | gene | 0,916567898 | 0,049773347 | 0,788217613 | ASCC2 | -0,125686338 |
| ENSG0000014145 | gene | 0,916891354 | 0,040170304 | 0,765722912 | NPC1 | -0,125177302 |
| ENSG0000009982 | gene | 0,918418219 | 0,036144184 | 0,765722912 | HCN2 | -0,122776833 |
| ENSG0000014044 | gene | 0,92181607 | 0,025586617 | 0,702826025 | IGF1R | -0,117449176 |
| ENSG0000011299 | gene | 0,923813383 | 0,041697921 | 0,767591009 | NNT | -0,114326649 |
| ENSG0000011759 | gene | 0,923907273 | 0,037803662 | 0,765722912 | PRDX6 | -0,11418003 |
| ENSG0000025499 | gene | 0,924651993 | 0,031526868 | 0,750135666 | BRK1 | -0,113017608 |
| ENSG0000003301 | gene | 0,926933457 | 0,005788646 | 0,546307419 | ALG1 | -0,109462321 |
| ENSG0000017160 | gene | 0,927071091 | 0,032985975 | 0,754653699 | CLSTN1 | -0,109248121 |
| ENSG0000017423 | gene | 0,92828994 | 0,046300297 | 0,783814537 | PITPNA | -0,107352611 |
| ENSG0000017537 | gene | 0,929162987 | 0,043771673 | 0,771136021 | EIF1AD | -0,105996409 |
| ENSG0000020456 | gene | 0,936171925 | 0,010330278 | 0,575745644 | MRPS18B | -0,095154595 |
| ENSG0000000405 | gene | 0,937695592 | 0,007000489 | 0,560473839 | ARF5 | -0,092808444 |
| ENSG0000013138 | gene | 0,938160363 | 0,033567281 | 0,754653699 | SLC6A6 | -0,092093546 |
| ENSG0000012458 | gene | 0,941148679 | 0,03920324 | 0,765722912 | PEX6 | -0,087505443 |
| ENSG0000013874 | gene | 0,944829358 | 0,013881801 | 0,621226599 | NAAA | -0,081874301 |
| ENSG0000014691 | gene | 0,947591454 | 0,018952512 | 0,673063319 | NCAPG2 | -0,077662908 |
| ENSG0000018325 | gene | 0,951472096 | 0,030318066 | 0,749158036 | PTTG1IP | -0,071766748 |
| ENSG0000028282 | gene | 0,966748422 | 0,028411772 | 0,729502277 | FRG1CP | -0,048787591 |
| ENSG0000018617 | gene | 0,966990782 | 0,00342782 | 0,493619508 | BCL9L | -0,048425957 |
| ENSG0000020558 | gene | 1,031677694 | 0,023143441 | 0,700392548 | HMGN1 | 0,044992329 |
| ENSG0000010142 | gene | 1,046415276 | 0,023347835 | 0,700392548 | CHMP4B | 0,065455508 |
| ENSG0000008829 | gene | 1,066047194 | 0,03504859 | 0,761794317 | EDEM2 | 0,092271307 |

| | | | | | | |
|---|---|---|---|---|---|---|
| ENSG0000015748 | gene | 1,066107005 | 0,021953421 | 0,700392548 | MYO1E | 0,092352248 |
| ENSG0000019745 | gene | 1,067971647 | 0,012838697 | 0,605697652 | HNRNPAB | 0,094873346 |
| ENSG0000000808 | gene | 1,069991807 | 0,048383091 | 0,785311848 | JARID2 | 0,09759975 |
| ENSG0000009113 | gene | 1,073710116 | 0,02348464 | 0,700392548 | LAMB1 | 0,102604542 |
| ENSG0000013921 | gene | 1,076610668 | 0,007767082 | 0,560473839 | AMIGO2 | 0,106496626 |
| ENSG0000016861 | gene | 1,079163816 | 0,04694865 | 0,784665672 | STAT3 | 0,109913881 |
| ENSG0000010211 | gene | 1,082871652 | 0,031350131 | 0,749219164 | EMD | 0,114862256 |
| ENSG0000013817 | gene | 1,08463965 | 0,042138938 | 0,767591009 | ARL3 | 0,117215816 |
| ENSG0000014667 | gene | 1,085311035 | 0,030975495 | 0,749158036 | CDCA5 | 0,118108558 |
| ENSG0000016772 | gene | 1,086741245 | 0,01090899 | 0,575745644 | TSR1 | 0,120008473 |
| ENSG0000027655 | gene | 1,087735141 | 0,014807988 | 0,630703367 | HERC2P2 | 0,12132731 |
| ENSG0000007515 | gene | 1,088725542 | 0,03672353 | 0,765722912 | EIF4G3 | 0,122640309 |
| ENSG0000007441 | gene | 1,09183806 | 0,025305368 | 0,702826025 | CA12 | 0,126758893 |
| ENSG0000012069 | gene | 1,093090079 | 0,036931211 | 0,765722912 | HSPH1 | 0,128412295 |
| ENSG0000011270 | gene | 1,094995068 | 0,030784697 | 0,749158036 | SENP6 | 0,130924372 |
| ENSG0000016616 | gene | 1,096966115 | 0,048175556 | 0,785311848 | BRD7 | 0,133518962 |
| ENSG0000010597 | gene | 1,097765819 | 0,035077304 | 0,761794317 | CAV2 | 0,134570324 |
| ENSG0000008423 | gene | 1,099693276 | 0,010618224 | 0,575745644 | APLP2 | 0,137101187 |
| ENSG0000017155 | gene | 1,101651514 | 0,039289253 | 0,765722912 | BCL2L1 | 0,139667927 |
| ENSG0000016800 | gene | 1,104266847 | 0,008815922 | 0,566479818 | POLR2G | 0,143088843 |
| ENSG0000012087 | gene | 1,105952172 | 0,044208034 | 0,773521117 | DUSP4 | 0,145288996 |
| ENSG0000016468 | gene | 1,108871239 | 0,014759608 | 0,630703367 | FABP5 | 0,149091851 |
| ENSG0000018804 | gene | 1,110702023 | 0,00450808 | 0,510612383 | ARL4C | 0,151471825 |
| ENSG0000008832 | gene | 1,111588169 | 0,040212114 | 0,765722912 | TPX2 | 0,152622384 |
| ENSG0000014351 | gene | 1,111916565 | 0,031976947 | 0,750135666 | TP53BP2 | 0,153048537 |
| ENSG0000006532 | gene | 1,113452779 | 0,021366408 | 0,700392548 | GLP2R | 0,155040375 |
| ENSG0000005808 | gene | 1,1171933 | 0,040933716 | 0,765722912 | LAMC2 | 0,159878826 |
| ENSG0000017904 | gene | 1,117697244 | 0,032960545 | 0,754653699 | RRS1 | 0,160529452 |
| ENSG0000000448 | gene | 1,121092249 | 0,011854425 | 0,597348252 | KDM1A | 0,164904995 |
| ENSG0000012421 | gene | 1,12113476 | 0,018251681 | 0,667092139 | STAU1 | 0,1649597 |
| ENSG0000008598 | gene | 1,121186625 | 0,040337659 | 0,765722912 | USP40 | 0,16502644 |
| ENSG0000007260 | gene | 1,121607513 | 0,01424663 | 0,626651758 | CHFR | 0,165567918 |
| ENSG0000016845 | gene | 1,12361015 | 0,041014892 | 0,765722912 | HR | 0,168141562 |
| ENSG0000016063 | gene | 1,123659507 | 0,048435557 | 0,785311848 | SAFB | 0,168204934 |
| ENSG0000017437 | gene | 1,123841905 | 0,038588941 | 0,765722912 | EXO1 | 0,1684391 |
| ENSG0000011541 | gene | 1,127869558 | 0,042989538 | 0,767591009 | FN1 | 0,173600225 |
| ENSG0000005793 | gene | 1,127997883 | 0,038301966 | 0,765722912 | MTA3 | 0,17376436 |
| ENSG0000013313 | gene | 1,128016265 | 0,030827645 | 0,749158036 | MORC4 | 0,17378787 |
| ENSG0000021012 | gene | 1,130271977 | 0,01797104 | 0,665808244 | MT-TA | 0,17666997 |
| ENSG0000010654 | gene | 1,13131841 | 0,041325343 | 0,765859559 | AGR2 | 0,178005034 |
| ENSG0000016351 | gene | 1,132019263 | 0,017609486 | 0,663077244 | TGFBR2 | 0,178898508 |
| ENSG0000010659 | gene | 1,132340107 | 0,006698947 | 0,560473839 | MRPL32 | 0,179307348 |
| ENSG0000015376 | gene | 1,133118032 | 0,046939148 | 0,784665672 | GTF2E1 | 0,180298149 |
| ENSG0000016552 | gene | 1,134531368 | 0,010556763 | 0,575745644 | ARF6 | 0,182096498 |
| ENSG0000017271 | gene | 1,136703652 | 0,01948171 | 0,680528353 | SLFN11 | 0,184856181 |
| ENSG0000001918 | gene | 1,136712515 | 0,021449794 | 0,700392548 | CYP24A1 | 0,184867429 |
| ENSG0000014683 | gene | 1,136881547 | 0,014495373 | 0,628093865 | TRIM4 | 0,185081946 |
| ENSG0000009990 | gene | 1,139768804 | 0,022048885 | 0,700392548 | RANBP1 | 0,188741211 |
| ENSG0000016645 | gene | 1,141347339 | 0,003587934 | 0,493619508 | ATMIN | 0,190737904 |
| ENSG0000018362 | gene | 1,141392637 | 0,02107691 | 0,700392548 | HMCES | 0,190795162 |
| ENSG0000015566 | gene | 1,14405219 | 0,039440591 | 0,765722912 | PDIA4 | 0,194152868 |
| ENSG0000015801 | gene | 1,145170352 | 0,039150643 | 0,765722912 | BABAM2 | 0,195562225 |
| ENSG0000011948 | gene | 1,145876762 | 0,024776748 | 0,702826025 | MAPKAP1 | 0,196451892 |
| ENSG0000016301 | gene | 1,147791013 | 0,009164368 | 0,570481883 | FBXO41 | 0,198859984 |
| ENSG0000011901 | gene | 1,148747638 | 0,039905077 | 0,765722912 | NDUFB3 | 0,200061895 |
| ENSG0000010794 | gene | 1,149698968 | 0,035880672 | 0,765722912 | BCCIP | 0,201256162 |
| ENSG0000011606 | gene | 1,15223069 | 0,046934456 | 0,784665672 | MSH6 | 0,20442959 |

| | | | | | | |
|---|---|---|---|---|---|---|
| ENSG0000016354 | gene | 1,152475406 | 0,012410814 | 0,605635271 | NUAK2 | 0,204735963 |
| ENSG0000016690 | gene | 1,155637121 | 0,018341427 | 0,667092139 | MRPL16 | 0,208688452 |
| ENSG0000017348 | gene | 1,15608379 | 0,004627113 | 0,510612383 | PTPRM | 0,209245965 |
| ENSG0000012422 | gene | 1,158709772 | 0,033775546 | 0,755747289 | DDX27 | 0,212519252 |
| ENSG0000016693 | gene | 1,159895738 | 0,046491316 | 0,783814537 | DIS3L | 0,213995129 |
| ENSG0000013539 | gene | 1,160642244 | 0,048409991 | 0,785311848 | DNAJC14 | 0,214923344 |
| ENSG0000016668 | gene | 1,160671561 | 0,04030979 | 0,765722912 | BEX3 | 0,214959786 |
| ENSG0000016684 | gene | 1,160852961 | 0,002810516 | 0,493619508 | TERF2IP | 0,215185245 |
| ENSG0000008060 | gene | 1,160885932 | 0,027874022 | 0,727417021 | PUM3 | 0,215226221 |
| ENSG0000016490 | gene | 1,161470613 | 0,047514223 | 0,785311848 | PHAX | 0,215952652 |
| ENSG0000017796 | gene | 1,1627004 | 0,011719263 | 0,597348252 | RIC8A | 0,217479396 |
| ENSG0000012621 | gene | 1,162713641 | 0,037180808 | 0,765722912 | KLC1 | 0,217495827 |
| ENSG0000015531 | gene | 1,163243375 | 0,04259359 | 0,767591009 | USP25 | 0,21815297 |
| ENSG0000013602 | gene | 1,163332134 | 0,034634263 | 0,761794317 | CKAP4 | 0,218263048 |
| ENSG0000017579 | gene | 1,163913488 | 0,03364896 | 0,754653699 | RUVBL1 | 0,218983829 |
| ENSG0000023969 | gene | 1,164321576 | 0,043928744 | 0,771136021 | TNFSF12 | 0,219489574 |
| ENSG0000016334 | gene | 1,164603853 | 0,0314007 | 0,749219164 | CLDN1 | 0,219839297 |
| ENSG0000013101 | gene | 1,165477869 | 0,014296415 | 0,626651758 | AKAP12 | 0,22092161 |
| ENSG0000020426 | gene | 1,166063893 | 0,030732392 | 0,749158036 | COL5A2 | 0,221646841 |
| ENSG0000014667 | gene | 1,166526417 | 0,009599271 | 0,57333908 | IGFBP3 | 0,222218979 |
| ENSG0000011884 | gene | 1,16747368 | 0,023153935 | 0,700392548 | RARRES1 | 0,223390025 |
| ENSG0000014760 | gene | 1,167806814 | 0,033150326 | 0,754653699 | TERF1 | 0,223801634 |
| ENSG0000016634 | gene | 1,168777787 | 0,044665447 | 0,776242869 | CYB5A | 0,225000665 |
| ENSG0000022186 | gene | 1,168898273 | 0,035331841 | 0,764159259 | CEBPD | 0,22514938 |
| ENSG0000019895 | gene | 1,169199458 | 0,045266984 | 0,776242869 | TGM2 | 0,225521066 |
| ENSG0000010404 | gene | 1,171830709 | 0,019651432 | 0,682146839 | ATP8B4 | 0,228764163 |
| ENSG0000013092 | gene | 1,173033678 | 0,047901202 | 0,785311848 | C12orf65 | 0,230244433 |
| ENSG0000013781 | gene | 1,175246543 | 0,014135068 | 0,626651758 | PAQR5 | 0,232963437 |
| ENSG0000012584 | gene | 1,175491347 | 0,007640386 | 0,560473839 | RRBP1 | 0,233263919 |
| ENSG0000004457 | gene | 1,176746543 | 0,016284811 | 0,658924168 | HSPA5 | 0,234803615 |
| ENSG0000023765 | gene | 1,177196797 | 0,020090001 | 0,682146839 | C2orf74 | 0,235355522 |
| ENSG0000011043 | gene | 1,179887008 | 0,025105087 | 0,702826025 | PDHX | 0,238648706 |
| ENSG0000011105 | gene | 1,18052875 | 0,005794425 | 0,546307419 | KRT18 | 0,239433177 |
| ENSG0000009092 | gene | 1,181876005 | 0,007343017 | 0,560473839 | PLEKHG2 | 0,241078685 |
| ENSG0000010056 | gene | 1,182946979 | 0,009246541 | 0,571930932 | PIGH | 0,242385411 |
| ENSG0000012573 | gene | 1,18296156 | 0,017818348 | 0,663077244 | SH2D3A | 0,242403194 |
| ENSG0000015314 | gene | 1,183281334 | 0,017402562 | 0,663077244 | SMARCA5 | 0,242793126 |
| ENSG0000024069 | gene | 1,186633183 | 0,019903583 | 0,682146839 | PNMA2 | 0,246874031 |
| ENSG0000006611 | gene | 1,186758569 | 0,030930818 | 0,749158036 | SMARCD1 | 0,247026466 |
| ENSG0000018034 | gene | 1,187259059 | 0,026241367 | 0,702826025 | TIGD2 | 0,247634765 |
| ENSG0000012043 | gene | 1,187925422 | 0,047761519 | 0,785311848 | TCP1 | 0,248444267 |
| ENSG0000017393 | gene | 1,188257404 | 0,038522934 | 0,765722912 | RBM4 | 0,248847391 |
| ENSG0000013146 | gene | 1,188701867 | 0,016489312 | 0,66168473 | PSME3 | 0,249386925 |
| ENSG0000010101 | gene | 1,189318609 | 0,040684578 | 0,765722912 | CD40 | 0,250135253 |
| ENSG0000015314 | gene | 1,1903743 | 0,023295816 | 0,700392548 | CETN3 | 0,251415284 |
| ENSG0000018275 | gene | 1,190768261 | 0,005502938 | 0,544332311 | PAPPA | 0,251892673 |
| ENSG0000011087 | gene | 1,191211752 | 0,024534094 | 0,702826025 | COQ5 | 0,252429892 |
| ENSG0000016843 | gene | 1,192459802 | 0,005648285 | 0,546307419 | STIP1 | 0,253940634 |
| ENSG0000024991 | gene | 1,193640393 | 0,004243918 | 0,505536224 | PDCD6 | 0,255368263 |
| ENSG0000011084 | gene | 1,193922631 | 0,025470505 | 0,702826025 | PRPF40B | 0,25570935 |
| ENSG0000017223 | gene | 1,19623304 | 0,031212988 | 0,749219164 | PAIP1 | 0,258498471 |
| ENSG0000017971 | gene | 1,19685971 | 0,024841432 | 0,702826025 | PCED1B | 0,259254056 |
| ENSG0000008623 | gene | 1,197108663 | 0,010708992 | 0,575745644 | EIF2AK1 | 0,259554113 |
| ENSG0000017495 | gene | 1,198200856 | 0,035575343 | 0,765722912 | DHX36 | 0,260869769 |
| ENSG0000021008 | gene | 1,200926339 | 0,036648516 | 0,765722912 | MT-RNR2 | 0,264147663 |
| ENSG0000020459 | gene | 1,201279759 | 0,018756749 | 0,669657331 | HLA-E | 0,264572171 |
| ENSG0000019629 | gene | 1,202714265 | 0,035662712 | 0,765722912 | NIF3L1 | 0,266293935 |

| | | | | | | |
|---|---|---|---|---|---|---|
| ENSG0000013147 | gene | 1,204167265 | 0,040167052 | 0,765722912 | PSMC3IP | 0,268035803 |
| ENSG0000014787 | gene | 1,209064282 | 0,040284495 | 0,765722912 | PLIN2 | 0,273890949 |
| ENSG0000011081 | gene | 1,209651345 | 0,007593278 | 0,560473839 | P3H3 | 0,274591283 |
| ENSG0000014563 | gene | 1,210368746 | 0,017691161 | 0,663077244 | PLK2 | 0,27544664 |
| ENSG0000028592 | gene | 1,211478636 | 0,02546473 | 0,702826025 | AC005082.2 | 0,276768963 |
| ENSG0000009201 | gene | 1,215812375 | 0,043473455 | 0,767591009 | PSME1 | 0,281920609 |
| ENSG0000012212 | gene | 1,220836489 | 0,003583431 | 0,493619508 | OCRL | 0,287869988 |
| ENSG0000021370 | gene | 1,2214739 | 0,00679011 | 0,560473839 | HMGB1P10 | 0,288623037 |
| ENSG0000018039 | gene | 1,222859409 | 0,020770829 | 0,697942964 | MCFD2 | 0,290258548 |
| ENSG0000014052 | gene | 1,223583294 | 0,048841347 | 0,787085197 | FANCI | 0,291112315 |
| ENSG0000026712 | gene | 1,22402237 | 0,00311723 | 0,493619508 | AC012615.3 | 0,291629925 |
| ENSG0000014740 | gene | 1,224364977 | 0,036658297 | 0,765722912 | CSGALNACT1 | 0,292033682 |
| ENSG0000024366 | gene | 1,224389525 | 0,010534006 | 0,575745644 | WDR92 | 0,292062607 |
| ENSG0000013433 | gene | 1,225061326 | 0,039554899 | 0,765722912 | LDHA | 0,292853971 |
| ENSG0000011114 | gene | 1,225509565 | 0,033464824 | 0,754653699 | LTA4H | 0,293381744 |
| ENSG0000002360 | gene | 1,225632004 | 0,040296832 | 0,765722912 | SNAPC1 | 0,293525875 |
| ENSG0000016792 | gene | 1,22610484 | 0,022145523 | 0,700392548 | GHDC | 0,294082344 |
| ENSG0000013552 | gene | 1,232624382 | 0,017530687 | 0,663077244 | LTV1 | 0,301733234 |
| ENSG0000013720 | gene | 1,234896159 | 0,048120439 | 0,785311848 | CMTR1 | 0,304389732 |
| ENSG0000013838 | gene | 1,235441315 | 0,037305592 | 0,765722912 | SSB | 0,305026482 |
| ENSG0000006419 | gene | 1,236104825 | 0,000627211 | 0,273995771 | SPA17 | 0,305801093 |
| ENSG0000015646 | gene | 1,240740507 | 0,041081417 | 0,765722912 | UQCRB | 0,311201417 |
| ENSG0000016659 | gene | 1,241950497 | 0,028245909 | 0,727575645 | HSP90B1 | 0,312607671 |
| ENSG0000012680 | gene | 1,242355552 | 0,007977434 | 0,560473839 | HSPA2 | 0,31307812 |
| ENSG0000027285 | gene | 1,24461651 | 0,046430975 | 0,783814537 | AC096772.2 | 0,315701289 |
| ENSG0000024253 | gene | 1,247288447 | 0,037795061 | 0,765722912 | AC007620.2 | 0,318795141 |
| ENSG0000013733 | gene | 1,248098337 | 0,003418671 | 0,493619508 | IER3 | 0,319731608 |
| ENSG0000016498 | gene | 1,24938425 | 0,017427446 | 0,663077244 | PSIP1 | 0,321217248 |
| ENSG0000011889 | gene | 1,250688901 | 0,04625539 | 0,783814537 | EEF2KMT | 0,322722976 |
| ENSG0000016626 | gene | 1,254715944 | 0,007799662 | 0,560473839 | CUL5 | 0,327360788 |
| ENSG0000010798 | gene | 1,254788457 | 0,049880023 | 0,788217613 | DKK1 | 0,327444163 |
| ENSG0000014387 | gene | 1,255071716 | 0,023105158 | 0,700392548 | RHOB | 0,327769804 |
| ENSG0000018568 | gene | 1,256472898 | 0,004320822 | 0,505536224 | PRAME | 0,329379553 |
| ENSG0000001358 | gene | 1,258239206 | 0,049614812 | 0,788217613 | GPRC5A | 0,331406221 |
| ENSG0000021145 | gene | 1,260668691 | 0,043210151 | 0,767591009 | MT-RNR1 | 0,334189179 |
| ENSG0000022441 | gene | 1,260681645 | 0,0233899 | 0,700392548 | HSP90AA2P | 0,334204004 |
| ENSG0000005504 | gene | 1,262154968 | 0,042624285 | 0,767591009 | NOP58 | 0,335889056 |
| ENSG0000015957 | gene | 1,262251772 | 0,009814511 | 0,57333908 | RSPRY1 | 0,335999702 |
| ENSG0000010090 | gene | 1,263833981 | 0,039433159 | 0,765722912 | NFKBIA | 0,337806961 |
| ENSG0000015432 | gene | 1,267783662 | 0,036158963 | 0,765722912 | NEIL2 | 0,342308581 |
| ENSG0000013229 | gene | 1,272642578 | 0,017726841 | 0,663077244 | EFR3A | 0,347827295 |
| ENSG0000013777 | gene | 1,273171118 | 0,026344215 | 0,702826025 | SLTM | 0,348426335 |
| ENSG0000009033 | gene | 1,274417164 | 0,043473901 | 0,767591009 | ICAM1 | 0,349837602 |
| ENSG0000028572 | gene | 1,275809625 | 0,046647667 | 0,784665672 | AL034430.2 | 0,351413068 |
| ENSG0000021312 | gene | 1,279336417 | 0,002009591 | 0,475979056 | TCTEX1D2 | 0,355395688 |
| ENSG0000021441 | gene | 1,285436888 | 0,02853282 | 0,729502277 | BBIP1 | 0,362258779 |
| ENSG0000020532 | gene | 1,285485409 | 0,028164844 | 0,727417021 | SARNP | 0,362313235 |
| ENSG0000010047 | gene | 1,28760916 | 0,008253275 | 0,560473839 | COCH | 0,364694745 |
| ENSG0000019812 | gene | 1,287680047 | 0,025632656 | 0,702826025 | MB | 0,364774168 |
| ENSG0000011049 | gene | 1,288242653 | 0,038907914 | 0,765722912 | MDK | 0,365404365 |
| ENSG0000024048 | gene | 1,289720792 | 0,003125175 | 0,493619508 | SETP14 | 0,367058774 |
| ENSG0000014833 | gene | 1,290623805 | 0,003475149 | 0,493619508 | NTMT1 | 0,368068541 |
| ENSG0000010715 | gene | 1,291207098 | 0,00479017 | 0,516859327 | CA9 | 0,368720414 |
| ENSG0000010203 | gene | 1,292534933 | 0,020553716 | 0,695460397 | SMARCA1 | 0,370203272 |
| ENSG0000020452 | gene | 1,294916943 | 0,013569551 | 0,620457054 | HLA-C | 0,372859565 |
| ENSG0000016700 | gene | 1,298636685 | 0,018130065 | 0,666897976 | PDIA3 | 0,37699787 |
| ENSG0000012818 | gene | 1,303194445 | 0,045203615 | 0,776242869 | DGCR6L | 0,382052359 |

| | | | | | | |
|---|---|---|---|---|---|---|
| ENSG000002366 | gene | 1,30353313 | 0,010036829 | 0,57333908 | PITPNA-AS1 | 0,38242725 |
| ENSG000000890 | gene | 1,304394904 | 0,003946985 | 0,497781395 | ESF1 | 0,383380709 |
| ENSG000001658 | gene | 1,304832525 | 0,001455719 | 0,425554113 | CASP7 | 0,383864649 |
| ENSG000001294 | gene | 1,305663068 | 0,022420792 | 0,700392548 | AJUBA | 0,384782651 |
| ENSG000001690 | gene | 1,306858364 | 0,039121272 | 0,765722912 | HNRNPH1 | 0,386102791 |
| ENSG000001787 | gene | 1,308231858 | 0,034799965 | 0,761794317 | THBD | 0,387618252 |
| ENSG000001578 | gene | 1,314444868 | 0,022990345 | 0,700392548 | SLC37A3 | 0,394453632 |
| ENSG000001554 | gene | 1,316801421 | 0,020000604 | 0,682146839 | NIFK | 0,397037798 |
| ENSG000002719 | gene | 1,317152052 | 0,042327491 | 0,767591009 | AC098614.3 | 0,3974219 |
| ENSG000001825 | gene | 1,317416922 | 0,028005106 | 0,727417021 | FAM104B | 0,397711987 |
| ENSG000001704 | gene | 1,324810742 | 0,042441796 | 0,767591009 | KRT8 | 0,405786275 |
| ENSG000001716 | gene | 1,325823971 | 0,002963873 | 0,493619508 | ZNF274 | 0,406889242 |
| ENSG000000784 | gene | 1,326998114 | 0,005917436 | 0,550470362 | EDN1 | 0,40816632 |
| ENSG000001715 | gene | 1,327007095 | 0,026015137 | 0,702826025 | FGB | 0,408176084 |
| ENSG000001742 | gene | 1,327757345 | 0,048085781 | 0,785311848 | ZBTB4 | 0,40899151 |
| ENSG000001503 | gene | 1,32934585 | 0,038864617 | 0,765722912 | CWC15 | 0,410716493 |
| ENSG000002260 | gene | 1,337912676 | 0,048959812 | 0,787166786 | SEC11B | 0,419983956 |
| ENSG000001375 | gene | 1,338419447 | 0,023651921 | 0,700392548 | RNF121 | 0,420530313 |
| ENSG000001872 | gene | 1,339854429 | 0,00321004 | 0,493619508 | BCAM | 0,422076265 |
| ENSG000001769 | gene | 1,341176473 | 0,033648804 | 0,754653699 | TCIM | 0,423499081 |
| ENSG000002218 | gene | 1,342522307 | 0,004583157 | 0,510612383 | C6orf226 | 0,42494606 |
| ENSG000002671 | gene | 1,348681891 | 0,012240547 | 0,60034321 | AD000091.1 | 0,431550105 |
| ENSG000001688 | gene | 1,352622037 | 0,027584674 | 0,727417021 | CHTF8 | 0,435758763 |
| ENSG000001353 | gene | 1,353009298 | 0,022424069 | 0,700392548 | PHF21A | 0,436171754 |
| ENSG000001456 | gene | 1,353043435 | 0,012821293 | 0,605697652 | SKP2 | 0,436208153 |
| ENSG000000880 | gene | 1,353481197 | 0,042015951 | 0,767591009 | SULT2B1 | 0,436674846 |
| ENSG000001246 | gene | 1,358072297 | 0,006342211 | 0,560473839 | MED20 | 0,441560283 |
| ENSG000002722 | gene | 1,358903244 | 0,048324611 | 0,785311848 | IER3-AS1 | 0,442442737 |
| ENSG000002056 | gene | 1,361057936 | 0,017821353 | 0,663077244 | SMIM11A | 0,44472848 |
| ENSG000001683 | gene | 1,362779913 | 0,023977192 | 0,700392548 | TAP1 | 0,446552588 |
| ENSG000001382 | gene | 1,363069904 | 0,044651463 | 0,776242869 | RBP4 | 0,446859552 |
| ENSG000001226 | gene | 1,363333093 | 0,031979834 | 0,750135666 | MRM2 | 0,447138088 |
| ENSG000000474 | gene | 1,364003624 | 0,024351265 | 0,702826025 | CP | 0,447847478 |
| ENSG000000113 | gene | 1,368216335 | 0,027512742 | 0,727417021 | LARS2 | 0,452296359 |
| ENSG000001813 | gene | 1,368517481 | 0,015346739 | 0,636889676 | DDX60L | 0,452613863 |
| ENSG000001321 | gene | 1,368934397 | 0,01723117 | 0,663077244 | TRIM21 | 0,45305331 |
| ENSG000001961 | gene | 1,37796219 | 0,019812582 | 0,682146839 | TDRD7 | 0,462536303 |
| ENSG000001242 | gene | 1,37896036 | 0,012827486 | 0,605697652 | ZNFX1 | 0,463580985 |
| ENSG000002755 | gene | 1,382450231 | 0,041986253 | 0,767591009 | SNORD116-4 | 0,467227543 |
| ENSG000002621 | gene | 1,394201159 | 0,027659548 | 0,727417021 | AC015912.1 | 0,479438732 |
| ENSG000001259 | gene | 1,395074841 | 0,005735389 | 0,546307419 | ID1 | 0,48034252 |
| ENSG000001059 | gene | 1,398886732 | 0,013657924 | 0,620457054 | GSDME | 0,484279153 |
| ENSG000001060 | gene | 1,401220991 | 0,031012412 | 0,749158036 | ABHD11 | 0,486684505 |
| ENSG000001093 | gene | 1,420815754 | 0,003144237 | 0,493619508 | AREG | 0,506719483 |
| ENSG000001530 | gene | 1,422072138 | 0,042782713 | 0,767591009 | CARHSP1 | 0,507994651 |
| ENSG000001832 | gene | 1,427535313 | 0,031306127 | 0,749219164 | DAZAP2 | 0,513526434 |
| ENSG000002357 | gene | 1,430286995 | 0,03775359 | 0,765722912 | HMGN1P36 | 0,51630466 |
| ENSG000001300 | gene | 1,433373862 | 0,010233417 | 0,575745644 | SAT1 | 0,519414952 |
| ENSG000001472 | gene | 1,435074431 | 0,006415706 | 0,560473839 | RBMX | 0,521125565 |
| ENSG000001210 | gene | 1,437993521 | 0,000648945 | 0,273995771 | TRIM25 | 0,524057175 |
| ENSG000001009 | gene | 1,444544378 | 0,002214827 | 0,487094782 | PSME2 | 0,530614525 |
| ENSG000000390 | gene | 1,447941036 | 0,000759653 | 0,295079662 | CDH1 | 0,534002853 |
| ENSG000000047 | gene | 1,451505046 | 0,040040794 | 0,765722912 | PDK4 | 0,537549588 |
| ENSG000002137 | gene | 1,464343387 | 0,000201246 | 0,139592754 | SERBP1P1 | 0,550253904 |
| ENSG000002266 | gene | 1,470342483 | 0,048873687 | 0,787085197 | TEX41 | 0,556152237 |
| ENSG000002259 | gene | 1,478534462 | 0,022882364 | 0,700392548 | CKS1BP2 | 0,564167871 |
| ENSG000001715 | gene | 1,485658294 | 0,021797648 | 0,700392548 | FGA | 0,571102329 |

| | | | | | |
|---|---|---|---|---|---|
| ENSG0000010992 | gene | 1,489136812 | 0,0137666 | 0,620457054 | FNBP4 | 0,574476305 |
| ENSG0000016942 | gene | 1,492243921 | 0,022548572 | 0,700392548 | CXCL8 | 0,577483377 |
| ENSG0000009999 | gene | 1,50731792 | 0,004739665 | 0,516859327 | SUSD2 | 0,591983739 |
| ENSG0000000643 | gene | 1,508650947 | 0,035144048 | 0,761794317 | MAP3K9 | 0,593259051 |
| ENSG0000016373 | gene | 1,549134805 | 0,007659205 | 0,560473839 | CXCL1 | 0,631462693 |
| ENSG0000023667 | gene | 1,551671777 | 0,000134698 | 0,118913539 | PRKG1-AS1 | 0,633823419 |
| ENSG0000010867 | gene | 1,55674517 | 0,017132433 | 0,663077244 | LGALS3BP | 0,638532803 |
| ENSG0000016113 | gene | 1,556810989 | 0,028146026 | 0,727417021 | USP41 | 0,638593799 |
| ENSG0000012573 | gene | 1,570240462 | 0,033259839 | 0,754653699 | C3 | 0,650985506 |
| ENSG0000019613 | gene | 1,570756179 | 0,01176859 | 0,597348252 | SERPINA3 | 0,651459255 |
| ENSG0000017319 | gene | 1,571785927 | 0,040727704 | 0,765722912 | PARP14 | 0,65240474 |
| ENSG0000002570 | gene | 1,577459369 | 0,003839065 | 0,497082126 | TYMP | 0,657602846 |
| ENSG0000015658 | gene | 1,581032479 | 0,008474698 | 0,566479818 | UBE2L6 | 0,660867005 |
| ENSG0000018248 | gene | 1,590185655 | 0,013013039 | 0,607546277 | WASH6P | 0,669195211 |
| ENSG0000016373 | gene | 1,593583433 | 0,015223856 | 0,636410574 | CXCL3 | 0,672274554 |
| ENSG0000013081 | gene | 1,59634168 | 0,043274812 | 0,767591009 | SHFL | 0,674769479 |
| ENSG0000013864 | gene | 1,60626865 | 0,008037056 | 0,560473839 | HERC5 | 0,683713205 |
| ENSG0000014037 | gene | 1,613183658 | 0,00371066 | 0,493619508 | BCL2A1 | 0,689910696 |
| ENSG0000016373 | gene | 1,614025345 | 0,00811706 | 0,560473839 | CXCL5 | 0,690663234 |
| ENSG0000006706 | gene | 1,630458043 | 0,017667626 | 0,663077244 | SP100 | 0,705277316 |
| ENSG0000024100 | gene | 1,64053329 | 0,00361341 | 0,493619508 | SEPTIN7P6 | 0,714164869 |
| ENSG0000020541 | gene | 1,64346026 | 0,029797748 | 0,745621897 | SAMD9 | 0,716736572 |
| ENSG0000015277 | gene | 1,671074786 | 0,030642948 | 0,749158036 | IFIT5 | 0,7407763 |
| ENSG0000018497 | gene | 1,681517036 | 0,014325681 | 0,626651758 | USP18 | 0,749763396 |
| ENSG0000028703 | gene | 1,714446231 | 0,000800706 | 0,297935015 | AC097709.1 | 0,777742659 |
| ENSG0000011500 | gene | 1,7603319 | 0,01374821 | 0,620457054 | CCL20 | 0,815847466 |
| ENSG0000007524 | gene | 1,774171283 | 0,040759387 | 0,765722912 | GRAMD4 | 0,827145298 |
| ENSG0000012687 | gene | 1,781246251 | 0,044817761 | 0,776242869 | WDR60 | 0,832886978 |
| ENSG0000025015 | gene | 1,783510023 | 0,001692193 | 0,444131997 | ARPC4-TTLL3 | 0,834719324 |
| ENSG0000016432 | gene | 1,7852458 | 0,038086622 | 0,765722912 | CFAP97 | 0,836122724 |
| ENSG0000021007 | gene | 1,788345405 | 0,005235968 | 0,524190612 | MT-TV | 0,838625409 |
| ENSG0000015940 | gene | 1,795780052 | 0,028989103 | 0,735782087 | C1R | 0,844610659 |
| ENSG0000010844 | gene | 1,8127658 | 0,006934586 | 0,560473839 | TRIM16L | 0,858192549 |
| ENSG0000008104 | gene | 1,821326773 | 0,00019559 | 0,139592754 | CXCL2 | 0,864989786 |
| ENSG0000018342 | gene | 1,836750153 | 0,024017168 | 0,700392548 | NPIPA1 | 0,877155395 |
| ENSG0000020499 | gene | 1,84250797 | 0,025346057 | 0,702826025 | SPIRE2 | 0,88167086 |
| ENSG0000017058 | gene | 1,847958232 | 0,041262498 | 0,765859559 | STAT2 | 0,885932149 |
| ENSG0000005533 | gene | 1,852429829 | 0,010684132 | 0,575745644 | EIF2AK2 | 0,889418894 |
| ENSG0000021004 | gene | 1,853024861 | 0,044900061 | 0,776242869 | MT-TF | 0,889882237 |
| ENSG0000012577 | gene | 1,856722152 | 0,01187192 | 0,597348252 | GPCPD1 | 0,89275794 |
| ENSG0000013589 | gene | 1,927353171 | 0,001533765 | 0,425554113 | SP110 | 0,946620958 |
| ENSG0000000254 | gene | 1,940025745 | 0,000695032 | 0,281227497 | LAP3 | 0,956075798 |
| ENSG0000006807 | gene | 1,959136887 | 0,010821084 | 0,575745644 | IFI35 | 0,970218204 |
| ENSG0000010678 | gene | 1,968438831 | 0,001040527 | 0,336818531 | TRIM14 | 0,977051882 |
| ENSG0000018831 | gene | 1,981264471 | 0,003468149 | 0,493619508 | PLSCR1 | 0,986421473 |
| ENSG0000016416 | gene | 2,016431095 | 0,017370894 | 0,663077244 | LSM6 | 1,011804107 |
| ENSG0000010134 | gene | 2,040486939 | 0,022869347 | 0,700392548 | SAMHD1 | 1,028913476 |
| ENSG0000011532 | gene | 2,056018617 | 0,038576944 | 0,765722912 | DOK1 | 1,039853328 |
| ENSG0000014880 | gene | 2,067204244 | 0,012184913 | 0,60034321 | FUOM | 1,047680937 |
| ENSG0000022191 | gene | 2,077111395 | 0,004003031 | 0,498377381 | PPP2R2A | 1,05457859 |
| ENSG0000025865 | gene | 2,092848592 | 0,025920022 | 0,702826025 | TRIM34 | 1,065467943 |
| ENSG0000025836 | gene | 2,101149405 | 0,038817718 | 0,765722912 | RTEL1 | 1,071178751 |
| ENSG0000014208 | gene | 2,1577809 | 0,00457446 | 0,510612383 | IFITM3 | 1,109548381 |
| ENSG0000025745 | gene | 2,186659775 | 0,002205197 | 0,487094782 | AC004551.1 | 1,128728767 |
| ENSG0000006193 | gene | 2,219487327 | 0,030721587 | 0,749158036 | TNK2 | 1,150226471 |
| ENSG0000016749 | gene | 2,223412679 | 0,032432901 | 0,751763165 | GATAD2A | 1,152775746 |
| ENSG0000004242 | gene | 2,300980042 | 0,048482997 | 0,785311848 | MED17 | 1,20224847 |

| | | | | | | |
|---|---|---|---|---|---|---|
| ENSG000001072( | gene | 2,333350346 | 0,011825398 | 0,597348252 | DDX58 | 1,22240294 |
| ENSG0000008912 | gene | 2,363596961 | 8,46528E-05 | 0,092192886 | OAS1 | 1,240984049 |
| ENSG000001638 | gene | 2,389650262 | 0,00087474 | 0,298954595 | DTX3L | 1,256799488 |
| ENSG000001154 | gene | 2,416802548 | 0,000257475 | 0,166689268 | STAT1 | 1,27309961 |
| ENSG00000115 2 | gene | 2,478457286 | 7,30275E-05 | 0,092192886 | IFIH1 | 1,309442395 |
| ENSG000001376 | gene | 2,484516625 | 0,001767185 | 0,451608845 | DDX60 | 1,312965196 |
| ENSG000002857 | gene | 2,494387259 | 0,049526363 | 0,788217613 | BX890604.2 | 1,318685464 |
| ENSG000001113 | gene | 2,587800867 | 0,000390059 | 0,210436714 | OAS3 | 1,371726605 |
| ENSG000001305 | gene | 2,662065559 | 0,000314748 | 0,179795344 | HELZ2 | 1,412546101 |
| ENSG000001161 | gene | 2,749927439 | 0,004075441 | 0,500969672 | DNAJC16 | 1,459393552 |
| ENSG000000772 | gene | 2,847216585 | 0,003499812 | 0,493619508 | IL4R | 1,509552242 |
| ENSG000001963 | gene | 2,923486382 | 0,049231457 | 0,788217613 | MFSD14C | 1,547689871 |
| ENSG000001985 | gene | 2,940647796 | 0,049604428 | 0,788217613 | MMP17 | 1,556134002 |
| ENSG000001056 | gene | 2,940782117 | 0,002509136 | 0,493619508 | TMEM59L | 1,556199898 |
| ENSG000001961 | gene | 2,949093513 | 0,025449586 | 0,702826025 | ZNF250 | 1,56027157 |
| ENSG000001199 | gene | 2,951200758 | 0,002242991 | 0,487094782 | IFIT3 | 1,561302065 |
| ENSG000001677 | gene | 3,013928058 | 0,031900387 | 0,750135666 | HSD11B1L | 1,59164498 |
| ENSG000002139 | gene | 3,053606189 | 0,003405598 | 0,493619508 | IRF9 | 1,610514016 |
| ENSG000002737 | gene | 3,188139641 | 0,014552685 | 0,628093865 | AL137058.2 | 1,672714821 |
| ENSG000002854 | gene | 3,414944736 | 0,002592649 | 0,493619508 | POLR2J3 | 1,771862232 |
| ENSG000001860 | gene | 3,434065909 | 0,025884119 | 0,702826025 | ZNF566 | 1,779917729 |
| ENSG000001443 | gene | 3,442209366 | 0,00595195 | 0,550470362 | CCDC150 | 1,78333485 |
| ENSG000001965( | gene | 3,448024601 | 0,009647437 | 0,57333908 | SULT1A1 | 1,785770068 |
| ENSG000002281( | gene | 3,477577367 | 0,017324422 | 0,663077244 | AL392172.1 | 1,798082611 |
| ENSG000001570 | gene | 3,543260864 | 0,001879868 | 0,468087114 | EXOG | 1,825077684 |
| ENSG000001175 | gene | 3,59060614 | 0,032197381 | 0,751763165 | PTBP2 | 1,84422741 |
| ENSG000001789( | gene | 3,659752756 | 0,005235647 | 0,524190612 | DPY19L3 | 1,871746187 |
| ENSG000001858 | gene | 3,72707252 | 0,003401096 | 0,493619508 | IFITM1 | 1,898042891 |
| ENSG000001757 | gene | 3,87233386 | 0,018990768 | 0,673063319 | LINC01106 | 1,953203343 |
| ENSG000001673 | gene | 4,151084547 | 0,021558176 | 0,700392548 | STIM1 | 2,053488316 |
| ENSG000002071 | gene | 4,151336543 | 0,00442352 | 0,510612383 | SNORD14D | 2,053575894 |
| ENSG000001386 | gene | 4,2951874 | 0,005135044 | 0,524190612 | HERC6 | 2,102721078 |
| ENSG000001382 | gene | 4,504370644 | 0,007789075 | 0,560473839 | DNAJC13 | 2,171325545 |
| ENSG000002591 | gene | 4,577130303 | 0,004873672 | 0,52009038 | AL163636.3 | 2,194443364 |
| ENSG000001384 | gene | 5,11465057 | 0,000828364 | 0,297935015 | PARP9 | 2,354635679 |
| ENSG000001425 | gene | 5,127134404 | 0,00415356 | 0,504190259 | FAM71E1 | 2,358152717 |
| ENSG000002048 | gene | 5,635992295 | 0,009878528 | 0,57333908 | MRPL53 | 2,494669639 |
| ENSG000001650 | gene | 6,017196653 | 0,041030368 | 0,765722912 | FXN | 2,589091507 |
| ENSG000001855( | gene | 6,299655065 | 0,009976856 | 0,57333908 | IRF7 | 2,655272837 |
| ENSG000001659 | gene | 6,482690112 | 3,55723E-05 | 0,086360569 | IFI27 | 2,69659261 |
| ENSG000001857 | gene | 7,759071306 | 9,49366E-05 | 0,092192886 | IFIT1 | 2,955883985 |
| ENSG000002016 | gene | 8,693482056 | 7,56732E-05 | 0,092192886 | SNORD32A | 3,119934145 |
| ENSG000002196 | gene | 8,829039613 | 0,003442486 | 0,493619508 | FAM228B | 3,142256516 |
| ENSG000002760 | gene | 9,032413854 | 0,004311124 | 0,505536224 | AC004542.4 | 3,17511159 |
| ENSG000001876( | gene | 12,18582067 | 1,42232E-05 | 0,067991613 | ISG15 | 3,60713151 |
| ENSG000001576( | gene | 13,20487301 | 1,74401E-05 | 0,067991613 | MX1 | 3,722998522 |
| ENSG000001267( | gene | 22,87539102 | 9,30515E-05 | 0,092192886 | IFI6 | 4,515724499 |

**SARS-CoV_DEGs**

| id | feature | fc | pval | qval | gene_name | de |
|---|---|---|---|---|---|---|
| ENSG00000 | gene | 0,79369111 | 4,0268E-09 | 2,0114E-05 | RPS2P5 | -0,3333504 |
| ENSG00000 | gene | 0,75355492 | 4,0568E-09 | 2,0114E-05 | ATP6V1F | -0,4082154 |
| ENSG00000 | gene | 1,3098708 | 1,0461E-08 | 2,7523E-05 | UGGT2 | 0,38942452 |
| ENSG00000 | gene | 0,49580144 | 1,1102E-08 | 2,7523E-05 | H2AZ1-DT | -1,0121656 |
| ENSG00000 | gene | 2,0038608 | 2,89E-08 | 4,1263E-05 | MT-TG | 1,0027823 |
| ENSG00000 | gene | 1,75250561 | 2,9987E-08 | 4,1263E-05 | MT-TM | 0,80941906 |
| ENSG00000 | gene | 1,38403885 | 3,7928E-08 | 4,1263E-05 | TWF1P1 | 0,46888444 |
| ENSG00000 | gene | 1,18385403 | 3,8243E-08 | 4,1263E-05 | CUL2 | 0,24349121 |
| ENSG00000 | gene | 1,30824336 | 4,2179E-08 | 4,1263E-05 | MAP1B | 0,38763093 |
| ENSG00000 | gene | 1,25715735 | 4,2437E-08 | 4,1263E-05 | ASCC3 | 0,33016523 |
| ENSG00000 | gene | 0,89589574 | 4,5773E-08 | 4,1263E-05 | HNRNPF | -0,1585972 |
| ENSG00000 | gene | 0,82062774 | 6,2748E-08 | 5,185E-05 | POLR2F | -0,2852002 |
| ENSG00000 | gene | 1,61641016 | 6,8709E-08 | 5,2409E-05 | AC105020.6 | 0,69279332 |
| ENSG00000 | gene | 0,7985016 | 1,0282E-07 | 7,2825E-05 | PFDN6 | -0,3246328 |
| ENSG00000 | gene | 1,23217337 | 1,492E-07 | 9,4083E-05 | USP47 | 0,30120526 |
| ENSG00000 | gene | 0,87537307 | 1,5181E-07 | 9,4083E-05 | SSU72 | -0,1920301 |
| ENSG00000 | gene | 1,24692453 | 1,7844E-07 | 9,903E-05 | DNAJC13 | 0,31837414 |
| ENSG00000 | gene | 1,11024637 | 1,7976E-07 | 9,903E-05 | IARS2 | 0,15087985 |
| ENSG00000 | gene | 0,91332152 | 1,9602E-07 | 9,9669E-05 | YBX1 | -0,1308053 |
| ENSG00000 | gene | 0,87274965 | 2,1716E-07 | 9,9669E-05 | DRG1 | -0,1963602 |
| ENSG00000 | gene | 0,62340607 | 2,2501E-07 | 9,9669E-05 | UBB | -0,6817559 |
| ENSG00000 | gene | 0,77639844 | 2,3608E-07 | 9,9669E-05 | RBM8A | -0,3651309 |
| ENSG00000 | gene | 0,49350617 | 2,379E-07 | 9,9669E-05 | IFITM1 | -1,01886 |
| ENSG00000 | gene | 1,18325101 | 2,4123E-07 | 9,9669E-05 | NORAD | 0,24275616 |
| ENSG00000 | gene | 0,86043866 | 2,5605E-07 | 0,00010156 | DDX27 | -0,2168557 |
| ENSG00000 | gene | 1,33734523 | 2,734E-07 | 0,00010427 | TTC37 | 0,41937194 |
| ENSG00000 | gene | 0,84624483 | 3,6064E-07 | 0,00013245 | RBM4 | -0,240853 |
| ENSG00000 | gene | 0,84962358 | 3,8081E-07 | 0,00013392 | MRI1 | -0,2351043 |
| ENSG00000 | gene | 1,18983576 | 4,0439E-07 | 0,00013392 | SMARCA1 | 0,25076244 |
| ENSG00000 | gene | 0,77673984 | 4,0664E-07 | 0,00013392 | PSME2 | -0,3644966 |
| ENSG00000 | gene | 0,71296936 | 4,2824E-07 | 0,00013392 | PQBP1 | -0,488088 |
| ENSG00000 | gene | 1,22269462 | 4,3331E-07 | 0,00013392 | ZNF83 | 0,29006412 |
| ENSG00000 | gene | 0,51586706 | 4,4567E-07 | 0,00013392 | ISG15 | -0,9549288 |
| ENSG00000 | gene | 0,81405607 | 4,6539E-07 | 0,00013573 | MT-ATP8 | -0,2967999 |
| ENSG00000 | gene | 2,02809429 | 4,9283E-07 | 0,00013962 | MT-TS2 | 1,02012473 |
| ENSG00000 | gene | 0,76147769 | 5,1624E-07 | 0,00014219 | FDPS | -0,3931263 |
| ENSG00000 | gene | 1,28467291 | 5,3315E-07 | 0,00014288 | ZNF638 | 0,36140108 |
| ENSG00000 | gene | 1,22548498 | 5,9373E-07 | 0,00015493 | NF1 | 0,29335281 |
| ENSG00000 | gene | 0,82410181 | 7,6627E-07 | 0,00018781 | MRPL14 | -0,2791055 |
| ENSG00000 | gene | 0,75943814 | 7,7087E-07 | 0,00018781 | HLA-A | -0,3969956 |
| ENSG00000 | gene | 1,17265144 | 7,7656E-07 | 0,00018781 | FAM20B | 0,22977425 |
| ENSG00000 | gene | 0,60804001 | 8,9437E-07 | 0,00021116 | IFI35 | -0,7177618 |
| ENSG00000 | gene | 0,7449423 | 1,0171E-06 | 0,00023455 | UBE2L6 | -0,4247994 |
| ENSG00000 | gene | 0,82693754 | 1,0485E-06 | 0,00023629 | NOSIP | -0,2741497 |
| ENSG00000 | gene | 0,74880277 | 1,0792E-06 | 0,0002378 | TMUB2 | -0,4173423 |
| ENSG00000 | gene | 0,83421705 | 1,1381E-06 | 0,00024247 | SAP18 | -0,2615053 |
| ENSG00000 | gene | 0,82112284 | 1,1656E-06 | 0,00024247 | CIB1 | -0,28433 |
| ENSG00000 | gene | 1,32617922 | 1,1737E-06 | 0,00024247 | SMC3 | 0,40727576 |
| ENSG00000 | gene | 1,16481563 | 1,2769E-06 | 0,0002584 | DNAJC10 | 0,22010161 |
| ENSG00000 | gene | 0,8586852 | 1,3053E-06 | 0,00025887 | ANP32B | -0,2197988 |
| ENSG00000 | gene | 0,80177549 | 1,6167E-06 | 0,00030298 | COA6 | -0,3187298 |
| ENSG00000 | gene | 0,85828017 | 1,6424E-06 | 0,00030298 | XRCC1 | -0,2204794 |
| ENSG00000 | gene | 0,86182195 | 1,7126E-06 | 0,00030298 | CCDC124 | -0,2145382 |
| ENSG00000 | gene | 1,15950287 | 1,7193E-06 | 0,00030298 | USP9X | 0,21350639 |
| ENSG00000 | gene | 0,66822184 | 1,7557E-06 | 0,00030298 | AL139095.2 | -0,581601 |

| | | | | | | |
|---|---|---|---|---|---|---|
| ENSG00000' | gene | 1,39218763 | 1,7653E-06 | 0,00030298 | DST | 0,47735366 |
| ENSG00000' | gene | 1,2642156 | 1,7953E-06 | 0,00030298 | C6orf120 | 0,33824253 |
| ENSG00000' | gene | 1,12014149 | 1,7959E-06 | 0,00030298 | ANKMY2 | 0,16368097 |
| ENSG00000' | gene | 1,35968144 | 1,8027E-06 | 0,00030298 | TMTC3 | 0,44326868 |
| ENSG00000' | gene | 1,17665266 | 1,8374E-06 | 0,00030366 | DDX6 | 0,23468851 |
| ENSG00000' | gene | 0,81919876 | 1,8846E-06 | 0,00030635 | PLEKHA4 | -0,2877146 |
| ENSG00000: | gene | 0,73179109 | 1,9873E-06 | 0,00031062 | HLA-H | -0,4504962 |
| ENSG00000: | gene | 2,10721862 | 1,9958E-06 | 0,00031062 | MT-TL2 | 1,07534 |
| ENSG00000' | gene | 1,15747715 | 2,0048E-06 | 0,00031062 | DMTF1 | 0,21098371 |
| ENSG00000' | gene | 0,82788085 | 2,0405E-06 | 0,00031129 | ISG20L2 | -0,2725049 |
| ENSG00000: | gene | 1,83303784 | 2,1882E-06 | 0,00032877 | MT-TR | 0,87423657 |
| ENSG00000' | gene | 1,22336272 | 2,4899E-06 | 0,0003558 | USO1 | 0,29085222 |
| ENSG00000' | gene | 0,89455174 | 2,5034E-06 | 0,0003558 | TUFM | -0,1607632 |
| ENSG00000' | gene | 1,19383097 | 2,5285E-06 | 0,0003558 | ATL3 | 0,25559858 |
| ENSG00000' | gene | 0,88986899 | 2,5351E-06 | 0,0003558 | CREB3 | -0,1683351 |
| ENSG00000' | gene | 0,82665115 | 2,5476E-06 | 0,0003558 | RPS9 | -0,2746495 |
| ENSG00000: | gene | 0,77382858 | 2,6415E-06 | 0,0003638 | HLA-C | -0,3699141 |
| ENSG00000' | gene | 0,83900944 | 2,6989E-06 | 0,00036401 | PES1 | -0,2532411 |
| ENSG00000' | gene | 1,19946607 | 2,7165E-06 | 0,00036401 | RAB3B | 0,26239234 |
| ENSG00000' | gene | 1,25157855 | 2,9966E-06 | 0,00039619 | ROCK2 | 0,32374884 |
| ENSG00000' | gene | 1,21348961 | 3,3363E-06 | 0,00043171 | NOC3L | 0,27916175 |
| ENSG00000' | gene | 0,87782856 | 3,3583E-06 | 0,00043171 | TALDO1 | -0,1879889 |
| ENSG00000' | gene | 1,15352198 | 3,3959E-06 | 0,00043171 | PHAX | 0,20604549 |
| ENSG00000( | gene | 1,2098056 | 3,6229E-06 | 0,00045474 | DHX29 | 0,27477524 |
| ENSG00000( | gene | 0,83437969 | 4,0366E-06 | 0,00050034 | UFD1 | -0,2612241 |
| ENSG00000' | gene | 1,16791301 | 4,3308E-06 | 0,00051546 | ABCD3 | 0,22393282 |
| ENSG00000' | gene | 0,87400382 | 4,3792E-06 | 0,00051546 | ENSA | -0,1942885 |
| ENSG00000' | gene | 0,84775674 | 4,3868E-06 | 0,00051546 | PRPF31 | -0,2382777 |
| ENSG00000' | gene | 1,14050968 | 4,4099E-06 | 0,00051546 | SNAPC3 | 0,18967869 |
| ENSG00000' | gene | 1,19044612 | 4,464E-06 | 0,00051546 | SLF2 | 0,25150233 |
| ENSG00000' | gene | 0,82701428 | 4,4705E-06 | 0,00051546 | PRCC | -0,2740159 |
| ENSG00000' | gene | 0,87383071 | 4,6101E-06 | 0,00052545 | PSMF1 | -0,1945743 |
| ENSG00000' | gene | 0,88490309 | 4,721E-06 | 0,00053198 | MED8 | -0,1764086 |
| ENSG00000( | gene | 1,33239589 | 4,8388E-06 | 0,00053911 | UFL1 | 0,41402281 |
| ENSG00000' | gene | 1,20502781 | 4,9947E-06 | 0,00055031 | USP16 | 0,26906644 |
| ENSG00000' | gene | 0,72906058 | 5,2283E-06 | 0,00056971 | SHFL | -0,4558894 |
| ENSG00000' | gene | 1,37542542 | 5,3959E-06 | 0,00058158 | POSTN | 0,45987792 |
| ENSG00000' | gene | 1,11186343 | 5,5921E-06 | 0,00059213 | COPB1 | 0,1529796 |
| ENSG00000' | gene | 0,90666186 | 5,6132E-06 | 0,00059213 | PTGR1 | -0,1413635 |
| ENSG00000( | gene | 1,28344795 | 5,7998E-06 | 0,00060534 | TPR | 0,36002479 |
| ENSG00000' | gene | 0,80240739 | 5,8605E-06 | 0,00060534 | MRPL9 | -0,3175932 |
| ENSG00000' | gene | 1,53923984 | 5,9817E-06 | 0,00061082 | UACA | 0,62221804 |
| ENSG00000: | gene | 0,68890565 | 6,0478E-06 | 0,00061082 | AC018638.1 | -0,5376217 |
| ENSG00000' | gene | 0,86042347 | 6,0984E-06 | 0,00061082 | PCBP2 | -0,2168812 |
| ENSG00000( | gene | 1,22229726 | 6,4258E-06 | 0,00063719 | PCM1 | 0,28959519 |
| ENSG00000' | gene | 1,26480473 | 7,0278E-06 | 0,00068998 | MOSPD2 | 0,33891466 |
| ENSG00000' | gene | 0,80192771 | 7,2822E-06 | 0,00070794 | CTBP1 | -0,3184559 |
| ENSG00000' | gene | 0,63355873 | 7,4788E-06 | 0,00072 | PARP10 | -0,6584497 |
| ENSG00000( | gene | 1,26111838 | 7,7603E-06 | 0,00073739 | REV3L | 0,33470371 |
| ENSG00000( | gene | 1,18469107 | 7,8082E-06 | 0,00073739 | TOP2B | 0,2445109 |
| ENSG00000' | gene | 1,21237902 | 7,918E-06 | 0,0007407 | NAA15 | 0,2778408 |
| ENSG00000' | gene | 0,85498911 | 8,0483E-06 | 0,00074586 | PSMD13 | -0,226022 |
| ENSG00000' | gene | 0,80355778 | 8,1817E-06 | 0,0007512 | MIIP | -0,3155263 |
| ENSG00000' | gene | 1,35754275 | 8,4665E-06 | 0,00077022 | SPATA21 | 0,44099763 |
| ENSG00000' | gene | 0,81238487 | 8,6357E-06 | 0,00077636 | TAF6 | -0,2997647 |
| ENSG00000' | gene | 1,26177254 | 8,6907E-06 | 0,00077636 | U2SURP | 0,33545186 |
| ENSG00000' | gene | 0,83786689 | 8,7705E-06 | 0,00077636 | PHB | -0,255207 |

| | | | | | | |
|---|---|---|---|---|---|---|
| ENSG00000 | gene | 0,71728924 | 8,8644E-06 | 0,00077636 | AL035427.1 | -0,4793731 |
| ENSG00000 | gene | 0,85703308 | 8,9255E-06 | 0,00077636 | CSNK1D | -0,2225772 |
| ENSG00000 | gene | 0,90774956 | 9,099E-06 | 0,00078457 | DDX23 | -0,1396338 |
| ENSG00000 | gene | 0,80718699 | 9,2201E-06 | 0,00078816 | RAD23A | -0,3090252 |
| ENSG00000 | gene | 0,84295289 | 9,3554E-06 | 0,00079289 | BAX | -0,2464761 |
| ENSG00000 | gene | 1,15167729 | 9,4953E-06 | 0,00079793 | PTPN11 | 0,20373651 |
| ENSG00000 | gene | 1,10991266 | 9,8274E-06 | 0,0008189 | DDX50 | 0,15044615 |
| ENSG00000 | gene | 1,23403331 | 1,0126E-05 | 0,00083674 | ORC3 | 0,30338133 |
| ENSG00000 | gene | 0,88300184 | 1,0574E-05 | 0,0008615 | TWF2 | -0,1795117 |
| ENSG00000 | gene | 1,28827351 | 1,0599E-05 | 0,0008615 | KIF5B | 0,36543892 |
| ENSG00000 | gene | 1,1547811 | 1,0892E-05 | 0,0008781 | RASA1 | 0,2076194 |
| ENSG00000 | gene | 1,24414432 | 1,1374E-05 | 0,00090348 | SMC6 | 0,31515385 |
| ENSG00000 | gene | 0,85510733 | 1,1389E-05 | 0,00090348 | PRKCSH | -0,2258226 |
| ENSG00000 | gene | 1,24864673 | 1,193E-05 | 0,00093414 | CDK6 | 0,32036537 |
| ENSG00000 | gene | 0,84340178 | 1,1964E-05 | 0,00093414 | PPP1CA | -0,245708 |
| ENSG00000 | gene | 0,64192067 | 1,2102E-05 | 0,00093751 | SAMHD1 | -0,6395331 |
| ENSG00000 | gene | 0,87330453 | 1,2817E-05 | 0,00098268 | ELOF1 | -0,1954433 |
| ENSG00000 | gene | 1,26741719 | 1,2883E-05 | 0,00098268 | SMCHD1 | 0,34189149 |
| ENSG00000 | gene | 0,84391727 | 1,3179E-05 | 0,00099253 | TECR | -0,2448265 |
| ENSG00000 | gene | 1,17153099 | 1,3212E-05 | 0,00099253 | IPO8 | 0,22839512 |
| ENSG00000 | gene | 1,64846692 | 1,3401E-05 | 0,0009991 | MT-TW | 0,72112493 |
| ENSG00000 | gene | 1,14884017 | 1,3512E-05 | 0,00099986 | MET | 0,2001781 |
| ENSG00000 | gene | 1,65223919 | 1,4098E-05 | 0,00103555 | RPSAP8 | 0,72442256 |
| ENSG00000 | gene | 0,85021829 | 1,4363E-05 | 0,00104725 | ELK1 | -0,2340948 |
| ENSG00000 | gene | 0,74729292 | 1,4746E-05 | 0,00105111 | TRMT112 | -0,4202542 |
| ENSG00000 | gene | 0,49415538 | 1,481E-05 | 0,00105111 | IFI27 | -1,0169633 |
| ENSG00000 | gene | 1,15579821 | 1,485E-05 | 0,00105111 | KCTD9 | 0,20888954 |
| ENSG00000 | gene | 0,88715012 | 1,4892E-05 | 0,00105111 | PGM1 | -0,1727498 |
| ENSG00000 | gene | 1,42738855 | 1,4946E-05 | 0,00105111 | KIAA1109 | 0,51337811 |
| ENSG00000 | gene | 1,22619146 | 1,5498E-05 | 0,00108182 | CLOCK | 0,29418426 |
| ENSG00000 | gene | 0,87943487 | 1,5703E-05 | 0,00108182 | TMEM9 | -0,1853514 |
| ENSG00000 | gene | 1,32700604 | 1,571E-05 | 0,00108182 | ARHGAP18 | 0,40817493 |
| ENSG00000 | gene | 0,80984282 | 1,5945E-05 | 0,00109043 | C1orf43 | -0,3042862 |
| ENSG00000 | gene | 0,84332266 | 1,6167E-05 | 0,00109805 | FARSA | -0,2458434 |
| ENSG00000 | gene | 0,83765505 | 1,6695E-05 | 0,0011262 | PPIH | -0,2555718 |
| ENSG00000 | gene | 1,24767577 | 1,7472E-05 | 0,00117063 | ICK | 0,31924308 |
| ENSG00000 | gene | 0,82079306 | 1,7824E-05 | 0,00118447 | PLEKHO1 | -0,2849096 |
| ENSG00000 | gene | 1,3344854 | 1,7974E-05 | 0,00118447 | AC004542.2 | 0,41628352 |
| ENSG00000 | gene | 1,26161765 | 1,8037E-05 | 0,00118447 | AKAP11 | 0,33527475 |
| ENSG00000 | gene | 0,87698161 | 1,8396E-05 | 0,00120011 | SCAMP2 | -0,1893815 |
| ENSG00000 | gene | 1,09633451 | 1,8559E-05 | 0,0012028 | LIX1L | 0,13268806 |
| ENSG00000 | gene | 0,78435797 | 1,8805E-05 | 0,00121084 | UBXN6 | -0,3504159 |
| ENSG00000 | gene | 0,86459407 | 1,9469E-05 | 0,00124202 | TRAP1 | -0,2099052 |
| ENSG00000 | gene | 0,89146914 | 1,954E-05 | 0,00124202 | GET3 | -0,1657432 |
| ENSG00000 | gene | 0,83994829 | 1,9737E-05 | 0,00124659 | CDK4 | -0,2516276 |
| ENSG00000 | gene | 1,22265145 | 1,9914E-05 | 0,00124981 | UGGT1 | 0,29001318 |
| ENSG00000 | gene | 0,80091098 | 2,0452E-05 | 0,00127547 | POLR2E | -0,3202862 |
| ENSG00000 | gene | 0,90263887 | 2,1281E-05 | 0,00131039 | TIMM50 | -0,1477792 |
| ENSG00000 | gene | 0,85734588 | 2,1401E-05 | 0,00131039 | GTF2F1 | -0,2220507 |
| ENSG00000 | gene | 0,85117218 | 2,1568E-05 | 0,00131039 | PSMC5 | -0,2324771 |
| ENSG00000 | gene | 1,40886454 | 2,1615E-05 | 0,00131039 | EEA1 | 0,49453291 |
| ENSG00000 | gene | 0,84818285 | 2,1672E-05 | 0,00131039 | UBE2M | -0,2375528 |
| ENSG00000 | gene | 0,91445555 | 2,214E-05 | 0,00133054 | CRTAP | -0,129015 |
| ENSG00000 | gene | 0,85137256 | 2,2429E-05 | 0,0013398 | CSNK1E | -0,2321375 |
| ENSG00000 | gene | 1,29110428 | 2,3059E-05 | 0,00136919 | RANBP2 | 0,36860553 |
| ENSG00000 | gene | 1,24560978 | 2,3707E-05 | 0,00139925 | ERGIC2 | 0,31685218 |
| ENSG00000 | gene | 0,81458779 | 2,3855E-05 | 0,0013997 | TPM3 | -0,2958579 |

| | | | | | | |
|---|---|---|---|---|---|---|
| ENSG00000 | gene | 0,77219132 | 2,4071E-05 | 0,0014025 | PREB | -0,3729698 |
| ENSG00000 | gene | 0,73210309 | 2,4186E-05 | 0,0014025 | EIF1 | -0,4498813 |
| ENSG00000 | gene | 0,87761071 | 2,4966E-05 | 0,00143929 | SH3GL1 | -0,188347 |
| ENSG00000 | gene | 1,21556513 | 2,5383E-05 | 0,00144811 | NBN | 0,28162719 |
| ENSG00000 | gene | 1,27212575 | 2,5411E-05 | 0,00144811 | ITGA4 | 0,34724129 |
| ENSG00000 | gene | 0,84130435 | 2,5678E-05 | 0,00145085 | PRMT5 | -0,2493003 |
| ENSG00000 | gene | 0,83688286 | 2,5907E-05 | 0,00145085 | ENY2 | -0,2569024 |
| ENSG00000 | gene | 1,42327179 | 2,5928E-05 | 0,00145085 | AC020978.2 | 0,50921119 |
| ENSG00000 | gene | 0,81337032 | 2,6044E-05 | 0,00145085 | CCDC12 | -0,2980157 |
| ENSG00000 | gene | 0,64983887 | 2,6403E-05 | 0,00145623 | IFITM3 | -0,6218461 |
| ENSG00000 | gene | 0,87516571 | 2,6434E-05 | 0,00145623 | HRAS | -0,1923719 |
| ENSG00000 | gene | 0,60149714 | 2,8047E-05 | 0,00153654 | BST2 | -0,7333702 |
| ENSG00000 | gene | 0,76100657 | 2,8429E-05 | 0,00154799 | HLA-B | -0,3940192 |
| ENSG00000 | gene | 1,1969867 | 2,8644E-05 | 0,00154799 | PUM3 | 0,25940713 |
| ENSG00000 | gene | 0,85999588 | 2,8724E-05 | 0,00154799 | CDC123 | -0,2175983 |
| ENSG00000 | gene | 0,86199789 | 2,8994E-05 | 0,00155407 | STX10 | -0,2142438 |
| ENSG00000 | gene | 1,14055438 | 2,9835E-05 | 0,00159054 | SF3B1 | 0,18973523 |
| ENSG00000 | gene | 0,84527698 | 3,025E-05 | 0,00160403 | DAZAP1 | -0,2425039 |
| ENSG00000 | gene | 0,54694064 | 3,0478E-05 | 0,00160758 | NECAP1 | -0,8705438 |
| ENSG00000 | gene | 0,87727945 | 3,0934E-05 | 0,00162298 | PFKP | -0,1888916 |
| ENSG00000 | gene | 0,81833342 | 3,1194E-05 | 0,00162797 | BCKDHA | -0,2892393 |
| ENSG00000 | gene | 0,77864895 | 3,1367E-05 | 0,00162843 | ECSIT | -0,3609551 |
| ENSG00000 | gene | 1,26271566 | 3,1825E-05 | 0,00164157 | UTRN | 0,3365298 |
| ENSG00000 | gene | 0,86947614 | 3,2029E-05 | 0,00164157 | LAP3 | -0,2017817 |
| ENSG00000 | gene | 0,87157991 | 3,231E-05 | 0,00164157 | MTA1 | -0,1982951 |
| ENSG00000 | gene | 1,31750118 | 3,262E-05 | 0,00164157 | LRRC40 | 0,39780425 |
| ENSG00000 | gene | 1,25856368 | 3,2674E-05 | 0,00164157 | TAF9B | 0,33177822 |
| ENSG00000 | gene | 0,79660341 | 3,2696E-05 | 0,00164157 | EMD | -0,3280664 |
| ENSG00000 | gene | 1,26130434 | 3,2778E-05 | 0,00164157 | LIMD1-AS1 | 0,33491642 |
| ENSG00000 | gene | 0,89320458 | 3,3338E-05 | 0,00166121 | PSMD4 | -0,1629374 |
| ENSG00000 | gene | 1,21576593 | 3,3725E-05 | 0,00166728 | AP002784.2 | 0,28186549 |
| ENSG00000 | gene | 0,75991784 | 3,3814E-05 | 0,00166728 | VDR | -0,3960846 |
| ENSG00000 | gene | 0,819596 | 3,3964E-05 | 0,00166728 | GTF2A2 | -0,2870152 |
| ENSG00000 | gene | 0,89092657 | 3,4276E-05 | 0,00167431 | LAPTM4A | -0,1666216 |
| ENSG00000 | gene | 1,23509151 | 3,484E-05 | 0,00169351 | ROCK1 | 0,30461794 |
| ENSG00000 | gene | 1,20691815 | 3,5736E-05 | 0,00172858 | USP9Y | 0,27132784 |
| ENSG00000 | gene | 1,24963641 | 3,6665E-05 | 0,00176492 | CD109 | 0,32150839 |
| ENSG00000 | gene | 0,72251541 | 3,7542E-05 | 0,00179549 | PTMS | -0,4688997 |
| ENSG00000 | gene | 0,79809478 | 3,7666E-05 | 0,00179549 | RNF220 | -0,325368 |
| ENSG00000 | gene | 0,85006595 | 3,7844E-05 | 0,00179549 | CEP164 | -0,2343533 |
| ENSG00000 | gene | 0,88824101 | 3,8143E-05 | 0,00180106 | USB1 | -0,1709769 |
| ENSG00000 | gene | 1,23859279 | 3,8615E-05 | 0,00181474 | MAN1A2 | 0,30870195 |
| ENSG00000 | gene | 0,76382967 | 3,9225E-05 | 0,00183469 | HSF1 | -0,3886771 |
| ENSG00000 | gene | 0,83022516 | 4,2749E-05 | 0,00197947 | SHARPIN | -0,2684254 |
| ENSG00000 | gene | 0,92568541 | 4,2882E-05 | 0,00197947 | TMEM50A | -0,1114061 |
| ENSG00000 | gene | 0,8397528 | 4,2919E-05 | 0,00197947 | MFAP2 | -0,2519634 |
| ENSG00000 | gene | 0,88211652 | 4,4374E-05 | 0,00203709 | MLF2 | -0,1809589 |
| ENSG00000 | gene | 1,25280399 | 4,4623E-05 | 0,00203907 | THOC2 | 0,32516071 |
| ENSG00000 | gene | 0,89751702 | 4,5432E-05 | 0,00205904 | NLE1 | -0,1559888 |
| ENSG00000 | gene | 0,65115557 | 4,5475E-05 | 0,00205904 | MYD88 | -0,6189258 |
| ENSG00000 | gene | 0,75384328 | 4,8276E-05 | 0,00217101 | RPP21 | -0,4076635 |
| ENSG00000 | gene | 0,92510882 | 4,8386E-05 | 0,00217101 | GRB2 | -0,112305 |
| ENSG00000 | gene | 1,18306201 | 4,8702E-05 | 0,00217536 | ZBED5 | 0,24252569 |
| ENSG00000 | gene | 1,16077856 | 4,9562E-05 | 0,00220385 | MTREX | 0,21509278 |
| ENSG00000 | gene | 1,12011689 | 5,0742E-05 | 0,00224625 | TOMM70 | 0,1636493 |
| ENSG00000 | gene | 0,88308059 | 5,1001E-05 | 0,00224741 | SLC25A6 | -0,179383 |
| ENSG00000 | gene | 1,26389267 | 5,1478E-05 | 0,00224741 | UBLCP1 | 0,33787395 |

| | | | | | | |
|---|---|---|---|---|---|---|
| ENSG000002 | gene | 1,10652655 | 5,1562E-05 | 0,00224741 | AC022149.1 | 0,14603807 |
| ENSG000001 | gene | 0,85673642 | 5,1675E-05 | 0,00224741 | BSG | -0,2230767 |
| ENSG000001 | gene | 1,23664335 | 5,2171E-05 | 0,00225905 | SEPTIN7 | 0,30642948 |
| ENSG000001 | gene | 0,90634994 | 5,2672E-05 | 0,00227087 | SRSF3 | -0,1418599 |
| ENSG000002 | gene | 0,77192334 | 5,3541E-05 | 0,00229831 | SERF1B | -0,3734705 |
| ENSG000001 | gene | 0,88312351 | 5,3865E-05 | 0,00230226 | COX6A1 | -0,1793129 |
| ENSG000001 | gene | 0,70316755 | 5,4106E-05 | 0,00230263 | FLAD1 | -0,5080596 |
| ENSG000001 | gene | 0,87185879 | 5,5315E-05 | 0,00234279 | PEPD | -0,1978336 |
| ENSG000002 | gene | 0,65213451 | 5,5745E-05 | 0,00234279 | HLA-DPB1 | -0,6167585 |
| ENSG000002 | gene | 0,64599408 | 5,5758E-05 | 0,00234279 | AL136295.5 | -0,6304071 |
| ENSG000001 | gene | 0,79748886 | 5,6051E-05 | 0,00234516 | CARD19 | -0,3264637 |
| ENSG000000 | gene | 0,92024231 | 5,6909E-05 | 0,00236148 | MAPRE3 | -0,1199143 |
| ENSG000001 | gene | 1,19803808 | 5,6917E-05 | 0,00236148 | MICU2 | 0,26067377 |
| ENSG000001 | gene | 0,80758987 | 5,7497E-05 | 0,00237556 | BUD31 | -0,3083053 |
| ENSG000000 | gene | 1,23941611 | 5,8585E-05 | 0,0024105 | SEMA3A | 0,30966063 |
| ENSG000001 | gene | 0,85851144 | 5,9464E-05 | 0,00242896 | CAMTA2 | -0,2200907 |
| ENSG000001 | gene | 1,19401404 | 5,9524E-05 | 0,00242896 | SOCS5 | 0,2558198 |
| ENSG000000 | gene | 1,18041556 | 6,0128E-05 | 0,00244358 | CCAR1 | 0,23929484 |
| ENSG000001 | gene | 0,86985134 | 6,0781E-05 | 0,00246003 | CD63 | -0,2011592 |
| ENSG000001 | gene | 0,34868849 | 6,1905E-05 | 0,00247816 | MX1 | -1,5199894 |
| ENSG000000 | gene | 1,26966176 | 6,1992E-05 | 0,00247816 | PIK3C2A | 0,34444421 |
| ENSG000001 | gene | 1,29402049 | 6,2022E-05 | 0,00247816 | SMARCAD1 | 0,37186046 |
| ENSG000001 | gene | 0,76581683 | 6,2229E-05 | 0,00247816 | CDK2 | -0,3849287 |
| ENSG000001 | gene | 0,8755869 | 6,262E-05 | 0,00248377 | SNRPB | -0,1916777 |
| ENSG000002 | gene | 0,76560931 | 6,3377E-05 | 0,00249665 | TMA7 | -0,3853197 |
| ENSG000001 | gene | 0,85137176 | 6,3449E-05 | 0,00249665 | SRSF7 | -0,2321389 |
| ENSG000001 | gene | 0,90820242 | 6,4039E-05 | 0,00250994 | FBXO7 | -0,1389142 |
| ENSG000001 | gene | 0,77190804 | 6,4299E-05 | 0,00251018 | AGTRAP | -0,3734991 |
| ENSG000001 | gene | 0,83941292 | 6,5624E-05 | 0,00254753 | RIC8A | -0,2525474 |
| ENSG000001 | gene | 1,22184457 | 6,5805E-05 | 0,00254753 | PPIG | 0,28906078 |
| ENSG000001 | gene | 1,1504013 | 6,6189E-05 | 0,00254753 | LRPPRC | 0,20213721 |
| ENSG000001 | gene | 1,38368267 | 6,6402E-05 | 0,00254753 | ECT2 | 0,46851311 |
| ENSG000001 | gene | 0,81893553 | 6,654E-05 | 0,00254753 | ZMAT5 | -0,2881782 |
| ENSG000001 | gene | 1,16174501 | 6,7856E-05 | 0,00258792 | SHLD2 | 0,21629344 |
| ENSG000002 | gene | 1,55185176 | 6,9468E-05 | 0,00263925 | IGFBP7-AS1 | 0,63399075 |
| ENSG000001 | gene | 0,86530213 | 7,0634E-05 | 0,00267331 | ALG12 | -0,2087241 |
| ENSG000001 | gene | 0,80525652 | 7,1279E-05 | 0,00267642 | NFATC2IP | -0,3124797 |
| ENSG000002 | gene | 1,31736947 | 7,1338E-05 | 0,00267642 | CHML | 0,39766002 |
| ENSG000001 | gene | 0,48912162 | 7,1526E-05 | 0,00267642 | IFIT1 | -1,0317349 |
| ENSG000001 | gene | 0,72434677 | 7,204E-05 | 0,00268551 | AKT1 | -0,4652476 |
| ENSG000000 | gene | 1,40803918 | 7,5879E-05 | 0,00281802 | ATRX | 0,49368748 |
| ENSG000001 | gene | 0,88722559 | 7,6586E-05 | 0,00282983 | ZMAT2 | -0,1726271 |
| ENSG000002 | gene | 0,8261019 | 7,6767E-05 | 0,00282983 | UBA52 | -0,2756083 |
| ENSG000000 | gene | 0,84411476 | 7,9052E-05 | 0,00290326 | CD9 | -0,2444889 |
| ENSG000001 | gene | 0,68177268 | 7,9971E-05 | 0,00292617 | KSR1 | -0,5526373 |
| ENSG000002 | gene | 0,51060263 | 8,0957E-05 | 0,00294853 | FP236383.5 | -0,9697271 |
| ENSG000001 | gene | 0,8505547 | 8,1208E-05 | 0,00294853 | NOB1 | -0,2335241 |
| ENSG000001 | gene | 0,83939509 | 8,1474E-05 | 0,00294853 | EIF3G | -0,2525781 |
| ENSG000002 | gene | 1,18426241 | 8,196E-05 | 0,00295534 | EIF4EP2 | 0,24398879 |
| ENSG000000 | gene | 0,86011932 | 8,2482E-05 | 0,00295815 | NSFL1C | -0,2173913 |
| ENSG000001 | gene | 0,92709007 | 8,2927E-05 | 0,00295815 | NDUFA10 | -0,1092186 |
| ENSG000001 | gene | 0,91335786 | 8,2933E-05 | 0,00295815 | PSMC3 | -0,1307479 |
| ENSG000001 | gene | 1,28724461 | 8,3391E-05 | 0,00296382 | SMC5 | 0,36428623 |
| ENSG000002 | gene | 0,85791359 | 8,3861E-05 | 0,00296988 | SRA1 | -0,2210958 |
| ENSG000001 | gene | 0,80501042 | 8,4509E-05 | 0,00298219 | TMEM258 | -0,3129206 |
| ENSG000001 | gene | 0,87272008 | 8,5634E-05 | 0,00300771 | PRDX1 | -0,1964091 |
| ENSG000000 | gene | 0,83973487 | 8,5839E-05 | 0,00300771 | STRN4 | -0,2519942 |

| | | | | | | |
|---|---|---|---|---|---|---|
| ENSG00000 | gene | 1,37764873 | 8,6179E-05 | 0,00300899 | KIF18A | 0,46220808 |
| ENSG00000 | gene | 0,84195604 | 8,6814E-05 | 0,00301881 | TFIP11 | -0,2481832 |
| ENSG00000 | gene | 0,86254868 | 8,7069E-05 | 0,00301881 | TKT | -0,2133222 |
| ENSG00000 | gene | 1,09501067 | 8,7722E-05 | 0,00303084 | WASHC5 | 0,13094493 |
| ENSG00000 | gene | 1,16046341 | 9,0011E-05 | 0,00309914 | CAPZA1 | 0,21470103 |
| ENSG00000( | gene | 0,87086686 | 9,0819E-05 | 0,00311436 | PKM | -0,1994759 |
| ENSG00000 | gene | 0,84144161 | 9,1082E-05 | 0,00311436 | VPS4A | -0,2490649 |
| ENSG00000 | gene | 1,14979153 | 9,2239E-05 | 0,00313049 | IKBIP | 0,2013723 |
| ENSG00000 | gene | 0,87929916 | 9,2646E-05 | 0,00313049 | POFUT2 | -0,185574 |
| ENSG00000 | gene | 1,17839851 | 9,2678E-05 | 0,00313049 | KLHL9 | 0,23682752 |
| ENSG00000 | gene | 1,1942844 | 9,2816E-05 | 0,00313049 | NUP107 | 0,25614644 |
| ENSG00000 | gene | 0,68082283 | 9,4083E-05 | 0,00315343 | UBALD1 | -0,5546487 |
| ENSG00000( | gene | 1,06931909 | 9,4298E-05 | 0,00315343 | TXLNA | 0,09669243 |
| ENSG00000 | gene | 0,87685116 | 9,4776E-05 | 0,00315343 | USP11 | -0,1895961 |
| ENSG00000 | gene | 0,7915802 | 9,4925E-05 | 0,00315343 | NDUFS7 | -0,3371926 |
| ENSG00000 | gene | 0,83728623 | 9,5086E-05 | 0,00315343 | MRPL51 | -0,2562072 |
| ENSG00000 | gene | 1,21469694 | 9,5735E-05 | 0,00315623 | IFT81 | 0,28059642 |
| ENSG00000; | gene | 1,22327405 | 9,5807E-05 | 0,00315623 | PRKDC | 0,29074765 |
| ENSG00000 | gene | 0,87576811 | 9,7063E-05 | 0,00318701 | SCPEP1 | -0,1913792 |
| ENSG00000 | gene | 1,14903025 | 9,7394E-05 | 0,00318734 | NOL11 | 0,20041677 |
| ENSG00000 | gene | 1,14235015 | 9,8314E-05 | 0,0031985 | ZNF106 | 0,19200493 |
| ENSG00000( | gene | 0,87719706 | 9,8963E-05 | 0,0031985 | SMG6 | -0,1890271 |
| ENSG00000 | gene | 1,52741217 | 9,8967E-05 | 0,0031985 | KIF20B | 0,61108942 |
| ENSG00000 | gene | 0,90992537 | 9,9026E-05 | 0,0031985 | TMEM59 | -0,1361799 |
| ENSG00000 | gene | 0,87082866 | 0,00010041 | 0,00322412 | PMM1 | -0,1995392 |
| ENSG00000 | gene | 1,47352865 | 0,00010047 | 0,00322412 | CENPE | 0,55927511 |
| ENSG00000 | gene | 1,11783199 | 0,00010212 | 0,003266 | EPRS1 | 0,16070336 |
| ENSG00000 | gene | 1,35282286 | 0,00010316 | 0,003266 | LHPP | 0,43597294 |
| ENSG00000 | gene | 0,92276696 | 0,00010322 | 0,003266 | MTHFD1L | -0,1159618 |
| ENSG00000 | gene | 0,6421703 | 0,00010365 | 0,003266 | COA1 | -0,6389721 |
| ENSG00000 | gene | 1,47434245 | 0,00010374 | 0,003266 | SGO2 | 0,56007166 |
| ENSG00000( | gene | 1,22049564 | 0,00010375 | 0,003266 | IBTK | 0,28746714 |
| ENSG00000 | gene | 1,23178074 | 0,00010414 | 0,00326796 | SCYL2 | 0,30074547 |
| ENSG00000 | gene | 0,59414623 | 0,00010587 | 0,0033118 | PLSCR1 | -0,7511101 |
| ENSG00000 | gene | 0,79969575 | 0,00010675 | 0,0033286 | H3-3B | -0,3224769 |
| ENSG00000 | gene | 0,31775337 | 0,00010747 | 0,00334062 | IFI6 | -1,6540207 |
| ENSG00000 | gene | 1,46077204 | 0,0001098 | 0,00340254 | CENPF | 0,54673106 |
| ENSG00000 | gene | 0,85820815 | 0,00011223 | 0,00343546 | CCDC86 | -0,2206005 |
| ENSG00000 | gene | 0,88124566 | 0,00011253 | 0,00343546 | TMBIM1 | -0,1823839 |
| ENSG00000 | gene | 0,78745567 | 0,00011288 | 0,00343546 | CLN6 | -0,3447294 |
| ENSG00000 | gene | 0,88100879 | 0,00011291 | 0,00343546 | ERH | -0,1827717 |
| ENSG00000; | gene | 0,80677153 | 0,00011301 | 0,00343546 | UQCRHL | -0,3097679 |
| ENSG00000 | gene | 0,80819818 | 0,00011313 | 0,00343546 | B4GALT3 | -0,307219 |
| ENSG00000 | gene | 0,85353341 | 0,00011329 | 0,00343546 | APH1A | -0,2284805 |
| ENSG00000 | gene | 0,84275977 | 0,00011456 | 0,00346153 | STX4 | -0,2468066 |
| ENSG00000 | gene | 1,11156129 | 0,00011489 | 0,00346153 | STK17A | 0,1525875 |
| ENSG00000 | gene | 0,72809339 | 0,0001152 | 0,00346153 | LAMTOR5 | -0,4578046 |
| ENSG00000 | gene | 0,92947585 | 0,00011625 | 0,00348254 | TXNDC12 | -0,1055107 |
| ENSG00000 | gene | 0,88268557 | 0,00011753 | 0,00350515 | EIF3D | -0,1800285 |
| ENSG00000 | gene | 0,86683698 | 0,0001178 | 0,00350515 | KRT10 | -0,2061674 |
| ENSG00000 | gene | 0,87154687 | 0,00011868 | 0,00350515 | ATP6V1B2 | -0,1983498 |
| ENSG00000( | gene | 0,79355041 | 0,0001188 | 0,00350515 | FDFT1 | -0,3336062 |
| ENSG00000 | gene | 0,90749756 | 0,00011897 | 0,00350515 | RAMAC | -0,1400343 |
| ENSG00000 | gene | 1,16785988 | 0,00011924 | 0,00350515 | IDE | 0,2238672 |
| ENSG00000; | gene | 1,1834053 | 0,00011964 | 0,00350515 | AP002383.2 | 0,24294426 |
| ENSG00000 | gene | 0,87377728 | 0,00011983 | 0,00350515 | TIMM44 | -0,1946625 |
| ENSG00000 | gene | 0,83955905 | 0,00012032 | 0,00350924 | AAMP | -0,2522963 |

| | | | | | | |
|---|---|---|---|---|---|---|
| ENSG00000 | gene | 0,80314702 | 0,00012079 | 0,00351251 | RPS3 | -0,316264 |
| ENSG00002 | gene | 1,38472029 | 0,00012268 | 0,00354334 | MT-TI | 0,46959458 |
| ENSG00000 | gene | 1,10597319 | 0,00012276 | 0,00354334 | CTTNBP2NL | 0,14531641 |
| ENSG00000 | gene | 0,87259825 | 0,00012292 | 0,00354334 | ASMTL | -0,1966105 |
| ENSG00000 | gene | 0,88393996 | 0,00012511 | 0,00358559 | BET1L | -0,1779797 |
| ENSG00000 | gene | 0,9149381 | 0,00012511 | 0,00358559 | PSMA7 | -0,128254 |
| ENSG00000 | gene | 0,8222051 | 0,00012665 | 0,00360363 | DEDD | -0,2824298 |
| ENSG00000 | gene | 0,73108868 | 0,0001269 | 0,00360363 | KCP | -0,4518817 |
| ENSG00002 | gene | 1,24635204 | 0,00012716 | 0,00360363 | GTF2IRD1P | 0,31771163 |
| ENSG00002 | gene | 0,73895673 | 0,0001272 | 0,00360363 | PSMB9 | -0,4364382 |
| ENSG00000 | gene | 0,87929882 | 0,00012859 | 0,00363283 | FAM118B | -0,1855746 |
| ENSG00000 | gene | 1,11780323 | 0,00013022 | 0,00366821 | NAE1 | 0,16066626 |
| ENSG00000 | gene | 0,84313396 | 0,00013058 | 0,00366821 | RABEPK | -0,2461662 |
| ENSG00000 | gene | 1,13712815 | 0,00013136 | 0,00367968 | MACF1 | 0,18539484 |
| ENSG00000 | gene | 0,67785254 | 0,00013234 | 0,00368949 | JSRP1 | -0,5609566 |
| ENSG00000 | gene | 0,85835919 | 0,0001328 | 0,00368949 | AKR1A1 | -0,2203466 |
| ENSG00000 | gene | 1,14266924 | 0,00013283 | 0,00368949 | BBS7 | 0,19240785 |
| ENSG00000 | gene | 0,89900563 | 0,00013352 | 0,00369835 | PRMT1 | -0,153598 |
| ENSG00000 | gene | 0,87001707 | 0,00013417 | 0,00370592 | PSMA5 | -0,2008844 |
| ENSG00000 | gene | 0,87005822 | 0,00013611 | 0,00374918 | SCNM1 | -0,2008162 |
| ENSG00002 | gene | 0,73035948 | 0,00013771 | 0,00378272 | AL138963.4 | -0,4533214 |
| ENSG00002 | gene | 1,40919235 | 0,00013896 | 0,00378962 | AC006369.1 | 0,49486855 |
| ENSG00000 | gene | 0,821514 | 0,00013962 | 0,00378962 | CCDC130 | -0,2836429 |
| ENSG00000 | gene | 0,86788504 | 0,00013963 | 0,00378962 | POLD2 | -0,2044241 |
| ENSG00000 | gene | 0,72138348 | 0,00013969 | 0,00378962 | DNAJB2 | -0,4711617 |
| ENSG00000 | gene | 1,12329462 | 0,00013987 | 0,00378962 | EIF2S3 | 0,16773637 |
| ENSG00000 | gene | 1,14010017 | 0,00014054 | 0,00379733 | SMARCA5 | 0,18916059 |
| ENSG00000 | gene | 0,87643107 | 0,00014213 | 0,00382145 | BABAM1 | -0,1902875 |
| ENSG00000 | gene | 0,72333241 | 0,00014221 | 0,00382145 | IER2 | -0,4672693 |
| ENSG00002 | gene | 1,32716544 | 0,00014436 | 0,00386333 | AL159978.1 | 0,40834822 |
| ENSG00000 | gene | 0,87574938 | 0,00014454 | 0,00386333 | PROCR | -0,19141 |
| ENSG00000 | gene | 0,7645601 | 0,00014508 | 0,0038673 | ANKRD13D | -0,3872982 |
| ENSG00000 | gene | 0,83887 | 0,00014581 | 0,00387624 | TCF7L1 | -0,2534808 |
| ENSG00000 | gene | 0,87952464 | 0,0001469 | 0,00388079 | BAP1 | -0,1852041 |
| ENSG00000 | gene | 0,87950283 | 0,00014704 | 0,00388079 | DRAP1 | -0,1852399 |
| ENSG00000 | gene | 0,84815699 | 0,00014715 | 0,00388079 | EIF4E2 | -0,2375968 |
| ENSG00002 | gene | 1,3576965 | 0,0001478 | 0,00388594 | GAS6-AS1 | 0,44116101 |
| ENSG00002 | gene | 0,74882203 | 0,00014849 | 0,00388594 | MCRIP1 | -0,4173052 |
| ENSG00000 | gene | 1,29245451 | 0,00014871 | 0,00388594 | STAG2 | 0,37011351 |
| ENSG00000 | gene | 0,88047759 | 0,00014892 | 0,00388594 | TRA2B | -0,1836418 |
| ENSG00000 | gene | 1,19931068 | 0,00014994 | 0,00389314 | ABCE1 | 0,26220544 |
| ENSG00000 | gene | 0,90089436 | 0,00014998 | 0,00389314 | HNRNPM | -0,1505702 |
| ENSG00000 | gene | 0,76262167 | 0,0001527 | 0,00395336 | HCFC1R1 | -0,3909606 |
| ENSG00000 | gene | 0,85262376 | 0,00015331 | 0,00395891 | PSMD9 | -0,2300188 |
| ENSG00000 | gene | 0,65802317 | 0,00015399 | 0,00396614 | LY6E | -0,6037897 |
| ENSG00002 | gene | 0,85798797 | 0,00015577 | 0,00400164 | FBXO17 | -0,2209707 |
| ENSG00000 | gene | 0,76608129 | 0,00015755 | 0,00403676 | STAT2 | -0,3844306 |
| ENSG00000 | gene | 1,13524282 | 0,00015941 | 0,00407402 | RFC1 | 0,18300092 |
| ENSG00000 | gene | 0,84846713 | 0,0001599 | 0,00407589 | EXOSC1 | -0,2370693 |
| ENSG00000 | gene | 0,84910326 | 0,00016037 | 0,00407686 | PPP5C | -0,2359881 |
| ENSG00000 | gene | 0,81065952 | 0,00016108 | 0,00407686 | KRTCAP2 | -0,302832 |
| ENSG00000 | gene | 0,87768586 | 0,00016126 | 0,00407686 | ELOVL1 | -0,1882234 |
| ENSG00000 | gene | 0,85305931 | 0,00016169 | 0,00407686 | ARHGDIA | -0,2292821 |
| ENSG00000 | gene | 0,88503699 | 0,00016199 | 0,00407686 | RBM6 | -0,1761903 |
| ENSG00000 | gene | 1,20057426 | 0,00016299 | 0,00409176 | ARFGEF1 | 0,26372465 |
| ENSG00000 | gene | 1,18092727 | 0,00016458 | 0,00410562 | ZFC3H1 | 0,23992012 |
| ENSG00000 | gene | 0,88358345 | 0,00016512 | 0,00410562 | GLB1 | -0,1785617 |

| | | | | | | |
|---|---|---|---|---|---|---|
| ENSG00000' | gene | 0,83766802 | 0,00016516 | 0,00410562 | MRPL21 | -0,2555495 |
| ENSG00000' | gene | 1,12754504 | 0,00016589 | 0,00410562 | GNPTAB | 0,17318506 |
| ENSG00000' | gene | 0,7419038 | 0,0001659 | 0,00410562 | STAT1 | -0,430696 |
| ENSG00000' | gene | 0,91016072 | 0,00016656 | 0,00410562 | R3HDM4 | -0,1358068 |
| ENSG00000( | gene | 1,18255621 | 0,00016663 | 0,00410562 | SCFD1 | 0,24190876 |
| ENSG00000; | gene | 1,3722992 | 0,00016686 | 0,00410562 | TRHDE-AS1 | 0,45659506 |
| ENSG00000' | gene | 0,76231892 | 0,00016842 | 0,00413099 | TAP1 | -0,3915334 |
| ENSG00000( | gene | 0,83291662 | 0,00016872 | 0,00413099 | TRAPPC3 | -0,263756 |
| ENSG00000' | gene | 0,76413735 | 0,00017077 | 0,00417083 | USP18 | -0,3880961 |
| ENSG00000; | gene | 0,84246231 | 0,00017143 | 0,0041751 | FAM174C | -0,2473159 |
| ENSG00000' | gene | 1,18481234 | 0,00017179 | 0,0041751 | NEMF | 0,24465857 |
| ENSG00000; | gene | 1,58774257 | 0,00017282 | 0,00418984 | AF230666.1 | 0,66697702 |
| ENSG00000; | gene | 1,5128389 | 0,00017518 | 0,0042353 | TMSB4XP4 | 0,59725837 |
| ENSG00000' | gene | 0,90840888 | 0,00017555 | 0,0042353 | KPNB1 | -0,1385863 |
| ENSG00000' | gene | 0,81698198 | 0,00017682 | 0,00425569 | POLR2H | -0,2916238 |
| ENSG00000' | gene | 1,14282998 | 0,0001776 | 0,00426419 | VPS13B | 0,19261079 |
| ENSG00000' | gene | 0,84877192 | 0,00017987 | 0,00429085 | MTMR3 | -0,2365512 |
| ENSG00000' | gene | 0,86364453 | 0,00017999 | 0,00429085 | TMEM45A | -0,2114905 |
| ENSG00000' | gene | 1,1382988 | 0,00018001 | 0,00429085 | MMGT1 | 0,1868793 |
| ENSG00000' | gene | 0,78100241 | 0,00018116 | 0,00430778 | UBXN1 | -0,3566011 |
| ENSG00000; | gene | 0,84899693 | 0,00018204 | 0,00431846 | CPNE1 | -0,2361687 |
| ENSG00000' | gene | 0,86300592 | 0,00018297 | 0,00433023 | MAP2K2 | -0,2125576 |
| ENSG00000; | gene | 1,38801628 | 0,00018463 | 0,00434655 | AC092567.1 | 0,47302449 |
| ENSG00000; | gene | 0,77657977 | 0,00018481 | 0,00434655 | ARL16 | -0,364794 |
| ENSG00000' | gene | 0,92025497 | 0,00018498 | 0,00434655 | CCT7 | -0,1198945 |
| ENSG00000; | gene | 0,84380554 | 0,00018683 | 0,00437962 | MT-TN | -0,2450175 |
| ENSG00000' | gene | 0,85110385 | 0,00018957 | 0,0044335 | EIF5A | -0,2325929 |
| ENSG00000' | gene | 0,8232177 | 0,00019069 | 0,00444922 | CDC37 | -0,2806541 |
| ENSG00000' | gene | 0,88709401 | 0,00019463 | 0,0045258 | STOML2 | -0,1728411 |
| ENSG00000; | gene | 0,84243563 | 0,00019489 | 0,0045258 | MICAL3 | -0,2473616 |
| ENSG00000' | gene | 0,89448851 | 0,00019542 | 0,00452745 | DDX56 | -0,1608651 |
| ENSG00000; | gene | 1,40815829 | 0,00019641 | 0,00453997 | AC018521.2 | 0,49380952 |
| ENSG00000' | gene | 0,70845506 | 0,00019836 | 0,00456946 | POLR3C | -0,4972518 |
| ENSG00000' | gene | 0,83285048 | 0,00019861 | 0,00456946 | NTMT1 | -0,2638706 |
| ENSG00000' | gene | 0,86204709 | 0,00019922 | 0,00457289 | PSMC1 | -0,2141614 |
| ENSG00000( | gene | 0,86660715 | 0,00020061 | 0,004594 | NANS | -0,20655 |
| ENSG00000' | gene | 0,88505784 | 0,00020171 | 0,00460868 | CSTB | -0,1761564 |
| ENSG00000' | gene | 0,88233262 | 0,00020282 | 0,00462336 | TSC22D4 | -0,1806055 |
| ENSG00000' | gene | 0,69090457 | 0,0002038 | 0,00462521 | SAT2 | -0,5334416 |
| ENSG00000' | gene | 0,86653696 | 0,00020399 | 0,00462521 | ADPRHL2 | -0,2066668 |
| ENSG00000; | gene | 0,84545573 | 0,00020457 | 0,00462521 | PBX2 | -0,2421989 |
| ENSG00000' | gene | 0,84320028 | 0,00020477 | 0,00462521 | P4HB | -0,2460528 |
| ENSG00000' | gene | 0,89551145 | 0,00020664 | 0,00465681 | YWHAE | -0,1592162 |
| ENSG00000' | gene | 0,89950125 | 0,00020844 | 0,00468679 | DNAJB12 | -0,1528028 |
| ENSG00000; | gene | 1,14630048 | 0,00021071 | 0,00472713 | HERC2P2 | 0,19698526 |
| ENSG00000' | gene | 1,21944365 | 0,00021171 | 0,00473895 | USP1 | 0,28622309 |
| ENSG00000; | gene | 0,82636583 | 0,00021237 | 0,00474292 | ARL2 | -0,2751475 |
| ENSG00000' | gene | 1,237504 | 0,0002146 | 0,00478206 | COPS2 | 0,30743319 |
| ENSG00000' | gene | 0,72145997 | 0,00021724 | 0,00482989 | PPP1R18 | -0,4710087 |
| ENSG00000; | gene | 1,54471367 | 0,00021986 | 0,00487724 | CATSPER2P | 0,62733945 |
| ENSG00000' | gene | 0,88594876 | 0,00022218 | 0,0049177 | ATP1A1 | -0,1747048 |
| ENSG00000' | gene | 0,88205287 | 0,00022313 | 0,00491818 | SLC31A1 | -0,181063 |
| ENSG00000; | gene | 1,75394403 | 0,00022319 | 0,00491818 | AC105105.1 | 0,81060271 |
| ENSG00000; | gene | 0,90153306 | 0,00022401 | 0,00492521 | BRK1 | -0,1495477 |
| ENSG00000' | gene | 1,14496091 | 0,00022947 | 0,00501815 | TTC21B | 0,19529835 |
| ENSG00000; | gene | 1,20110479 | 0,0002298 | 0,00501815 | CEBPZOS | 0,26436202 |
| ENSG00000' | gene | 0,83904009 | 0,00023047 | 0,00501815 | GAPDH | -0,2531883 |

| | | | | | | |
|---|---|---|---|---|---|---|
| ENSG00000( | gene | 0,75238071 | 0,00023082 | 0,00501815 | USE1 | -0,4104652 |
| ENSG00002 | gene | 0,89577563 | 0,00023114 | 0,00501815 | TAPBP | -0,1587907 |
| ENSG00002 | gene | 1,45797418 | 0,00023127 | 0,00501815 | MT-TH | 0,54396517 |
| ENSG00000 | gene | 1,20916633 | 0,00023332 | 0,00505157 | RANBP6 | 0,27401271 |
| ENSG00002 | gene | 1,41126847 | 0,00023412 | 0,00505774 | AC145207.4 | 0,49699247 |
| ENSG00002 | gene | 1,95177302 | 0,00023604 | 0,00508825 | AC004528.2 | 0,96478529 |
| ENSG00000 | gene | 0,84161893 | 0,0002373 | 0,0050949 | MEIS3 | -0,2487609 |
| ENSG00002 | gene | 1,60727825 | 0,00023738 | 0,0050949 | MT-TQ | 0,68461971 |
| ENSG00000 | gene | 1,14519694 | 0,00023913 | 0,00512144 | SENP6 | 0,19559572 |
| ENSG00000 | gene | 0,79847684 | 0,00024211 | 0,00517414 | USF2 | -0,3246775 |
| ENSG00000 | gene | 0,8101686 | 0,00024325 | 0,00517869 | B2M | -0,3037059 |
| ENSG00000 | gene | 0,87722386 | 0,00024362 | 0,00517869 | IFNGR2 | -0,188983 |
| ENSG00000 | gene | 0,86235763 | 0,00024395 | 0,00517869 | DDX41 | -0,2136418 |
| ENSG00002 | gene | 1,56957884 | 0,00024442 | 0,00517869 | AC068888.1 | 0,6503775 |
| ENSG00000( | gene | 0,69738214 | 0,00024548 | 0,00518173 | PDE4A | -0,5199787 |
| ENSG00002 | gene | 1,94931253 | 0,00024624 | 0,00518173 | AC008914.1 | 0,96296541 |
| ENSG00000 | gene | 0,81603421 | 0,00024679 | 0,00518173 | TMEM54 | -0,2932985 |
| ENSG00002 | gene | 0,84700415 | 0,00024687 | 0,00518173 | HLA-E | -0,2395591 |
| ENSG00000 | gene | 0,63086956 | 0,00024717 | 0,00518173 | ABTB1 | -0,6645864 |
| ENSG00000( | gene | 0,84838705 | 0,00024962 | 0,00520488 | SMARCD1 | -0,2372055 |
| ENSG00000 | gene | 0,85008988 | 0,00025023 | 0,00520488 | SAP30BP | -0,2343127 |
| ENSG00000( | gene | 0,83865323 | 0,00025029 | 0,00520488 | OTUD5 | -0,2538537 |
| ENSG00000 | gene | 1,10478031 | 0,00025047 | 0,00520488 | ENC1 | 0,14375951 |
| ENSG00000 | gene | 0,90095457 | 0,00025144 | 0,00520488 | CD2BP2 | -0,1504737 |
| ENSG00000 | gene | 0,76826489 | 0,00025151 | 0,00520488 | DIABLO | -0,3803243 |
| ENSG00000 | gene | 0,86958451 | 0,00025195 | 0,00520488 | NUBP1 | -0,2016019 |
| ENSG00000 | gene | 0,92733172 | 0,00025383 | 0,00523277 | HDGF | -0,1088426 |
| ENSG00000 | gene | 0,86827219 | 0,00025678 | 0,00528273 | MACROH2A | -0,2037807 |
| ENSG00000 | gene | 0,84433654 | 0,00025955 | 0,00532863 | TJAP1 | -0,2441099 |
| ENSG00000 | gene | 1,29944465 | 0,00026037 | 0,00533076 | MYSM1 | 0,37789519 |
| ENSG00000 | gene | 1,23182761 | 0,00026073 | 0,00533076 | NIPBL | 0,30080037 |
| ENSG00000 | gene | 0,85465894 | 0,00026159 | 0,00533722 | MAPKAPK2 | -0,2265793 |
| ENSG00000( | gene | 1,41601809 | 0,00026514 | 0,00539855 | ARVCF | 0,50183969 |
| ENSG00000 | gene | 0,86868794 | 0,00026629 | 0,00541091 | STX5 | -0,2030901 |
| ENSG00000( | gene | 0,83648264 | 0,00026953 | 0,00546563 | RFXANK | -0,2575925 |
| ENSG00000( | gene | 0,92450329 | 0,00027231 | 0,0055107 | RHOA | -0,1132496 |
| ENSG00002 | gene | 2,20685233 | 0,00027303 | 0,00551393 | AC093752.1 | 1,1419901 |
| ENSG00002 | gene | 0,86574085 | 0,00027413 | 0,00552487 | ABHD16A | -0,2079929 |
| ENSG00002 | gene | 0,78440032 | 0,00027518 | 0,00553491 | AC104563.1 | -0,350338 |
| ENSG00000 | gene | 0,89642111 | 0,00027739 | 0,005548 | BRD3 | -0,1577515 |
| ENSG00000 | gene | 0,83181809 | 0,0002774 | 0,005548 | BIN3 | -0,26566 |
| ENSG00000( | gene | 0,86725653 | 0,00027751 | 0,005548 | CHERP | -0,2054693 |
| ENSG00000( | gene | 1,24699415 | 0,00028031 | 0,00559272 | NEDD4 | 0,3184547 |
| ENSG00000( | gene | 0,85947657 | 0,00028135 | 0,0055985 | NCAPH2 | -0,2184698 |
| ENSG00000( | gene | 1,28672303 | 0,00028213 | 0,0055985 | IFT80 | 0,36370155 |
| ENSG00000 | gene | 0,89576467 | 0,0002823 | 0,0055985 | DUS1L | -0,1588083 |
| ENSG00000( | gene | 1,21861552 | 0,00028399 | 0,00561886 | RPAP3 | 0,28524302 |
| ENSG00000 | gene | 0,88506916 | 0,00028446 | 0,00561886 | PEX14 | -0,1761379 |
| ENSG00002 | gene | 0,87038573 | 0,00028908 | 0,00568965 | TRIM26 | -0,2002732 |
| ENSG00000 | gene | 0,8602805 | 0,00028919 | 0,00568965 | SLC4A2 | -0,217121 |
| ENSG00000 | gene | 0,83935502 | 0,00029173 | 0,0057208 | RILPL1 | -0,2526469 |
| ENSG00000 | gene | 0,8657096 | 0,00029192 | 0,0057208 | SINHCAF | -0,2080449 |
| ENSG00000( | gene | 0,89697291 | 0,00029314 | 0,00573329 | FAM50A | -0,1568637 |
| ENSG00000 | gene | 1,21258256 | 0,00029529 | 0,00576396 | CSE1L | 0,27808297 |
| ENSG00002 | gene | 0,90733387 | 0,00030218 | 0,00588685 | RNF5 | -0,1402946 |
| ENSG00000 | gene | 1,16870227 | 0,00030767 | 0,00598203 | GALNT1 | 0,22490744 |
| ENSG00000 | gene | 0,89399829 | 0,00030856 | 0,00598764 | POP7 | -0,161656 |

| | | | | | | |
|---|---|---|---|---|---|---|
| ENSG00000' | gene | 0,8221816 | 0,00030973 | 0,00599852 | BCAP31 | -0,282471 |
| ENSG00000' | gene | 1,22994848 | 0,00031064 | 0,00600458 | ATP11C | 0,29859789 |
| ENSG00000' | gene | 0,86367937 | 0,00031155 | 0,00601046 | DNPEP | -0,2114323 |
| ENSG00000' | gene | 0,86405351 | 0,00031314 | 0,0060294 | OS9 | -0,2108074 |
| ENSG00000' | gene | 0,84208733 | 0,00031571 | 0,006067 | CHCHD6 | -0,2479582 |
| ENSG00000' | gene | 1,80582943 | 0,00031756 | 0,00609076 | ACAD10 | 0,85266163 |
| ENSG00000: | gene | 1,36262503 | 0,00031924 | 0,00611111 | AC044849.1 | 0,44638862 |
| ENSG00000' | gene | 0,89874415 | 0,00032034 | 0,00612034 | DKC1 | -0,1540176 |
| ENSG00000' | gene | 1,14029044 | 0,00032273 | 0,00615416 | GTF2F2 | 0,18940134 |
| ENSG00000' | gene | 0,77104941 | 0,0003251 | 0,00618625 | SSNA1 | -0,3751048 |
| ENSG00000' | gene | 1,14442528 | 0,00032599 | 0,00618625 | NCBP1 | 0,19462327 |
| ENSG00000: | gene | 1,3911744 | 0,0003263 | 0,00618625 | AL354740.1 | 0,47630329 |
| ENSG00000' | gene | 0,85360205 | 0,00032691 | 0,00618625 | MRPL2 | -0,2283645 |
| ENSG00000( | gene | 0,90259645 | 0,00032761 | 0,0061877 | ENO1 | -0,147847 |
| ENSG00000' | gene | 1,88824793 | 0,00033068 | 0,00622545 | DMXL1 | 0,9170482 |
| ENSG00000' | gene | 0,82968711 | 0,00033086 | 0,00622545 | NUMA1 | -0,2693607 |
| ENSG00000( | gene | 1,13541937 | 0,00033936 | 0,00637336 | UBA5 | 0,18322525 |
| ENSG00000: | gene | 1,87786738 | 0,00034159 | 0,00640311 | AC145207.3 | 0,90909518 |
| ENSG00000' | gene | 0,86015522 | 0,00034381 | 0,00643245 | PIP5K1A | -0,2173311 |
| ENSG00000' | gene | 0,7899032 | 0,00034479 | 0,00643873 | PIH1D1 | -0,3402522 |
| ENSG00000' | gene | 1,13645121 | 0,00034733 | 0,00647399 | DDX46 | 0,18453575 |
| ENSG00000' | gene | 0,86529788 | 0,00034935 | 0,00649929 | FAM160B2 | -0,2087312 |
| ENSG00000: | gene | 0,78547171 | 0,00035315 | 0,00655777 | SMIM7 | -0,3483688 |
| ENSG00000' | gene | 0,84526844 | 0,00035454 | 0,00656658 | VPS28 | -0,2425185 |
| ENSG00000' | gene | 0,79288796 | 0,00035495 | 0,00656658 | DHRS4 | -0,3348111 |
| ENSG00000' | gene | 1,14020906 | 0,00035659 | 0,00658466 | CCDC50 | 0,18929837 |
| ENSG00000: | gene | 1,34193457 | 0,00035828 | 0,00660287 | GAS5-AS1 | 0,42431433 |
| ENSG00000' | gene | 0,82570351 | 0,00035903 | 0,00660287 | PHF11 | -0,2763043 |
| ENSG00000' | gene | 0,85753212 | 0,00035958 | 0,00660287 | TBC1D9B | -0,2217374 |
| ENSG00000( | gene | 0,84844519 | 0,00036162 | 0,00662074 | RIPOR1 | -0,2371066 |
| ENSG00000' | gene | 0,85524856 | 0,00036288 | 0,00662074 | KIAA0930 | -0,2255843 |
| ENSG00000' | gene | 0,8347393 | 0,00036313 | 0,00662074 | ARL8A | -0,2606024 |
| ENSG00000' | gene | 0,84158741 | 0,00036367 | 0,00662074 | YIPF3 | -0,248815 |
| ENSG00000' | gene | 0,83593315 | 0,00036389 | 0,00662074 | NAPA | -0,2585405 |
| ENSG00000' | gene | 0,66309352 | 0,00036995 | 0,00671867 | UPP1 | -0,5927157 |
| ENSG00000' | gene | 0,90324913 | 0,000372 | 0,00674359 | PTDSS1 | -0,1468041 |
| ENSG00000( | gene | 0,85428694 | 0,0003753 | 0,00678698 | RNH1 | -0,2272074 |
| ENSG00000( | gene | 1,20519186 | 0,00037576 | 0,00678698 | BCLAF1 | 0,26926284 |
| ENSG00000( | gene | 0,90142772 | 0,00037682 | 0,00679366 | MAST2 | -0,1497163 |
| ENSG00000' | gene | 0,90658227 | 0,00037935 | 0,00682689 | KLHDC3 | -0,1414901 |
| ENSG00000: | gene | 2,09339132 | 0,00038103 | 0,0068447 | AC087521.4 | 1,06584202 |
| ENSG00000' | gene | 1,12187157 | 0,00038186 | 0,00684724 | MFN1 | 0,16590752 |
| ENSG00000' | gene | 0,74666485 | 0,00038697 | 0,00692643 | SLC3A2 | -0,4214673 |
| ENSG00000' | gene | 0,68688012 | 0,00038843 | 0,00693986 | FKBP1B | -0,5418698 |
| ENSG00000: | gene | 1,57896073 | 0,00038953 | 0,00694717 | AC008878.4 | 0,65897529 |
| ENSG00000' | gene | 1,2589958 | 0,00039271 | 0,00699121 | MBNL1 | 0,33227347 |
| ENSG00000: | gene | 1,32614428 | 0,00039662 | 0,00704811 | MED4-AS1 | 0,40723774 |
| ENSG00000' | gene | 0,8842506 | 0,00039896 | 0,00706353 | MRGBP | -0,1774728 |
| ENSG00000( | gene | 0,80421345 | 0,00039952 | 0,00706353 | BLVRB | -0,3143496 |
| ENSG00000' | gene | 0,85554378 | 0,00039962 | 0,00706353 | FADS1 | -0,2250864 |
| ENSG00000' | gene | 1,20337104 | 0,0004028 | 0,00710282 | CUL5 | 0,26708154 |
| ENSG00000' | gene | 1,18153474 | 0,00040328 | 0,00710282 | SUCLA2 | 0,24066205 |
| ENSG00000: | gene | 0,88038658 | 0,00040725 | 0,00714252 | ABCF1 | -0,1837909 |
| ENSG00000' | gene | 0,65115736 | 0,00040767 | 0,00714252 | FMR1 | -0,6189219 |
| ENSG00000: | gene | 0,81935117 | 0,00040813 | 0,00714252 | AC017083.3 | -0,2874462 |
| ENSG00000' | gene | 0,87874164 | 0,00040841 | 0,00714252 | IMP4 | -0,186489 |
| ENSG00000( | gene | 0,79365387 | 0,00041209 | 0,00719423 | PSME1 | -0,3334181 |

| | | | | | | |
|---|---|---|---|---|---|---|
| ENSG00000( | gene | 0,77909069 | 0,00041362 | 0,00720122 | NDUFB4 | -0,3601368 |
| ENSG00000' | gene | 1,16072793 | 0,00041426 | 0,00720122 | ARPP19 | 0,21502985 |
| ENSG00000: | gene | 1,48686689 | 0,00041467 | 0,00720122 | MT-TT | 0,5722755 |
| ENSG00000( | gene | 0,90860688 | 0,00041557 | 0,00720414 | SNX11 | -0,1382719 |
| ENSG00000: | gene | 1,28672588 | 0,00041724 | 0,00722047 | AC012363.2 | 0,36370473 |
| ENSG00000: | gene | 1,15215163 | 0,00041936 | 0,00724447 | AL031587.5 | 0,20433059 |
| ENSG00000: | gene | 0,82734372 | 0,00042145 | 0,00726803 | RPL4P4 | -0,2734413 |
| ENSG00000' | gene | 1,33961573 | 0,00042292 | 0,00726997 | SMC4 | 0,42181923 |
| ENSG00000( | gene | 0,85800169 | 0,00042303 | 0,00726997 | BUD23 | -0,2209476 |
| ENSG00000: | gene | 2,20548586 | 0,00042439 | 0,00728064 | TMSB4XP6 | 1,14109651 |
| ENSG00000: | gene | 2,371688 | 0,0004268 | 0,00730947 | SNORA18 | 1,24591423 |
| ENSG00000' | gene | 1,09360958 | 0,00042814 | 0,00731973 | DCAF13 | 0,12909778 |
| ENSG00000' | gene | 1,12213865 | 0,00042898 | 0,00732153 | CPD | 0,16625095 |
| ENSG00000' | gene | 0,82571836 | 0,00043135 | 0,00734924 | ZFPL1 | -0,2762783 |
| ENSG00000' | gene | 0,86675569 | 0,00043438 | 0,00738351 | AUP1 | -0,2063027 |
| ENSG00000( | gene | 1,3523842 | 0,00043527 | 0,00738351 | EVI5 | 0,43550506 |
| ENSG00000' | gene | 0,80455159 | 0,000436 | 0,00738351 | ATP5MC1 | -0,3137432 |
| ENSG00000: | gene | 0,91562057 | 0,00043634 | 0,00738351 | GNL1 | -0,1271782 |
| ENSG00000' | gene | 1,17597277 | 0,00043952 | 0,00741357 | AHNAK | 0,23385466 |
| ENSG00000' | gene | 1,20012675 | 0,00043961 | 0,00741357 | PIGK | 0,26318679 |
| ENSG00000( | gene | 0,7182271 | 0,00044211 | 0,00743738 | IL4R | -0,477488 |
| ENSG00000' | gene | 0,84107853 | 0,00044252 | 0,00743738 | TARBP2 | -0,2496876 |
| ENSG00000' | gene | 1,08947634 | 0,00044668 | 0,00749459 | UGP2 | 0,12363487 |
| ENSG00000' | gene | 0,85358284 | 0,00044845 | 0,00751155 | GLOD4 | -0,2283969 |
| ENSG00000' | gene | 0,85960103 | 0,0004499 | 0,0075232 | MPDU1 | -0,2182609 |
| ENSG00000' | gene | 1,10263592 | 0,00045233 | 0,0075511 | CNOT11 | 0,14095651 |
| ENSG00000' | gene | 0,86716946 | 0,00045326 | 0,00755376 | ARFIP2 | -0,2056141 |
| ENSG00000' | gene | 1,30837682 | 0,00045518 | 0,00757303 | MANEA | 0,38777811 |
| ENSG00000: | gene | 1,29524574 | 0,0004566 | 0,00758333 | AC130304.1 | 0,37322584 |
| ENSG00000' | gene | 0,78700385 | 0,00045732 | 0,00758333 | LGALS1 | -0,3455574 |
| ENSG00000' | gene | 1,26265156 | 0,00045954 | 0,00759425 | CD2AP | 0,33645657 |
| ENSG00000' | gene | 0,77164047 | 0,00046022 | 0,00759425 | CUTA | -0,3739993 |
| ENSG00000' | gene | 1,20117357 | 0,00046028 | 0,00759425 | TMX3 | 0,26444463 |
| ENSG00000: | gene | 0,89152782 | 0,00046787 | 0,00769931 | CLIC1 | -0,1656483 |
| ENSG00000' | gene | 0,89895712 | 0,0004682 | 0,00769931 | HARS1 | -0,1536758 |
| ENSG00000: | gene | 0,77384984 | 0,000469 | 0,00769976 | RPL10P9 | -0,3698744 |
| ENSG00000' | gene | 0,66686496 | 0,00047042 | 0,00771028 | TMEM51 | -0,5845334 |
| ENSG00000: | gene | 1,46748867 | 0,00047796 | 0,00782094 | SMARCA5-A | 0,55334936 |
| ENSG00000: | gene | 0,89269084 | 0,00048208 | 0,00787532 | PISD | -0,1637675 |
| ENSG00000( | gene | 1,19228099 | 0,00048399 | 0,00789356 | CBX5 | 0,25372428 |
| ENSG00000' | gene | 1,17993076 | 0,00048501 | 0,00789708 | TCAIM | 0,2387022 |
| ENSG00000' | gene | 0,80819557 | 0,00048638 | 0,00790639 | SRSF5 | -0,3072237 |
| ENSG00000' | gene | 0,80219864 | 0,00048967 | 0,00794685 | STARD3 | -0,3179686 |
| ENSG00000' | gene | 1,25346829 | 0,00049071 | 0,00795082 | KTN1 | 0,3259255 |
| ENSG00000: | gene | 1,40141893 | 0,00049213 | 0,00795403 | AC135507.1 | 0,48688829 |
| ENSG00000: | gene | 0,91210396 | 0,00049251 | 0,00795403 | AGAP2-AS1 | -0,1327298 |
| ENSG00000' | gene | 1,68316041 | 0,000495 | 0,00798122 | AHI1 | 0,75117268 |
| ENSG00000' | gene | 0,80174998 | 0,00049604 | 0,00798489 | TRIM25 | -0,3187757 |
| ENSG00000' | gene | 0,82695201 | 0,000497 | 0,00798738 | ALDOA | -0,2741245 |
| ENSG00000' | gene | 0,75356097 | 0,00050068 | 0,00803358 | C8orf76 | -0,4082038 |
| ENSG00000( | gene | 0,79225446 | 0,00050186 | 0,00803946 | MRPS24 | -0,3359642 |
| ENSG00000' | gene | 0,86301778 | 0,00050349 | 0,00804529 | CHTOP | -0,2125378 |
| ENSG00000' | gene | 0,89340457 | 0,00050384 | 0,00804529 | CYC1 | -0,1626145 |
| ENSG00000' | gene | 0,85986802 | 0,00050559 | 0,00806011 | NUCB1 | -0,2178129 |
| ENSG00000' | gene | 1,11486002 | 0,00051344 | 0,00817213 | WWP1 | 0,15686257 |
| ENSG00000' | gene | 0,86651111 | 0,00052235 | 0,00830074 | TFPT | -0,2067098 |
| ENSG00000' | gene | 1,21762802 | 0,00052546 | 0,00833674 | GOLGB1 | 0,28407346 |

| | | | | | | |
|---|---|---|---|---|---|---|
| ENSG000002 | gene | 2,42327817 | 0,00052974 | 0,00839114 | PDIA3P2 | 1,27696002 |
| ENSG000001 | gene | 0,72597935 | 0,00053529 | 0,00846022 | JAM3 | -0,4619996 |
| ENSG000002 | gene | 1,83186117 | 0,0005358 | 0,00846022 | RNF185-AS1 | 0,87331017 |
| ENSG000000 | gene | 1,43036627 | 0,00054021 | 0,00850973 | ASPM | 0,51638463 |
| ENSG000001 | gene | 1,21086906 | 0,00054125 | 0,00850973 | UHRF1BP1L | 0,27604286 |
| ENSG000001 | gene | 0,88122051 | 0,00054195 | 0,00850973 | ARAP1 | -0,182425 |
| ENSG000001 | gene | 0,76792091 | 0,00054237 | 0,00850973 | EEF1D | -0,3809704 |
| ENSG000002 | gene | 1,38308278 | 0,00054785 | 0,00856975 | AP001107.6 | 0,46788751 |
| ENSG000001 | gene | 0,85633826 | 0,00054876 | 0,00856975 | LMAN2 | -0,2237473 |
| ENSG000001 | gene | 1,09475489 | 0,00054879 | 0,00856975 | AP3M1 | 0,13060789 |
| ENSG000001 | gene | 0,88180788 | 0,0005544 | 0,00864375 | TMEM87A | -0,1814637 |
| ENSG000001 | gene | 1,13698005 | 0,00055669 | 0,00866582 | GLCE | 0,18520694 |
| ENSG000001 | gene | 1,19335028 | 0,00055804 | 0,00867324 | HAT1 | 0,25501757 |
| ENSG000000 | gene | 1,13454953 | 0,00056102 | 0,00870593 | WDFY1 | 0,1821196 |
| ENSG000001 | gene | 0,82444338 | 0,00056341 | 0,0087233 | GSE1 | -0,2785077 |
| ENSG000001 | gene | 0,84581413 | 0,00056402 | 0,0087233 | HDAC3 | -0,2415874 |
| ENSG000001 | gene | 0,88680827 | 0,00056478 | 0,0087233 | AIP | -0,1733059 |
| ENSG000001 | gene | 0,86788983 | 0,0005682 | 0,00876254 | MRPS7 | -0,2044162 |
| ENSG000001 | gene | 1,14737438 | 0,00057387 | 0,0088361 | VBP1 | 0,19833621 |
| ENSG000001 | gene | 0,88874088 | 0,00057669 | 0,00885447 | SEC11A | -0,1701652 |
| ENSG000001 | gene | 0,85098045 | 0,00057684 | 0,00885447 | PRMT2 | -0,2328021 |
| ENSG000002 | gene | 0,8235933 | 0,00058319 | 0,00891515 | AC074212.1 | -0,279996 |
| ENSG000001 | gene | 0,84103197 | 0,00058496 | 0,00891515 | ZFAND2B | -0,2497674 |
| ENSG000001 | gene | 0,65295056 | 0,00058533 | 0,00891515 | KHDC4 | -0,6149543 |
| ENSG000002 | gene | 0,8319878 | 0,00058627 | 0,00891515 | AC092115.4 | -0,2653657 |
| ENSG000000 | gene | 1,08678966 | 0,00058653 | 0,00891515 | GOLGA5 | 0,12007275 |
| ENSG000002 | gene | 0,88803617 | 0,00058674 | 0,00891515 | CCDC85C | -0,1713097 |
| ENSG000000 | gene | 0,88202731 | 0,00058709 | 0,00891515 | SPAG7 | -0,1811048 |
| ENSG000001 | gene | 0,8962778 | 0,00059024 | 0,00894687 | PTPA | -0,1579821 |
| ENSG000001 | gene | 0,75360748 | 0,00059098 | 0,00894687 | ATAD3B | -0,4081148 |
| ENSG000002 | gene | 0,32588851 | 0,00059307 | 0,00896474 | NDUFC2-KC | -1,6175496 |
| ENSG000001 | gene | 0,78719451 | 0,00060091 | 0,00906939 | RNF114 | -0,3452079 |
| ENSG000002 | gene | 1,4838853 | 0,00060263 | 0,00908164 | AL157871.1 | 0,56937958 |
| ENSG000001 | gene | 0,84450972 | 0,00060574 | 0,00911454 | CIZ1 | -0,2438141 |
| ENSG000001 | gene | 0,85849442 | 0,00061158 | 0,0091885 | NHP2 | -0,2201193 |
| ENSG000000 | gene | 0,76931646 | 0,0006146 | 0,00921696 | GADD45B | -0,3783509 |
| ENSG000000 | gene | 0,84097467 | 0,00061533 | 0,00921696 | P4HA2 | -0,2498658 |
| ENSG000001 | gene | 0,80275547 | 0,00061701 | 0,00921741 | CIAO3 | -0,3169675 |
| ENSG000001 | gene | 0,66857291 | 0,00061722 | 0,00921741 | OAS3 | -0,5808432 |
| ENSG000001 | gene | 1,091009 | 0,00061934 | 0,00923513 | CAPRIN1 | 0,125663 |
| ENSG000002 | gene | 1,27591469 | 0,00062189 | 0,00925928 | AC007383.3 | 0,35153188 |
| ENSG000001 | gene | 0,86008125 | 0,00062402 | 0,0092744 | MVD | -0,2174551 |
| ENSG000001 | gene | 1,12483866 | 0,00062478 | 0,0092744 | GFM2 | 0,16971809 |
| ENSG000001 | gene | 1,14309716 | 0,00062784 | 0,00930589 | STAG1 | 0,19294803 |
| ENSG000001 | gene | 0,8205029 | 0,00062968 | 0,00930661 | FOSL1 | -0,2854197 |
| ENSG000001 | gene | 1,40884118 | 0,0006303 | 0,00930661 | AFF3 | 0,49450898 |
| ENSG000001 | gene | 0,89563067 | 0,00063103 | 0,00930661 | GNA12 | -0,1590242 |
| ENSG000000 | gene | 0,91213935 | 0,00063184 | 0,00930661 | GNPTG | -0,1326738 |
| ENSG000001 | gene | 0,88222266 | 0,00063258 | 0,00930661 | GUSB | -0,1807853 |
| ENSG000001 | gene | 0,89950703 | 0,00063509 | 0,00932338 | USF1 | -0,1527935 |
| ENSG000000 | gene | 0,92267309 | 0,0006356 | 0,00932338 | CRAT | -0,1161085 |
| ENSG000001 | gene | 0,81295646 | 0,00064082 | 0,00936599 | PTOV1 | -0,29875 |
| ENSG000001 | gene | 0,9235947 | 0,00064099 | 0,00936599 | IK | -0,1146682 |
| ENSG000002 | gene | 1,53959843 | 0,00064134 | 0,00936599 | AC137630.3 | 0,62255411 |
| ENSG000001 | gene | 1,19489064 | 0,00064361 | 0,00937374 | NMD3 | 0,25687859 |
| ENSG000001 | gene | 1,1516007 | 0,00064376 | 0,00937374 | CD47 | 0,20364057 |
| ENSG000001 | gene | 0,86460333 | 0,00064663 | 0,00940172 | SRSF4 | -0,2098897 |

| | | | | | | |
|---|---|---|---|---|---|---|
| ENSG00002 | gene | 1,38262498 | 0,00064899 | 0,00941752 | AL109628.2 | 0,4674099 |
| ENSG00000 | gene | 0,90090281 | 0,00064996 | 0,00941752 | SCAMP3 | -0,1505566 |
| ENSG00002 | gene | 0,81118189 | 0,00065056 | 0,00941752 | TMEM199 | -0,3019027 |
| ENSG00000 | gene | 0,76573708 | 0,00065793 | 0,00951023 | RPS16 | -0,385079 |
| ENSG00000 | gene | 0,78035438 | 0,00065918 | 0,00951447 | ARL6IP4 | -0,3577987 |
| ENSG00000 | gene | 0,82608954 | 0,00066195 | 0,00954049 | MAF1 | -0,2756299 |
| ENSG00000 | gene | 0,84970151 | 0,00066713 | 0,00958884 | SF3A2 | -0,234972 |
| ENSG00002 | gene | 1,72424226 | 0,00066724 | 0,00958884 | AC009630.2 | 0,78596249 |
| ENSG00000 | gene | 0,85411904 | 0,0006688 | 0,00959001 | ERCC3 | -0,2274909 |
| ENSG00000 | gene | 0,77550605 | 0,00066925 | 0,00959001 | TMEM159 | -0,3667901 |
| ENSG00002 | gene | 1,48562616 | 0,00067124 | 0,00960471 | AFAP1-AS1 | 0,57107112 |
| ENSG00000 | gene | 1,20592954 | 0,00067249 | 0,00960869 | PPP2R3A | 0,27014562 |
| ENSG00000 | gene | 0,88726557 | 0,00067519 | 0,00962246 | ADPGK | -0,1725621 |
| ENSG00002 | gene | 0,81236155 | 0,0006754 | 0,00962246 | AC034102.1 | -0,2998061 |
| ENSG00002 | gene | 0,90289298 | 0,0006775 | 0,00963219 | RNPS1 | -0,1473731 |
| ENSG00002 | gene | 1,33792987 | 0,00067901 | 0,00963219 | BX322234.1 | 0,4200025 |
| ENSG00000 | gene | 1,18399214 | 0,00067925 | 0,00963219 | ATF1 | 0,2436595 |
| ENSG00000 | gene | 1,16124689 | 0,00067997 | 0,00963219 | GRIK2 | 0,21567473 |
| ENSG00000 | gene | 0,90841133 | 0,00068527 | 0,00968941 | RAB11B | -0,1385824 |
| ENSG00000 | gene | 0,78870016 | 0,00068596 | 0,00968941 | TXNRD1 | -0,3424512 |
| ENSG00000 | gene | 0,81039995 | 0,00068986 | 0,00972416 | ATP6V0B | -0,303294 |
| ENSG00000 | gene | 1,23789873 | 0,00069038 | 0,00972416 | MIS18BP1 | 0,3078933 |
| ENSG00000 | gene | 0,86678913 | 0,00069321 | 0,00973866 | GRN | -0,206247 |
| ENSG00000 | gene | 1,26767683 | 0,00069407 | 0,00973866 | IL6ST | 0,34218701 |
| ENSG00000 | gene | 0,83977049 | 0,00069519 | 0,00973866 | SFT2D1 | -0,251933 |
| ENSG00000 | gene | 0,93482855 | 0,00069639 | 0,00973866 | EIF4H | -0,0972263 |
| ENSG00000 | gene | 0,91345859 | 0,0006966 | 0,00973866 | TRAF7 | -0,1305888 |
| ENSG00000 | gene | 0,91092078 | 0,0006973 | 0,00973866 | FTSJ3 | -0,1346025 |
| ENSG00000 | gene | 0,79907942 | 0,0006986 | 0,00974307 | EIF4A1 | -0,3235892 |
| ENSG00000 | gene | 0,83506117 | 0,00070076 | 0,00975694 | NQO1 | -0,2600462 |
| ENSG00000 | gene | 0,84153922 | 0,00070156 | 0,00975694 | FBXO44 | -0,2488976 |
| ENSG00000 | gene | 0,76670614 | 0,00070844 | 0,00983878 | SERF2 | -0,3832544 |
| ENSG00000 | gene | 1,26865367 | 0,00070969 | 0,00984241 | TBC1D23 | 0,34329828 |
| ENSG00000 | gene | 0,83553634 | 0,00071381 | 0,00988565 | CYB5R1 | -0,2592255 |
| ENSG00002 | gene | 0,78606326 | 0,00072454 | 0,01000029 | ENSAP2 | -0,3472827 |
| ENSG00002 | gene | 0,73142332 | 0,0007248 | 0,01000029 | RPS7P10 | -0,4512215 |
| ENSG00000 | gene | 1,1202873 | 0,00072511 | 0,01000029 | ARHGEF12 | 0,16386877 |
| ENSG00000 | gene | 0,92998472 | 0,00072872 | 0,01003131 | OGFOD1 | -0,1047211 |
| ENSG00000 | gene | 0,83547891 | 0,00072938 | 0,01003131 | GGA1 | -0,2593247 |
| ENSG00002 | gene | 2,35190185 | 0,00073097 | 0,01003922 | MATR3 | 1,23382785 |
| ENSG00000 | gene | 0,84001751 | 0,00073244 | 0,01004547 | CD99 | -0,2515087 |
| ENSG00002 | gene | 0,82924664 | 0,00073386 | 0,01005105 | BRD2 | -0,2701268 |
| ENSG00000 | gene | 0,7517128 | 0,00073692 | 0,01007875 | INO80B | -0,4117465 |
| ENSG00000 | gene | 0,85461017 | 0,0007391 | 0,01007875 | C16orf58 | -0,2266616 |
| ENSG00000 | gene | 0,78523442 | 0,0007393 | 0,01007875 | PPP2R3B | -0,3488047 |
| ENSG00002 | gene | 0,42508825 | 0,0007403 | 0,01007875 | RNA5-8SN2 | -1,2341657 |
| ENSG00000 | gene | 0,82566821 | 0,00074132 | 0,01007875 | C7orf26 | -0,2763659 |
| ENSG00000 | gene | 1,45624371 | 0,00074198 | 0,01007875 | ATAD2 | 0,54225182 |
| ENSG00000 | gene | 1,29702138 | 0,00074464 | 0,01009057 | KNTC1 | 0,37520226 |
| ENSG00000 | gene | 1,17611171 | 0,00074489 | 0,01009057 | TOPBP1 | 0,2340251 |
| ENSG00000 | gene | 0,87507629 | 0,00074844 | 0,0101132 | EMC4 | -0,1925193 |
| ENSG00002 | gene | 0,62254436 | 0,0007486 | 0,0101132 | AC005041.5 | -0,6837514 |
| ENSG00000 | gene | 0,89234919 | 0,00075215 | 0,01014735 | YJU2 | -0,1643197 |
| ENSG00000 | gene | 0,87604094 | 0,0007582 | 0,01021514 | TMEM11 | -0,1909298 |
| ENSG00002 | gene | 0,88232425 | 0,00076288 | 0,01026414 | GOLGA2P1C | -0,1806192 |
| ENSG00000 | gene | 0,66432339 | 0,00076455 | 0,0102727 | EPSTI1 | -0,5900424 |
| ENSG00000 | gene | 1,28182203 | 0,00077044 | 0,01032894 | CFAP73 | 0,35819597 |

| | | | | | | |
|---|---|---|---|---|---|---|
| ENSG00000' | gene | 1,26155697 | 0,00077082 | 0,01032894 | NEDD1 | 0,33520535 |
| ENSG00000' | gene | 0,87500482 | 0,00077395 | 0,01034784 | SEC13 | -0,1926371 |
| ENSG00000( | gene | 0,80692792 | 0,00077431 | 0,01034784 | PTBP1 | -0,3094883 |
| ENSG00000' | gene | 0,82512289 | 0,00077732 | 0,01037205 | TRIM41 | -0,2773191 |
| ENSG00000' | gene | 0,86259835 | 0,00077822 | 0,01037205 | PRDX2 | -0,2132391 |
| ENSG00000' | gene | 1,20265877 | 0,00078549 | 0,01044643 | TWISTNB | 0,26622736 |
| ENSG00000; | gene | 0,41380242 | 0,00078591 | 0,01044643 | RNA5-8SN4 | -1,272986 |
| ENSG00000; | gene | 1,49508579 | 0,00079455 | 0,0105324 | AC008264.2 | 0,58022827 |
| ENSG00000' | gene | 1,2408793 | 0,0007956 | 0,0105324 | CTNNAL1 | 0,31136279 |
| ENSG00000' | gene | 0,89377762 | 0,00079586 | 0,0105324 | H1-0 | -0,1620122 |
| ENSG00000( | gene | 0,84817382 | 0,00079662 | 0,0105324 | LYPLA2 | -0,2375681 |
| ENSG00000' | gene | 0,78978642 | 0,00080172 | 0,01058574 | RPL13A | -0,3404655 |
| ENSG00000' | gene | 0,76236491 | 0,00081091 | 0,01069078 | ADCY7 | -0,3914464 |
| ENSG00000; | gene | 0,85407794 | 0,00081184 | 0,01069078 | NUP62 | -0,2275604 |
| ENSG00000' | gene | 1,52293435 | 0,00081516 | 0,0107203 | TCHP | 0,60685375 |
| ENSG00000( | gene | 1,1322556 | 0,00082039 | 0,01077478 | CDC27 | 0,17919967 |
| ENSG00000; | gene | 0,51133565 | 0,00082197 | 0,01078081 | SNORD3A | -0,9676575 |
| ENSG00000' | gene | 0,82159386 | 0,00082382 | 0,01078081 | INO80E | -0,2835027 |
| ENSG00000( | gene | 1,11490437 | 0,00082411 | 0,01078081 | OSBPL3 | 0,15691997 |
| ENSG00000' | gene | 0,80433325 | 0,00083026 | 0,01084697 | EIF1B | -0,3141347 |
| ENSG00000' | gene | 0,8047288 | 0,00083642 | 0,01091302 | TMBIM6 | -0,3134254 |
| ENSG00000' | gene | 0,79299835 | 0,00084701 | 0,01103243 | ITM2C | -0,3346102 |
| ENSG00000' | gene | 0,85843364 | 0,00084779 | 0,01103243 | TBL2 | -0,2202215 |
| ENSG00000' | gene | 0,86548881 | 0,00085231 | 0,01107429 | SHMT2 | -0,2084129 |
| ENSG00000' | gene | 0,69000412 | 0,00085324 | 0,01107429 | NPIPB3 | -0,5353231 |
| ENSG00000( | gene | 1,19065089 | 0,00085464 | 0,01107791 | RECQL | 0,25175047 |
| ENSG00000( | gene | 1,15644024 | 0,00085881 | 0,01109404 | HLTF | 0,20969071 |
| ENSG00000' | gene | 0,80013894 | 0,0008594 | 0,01109404 | RPL28 | -0,3216776 |
| ENSG00000' | gene | 1,18218782 | 0,00085954 | 0,01109404 | PRRX1 | 0,24145926 |
| ENSG00000; | gene | 1,38276091 | 0,00086235 | 0,01109404 | AC016588.1 | 0,46755172 |
| ENSG00000' | gene | 1,17063787 | 0,00086268 | 0,01109404 | PRPF40A | 0,22729485 |
| ENSG00000' | gene | 1,11394478 | 0,00086329 | 0,01109404 | MAP1A | 0,15567771 |
| ENSG00000( | gene | 0,87454779 | 0,00086372 | 0,01109404 | GSTP1 | -0,1933909 |
| ENSG00000' | gene | 1,11698213 | 0,00086797 | 0,01113427 | FKBP14 | 0,15960611 |
| ENSG00000' | gene | 0,82467485 | 0,00086951 | 0,01113699 | LSM6 | -0,2781027 |
| ENSG00000; | gene | 1,63516565 | 0,00087144 | 0,01113699 | AC130456.5 | 0,70943679 |
| ENSG00000; | gene | 0,90150442 | 0,00087155 | 0,01113699 | AATF | -0,1495935 |
| ENSG00000' | gene | 1,51589995 | 0,00087894 | 0,0112168 | ATR | 0,60017454 |
| ENSG00000' | gene | 0,8500071 | 0,00088006 | 0,0112168 | IDH3B | -0,2344532 |
| ENSG00000' | gene | 0,66389312 | 0,00088619 | 0,01128044 | MX2 | -0,5909771 |
| ENSG00000' | gene | 0,8881446 | 0,00088973 | 0,01131103 | FADD | -0,1711335 |
| ENSG00000' | gene | 0,77982282 | 0,00089169 | 0,01131382 | TMEM205 | -0,3587817 |
| ENSG00000' | gene | 0,87128263 | 0,00089224 | 0,01131382 | NAT9 | -0,1987873 |
| ENSG00000; | gene | 1,54742223 | 0,00089786 | 0,0113699 | AC005726.2 | 0,62986691 |
| ENSG00000' | gene | 1,23224792 | 0,00089895 | 0,0113699 | GABPA | 0,30129254 |
| ENSG00000' | gene | 0,82635825 | 0,00090164 | 0,01138938 | ADRM1 | -0,2751607 |
| ENSG00000' | gene | 0,87371833 | 0,0009098 | 0,01147781 | NPM3 | -0,1947598 |
| ENSG00000; | gene | 1,30643225 | 0,00091712 | 0,01154444 | AL050331.2 | 0,38563231 |
| ENSG00000' | gene | 0,87589614 | 0,00091793 | 0,01154444 | ZC3H18 | -0,1911683 |
| ENSG00000' | gene | 0,87720275 | 0,00091961 | 0,01154444 | MSRB1 | -0,1890178 |
| ENSG00000' | gene | 0,80120424 | 0,00091974 | 0,01154444 | ATG101 | -0,319758 |
| ENSG00000; | gene | 1,80940301 | 0,00092155 | 0,01155254 | AC025262.3 | 0,85551378 |
| ENSG00000; | gene | 1,32743926 | 0,00092862 | 0,01162094 | SPTLC1P1 | 0,40864585 |
| ENSG00000; | gene | 0,6164614 | 0,00092935 | 0,01162094 | RPL39P3 | -0,6979175 |
| ENSG00000' | gene | 1,15866266 | 0,00093548 | 0,01168293 | ATF7IP | 0,21246059 |
| ENSG00000; | gene | 0,65558341 | 0,000938 | 0,01169098 | RPL24P2 | -0,6091487 |
| ENSG00000' | gene | 0,88660937 | 0,00093849 | 0,01169098 | CMIP | -0,1736295 |

| | | | | | | |
|---|---|---|---|---|---|---|
| ENSG000002 | gene | 1,78009251 | 0,00094412 | 0,01174647 | SNORA8 | 0,83195222 |
| ENSG000001 | gene | 0,86195735 | 0,00094866 | 0,01178812 | MKRN1 | -0,2143116 |
| ENSG000002 | gene | 0,63472785 | 0,00095295 | 0,01182659 | TEN1 | -0,6557899 |
| ENSG000002 | gene | 1,47265054 | 0,00095596 | 0,01183476 | AL450306.1 | 0,55841512 |
| ENSG000001 | gene | 1,16959685 | 0,00095618 | 0,01183476 | UAP1 | 0,22601133 |
| ENSG000001 | gene | 0,8215313 | 0,00095719 | 0,01183476 | GMPR | -0,2836126 |
| ENSG000001 | gene | 0,79702667 | 0,00096447 | 0,01189692 | IST1 | -0,3273001 |
| ENSG000000 | gene | 0,87571785 | 0,00096462 | 0,01189692 | POLR2J | -0,191462 |
| ENSG000002 | gene | 1,67518924 | 0,00097252 | 0,01197949 | AC005045.1 | 0,74432408 |
| ENSG000001 | gene | 0,86182124 | 0,00097648 | 0,01199715 | SELENOK | -0,2145394 |
| ENSG000000 | gene | 0,85661262 | 0,0009774 | 0,01199715 | COASY | -0,2232852 |
| ENSG000001 | gene | 0,83954168 | 0,00097758 | 0,01199715 | LMBR1L | -0,2523261 |
| ENSG000000 | gene | 1,16886333 | 0,00097894 | 0,01199896 | TRPM7 | 0,22510626 |
| ENSG000002 | gene | 0,74239475 | 0,00098167 | 0,01201762 | PPME1 | -0,4297416 |
| ENSG000002 | gene | 1,67240494 | 0,00098364 | 0,01202689 | AC243964.4 | 0,74192421 |
| ENSG000001 | gene | 0,69143968 | 0,00098633 | 0,01204485 | SPSB3 | -0,5323247 |
| ENSG000001 | gene | 0,84141248 | 0,00098946 | 0,01206822 | ST6GALNAC | -0,2491149 |
| ENSG000001 | gene | 0,73939931 | 0,00099237 | 0,01208885 | NPIPB4 | -0,4355744 |
| ENSG000001 | gene | 0,81324586 | 0,0009952 | 0,01210849 | CYB5R3 | -0,2982365 |
| ENSG000002 | gene | 0,88704663 | 0,00100043 | 0,01215721 | PHB2 | -0,1729182 |
| ENSG000001 | gene | 1,25691727 | 0,00101436 | 0,01231139 | IMPACT | 0,32988969 |
| ENSG000001 | gene | 0,78292588 | 0,00101803 | 0,01233704 | SLC41A3 | -0,3530524 |
| ENSG000000 | gene | 0,82377999 | 0,00101896 | 0,01233704 | BCL3 | -0,279669 |
| ENSG000001 | gene | 0,86736438 | 0,00102827 | 0,01243457 | CTDNEP1 | -0,2052899 |
| ENSG000000 | gene | 0,86212217 | 0,00103546 | 0,01249361 | CHMP5 | -0,2140358 |
| ENSG000002 | gene | 1,35564941 | 0,00103567 | 0,01249361 | AC020651.1 | 0,43898413 |
| ENSG000001 | gene | 0,82854734 | 0,00103866 | 0,01249991 | EMC9 | -0,271344 |
| ENSG000001 | gene | 1,10545322 | 0,00103872 | 0,01249991 | SGCB | 0,14463798 |
| ENSG000001 | gene | 0,84259704 | 0,00104088 | 0,01250402 | SERPINB8 | -0,2470853 |
| ENSG000000 | gene | 0,80923479 | 0,00104158 | 0,01250402 | PHPT1 | -0,3053697 |
| ENSG000000 | gene | 1,18889765 | 0,00104397 | 0,01251575 | MAP4K5 | 0,24962453 |
| ENSG000000 | gene | 1,35781719 | 0,00104508 | 0,01251575 | FAR2 | 0,44128926 |
| ENSG000001 | gene | 0,73360342 | 0,00105929 | 0,01267056 | STAG3L3 | -0,4469277 |
| ENSG000002 | gene | 0,65214261 | 0,00106298 | 0,0126994 | MRPL53 | -0,6167406 |
| ENSG000001 | gene | 0,90935253 | 0,00106833 | 0,01274163 | GPI | -0,1370884 |
| ENSG000001 | gene | 1,28510356 | 0,00106908 | 0,01274163 | CCDC14 | 0,36188462 |
| ENSG000002 | gene | 1,21639991 | 0,00107253 | 0,01276732 | CBX3P9 | 0,28261762 |
| ENSG000001 | gene | 0,91389719 | 0,00108165 | 0,01286052 | DNAJA3 | -0,1298962 |
| ENSG000000 | gene | 1,14559853 | 0,00108637 | 0,01290111 | DLD | 0,19610155 |
| ENSG000001 | gene | 2,00692708 | 0,00108775 | 0,01290206 | DGLUCY | 1,0049882 |
| ENSG000002 | gene | 1,43023938 | 0,00109287 | 0,01294728 | AC115618.2 | 0,51625664 |
| ENSG000002 | gene | 1,38838838 | 0,00109549 | 0,01296292 | AC107398.2 | 0,47341119 |
| ENSG000002 | gene | 0,78060099 | 0,00109979 | 0,01299824 | U2AF1L5 | -0,3573428 |
| ENSG000000 | gene | 0,82207773 | 0,00110374 | 0,01302942 | CYBA | -0,2826533 |
| ENSG000001 | gene | 1,10008424 | 0,00110962 | 0,0130832 | DDX17 | 0,137614 |
| ENSG000000 | gene | 0,73656816 | 0,00112668 | 0,01326865 | PARP12 | -0,4411091 |
| ENSG000002 | gene | 0,63347342 | 0,00112894 | 0,01327948 | FP236383.4 | -0,658644 |
| ENSG000000 | gene | 1,16225154 | 0,0011344 | 0,01332252 | MPP5 | 0,21692234 |
| ENSG000000 | gene | 0,88072384 | 0,00113663 | 0,01332252 | GRAMD1A | -0,1832384 |
| ENSG000002 | gene | 0,7428534 | 0,00113663 | 0,01332252 | MTFP1 | -0,4288506 |
| ENSG000001 | gene | 0,87990821 | 0,00114021 | 0,01334858 | GNB2 | -0,1845751 |
| ENSG000002 | gene | 2,02196188 | 0,00114155 | 0,01334858 | AC027319.1 | 1,0157558 |
| ENSG000001 | gene | 0,84842455 | 0,00114412 | 0,01336284 | CNPPD1 | -0,2371417 |
| ENSG000001 | gene | 0,83704153 | 0,0011509 | 0,0133993 | HAX1 | -0,2566289 |
| ENSG000001 | gene | 1,23268075 | 0,00115125 | 0,0133993 | NES | 0,3017992 |
| ENSG000002 | gene | 1,31801406 | 0,00115129 | 0,0133993 | AC083798.2 | 0,39836576 |
| ENSG000002 | gene | 1,75162392 | 0,00115605 | 0,01342319 | AL451085.3 | 0,80869306 |

| | | | | | | |
|---|---|---|---|---|---|---|
| ENSG00000( | gene | 1,09909398 | 0,00115605 | 0,01342319 | HECTD1 | 0,13631475 |
| ENSG00000 | gene | 0,83228297 | 0,00116659 | 0,01351448 | TERF2 | -0,264854 |
| ENSG00000 | gene | 1,35087988 | 0,00116664 | 0,01351448 | CPED1 | 0,4338994 |
| ENSG00000( | gene | 1,26312178 | 0,00117342 | 0,01357713 | CFH | 0,33699374 |
| ENSG00000 | gene | 0,89176281 | 0,00117809 | 0,0136153 | PSMB4 | -0,1652681 |
| ENSG00000 | gene | 0,82934709 | 0,00118106 | 0,01362879 | DNAJB6 | -0,2699521 |
| ENSG00000; | gene | 0,89918632 | 0,001182 | 0,01362879 | EIF6 | -0,153308 |
| ENSG00000 | gene | 1,2995416 | 0,00118898 | 0,0136933 | PLCB4 | 0,37800282 |
| ENSG00000 | gene | 0,81006211 | 0,00119238 | 0,01371651 | RPL23AP82 | -0,3038956 |
| ENSG00000 | gene | 0,84292983 | 0,00119386 | 0,01371763 | TOM1 | -0,2465156 |
| ENSG00000; | gene | 1,619471 | 0,00119836 | 0,01375345 | RBM5-AS1 | 0,69552263 |
| ENSG00000( | gene | 1,10820644 | 0,00120634 | 0,01379862 | SEC63 | 0,14822666 |
| ENSG00000( | gene | 0,89513056 | 0,00120815 | 0,01379862 | ERCC1 | -0,15983 |
| ENSG00000; | gene | 0,72033484 | 0,0012082 | 0,01379862 | PRR13 | -0,4732604 |
| ENSG00000 | gene | 1,28985797 | 0,00120931 | 0,01379862 | MRAP | 0,36721222 |
| ENSG00000; | gene | 1,50427883 | 0,00120991 | 0,01379862 | RNU6-26P | 0,58907201 |
| ENSG00000 | gene | 0,85205621 | 0,00121204 | 0,01379862 | CDC42SE1 | -0,2309795 |
| ENSG00000 | gene | 0,88792152 | 0,00121204 | 0,01379862 | PSMB7 | -0,1714959 |
| ENSG00000; | gene | 0,85705084 | 0,00121485 | 0,01380147 | GET4 | -0,2225473 |
| ENSG00000 | gene | 0,84315065 | 0,00121543 | 0,01380147 | TTC9C | -0,2461377 |
| ENSG00000; | gene | 1,72891397 | 0,00121647 | 0,01380147 | AC025370.1 | 0,78986608 |
| ENSG00000 | gene | 0,83888813 | 0,00121855 | 0,01380929 | ABHD4 | -0,2534497 |
| ENSG00000 | gene | 1,08915328 | 0,00122441 | 0,01385594 | CMAS | 0,123207 |
| ENSG00000 | gene | 1,6520151 | 0,00122546 | 0,01385594 | TRPC1 | 0,72422687 |
| ENSG00000 | gene | 0,55968295 | 0,0012312 | 0,01390495 | IFI44 | -0,8373183 |
| ENSG00000 | gene | 1,14242114 | 0,00124759 | 0,01407412 | NAA25 | 0,19209458 |
| ENSG00000 | gene | 1,12436805 | 0,00125509 | 0,01414259 | ANXA1 | 0,16911437 |
| ENSG00000 | gene | 1,12356562 | 0,00125748 | 0,01415348 | ZNF12 | 0,16808438 |
| ENSG00000( | gene | 1,17512807 | 0,00126365 | 0,01420677 | NUCKS1 | 0,232818 |
| ENSG00000 | gene | 0,81888414 | 0,00126673 | 0,0142107 | ANAPC11 | -0,2882687 |
| ENSG00000 | gene | 0,55136884 | 0,00126707 | 0,0142107 | AREG | -0,8589103 |
| ENSG00000 | gene | 0,92733063 | 0,0012683 | 0,0142107 | LRRC59 | -0,1088443 |
| ENSG00000 | gene | 0,90359891 | 0,00127814 | 0,01430026 | GTPBP6 | -0,1462456 |
| ENSG00000; | gene | 1,74186019 | 0,00127918 | 0,01430026 | AC004542.1 | 0,80062883 |
| ENSG00000 | gene | 0,85436148 | 0,00128251 | 0,01432139 | PGD | -0,2270815 |
| ENSG00000 | gene | 0,89793894 | 0,00128879 | 0,01437528 | CTNNBL1 | -0,1553108 |
| ENSG00000; | gene | 0,8746865 | 0,00129396 | 0,01440195 | HEXA | -0,1931621 |
| ENSG00000; | gene | 1,69073023 | 0,00129408 | 0,01440195 | AL513217.1 | 0,75764648 |
| ENSG00000 | gene | 0,64285832 | 0,00129743 | 0,014423 | CLN3 | -0,6374273 |
| ENSG00000; | gene | 0,8133834 | 0,0012998 | 0,01443312 | SNHG32 | -0,2979925 |
| ENSG00000 | gene | 0,92748826 | 0,001305 | 0,01447368 | VPS11 | -0,1085991 |
| ENSG00000 | gene | 1,16024099 | 0,00130637 | 0,01447368 | KPNA3 | 0,21442449 |
| ENSG00000 | gene | 0,85554202 | 0,00131008 | 0,01449866 | NSUN5 | -0,2250894 |
| ENSG00000 | gene | 1,25521903 | 0,00131563 | 0,01453022 | UBR3 | 0,32793913 |
| ENSG00000; | gene | 1,30516895 | 0,00131708 | 0,01453022 | RPL7P23 | 0,38423657 |
| ENSG00000 | gene | 0,88394104 | 0,00131733 | 0,01453022 | SELENOS | -0,177978 |
| ENSG00000 | gene | 1,23017735 | 0,00132207 | 0,01455154 | FAR1 | 0,29886632 |
| ENSG00000 | gene | 1,09574877 | 0,0013222 | 0,01455154 | MAK16 | 0,13191705 |
| ENSG00000 | gene | 0,83307705 | 0,0013404 | 0,01473551 | PITPNM1 | -0,2634782 |
| ENSG00000 | gene | 0,87280609 | 0,00134378 | 0,01475631 | CAMK2G | -0,1962669 |
| ENSG00000 | gene | 1,09925636 | 0,00134597 | 0,0147587 | PRKCA | 0,13652788 |
| ENSG00000 | gene | 0,89601394 | 0,00134698 | 0,0147587 | ATG4B | -0,1584069 |
| ENSG00000; | gene | 0,41680325 | 0,00135774 | 0,01483015 | RNA5-8SN3 | -1,2625616 |
| ENSG00000; | gene | 1,44112551 | 0,00135794 | 0,01483015 | AP003307.2 | 0,52719599 |
| ENSG00000 | gene | 0,8834395 | 0,00135976 | 0,01483015 | NELFA | -0,1787967 |
| ENSG00000 | gene | 0,75881433 | 0,00136081 | 0,01483015 | KXD1 | -0,3981812 |
| ENSG00000( | gene | 1,22881668 | 0,00136122 | 0,01483015 | LZTS1 | 0,2972697 |

| | | | | | | |
|---|---|---|---|---|---|---|
| ENSG00000' | gene | 0,77245879 | 0,00136247 | 0,01483015 | MYL6B | -0,3724701 |
| ENSG00002; | gene | 1,29934668 | 0,00136667 | 0,01485389 | AL353763.2 | 0,37778641 |
| ENSG00000( | gene | 1,21520086 | 0,00136845 | 0,01485389 | ESF1 | 0,2811948 |
| ENSG00002; | gene | 0,88739921 | 0,00136964 | 0,01485389 | BAG6 | -0,1723448 |
| ENSG00002; | gene | 1,74895287 | 0,00137064 | 0,01485389 | AC027601.4 | 0,80649141 |
| ENSG00000' | gene | 1,16747185 | 0,00137533 | 0,01488835 | GOLIM4 | 0,22338777 |
| ENSG00000' | gene | 0,81464425 | 0,00138339 | 0,01495066 | WDR74 | -0,2957579 |
| ENSG00000' | gene | 1,1636935 | 0,0013841 | 0,01495066 | PLS3 | 0,21871113 |
| ENSG00000( | gene | 0,88775935 | 0,00138785 | 0,01496336 | TCF3 | -0,1717594 |
| ENSG00000' | gene | 1,1881831 | 0,00138829 | 0,01496336 | DSEL | 0,24875718 |
| ENSG00002; | gene | 1,41231971 | 0,00139044 | 0,0149702 | AC026347.2 | 0,49806671 |
| ENSG00000' | gene | 1,12175841 | 0,00139266 | 0,01497792 | BICC1 | 0,165762 |
| ENSG00000' | gene | 0,88525046 | 0,0013957 | 0,01499438 | UNC50 | -0,1758424 |
| ENSG00002; | gene | 1,36611165 | 0,00140238 | 0,01504981 | AC018362.2 | 0,4500754 |
| ENSG00000' | gene | 1,10931605 | 0,00141185 | 0,01512646 | BAZ1A | 0,14967045 |
| ENSG00002; | gene | 0,79080457 | 0,00141258 | 0,01512646 | SNRPEP4 | -0,3386069 |
| ENSG00000' | gene | 0,85096494 | 0,0014159 | 0,01514566 | TBRG4 | -0,2328284 |
| ENSG00002; | gene | 1,60083914 | 0,00142076 | 0,01518136 | AL355816.2 | 0,67882835 |
| ENSG00000' | gene | 0,85044688 | 0,00142595 | 0,01522033 | NDUFV1 | -0,233707 |
| ENSG00000' | gene | 0,81130872 | 0,00142861 | 0,01523239 | ALDOC | -0,3016771 |
| ENSG00000' | gene | 1,11172076 | 0,00144062 | 0,01534392 | EPS8 | 0,15279447 |
| ENSG00000( | gene | 0,881891 | 0,00144787 | 0,0153936 | TRAF4 | -0,1813277 |
| ENSG00002; | gene | 1,40248636 | 0,00144839 | 0,0153936 | AL512791.1 | 0,48798674 |
| ENSG00002; | gene | 1,31888841 | 0,00145098 | 0,0154046 | VCAN-AS1 | 0,3993225 |
| ENSG00000( | gene | 1,21987059 | 0,00145929 | 0,01546852 | C12orf4 | 0,2867281 |
| ENSG00000' | gene | 0,83667944 | 0,00146042 | 0,01546852 | SNF8 | -0,2572531 |
| ENSG00000' | gene | 0,82887892 | 0,00146406 | 0,01546852 | SNHG29 | -0,2707667 |
| ENSG00000' | gene | 1,27513673 | 0,00146571 | 0,01546852 | SFT2D3 | 0,35065196 |
| ENSG00002; | gene | 1,20800208 | 0,00146712 | 0,01546852 | CD302 | 0,27262294 |
| ENSG00000' | gene | 0,83284177 | 0,00146765 | 0,01546852 | CDK2AP2 | -0,2638857 |
| ENSG00000( | gene | 0,92467601 | 0,00146792 | 0,01546852 | KARS1 | -0,1129801 |
| ENSG00000' | gene | 0,83675849 | 0,00147148 | 0,01548158 | GSTO1 | -0,2571168 |
| ENSG00000' | gene | 1,08688486 | 0,00147306 | 0,01548158 | RSPRY1 | 0,12019912 |
| ENSG00000' | gene | 1,24220169 | 0,00147508 | 0,01548158 | DIAPH3 | 0,31289943 |
| ENSG00000' | gene | 0,63537683 | 0,0014754 | 0,01548158 | IFIT3 | -0,6543156 |
| ENSG00000' | gene | 1,7568318 | 0,00147997 | 0,01550331 | CAPN12 | 0,81297607 |
| ENSG00000' | gene | 1,30809285 | 0,00148145 | 0,01550331 | CIP2A | 0,38746495 |
| ENSG00000' | gene | 1,43722478 | 0,00148216 | 0,01550331 | TCP11L1 | 0,52328572 |
| ENSG00000' | gene | 1,23252365 | 0,00148823 | 0,01555033 | CACNA2D1 | 0,30161532 |
| ENSG00000' | gene | 1,18169175 | 0,00149948 | 0,01563994 | CMPK1 | 0,24085376 |
| ENSG00000' | gene | 0,7407591 | 0,00149996 | 0,01563994 | ATP5MG | -0,4329237 |
| ENSG00000' | gene | 1,69200696 | 0,00150469 | 0,01567285 | PHLDB2 | 0,7587355 |
| ENSG00000' | gene | 0,87635886 | 0,00151047 | 0,01568512 | CDC26 | -0,1904063 |
| ENSG00000' | gene | 0,82702514 | 0,0015114 | 0,01568512 | RHOC | -0,2739969 |
| ENSG00000' | gene | 1,19288206 | 0,00151314 | 0,01568512 | SLC39A10 | 0,25445141 |
| ENSG00000' | gene | 1,11635653 | 0,00151364 | 0,01568512 | COG6 | 0,15879785 |
| ENSG00000' | gene | 1,14624169 | 0,00151378 | 0,01568512 | MPHOSPH1( | 0,19691128 |
| ENSG00000' | gene | 0,74542679 | 0,00151657 | 0,0156976 | CAPN10 | -0,4238614 |
| ENSG00000' | gene | 0,85070386 | 0,00151943 | 0,0157108 | MAPK8IP1 | -0,2332711 |
| ENSG00000' | gene | 1,18389966 | 0,00153113 | 0,01581527 | LANCL1 | 0,24354681 |
| ENSG00000' | gene | 0,77868489 | 0,00155279 | 0,01599786 | MIEN1 | -0,3608885 |
| ENSG00000' | gene | 0,89795349 | 0,00155306 | 0,01599786 | PHLDA2 | -0,1552874 |
| ENSG00000' | gene | 0,84947034 | 0,00155394 | 0,01599786 | TBC1D17 | -0,2353645 |
| ENSG00000' | gene | 0,81603978 | 0,00155583 | 0,01599786 | FADS2 | -0,2932886 |
| ENSG00002; | gene | 1,38180389 | 0,00155687 | 0,01599786 | AC006064.4 | 0,46655288 |
| ENSG00000' | gene | 0,85481026 | 0,00156811 | 0,01606241 | TMEM208 | -0,2263239 |
| ENSG00000' | gene | 1,1315864 | 0,00156817 | 0,01606241 | SLAIN2 | 0,17834674 |

| | | | | | | |
|---|---|---|---|---|---|---|
| ENSG00000 | gene | 1,20737475 | 0,00156822 | 0,01606241 | ELMOD2 | 0,27187354 |
| ENSG00000 | gene | 1,71856843 | 0,00157071 | 0,01606241 | AC090971.2 | 0,78120729 |
| ENSG00000 | gene | 0,73497004 | 0,00157283 | 0,01606241 | FBRSL1 | -0,4442427 |
| ENSG00000 | gene | 1,34385382 | 0,0015735 | 0,01606241 | NUF2 | 0,42637621 |
| ENSG00000 | gene | 0,86048887 | 0,00157449 | 0,01606241 | STK25 | -0,2167716 |
| ENSG00000 | gene | 1,28576987 | 0,00157793 | 0,01608092 | AP001001.1 | 0,36263245 |
| ENSG00000 | gene | 0,92832841 | 0,00158007 | 0,01608625 | MANBAL | -0,1072928 |
| ENSG00000 | gene | 0,82382178 | 0,0015825 | 0,01609447 | FYN | -0,2795958 |
| ENSG00000 | gene | 1,23575704 | 0,00158686 | 0,01612223 | TRHDE | 0,30539513 |
| ENSG00000 | gene | 0,77558097 | 0,00158851 | 0,01612249 | SPNS1 | -0,3666507 |
| ENSG00000 | gene | 0,703759 | 0,00160264 | 0,01624924 | PML | -0,5068466 |
| ENSG00000 | gene | 0,91196852 | 0,00161042 | 0,01629967 | NUDC | -0,1329441 |
| ENSG00000 | gene | 0,86052447 | 0,00161304 | 0,01629967 | GFER | -0,2167119 |
| ENSG00000 | gene | 1,37340083 | 0,00161495 | 0,01629967 | AL136038.4 | 0,45775274 |
| ENSG00000 | gene | 0,86962798 | 0,00161559 | 0,01629967 | WDR25 | -0,2015297 |
| ENSG00000 | gene | 1,87447538 | 0,00161583 | 0,01629967 | AC092115.3 | 0,90648688 |
| ENSG00000 | gene | 1,47648475 | 0,0016221 | 0,01634632 | AC004079.2 | 0,56216645 |
| ENSG00000 | gene | 1,20773038 | 0,0016255 | 0,01636397 | EDEM3 | 0,27229842 |
| ENSG00000 | gene | 0,89739293 | 0,00163782 | 0,01647118 | SRPRB | -0,1561883 |
| ENSG00000 | gene | 0,93279003 | 0,00164368 | 0,01651338 | CHMP4B | -0,1003757 |
| ENSG00000 | gene | 0,77764842 | 0,00164853 | 0,01654532 | AC008761.1 | -0,36281 |
| ENSG00000 | gene | 0,85043367 | 0,00165041 | 0,01654746 | STIM1 | -0,2337294 |
| ENSG00000 | gene | 0,83971538 | 0,00165596 | 0,01657428 | CIAO2B | -0,2520277 |
| ENSG00000 | gene | 0,87545466 | 0,00165677 | 0,01657428 | DNASE2 | -0,1918956 |
| ENSG00000 | gene | 1,24091273 | 0,00165961 | 0,01657428 | WASHC4 | 0,31140166 |
| ENSG00000 | gene | 0,88681846 | 0,00166107 | 0,01657428 | OSBPL2 | -0,1732893 |
| ENSG00000 | gene | 0,7972134 | 0,00166144 | 0,01657428 | FST | -0,3269621 |
| ENSG00000 | gene | 0,92807082 | 0,00166902 | 0,01663313 | GUCD1 | -0,1076932 |
| ENSG00000 | gene | 0,8985877 | 0,0016709 | 0,01663517 | PFDN2 | -0,1542688 |
| ENSG00000 | gene | 0,90307284 | 0,00167389 | 0,01664823 | TCTN2 | -0,1470857 |
| ENSG00000 | gene | 0,74407383 | 0,00168251 | 0,01670993 | NPR3 | -0,4264823 |
| ENSG00000 | gene | 1,39809732 | 0,00168346 | 0,01670993 | AC104117.2 | 0,48346478 |
| ENSG00000 | gene | 1,45411993 | 0,00168777 | 0,01673596 | AC005253.2 | 0,54014627 |
| ENSG00000 | gene | 0,92765197 | 0,00170114 | 0,01685163 | NELFE | -0,1083444 |
| ENSG00000 | gene | 1,13154896 | 0,00170647 | 0,01687717 | RPS6KB1 | 0,178299 |
| ENSG00000 | gene | 0,8445118 | 0,00170837 | 0,01687717 | DPM2 | -0,2438105 |
| ENSG00000 | gene | 0,9069156 | 0,00170882 | 0,01687717 | ZNF511 | -0,1409598 |
| ENSG00000 | gene | 0,86907081 | 0,00171386 | 0,01691013 | DHRS7 | -0,2024544 |
| ENSG00000 | gene | 1,1055146 | 0,00171607 | 0,01691502 | SDAD1 | 0,14471808 |
| ENSG00000 | gene | 0,90159844 | 0,00171821 | 0,01691935 | CLPP | -0,1494431 |
| ENSG00000 | gene | 0,42762245 | 0,00172161 | 0,01693596 | AL353644.2 | -1,2255905 |
| ENSG00000 | gene | 1,61074776 | 0,00172445 | 0,01694712 | AL359632.1 | 0,68773058 |
| ENSG00000 | gene | 1,74348412 | 0,00173911 | 0,01706691 | PORCN | 0,80197322 |
| ENSG00000 | gene | 1,77262761 | 0,0017401 | 0,01706691 | AC105020.4 | 0,82588949 |
| ENSG00000 | gene | 0,81222406 | 0,00174302 | 0,01706691 | BMERB1 | -0,3000503 |
| ENSG00000 | gene | 1,28244944 | 0,00174352 | 0,01706691 | AC021088.1 | 0,35890195 |
| ENSG00000 | gene | 1,14225875 | 0,0017477 | 0,01707469 | NAV3 | 0,19188949 |
| ENSG00000 | gene | 0,77217392 | 0,00174878 | 0,01707469 | MT-ATP6 | -0,3730023 |
| ENSG00000 | gene | 0,83629107 | 0,00175098 | 0,01707469 | PSMD7 | -0,2579229 |
| ENSG00000 | gene | 1,52549309 | 0,00175121 | 0,01707469 | AC096772.2 | 0,60927565 |
| ENSG00000 | gene | 1,38168908 | 0,0017615 | 0,01715821 | ACBD3-AS1 | 0,466433 |
| ENSG00000 | gene | 0,89232717 | 0,00176674 | 0,01719229 | MPRIP | -0,1643553 |
| ENSG00000 | gene | 1,33003198 | 0,00178282 | 0,01733182 | AC068831.5 | 0,41146093 |
| ENSG00000 | gene | 1,76047167 | 0,00179359 | 0,01741939 | AL138478.1 | 0,81596201 |
| ENSG00000 | gene | 1,16023877 | 0,00179544 | 0,01742034 | SNTB2 | 0,21442173 |
| ENSG00000 | gene | 0,7931179 | 0,0018131 | 0,01757453 | AC025165.2 | -0,3343928 |
| ENSG00000 | gene | 0,8880685 | 0,00181599 | 0,01758533 | ESS2 | -0,1712571 |

| | | | | | | |
|---|---|---|---|---|---|---|
| ENSG00000' | gene | 0,83343739 | 0,00182005 | 0,01760744 | DDB2 | -0,2628543 |
| ENSG00000( | gene | 0,84342558 | 0,00182329 | 0,01762161 | AK2 | -0,2456673 |
| ENSG00002 | gene | 1,24113538 | 0,00182745 | 0,0176389 | AC104116.1 | 0,31166048 |
| ENSG00000( | gene | 0,90583613 | 0,00182864 | 0,0176389 | UBE3C | -0,142678 |
| ENSG00000' | gene | 0,84901361 | 0,00183186 | 0,01765276 | CHMP7 | -0,2361404 |
| ENSG00000' | gene | 0,67552349 | 0,00183908 | 0,01770516 | ZNF667-AS1 | -0,5659222 |
| ENSG00000' | gene | 0,89568328 | 0,00184108 | 0,0177072 | RHOG | -0,1589394 |
| ENSG00000' | gene | 1,3726819 | 0,00184461 | 0,017724 | MKI67 | 0,45699734 |
| ENSG00000' | gene | 0,8367337 | 0,00187375 | 0,01798659 | EML2 | -0,2571595 |
| ENSG00000' | gene | 0,86742484 | 0,00187786 | 0,01798831 | SEC61G | -0,2051893 |
| ENSG00000( | gene | 0,85863693 | 0,00187932 | 0,01798831 | SLC9A3 | -0,2198799 |
| ENSG00000' | gene | 1,06523299 | 0,00187938 | 0,01798831 | BTBD10 | 0,09116902 |
| ENSG00000' | gene | 1,11518271 | 0,00188627 | 0,0180369 | NUDCD1 | 0,1572801 |
| ENSG00000' | gene | 0,84848322 | 0,0018893 | 0,01804843 | SPCS2 | -0,237042 |
| ENSG00000' | gene | 1,3730617 | 0,00190203 | 0,01814421 | JMJD1C | 0,45739646 |
| ENSG00000' | gene | 0,80711627 | 0,00190298 | 0,01814421 | WBP2 | -0,3091516 |
| ENSG00000' | gene | 1,07798943 | 0,00190854 | 0,01817356 | TRIP12 | 0,10834303 |
| ENSG00002 | gene | 0,92058003 | 0,00191118 | 0,01817356 | GDI1 | -0,1193849 |
| ENSG00002 | gene | 1,36154993 | 0,00191156 | 0,01817356 | ANO1-AS1 | 0,44524989 |
| ENSG00000' | gene | 0,7175758 | 0,00191916 | 0,0182283 | NICN1 | -0,4787969 |
| ENSG00002 | gene | 1,53689131 | 0,0019218 | 0,01823594 | AL353689.2 | 0,62001514 |
| ENSG00000' | gene | 0,71849459 | 0,0019279 | 0,0182763 | IFIH1 | -0,4769508 |
| ENSG00000' | gene | 0,88677801 | 0,00193019 | 0,01828062 | DPP9 | -0,1733551 |
| ENSG00000' | gene | 0,92480261 | 0,00193532 | 0,01829852 | FAM32A | -0,1127826 |
| ENSG00000' | gene | 0,72753467 | 0,00193577 | 0,01829852 | UBE2Z | -0,4589121 |
| ENSG00000' | gene | 1,2409096 | 0,00194231 | 0,01834283 | AIDA | 0,31139802 |
| ENSG00002 | gene | 1,57390193 | 0,00194553 | 0,01835575 | AC091117.2 | 0,65434565 |
| ENSG00000' | gene | 0,85858153 | 0,00194755 | 0,01835731 | GLIS2 | -0,219973 |
| ENSG00002 | gene | 0,85005089 | 0,00195124 | 0,01837465 | AL590714.1 | -0,2343789 |
| ENSG00000' | gene | 1,34071768 | 0,00195451 | 0,01838793 | SCN9A | 0,42300547 |
| ENSG00002 | gene | 1,45187571 | 0,00195987 | 0,01840864 | AC097534.1 | 0,53791795 |
| ENSG00002 | gene | 1,31776088 | 0,00196042 | 0,01840864 | AC116317.1 | 0,39808861 |
| ENSG00000' | gene | 1,0918876 | 0,00196279 | 0,01841348 | TYW3 | 0,12682436 |
| ENSG00000' | gene | 0,8849384 | 0,0019669 | 0,01843461 | ATP6V0E1 | -0,1763511 |
| ENSG00000' | gene | 0,78127483 | 0,00197077 | 0,01845338 | TPP1 | -0,356098 |
| ENSG00000' | gene | 0,81105556 | 0,00197618 | 0,0184787 | PTGES3 | -0,3021273 |
| ENSG00000' | gene | 1,11490514 | 0,0019772 | 0,0184787 | EID2 | 0,15692097 |
| ENSG00002 | gene | 0,59904509 | 0,00197912 | 0,01847928 | AARSD1 | -0,7392635 |
| ENSG00002 | gene | 1,57198332 | 0,00198473 | 0,01851422 | AC012640.6 | 0,65258591 |
| ENSG00002 | gene | 1,24022518 | 0,00199332 | 0,0185749 | AL450467.1 | 0,31060209 |
| ENSG00002 | gene | 1,59005575 | 0,00199683 | 0,0185749 | SNORA24 | 0,66907735 |
| ENSG00000' | gene | 0,91160659 | 0,00199686 | 0,0185749 | SPCS1 | -0,1335167 |
| ENSG00002 | gene | 1,24711907 | 0,00200451 | 0,01862865 | MT-TP | 0,31859922 |
| ENSG00000' | gene | 1,34085991 | 0,00201369 | 0,01869297 | INPP5F | 0,42315851 |
| ENSG00000' | gene | 0,70849067 | 0,00201521 | 0,01869297 | SLC35B2 | -0,4971792 |
| ENSG00000' | gene | 1,24820648 | 0,00201869 | 0,0187078 | SSX2IP | 0,3198566 |
| ENSG00000' | gene | 0,911883 | 0,00203347 | 0,01881967 | TMED9 | -0,1330794 |
| ENSG00002 | gene | 1,58806598 | 0,00203522 | 0,01881967 | AL110504.1 | 0,66727086 |
| ENSG00000' | gene | 0,82314161 | 0,00203646 | 0,01881967 | LGALS3BP | -0,2807875 |
| ENSG00000( | gene | 1,14053572 | 0,00205692 | 0,01899109 | NUCB2 | 0,18971163 |
| ENSG00000' | gene | 0,83960956 | 0,00206504 | 0,01904832 | RPS11 | -0,2522095 |
| ENSG00000' | gene | 0,91082946 | 0,00207984 | 0,01916699 | THAP4 | -0,1347471 |
| ENSG00000' | gene | 0,90452342 | 0,00208387 | 0,01918635 | KANSL2 | -0,1447702 |
| ENSG00000' | gene | 0,87537276 | 0,00211167 | 0,01938886 | SEM1 | -0,1920306 |
| ENSG00000' | gene | 0,88484221 | 0,00211247 | 0,01938886 | PCGF3 | -0,1765079 |
| ENSG00000' | gene | 0,8232434 | 0,00211331 | 0,01938886 | SSR2 | -0,2806091 |
| ENSG00002 | gene | 1,35834354 | 0,00211369 | 0,01938886 | ZNF814 | 0,4418484 |

| | | | | | | |
|---|---|---|---|---|---|---|
| ENSG00000 | gene | 0,89047629 | 0,00211614 | 0,01939338 | MAGOH | -0,1673509 |
| ENSG00000 | gene | 0,62434437 | 0,00211829 | 0,01939521 | SOD2 | -0,6795861 |
| ENSG00000 | gene | 1,19060688 | 0,00212145 | 0,01940614 | ARID5B | 0,25169714 |
| ENSG00000 | gene | 0,8721079 | 0,00212819 | 0,01944985 | DLG4 | -0,1974214 |
| ENSG00000 | gene | 1,32016502 | 0,00213656 | 0,01950844 | CASK-AS1 | 0,40071828 |
| ENSG00000 | gene | 1,10255851 | 0,00214015 | 0,01952324 | CTR9 | 0,14085522 |
| ENSG00000 | gene | 1,23041298 | 0,00214464 | 0,01953894 | DOCK7 | 0,29914263 |
| ENSG00000 | gene | 0,87343305 | 0,00214582 | 0,01953894 | SLC25A39 | -0,195231 |
| ENSG00000 | gene | 1,13136725 | 0,00215789 | 0,01955413 | NDUFAB1 | 0,17806732 |
| ENSG00000 | gene | 1,43058164 | 0,00215855 | 0,01955413 | AC012615.2 | 0,51660183 |
| ENSG00000 | gene | 1,30372587 | 0,0021588 | 0,01955413 | AC011497.2 | 0,38264055 |
| ENSG00000 | gene | 0,84377592 | 0,00215971 | 0,01955413 | AC141424.1 | -0,2450682 |
| ENSG00000 | gene | 0,86005565 | 0,00216005 | 0,01955413 | POR | -0,2174981 |
| ENSG00000 | gene | 1,22611158 | 0,00216055 | 0,01955413 | AC016026.1 | 0,29409027 |
| ENSG00000 | gene | 0,90716043 | 0,00216129 | 0,01955413 | FHL3 | -0,1405704 |
| ENSG00000 | gene | 0,81358379 | 0,00216373 | 0,0195584 | SNAPIN | -0,2976372 |
| ENSG00000 | gene | 0,83659409 | 0,0021716 | 0,01961162 | RPL24 | -0,2574003 |
| ENSG00000 | gene | 0,85857019 | 0,00218425 | 0,01968554 | YEATS2 | -0,219992 |
| ENSG00000 | gene | 1,25884104 | 0,00218541 | 0,01968554 | G2E3 | 0,33209612 |
| ENSG00000 | gene | 1,25865443 | 0,00218574 | 0,01968554 | TWF1 | 0,33188224 |
| ENSG00000 | gene | 0,80300873 | 0,00218919 | 0,0196987 | NUPR1 | -0,3165124 |
| ENSG00000 | gene | 0,88191513 | 0,00219494 | 0,0197326 | BCAR1 | -0,1812883 |
| ENSG00000 | gene | 1,23347583 | 0,00220163 | 0,01977475 | NUDT17 | 0,30272945 |
| ENSG00000 | gene | 1,32516372 | 0,00220831 | 0,01980979 | AC019257.9 | 0,40617061 |
| ENSG00000 | gene | 0,89755115 | 0,00220952 | 0,01980979 | GRK6 | -0,1559339 |
| ENSG00000 | gene | 0,88010574 | 0,00221449 | 0,0198364 | TIMM17B | -0,1842512 |
| ENSG00000 | gene | 1,10295661 | 0,00222071 | 0,01983817 | INTS13 | 0,14137604 |
| ENSG00000 | gene | 0,87990337 | 0,00222151 | 0,01983817 | GSK3A | -0,184583 |
| ENSG00000 | gene | 0,90757853 | 0,00222256 | 0,01983817 | FBH1 | -0,1399056 |
| ENSG00000 | gene | 0,6984419 | 0,00222269 | 0,01983817 | TCEAL4 | -0,517788 |
| ENSG00000 | gene | 0,86327115 | 0,00222672 | 0,01985208 | HPCAL1 | -0,2121143 |
| ENSG00000 | gene | 0,85049127 | 0,00223036 | 0,01985208 | GEMIN7 | -0,2336317 |
| ENSG00000 | gene | 0,82439017 | 0,00223292 | 0,01985208 | NPC2 | -0,2786008 |
| ENSG00000 | gene | 1,64561957 | 0,00223367 | 0,01985208 | XRN1 | 0,71863086 |
| ENSG00000 | gene | 0,79516189 | 0,00223648 | 0,01985208 | CFL1 | -0,3306795 |
| ENSG00000 | gene | 0,87585941 | 0,00223772 | 0,01985208 | DNAJC11 | -0,1912288 |
| ENSG00000 | gene | 0,77566502 | 0,00223826 | 0,01985208 | C11orf24 | -0,3664944 |
| ENSG00000 | gene | 0,78945022 | 0,00224555 | 0,01989886 | ENDOV | -0,3410798 |
| ENSG00000 | gene | 1,09281755 | 0,00225188 | 0,01991929 | ADSS2 | 0,12805255 |
| ENSG00000 | gene | 0,83511147 | 0,00225331 | 0,01991929 | PLEKHM2 | -0,2599593 |
| ENSG00000 | gene | 0,83234401 | 0,00225388 | 0,01991929 | CXXC1 | -0,2647482 |
| ENSG00000 | gene | 1,19375728 | 0,00225773 | 0,01993554 | UBA6 | 0,25550954 |
| ENSG00000 | gene | 0,82972887 | 0,00226169 | 0,01994687 | ZSWIM8 | -0,2692881 |
| ENSG00000 | gene | 0,88631874 | 0,0022649 | 0,01994687 | ZNF622 | -0,1741025 |
| ENSG00000 | gene | 0,83367864 | 0,00226798 | 0,01994687 | NOP16 | -0,2624367 |
| ENSG00000 | gene | 1,3924218 | 0,00227392 | 0,01994687 | AL354956.1 | 0,4775963 |
| ENSG00000 | gene | 1,3236954 | 0,0022745 | 0,01994687 | STEAP1 | 0,40457118 |
| ENSG00000 | gene | 0,89020198 | 0,00227471 | 0,01994687 | DAXX | -0,1677954 |
| ENSG00000 | gene | 1,1344875 | 0,00227546 | 0,01994687 | MAN2A1 | 0,18204072 |
| ENSG00000 | gene | 0,68135765 | 0,00227601 | 0,01994687 | PRUNE1 | -0,5535158 |
| ENSG00000 | gene | 0,89024018 | 0,00227711 | 0,01994687 | MGAT4B | -0,1677335 |
| ENSG00000 | gene | 1,32675603 | 0,00228086 | 0,01995525 | AL451165.2 | 0,40790311 |
| ENSG00000 | gene | 1,29872347 | 0,00228209 | 0,01995525 | AP001496.3 | 0,37709428 |
| ENSG00000 | gene | 0,78048663 | 0,0022915 | 0,02001982 | ALKBH2 | -0,3575542 |
| ENSG00000 | gene | 1,33041104 | 0,00229365 | 0,02002096 | AC008569.1 | 0,41187205 |
| ENSG00000 | gene | 1,65390261 | 0,00229673 | 0,02003026 | CCDC142 | 0,72587429 |
| ENSG00000 | gene | 0,89290015 | 0,00230951 | 0,02012402 | DEF8 | -0,1634292 |

| | | | | | | |
|---|---|---|---|---|---|---|
| ENSG00000; | gene | 1,33901124 | 0,00231193 | 0,02012743 | AC090948.4 | 0,42116807 |
| ENSG00000 | gene | 0,84265037 | 0,00232658 | 0,02023714 | TLE5 | -0,2469939 |
| ENSG00000 | gene | 0,91210934 | 0,00233199 | 0,02025647 | PSAP | -0,1327213 |
| ENSG00000; | gene | 0,61670294 | 0,00233289 | 0,02025647 | AP000944.2 | -0,6973524 |
| ENSG00000 | gene | 1,44975963 | 0,00233675 | 0,02025948 | CCDC85B | 0,53581372 |
| ENSG00000( | gene | 0,84696688 | 0,00233732 | 0,02025948 | EPB41L3 | -0,2396225 |
| ENSG00000 | gene | 0,85707799 | 0,00233991 | 0,02026421 | RARA | -0,2225016 |
| ENSG00000; | gene | 1,37709139 | 0,00234233 | 0,02026747 | MIR4442 | 0,46162431 |
| ENSG00000 | gene | 0,93766303 | 0,00234951 | 0,02030525 | UBE2Q1 | -0,0928585 |
| ENSG00000 | gene | 0,93340302 | 0,00235079 | 0,02030525 | UBE2J2 | -0,099428 |
| ENSG00000 | gene | 1,124061 | 0,00236258 | 0,02038934 | ITGB1 | 0,16872032 |
| ENSG00000 | gene | 0,73543517 | 0,00236703 | 0,02040995 | APEX1 | -0,4433299 |
| ENSG00000( | gene | 0,91369998 | 0,00237669 | 0,02046377 | UIMC1 | -0,1302076 |
| ENSG00000; | gene | 1,48685263 | 0,0023774 | 0,02046377 | NAV2-AS1 | 0,57226166 |
| ENSG00000( | gene | 0,82086576 | 0,00238258 | 0,02047289 | MRPL43 | -0,2847818 |
| ENSG00000; | gene | 1,36949471 | 0,00238259 | 0,02047289 | AC011446.3 | 0,45364369 |
| ENSG00000 | gene | 0,88012952 | 0,00238815 | 0,02050293 | SUGP1 | -0,1842122 |
| ENSG00000 | gene | 1,08568978 | 0,00239352 | 0,02053131 | PPP3CB | 0,11861193 |
| ENSG00000 | gene | 0,8861509 | 0,00239658 | 0,02053902 | DAP3 | -0,1743757 |
| ENSG00000 | gene | 0,7517308 | 0,00240059 | 0,02053902 | RPAIN | -0,411712 |
| ENSG00000 | gene | 1,2444377 | 0,00240064 | 0,02053902 | ZNF146 | 0,315494 |
| ENSG00000 | gene | 1,21649356 | 0,00241021 | 0,02060313 | HAS2 | 0,28272869 |
| ENSG00000; | gene | 1,67821427 | 0,00241564 | 0,02063176 | AC015909.1 | 0,74692693 |
| ENSG00000 | gene | 0,72789629 | 0,00242081 | 0,02065811 | PFKFB3 | -0,4581952 |
| ENSG00000 | gene | 0,86620752 | 0,00242289 | 0,02065811 | APLP1 | -0,2072154 |
| ENSG00000; | gene | 1,36689955 | 0,00243323 | 0,02072687 | RNU6-395P | 0,45090722 |
| ENSG00000 | gene | 1,22396069 | 0,00243579 | 0,02072687 | FMN2 | 0,29155722 |
| ENSG00000; | gene | 1,21401029 | 0,00243723 | 0,02072687 | HMGA1P3 | 0,27978065 |
| ENSG00000( | gene | 1,06150495 | 0,00244113 | 0,0207423 | DDX1 | 0,0861111 |
| ENSG00000 | gene | 0,83733004 | 0,00245567 | 0,02083516 | IFT20 | -0,2561317 |
| ENSG00000( | gene | 1,17475475 | 0,00245766 | 0,02083516 | RRM2B | 0,2323596 |
| ENSG00000 | gene | 0,92509283 | 0,00245836 | 0,02083516 | HDLBP | -0,11233 |
| ENSG00000 | gene | 0,76612019 | 0,00246193 | 0,02084756 | PHF21A | -0,3843574 |
| ENSG00000 | gene | 0,8613739 | 0,00247388 | 0,0209309 | RAC3 | -0,2152885 |
| ENSG00000; | gene | 1,60330924 | 0,002478 | 0,02093237 | AC004079.1 | 0,68105271 |
| ENSG00000 | gene | 1,10734288 | 0,00247828 | 0,02093237 | CNOT7 | 0,14710201 |
| ENSG00000 | gene | 0,87934692 | 0,0024838 | 0,02096116 | RNF113A | -0,1854956 |
| ENSG00000 | gene | 0,81907521 | 0,00250025 | 0,02108205 | CALU | -0,2879322 |
| ENSG00000 | gene | 0,83157899 | 0,00251054 | 0,02115085 | ABCB9 | -0,2660748 |
| ENSG00000 | gene | 1,16558101 | 0,00252094 | 0,02122043 | CENPU | 0,22104927 |
| ENSG00000 | gene | 0,86004699 | 0,00253826 | 0,02134812 | UBAP1 | -0,2175126 |
| ENSG00000 | gene | 0,74282255 | 0,00254437 | 0,02136324 | CHTF8 | -0,4289105 |
| ENSG00000 | gene | 1,1931497 | 0,00254464 | 0,02136324 | CFAP97 | 0,25477506 |
| ENSG00000( | gene | 0,84479465 | 0,00255029 | 0,02136324 | MRPS18A | -0,2433274 |
| ENSG00000 | gene | 1,09831185 | 0,0025506 | 0,02136324 | GFM1 | 0,13528775 |
| ENSG00000 | gene | 1,55234097 | 0,00255137 | 0,02136324 | LRRC49 | 0,63444548 |
| ENSG00000 | gene | 1,40698186 | 0,00255299 | 0,02136324 | CTRL | 0,49260373 |
| ENSG00000 | gene | 1,06739982 | 0,00256684 | 0,021461 | CDC16 | 0,09410067 |
| ENSG00000; | gene | 1,88432694 | 0,00256921 | 0,02146274 | AC017083.2 | 0,9140493 |
| ENSG00000 | gene | 0,93160249 | 0,00257853 | 0,02152251 | IDH1 | -0,1022136 |
| ENSG00000 | gene | 0,73485913 | 0,00259015 | 0,02157709 | UBC | -0,4444604 |
| ENSG00000 | gene | 0,89893247 | 0,00259323 | 0,02157709 | MAGEH1 | -0,1537154 |
| ENSG00000 | gene | 0,66794492 | 0,00259374 | 0,02157709 | ZSCAN9 | -0,5821989 |
| ENSG00000( | gene | 0,84543222 | 0,00259378 | 0,02157709 | PHGDH | -0,242239 |
| ENSG00000; | gene | 1,62913679 | 0,00259756 | 0,02159047 | AC005540.1 | 0,70410774 |
| ENSG00000 | gene | 1,09283446 | 0,00260685 | 0,02164951 | CAND1 | 0,12807488 |
| ENSG00000 | gene | 0,94907093 | 0,00262124 | 0,02175085 | GLYR1 | -0,0754122 |

| | | | | | | |
|---|---|---|---|---|---|---|
| ENSG00000' | gene | 0,78322198 | 0,0026323 | 0,02182433 | USP3 | -0,3525068 |
| ENSG00000' | gene | 1,09578722 | 0,00263461 | 0,02182524 | PAK2 | 0,13196768 |
| ENSG00000' | gene | 0,83544383 | 0,00264402 | 0,02188489 | RAB13 | -0,2593853 |
| ENSG00000' | gene | 1,34580214 | 0,00266034 | 0,02199063 | CEP55 | 0,42846632 |
| ENSG00000; | gene | 1,66053364 | 0,00266123 | 0,02199063 | SNORA4 | 0,73164695 |
| ENSG00000' | gene | 0,87277781 | 0,00266604 | 0,02201199 | NARF | -0,1963137 |
| ENSG00000; | gene | 0,83544489 | 0,00267125 | 0,0220367 | PSMB8 | -0,2593834 |
| ENSG00000' | gene | 1,24315748 | 0,00267683 | 0,0220644 | ARMT1 | 0,31400906 |
| ENSG00000' | gene | 1,20155599 | 0,00268335 | 0,02209482 | FN1 | 0,26490388 |
| ENSG00000; | gene | 1,40263105 | 0,00268692 | 0,02209482 | AC078909.1 | 0,48813557 |
| ENSG00000; | gene | 1,53031998 | 0,00268721 | 0,02209482 | AF127577.1 | 0,61383334 |
| ENSG00000' | gene | 0,87383975 | 0,00269786 | 0,022164 | NBL1 | -0,1945594 |
| ENSG00000' | gene | 0,83467553 | 0,0027033 | 0,02219032 | SH3GLB2 | -0,2607126 |
| ENSG00000; | gene | 1,88240774 | 0,00272689 | 0,02236547 | AC008894.3 | 0,91257915 |
| ENSG00000' | gene | 1,15175666 | 0,00273196 | 0,02238851 | DICER1 | 0,20383594 |
| ENSG00000' | gene | 1,40529936 | 0,00275068 | 0,02252329 | SMC2 | 0,49087749 |
| ENSG00000' | gene | 1,50412294 | 0,00275536 | 0,02254306 | C17orf58 | 0,58892249 |
| ENSG00000' | gene | 1,49024425 | 0,00276188 | 0,02257419 | PBLD | 0,57554881 |
| ENSG00000' | gene | 1,12387605 | 0,00276464 | 0,02257419 | NFXL1 | 0,16848293 |
| ENSG00000' | gene | 0,87456062 | 0,002766 | 0,02257419 | CAMLG | -0,1933697 |
| ENSG00000( | gene | 0,82890017 | 0,00276934 | 0,02258287 | NDUFB2 | -0,2707297 |
| ENSG00000( | gene | 1,12114331 | 0,00278273 | 0,02267344 | ANAPC4 | 0,1649707 |
| ENSG00000( | gene | 1,06007611 | 0,00278586 | 0,02268028 | EIF4G3 | 0,08416785 |
| ENSG00000( | gene | 0,91961641 | 0,00278909 | 0,02268795 | IP6K2 | -0,1208959 |
| ENSG00000' | gene | 0,8946505 | 0,00279737 | 0,02273668 | PPP2R5D | -0,1606039 |
| ENSG00000' | gene | 0,91582281 | 0,00280501 | 0,02278004 | DENND5A | -0,1268596 |
| ENSG00000' | gene | 0,88166869 | 0,00281118 | 0,02281147 | PGK1 | -0,1816915 |
| ENSG00000' | gene | 0,91370031 | 0,00282305 | 0,02288913 | CNPY3 | -0,130207 |
| ENSG00000; | gene | 1,55065014 | 0,0028271 | 0,02290319 | AL133453.1 | 0,63287322 |
| ENSG00000' | gene | 0,84492147 | 0,00284109 | 0,02299776 | MAP2K1 | -0,2431108 |
| ENSG00000( | gene | 0,89079096 | 0,00285042 | 0,02304152 | TEAD2 | -0,1668412 |
| ENSG00000; | gene | 1,31321319 | 0,00285114 | 0,02304152 | ZNF32-AS1 | 0,39310115 |
| ENSG00000; | gene | 2,19071736 | 0,00285848 | 0,02308197 | SNORD58B | 1,13140337 |
| ENSG00000' | gene | 0,91562593 | 0,00286195 | 0,02308719 | C12orf10 | -0,1271698 |
| ENSG00000' | gene | 0,88184628 | 0,00286378 | 0,02308719 | LDHA | -0,1814009 |
| ENSG00000' | gene | 0,89118651 | 0,00287112 | 0,02311386 | ANXA5 | -0,1662007 |
| ENSG00000' | gene | 1,29473905 | 0,00287351 | 0,02311386 | CMBL | 0,37266136 |
| ENSG00000' | gene | 0,80955829 | 0,00287408 | 0,02311386 | ZNRF1 | -0,3047931 |
| ENSG00000' | gene | 0,85474648 | 0,00288016 | 0,0231311 | SIPA1L1 | -0,2264315 |
| ENSG00000' | gene | 0,80958543 | 0,00288089 | 0,0231311 | BFAR | -0,3047448 |
| ENSG00000; | gene | 1,22869952 | 0,00288745 | 0,02316505 | AC005841.1 | 0,29713214 |
| ENSG00000' | gene | 0,90414542 | 0,00289194 | 0,02318228 | SF3B4 | -0,1453733 |
| ENSG00000' | gene | 1,10762771 | 0,00289734 | 0,0231932 | SYNCRIP | 0,14747306 |
| ENSG00000; | gene | 2,40206587 | 0,00289873 | 0,0231932 | AC125257.2 | 1,26427571 |
| ENSG00000; | gene | 0,89451981 | 0,00290046 | 0,0231932 | E2F4 | -0,1608147 |
| ENSG00000' | gene | 1,10620866 | 0,00290472 | 0,0231932 | SSR3 | 0,14562354 |
| ENSG00000; | gene | 1,27886395 | 0,002905 | 0,0231932 | RPL21P75 | 0,35486279 |
| ENSG00000; | gene | 1,40889225 | 0,0029316 | 0,02338369 | AL606534.2 | 0,49456128 |
| ENSG00000; | gene | 1,42323621 | 0,00293357 | 0,02338369 | AP003392.3 | 0,50917512 |
| ENSG00000; | gene | 1,3305994 | 0,00294529 | 0,02345826 | AC108449.3 | 0,41207629 |
| ENSG00000( | gene | 0,85045501 | 0,00294807 | 0,02346153 | RHBDD2 | -0,2336932 |
| ENSG00000' | gene | 1,56158584 | 0,00295308 | 0,02346566 | TTK | 0,64301188 |
| ENSG00000' | gene | 0,85958388 | 0,00295365 | 0,02346566 | CBY1 | -0,2182897 |
| ENSG00000' | gene | 1,11388365 | 0,00295829 | 0,02346566 | ZCRB1 | 0,15559855 |
| ENSG00000' | gene | 0,93089557 | 0,00296807 | 0,02346566 | SNRNP70 | -0,1033088 |
| ENSG00000' | gene | 0,91019382 | 0,00296862 | 0,02346566 | MRPS5 | -0,1357543 |
| ENSG00000' | gene | 0,83098916 | 0,00296868 | 0,02346566 | AHSA1 | -0,2670984 |

| | | | | | | |
|---|---|---|---|---|---|---|
| ENSG00000 | gene | 0,9078469 | 0,00297109 | 0,02346566 | NOP56 | -0,1394791 |
| ENSG00000 | gene | 1,11252245 | 0,00297276 | 0,02346566 | TTC8 | 0,15383445 |
| ENSG00000 | gene | 1,08106792 | 0,00297285 | 0,02346566 | TPP2 | 0,11245716 |
| ENSG00000 | gene | 0,87500091 | 0,00297314 | 0,02346566 | GRHPR | -0,1926436 |
| ENSG00000 | gene | 0,92499214 | 0,00297462 | 0,02346566 | MAD2L2 | -0,112487 |
| ENSG00000 | gene | 1,11028706 | 0,00297926 | 0,02348361 | PEAK1 | 0,15093273 |
| ENSG00000 | gene | 1,16006494 | 0,00298621 | 0,02350929 | PJA2 | 0,21420557 |
| ENSG00000 | gene | 0,86553771 | 0,00298785 | 0,02350929 | EXOSC7 | -0,2083314 |
| ENSG00000 | gene | 1,44606168 | 0,00299159 | 0,02350929 | AC010618.3 | 0,53212909 |
| ENSG00000 | gene | 1,30506028 | 0,002992 | 0,02350929 | MICU3 | 0,38411644 |
| ENSG00000 | gene | 0,80091032 | 0,00302547 | 0,02375338 | POLR2I | -0,3202874 |
| ENSG00000 | gene | 0,86820597 | 0,0030379 | 0,02382621 | YBX3 | -0,2038908 |
| ENSG00000 | gene | 1,07654234 | 0,00303955 | 0,02382621 | PTPN21 | 0,10640506 |
| ENSG00000 | gene | 1,14725554 | 0,00304362 | 0,02383929 | PPIL4 | 0,19818678 |
| ENSG00000 | gene | 1,15510419 | 0,00304977 | 0,02386859 | SREK1 | 0,20802299 |
| ENSG00000 | gene | 0,81995047 | 0,00308306 | 0,0241101 | NEDD8 | -0,2863913 |
| ENSG00000 | gene | 0,87100285 | 0,00309339 | 0,02417182 | ZDHHC4 | -0,1992507 |
| ENSG00000 | gene | 1,40187988 | 0,00309943 | 0,02419993 | FERP1 | 0,48736274 |
| ENSG00000 | gene | 0,87415523 | 0,0031035 | 0,02421267 | SLC25A25 | -0,1940386 |
| ENSG00000 | gene | 1,3126748 | 0,00310981 | 0,02424285 | TREH | 0,39250955 |
| ENSG00000 | gene | 1,30132119 | 0,00311952 | 0,02429942 | MSH2 | 0,37997709 |
| ENSG00000 | gene | 1,36294928 | 0,00313169 | 0,0243726 | AC010335.1 | 0,44673187 |
| ENSG00000 | gene | 1,34018094 | 0,00313383 | 0,0243726 | AC022400.9 | 0,42242779 |
| ENSG00000 | gene | 0,91284066 | 0,00314791 | 0,02444539 | NUB1 | -0,131565 |
| ENSG00000 | gene | 0,88016655 | 0,00314812 | 0,02444539 | PDCD5 | -0,1841515 |
| ENSG00000 | gene | 1,3109561 | 0,0031622 | 0,02452833 | AC064836.4 | 0,39061937 |
| ENSG00000 | gene | 0,88265986 | 0,00316574 | 0,02452833 | ANAPC2 | -0,1800705 |
| ENSG00000 | gene | 0,93555121 | 0,00317 | 0,02452833 | ARAF | -0,0961115 |
| ENSG00000 | gene | 0,71513761 | 0,00317079 | 0,02452833 | U2AF1 | -0,4837072 |
| ENSG00000 | gene | 0,91179057 | 0,00317117 | 0,02452833 | AGPAT2 | -0,1332256 |
| ENSG00000 | gene | 0,86836765 | 0,00317523 | 0,02454063 | TOP1MT | -0,2036221 |
| ENSG00000 | gene | 1,54853088 | 0,00319257 | 0,02465539 | SNORA25 | 0,63090015 |
| ENSG00000 | gene | 1,76220626 | 0,00319694 | 0,02466213 | AC037198.2 | 0,8173828 |
| ENSG00000 | gene | 0,8516733 | 0,00319842 | 0,02466213 | STIP1 | -0,231628 |
| ENSG00000 | gene | 0,89925842 | 0,00320709 | 0,0246927 | NSDHL | -0,1531923 |
| ENSG00000 | gene | 0,84297856 | 0,00320736 | 0,0246927 | RPL15P3 | -0,2464321 |
| ENSG00000 | gene | 0,60518298 | 0,00321484 | 0,02473106 | GABARAPL | -0,7245567 |
| ENSG00000 | gene | 1,1154563 | 0,00322402 | 0,02478243 | RAD21 | 0,15763399 |
| ENSG00000 | gene | 1,19620086 | 0,00322659 | 0,024783 | NFYB | 0,25845966 |
| ENSG00000 | gene | 0,93344136 | 0,00324278 | 0,02488024 | CAP1 | -0,0993687 |
| ENSG00000 | gene | 0,76996644 | 0,00324427 | 0,02488024 | AC010343.1 | -0,3771325 |
| ENSG00000 | gene | 0,83430254 | 0,00324919 | 0,02489876 | NUDT14 | -0,2613575 |
| ENSG00000 | gene | 1,23584587 | 0,00326324 | 0,02498711 | UPF2 | 0,30549882 |
| ENSG00000 | gene | 1,34930305 | 0,00327257 | 0,02501135 | AP003692.1 | 0,43221441 |
| ENSG00000 | gene | 0,76157291 | 0,00327349 | 0,02501135 | RPS17 | -0,3929459 |
| ENSG00000 | gene | 0,89293094 | 0,00327397 | 0,02501135 | UBE2I | -0,1633795 |
| ENSG00000 | gene | 0,86244697 | 0,00328407 | 0,02506918 | RPL27 | -0,2134923 |
| ENSG00000 | gene | 0,68844904 | 0,00328935 | 0,02509012 | C11orf98 | -0,5385782 |
| ENSG00000 | gene | 0,86361838 | 0,00329192 | 0,02509042 | MT-ND6 | -0,2115341 |
| ENSG00000 | gene | 1,19690964 | 0,00330392 | 0,02514444 | MALSU1 | 0,25931424 |
| ENSG00000 | gene | 0,78476069 | 0,00330407 | 0,02514444 | AKR1B1 | -0,3496753 |
| ENSG00000 | gene | 0,90894782 | 0,00330816 | 0,02515619 | TARS2 | -0,1377306 |
| ENSG00000 | gene | 1,11700347 | 0,00332168 | 0,0252397 | GPSM2 | 0,15963367 |
| ENSG00000 | gene | 0,89771556 | 0,00333283 | 0,02530502 | RNF167 | -0,1556697 |
| ENSG00000 | gene | 0,8506779 | 0,00333721 | 0,02531887 | FOXK2 | -0,2333151 |
| ENSG00000 | gene | 0,82059822 | 0,00335449 | 0,02543052 | NPIPB5 | -0,2852521 |
| ENSG00000 | gene | 0,88013911 | 0,00337608 | 0,02556647 | ATF5 | -0,1841965 |

| | | | | | | |
|---|---|---|---|---|---|---|
| ENSG00002 | gene | 1,6601044 | 0,00337758 | 0,02556647 | U62317.2 | 0,73127397 |
| ENSG00002 | gene | 1,25301465 | 0,0033851 | 0,02560382 | AC087164.2 | 0,32540328 |
| ENSG00000 | gene | 0,89200046 | 0,00339266 | 0,0256415 | GABARAPL2 | -0,1648836 |
| ENSG00000 | gene | 0,79273185 | 0,0033959 | 0,02564641 | RPL18A | -0,3350951 |
| ENSG00000 | gene | 0,87230402 | 0,00339955 | 0,02565445 | IGSF8 | -0,1970971 |
| ENSG00000 | gene | 0,82268678 | 0,0034061 | 0,02568435 | MAZ | -0,2815848 |
| ENSG00000 | gene | 0,8537818 | 0,00342089 | 0,02577624 | TMEM120A | -0,2280607 |
| ENSG00000 | gene | 0,91154979 | 0,00342616 | 0,02579639 | PITPNA | -0,1336066 |
| ENSG00000 | gene | 0,89435345 | 0,00343597 | 0,02585062 | LSM7 | -0,161083 |
| ENSG00002 | gene | 1,3217286 | 0,00344124 | 0,02587064 | AC112497.2 | 0,40242597 |
| ENSG00000 | gene | 1,4466009 | 0,0034574 | 0,02597244 | NT5DC4 | 0,53266695 |
| ENSG00000 | gene | 0,83095605 | 0,00346172 | 0,02598112 | CERS2 | -0,2671559 |
| ENSG00000 | gene | 0,8493648 | 0,0034638 | 0,02598112 | DDA1 | -0,2355438 |
| ENSG00000 | gene | 0,88845778 | 0,00347664 | 0,02604746 | MTFR1L | -0,1706249 |
| ENSG00002 | gene | 1,17631136 | 0,0034779 | 0,02604746 | C22orf39 | 0,23426998 |
| ENSG00000 | gene | 0,93262702 | 0,00348124 | 0,02605023 | DNAJC8 | -0,1006279 |
| ENSG00000 | gene | 1,20807198 | 0,00348355 | 0,02605023 | MBNL2 | 0,27270642 |
| ENSG00002 | gene | 0,8646274 | 0,00348615 | 0,02605023 | AC134312.5 | -0,2098495 |
| ENSG00000 | gene | 0,92302007 | 0,00349205 | 0,02606 | TBC1D14 | -0,1155661 |
| ENSG00000 | gene | 0,90881297 | 0,00349271 | 0,02606 | GALNS | -0,1379447 |
| ENSG00000 | gene | 0,84172471 | 0,00349755 | 0,02607644 | CDK2AP1 | -0,2485796 |
| ENSG00000 | gene | 1,20448631 | 0,00350114 | 0,02608362 | BLZF1 | 0,26841799 |
| ENSG00000 | gene | 1,16232225 | 0,0035155 | 0,02617097 | EMC2 | 0,2170101 |
| ENSG00000 | gene | 1,11477627 | 0,00352108 | 0,0261928 | TASOR2 | 0,1567542 |
| ENSG00000 | gene | 1,33889838 | 0,00352438 | 0,02619774 | FAM214A | 0,42104647 |
| ENSG00000 | gene | 0,87603969 | 0,00353155 | 0,02623133 | PSMA6 | -0,1909319 |
| ENSG00000 | gene | 0,80953897 | 0,00353779 | 0,02625805 | WDR45 | -0,3048276 |
| ENSG00002 | gene | 1,40246991 | 0,00355743 | 0,02638402 | MIR6080 | 0,48796982 |
| ENSG00002 | gene | 1,99520189 | 0,00356683 | 0,02643402 | AC007378.1 | 0,99653474 |
| ENSG00000 | gene | 0,88048297 | 0,00359158 | 0,02659754 | KLC2 | -0,183633 |
| ENSG00000 | gene | 0,59182595 | 0,00359943 | 0,02663575 | CBR4 | -0,7567551 |
| ENSG00002 | gene | 0,59100819 | 0,00360401 | 0,02664975 | AL035699.1 | -0,75875 |
| ENSG00002 | gene | 1,52431728 | 0,00361566 | 0,02670314 | AC007193.3 | 0,60816323 |
| ENSG00000 | gene | 0,68337909 | 0,00361661 | 0,02670314 | XAF1 | -0,549242 |
| ENSG00002 | gene | 1,36872028 | 0,00362772 | 0,02675706 | AC005288.1 | 0,45282764 |
| ENSG00000 | gene | 1,26662996 | 0,00362931 | 0,02675706 | CFAP126 | 0,34099511 |
| ENSG00000 | gene | 1,33126141 | 0,00363329 | 0,02676646 | DLGAP5 | 0,41279389 |
| ENSG00002 | gene | 0,83218712 | 0,0036479 | 0,02685415 | QTRT1 | -0,2650201 |
| ENSG00000 | gene | 1,26999863 | 0,00365581 | 0,02687383 | CALD1 | 0,34482695 |
| ENSG00000 | gene | 1,0984903 | 0,00365803 | 0,02687383 | GNAI3 | 0,13552213 |
| ENSG00000 | gene | 1,07628676 | 0,00366019 | 0,02687383 | MAVS | 0,10606251 |
| ENSG00000 | gene | 1,34238967 | 0,00366349 | 0,02687383 | NSD3 | 0,42480352 |
| ENSG00000 | gene | 0,74340041 | 0,00366412 | 0,02687383 | HELZ2 | -0,4277886 |
| ENSG00000 | gene | 0,70225047 | 0,00367292 | 0,02691848 | RPL12P4 | -0,5099424 |
| ENSG00000 | gene | 1,06409212 | 0,00367705 | 0,02692367 | SEC24D | 0,08962305 |
| ENSG00000 | gene | 0,87028287 | 0,00368382 | 0,02692367 | SPATS2 | -0,2004437 |
| ENSG00000 | gene | 1,07618042 | 0,0036839 | 0,02692367 | AZIN1 | 0,10591996 |
| ENSG00000 | gene | 1,08255823 | 0,00368449 | 0,02692367 | CPSF3 | 0,11444463 |
| ENSG00002 | gene | 0,65742319 | 0,00369589 | 0,02698706 | HNRNPA1P4 | -0,6051057 |
| ENSG00000 | gene | 0,79958664 | 0,0036988 | 0,02698843 | ANKZF1 | -0,3226737 |
| ENSG00000 | gene | 1,15753545 | 0,00371885 | 0,02708527 | SP3 | 0,21105638 |
| ENSG00000 | gene | 0,83341019 | 0,00371911 | 0,02708527 | TPI1 | -0,2629014 |
| ENSG00000 | gene | 0,83180962 | 0,00372026 | 0,02708527 | USP10 | -0,2656747 |
| ENSG00002 | gene | 1,29076054 | 0,00373552 | 0,02717636 | AC009019.1 | 0,36822138 |
| ENSG00000 | gene | 1,11710448 | 0,00374225 | 0,02720537 | XPA | 0,15976413 |
| ENSG00000 | gene | 0,85253745 | 0,00375283 | 0,0272623 | RPL10 | -0,2301649 |
| ENSG00000 | gene | 1,10334999 | 0,0037595 | 0,02729081 | BIRC6 | 0,1418905 |

| | | | | | | |
|---|---|---|---|---|---|---|
| ENSG00000' | gene | 0,86221174 | 0,00376939 | 0,02734167 | MTSS2 | -0,2138859 |
| ENSG00000' | gene | 0,80546118 | 0,00377416 | 0,02734167 | FBXL12 | -0,312113 |
| ENSG00000' | gene | 1,12433545 | 0,00377757 | 0,02734167 | TIGD2 | 0,16907254 |
| ENSG00000' | gene | 0,91482268 | 0,0037782 | 0,02734167 | ARHGEF11 | -0,128436 |
| ENSG00000( | gene | 0,89442466 | 0,0037803 | 0,02734167 | PSMC4 | -0,1609681 |
| ENSG00000' | gene | 0,87220334 | 0,0037892 | 0,02737091 | PEMT | -0,1972636 |
| ENSG00000' | gene | 0,93208935 | 0,00378986 | 0,02737091 | EFTUD2 | -0,1014598 |
| ENSG00002 | gene | 2,40446073 | 0,00379298 | 0,02737353 | AC022966.2 | 1,26571336 |
| ENSG00000' | gene | 0,67907483 | 0,00379597 | 0,02737516 | USP20 | -0,5583575 |
| ENSG00000' | gene | 1,17391197 | 0,00379982 | 0,02738297 | ACAP2 | 0,23132422 |
| ENSG00002 | gene | 1,292549 | 0,00380852 | 0,02742576 | AC087284.2 | 0,37021897 |
| ENSG00002 | gene | 1,74948173 | 0,00382348 | 0,02750011 | AL035252.5 | 0,8069276 |
| ENSG00000( | gene | 0,81096125 | 0,0038257 | 0,02750011 | RPL31 | -0,3022951 |
| ENSG00000' | gene | 1,09443191 | 0,00382716 | 0,02750011 | PTPN14 | 0,1301822 |
| ENSG00000' | gene | 0,8270476 | 0,00384446 | 0,02760433 | SPINDOC | -0,2739577 |
| ENSG00002 | gene | 0,51757756 | 0,00384724 | 0,02760433 | AL161909.1 | -0,950153 |
| ENSG00000' | gene | 0,86463364 | 0,0038625 | 0,02769383 | ZFP36L1 | -0,2098391 |
| ENSG00002 | gene | 1,71231299 | 0,00387837 | 0,02776473 | AC005696.2 | 0,77594643 |
| ENSG00000' | gene | 0,85762706 | 0,00387949 | 0,02776473 | PLEKHJ1 | -0,2215777 |
| ENSG00002 | gene | 0,50017166 | 0,00388081 | 0,02776473 | AC015813.2 | -0,9995048 |
| ENSG00000( | gene | 0,93203849 | 0,00388359 | 0,02776473 | PHF23 | -0,1015386 |
| ENSG00002 | gene | 1,60046375 | 0,00389356 | 0,02781594 | PAPPA-AS1 | 0,67849 |
| ENSG00000' | gene | 1,17698796 | 0,00389955 | 0,02783868 | RBBP8 | 0,23509956 |
| ENSG00002 | gene | 1,20347569 | 0,00391415 | 0,02792279 | AC002524.1 | 0,267207 |
| ENSG00000' | gene | 0,78360296 | 0,00392099 | 0,02795152 | CENPX | -0,3518052 |
| ENSG00000' | gene | 1,24923805 | 0,00393987 | 0,02806589 | SLC19A1 | 0,32104842 |
| ENSG00000' | gene | 0,92848012 | 0,0039445 | 0,0280758 | LSS | -0,1070571 |
| ENSG00000' | gene | 0,92389856 | 0,00394692 | 0,0280758 | POM121 | -0,1141936 |
| ENSG00002 | gene | 1,60560871 | 0,00396899 | 0,02821255 | AC005104.1 | 0,68312035 |
| ENSG00000( | gene | 1,16304476 | 0,00397543 | 0,02823806 | BCAT1 | 0,21790662 |
| ENSG00000' | gene | 0,89567853 | 0,00398226 | 0,02826632 | CASP7 | -0,1589471 |
| ENSG00000' | gene | 0,82738307 | 0,0039876 | 0,02828397 | GLMP | -0,2733727 |
| ENSG00000' | gene | 0,88123987 | 0,00400098 | 0,02835866 | FKBP8 | -0,1823933 |
| ENSG00000' | gene | 1,06575372 | 0,00401461 | 0,02842118 | TOR1AIP1 | 0,09187409 |
| ENSG00002 | gene | 1,32327071 | 0,00402131 | 0,02842118 | AC016825.1 | 0,40410824 |
| ENSG00000' | gene | 0,8674905 | 0,00402314 | 0,02842118 | PLD3 | -0,2050801 |
| ENSG00000( | gene | 0,90599904 | 0,00402489 | 0,02842118 | APLP2 | -0,1424186 |
| ENSG00000' | gene | 0,7330094 | 0,00402807 | 0,02842118 | AL450998.1 | -0,4480964 |
| ENSG00002 | gene | 1,26173741 | 0,00402935 | 0,02842118 | AC020763.4 | 0,33541169 |
| ENSG00000' | gene | 0,82317223 | 0,00402987 | 0,02842118 | COX7A2 | -0,2807338 |
| ENSG00000' | gene | 0,83921933 | 0,00404398 | 0,02850046 | NOP53 | -0,2528802 |
| ENSG00000' | gene | 1,20369029 | 0,00405485 | 0,02855671 | PRIM1 | 0,26746423 |
| ENSG00000' | gene | 0,84756552 | 0,0040637 | 0,02859879 | NDUFB8 | -0,2386032 |
| ENSG00002 | gene | 2,07725071 | 0,00408491 | 0,02872764 | AL136988.1 | 1,05467535 |
| ENSG00000' | gene | 0,83469 | 0,00410386 | 0,02883171 | KIAA1191 | -0,2606876 |
| ENSG00000( | gene | 0,91040078 | 0,00410552 | 0,02883171 | SARS1 | -0,1354263 |
| ENSG00002 | gene | 1,34197817 | 0,00411629 | 0,02888686 | AC012157.2 | 0,42436121 |
| ENSG00002 | gene | 1,32483214 | 0,0041244 | 0,0289233 | AC093110.1 | 0,40580958 |
| ENSG00000' | gene | 0,7948069 | 0,00414871 | 0,0290732 | CAMTA1 | -0,3313237 |
| ENSG00000' | gene | 1,09474593 | 0,00415265 | 0,02908028 | NUP88 | 0,13059609 |
| ENSG00000' | gene | 0,67085905 | 0,00416285 | 0,02910979 | RNF216P1 | -0,5759184 |
| ENSG00000' | gene | 1,11770925 | 0,00416381 | 0,02910979 | PPP3CA | 0,16054495 |
| ENSG00000' | gene | 0,85657277 | 0,00416781 | 0,02910979 | SREBF2 | -0,2233523 |
| ENSG00000( | gene | 0,84892496 | 0,00416973 | 0,02910979 | TMED2 | -0,2362911 |
| ENSG00000' | gene | 1,159917 | 0,00417154 | 0,02910979 | FAM98B | 0,21402157 |
| ENSG00000' | gene | 0,86269775 | 0,00418591 | 0,0291895 | ITPA | -0,2130729 |
| ENSG00000' | gene | 1,57842712 | 0,00418887 | 0,0291896 | ZNF75A | 0,65848765 |

| | | | | | | |
|---|---|---|---|---|---|---|
| ENSG00000 | gene | 0,97069128 | 0,00419874 | 0,02922589 | PTTG1IP | -0,0429156 |
| ENSG00000 | gene | 1,38580585 | 0,00420277 | 0,02922589 | AC091180.4 | 0,47072515 |
| ENSG00000 | gene | 1,30437645 | 0,00420362 | 0,02922589 | KIF11 | 0,3833603 |
| ENSG00000 | gene | 1,21904651 | 0,00420736 | 0,02922589 | RAB11FIP1F | 0,28575316 |
| ENSG00000 | gene | 0,76848776 | 0,00421007 | 0,02922589 | AKR1C1 | -0,3799058 |
| ENSG00000 | gene | 0,91175782 | 0,00421176 | 0,02922589 | EDC3 | -0,1332774 |
| ENSG00000 | gene | 0,92972522 | 0,00422423 | 0,02926626 | PLOD1 | -0,1051237 |
| ENSG00000 | gene | 1,32981866 | 0,0042256 | 0,02926626 | RFC3 | 0,41122952 |
| ENSG00000 | gene | 0,82893178 | 0,0042295 | 0,02926626 | MYADM | -0,2706747 |
| ENSG00000 | gene | 1,39866011 | 0,00423132 | 0,02926626 | AC004974.1 | 0,48404541 |
| ENSG00000 | gene | 1,32066372 | 0,00423233 | 0,02926626 | KF459542.1 | 0,40126316 |
| ENSG00000 | gene | 1,33439139 | 0,00423867 | 0,02928611 | ZMYM4-AS1 | 0,41618188 |
| ENSG00000 | gene | 0,85288099 | 0,00424272 | 0,02928611 | CLP1 | -0,2295836 |
| ENSG00000 | gene | 1,33935101 | 0,00424406 | 0,02928611 | AC087190.1 | 0,42153411 |
| ENSG00000 | gene | 0,9256802 | 0,00427822 | 0,0294828 | CMTR1 | -0,1114142 |
| ENSG00000 | gene | 0,72793113 | 0,00427851 | 0,0294828 | DGKZ | -0,4581261 |
| ENSG00000 | gene | 0,83773252 | 0,00428156 | 0,0294833 | ST7 | -0,2554384 |
| ENSG00000 | gene | 0,92494723 | 0,00428574 | 0,02949162 | KCTD17 | -0,112557 |
| ENSG00000 | gene | 0,6987013 | 0,00432913 | 0,02975279 | PARP9 | -0,5172523 |
| ENSG00000 | gene | 0,8663731 | 0,0043297 | 0,02975279 | TPST1 | -0,2069397 |
| ENSG00000 | gene | 1,46339631 | 0,00433656 | 0,02977933 | SNORA63B | 0,54932053 |
| ENSG00000 | gene | 1,85681918 | 0,00434214 | 0,02979697 | AP002770.1 | 0,89283333 |
| ENSG00000 | gene | 1,78452199 | 0,00435677 | 0,02986476 | SNORA67 | 0,83553768 |
| ENSG00000 | gene | 0,75792628 | 0,00435804 | 0,02986476 | PEA15 | -0,3998706 |
| ENSG00000 | gene | 1,96684582 | 0,00436186 | 0,0298703 | AC006001.4 | 0,97588387 |
| ENSG00000 | gene | 1,13020393 | 0,00438272 | 0,02999243 | TRMT10C | 0,17658311 |
| ENSG00000 | gene | 1,20962166 | 0,00438937 | 0,03001724 | RPL7P9 | 0,27455588 |
| ENSG00000 | gene | 0,90020658 | 0,00440637 | 0,03009535 | HNRNPH1 | -0,151672 |
| ENSG00000 | gene | 0,76455556 | 0,00440899 | 0,03009535 | PTGES | -0,3873067 |
| ENSG00000 | gene | 0,93074115 | 0,00441513 | 0,03009535 | TMEM183A | -0,1035481 |
| ENSG00000 | gene | 1,26772122 | 0,0044194 | 0,03009535 | PHF3 | 0,34223753 |
| ENSG00000 | gene | 0,79902951 | 0,00441964 | 0,03009535 | TP73-AS1 | -0,3236793 |
| ENSG00000 | gene | 1,20680775 | 0,00441972 | 0,03009535 | AC105020.2 | 0,27119587 |
| ENSG00000 | gene | 1,12679269 | 0,00442204 | 0,03009535 | PPP1R12A | 0,17222211 |
| ENSG00000 | gene | 1,19599645 | 0,00442815 | 0,03010691 | PTGER2 | 0,2582131 |
| ENSG00000 | gene | 1,21292064 | 0,00442981 | 0,03010691 | TBL1XR1 | 0,27848516 |
| ENSG00000 | gene | 0,93062633 | 0,00443345 | 0,03011103 | CLPTM1L | -0,1037261 |
| ENSG00000 | gene | 0,79009515 | 0,00443767 | 0,03011248 | RACK1 | -0,3399017 |
| ENSG00000 | gene | 1,51963803 | 0,00443974 | 0,03011248 | NRF1 | 0,60372773 |
| ENSG00000 | gene | 0,84345906 | 0,00444754 | 0,03014479 | SLC25A37 | -0,24561 |
| ENSG00000 | gene | 1,44073071 | 0,00445218 | 0,03015564 | DIO2 | 0,5268007 |
| ENSG00000 | gene | 1,35828389 | 0,00446634 | 0,03019577 | AC107072.2 | 0,44178505 |
| ENSG00000 | gene | 1,51288786 | 0,00446792 | 0,03019577 | AL137060.1 | 0,59730505 |
| ENSG00000 | gene | 0,81009979 | 0,00446954 | 0,03019577 | XBP1 | -0,3038285 |
| ENSG00000 | gene | 1,32389021 | 0,00447029 | 0,03019577 | AL929472.2 | 0,40478349 |
| ENSG00000 | gene | 1,30640406 | 0,00447499 | 0,03020692 | AC010761.1 | 0,38560118 |
| ENSG00000 | gene | 0,72506069 | 0,0044898 | 0,03026686 | AL158847.1 | -0,4638263 |
| ENSG00000 | gene | 1,29162177 | 0,00448997 | 0,03026686 | ANKRD17 | 0,36918366 |
| ENSG00000 | gene | 1,34503422 | 0,00449382 | 0,03027221 | AC010776.2 | 0,42764288 |
| ENSG00000 | gene | 0,84652341 | 0,00449976 | 0,03027478 | UCHL1 | -0,2403781 |
| ENSG00000 | gene | 0,78279714 | 0,00450545 | 0,03027478 | GUK1 | -0,3532896 |
| ENSG00000 | gene | 0,85109987 | 0,00450616 | 0,03027478 | TST | -0,2325997 |
| ENSG00000 | gene | 1,39780803 | 0,00450641 | 0,03027478 | P4HA2-AS1 | 0,48316624 |
| ENSG00000 | gene | 0,92418049 | 0,00453272 | 0,0304177 | TRIP10 | -0,1137535 |
| ENSG00000 | gene | 0,7894321 | 0,00453382 | 0,0304177 | TRIM5 | -0,3411129 |
| ENSG00000 | gene | 1,27286051 | 0,00453758 | 0,03042233 | OPN1SW | 0,34807433 |
| ENSG00000 | gene | 1,2114886 | 0,00454467 | 0,03044931 | C1orf56 | 0,27678083 |

| | | | | | | |
|---|---|---|---|---|---|---|
| ENSG00000( | gene | 1,08540746 | 0,00454831 | 0,03045093 | UBR2 | 0,11823673 |
| ENSG00000' | gene | 0,8456645 | 0,00455106 | 0,03045093 | TBCA | -0,2418427 |
| ENSG00000' | gene | 0,90657281 | 0,00455987 | 0,03048467 | PINK1 | -0,1415052 |
| ENSG00000' | gene | 0,71895161 | 0,00456225 | 0,03048467 | TPT1 | -0,4760334 |
| ENSG00000: | gene | 1,55032886 | 0,00458663 | 0,03062693 | ATP2A1-AS' | 0,63257428 |
| ENSG00000: | gene | 1,43713111 | 0,00459274 | 0,03064712 | AC005606.1 | 0,52319168 |
| ENSG00000( | gene | 0,88009184 | 0,00460509 | 0,03070887 | PAFAH1B3 | -0,184274 |
| ENSG00000' | gene | 0,72396054 | 0,00461 | 0,03072097 | NRSN2 | -0,466017 |
| ENSG00000' | gene | 0,91048535 | 0,00462255 | 0,03078387 | ENTPD6 | -0,1352923 |
| ENSG00000' | gene | 1,08294703 | 0,00463745 | 0,03086236 | METAP2 | 0,11496268 |
| ENSG00000' | gene | 0,81358064 | 0,00464759 | 0,03090915 | COX6B1 | -0,2976428 |
| ENSG00000' | gene | 0,86131016 | 0,00466882 | 0,03102948 | RFNG | -0,2153952 |
| ENSG00000' | gene | 0,90487096 | 0,00467673 | 0,03106128 | SF1 | -0,144216 |
| ENSG00000: | gene | 1,27277328 | 0,0046872 | 0,03110273 | AC099508.1 | 0,34797546 |
| ENSG00000: | gene | 1,98678823 | 0,00469321 | 0,03110273 | MIR22 | 0,99043811 |
| ENSG00000' | gene | 1,1283357 | 0,0046933 | 0,03110273 | ARL14EP | 0,17419636 |
| ENSG00000' | gene | 1,09570011 | 0,00469598 | 0,03110273 | TMEM131L | 0,131853 |
| ENSG00000' | gene | 0,84810133 | 0,00469866 | 0,03110273 | PRRC2B | -0,2376914 |
| ENSG00000' | gene | 1,16085065 | 0,00471042 | 0,0311598 | CUL4B | 0,21518238 |
| ENSG00000: | gene | 1,37078356 | 0,00471405 | 0,03116298 | SNORD17 | 0,45500079 |
| ENSG00000: | gene | 1,42218172 | 0,00475233 | 0,03139368 | AC097451.1 | 0,50810582 |
| ENSG00000' | gene | 1,26663457 | 0,00475527 | 0,03139368 | SYTL4 | 0,34100037 |
| ENSG00000' | gene | 0,80329574 | 0,00476466 | 0,0314347 | NDUFA2 | -0,3159969 |
| ENSG00000' | gene | 1,25052865 | 0,00479955 | 0,03164386 | NCAPG | 0,32253811 |
| ENSG00000' | gene | 0,83286944 | 0,00481085 | 0,03167539 | ARFGAP1 | -0,2638377 |
| ENSG00000' | gene | 0,77582756 | 0,0048126 | 0,03167539 | C15orf40 | -0,3661921 |
| ENSG00000' | gene | 0,87362246 | 0,00481392 | 0,03167539 | TMEM115 | -0,1949181 |
| ENSG00000' | gene | 0,72911936 | 0,004827 | 0,03174039 | IRF1-AS1 | -0,4557731 |
| ENSG00000: | gene | 1,22560337 | 0,00483428 | 0,03175872 | AC002094.2 | 0,29349217 |
| ENSG00000' | gene | 0,82303121 | 0,00483741 | 0,03175872 | AVPI1 | -0,280981 |
| ENSG00000' | gene | 0,85554176 | 0,00483939 | 0,03175872 | BTBD6 | -0,2250898 |
| ENSG00000: | gene | 1,32317223 | 0,00484643 | 0,03178388 | AL606534.1 | 0,40400086 |
| ENSG00000( | gene | 0,8840473 | 0,00486101 | 0,03185844 | PTPN23 | -0,1778045 |
| ENSG00000: | gene | 1,43805252 | 0,00488855 | 0,03201773 | AC124283.1 | 0,52411637 |
| ENSG00000' | gene | 0,86203159 | 0,00489657 | 0,03203836 | SNAPC2 | -0,2141874 |
| ENSG00000' | gene | 0,84877876 | 0,00490064 | 0,03203836 | NR1H2 | -0,2365395 |
| ENSG00000' | gene | 0,79073125 | 0,00490139 | 0,03203836 | NDUFB9 | -0,3387407 |
| ENSG00000: | gene | 1,30392226 | 0,00491637 | 0,032113 | AC114947.2 | 0,38285786 |
| ENSG00000' | gene | 0,90798645 | 0,00491929 | 0,032113 | POLDIP3 | -0,1392573 |
| ENSG00000( | gene | 0,9550589 | 0,00493111 | 0,03215133 | EIF2AK1 | -0,0663384 |
| ENSG00000' | gene | 1,09790999 | 0,00493164 | 0,03215133 | IQGAP1 | 0,13475979 |
| ENSG00000' | gene | 0,83237261 | 0,004942 | 0,03218334 | PYCR1 | -0,2646986 |
| ENSG00000' | gene | 1,19492357 | 0,00494374 | 0,03218334 | SEPTIN11 | 0,25691835 |
| ENSG00000( | gene | 0,79148696 | 0,00494629 | 0,03218334 | CDC34 | -0,3373625 |
| ENSG00000: | gene | 0,75025667 | 0,00497097 | 0,03232271 | HMGA1P2 | -0,4145439 |
| ENSG00000' | gene | 0,83200954 | 0,00497975 | 0,03235861 | SMIM29 | -0,265328 |
| ENSG00000: | gene | 0,90343888 | 0,00500213 | 0,03248275 | AC067852.2 | -0,1465011 |
| ENSG00000: | gene | 1,31088972 | 0,00502035 | 0,03257971 | MIR3074 | 0,39054632 |
| ENSG00000: | gene | 0,81939143 | 0,00503296 | 0,03263549 | SNHG5 | -0,2873753 |
| ENSG00000' | gene | 0,88944054 | 0,00503731 | 0,03263549 | RPL3 | -0,1690299 |
| ENSG00000: | gene | 1,33442619 | 0,00503882 | 0,03263549 | AP000926.2 | 0,41621951 |
| ENSG00000' | gene | 1,3323853 | 0,00505041 | 0,0326892 | TOP2A | 0,41401135 |
| ENSG00000' | gene | 1,13217895 | 0,00506355 | 0,03275289 | ADAMTS12 | 0,179102 |
| ENSG00000( | gene | 0,82309981 | 0,00507289 | 0,03279188 | AIFM2 | -0,2808607 |
| ENSG00000( | gene | 1,34510118 | 0,00508388 | 0,0328407 | ANLN | 0,4277147 |
| ENSG00000: | gene | 1,19211966 | 0,00508706 | 0,0328407 | BMPR2 | 0,25352906 |
| ENSG00000' | gene | 1,07384395 | 0,00510124 | 0,03291082 | HBS1L | 0,10278436 |

| | | | | | | |
|---|---|---|---|---|---|---|
| ENSG00000' | gene | 0,89405007 | 0,00510818 | 0,03293413 | APP | -0,1615725 |
| ENSG00002 | gene | 0,63454681 | 0,00511972 | 0,03298711 | AL135905.2 | -0,6562015 |
| ENSG00000' | gene | 0,88722892 | 0,00514031 | 0,0330822 | MICOS10 | -0,1726217 |
| ENSG00000( | gene | 1,16025819 | 0,0051442 | 0,0330822 | LMAN1 | 0,21444588 |
| ENSG00000' | gene | 0,7239899 | 0,0051472 | 0,0330822 | SDR39U1 | -0,4659585 |
| ENSG00000' | gene | 0,8509028 | 0,00515064 | 0,0330822 | NXF1 | -0,2329338 |
| ENSG00002 | gene | 1,85645378 | 0,005153 | 0,0330822 | AL669831.3 | 0,89254939 |
| ENSG00002 | gene | 1,36370964 | 0,0051545 | 0,0330822 | AC127496.6 | 0,4475365 |
| ENSG00000' | gene | 1,08863952 | 0,00517567 | 0,03317942 | MOB3A | 0,12252631 |
| ENSG00002 | gene | 1,50238841 | 0,00517634 | 0,03317942 | AC091053.1 | 0,58725784 |
| ENSG00002 | gene | 1,26126378 | 0,00519025 | 0,03324713 | AC009065.8 | 0,33487003 |
| ENSG00002 | gene | 1,28630609 | 0,00519986 | 0,03327557 | TAB3-AS2 | 0,36323398 |
| ENSG00000' | gene | 0,92501026 | 0,00520141 | 0,03327557 | NRBP1 | -0,1124587 |
| ENSG00002 | gene | 1,32085649 | 0,00521213 | 0,03328736 | AC120114.2 | 0,40147373 |
| ENSG00000' | gene | 1,10637832 | 0,0052136 | 0,03328736 | VPS54 | 0,14584479 |
| ENSG00002 | gene | 1,32543293 | 0,00521448 | 0,03328736 | FMN1 | 0,40646367 |
| ENSG00000' | gene | 0,91847819 | 0,00521668 | 0,03328736 | RRP7A | -0,1226826 |
| ENSG00000( | gene | 0,89182943 | 0,0052229 | 0,03330564 | SDF4 | -0,1651603 |
| ENSG00000' | gene | 0,90757621 | 0,00523535 | 0,03334964 | POLR2D | -0,1399093 |
| ENSG00000' | gene | 0,87338159 | 0,00523653 | 0,03334964 | TEX264 | -0,195316 |
| ENSG00000' | gene | 0,76396836 | 0,00527653 | 0,03358283 | FAM219A | -0,3884152 |
| ENSG00000' | gene | 0,86450893 | 0,00528182 | 0,03359498 | SHC1 | -0,2100472 |
| ENSG00000' | gene | 0,86649199 | 0,0052871 | 0,03360699 | MRPS11 | -0,2067417 |
| ENSG00000' | gene | 0,81467459 | 0,00529071 | 0,03360837 | NECAP2 | -0,2957042 |
| ENSG00002 | gene | 1,50960432 | 0,00529766 | 0,03363097 | AC006547.2 | 0,59417045 |
| ENSG00000( | gene | 0,89705681 | 0,00530583 | 0,03366131 | PDRG1 | -0,1567287 |
| ENSG00000' | gene | 0,8240723 | 0,00532618 | 0,03374813 | EGR1 | -0,2791572 |
| ENSG00000( | gene | 0,83685846 | 0,00532789 | 0,03374813 | CIRBP | -0,2569445 |
| ENSG00000' | gene | 1,08771031 | 0,00533184 | 0,03374813 | KMT5B | 0,12129437 |
| ENSG00000' | gene | 0,91286656 | 0,00533822 | 0,03374813 | CDK7 | -0,1315241 |
| ENSG00002 | gene | 1,37584258 | 0,00533844 | 0,03374813 | AP002490.1 | 0,46031541 |
| ENSG00002 | gene | 1,11489237 | 0,00533994 | 0,03374813 | SACM1L | 0,15690445 |
| ENSG00000' | gene | 0,92312662 | 0,00536246 | 0,03386887 | MORF4L2 | -0,1153995 |
| ENSG00000' | gene | 0,87607136 | 0,00537159 | 0,03390494 | SOD1 | -0,1908797 |
| ENSG00000( | gene | 0,84743795 | 0,00537987 | 0,03393564 | SP100 | -0,2388204 |
| ENSG00002 | gene | 1,27880232 | 0,00539959 | 0,03402173 | SZT2-AS1 | 0,35479326 |
| ENSG00000' | gene | 0,91049277 | 0,00540038 | 0,03402173 | MRTFA | -0,1352805 |
| ENSG00000' | gene | 0,8300703 | 0,00541054 | 0,03405396 | CHKA | -0,2686946 |
| ENSG00000' | gene | 0,75463385 | 0,00541237 | 0,03405396 | NPAS2 | -0,4061513 |
| ENSG00002 | gene | 1,33208731 | 0,00541869 | 0,03407209 | TRBV12-4 | 0,41368864 |
| ENSG00000' | gene | 0,90334784 | 0,0054291 | 0,03410034 | BCL2L12 | -0,1466465 |
| ENSG00000' | gene | 0,91558514 | 0,00543006 | 0,03410034 | POLR2G | -0,127234 |
| ENSG00000' | gene | 0,91819682 | 0,00544047 | 0,03414411 | QARS1 | -0,1231247 |
| ENSG00002 | gene | 0,60283775 | 0,00545228 | 0,0341789 | AL162151.2 | -0,7301583 |
| ENSG00000' | gene | 0,84335138 | 0,00545395 | 0,0341789 | FAM89B | -0,2457942 |
| ENSG00000( | gene | 0,90006755 | 0,00545635 | 0,0341789 | FXYD5 | -0,1518948 |
| ENSG00000' | gene | 1,22577401 | 0,00546854 | 0,03423362 | CKAP2L | 0,29369302 |
| ENSG00000( | gene | 0,89097346 | 0,00547287 | 0,03423912 | RAB7A | -0,1665456 |
| ENSG00002 | gene | 0,80355617 | 0,00550659 | 0,03440134 | AC016876.3 | -0,3155292 |
| ENSG00002 | gene | 1,26805383 | 0,00550949 | 0,03440134 | AL354920.1 | 0,34261599 |
| ENSG00000' | gene | 1,20840359 | 0,00551077 | 0,03440134 | PTAR1 | 0,27310237 |
| ENSG00002 | gene | 1,35545232 | 0,00551268 | 0,03440134 | AC027277.1 | 0,43877436 |
| ENSG00002 | gene | 0,7451641 | 0,00552755 | 0,03447246 | AC069288.1 | -0,4243699 |
| ENSG00000' | gene | 0,89143889 | 0,00553196 | 0,03447828 | PSMB6 | -0,1657922 |
| ENSG00000' | gene | 0,84737402 | 0,00553991 | 0,03450611 | FBRS | -0,2389292 |
| ENSG00000' | gene | 1,08577402 | 0,00554363 | 0,03450761 | LTV1 | 0,11872387 |
| ENSG00002 | gene | 1,44893819 | 0,00554759 | 0,03451062 | AC137590.1 | 0,53499605 |

| | | | | | | |
|---|---|---|---|---|---|---|
| ENSG00002 | gene | 1,41519974 | 0,0055559 | 0,03454062 | AC027644.3 | 0,50100569 |
| ENSG00000 | gene | 0,80658625 | 0,0056012 | 0,03480042 | SORBS3 | -0,3100993 |
| ENSG00000 | gene | 0,8739599 | 0,00560676 | 0,03481174 | NT5C3B | -0,194361 |
| ENSG00000 | gene | 0,92578403 | 0,00561009 | 0,03481174 | MT-CYB | -0,1112524 |
| ENSG00002 | gene | 1,4478141 | 0,00561355 | 0,03481174 | AL035661.2 | 0,53387638 |
| ENSG00000 | gene | 0,69256118 | 0,00562261 | 0,03484613 | MOB2 | -0,5299866 |
| ENSG00000 | gene | 1,07749863 | 0,00562942 | 0,03486655 | TRAK2 | 0,10768604 |
| ENSG000000 | gene | 0,74013866 | 0,00564306 | 0,03492923 | BTN3A1 | -0,4341325 |
| ENSG00000 | gene | 0,94396506 | 0,0056499 | 0,03494969 | SFXN3 | -0,0831946 |
| ENSG00002 | gene | 1,35398477 | 0,00565635 | 0,03496783 | AC104758.1 | 0,43721151 |
| ENSG00002 | gene | 0,84629517 | 0,00568054 | 0,03509545 | NDUFS3 | -0,2407672 |
| ENSG00002 | gene | 1,33055454 | 0,00571275 | 0,03525746 | AC036103.1 | 0,41202765 |
| ENSG00002 | gene | 1,452472 | 0,00571593 | 0,03525746 | ZNF271P | 0,53851035 |
| ENSG00000 | gene | 0,82878496 | 0,00571743 | 0,03525746 | OAZ1 | -0,2709303 |
| ENSG00002 | gene | 1,19735056 | 0,00572648 | 0,03529137 | TRNP1 | 0,2598456 |
| ENSG00000 | gene | 1,17356885 | 0,00573459 | 0,03531936 | OGFRL1 | 0,23090248 |
| ENSG00000 | gene | 0,79684003 | 0,00575349 | 0,03541171 | NUDT22 | -0,327638 |
| ENSG00000 | gene | 1,2006131 | 0,00576115 | 0,03541171 | FBN1 | 0,26377131 |
| ENSG00000 | gene | 1,1941803 | 0,00576235 | 0,03541171 | OPA1 | 0,25602068 |
| ENSG00000 | gene | 1,14293759 | 0,00576387 | 0,03541171 | FAM160B1 | 0,19274662 |
| ENSG00002 | gene | 1,1489678 | 0,00577253 | 0,03544297 | SBDSP1 | 0,20033836 |
| ENSG00000 | gene | 0,94223894 | 0,00578353 | 0,03548857 | FKBP15 | -0,0858351 |
| ENSG00000 | gene | 0,90690567 | 0,00579353 | 0,03552157 | SERTAD3 | -0,1409756 |
| ENSG00000 | gene | 0,80302545 | 0,00579608 | 0,03552157 | TARDBP | -0,3164824 |
| ENSG00000 | gene | 0,84744988 | 0,00580963 | 0,03558261 | TIMM9 | -0,2388 |
| ENSG00000 | gene | 0,85665948 | 0,00581402 | 0,03558757 | TRIM11 | -0,2232062 |
| ENSG00000 | gene | 1,07495063 | 0,00582557 | 0,03563622 | WDR75 | 0,1042704 |
| ENSG00000 | gene | 0,8579557 | 0,00584386 | 0,03570834 | H2AZ1 | -0,2210249 |
| ENSG00000 | gene | 1,10353417 | 0,00584456 | 0,03570834 | AXL | 0,1421313 |
| ENSG000000 | gene | 0,8819738 | 0,00587558 | 0,0358556 | SELENOO | -0,1811923 |
| ENSG00000 | gene | 0,77220386 | 0,00587589 | 0,0358556 | COX7A2L | -0,3729463 |
| ENSG00000 | gene | 0,90398714 | 0,0059059 | 0,03601653 | NPRL3 | -0,1456258 |
| ENSG00002 | gene | 0,83241441 | 0,00592054 | 0,03606179 | PAGR1 | -0,2646262 |
| ENSG000000 | gene | 0,83675469 | 0,00592228 | 0,03606179 | SLC66A1 | -0,2571234 |
| ENSG00002 | gene | 1,40563791 | 0,00592423 | 0,03606179 | AC015688.6 | 0,491225 |
| ENSG00000 | gene | 1,12647389 | 0,00594172 | 0,03614608 | CCND1 | 0,17181388 |
| ENSG00000 | gene | 0,86818021 | 0,00596985 | 0,03628572 | NAXE | -0,2039336 |
| ENSG00000 | gene | 0,91149047 | 0,00597199 | 0,03628572 | ZCCHC17 | -0,1337005 |
| ENSG00000 | gene | 0,80018156 | 0,00598718 | 0,03634076 | RAB5IF | -0,3216007 |
| ENSG00000 | gene | 1,10847012 | 0,00598838 | 0,03634076 | ZBTB18 | 0,14856988 |
| ENSG00000 | gene | 0,8756123 | 0,00600731 | 0,03643332 | CHMP1A | -0,1916359 |
| ENSG000000 | gene | 0,86254275 | 0,00603384 | 0,03655365 | BAD | -0,2133321 |
| ENSG00000 | gene | 0,91018065 | 0,00603452 | 0,03655365 | KAT2A | -0,1357752 |
| ENSG00002 | gene | 1,35979045 | 0,00605151 | 0,03663418 | AC025442.1 | 0,44338435 |
| ENSG00002 | gene | 1,4565083 | 0,00606852 | 0,03669305 | AP003071.3 | 0,54251392 |
| ENSG00000 | gene | 0,92996749 | 0,00606864 | 0,03669305 | ZFAND3 | -0,1047478 |
| ENSG00000 | gene | 1,14155964 | 0,00607632 | 0,03669606 | UEVLD | 0,19100623 |
| ENSG00000 | gene | 0,84119504 | 0,00607654 | 0,03669606 | RNF25 | -0,2494878 |
| ENSG000000 | gene | 1,09925862 | 0,00608093 | 0,03670027 | PSME4 | 0,13653085 |
| ENSG00000 | gene | 0,86754425 | 0,006101 | 0,036799 | PPIB | -0,2049907 |
| ENSG00000 | gene | 1,11832169 | 0,00611494 | 0,03686064 | PNN | 0,16133525 |
| ENSG00000 | gene | 1,17228664 | 0,00612295 | 0,03687645 | ALCAM | 0,22932537 |
| ENSG000000 | gene | 1,0894347 | 0,006125 | 0,03687645 | SUPT16H | 0,12357973 |
| ENSG00000 | gene | 0,86877438 | 0,00613386 | 0,03690738 | ELP5 | -0,2029465 |
| ENSG000000 | gene | 0,88741824 | 0,00613994 | 0,03692155 | DGCR2 | -0,1723139 |
| ENSG000000 | gene | 0,87698235 | 0,00614533 | 0,03693156 | GNAS | -0,1893803 |
| ENSG00000 | gene | 0,8607788 | 0,00617055 | 0,03706065 | RPL4 | -0,2162856 |

| | | | | | | |
|---|---|---|---|---|---|---|
| ENSG00000( | gene | 1,09504745 | 0,00618085 | 0,03708256 | SEL1L | 0,13099339 |
| ENSG00000' | gene | 0,83716798 | 0,00618357 | 0,03708256 | NABP2 | -0,256411 |
| ENSG00000: | gene | 1,27524744 | 0,00618541 | 0,03708256 | SMAD1-AS1 | 0,35077721 |
| ENSG00000: | gene | 1,60260269 | 0,00622226 | 0,03725655 | AL020996.3 | 0,6804168 |
| ENSG00000( | gene | 0,85374166 | 0,00622837 | 0,03725655 | NCK2 | -0,2281285 |
| ENSG00000' | gene | 0,68025188 | 0,00623316 | 0,03725655 | UBFD1 | -0,5558591 |
| ENSG00000' | gene | 0,84745124 | 0,00623368 | 0,03725655 | MOSPD3 | -0,2387977 |
| ENSG00000' | gene | 0,81962345 | 0,00623538 | 0,03725655 | MGRN1 | -0,2869668 |
| ENSG00000' | gene | 0,85752439 | 0,00623838 | 0,03725655 | AMPD2 | -0,2217504 |
| ENSG00000' | gene | 1,14089239 | 0,00624073 | 0,03725655 | ZC3H13 | 0,19016272 |
| ENSG00000( | gene | 0,88268525 | 0,00624904 | 0,03726215 | LRCH4 | -0,180029 |
| ENSG00000' | gene | 0,93730385 | 0,00624919 | 0,03726215 | CTDSP2 | -0,0934113 |
| ENSG00000' | gene | 0,91838176 | 0,00625866 | 0,03729369 | ARF3 | -0,1228341 |
| ENSG00000( | gene | 0,79742015 | 0,006262 | 0,03729369 | RPLP0 | -0,326588 |
| ENSG00000: | gene | 0,87889303 | 0,00628264 | 0,03736362 | AC133644.3 | -0,1862405 |
| ENSG00000' | gene | 0,58592284 | 0,00628575 | 0,03736362 | DGCR6 | -0,7712174 |
| ENSG00000: | gene | 0,92563235 | 0,00628863 | 0,03736362 | MLLT6 | -0,1114888 |
| ENSG00000' | gene | 0,8694057 | 0,00628881 | 0,03736362 | IRF3 | -0,2018985 |
| ENSG00000' | gene | 0,87842771 | 0,00633038 | 0,03757087 | NECAB3 | -0,1870045 |
| ENSG00000' | gene | 0,77099314 | 0,00633128 | 0,03757087 | ERVK3-1 | -0,3752101 |
| ENSG00000: | gene | 0,64943479 | 0,00634138 | 0,0376083 | MIR1244-3 | -0,6227434 |
| ENSG00000' | gene | 0,71007851 | 0,00636129 | 0,03770387 | SRRD | -0,4939495 |
| ENSG00000' | gene | 0,84803632 | 0,00638164 | 0,03780186 | SSR4 | -0,237802 |
| ENSG00000' | gene | 0,88869846 | 0,0063884 | 0,03781934 | RABGGTA | -0,1702341 |
| ENSG00000: | gene | 0,81495445 | 0,00639865 | 0,0378574 | CBSL | -0,2952087 |
| ENSG00000' | gene | 0,89660784 | 0,00640759 | 0,03785994 | COL27A1 | -0,157451 |
| ENSG00000' | gene | 0,70736433 | 0,00641237 | 0,03785994 | TMEM44 | -0,4994746 |
| ENSG00000: | gene | 1,24069754 | 0,00641385 | 0,03785994 | EEF1AKMT2 | 0,31115145 |
| ENSG00000' | gene | 0,8961289 | 0,00641667 | 0,03785994 | VRK3 | -0,1582218 |
| ENSG00000' | gene | 0,91001891 | 0,00641817 | 0,03785994 | PSMD6 | -0,1360316 |
| ENSG00000' | gene | 0,89092691 | 0,00644945 | 0,03802185 | RPS4Y1 | -0,166621 |
| ENSG00000' | gene | 1,06528073 | 0,00646432 | 0,03804953 | EHBP1 | 0,09123367 |
| ENSG00000' | gene | 0,85589774 | 0,00646498 | 0,03804953 | HMOX1 | -0,2244897 |
| ENSG00000( | gene | 0,80554064 | 0,00646566 | 0,03804953 | MAP2K7 | -0,3119707 |
| ENSG00000( | gene | 0,877945 | 0,00647152 | 0,03806142 | GDE1 | -0,1877975 |
| ENSG00000: | gene | 0,73146067 | 0,00647622 | 0,0380665 | AC018630.2 | -0,4511478 |
| ENSG00000: | gene | 0,66267123 | 0,00649163 | 0,03813451 | BCL2L2-PAE | -0,5936348 |
| ENSG00000' | gene | 1,13850157 | 0,00651896 | 0,03827158 | USP34 | 0,18713628 |
| ENSG00000' | gene | 0,77532434 | 0,00652326 | 0,03827158 | ASXL1 | -0,3671281 |
| ENSG00000( | gene | 1,13170417 | 0,00652655 | 0,03827158 | FAP | 0,17849689 |
| ENSG00000' | gene | 1,09232005 | 0,0065415 | 0,03832901 | SLC20A1 | 0,12739563 |
| ENSG00000' | gene | 0,90233104 | 0,00654407 | 0,03832901 | RHOT2 | -0,1482713 |
| ENSG00000' | gene | 1,09451374 | 0,00655377 | 0,03836318 | SEC23A | 0,13029007 |
| ENSG00000: | gene | 0,77598226 | 0,00656142 | 0,0383853 | ISY1 | -0,3659044 |
| ENSG00000' | gene | 0,80374646 | 0,00657797 | 0,03845938 | MSANTD2 | -0,3151876 |
| ENSG00000' | gene | 1,11397075 | 0,00658652 | 0,03847593 | AASDHPPT | 0,15571135 |
| ENSG00000' | gene | 1,5648206 | 0,00659043 | 0,03847593 | EME1 | 0,64599727 |
| ENSG00000' | gene | 0,93392737 | 0,00659244 | 0,03847593 | ASCC2 | -0,0986177 |
| ENSG00000' | gene | 0,90943051 | 0,00660561 | 0,03853016 | BTBD2 | -0,1369647 |
| ENSG00000' | gene | 0,94320676 | 0,00660998 | 0,03853296 | KIF1C | -0,084354 |
| ENSG00000' | gene | 0,91907026 | 0,00664429 | 0,03871023 | TUBG1 | -0,1217529 |
| ENSG00000' | gene | 1,32806517 | 0,00664846 | 0,03871177 | RAD17 | 0,40932595 |
| ENSG00000' | gene | 0,76987444 | 0,00665298 | 0,03871533 | KIFC2 | -0,3773049 |
| ENSG00000' | gene | 0,68930742 | 0,0066847 | 0,03886268 | MFAP4 | -0,5367806 |
| ENSG00000: | gene | 1,64063847 | 0,00668614 | 0,03886268 | AC124068.1 | 0,71425737 |
| ENSG00000' | gene | 0,94831557 | 0,00669145 | 0,03886888 | EHD4 | -0,0765609 |
| ENSG00000' | gene | 1,52794024 | 0,00670183 | 0,03886888 | TPGS1 | 0,61158812 |

| | | | | | | |
|---|---|---|---|---|---|---|
| ENSG00000; | gene | 1,39512179 | 0,00670284 | 0,03886888 | AC096536.2 | 0,48039107 |
| ENSG00000 | gene | 1,37902419 | 0,00670527 | 0,03886888 | ZNF644 | 0,46364776 |
| ENSG00000; | gene | 1,65775399 | 0,00671246 | 0,03886888 | AC090197.1 | 0,72922993 |
| ENSG00000 | gene | 1,11540478 | 0,00671456 | 0,03886888 | PRPF4B | 0,15756736 |
| ENSG00000; | gene | 1,30890136 | 0,00671464 | 0,03886888 | AL357140.1 | 0,38835638 |
| ENSG00000; | gene | 1,61104023 | 0,00671941 | 0,03887155 | AL022312.1 | 0,68799252 |
| ENSG00000 | gene | 1,26893319 | 0,00672294 | 0,03887155 | ENOX2 | 0,34361611 |
| ENSG00000 | gene | 0,84630866 | 0,00674011 | 0,03894809 | ACSF2 | -0,2407442 |
| ENSG00000 | gene | 0,87081325 | 0,00676896 | 0,039092 | STC2 | -0,1995647 |
| ENSG00000 | gene | 0,8430936 | 0,00678033 | 0,03913488 | FRMD8 | -0,2462353 |
| ENSG00000 | gene | 0,93769964 | 0,00679832 | 0,03921591 | ANO10 | -0,0928022 |
| ENSG00000 | gene | 0,73861982 | 0,00681817 | 0,03930756 | DEDD2 | -0,4370961 |
| ENSG00000 | gene | 0,70661627 | 0,00682332 | 0,03931438 | PNRC2 | -0,5010011 |
| ENSG00000( | gene | 0,8123169 | 0,00683902 | 0,03937037 | CD82 | -0,2998854 |
| ENSG00000 | gene | 0,88656272 | 0,00684098 | 0,03937037 | TRIM21 | -0,1737054 |
| ENSG00000 | gene | 1,14076412 | 0,00684551 | 0,03937361 | CDC73 | 0,19000051 |
| ENSG00000; | gene | 1,47808396 | 0,00685929 | 0,0394158 | AC007620.2 | 0,56372822 |
| ENSG00000 | gene | 0,82691298 | 0,00686815 | 0,0394158 | MZT2B | -0,2741926 |
| ENSG00000; | gene | 1,10408568 | 0,00686835 | 0,0394158 | AC240274.1 | 0,14285213 |
| ENSG00000 | gene | 0,85066741 | 0,00686875 | 0,0394158 | MAN2C1 | -0,2333329 |
| ENSG00000 | gene | 1,75285653 | 0,0068845 | 0,03948334 | NEK3 | 0,80970792 |
| ENSG00000( | gene | 0,91210155 | 0,00689481 | 0,03951963 | EIPR1 | -0,1327336 |
| ENSG00000 | gene | 1,14919524 | 0,00690821 | 0,03956691 | MRPL1 | 0,20062392 |
| ENSG00000( | gene | 1,22342861 | 0,00691126 | 0,03956691 | NDUFAF7 | 0,29092992 |
| ENSG00000 | gene | 0,81001921 | 0,00691503 | 0,03956691 | RPLP1 | -0,303972 |
| ENSG00000; | gene | 0,85428433 | 0,0069244 | 0,03959596 | MEG3 | -0,2272118 |
| ENSG00000( | gene | 0,86416228 | 0,00692809 | 0,03959596 | FTL | -0,2106258 |
| ENSG00000 | gene | 1,33847277 | 0,00693748 | 0,03962677 | CDC42EP3 | 0,42058779 |
| ENSG00000 | gene | 0,77790916 | 0,0069689 | 0,03978333 | JDP2 | -0,3623264 |
| ENSG00000 | gene | 0,87052112 | 0,00698098 | 0,03982933 | CCDC9 | -0,2000488 |
| ENSG00000 | gene | 0,80671459 | 0,00699481 | 0,03986323 | SGF29 | -0,3098697 |
| ENSG00000 | gene | 1,16478688 | 0,00699496 | 0,03986323 | SDE2 | 0,22006601 |
| ENSG00000; | gene | 1,12124907 | 0,00701739 | 0,03996806 | YAE1 | 0,16510679 |
| ENSG00000 | gene | 0,91892372 | 0,00703586 | 0,04005029 | GALNT2 | -0,121983 |
| ENSG00000 | gene | 0,86500777 | 0,00706063 | 0,04016821 | NMT1 | -0,209215 |
| ENSG00000 | gene | 1,26996649 | 0,00707929 | 0,04025131 | TMEM47 | 0,34479043 |
| ENSG00000( | gene | 1,14258685 | 0,00709332 | 0,04030792 | SUCO | 0,19230383 |
| ENSG00000 | gene | 1,07935128 | 0,00710029 | 0,0403225 | ITFG1 | 0,11016447 |
| ENSG00000; | gene | 1,30174023 | 0,00710401 | 0,0403225 | AL121845.1 | 0,38044158 |
| ENSG00000 | gene | 0,90618464 | 0,00715549 | 0,04059145 | AKT2 | -0,1421231 |
| ENSG00000 | gene | 0,81040663 | 0,0071763 | 0,04068622 | DTX3L | -0,3032821 |
| ENSG00000; | gene | 0,62333403 | 0,00719292 | 0,04075715 | AC002985.1 | -0,6819226 |
| ENSG00000; | gene | 0,83569779 | 0,0072018 | 0,04078415 | ELOCP2 | -0,2589468 |
| ENSG00000( | gene | 1,08282502 | 0,00721002 | 0,04080741 | RABEP1 | 0,11480013 |
| ENSG00000; | gene | 0,90231069 | 0,00721719 | 0,04082469 | RXRB | -0,1483038 |
| ENSG00000; | gene | 1,5377336 | 0,00723052 | 0,04087676 | AC011510.1 | 0,62080559 |
| ENSG00000 | gene | 0,89436041 | 0,0072371 | 0,04089062 | DDX42 | -0,1610718 |
| ENSG00000 | gene | 0,85124854 | 0,00724691 | 0,04092273 | ATXN2L | -0,2323477 |
| ENSG00000; | gene | 1,18579733 | 0,007257 | 0,04095195 | PPP1CB | 0,24585745 |
| ENSG00000 | gene | 0,89700426 | 0,00726604 | 0,04095195 | THUMPD3 | -0,1568133 |
| ENSG00000 | gene | 0,88234635 | 0,00727047 | 0,04095195 | DCTN2 | -0,180583 |
| ENSG00000 | gene | 0,9052526 | 0,0072709 | 0,04095195 | FBLN5 | -0,1436077 |
| ENSG00000 | gene | 1,09011247 | 0,00727273 | 0,04095195 | MIDN | 0,12447699 |
| ENSG00000; | gene | 1,42385996 | 0,00730043 | 0,04108461 | FAM200B | 0,50980726 |
| ENSG00000 | gene | 0,87678381 | 0,00732141 | 0,0411793 | DENR | -0,1897069 |
| ENSG00000 | gene | 0,90794513 | 0,00732882 | 0,0411976 | SLC2A4RG | -0,139323 |
| ENSG00000; | gene | 0,81977017 | 0,00733967 | 0,04123524 | FTH1P20 | -0,2867086 |

| | | | | | | |
|---|---|---|---|---|---|---|
| ENSG000001 | gene | 1,12403194 | 0,00739294 | 0,04151101 | PEX10 | 0,16868303 |
| ENSG000001 | gene | 1,11829029 | 0,00739987 | 0,04152636 | CHST3 | 0,16129474 |
| ENSG000001 | gene | 1,31832823 | 0,00740602 | 0,04153737 | NEXN | 0,39870961 |
| ENSG000000 | gene | 1,11299777 | 0,00742565 | 0,04162396 | TNRC6A | 0,15445071 |
| ENSG000001 | gene | 0,93622038 | 0,00743252 | 0,04163889 | COQ5 | -0,0950799 |
| ENSG000001 | gene | 1,22898427 | 0,00744316 | 0,04167497 | PRDM2 | 0,29746645 |
| ENSG000001 | gene | 0,81123377 | 0,00747276 | 0,0418171 | SHISA5 | -0,3018104 |
| ENSG000000 | gene | 0,82061496 | 0,0074777 | 0,04182113 | WIPI1 | -0,2852226 |
| ENSG000002 | gene | 0,75993506 | 0,00749648 | 0,04182767 | AC115223.1 | -0,3960519 |
| ENSG000001 | gene | 1,38641214 | 0,00749738 | 0,04182767 | USP25 | 0,47135619 |
| ENSG000002 | gene | 1,21672074 | 0,0074992 | 0,04182767 | CCDC39 | 0,28299808 |
| ENSG000001 | gene | 0,92436269 | 0,00750022 | 0,04182767 | UBAC1 | -0,1134691 |
| ENSG000002 | gene | 0,71284397 | 0,00750382 | 0,04182767 | LINC00667 | -0,4883418 |
| ENSG000002 | gene | 1,3324197 | 0,00750884 | 0,04182767 | AL049829.2 | 0,41404859 |
| ENSG000002 | gene | 1,83152891 | 0,00750946 | 0,04182767 | MYLK-AS1 | 0,87304847 |
| ENSG000001 | gene | 0,81352065 | 0,00751976 | 0,04182767 | PFDN1 | -0,2977491 |
| ENSG000002 | gene | 1,51111783 | 0,00752072 | 0,04182767 | AC008764.3 | 0,59561616 |
| ENSG000002 | gene | 1,31390308 | 0,00752105 | 0,04182767 | AL445183.1 | 0,39385886 |
| ENSG000002 | gene | 1,27245125 | 0,00753357 | 0,04187384 | AC093503.2 | 0,34761038 |
| ENSG000001 | gene | 0,91638626 | 0,00755646 | 0,04195626 | DPF2 | -0,1259723 |
| ENSG000001 | gene | 0,70165843 | 0,00755842 | 0,04195626 | TRABD2A | -0,5111592 |
| ENSG000000 | gene | 1,13550608 | 0,0075611 | 0,04195626 | OXCT1 | 0,18333543 |
| ENSG000001 | gene | 0,86384285 | 0,00756731 | 0,04196389 | DHCR7 | -0,2111592 |
| ENSG000002 | gene | 0,92178977 | 0,00757094 | 0,04196389 | CDK11B | -0,1174903 |
| ENSG000000 | gene | 0,82694515 | 0,00757767 | 0,04197774 | MKS1 | -0,2741364 |
| ENSG000001 | gene | 1,09194241 | 0,00762001 | 0,04216918 | DPYSL3 | 0,12689676 |
| ENSG000001 | gene | 1,07870841 | 0,00762073 | 0,04216918 | AP3B1 | 0,10930494 |
| ENSG000000 | gene | 1,24344174 | 0,00763325 | 0,04221489 | VPS50 | 0,31433891 |
| ENSG000002 | gene | 1,35766823 | 0,00765026 | 0,04228536 | AC073254.1 | 0,44113098 |
| ENSG000001 | gene | 0,89332597 | 0,00765831 | 0,0423063 | FHOD1 | -0,1627414 |
| ENSG000002 | gene | 1,25050943 | 0,0076742 | 0,0423705 | AL031775.1 | 0,32251594 |
| ENSG000000 | gene | 0,80713768 | 0,00769471 | 0,04245808 | SAMD4A | -0,3091133 |
| ENSG000002 | gene | 1,53577441 | 0,00769863 | 0,04245808 | AC087292.2 | 0,61896631 |
| ENSG000000 | gene | 0,84520907 | 0,00772076 | 0,04255643 | WDR54 | -0,2426198 |
| ENSG000000 | gene | 0,91466611 | 0,00774202 | 0,04264992 | TSR3 | -0,1286829 |
| ENSG000001 | gene | 0,90048528 | 0,00775207 | 0,04268159 | SGSM3 | -0,1512254 |
| ENSG000000 | gene | 0,92396223 | 0,00776221 | 0,04271369 | RBM5 | -0,1140942 |
| ENSG000001 | gene | 1,07007153 | 0,00777843 | 0,04277924 | VAPA | 0,09770724 |
| ENSG000001 | gene | 0,8857857 | 0,00779456 | 0,04282418 | GIPC1 | -0,1749704 |
| ENSG000001 | gene | 0,73130222 | 0,00779525 | 0,04282418 | MEN1 | -0,4514603 |
| ENSG000000 | gene | 0,87611649 | 0,00780645 | 0,04286197 | GNB5 | -0,1908054 |
| ENSG000001 | gene | 1,07899725 | 0,00782896 | 0,04296181 | THAP12 | 0,10969118 |
| ENSG000001 | gene | 0,74409174 | 0,00783453 | 0,04296861 | TUBB6 | -0,4264476 |
| ENSG000000 | gene | 0,85882286 | 0,00784544 | 0,04300466 | ALAS1 | -0,2195675 |
| ENSG000001 | gene | 1,58873949 | 0,00785837 | 0,0430517 | PDE5A | 0,66788258 |
| ENSG000001 | gene | 0,91876118 | 0,0078716 | 0,04310037 | DDX54 | -0,1222382 |
| ENSG000001 | gene | 1,28510091 | 0,00788313 | 0,04313969 | USP32P1 | 0,36188164 |
| ENSG000001 | gene | 1,08772927 | 0,00788751 | 0,04313984 | YWHAQ | 0,12131952 |
| ENSG000002 | gene | 1,47964988 | 0,00789572 | 0,04314682 | ABALON | 0,56525584 |
| ENSG000001 | gene | 0,8402004 | 0,00789749 | 0,04314682 | SAMD4B | -0,2511946 |
| ENSG000001 | gene | 0,70707478 | 0,00795971 | 0,0434628 | DDIT3 | -0,5000653 |
| ENSG000002 | gene | 1,41696531 | 0,00797023 | 0,04349631 | AC108863.2 | 0,50280443 |
| ENSG000001 | gene | 0,86907039 | 0,00799209 | 0,0435916 | PTGES2 | -0,2024551 |
| ENSG000001 | gene | 0,91801143 | 0,00801445 | 0,04368955 | IMPDH1 | -0,123416 |
| ENSG000001 | gene | 1,17381816 | 0,00805616 | 0,04389281 | SPAG16 | 0,23120894 |
| ENSG000001 | gene | 1,11743595 | 0,00806388 | 0,04391072 | MTERF3 | 0,16019215 |
| ENSG000001 | gene | 0,7404046 | 0,00807369 | 0,04394 | ADARB1 | -0,4336142 |

| | | | | | | |
|---|---|---|---|---|---|---|
| ENSG00002 | gene | 1,2669694 | 0,00808836 | 0,04397248 | ANKRD36BF | 0,34138168 |
| ENSG00000 | gene | 0,68992249 | 0,00809172 | 0,04397248 | KDM6A | -0,5354938 |
| ENSG00002 | gene | 1,28669589 | 0,00809296 | 0,04397248 | AC010978.1 | 0,36367112 |
| ENSG00000 | gene | 0,93632393 | 0,00814972 | 0,04425665 | DNAJC5 | -0,0949204 |
| ENSG00000 | gene | 1,23832388 | 0,00815597 | 0,04426635 | FBXO5 | 0,30838869 |
| ENSG00000 | gene | 1,11302281 | 0,00819337 | 0,04444501 | TRAF5 | 0,15448316 |
| ENSG00000 | gene | 0,84400738 | 0,00823112 | 0,04462535 | CISD2 | -0,2446725 |
| ENSG00000 | gene | 0,85161251 | 0,00824161 | 0,04465782 | SLC6A9 | -0,231731 |
| ENSG00000 | gene | 0,91678654 | 0,00827196 | 0,04479776 | LZTS2 | -0,1253422 |
| ENSG00000 | gene | 1,27606377 | 0,00827962 | 0,04480127 | HEATR5B | 0,35170043 |
| ENSG00002 | gene | 0,62266792 | 0,0082844 | 0,04480127 | LINC00339 | -0,6834651 |
| ENSG00002 | gene | 0,78212295 | 0,00829079 | 0,04480127 | RPL23AP7 | -0,3545327 |
| ENSG00000 | gene | 0,9199904 | 0,00829138 | 0,04480127 | RAB11FIP3 | -0,1203093 |
| ENSG00000 | gene | 0,89984573 | 0,00829671 | 0,04480127 | RRP1 | -0,1522504 |
| ENSG00000 | gene | 1,22887747 | 0,00829971 | 0,04480127 | LHFPL5 | 0,29734107 |
| ENSG00000 | gene | 1,39171225 | 0,00830681 | 0,04480483 | FAM166A | 0,47686095 |
| ENSG00000 | gene | 1,1607087 | 0,00830941 | 0,04480483 | CASP3 | 0,21500595 |
| ENSG00000 | gene | 1,13875737 | 0,00832576 | 0,04486862 | GNG12 | 0,18746039 |
| ENSG00002 | gene | 1,23483874 | 0,00833091 | 0,04487199 | AC073641.1 | 0,30432265 |
| ENSG00000 | gene | 0,9317789 | 0,00833994 | 0,04489624 | CYP51A1 | -0,1019404 |
| ENSG00000 | gene | 0,85841734 | 0,00836002 | 0,0449799 | ZNF598 | -0,2202489 |
| ENSG00000 | gene | 0,87998035 | 0,00837112 | 0,04499221 | ASB13 | -0,1844568 |
| ENSG00000 | gene | 0,87733576 | 0,00837138 | 0,04499221 | BRMS1 | -0,188799 |
| ENSG00000 | gene | 0,94685345 | 0,00837879 | 0,04500763 | HNRNPK | -0,0787869 |
| ENSG00000 | gene | 0,86400937 | 0,00838842 | 0,045032 | CNIH4 | -0,2108811 |
| ENSG00000 | gene | 0,89257589 | 0,00839241 | 0,045032 | COMMD4 | -0,1639533 |
| ENSG00000 | gene | 0,87006634 | 0,00839895 | 0,04504273 | LAMTOR4 | -0,2008027 |
| ENSG00000 | gene | 0,81723069 | 0,00842277 | 0,04513084 | POLR1C | -0,2911847 |
| ENSG00000 | gene | 0,89618761 | 0,00842448 | 0,04513084 | YY1AP1 | -0,1581273 |
| ENSG00000 | gene | 0,92571762 | 0,00843516 | 0,04514288 | BRD8 | -0,1113559 |
| ENSG00002 | gene | 1,39884951 | 0,00843584 | 0,04514288 | RNU7-111P | 0,48424077 |
| ENSG00000 | gene | 0,80455277 | 0,00846791 | 0,04528946 | POMT2 | -0,3137411 |
| ENSG00000 | gene | 1,1549125 | 0,00847428 | 0,04528946 | ITGAV | 0,20778355 |
| ENSG00002 | gene | 1,52978906 | 0,00847693 | 0,04528946 | SNORA64 | 0,61333273 |
| ENSG00000 | gene | 0,84487354 | 0,00849987 | 0,04538082 | MPV17 | -0,2431927 |
| ENSG00000 | gene | 0,81472854 | 0,00850318 | 0,04538082 | ING1 | -0,2956086 |
| ENSG00002 | gene | 1,58005377 | 0,00855324 | 0,04560324 | AC079328.2 | 0,65997366 |
| ENSG00000 | gene | 0,74243601 | 0,00855406 | 0,04560324 | MAPK7 | -0,4296614 |
| ENSG00000 | gene | 0,87315302 | 0,00856304 | 0,0456266 | RPS5 | -0,1956936 |
| ENSG00000 | gene | 0,60803515 | 0,00858468 | 0,04571732 | ZNF7 | -0,7177734 |
| ENSG00000 | gene | 0,89232246 | 0,00861524 | 0,04585544 | CFDP1 | -0,1643629 |
| ENSG00000 | gene | 0,9350558 | 0,00862839 | 0,04590081 | NCL | -0,0968756 |
| ENSG00002 | gene | 1,46111214 | 0,00865391 | 0,04601191 | AL031600.3 | 0,54706691 |
| ENSG00002 | gene | 1,28189675 | 0,00871213 | 0,04628916 | AL022328.4 | 0,35828007 |
| ENSG00000 | gene | 1,15537904 | 0,00871787 | 0,04628916 | TSC2 | 0,20836623 |
| ENSG00002 | gene | 1,24941931 | 0,00872232 | 0,04628916 | AL391069.1 | 0,32125773 |
| ENSG00000 | gene | 1,09922958 | 0,00872743 | 0,04628916 | HIPK2 | 0,13649274 |
| ENSG00000 | gene | 0,94714775 | 0,00873204 | 0,04628916 | ADIPOR1 | -0,0783386 |
| ENSG00002 | gene | 1,37800078 | 0,00873407 | 0,04628916 | AC139530.1 | 0,4625767 |
| ENSG00000 | gene | 0,92436698 | 0,00874786 | 0,04633748 | CORO1B | -0,1134624 |
| ENSG00000 | gene | 1,29341716 | 0,00875313 | 0,04634066 | GLIPR1 | 0,37118766 |
| ENSG00000 | gene | 0,82964796 | 0,00877384 | 0,04642549 | RPS21 | -0,2694288 |
| ENSG00002 | gene | 1,58742365 | 0,00878213 | 0,04644458 | MIR6720 | 0,66668721 |
| ENSG00000 | gene | 1,10300251 | 0,00878917 | 0,04645706 | DPH3 | 0,14143608 |
| ENSG00000 | gene | 0,94916753 | 0,00881336 | 0,04656011 | CPSF6 | -0,0752653 |
| ENSG00000 | gene | 0,92179635 | 0,00882953 | 0,04662064 | EFNB2 | -0,11748 |
| ENSG00002 | gene | 0,85590641 | 0,00883612 | 0,04663066 | RPS28 | -0,224475 |

| | | | | | | |
|---|---|---|---|---|---|---|
| ENSG00000 | gene | 0,84123439 | 0,00884278 | 0,04664098 | RPLP2 | -0,2494203 |
| ENSG00002 | gene | 1,99056805 | 0,00885576 | 0,04668459 | TRIM13 | 0,99318019 |
| ENSG00000 | gene | 0,89457652 | 0,00887914 | 0,04674795 | BAMBI | -0,1607232 |
| ENSG00000 | gene | 0,88749549 | 0,00887983 | 0,04674795 | UBTD1 | -0,1721883 |
| ENSG00000 | gene | 0,67556087 | 0,00888426 | 0,04674795 | TGIF1 | -0,5658423 |
| ENSG00000 | gene | 0,88341364 | 0,00888713 | 0,04674795 | SNX3 | -0,178839 |
| ENSG00000 | gene | 1,13194692 | 0,00889302 | 0,04674795 | TCERG1 | 0,17880631 |
| ENSG00002 | gene | 1,58853604 | 0,0088997 | 0,04674795 | AL121748.1 | 0,66769782 |
| ENSG00000 | gene | 0,90123314 | 0,00890894 | 0,04674795 | ATG13 | -0,1500277 |
| ENSG00002 | gene | 0,80156862 | 0,00891486 | 0,04674795 | TTLL3 | -0,3191021 |
| ENSG00000 | gene | 0,87381637 | 0,00891499 | 0,04674795 | HRH1 | -0,194598 |
| ENSG00000 | gene | 0,84655783 | 0,00891606 | 0,04674795 | NGDN | -0,2403195 |
| ENSG00000 | gene | 1,1819348 | 0,00891964 | 0,04674795 | TMEM123 | 0,24115045 |
| ENSG00000 | gene | 1,06699489 | 0,0089287 | 0,04677073 | PDCD6IP | 0,09355327 |
| ENSG00000 | gene | 0,81188215 | 0,0089561 | 0,04688946 | COX4I1 | -0,3006578 |
| ENSG00002 | gene | 1,30407063 | 0,00897743 | 0,04697636 | EML4-AS1 | 0,38302201 |
| ENSG00002 | gene | 0,81982388 | 0,00898378 | 0,04698477 | PCGF2 | -0,2866141 |
| ENSG00000 | gene | 0,86883316 | 0,00899384 | 0,04699212 | RNF181 | -0,2028489 |
| ENSG00000 | gene | 1,41953909 | 0,00899466 | 0,04699212 | METTL14 | 0,50542258 |
| ENSG00000 | gene | 1,27237775 | 0,00902604 | 0,04711324 | HMMR | 0,34752705 |
| ENSG00000 | gene | 0,65982967 | 0,00902735 | 0,04711324 | UNC119 | -0,5998345 |
| ENSG00000 | gene | 0,88163947 | 0,00904746 | 0,04719336 | GTF3C2 | -0,1817393 |
| ENSG00000 | gene | 0,91962497 | 0,00905485 | 0,04720707 | PPM1G | -0,1208825 |
| ENSG00000 | gene | 0,78527913 | 0,00906016 | 0,04720998 | HDDC3 | -0,3487225 |
| ENSG00000 | gene | 0,93958621 | 0,00908417 | 0,04727262 | LRPAP1 | -0,0899026 |
| ENSG00000 | gene | 0,86920851 | 0,00908815 | 0,04727262 | SELPLG | -0,2022258 |
| ENSG00000 | gene | 0,75333255 | 0,00909338 | 0,04727262 | LDLR | -0,4086412 |
| ENSG00002 | gene | 1,54048411 | 0,00909576 | 0,04727262 | RCC2-AS1 | 0,6233838 |
| ENSG00000 | gene | 0,94189766 | 0,00909602 | 0,04727262 | AAR2 | -0,0863578 |
| ENSG00000 | gene | 0,90937361 | 0,0091039 | 0,04728877 | EPHB4 | -0,137055 |
| ENSG00002 | gene | 1,33458681 | 0,00918073 | 0,04762325 | AC007308.1 | 0,41639315 |
| ENSG00000 | gene | 1,30681245 | 0,00918157 | 0,04762325 | TUBGCP4 | 0,3860521 |
| ENSG00000 | gene | 0,90984691 | 0,00918544 | 0,04762325 | GNPDA1 | -0,1363043 |
| ENSG00002 | gene | 1,78503187 | 0,0091875 | 0,04762325 | AP000942.2 | 0,83594983 |
| ENSG00000 | gene | 0,87520362 | 0,00921361 | 0,04773361 | CASP4 | -0,1923094 |
| ENSG00000 | gene | 0,86587617 | 0,00923405 | 0,04781455 | MNT | -0,2077674 |
| ENSG00000 | gene | 1,21755154 | 0,00924221 | 0,04783182 | SLC4A7 | 0,28398284 |
| ENSG00002 | gene | 0,89330441 | 0,00925666 | 0,0478816 | CISD3 | -0,1627762 |
| ENSG00000 | gene | 0,73241011 | 0,00926479 | 0,04789867 | PEX1 | -0,4492764 |
| ENSG00000 | gene | 1,14627132 | 0,0092771 | 0,04793731 | SERPINE1 | 0,19694857 |
| ENSG00000 | gene | 1,07560017 | 0,00929832 | 0,04800346 | TTC27 | 0,10514189 |
| ENSG00000 | gene | 0,81792149 | 0,00930277 | 0,04800346 | ETV4 | -0,2899657 |
| ENSG00000 | gene | 0,84538439 | 0,00930866 | 0,04800346 | CD68 | -0,2423206 |
| ENSG00000 | gene | 0,735048 | 0,00930926 | 0,04800346 | ASB6 | -0,4440896 |
| ENSG00000 | gene | 0,88386564 | 0,00932795 | 0,04807483 | NFE2L1 | -0,178101 |
| ENSG00002 | gene | 1,37695277 | 0,00934878 | 0,04813974 | AL354811.1 | 0,46147907 |
| ENSG00000 | gene | 0,87119374 | 0,00935179 | 0,04813974 | SLC2A1 | -0,1989345 |
| ENSG00000 | gene | 0,88009529 | 0,00935511 | 0,04813974 | HMG20B | -0,1842684 |
| ENSG00000 | gene | 1,13981447 | 0,00936859 | 0,04818411 | ADAM10 | 0,18879902 |
| ENSG00002 | gene | 0,67455222 | 0,00938909 | 0,04824631 | AP000944.5 | -0,567998 |
| ENSG00000 | gene | 0,91947327 | 0,00939042 | 0,04824631 | DCAF7 | -0,1211205 |
| ENSG00000 | gene | 0,86830099 | 0,00939968 | 0,0482689 | MRPS28 | -0,2037329 |
| ENSG00000 | gene | 0,84914865 | 0,00940931 | 0,04829334 | TNIP1 | -0,235911 |
| ENSG00002 | gene | 1,27086655 | 0,00941626 | 0,04830401 | AL137060.5 | 0,34581255 |
| ENSG00000 | gene | 0,87321621 | 0,00943024 | 0,04835071 | TRAF2 | -0,1955892 |
| ENSG00000 | gene | 0,87893397 | 0,00945593 | 0,04845735 | UBL5 | -0,1861733 |
| ENSG00000 | gene | 0,89416894 | 0,00948967 | 0,04860517 | ATP6V1G1 | -0,1613807 |

| | | | | | | |
|---|---|---|---|---|---|---|
| ENSG00000( | gene | 0,92382513 | 0,00950667 | 0,04864553 | ERP29 | -0,1143083 |
| ENSG00000' | gene | 0,90620826 | 0,00950737 | 0,04864553 | GMPPA | -0,1420855 |
| ENSG00002 | gene | 0,71802116 | 0,00956889 | 0,04893508 | TMEM185A | -0,4779017 |
| ENSG00000' | gene | 0,87970826 | 0,00959234 | 0,0490297 | SNUPN | -0,1849029 |
| ENSG00000' | gene | 0,81533813 | 0,00960038 | 0,0490455 | PCBP1 | -0,2945296 |
| ENSG00000' | gene | 0,9319787 | 0,00961012 | 0,04907003 | C19orf12 | -0,1016311 |
| ENSG00002 | gene | 1,35564154 | 0,00962827 | 0,04913737 | HIF1A-AS1 | 0,43897575 |
| ENSG00000' | gene | 0,82477878 | 0,00963451 | 0,04914393 | ATF4 | -0,2779209 |
| ENSG00000' | gene | 0,92178454 | 0,00966655 | 0,04924862 | ACO2 | -0,1174985 |
| ENSG00000' | gene | 0,77544587 | 0,00966967 | 0,04924862 | NDRG1 | -0,366902 |
| ENSG00000' | gene | 0,86437317 | 0,00966993 | 0,04924862 | SLC35A2 | -0,2102738 |
| ENSG00000' | gene | 0,8556834 | 0,00967545 | 0,04925144 | FCGRT | -0,224851 |
| ENSG00000' | gene | 0,87988569 | 0,00969048 | 0,04930259 | BAHD1 | -0,184612 |
| ENSG00000' | gene | 0,8793158 | 0,00970013 | 0,04932642 | PLIN3 | -0,1855467 |
| ENSG00000' | gene | 0,8130136 | 0,00972464 | 0,04941716 | RSRP1 | -0,2986486 |
| ENSG00000' | gene | 0,91338621 | 0,00972795 | 0,04941716 | EDF1 | -0,1307031 |
| ENSG00000' | gene | 0,86817076 | 0,009778 | 0,049646 | TNIP2 | -0,2039493 |
| ENSG00000' | gene | 0,88720914 | 0,00983241 | 0,04988602 | PMM2 | -0,1726539 |
| ENSG00000' | gene | 1,49610423 | 0,00983533 | 0,04988602 | PARPBP | 0,58121068 |
| ENSG00000( | gene | 1,13614756 | 0,00984806 | 0,04992504 | MAPK6 | 0,18415022 |
| ENSG00000' | gene | 0,8076246 | 0,00988737 | 0,05009868 | PLEKHB2 | -0,3082432 |
| ENSG00000' | gene | 1,52444685 | 0,00991886 | 0,0502326 | EXD3 | 0,60828585 |
| ENSG00000' | gene | 0,89397842 | 0,00998352 | 0,05053423 | EMC8 | -0,1616881 |
| ENSG00000' | gene | 1,20008323 | 0,01001201 | 0,05065262 | OBI1 | 0,26313447 |
| ENSG00000' | gene | 0,86373474 | 0,01002981 | 0,05071679 | FUT11 | -0,2113398 |
| ENSG00000' | gene | 0,84291929 | 0,010047 | 0,0507778 | GATD3A | -0,2465336 |
| ENSG00000' | gene | 0,94648038 | 0,01006304 | 0,05083294 | CASC3 | -0,0793555 |
| ENSG00000' | gene | 1,18680222 | 0,01007838 | 0,05086757 | CSNK1G3 | 0,24707953 |
| ENSG00000' | gene | 0,60180318 | 0,01008015 | 0,05086757 | INSIG2 | -0,7326364 |
| ENSG00000' | gene | 0,83713444 | 0,01008638 | 0,05087311 | RAB5C | -0,2564688 |
| ENSG00000' | gene | 0,83671299 | 0,01014074 | 0,05108996 | PHF1 | -0,2571953 |
| ENSG00000( | gene | 0,89111568 | 0,01014362 | 0,05108996 | XAB2 | -0,1663154 |
| ENSG00000' | gene | 0,89601769 | 0,01014539 | 0,05108996 | ZMYND19 | -0,1584009 |
| ENSG00000' | gene | 0,86983369 | 0,01015335 | 0,05108996 | PCOLCE | -0,2011885 |
| ENSG00000' | gene | 0,88504229 | 0,01015513 | 0,05108996 | JOSD2 | -0,1761817 |
| ENSG00000' | gene | 1,11428401 | 0,01017172 | 0,05114747 | EIF1AY | 0,15611699 |
| ENSG00002 | gene | 1,5588784 | 0,01017819 | 0,05115407 | AC025423.4 | 0,6405084 |
| ENSG00000' | gene | 1,33410227 | 0,01019519 | 0,05121355 | CNOT6 | 0,41586926 |
| ENSG00000' | gene | 1,40855129 | 0,0102293 | 0,05135887 | CENPBD1 | 0,4942121 |
| ENSG00000' | gene | 1,1544788 | 0,01023459 | 0,05135939 | PRKAA1 | 0,20724168 |
| ENSG00002 | gene | 1,34553788 | 0,01024222 | 0,05137171 | MIR1184-1 | 0,42818301 |
| ENSG00000' | gene | 0,82639872 | 0,01025376 | 0,05140357 | MRPL10 | -0,2750901 |
| ENSG00002 | gene | 1,21604646 | 0,01029861 | 0,05159272 | AC123768.5 | 0,28219835 |
| ENSG00000' | gene | 0,88720239 | 0,01030189 | 0,05159272 | SRSF9 | -0,1726649 |
| ENSG00000' | gene | 0,9122124 | 0,01032321 | 0,05160715 | DDRGK1 | -0,1325583 |
| ENSG00000' | gene | 1,40214724 | 0,01032461 | 0,05160715 | AMN | 0,48763786 |
| ENSG00002 | gene | 0,89695153 | 0,01033015 | 0,05160715 | MRPL20 | -0,1568981 |
| ENSG00000' | gene | 0,68924853 | 0,01033223 | 0,05160715 | SNX15 | -0,5369038 |
| ENSG00000' | gene | 0,89543739 | 0,01033256 | 0,05160715 | HTATIP2 | -0,1593355 |
| ENSG00000' | gene | 0,78204038 | 0,0103406 | 0,05160715 | CFAP298 | -0,354685 |
| ENSG00000' | gene | 1,30802908 | 0,01034121 | 0,05160715 | NABP1 | 0,38739462 |
| ENSG00000' | gene | 1,3270938 | 0,01036168 | 0,0516485 | FBXL17 | 0,40827034 |
| ENSG00000' | gene | 0,92119498 | 0,01036764 | 0,0516485 | FIP1L1 | -0,1184215 |
| ENSG00000' | gene | 0,81059595 | 0,01036923 | 0,0516485 | TIMM10 | -0,3029451 |
| ENSG00000' | gene | 0,5566027 | 0,01037033 | 0,0516485 | IL6 | -0,8452802 |
| ENSG00000( | gene | 1,16644745 | 0,01047133 | 0,05212536 | SEC62 | 0,22212131 |
| ENSG00000( | gene | 1,48568045 | 0,0104978 | 0,05223089 | WDR47 | 0,57112385 |

| | | | | | | |
|---|---|---|---|---|---|---|
| ENSG00000 | gene | 0,7891434 | 0,01050517 | 0,05224137 | C12orf57 | -0,3416406 |
| ENSG00000 | gene | 1,10504604 | 0,01051408 | 0,05225944 | SBDS | 0,14410648 |
| ENSG00000 | gene | 1,66918508 | 0,01052082 | 0,05226678 | AC005943.2 | 0,73914393 |
| ENSG00000 | gene | 0,81656289 | 0,01053066 | 0,05228872 | DAD1 | -0,2923641 |
| ENSG00000 | gene | 1,22002688 | 0,01053579 | 0,05228872 | DOC2A | 0,28691293 |
| ENSG00000 | gene | 0,60437516 | 0,01054502 | 0,05230835 | AC242498.1 | -0,7264837 |
| ENSG00000 | gene | 0,89156296 | 0,01056384 | 0,0523755 | FDX2 | -0,1655914 |
| ENSG00000 | gene | 1,22597482 | 0,01057647 | 0,0523951 | IDI2 | 0,29392935 |
| ENSG00000 | gene | 0,86904822 | 0,01057841 | 0,0523951 | WBP1 | -0,2024919 |
| ENSG00000 | gene | 0,89064924 | 0,0105871 | 0,0523951 | UQCRH | -0,1670707 |
| ENSG00000 | gene | 0,94255519 | 0,01059084 | 0,0523951 | REXO4 | -0,085351 |
| ENSG00000 | gene | 0,65700453 | 0,01059951 | 0,0523951 | HAUS7 | -0,6060248 |
| ENSG00000 | gene | 1,37655655 | 0,01060413 | 0,0523951 | AL133410.2 | 0,46106387 |
| ENSG00000 | gene | 0,90526435 | 0,01060478 | 0,0523951 | GTF3C5 | -0,1435889 |
| ENSG00000 | gene | 0,92601809 | 0,01061414 | 0,05241525 | PLD2 | -0,1108877 |
| ENSG00000 | gene | 0,89195257 | 0,01062316 | 0,05243048 | CUEDC2 | -0,1649611 |
| ENSG00000 | gene | 0,87228252 | 0,0106278 | 0,05243048 | COX20P1 | -0,1971326 |
| ENSG00000 | gene | 1,42533402 | 0,01064215 | 0,05244811 | AL355987.6 | 0,51130005 |
| ENSG00000 | gene | 0,89013351 | 0,01064232 | 0,05244811 | TNKS1BP1 | -0,1679064 |
| ENSG00000 | gene | 1,15072634 | 0,01064901 | 0,05244811 | STXBP3 | 0,20254477 |
| ENSG00000 | gene | 0,81529299 | 0,01065253 | 0,05244811 | NKAPD1 | -0,2946095 |
| ENSG00000 | gene | 1,30439116 | 0,01067519 | 0,05250071 | MIR1279 | 0,38337657 |
| ENSG00000 | gene | 0,92279403 | 0,01067754 | 0,05250071 | EBNA1BP2 | -0,1159194 |
| ENSG00000 | gene | 0,84918434 | 0,0106791 | 0,05250071 | ZFYVE19 | -0,2358503 |
| ENSG00000 | gene | 1,40574112 | 0,0106848 | 0,05250271 | AL358333.3 | 0,49133093 |
| ENSG00000 | gene | 1,09891277 | 0,01069313 | 0,05251761 | RPF2 | 0,13607687 |
| ENSG00000 | gene | 1,48865351 | 0,01072482 | 0,05264718 | AL137782.1 | 0,574008 |
| ENSG00000 | gene | 0,91805301 | 0,01073287 | 0,05266061 | PDE6D | -0,1233506 |
| ENSG00000 | gene | 1,13588884 | 0,01074079 | 0,05267341 | BTAF1 | 0,18382165 |
| ENSG00000 | gene | 0,82456575 | 0,01076232 | 0,05275294 | CCDC84 | -0,2782936 |
| ENSG00000 | gene | 1,20530488 | 0,01080922 | 0,05292117 | SH3BGRL | 0,26939812 |
| ENSG00000 | gene | 0,90227807 | 0,01081186 | 0,05292117 | DDAH2 | -0,148356 |
| ENSG00000 | gene | 1,27739546 | 0,01081266 | 0,05292117 | SNORD4A | 0,35320523 |
| ENSG00000 | gene | 0,53188877 | 0,01082358 | 0,05293342 | NR4A1 | -0,9108035 |
| ENSG00000 | gene | 1,23638744 | 0,01082584 | 0,05293342 | AC069234.3 | 0,3061309 |
| ENSG00000 | gene | 0,51237977 | 0,01087286 | 0,05313714 | NPIPP1 | -0,9647146 |
| ENSG00000 | gene | 1,53098434 | 0,01088963 | 0,05319287 | AC013451.1 | 0,61445953 |
| ENSG00000 | gene | 1,15320326 | 0,01090242 | 0,05319439 | MOB4 | 0,20564683 |
| ENSG00000 | gene | 1,33678924 | 0,01090481 | 0,05319439 | AL355032.1 | 0,41877203 |
| ENSG00000 | gene | 1,25000959 | 0,01090603 | 0,05319439 | AL391839.2 | 0,32193916 |
| ENSG00000 | gene | 1,4068842 | 0,01091259 | 0,05320022 | SNORD68 | 0,49250359 |
| ENSG00000 | gene | 0,93098814 | 0,01094101 | 0,05330984 | PHC2 | -0,1031653 |
| ENSG00000 | gene | 1,28651981 | 0,01094583 | 0,05330984 | NCBP3 | 0,36347367 |
| ENSG00000 | gene | 0,78693804 | 0,01095932 | 0,05334934 | PDPR | -0,3456781 |
| ENSG00000 | gene | 1,08884456 | 0,01096795 | 0,05336515 | RTRAF | 0,12279802 |
| ENSG00000 | gene | 0,88768837 | 0,01099182 | 0,05344524 | GALE | -0,1718748 |
| ENSG00000 | gene | 0,83474224 | 0,01099519 | 0,05344524 | FGFR1 | -0,2605973 |
| ENSG00000 | gene | 0,9038822 | 0,01100569 | 0,05347007 | NELFCD | -0,1457933 |
| ENSG00000 | gene | 0,90848102 | 0,01106756 | 0,05374432 | WASF3 | -0,1384717 |
| ENSG00000 | gene | 0,8154456 | 0,01107332 | 0,05374597 | C19orf47 | -0,2943395 |
| ENSG00000 | gene | 0,71103605 | 0,01112109 | 0,05394131 | PCTP | -0,4920054 |
| ENSG00000 | gene | 1,24648119 | 0,01112444 | 0,05394131 | SLC25A5-AS | 0,31786111 |
| ENSG00000 | gene | 0,89204754 | 0,0111627 | 0,05410035 | PRDX5 | -0,1648075 |
| ENSG00000 | gene | 0,82892082 | 0,01119717 | 0,05424089 | ILK | -0,2706938 |
| ENSG00000 | gene | 1,14609206 | 0,0112087 | 0,05427025 | BAG2 | 0,19672293 |
| ENSG00000 | gene | 0,82221136 | 0,01122033 | 0,05430004 | NINJ1 | -0,2824188 |
| ENSG00000 | gene | 0,73028777 | 0,01122858 | 0,05431348 | MIR570 | -0,453463 |

| | | | | | | |
|---|---|---|---|---|---|---|
| ENSG00000 | gene | 0,93694899 | 0,01123533 | 0,05431961 | DNTTIP1 | -0,0939576 |
| ENSG00000 | gene | 1,43870486 | 0,01125448 | 0,05437242 | SLC35A3 | 0,52477067 |
| ENSG00002 | gene | 1,20557252 | 0,01125722 | 0,05437242 | AC015802.6 | 0,26971844 |
| ENSG00000 | gene | 0,88966643 | 0,01126332 | 0,05437542 | PIGT | -0,1686636 |
| ENSG00000 | gene | 1,16542867 | 0,01129217 | 0,05448815 | STYX | 0,2208607 |
| ENSG00000 | gene | 1,35599783 | 0,01132757 | 0,05463239 | THBS1 | 0,43935487 |
| ENSG00000 | gene | 1,0866277 | 0,01135494 | 0,05473183 | DDX18 | 0,11985772 |
| ENSG00000 | gene | 0,75790162 | 0,01136841 | 0,05473183 | COX7C | -0,3999175 |
| ENSG00000 | gene | 0,89051244 | 0,01137132 | 0,05473183 | CTSL | -0,1672923 |
| ENSG00002 | gene | 0,89829794 | 0,01137265 | 0,05473183 | MT-RNR1 | -0,1547341 |
| ENSG00000 | gene | 0,7982343 | 0,01137579 | 0,05473183 | CLCN7 | -0,3251158 |
| ENSG00000 | gene | 0,88836427 | 0,01139351 | 0,0547905 | CARS1 | -0,1707767 |
| ENSG00000 | gene | 1,28816541 | 0,01140788 | 0,05483304 | MYBL1 | 0,36531786 |
| ENSG00000 | gene | 1,08066789 | 0,01143072 | 0,05486859 | PKD2 | 0,11192322 |
| ENSG00000 | gene | 1,12175974 | 0,01143301 | 0,05486859 | RDX | 0,16576371 |
| ENSG00000 | gene | 1,21604075 | 0,01144115 | 0,05486859 | TMEM267 | 0,28219158 |
| ENSG00000 | gene | 1,1291587 | 0,01144237 | 0,05486859 | HERC4 | 0,17524827 |
| ENSG00000 | gene | 0,9416057 | 0,01144625 | 0,05486859 | UBL4A | -0,086805 |
| ENSG00000 | gene | 0,84785223 | 0,01144848 | 0,05486859 | SNX32 | -0,2381152 |
| ENSG00000 | gene | 1,232199 | 0,01145865 | 0,05488355 | WDR59 | 0,30123527 |
| ENSG00000 | gene | 1,10912646 | 0,01146613 | 0,05488355 | FOCAD | 0,14942387 |
| ENSG00000 | gene | 1,18713877 | 0,0114682 | 0,05488355 | PKN2 | 0,24748859 |
| ENSG00000 | gene | 0,89142125 | 0,01148359 | 0,05493068 | INTS11 | -0,1658207 |
| ENSG00000 | gene | 0,72254042 | 0,01150673 | 0,0550148 | SQSTM1 | -0,4688498 |
| ENSG00000 | gene | 1,07052443 | 0,01151855 | 0,0550448 | GSPT1 | 0,09831772 |
| ENSG00000 | gene | 0,924107 | 0,01153211 | 0,05508305 | PLBD2 | -0,1138682 |
| ENSG00000 | gene | 0,87901308 | 0,01153833 | 0,05508623 | MAP11 | -0,1860435 |
| ENSG00002 | gene | 1,29910969 | 0,01155473 | 0,05513799 | AL356652.1 | 0,37752325 |
| ENSG00002 | gene | 1,65640095 | 0,01163262 | 0,05548295 | COL4A2-AS | 0,72805193 |
| ENSG00000 | gene | 0,92637576 | 0,01164892 | 0,05553399 | PDAP1 | -0,1103306 |
| ENSG00000 | gene | 0,89653168 | 0,01167325 | 0,05560679 | TBC1D22A | -0,1575735 |
| ENSG00000 | gene | 0,79885673 | 0,01167541 | 0,05560679 | PARP14 | -0,3239913 |
| ENSG00002 | gene | 1,13727109 | 0,01172737 | 0,05582747 | AL365203.2 | 0,18557619 |
| ENSG00002 | gene | 1,34625635 | 0,01176143 | 0,05596273 | AC106760.1 | 0,42895315 |
| ENSG00000 | gene | 0,92832361 | 0,0117815 | 0,05603136 | EIF3B | -0,1073003 |
| ENSG00000 | gene | 1,15260078 | 0,01178825 | 0,05603656 | MORC3 | 0,2048929 |
| ENSG00000 | gene | 0,78923857 | 0,01183221 | 0,05621858 | DRG2 | -0,3414666 |
| ENSG00000 | gene | 1,11803954 | 0,01184637 | 0,05625704 | FJX1 | 0,16097121 |
| ENSG00002 | gene | 1,20484892 | 0,01185165 | 0,05625704 | AP001267.1 | 0,26885225 |
| ENSG00000 | gene | 1,17072222 | 0,01191065 | 0,05651003 | NRIP1 | 0,2273988 |
| ENSG00000 | gene | 1,3381731 | 0,01192202 | 0,0565139 | GALNT5 | 0,42026475 |
| ENSG00000 | gene | 0,82178018 | 0,01192286 | 0,0565139 | MRPS16 | -0,2831756 |
| ENSG00002 | gene | 0,87441731 | 0,0119308 | 0,05652453 | RPLP0P6 | -0,1936061 |
| ENSG00000 | gene | 1,39468126 | 0,01199597 | 0,05676314 | FAM86C1 | 0,47993545 |
| ENSG00002 | gene | 1,25673549 | 0,01200104 | 0,05676314 | AC021078.1 | 0,32968103 |
| ENSG00000 | gene | 0,92488532 | 0,01200195 | 0,05676314 | OAF | -0,1126536 |
| ENSG00000 | gene | 1,0586107 | 0,01201115 | 0,05676314 | EFCAB14 | 0,08217214 |
| ENSG00000 | gene | 0,8865511 | 0,01201307 | 0,05676314 | ERICH1 | -0,1737243 |
| ENSG00002 | gene | 0,90719807 | 0,01201792 | 0,05676314 | ITPRIPL2 | -0,1405105 |
| ENSG00000 | gene | 0,73432065 | 0,01202124 | 0,05676314 | MRPL22 | -0,4455179 |
| ENSG00000 | gene | 1,25619116 | 0,01204567 | 0,05685144 | SMURF2 | 0,32905603 |
| ENSG00000 | gene | 1,20891718 | 0,01206074 | 0,0568878 | TMPO | 0,27371541 |
| ENSG00000 | gene | 0,70598313 | 0,01206485 | 0,0568878 | E2F6 | -0,5022944 |
| ENSG00000 | gene | 1,30640981 | 0,01207425 | 0,05690508 | PTPRU | 0,38560753 |
| ENSG00000 | gene | 0,91707216 | 0,01209262 | 0,05695431 | SEC24C | -0,1248928 |
| ENSG00000 | gene | 0,92268062 | 0,01209619 | 0,05695431 | AACS | -0,1160967 |
| ENSG00000 | gene | 0,81798769 | 0,01211181 | 0,05700079 | NFKBIA | -0,289849 |

| | | | | | | |
|---|---|---|---|---|---|---|
| ENSG00000: | gene | 1,20979471 | 0,01214369 | 0,05712371 | WNT5A-AS1 | 0,27476226 |
| ENSG00000( | gene | 0,90486776 | 0,01215939 | 0,05716639 | UBE2A | -0,1442211 |
| ENSG00000 | gene | 0,89043223 | 0,01216997 | 0,05716639 | DONSON | -0,1674223 |
| ENSG00000 | gene | 0,86296178 | 0,01217425 | 0,05716639 | CDYL | -0,2126314 |
| ENSG00000: | gene | 1,68521622 | 0,01217582 | 0,05716639 | AC092069.1 | 0,75293371 |
| ENSG00000: | gene | 1,48782307 | 0,01219235 | 0,05721693 | AC010245.1 | 0,57320297 |
| ENSG00000( | gene | 0,91193772 | 0,01223352 | 0,05736384 | SRI | -0,1329928 |
| ENSG00000( | gene | 1,10987607 | 0,01224068 | 0,05736384 | SNAPC1 | 0,1503986 |
| ENSG00000 | gene | 0,71646741 | 0,01224629 | 0,05736384 | HMG20A | -0,481027 |
| ENSG00000 | gene | 0,86877133 | 0,0122468 | 0,05736384 | DMWD | -0,2029516 |
| ENSG00000: | gene | 1,1436735 | 0,01226681 | 0,05743045 | MYCBP | 0,19367525 |
| ENSG00000 | gene | 0,87234851 | 0,01232209 | 0,05766202 | TRAPPC4 | -0,1970235 |
| ENSG00000 | gene | 0,87348718 | 0,0123407 | 0,05772187 | FASTK | -0,1951416 |
| ENSG00000: | gene | 1,27780356 | 0,01235725 | 0,05776585 | SNORD6 | 0,35366607 |
| ENSG00000 | gene | 1,37978326 | 0,01237386 | 0,05776585 | NPTX1 | 0,46444166 |
| ENSG00000: | gene | 1,60413148 | 0,01237937 | 0,05776585 | KF456478.1 | 0,68179239 |
| ENSG00000: | gene | 0,63435838 | 0,01238002 | 0,05776585 | AC087289.3 | -0,65663 |
| ENSG00000 | gene | 1,41881993 | 0,01238143 | 0,05776585 | SPC24 | 0,5046915 |
| ENSG00000 | gene | 0,92103718 | 0,01238729 | 0,05776585 | DDX49 | -0,1186687 |
| ENSG00000: | gene | 1,43451146 | 0,01239088 | 0,05776585 | ARHGEF28 | 0,52055949 |
| ENSG00000 | gene | 0,72535337 | 0,01244225 | 0,05797806 | SGK1 | -0,4632441 |
| ENSG00000 | gene | 1,14612954 | 0,01245848 | 0,05799488 | TIA1 | 0,19677012 |
| ENSG00000: | gene | 0,93225192 | 0,01246295 | 0,05799488 | NAXD | -0,1012082 |
| ENSG00000: | gene | 0,89824742 | 0,0124634 | 0,05799488 | HMBS | -0,1548152 |
| ENSG00000 | gene | 1,1226116 | 0,0124812 | 0,05802652 | PHKB | 0,16685888 |
| ENSG00000: | gene | 1,13420789 | 0,0124819 | 0,05802652 | CCDC71L | 0,1816851 |
| ENSG00000 | gene | 1,30151134 | 0,01250544 | 0,0581087 | LNPK | 0,38018788 |
| ENSG00000 | gene | 1,12484412 | 0,01251985 | 0,0581484 | PLCB3 | 0,16972508 |
| ENSG00000 | gene | 1,46951537 | 0,01253938 | 0,05819957 | NOL3 | 0,55534044 |
| ENSG00000 | gene | 0,8808061 | 0,01254261 | 0,05819957 | MAN1B1 | -0,1831036 |
| ENSG00000( | gene | 1,11391747 | 0,01255307 | 0,05822089 | HPF1 | 0,15564235 |
| ENSG00000 | gene | 1,1047287 | 0,01256641 | 0,05825549 | NUDT21 | 0,14369212 |
| ENSG00000 | gene | 0,89564519 | 0,01258513 | 0,05826742 | RPL8 | -0,1590008 |
| ENSG00000 | gene | 0,92145064 | 0,01258641 | 0,05826742 | DEAF1 | -0,1180212 |
| ENSG00000( | gene | 0,88931467 | 0,01258661 | 0,05826742 | RDH11 | -0,1692341 |
| ENSG00000 | gene | 0,96307833 | 0,01260171 | 0,05829372 | TPD52L2 | -0,054275 |
| ENSG00000: | gene | 1,34641138 | 0,01260405 | 0,05829372 | AC130324.3 | 0,42911927 |
| ENSG00000: | gene | 1,30436065 | 0,01263553 | 0,05840629 | AC091564.4 | 0,38334283 |
| ENSG00000 | gene | 1,40118014 | 0,01264017 | 0,05840629 | SCLT1 | 0,48664244 |
| ENSG00000( | gene | 0,90498558 | 0,01267826 | 0,05853295 | NOP14 | -0,1440333 |
| ENSG00000( | gene | 0,8408147 | 0,01268508 | 0,05853295 | CRLF1 | -0,2501402 |
| ENSG00000 | gene | 1,13413649 | 0,01269106 | 0,05853295 | EIF1AX | 0,18159428 |
| ENSG00000 | gene | 1,08679438 | 0,01269119 | 0,05853295 | SCRN1 | 0,12007901 |
| ENSG00000 | gene | 0,89607895 | 0,01276136 | 0,05882921 | SURF2 | -0,1583023 |
| ENSG00000( | gene | 0,87929739 | 0,01277822 | 0,05886737 | EPN2 | -0,1855769 |
| ENSG00000 | gene | 0,90796566 | 0,01278151 | 0,05886737 | RAE1 | -0,1392904 |
| ENSG00000 | gene | 0,87748371 | 0,01279165 | 0,05887691 | ATP5MC2 | -0,1885558 |
| ENSG00000 | gene | 0,9353997 | 0,01279893 | 0,05887691 | CDC42BPB | -0,0963451 |
| ENSG00000 | gene | 0,92108235 | 0,01280139 | 0,05887691 | ARFGAP2 | -0,1185979 |
| ENSG00000: | gene | 1,60362406 | 0,01281862 | 0,05892882 | AC004477.2 | 0,68133597 |
| ENSG00000: | gene | 1,23719161 | 0,01284914 | 0,05904172 | AC139887.2 | 0,30706896 |
| ENSG00000( | gene | 1,09249058 | 0,01285669 | 0,05904905 | EXOC1 | 0,12762084 |
| ENSG00000( | gene | 1,09690654 | 0,01287004 | 0,05908304 | NOP58 | 0,13344061 |
| ENSG00000 | gene | 0,89374833 | 0,01288356 | 0,05909072 | GLUL | -0,1620595 |
| ENSG00000 | gene | 0,89899351 | 0,01288364 | 0,05909072 | GPR108 | -0,1536174 |
| ENSG00000: | gene | 1,54312384 | 0,01289978 | 0,05912557 | AL358790.1 | 0,62585385 |
| ENSG00000: | gene | 0,77867172 | 0,01290643 | 0,05912557 | LIX1L-AS1 | -0,3609129 |

| | | | | | | |
|---|---|---|---|---|---|---|
| ENSG00002 | gene | 0,84329638 | 0,01290912 | 0,05912557 | HNRNPCP2 | -0,2458883 |
| ENSG00002 | gene | 1,24074275 | 0,01293508 | 0,05917121 | AC010735.1 | 0,31120402 |
| ENSG00002 | gene | 0,86454611 | 0,01293769 | 0,05917121 | RPL41 | -0,2099852 |
| ENSG00001 | gene | 0,89689714 | 0,01294194 | 0,05917121 | MRPL52 | -0,1569856 |
| ENSG00002 | gene | 1,4771339 | 0,01294296 | 0,05917121 | AC010976.1 | 0,56280061 |
| ENSG00001 | gene | 0,87384407 | 0,01296046 | 0,05922391 | FNTA | -0,1945522 |
| ENSG00002 | gene | 1,28912072 | 0,01297767 | 0,05927527 | MIR1282 | 0,36638737 |
| ENSG00001 | gene | 0,90339353 | 0,01303315 | 0,05950124 | PFN1 | -0,1465735 |
| ENSG00001 | gene | 0,90381415 | 0,01306964 | 0,05964037 | WDR45B | -0,1459019 |
| ENSG00001 | gene | 1,21135423 | 0,01308056 | 0,05966274 | ZEB2 | 0,27662081 |
| ENSG00001 | gene | 0,85662889 | 0,01309623 | 0,05970678 | UXT | -0,2232578 |
| ENSG00001 | gene | 0,87961737 | 0,01310501 | 0,05971932 | IFT27 | -0,185052 |
| ENSG00002 | gene | 0,82439591 | 0,01311343 | 0,05973025 | C5orf66 | -0,2785907 |
| ENSG00001 | gene | 0,69639397 | 0,01313961 | 0,0598058 | DDIT4 | -0,5220244 |
| ENSG00000 | gene | 1,17431363 | 0,01314208 | 0,0598058 | THAP3 | 0,23181777 |
| ENSG00001 | gene | 0,91803049 | 0,01318005 | 0,05992928 | HSPBP1 | -0,123386 |
| ENSG00001 | gene | 1,13537019 | 0,0131813 | 0,05992928 | ZNF22 | 0,18316277 |
| ENSG00001 | gene | 1,27294625 | 0,01320552 | 0,06001188 | TYW1 | 0,34817151 |
| ENSG00002 | gene | 1,17588047 | 0,01323417 | 0,0601098 | AL391056.2 | 0,23374141 |
| ENSG00001 | gene | 0,86551991 | 0,01324235 | 0,0601098 | RELA | -0,2083611 |
| ENSG00001 | gene | 0,89995783 | 0,01325009 | 0,0601098 | CNPY4 | -0,1520707 |
| ENSG00002 | gene | 1,43936013 | 0,01325131 | 0,0601098 | AC006449.9 | 0,52542761 |
| ENSG00001 | gene | 1,21142541 | 0,01326912 | 0,06016303 | NIN | 0,27670558 |
| ENSG00002 | gene | 0,7372757 | 0,01327855 | 0,06017827 | AC016739.1 | -0,4397239 |
| ENSG00001 | gene | 1,11408556 | 0,01330696 | 0,06027951 | MYOF | 0,15586003 |
| ENSG00001 | gene | 0,88364909 | 0,01332434 | 0,06032506 | BRD4 | -0,1784545 |
| ENSG00002 | gene | 1,28017393 | 0,01332919 | 0,06032506 | ADAMTSL4- | 0,35633984 |
| ENSG00002 | gene | 0,80741141 | 0,0133416 | 0,06035369 | HGH1 | -0,3086241 |
| ENSG00001 | gene | 1,15774269 | 0,01337565 | 0,06048016 | EDIL3 | 0,21131465 |
| ENSG00001 | gene | 0,92143368 | 0,01341023 | 0,06060887 | CSK | -0,1180478 |
| ENSG00001 | gene | 0,86634209 | 0,0134279 | 0,06066107 | TCEA2 | -0,2069913 |
| ENSG00001 | gene | 0,90426073 | 0,0134812 | 0,06087415 | ATP6V0D1 | -0,1451893 |
| ENSG00002 | gene | 0,74081626 | 0,01352824 | 0,06105871 | SVIL-AS1 | -0,4328123 |
| ENSG00002 | gene | 1,68275998 | 0,01363105 | 0,06149475 | AC087393.2 | 0,75082941 |
| ENSG00002 | gene | 0,8801375 | 0,01365427 | 0,06156528 | AC006001.3 | -0,1841992 |
| ENSG00002 | gene | 1,06430199 | 0,0136591 | 0,06156528 | MTND2P28 | 0,08990757 |
| ENSG00001 | gene | 1,27814571 | 0,01369617 | 0,06170434 | RASGRF2 | 0,35405232 |
| ENSG00001 | gene | 0,89895738 | 0,01370367 | 0,0617101 | CFAP36 | -0,1536754 |
| ENSG00001 | gene | 0,88225576 | 0,01371493 | 0,06171147 | GIT1 | -0,1807311 |
| ENSG00002 | gene | 0,89364203 | 0,01371642 | 0,06171147 | AC092368.3 | -0,1622311 |
| ENSG00001 | gene | 1,21032229 | 0,01374433 | 0,06180897 | ZMAT3 | 0,27539126 |
| ENSG00001 | gene | 0,91358154 | 0,01376453 | 0,06187177 | SCYL1 | -0,1303946 |
| ENSG00002 | gene | 1,22605045 | 0,01377639 | 0,06189701 | LINC00106 | 0,29401835 |
| ENSG00002 | gene | 1,70069539 | 0,01379185 | 0,06193839 | AL158151.1 | 0,76612477 |
| ENSG00001 | gene | 1,10269498 | 0,01380975 | 0,06199069 | HPRT1 | 0,14103377 |
| ENSG00001 | gene | 0,85753486 | 0,01383252 | 0,06206482 | E4F1 | -0,2217328 |
| ENSG00001 | gene | 0,89884851 | 0,01384997 | 0,06208462 | MARS1 | -0,1538501 |
| ENSG00001 | gene | 0,92770639 | 0,01385328 | 0,06208462 | B4GALT2 | -0,1082598 |
| ENSG00002 | gene | 1,21380682 | 0,01385571 | 0,06208462 | AL031282.1 | 0,27953883 |
| ENSG00001 | gene | 1,08116213 | 0,01386759 | 0,06210976 | NETO2 | 0,11258289 |
| ENSG00001 | gene | 0,86641794 | 0,01388658 | 0,06215325 | CARHSP1 | -0,206865 |
| ENSG00000 | gene | 0,94271515 | 0,01388984 | 0,06215325 | RANBP3 | -0,0851062 |
| ENSG00000 | gene | 0,91285152 | 0,01391693 | 0,0622464 | HSD17B10 | -0,1315479 |
| ENSG00001 | gene | 0,87062383 | 0,01393457 | 0,0622972 | BCL2L1 | -0,1998786 |
| ENSG00001 | gene | 1,13221642 | 0,01394915 | 0,06233428 | ZDHHC2 | 0,17914975 |
| ENSG00001 | gene | 0,81071516 | 0,01395869 | 0,06234883 | CLIP2 | -0,302733 |
| ENSG00002 | gene | 0,93319003 | 0,01397816 | 0,06238801 | ZBTB22 | -0,0997572 |

| | | | | | | |
|---|---|---|---|---|---|---|
| ENSG00000( | gene | 0,85852192 | 0,01398005 | 0,06238801 | CRAMP1 | -0,2200731 |
| ENSG00000 | gene | 0,77498296 | 0,01399651 | 0,06243337 | LDLRAP1 | -0,3677635 |
| ENSG00000 | gene | 0,76838703 | 0,01401317 | 0,06246107 | RPL17 | -0,3800949 |
| ENSG00000 | gene | 0,90473298 | 0,0140196 | 0,06246107 | ACP1 | -0,144436 |
| ENSG00000( | gene | 0,94098177 | 0,01402162 | 0,06246107 | EIF3I | -0,0877613 |
| ENSG00000 | gene | 0,94115629 | 0,01405563 | 0,06258031 | RBM17 | -0,0874938 |
| ENSG00000 | gene | 1,19631458 | 0,014061 | 0,06258031 | CAPZA2 | 0,25859681 |
| ENSG00000 | gene | 1,06687587 | 0,01408039 | 0,06263846 | LONP2 | 0,09339232 |
| ENSG00000 | gene | 1,30716074 | 0,01409185 | 0,06266135 | AC135457.1 | 0,38643655 |
| ENSG00000 | gene | 0,82440052 | 0,01410874 | 0,06270832 | C1R | -0,2785827 |
| ENSG00000 | gene | 0,64581726 | 0,0141195 | 0,06272801 | ATG10 | -0,6308021 |
| ENSG00000 | gene | 0,93916633 | 0,01413642 | 0,06275983 | EEF2 | -0,0905474 |
| ENSG00000 | gene | 0,86915133 | 0,01413932 | 0,06275983 | UBA7 | -0,2023207 |
| ENSG00000 | gene | 1,38064404 | 0,01415664 | 0,06280859 | TMSB4XP1 | 0,46534141 |
| ENSG00000 | gene | 1,12737155 | 0,01417612 | 0,06286692 | PRRC2C | 0,17296306 |
| ENSG00000 | gene | 1,054457 | 0,01418869 | 0,06289452 | CTNNB1 | 0,07650027 |
| ENSG00000 | gene | 1,16010951 | 0,01421116 | 0,06291384 | CDH6 | 0,214261 |
| ENSG00000 | gene | 0,94464781 | 0,01422295 | 0,06291384 | SAMM50 | -0,0821515 |
| ENSG00000 | gene | 1,36950057 | 0,01422698 | 0,06291384 | LEMD3 | 0,45364987 |
| ENSG00000( | gene | 0,91907192 | 0,01422876 | 0,06291384 | FTSJ1 | -0,1217503 |
| ENSG00000 | gene | 0,82727019 | 0,01423069 | 0,06291384 | OSGIN1 | -0,2735695 |
| ENSG00000( | gene | 0,83021175 | 0,01423161 | 0,06291384 | EPB41L1 | -0,2684487 |
| ENSG00000 | gene | 0,87563736 | 0,01423746 | 0,06291384 | G6PD | -0,1915946 |
| ENSG00000 | gene | 1,44644957 | 0,0142577 | 0,0629274 | AMBRA1 | 0,53251602 |
| ENSG00000 | gene | 1,13387987 | 0,01426905 | 0,0629274 | FLNC | 0,1812678 |
| ENSG00000 | gene | 1,14517167 | 0,01426968 | 0,0629274 | LARP7 | 0,19556389 |
| ENSG00000 | gene | 0,89219003 | 0,01427181 | 0,0629274 | HEIH | -0,1645771 |
| ENSG00000 | gene | 1,18690138 | 0,01427226 | 0,0629274 | SPRN | 0,24720006 |
| ENSG00000 | gene | 1,17215278 | 0,01430281 | 0,06303406 | DEK | 0,22916062 |
| ENSG00000 | gene | 1,05961984 | 0,01431746 | 0,06306498 | DYNC1H1 | 0,08354677 |
| ENSG00000 | gene | 1,19871426 | 0,01432254 | 0,06306498 | KISS1 | 0,2614878 |
| ENSG00000 | gene | 1,109443 | 0,01437675 | 0,06327555 | RCN2 | 0,14983555 |
| ENSG00000 | gene | 0,92710053 | 0,01438405 | 0,06327963 | GLRX5 | -0,1092023 |
| ENSG00000 | gene | 1,28001825 | 0,01440853 | 0,06334859 | FGF5 | 0,35616438 |
| ENSG00000 | gene | 1,16395242 | 0,01441251 | 0,06334859 | GNG10 | 0,21903208 |
| ENSG00000 | gene | 0,86920287 | 0,01442559 | 0,06336018 | RPS28P7 | -0,2022352 |
| ENSG00000 | gene | 0,92031322 | 0,01442792 | 0,06336018 | DUSP11 | -0,1198032 |
| ENSG00000 | gene | 0,80939294 | 0,01444654 | 0,063391 | C19orf25 | -0,3050878 |
| ENSG00000 | gene | 0,91285687 | 0,01444773 | 0,063391 | DPY30 | -0,1315394 |
| ENSG00000 | gene | 0,92059816 | 0,01446823 | 0,0634529 | PSMB3 | -0,1193565 |
| ENSG00000( | gene | 0,76434923 | 0,01447789 | 0,0634672 | MKNK2 | -0,3876961 |
| ENSG00000 | gene | 1,44467483 | 0,01450073 | 0,06351621 | MIR6126 | 0,53074481 |
| ENSG00000 | gene | 0,77606757 | 0,01450189 | 0,06351621 | NSUN4 | -0,3657458 |
| ENSG00000 | gene | 1,12926635 | 0,01453919 | 0,06365146 | AL671710.1 | 0,17538581 |
| ENSG00000( | gene | 1,33403628 | 0,01455649 | 0,06369911 | SLC9A7 | 0,4157979 |
| ENSG00000 | gene | 0,75126099 | 0,01459535 | 0,06384098 | MAFG | -0,4126139 |
| ENSG00000( | gene | 0,86856703 | 0,01461637 | 0,06390473 | REXO2 | -0,2032909 |
| ENSG00000( | gene | 0,89288302 | 0,0146273 | 0,06392434 | TMED1 | -0,1634569 |
| ENSG00000 | gene | 0,88797343 | 0,01464935 | 0,0639925 | CCDC174 | -0,1714116 |
| ENSG00000 | gene | 0,83696608 | 0,01466103 | 0,06401531 | SDF2 | -0,2567589 |
| ENSG00000 | gene | 0,76070021 | 0,01467678 | 0,06403891 | FBLIM1 | -0,3946001 |
| ENSG00000 | gene | 0,8765056 | 0,01467935 | 0,06403891 | TRMT1 | -0,1901648 |
| ENSG00000( | gene | 0,87325475 | 0,01469596 | 0,06408316 | PGS1 | -0,1955255 |
| ENSG00000( | gene | 0,93353402 | 0,01471057 | 0,06411871 | SF3B2 | -0,0992255 |
| ENSG00000 | gene | 1,10005968 | 0,0147179 | 0,06412243 | PDE4DIP | 0,13758179 |
| ENSG00000 | gene | 0,86144814 | 0,01472553 | 0,06412752 | COQ4 | -0,2151641 |
| ENSG00000 | gene | 0,61134661 | 0,0147393 | 0,06413779 | RPPH1 | -0,7099375 |

| | | | | | | |
|---|---|---|---|---|---|---|
| ENSG00002 | gene | 1,28280616 | 0,01474083 | 0,06413779 | AC012615.3 | 0,35930319 |
| ENSG00000 | gene | 1,24804862 | 0,01477151 | 0,06424309 | ITGA6 | 0,31967414 |
| ENSG00000 | gene | 1,06174141 | 0,01480205 | 0,06434773 | TSPYL1 | 0,08643244 |
| ENSG00002 | gene | 0,81857577 | 0,01482261 | 0,06440886 | RPS29 | -0,2888121 |
| ENSG00000 | gene | 1,09780899 | 0,01485621 | 0,0644705 | NCKAP1 | 0,13462705 |
| ENSG00000 | gene | 0,93665557 | 0,01485981 | 0,0644705 | CCNY | -0,0944095 |
| ENSG00000 | gene | 1,1523588 | 0,01486067 | 0,0644705 | UBL3 | 0,20458998 |
| ENSG00000 | gene | 0,90317953 | 0,0148628 | 0,0644705 | PPP6R1 | -0,1469153 |
| ENSG00000 | gene | 0,80814034 | 0,01490331 | 0,06461716 | RPL30 | -0,3073222 |
| ENSG00000 | gene | 1,11523509 | 0,01491158 | 0,06461716 | RAP1B | 0,15734786 |
| ENSG00000 | gene | 0,68214322 | 0,01491616 | 0,06461716 | ARSG | -0,5518534 |
| ENSG00000 | gene | 1,09696222 | 0,01494577 | 0,06471714 | TLN1 | 0,13351384 |
| ENSG00000 | gene | 0,83954279 | 0,01497013 | 0,06475636 | YIF1B | -0,2523242 |
| ENSG00000 | gene | 1,15611896 | 0,01498478 | 0,06475636 | GTF2A1 | 0,20928986 |
| ENSG00000 | gene | 0,7899862 | 0,01498797 | 0,06475636 | GGA3 | -0,3401006 |
| ENSG00000 | gene | 0,85608499 | 0,01498888 | 0,06475636 | ARRDC1 | -0,2241741 |
| ENSG00000 | gene | 0,88948732 | 0,01499017 | 0,06475636 | LONP1 | -0,1689541 |
| ENSG00000 | gene | 0,87270634 | 0,01499401 | 0,06475636 | FAU | -0,1964318 |
| ENSG00000 | gene | 0,77523708 | 0,01500489 | 0,06477514 | NDUFA13 | -0,3672905 |
| ENSG00000 | gene | 1,12998128 | 0,01503171 | 0,06486266 | MGAT2 | 0,17629887 |
| ENSG00000 | gene | 0,89268224 | 0,01504337 | 0,06488474 | CCDC28B | -0,1637814 |
| ENSG00000 | gene | 0,76731711 | 0,0150591 | 0,0649034 | ARL10 | -0,3821052 |
| ENSG00000 | gene | 1,38786845 | 0,01506078 | 0,0649034 | ARID1B | 0,47287082 |
| ENSG00002 | gene | 0,67483698 | 0,01507836 | 0,06492554 | AC009690.2 | -0,5673891 |
| ENSG00000 | gene | 1,09882353 | 0,01508014 | 0,06492554 | LAMC1 | 0,1359597 |
| ENSG00000 | gene | 0,81544981 | 0,01508556 | 0,06492554 | HS1BP3 | -0,294332 |
| ENSG00000 | gene | 1,09502531 | 0,01509757 | 0,06494904 | STK39 | 0,13096422 |
| ENSG00000 | gene | 0,84629761 | 0,01511688 | 0,06500388 | CYB561D2 | -0,240763 |
| ENSG00002 | gene | 0,86287624 | 0,01515017 | 0,0651188 | DDR1 | -0,2127744 |
| ENSG00000 | gene | 1,32458612 | 0,01518359 | 0,06523417 | SLC14A1 | 0,40554164 |
| ENSG00000 | gene | 0,9210646 | 0,01523386 | 0,06541507 | COPS3 | -0,1186258 |
| ENSG00000 | gene | 0,92034645 | 0,01524112 | 0,06541507 | DHX38 | -0,1197511 |
| ENSG00000 | gene | 0,89328498 | 0,01524548 | 0,06541507 | RPSA | -0,1628076 |
| ENSG00000 | gene | 1,07716023 | 0,01525921 | 0,06544563 | ARGLU1 | 0,10723287 |
| ENSG00002 | gene | 0,82021993 | 0,01527817 | 0,06547758 | RPS10P3 | -0,2859173 |
| ENSG00000 | gene | 0,87656962 | 0,01527986 | 0,06547758 | ATRAID | -0,1900594 |
| ENSG00000 | gene | 0,91404696 | 0,0153077 | 0,06556852 | ALYREF | -0,1296598 |
| ENSG00002 | gene | 1,16785678 | 0,01531465 | 0,06556997 | NEXN-AS1 | 0,22386336 |
| ENSG00002 | gene | 0,72316203 | 0,01533849 | 0,06561904 | AP001267.5 | -0,4676092 |
| ENSG00000 | gene | 0,9075658 | 0,01534442 | 0,06561904 | DHRSX | -0,1399258 |
| ENSG00000 | gene | 1,07276056 | 0,01534596 | 0,06561904 | TIPRL | 0,1013281 |
| ENSG00000 | gene | 0,75150675 | 0,01535985 | 0,06563953 | KHNYN | -0,412142 |
| ENSG00000 | gene | 0,93199648 | 0,01536399 | 0,06563953 | EXOC7 | -0,1016036 |
| ENSG00000 | gene | 0,75872756 | 0,01537701 | 0,06566684 | SNAP23 | -0,3983462 |
| ENSG00000 | gene | 0,85055448 | 0,01539507 | 0,06571567 | GATD1 | -0,2335245 |
| ENSG00002 | gene | 0,84047066 | 0,01543068 | 0,06583934 | AC010422.8 | -0,2507306 |
| ENSG00000 | gene | 0,89923548 | 0,01546026 | 0,06591918 | DPYSL2 | -0,1532291 |
| ENSG00000 | gene | 1,07631931 | 0,01546269 | 0,06591918 | DHX9 | 0,10610615 |
| ENSG00000 | gene | 0,96098186 | 0,01553202 | 0,06616231 | PELO | -0,0574189 |
| ENSG00000 | gene | 0,91421012 | 0,01553306 | 0,06616231 | ATP2A2 | -0,1294023 |
| ENSG00000 | gene | 0,92153403 | 0,01555101 | 0,0662103 | METTL17 | -0,1178907 |
| ENSG00000 | gene | 0,80830062 | 0,01556596 | 0,06624554 | TSSK3 | -0,3070361 |
| ENSG00000 | gene | 1,2184267 | 0,0155789 | 0,06624582 | CDK1 | 0,28501946 |
| ENSG00000 | gene | 0,93117516 | 0,01557939 | 0,06624582 | ABL1 | -0,1028755 |
| ENSG00000 | gene | 0,88885528 | 0,01559485 | 0,06628313 | BCL6 | -0,1699796 |
| ENSG00000 | gene | 0,72293698 | 0,01561152 | 0,06632556 | LIF | -0,4680582 |
| ENSG00000 | gene | 0,86690104 | 0,01565172 | 0,06644035 | HARS2 | -0,2060608 |

| | | | | | | |
|---|---|---|---|---|---|---|
| ENSG00002 | gene | 0,78140675 | 0,01565194 | 0,06644035 | RPL23AP42 | -0,3558544 |
| ENSG00000 | gene | 0,89528808 | 0,01566916 | 0,06646002 | SRP68 | -0,1595761 |
| ENSG00002 | gene | 1,1951078 | 0,01566998 | 0,06646002 | ARSD-AS1 | 0,25714076 |
| ENSG00000 | gene | 0,8800891 | 0,01568054 | 0,06646925 | SLC25A38 | -0,1842785 |
| ENSG00002 | gene | 0,66353211 | 0,01568556 | 0,06646925 | AC007229.1 | -0,5917618 |
| ENSG00000 | gene | 1,10707157 | 0,01575247 | 0,06671268 | CWC27 | 0,1467485 |
| ENSG00000 | gene | 0,86316231 | 0,01575646 | 0,06671268 | RPL27A | -0,2122962 |
| ENSG00002 | gene | 0,87167966 | 0,01576393 | 0,06671579 | ARHGEF25 | -0,1981301 |
| ENSG00002 | gene | 1,43762872 | 0,01577682 | 0,06674187 | AC005363.1 | 0,52369113 |
| ENSG00000 | gene | 0,8866643 | 0,01580317 | 0,06682485 | RBCK1 | -0,1735401 |
| ENSG00000 | gene | 1,12288552 | 0,01587628 | 0,06710535 | QSER1 | 0,16721085 |
| ENSG00002 | gene | 1,33339841 | 0,01593094 | 0,06728546 | AC090984.1 | 0,41510791 |
| ENSG00000 | gene | 1,05504865 | 0,01593856 | 0,06728546 | COPB2 | 0,07730953 |
| ENSG00000 | gene | 0,85048512 | 0,01594076 | 0,06728546 | ZFAND6 | -0,2336421 |
| ENSG00002 | gene | 1,51085241 | 0,01594603 | 0,06728546 | APOM | 0,59536274 |
| ENSG00000 | gene | 0,8447354 | 0,01596921 | 0,0673546 | ATP5MPL | -0,2434286 |
| ENSG00000 | gene | 0,78427748 | 0,01597663 | 0,06735724 | CD44 | -0,3505639 |
| ENSG00000 | gene | 0,77962562 | 0,01603457 | 0,0675728 | GPT2 | -0,3591466 |
| ENSG00000 | gene | 0,87382549 | 0,01604422 | 0,06758473 | GSTM3 | -0,1945829 |
| ENSG00002 | gene | 1,31603686 | 0,01606738 | 0,06765358 | AC005306.1 | 0,3961999 |
| ENSG00000 | gene | 0,57651402 | 0,01617041 | 0,06805335 | PTGS2 | -0,7945724 |
| ENSG00000 | gene | 0,91400766 | 0,01617605 | 0,06805335 | NUDT9 | -0,1297218 |
| ENSG00000 | gene | 1,16412542 | 0,01620373 | 0,06814086 | FANCL | 0,2192465 |
| ENSG00000 | gene | 0,93579124 | 0,01621816 | 0,06817263 | MT-ND1 | -0,0957414 |
| ENSG00000 | gene | 0,91727296 | 0,01624366 | 0,06825088 | TUBG2 | -0,124577 |
| ENSG00002 | gene | 1,23198051 | 0,01630247 | 0,06841836 | BACE1-AS | 0,30097943 |
| ENSG00000 | gene | 0,91185617 | 0,01630267 | 0,06841836 | INPP5K | -0,1331218 |
| ENSG00002 | gene | 0,81203077 | 0,01630421 | 0,06841836 | ST20-MTHFS | -0,3003937 |
| ENSG00000 | gene | 0,92764519 | 0,01635001 | 0,06858152 | FGD1 | -0,108355 |
| ENSG00002 | gene | 0,74031751 | 0,01637207 | 0,06864502 | TUBB2BP1 | -0,4337839 |
| ENSG00000 | gene | 0,86020539 | 0,0163892 | 0,06866339 | C19orf48 | -0,2172469 |
| ENSG00000 | gene | 1,07660135 | 0,01639665 | 0,06866339 | SRRM1 | 0,10648414 |
| ENSG00000 | gene | 0,91953106 | 0,01639723 | 0,06866339 | EEF1AKNMT | -0,1210298 |
| ENSG00000 | gene | 0,89586991 | 0,01641217 | 0,06869693 | SLC16A3 | -0,1586388 |
| ENSG00000 | gene | 0,92098442 | 0,01643167 | 0,06874955 | TSPAN9 | -0,1187513 |
| ENSG00002 | gene | 1,21658117 | 0,01644953 | 0,06879526 | AC087289.4 | 0,28283259 |
| ENSG00002 | gene | 1,1897323 | 0,01648961 | 0,06891759 | AL589765.7 | 0,25063699 |
| ENSG00000 | gene | 0,92808595 | 0,01649301 | 0,06891759 | PSMA2 | -0,1076697 |
| ENSG00000 | gene | 0,8756402 | 0,01649963 | 0,06891759 | RGS3 | -0,1915899 |
| ENSG00000 | gene | 0,82881323 | 0,01652182 | 0,06897183 | HDHD5 | -0,2708811 |
| ENSG00000 | gene | 0,88733027 | 0,01652653 | 0,06897183 | CIDECP1 | -0,1724569 |
| ENSG00000 | gene | 0,83263044 | 0,01653396 | 0,06897381 | VPS26C | -0,2642518 |
| ENSG00000 | gene | 1,15603794 | 0,01655252 | 0,06900806 | RAB8B | 0,20918875 |
| ENSG00002 | gene | 1,34449862 | 0,01655609 | 0,06900806 | AC104411.1 | 0,42706827 |
| ENSG00000 | gene | 0,88532371 | 0,01657075 | 0,0690152 | MAP2K3 | -0,175723 |
| ENSG00000 | gene | 0,89539743 | 0,01657172 | 0,0690152 | NTAN1 | -0,1593999 |
| ENSG00000 | gene | 0,76483223 | 0,0165903 | 0,06901762 | ST6GALNAC | -0,3867848 |
| ENSG00000 | gene | 0,73202826 | 0,01659122 | 0,06901762 | C15orf48 | -0,4500288 |
| ENSG00000 | gene | 0,95270088 | 0,01659318 | 0,06901762 | AHCY | -0,0699048 |
| ENSG00000 | gene | 0,89965711 | 0,01660785 | 0,06904968 | MEAF6 | -0,1525529 |
| ENSG00000 | gene | 0,91997092 | 0,01662008 | 0,06906598 | ITPK1 | -0,1203398 |
| ENSG00000 | gene | 0,91094953 | 0,01662571 | 0,06906598 | DHRS7B | -0,134557 |
| ENSG00000 | gene | 1,11053307 | 0,01667358 | 0,06923584 | MBIP | 0,15125236 |
| ENSG00000 | gene | 0,93696887 | 0,01672939 | 0,06943852 | EXOSC10 | -0,093927 |
| ENSG00000 | gene | 0,89282524 | 0,01674281 | 0,06946513 | KLC4 | -0,1635503 |
| ENSG00002 | gene | 0,64393605 | 0,01689213 | 0,07005536 | RNASEK | -0,6350107 |
| ENSG00002 | gene | 1,28354002 | 0,01693506 | 0,07020255 | AL662797.2 | 0,36012828 |

| | | | | | | |
|---|---|---|---|---|---|---|
| ENSG00000 | gene | 0,81271087 | 0,01694178 | 0,07020255 | ATP5ME | -0,2991859 |
| ENSG00002 | gene | 1,51568262 | 0,01695517 | 0,07022868 | AC004079.5 | 0,59996769 |
| ENSG00000 | gene | 0,91108161 | 0,01701566 | 0,0704498 | RCN3 | -0,1343478 |
| ENSG00002 | gene | 1,8111816 | 0,0170435 | 0,07047588 | AL034430.2 | 0,85693121 |
| ENSG00000 | gene | 1,11139968 | 0,01704499 | 0,07047588 | WBP4 | 0,15237773 |
| ENSG00000 | gene | 0,84609676 | 0,01704848 | 0,07047588 | H1-10 | -0,2411054 |
| ENSG00000 | gene | 0,89513186 | 0,01705039 | 0,07047588 | GPR137 | -0,1598279 |
| ENSG00000 | gene | 1,10109114 | 0,01708517 | 0,07059021 | MINDY3 | 0,13893389 |
| ENSG00000 | gene | 1,08378446 | 0,01710845 | 0,07065698 | PINK1-AS | 0,11607786 |
| ENSG00000 | gene | 0,92308225 | 0,01713309 | 0,07072927 | ACADVL | -0,1154689 |
| ENSG00000 | gene | 0,90915863 | 0,01715981 | 0,07077342 | NDUFAF1 | -0,1373961 |
| ENSG00000 | gene | 0,91593376 | 0,01716514 | 0,07077342 | BRI3 | -0,1266848 |
| ENSG00000 | gene | 0,7460871 | 0,01716519 | 0,07077342 | WHRN | -0,422584 |
| ENSG00000 | gene | 0,89829617 | 0,01717746 | 0,07079454 | ELP6 | -0,1547369 |
| ENSG00000 | gene | 1,20257047 | 0,01719438 | 0,07082034 | GMFB | 0,26612144 |
| ENSG00000 | gene | 0,85098612 | 0,01720473 | 0,07082034 | SYVN1 | -0,2327925 |
| ENSG00000 | gene | 0,91399694 | 0,01720514 | 0,07082034 | PPAN | -0,1297388 |
| ENSG00000 | gene | 0,92184362 | 0,01729043 | 0,07113255 | ANXA7 | -0,1174061 |
| ENSG00002 | gene | 2,41186295 | 0,01729534 | 0,07113255 | AL133346.1 | 1,27014793 |
| ENSG00002 | gene | 0,58949213 | 0,0173049 | 0,07114238 | AC068533.4 | -0,7624555 |
| ENSG00000 | gene | 0,88271773 | 0,01734056 | 0,07125942 | TMSB10 | -0,1799759 |
| ENSG00000 | gene | 0,88119656 | 0,01736822 | 0,07133005 | HAGH | -0,1824642 |
| ENSG00000 | gene | 0,94625978 | 0,01737659 | 0,07133005 | ANXA4 | -0,0796918 |
| ENSG00000 | gene | 0,81271303 | 0,01738585 | 0,07133005 | JMJD8 | -0,2991821 |
| ENSG00000 | gene | 0,84205715 | 0,01738652 | 0,07133005 | CRTC2 | -0,2480099 |
| ENSG00000 | gene | 0,87076726 | 0,01741156 | 0,07140324 | MBD1 | -0,1996409 |
| ENSG00000 | gene | 1,23246047 | 0,01748636 | 0,07168033 | KRR1 | 0,30154138 |
| ENSG00000 | gene | 0,91616942 | 0,0175133 | 0,07176113 | MBD6 | -0,1263137 |
| ENSG00000 | gene | 1,23904955 | 0,01752684 | 0,07178692 | TNFRSF10D | 0,30923388 |
| ENSG00002 | gene | 1,4376482 | 0,01753888 | 0,07180657 | KRTAP2-3 | 0,52371068 |
| ENSG00000 | gene | 0,90128266 | 0,01756499 | 0,07188378 | PTMA | -0,1499485 |
| ENSG00002 | gene | 1,3165858 | 0,01757831 | 0,07190864 | TRIM59 | 0,39680154 |
| ENSG00000 | gene | 1,12661683 | 0,01759 | 0,07192677 | ZDHHC17 | 0,17199693 |
| ENSG00000 | gene | 0,87257246 | 0,01760313 | 0,0719508 | URM1 | -0,1966532 |
| ENSG00000 | gene | 0,95268692 | 0,01768496 | 0,07225548 | RTCB | -0,0699259 |
| ENSG00002 | gene | 0,85106685 | 0,01770111 | 0,07229167 | FNTB | -0,2326556 |
| ENSG00000 | gene | 0,88692136 | 0,01772205 | 0,07234741 | TEPSIN | -0,1731219 |
| ENSG00000 | gene | 0,86587586 | 0,01773297 | 0,07236221 | UQCC2 | -0,2077679 |
| ENSG00000 | gene | 0,89045544 | 0,01777988 | 0,07251259 | DMPK | -0,1673847 |
| ENSG00000 | gene | 1,10043133 | 0,01778445 | 0,07251259 | CKAP5 | 0,13806912 |
| ENSG00000 | gene | 1,09716229 | 0,01782291 | 0,07263951 | PAN3 | 0,13377694 |
| ENSG00000 | gene | 1,1507543 | 0,01784349 | 0,07269353 | MIB1 | 0,20257984 |
| ENSG00000 | gene | 0,90467654 | 0,01788494 | 0,07283249 | RNF40 | -0,144526 |
| ENSG00000 | gene | 0,80804528 | 0,01807075 | 0,07355894 | SLC9A3R1 | -0,307492 |
| ENSG00002 | gene | 1,3310642 | 0,01808496 | 0,07357742 | AC080038.4 | 0,41258016 |
| ENSG00000 | gene | 1,1793635 | 0,0181008 | 0,07357742 | COL12A1 | 0,23800845 |
| ENSG00000 | gene | 1,17633205 | 0,01810082 | 0,07357742 | CYB5B | 0,23429536 |
| ENSG00000 | gene | 0,93290478 | 0,01810497 | 0,07357742 | METTL3 | -0,1001983 |
| ENSG00000 | gene | 1,09862171 | 0,01811389 | 0,0735835 | SDCCAG8 | 0,13569471 |
| ENSG00002 | gene | 1,24392714 | 0,01812558 | 0,07360085 | AL603839.1 | 0,31490199 |
| ENSG00000 | gene | 1,09041154 | 0,01813891 | 0,07362483 | BCCIP | 0,12487274 |
| ENSG00002 | gene | 1,36585498 | 0,01817681 | 0,07374847 | AL020996.1 | 0,44980431 |
| ENSG00000 | gene | 0,90926066 | 0,01821841 | 0,07388703 | PSMB5 | -0,1372342 |
| ENSG00000 | gene | 0,94278053 | 0,01824885 | 0,07393246 | CNDP2 | -0,0850061 |
| ENSG00000 | gene | 0,85502304 | 0,01825086 | 0,07393246 | AC106795.1 | -0,2259648 |
| ENSG00002 | gene | 1,20572289 | 0,01825198 | 0,07393246 | SRRM5 | 0,26989838 |
| ENSG00000 | gene | 1,14449949 | 0,01826752 | 0,07396516 | ZC3H15 | 0,19471683 |

| | | | | | | |
|---|---|---|---|---|---|---|
| ENSG00000 | gene | 0,89160411 | 0,01827514 | 0,07396582 | DHRS1 | -0,1655248 |
| ENSG00000 | gene | 0,87099715 | 0,01832858 | 0,07415185 | ASPHD1 | -0,1992601 |
| ENSG00000 | gene | 0,80759699 | 0,01835027 | 0,07420934 | PRKCZ | -0,3082926 |
| ENSG00000 | gene | 0,85308073 | 0,01837733 | 0,07427945 | FAM102A | -0,2292458 |
| ENSG00000 | gene | 0,79835862 | 0,01838259 | 0,07427945 | GLA | -0,3248911 |
| ENSG00000 | gene | 0,8576927 | 0,01839372 | 0,07428794 | ACD | -0,2214672 |
| ENSG00000 | gene | 0,89255925 | 0,01839968 | 0,07428794 | PRR14 | -0,1639801 |
| ENSG00000 | gene | 0,918914 | 0,01842115 | 0,0743182 | PHF5A | -0,1219982 |
| ENSG00000 | gene | 0,87614277 | 0,0184328 | 0,0743182 | NFIL3 | -0,1907621 |
| ENSG00000 | gene | 1,15758975 | 0,01843453 | 0,0743182 | NR1D2 | 0,21112405 |
| ENSG00000 | gene | 0,89881986 | 0,01843715 | 0,0743182 | MAU2 | -0,1538961 |
| ENSG00000 | gene | 1,20532586 | 0,01844515 | 0,07432023 | GOPC | 0,26942323 |
| ENSG00000 | gene | 0,65479428 | 0,01845894 | 0,0743456 | CXCL1 | -0,6108864 |
| ENSG00000 | gene | 0,89635646 | 0,01849734 | 0,0744601 | PPP1R12C | -0,1578555 |
| ENSG00000 | gene | 1,34945517 | 0,01850752 | 0,0744601 | AC027290.2 | 0,43237705 |
| ENSG00000 | gene | 1,1291833 | 0,0185099 | 0,0744601 | ERBIN | 0,1752797 |
| ENSG00000 | gene | 0,78491834 | 0,01856274 | 0,07459107 | ANKRD11 | -0,3493855 |
| ENSG00000 | gene | 1,23324285 | 0,01856303 | 0,07459107 | HGF | 0,30245692 |
| ENSG00000 | gene | 0,83706568 | 0,01857222 | 0,07459107 | ETHE1 | -0,2565873 |
| ENSG00000 | gene | 1,14762838 | 0,01857254 | 0,07459107 | SCARF2 | 0,19865556 |
| ENSG00000 | gene | 0,89754015 | 0,01862858 | 0,07478585 | DVL2 | -0,1559516 |
| ENSG00000 | gene | 1,10467501 | 0,01864456 | 0,07480101 | FBXO11 | 0,143622 |
| ENSG00000 | gene | 0,92833972 | 0,01864745 | 0,07480101 | POLR2C | -0,1072753 |
| ENSG00000 | gene | 0,83328114 | 0,01867467 | 0,07487991 | DDX58 | -0,2631248 |
| ENSG00000 | gene | 0,68289116 | 0,0186941 | 0,07492754 | IER3 | -0,5502724 |
| ENSG00000 | gene | 1,15373173 | 0,01870996 | 0,07496079 | PRTFDC1 | 0,20630781 |
| ENSG00000 | gene | 0,91666376 | 0,01872961 | 0,0750092 | TSEN34 | -0,1255355 |
| ENSG00000 | gene | 0,93052531 | 0,01873756 | 0,07501075 | PGAM1 | -0,1038827 |
| ENSG00000 | gene | 1,07419676 | 0,01876587 | 0,07509378 | RAP1GDS1 | 0,10325827 |
| ENSG00000 | gene | 0,80455588 | 0,01879036 | 0,07510304 | ZFAS1 | -0,3137355 |
| ENSG00000 | gene | 1,35377622 | 0,01879981 | 0,07510304 | AL117190.4 | 0,43698928 |
| ENSG00000 | gene | 1,33615339 | 0,01880229 | 0,07510304 | IFT122 | 0,41808564 |
| ENSG00000 | gene | 0,92940828 | 0,0188041 | 0,07510304 | PSMD8 | -0,1056156 |
| ENSG00000 | gene | 0,74501581 | 0,01880623 | 0,07510304 | ASB1 | -0,4246571 |
| ENSG00000 | gene | 1,23286269 | 0,01881705 | 0,07510304 | AP000779.1 | 0,30201213 |
| ENSG00000 | gene | 1,25447309 | 0,0188212 | 0,07510304 | OXR1 | 0,32708152 |
| ENSG00000 | gene | 0,94136642 | 0,01892743 | 0,07549304 | PACSIN2 | -0,0871717 |
| ENSG00000 | gene | 0,77764896 | 0,01893417 | 0,07549304 | TBC1D10A | -0,362809 |
| ENSG00000 | gene | 0,91437351 | 0,01895147 | 0,07553167 | ABR | -0,1291445 |
| ENSG00000 | gene | 1,40830051 | 0,0189706 | 0,07557752 | SNORD87 | 0,49395522 |
| ENSG00000 | gene | 0,89332116 | 0,01903885 | 0,07581899 | NDUFA6 | -0,1627492 |
| ENSG00000 | gene | 1,26620778 | 0,01905424 | 0,07584979 | LAMA5 | 0,34051416 |
| ENSG00000 | gene | 0,78903883 | 0,01907941 | 0,0759195 | BANF1 | -0,3418318 |
| ENSG00000 | gene | 0,81670577 | 0,01908981 | 0,07593041 | CRYAB | -0,2921117 |
| ENSG00000 | gene | 1,1184195 | 0,01910911 | 0,0759767 | RRS1 | 0,16146142 |
| ENSG00000 | gene | 1,09533908 | 0,01914443 | 0,07608663 | MT-ND5 | 0,13137755 |
| ENSG00000 | gene | 0,89977037 | 0,01915808 | 0,07611039 | BCAR3 | -0,1523712 |
| ENSG00000 | gene | 0,87893566 | 0,01920775 | 0,07627064 | TP53I3 | -0,1861705 |
| ENSG00000 | gene | 0,91630465 | 0,0192138 | 0,07627064 | SNX1 | -0,1261007 |
| ENSG00000 | gene | 0,9334602 | 0,01925103 | 0,07638786 | PRPF19 | -0,0993396 |
| ENSG00000 | gene | 0,85315303 | 0,01929339 | 0,07652529 | SUPT7L | -0,2291236 |
| ENSG00000 | gene | 0,89976034 | 0,01931924 | 0,07657995 | EHMT2 | -0,1523873 |
| ENSG00000 | gene | 0,91385295 | 0,01932261 | 0,07657995 | IRF2 | -0,1299661 |
| ENSG00000 | gene | 0,81799387 | 0,01934643 | 0,07664373 | GABARAP | -0,2898381 |
| ENSG00000 | gene | 1,32151104 | 0,01936098 | 0,07667071 | FO704657.1 | 0,40218847 |
| ENSG00000 | gene | 0,88128353 | 0,01938016 | 0,07668549 | AC245060.7 | -0,1823219 |
| ENSG00000 | gene | 0,94092806 | 0,0193848 | 0,07668549 | STAT3 | -0,0878437 |

| | | | | | | |
|---|---|---|---|---|---|---|
| ENSG00000 | gene | 0,93682928 | 0,01939912 | 0,07668549 | FAM149B1 | -0,0941419 |
| ENSG00000 | gene | 0,89906841 | 0,01940253 | 0,07668549 | DRAM2 | -0,1534972 |
| ENSG00000 | gene | 1,58952188 | 0,01941059 | 0,07668549 | AC004707.2 | 0,66859288 |
| ENSG00000 | gene | 1,40781433 | 0,01941111 | 0,07668549 | LRRC57 | 0,49345707 |
| ENSG00000 | gene | 0,92527609 | 0,01945356 | 0,07682258 | P3H1 | -0,1120442 |
| ENSG00000 | gene | 0,89473511 | 0,01946708 | 0,0768331 | PABPC1 | -0,1604675 |
| ENSG00000 | gene | 1,10484705 | 0,01948183 | 0,0768331 | CBX3 | 0,14384666 |
| ENSG00000 | gene | 0,86261157 | 0,01948699 | 0,0768331 | GRK2 | -0,213217 |
| ENSG00000 | gene | 1,061279 | 0,01948722 | 0,0768331 | PICALM | 0,08580398 |
| ENSG00000 | gene | 0,92109735 | 0,01951418 | 0,07690883 | DBN1 | -0,1185744 |
| ENSG00000 | gene | 0,81917868 | 0,01952576 | 0,07691226 | LINC00623 | -0,2877499 |
| ENSG00000 | gene | 1,15121246 | 0,01953056 | 0,07691226 | AC018557.1 | 0,20315411 |
| ENSG00000 | gene | 0,86133396 | 0,01956534 | 0,07701862 | HMBOX1 | -0,2153554 |
| ENSG00000 | gene | 0,92540556 | 0,01958739 | 0,07707481 | SNW1 | -0,1118423 |
| ENSG00000 | gene | 0,866884 | 0,01959955 | 0,07709207 | LTBP4 | -0,2060891 |
| ENSG00000 | gene | 1,23363433 | 0,01962429 | 0,07710777 | DEPDC1 | 0,30291482 |
| ENSG00000 | gene | 1,10853021 | 0,01962453 | 0,07710777 | FAM210B | 0,14864809 |
| ENSG00000 | gene | 1,53192025 | 0,01962687 | 0,07710777 | CELF2 | 0,61534119 |
| ENSG00000 | gene | 1,36387337 | 0,01966263 | 0,07721769 | MIR4253 | 0,4477097 |
| ENSG00000 | gene | 0,83893864 | 0,01969177 | 0,07730152 | AC016596.3 | -0,2533628 |
| ENSG00000 | gene | 1,16119749 | 0,01971749 | 0,07737185 | MAMLD1 | 0,21561336 |
| ENSG00000 | gene | 1,10092326 | 0,01972535 | 0,07737204 | IPO9 | 0,13871391 |
| ENSG00000 | gene | 0,86752362 | 0,01979928 | 0,07761801 | RPS14 | -0,2050251 |
| ENSG00000 | gene | 1,31670331 | 0,01980371 | 0,07761801 | AL139317.3 | 0,3969303 |
| ENSG00000 | gene | 1,45585391 | 0,01981276 | 0,07762282 | AC005045.2 | 0,54186559 |
| ENSG00000 | gene | 0,81552989 | 0,01986304 | 0,07778907 | CHMP1B | -0,2941903 |
| ENSG00000 | gene | 0,86313903 | 0,01989962 | 0,07786598 | AP000688.1 | -0,2123351 |
| ENSG00000 | gene | 0,85079775 | 0,01990215 | 0,07786598 | HSCB | -0,2331119 |
| ENSG00000 | gene | 0,86966604 | 0,01990624 | 0,07786598 | PRELID1 | -0,2014666 |
| ENSG00000 | gene | 1,39349284 | 0,01996111 | 0,07804983 | AMPH | 0,47870559 |
| ENSG00000 | gene | 0,90344175 | 0,01997367 | 0,07806817 | KIRREL3 | -0,1464965 |
| ENSG00000 | gene | 1,13677817 | 0,0200381 | 0,07828913 | ACTR2 | 0,18495075 |
| ENSG00000 | gene | 1,12649354 | 0,02005132 | 0,0783099 | F2R | 0,17183904 |
| ENSG00000 | gene | 1,39043663 | 0,02010877 | 0,07850336 | RAD51C | 0,47553799 |
| ENSG00000 | gene | 1,12922918 | 0,02014873 | 0,07862842 | MTMR1 | 0,17533832 |
| ENSG00000 | gene | 0,87178463 | 0,02016438 | 0,07863945 | RTF2 | -0,1979563 |
| ENSG00000 | gene | 1,16121703 | 0,02016742 | 0,07863945 | AL031284.1 | 0,21563764 |
| ENSG00000 | gene | 1,10016921 | 0,02018433 | 0,07867447 | CMTR2 | 0,13772543 |
| ENSG00000 | gene | 1,09801523 | 0,02024481 | 0,07885738 | DYNC2LI1 | 0,13489807 |
| ENSG00000 | gene | 0,84141905 | 0,02024716 | 0,07885738 | PIN1 | -0,2491036 |
| ENSG00000 | gene | 0,87233121 | 0,02026851 | 0,07890951 | RPS6 | -0,1970521 |
| ENSG00000 | gene | 1,20319708 | 0,02028123 | 0,07892804 | AC007314.1 | 0,26687298 |
| ENSG00000 | gene | 0,90922117 | 0,02029443 | 0,07893775 | CBX4 | -0,1372968 |
| ENSG00000 | gene | 0,94157839 | 0,02030223 | 0,07893775 | ERGIC3 | -0,0868469 |
| ENSG00000 | gene | 1,76542019 | 0,02031252 | 0,07893775 | CHFR | 0,8200116 |
| ENSG00000 | gene | 0,91919641 | 0,02031556 | 0,07893775 | DHX32 | -0,1215549 |
| ENSG00000 | gene | 0,85012128 | 0,02035657 | 0,07904228 | NEU1 | -0,2342594 |
| ENSG00000 | gene | 0,89337156 | 0,02035841 | 0,07904228 | TSTA3 | -0,1626678 |
| ENSG00000 | gene | 1,04506157 | 0,02040248 | 0,07915871 | BRD7 | 0,06358795 |
| ENSG00000 | gene | 1,09854804 | 0,02040436 | 0,07915871 | ITPRID2 | 0,13559796 |
| ENSG00000 | gene | 0,87273856 | 0,02043255 | 0,07923705 | CDIPT | -0,1963785 |
| ENSG00000 | gene | 0,90194771 | 0,02044559 | 0,07925663 | AC092849.1 | -0,1488843 |
| ENSG00000 | gene | 0,74763314 | 0,02046579 | 0,07929242 | RGS2 | -0,4195976 |
| ENSG00000 | gene | 0,89402537 | 0,02047082 | 0,07929242 | MAPK8IP3 | -0,1616123 |
| ENSG00000 | gene | 1,27503345 | 0,02049034 | 0,07933705 | SNORA44 | 0,3505351 |
| ENSG00000 | gene | 1,28526432 | 0,02052639 | 0,07944276 | AP003396.5 | 0,36206509 |
| ENSG00000 | gene | 0,89406616 | 0,02053366 | 0,07944276 | FKBP10 | -0,1615465 |

| | | | | | | |
|---|---|---|---|---|---|---|
| ENSG00000 | gene | 0,8945308 | 0,02056152 | 0,07951952 | QDPR | -0,1607969 |
| ENSG00000 | gene | 0,7695945 | 0,02058965 | 0,07959725 | AC005062.1 | -0,3778296 |
| ENSG00000 | gene | 0,89504418 | 0,020604 | 0,07962168 | SELENOH | -0,1599692 |
| ENSG00000 | gene | 1,1267017 | 0,02070646 | 0,07998645 | SOX9 | 0,1721056 |
| ENSG00000 | gene | 1,34600781 | 0,02077153 | 0,08020659 | AC112211.1 | 0,42868678 |
| ENSG00000 | gene | 0,94943671 | 0,02082861 | 0,08039569 | WDR82 | -0,0748563 |
| ENSG00000 | gene | 0,86770998 | 0,02085466 | 0,0804649 | RNASE4 | -0,2047152 |
| ENSG00000 | gene | 0,93355479 | 0,0208635 | 0,08046771 | DDX19B | -0,0991934 |
| ENSG00000 | gene | 1,13267993 | 0,02087403 | 0,08047703 | RLIM | 0,17974025 |
| ENSG00000 | gene | 0,62071001 | 0,02089919 | 0,08051936 | FP671120.4 | -0,6880087 |
| ENSG00000 | gene | 0,90391965 | 0,02090125 | 0,08051936 | KAT5 | -0,1457336 |
| ENSG00000 | gene | 0,86233455 | 0,02091936 | 0,08054072 | LRCH3 | -0,2136804 |
| ENSG00000 | gene | 0,86082639 | 0,02092304 | 0,08054072 | RPS7 | -0,2162058 |
| ENSG00000 | gene | 1,14388631 | 0,02094343 | 0,08056786 | PSTPIP2 | 0,19394367 |
| ENSG00000 | gene | 0,90970696 | 0,02095526 | 0,08056786 | GNG5 | -0,1365262 |
| ENSG00000 | gene | 0,83445841 | 0,02095867 | 0,08056786 | SPOP | -0,261088 |
| ENSG00000 | gene | 0,91996938 | 0,02096259 | 0,08056786 | FKBP11 | -0,1203423 |
| ENSG00000 | gene | 0,89401478 | 0,02100516 | 0,08068861 | DDX39B | -0,1616294 |
| ENSG00000 | gene | 1,32009342 | 0,02101827 | 0,08068861 | AC013489.3 | 0,40064003 |
| ENSG00000 | gene | 0,83027939 | 0,02102119 | 0,08068861 | CKS2 | -0,2683312 |
| ENSG00000 | gene | 0,87682885 | 0,02102656 | 0,08068861 | TMEM14C | -0,1896328 |
| ENSG00000 | gene | 1,15286484 | 0,02104758 | 0,08073801 | NECTIN3 | 0,20522338 |
| ENSG00000 | gene | 0,86070884 | 0,02106333 | 0,0807571 | KRI1 | -0,2164028 |
| ENSG00000 | gene | 1,09577347 | 0,02106884 | 0,0807571 | ANKIB1 | 0,13194958 |
| ENSG00000 | gene | 1,26549355 | 0,02111075 | 0,08088648 | LMO7 | 0,33970016 |
| ENSG00000 | gene | 0,82800092 | 0,02113565 | 0,0809031 | RPL14 | -0,2722957 |
| ENSG00000 | gene | 1,09107867 | 0,02113879 | 0,0809031 | ASF1A | 0,12575513 |
| ENSG00000 | gene | 0,88115919 | 0,02113957 | 0,0809031 | SMIM15-AS | -0,1825254 |
| ENSG00000 | gene | 0,92064083 | 0,02116497 | 0,08095032 | JOSD1 | -0,1192897 |
| ENSG00000 | gene | 0,79086597 | 0,02116823 | 0,08095032 | TIMM8B | -0,3384949 |
| ENSG00000 | gene | 1,14100313 | 0,02119452 | 0,08101959 | CLIC4 | 0,19030275 |
| ENSG00000 | gene | 0,90518479 | 0,02122224 | 0,08105057 | EPHX1 | -0,1437158 |
| ENSG00000 | gene | 0,93981526 | 0,02122607 | 0,08105057 | GTPBP2 | -0,0895509 |
| ENSG00000 | gene | 1,25741625 | 0,02122714 | 0,08105057 | WDR76 | 0,33046231 |
| ENSG00000 | gene | 0,90522727 | 0,02124481 | 0,08108682 | SMARCD3 | -0,1436481 |
| ENSG00000 | gene | 1,05908487 | 0,02125968 | 0,08111233 | SCCPDH | 0,0828182 |
| ENSG00000 | gene | 1,12262983 | 0,02131775 | 0,08130263 | PAXBP1 | 0,1668823 |
| ENSG00000 | gene | 0,89501574 | 0,02132597 | 0,08130271 | SNRPA | -0,160015 |
| ENSG00000 | gene | 1,20045241 | 0,02134964 | 0,08134671 | DYNLT3 | 0,26357821 |
| ENSG00000 | gene | 0,88414994 | 0,02135458 | 0,08134671 | TM2D2 | -0,177637 |
| ENSG00000 | gene | 1,17435245 | 0,02136213 | 0,08134671 | OBI1-AS1 | 0,23186546 |
| ENSG00000 | gene | 1,11879889 | 0,02141881 | 0,08153124 | TRNT1 | 0,16195073 |
| ENSG00000 | gene | 0,84881038 | 0,02143318 | 0,08155467 | RPS20 | -0,2364858 |
| ENSG00000 | gene | 1,23528892 | 0,02144935 | 0,08158489 | AP003096.1 | 0,30484851 |
| ENSG00000 | gene | 0,40487422 | 0,02151354 | 0,08179763 | CXCL8 | -1,3044543 |
| ENSG00000 | gene | 1,25017317 | 0,02154527 | 0,08188688 | CDKN2AIP | 0,32212794 |
| ENSG00000 | gene | 1,11183399 | 0,02155526 | 0,08189347 | MTMR6 | 0,15294139 |
| ENSG00000 | gene | 1,26136162 | 0,02160326 | 0,08204438 | MIR762 | 0,33498195 |
| ENSG00000 | gene | 0,92204478 | 0,02161812 | 0,08205518 | TBP | -0,1170913 |
| ENSG00000 | gene | 0,50011746 | 0,02162265 | 0,08205518 | AD000671.1 | -0,9996611 |
| ENSG00000 | gene | 1,38169531 | 0,02171289 | 0,08236611 | DLG1 | 0,4664395 |
| ENSG00000 | gene | 0,78716332 | 0,02173737 | 0,08242746 | PIGQ | -0,3452651 |
| ENSG00000 | gene | 1,08853891 | 0,02176518 | 0,08250134 | ZW10 | 0,12239298 |
| ENSG00000 | gene | 0,90428502 | 0,0217759 | 0,08251043 | CTBP2 | -0,1451505 |
| ENSG00000 | gene | 0,75287301 | 0,02180381 | 0,08258464 | NBEAL1 | -0,4095216 |
| ENSG00000 | gene | 0,93619772 | 0,02182362 | 0,0826281 | SRRT | -0,0951148 |
| ENSG00000 | gene | 0,85253874 | 0,02188779 | 0,08283224 | ABHD17C | -0,2301627 |

| | | | | | | |
|---|---|---|---|---|---|---|
| ENSG00000 | gene | 0,92705348 | 0,02189424 | 0,08283224 | DNAJC7 | -0,1092755 |
| ENSG00000( | gene | 1,06975232 | 0,02190671 | 0,0828478 | RNF19A | 0,0972768 |
| ENSG00000 | gene | 0,8997632 | 0,02197593 | 0,08307789 | GBA | -0,1523827 |
| ENSG00000 | gene | 0,94390461 | 0,02205782 | 0,08335571 | TAGLN2 | -0,083287 |
| ENSG00000 | gene | 1,2091383 | 0,02209086 | 0,08344306 | CCDC34 | 0,27397927 |
| ENSG00000 | gene | 0,90910987 | 0,02209777 | 0,08344306 | MPG | -0,1374734 |
| ENSG00000 | gene | 1,3077293 | 0,02210681 | 0,08344543 | RRP7BP | 0,38706394 |
| ENSG00000 | gene | 1,16261189 | 0,02212161 | 0,08346952 | MELK | 0,21736957 |
| ENSG00000 | gene | 1,34041996 | 0,02214436 | 0,08352357 | IMMP2L | 0,42268507 |
| ENSG00000 | gene | 1,31047394 | 0,02217638 | 0,08361256 | LRRN3 | 0,39008866 |
| ENSG00000 | gene | 0,7997194 | 0,02218973 | 0,08363106 | PHLDA1 | -0,3224342 |
| ENSG00000 | gene | 1,12467012 | 0,02220131 | 0,08364291 | PHYKPL | 0,16950191 |
| ENSG00000; | gene | 0,89422663 | 0,02221939 | 0,08367924 | PLEKHO2 | -0,1612876 |
| ENSG00000( | gene | 1,05781884 | 0,02232209 | 0,08399111 | MRPS10 | 0,08109258 |
| ENSG00000; | gene | 0,8790321 | 0,02232318 | 0,08399111 | LINC02637 | -0,1860123 |
| ENSG00000 | gene | 1,33707525 | 0,02232761 | 0,08399111 | LIG4 | 0,41908066 |
| ENSG00000 | gene | 0,92771703 | 0,02235683 | 0,08405391 | MLLT1 | -0,1082433 |
| ENSG00000; | gene | 1,38131077 | 0,02236956 | 0,08405391 | AC114498.2 | 0,46603794 |
| ENSG00000; | gene | 1,25346926 | 0,02236973 | 0,08405391 | COA6-AS1 | 0,32592661 |
| ENSG00000 | gene | 0,89012072 | 0,02238562 | 0,08408174 | LTBR | -0,1679271 |
| ENSG00000 | gene | 1,19052967 | 0,02243135 | 0,08421042 | RIN2 | 0,25160357 |
| ENSG00000 | gene | 0,84204718 | 0,02243686 | 0,08421042 | SARAF | -0,248027 |
| ENSG00000; | gene | 1,08122344 | 0,022456 | 0,08425035 | PCMTD2 | 0,11266469 |
| ENSG00000; | gene | 0,84097386 | 0,02248494 | 0,08432705 | STRADA | -0,2498671 |
| ENSG00000; | gene | 0,82800714 | 0,02254471 | 0,08451831 | FADS3 | -0,2722849 |
| ENSG00000 | gene | 0,88315567 | 0,02255953 | 0,08451831 | SIAH2 | -0,1792603 |
| ENSG00000 | gene | 1,07815234 | 0,02256849 | 0,08451831 | MFSD1 | 0,10856104 |
| ENSG00000; | gene | 0,61963094 | 0,02257004 | 0,08451831 | AC117378.1 | -0,6905189 |
| ENSG00000 | gene | 1,1807423 | 0,02259963 | 0,08459719 | CASKIN1 | 0,23969412 |
| ENSG00000 | gene | 1,51506002 | 0,02261515 | 0,08460761 | ADGRB2 | 0,59937495 |
| ENSG00000; | gene | 0,84146417 | 0,02262341 | 0,08460761 | CYTOR | -0,2490262 |
| ENSG00000 | gene | 1,21369312 | 0,02262801 | 0,08460761 | CKAP2 | 0,27940368 |
| ENSG00000 | gene | 1,07613482 | 0,02264945 | 0,08465583 | CEP89 | 0,10585883 |
| ENSG00000 | gene | 0,92609074 | 0,02267271 | 0,08468352 | MRPS22 | -0,1107745 |
| ENSG00000 | gene | 1,42691186 | 0,02267394 | 0,08468352 | TMEM241 | 0,51289622 |
| ENSG00000; | gene | 0,71229437 | 0,0227107 | 0,0847713 | MDP1 | -0,4894545 |
| ENSG00000 | gene | 0,88891281 | 0,02271454 | 0,0847713 | NONO | -0,1698862 |
| ENSG00000; | gene | 1,19643868 | 0,02274792 | 0,0848537 | AC005944.1 | 0,25874646 |
| ENSG00000( | gene | 0,76461741 | 0,0227555 | 0,0848537 | SLC2A3 | -0,38719 |
| ENSG00000 | gene | 0,93361457 | 0,02277615 | 0,0848537 | POLE3 | -0,099101 |
| ENSG00000( | gene | 0,93142886 | 0,02277885 | 0,0848537 | SIRT2 | -0,1024825 |
| ENSG00000 | gene | 0,87074944 | 0,0227794 | 0,0848537 | DPAGT1 | -0,1996705 |
| ENSG00000 | gene | 1,27492315 | 0,02281478 | 0,08495358 | NDST2 | 0,35041029 |
| ENSG00000( | gene | 1,14553797 | 0,02286273 | 0,08510016 | VRK2 | 0,19602527 |
| ENSG00000; | gene | 1,44932832 | 0,02287349 | 0,08510826 | RNU6-2 | 0,53538445 |
| ENSG00000( | gene | 1,09094121 | 0,02294358 | 0,08532627 | P3H2 | 0,12557336 |
| ENSG00000 | gene | 1,0739607 | 0,02294929 | 0,08532627 | TMEM181 | 0,1029412 |
| ENSG00000 | gene | 1,59247486 | 0,02297041 | 0,08537279 | DFFA | 0,67127059 |
| ENSG00000( | gene | 0,92155732 | 0,0229888 | 0,08537722 | SMARCB1 | -0,1178542 |
| ENSG00000 | gene | 1,14115366 | 0,0229965 | 0,08537722 | ZRANB2 | 0,19049306 |
| ENSG00000 | gene | 0,9145191 | 0,02299744 | 0,08537722 | SETDB1 | -0,1289148 |
| ENSG00000 | gene | 0,90098587 | 0,02302259 | 0,08543862 | TPRA1 | -0,1504236 |
| ENSG00000; | gene | 0,82243168 | 0,02305919 | 0,08554245 | PSMA6P1 | -0,2820323 |
| ENSG00000 | gene | 0,77359157 | 0,02309193 | 0,08562236 | MSC | -0,370356 |
| ENSG00000; | gene | 0,9084432 | 0,023098 | 0,08562236 | NPM1P27 | -0,1385318 |
| ENSG00000 | gene | 0,88183007 | 0,02314273 | 0,08574099 | CALCOCO2 | -0,1814274 |
| ENSG00000 | gene | 0,75972232 | 0,02315269 | 0,08574099 | RPS15 | -0,3964559 |

| | | | | | | |
|---|---|---|---|---|---|---|
| ENSG00000 | gene | 1,10444704 | 0,02315595 | 0,08574099 | SNX7 | 0,14332424 |
| ENSG00000 | gene | 0,8774332 | 0,02320013 | 0,08587251 | RABAC1 | -0,1886388 |
| ENSG00000 | gene | 0,79758728 | 0,02324905 | 0,08600123 | PNRC1 | -0,3262857 |
| ENSG00000( | gene | 1,11055923 | 0,02325225 | 0,08600123 | NFX1 | 0,15128634 |
| ENSG00000 | gene | 1,13170861 | 0,02334133 | 0,08629852 | TSPYL4 | 0,17850254 |
| ENSG00000 | gene | 1,21577175 | 0,02335405 | 0,0863032 | PBRM1 | 0,2818724 |
| ENSG00000 | gene | 1,13829252 | 0,02336 | 0,0863032 | ORMDL1 | 0,18687135 |
| ENSG00000 | gene | 0,8188577 | 0,02340468 | 0,08643607 | CNBP | -0,2883153 |
| ENSG00000 | gene | 1,33954466 | 0,02343811 | 0,08652727 | EPS8L2 | 0,42174268 |
| ENSG00000 | gene | 0,90668258 | 0,02348439 | 0,0866347 | SRSF1 | -0,1413305 |
| ENSG00000 | gene | 1,32338084 | 0,02348468 | 0,0866347 | STK36 | 0,40422829 |
| ENSG00000 | gene | 1,31777843 | 0,02354991 | 0,08684303 | TNFSF4 | 0,39810781 |
| ENSG00000: | gene | 1,23984083 | 0,0235926 | 0,08696666 | AP006621.5 | 0,31015492 |
| ENSG00000 | gene | 0,95537505 | 0,02360751 | 0,08696666 | TSR1 | -0,0658609 |
| ENSG00000 | gene | 0,89938169 | 0,02360975 | 0,08696666 | MFSD12 | -0,1529946 |
| ENSG00000 | gene | 1,2488661 | 0,02364822 | 0,08706994 | GTF2IRD2B | 0,32061881 |
| ENSG00000 | gene | 0,84024586 | 0,02365535 | 0,08706994 | DAB2IP | -0,2511166 |
| ENSG00000 | gene | 0,7020691 | 0,02368789 | 0,08713963 | GSDMD | -0,5103151 |
| ENSG00000 | gene | 1,15256426 | 0,02369712 | 0,08713963 | PRPF39 | 0,20484719 |
| ENSG00000: | gene | 0,80789101 | 0,02370086 | 0,08713963 | AL121845.3 | -0,3077674 |
| ENSG00000 | gene | 0,81083371 | 0,02370943 | 0,08713963 | FOXRED1 | -0,302522 |
| ENSG00000 | gene | 1,12936378 | 0,02375923 | 0,0872903 | ACAP3 | 0,17551027 |
| ENSG00000 | gene | 0,79804689 | 0,02378056 | 0,08730844 | RECQL5 | -0,3254546 |
| ENSG00000 | gene | 1,18370795 | 0,02378178 | 0,08730844 | WDR66 | 0,24331317 |
| ENSG00000 | gene | 0,91412739 | 0,02383625 | 0,08747501 | NDEL1 | -0,1295329 |
| ENSG00000 | gene | 0,91267096 | 0,02384479 | 0,08747501 | SEC61B | -0,1318333 |
| ENSG00000 | gene | 0,93908154 | 0,02386707 | 0,08752435 | UTP14A | -0,0906777 |
| ENSG00000 | gene | 0,81812797 | 0,02388605 | 0,08756157 | NRP2 | -0,2896016 |
| ENSG00000 | gene | 0,92599247 | 0,02391181 | 0,0876236 | XPC | -0,1109276 |
| ENSG00000 | gene | 0,88521818 | 0,02392164 | 0,08762376 | P4HTM | -0,175895 |
| ENSG00000( | gene | 0,93189882 | 0,02393252 | 0,08762376 | MUL1 | -0,1017548 |
| ENSG00000: | gene | 1,13147658 | 0,02393836 | 0,08762376 | MIR222HG | 0,17820672 |
| ENSG00000: | gene | 1,35955462 | 0,02401866 | 0,08785676 | SNORD14C | 0,44313411 |
| ENSG00000: | gene | 1,28210676 | 0,02401973 | 0,08785676 | AC009511.2 | 0,3585164 |
| ENSG00000 | gene | 1,07962304 | 0,02406694 | 0,08799698 | FUBP1 | 0,11052767 |
| ENSG00000 | gene | 0,94040507 | 0,02408239 | 0,08801977 | WIPI2 | -0,0886458 |
| ENSG00000 | gene | 0,92225109 | 0,02409266 | 0,08801977 | UBE2D3 | -0,1167685 |
| ENSG00000 | gene | 0,88812744 | 0,02409981 | 0,08801977 | PYM1 | -0,1711614 |
| ENSG00000: | gene | 1,29020084 | 0,02413381 | 0,08811152 | OCIAD1-AS1 | 0,36759566 |
| ENSG00000( | gene | 1,09832859 | 0,02416044 | 0,08817627 | SLC30A9 | 0,13530973 |
| ENSG00000 | gene | 0,86780899 | 0,02420896 | 0,08831867 | EIF4A2 | -0,2045506 |
| ENSG00000 | gene | 0,86003579 | 0,02422409 | 0,08831867 | PRPF38B | -0,2175314 |
| ENSG00000 | gene | 0,82919131 | 0,02422618 | 0,08831867 | ABCD4 | -0,2702231 |
| ENSG00000 | gene | 1,39136141 | 0,02423695 | 0,08832546 | PTBP2 | 0,47649721 |
| ENSG00000: | gene | 0,73521318 | 0,02438391 | 0,08882839 | SNHG15 | -0,4437655 |
| ENSG00000( | gene | 1,08407247 | 0,02440386 | 0,08886842 | CDC5L | 0,1164612 |
| ENSG00000 | gene | 0,73458218 | 0,02441562 | 0,08887859 | TMEM234 | -0,4450042 |
| ENSG00000: | gene | 1,17301345 | 0,02443544 | 0,08890865 | PRMT5-AS1 | 0,23021956 |
| ENSG00000 | gene | 1,14546577 | 0,02444181 | 0,08890865 | TRUB1 | 0,19593434 |
| ENSG00000 | gene | 0,92640511 | 0,02446068 | 0,08894466 | CHID1 | -0,1102849 |
| ENSG00000 | gene | 1,10850508 | 0,02447015 | 0,0889465 | TACC1 | 0,14861538 |
| ENSG00000 | gene | 1,13088047 | 0,02448837 | 0,08898011 | MCCC2 | 0,17744644 |
| ENSG00000( | gene | 0,90334125 | 0,02450034 | 0,08899099 | APBA2 | -0,146657 |
| ENSG00000 | gene | 0,94071716 | 0,02454641 | 0,08912214 | CHPT1 | -0,0881671 |
| ENSG00000 | gene | 1,34628571 | 0,02455807 | 0,08912214 | LIN7A | 0,42898461 |
| ENSG00000( | gene | 0,71492344 | 0,02456341 | 0,08912214 | MAP3K4 | -0,4841393 |
| ENSG00000 | gene | 1,10585853 | 0,02459205 | 0,0891934 | DDX21 | 0,14516684 |

| | | | | | | |
|---|---|---|---|---|---|---|
| ENSG00000( | gene | 0,90166822 | 0,0246267 | 0,08928642 | RNF10 | -0,1493314 |
| ENSG00000 | gene | 0,89171948 | 0,02469859 | 0,08951435 | GMPR2 | -0,1653382 |
| ENSG00000 | gene | 0,90578263 | 0,02476091 | 0,0897074 | BCL7B | -0,1427632 |
| ENSG00000 | gene | 1,51340024 | 0,02477693 | 0,08973267 | ZNF605 | 0,59779358 |
| ENSG00000: | gene | 0,87336223 | 0,02483439 | 0,08990794 | AP5Z1 | -0,195348 |
| ENSG00000 | gene | 0,67543767 | 0,02486364 | 0,08997231 | RAB40C | -0,5661055 |
| ENSG00000: | gene | 1,21833283 | 0,02487032 | 0,08997231 | AC018761.4 | 0,28490831 |
| ENSG00000 | gene | 0,92545391 | 0,024931 | 0,09013654 | CTSK | -0,1117669 |
| ENSG00000 | gene | 0,87270021 | 0,0249339 | 0,09013654 | AGPAT3 | -0,196442 |
| ENSG00000 | gene | 0,8930124 | 0,02495007 | 0,09014978 | SLC43A3 | -0,1632479 |
| ENSG00000 | gene | 0,92590466 | 0,0249576 | 0,09014978 | C11orf58 | -0,1110644 |
| ENSG00000: | gene | 1,25310076 | 0,02496853 | 0,09014978 | AC098595.1 | 0,32550243 |
| ENSG00000( | gene | 0,92393647 | 0,02497463 | 0,09014978 | IDH3G | -0,1141344 |
| ENSG00000 | gene | 0,86112385 | 0,02498476 | 0,09014978 | MID1IP1 | -0,2157073 |
| ENSG00000 | gene | 0,93996105 | 0,02499211 | 0,09014978 | TMCO3 | -0,0893271 |
| ENSG00000 | gene | 0,9198787 | 0,02502508 | 0,09023588 | TMEM219 | -0,1204845 |
| ENSG00000: | gene | 1,22728335 | 0,02506199 | 0,09033614 | RPL35P2 | 0,29546837 |
| ENSG00000 | gene | 0,86930422 | 0,02512049 | 0,09051408 | PPP1R16A | -0,2020669 |
| ENSG00000 | gene | 0,77999498 | 0,02519853 | 0,09075008 | RGCC | -0,3584633 |
| ENSG00000 | gene | 1,36047719 | 0,02520429 | 0,09075008 | MRM2 | 0,44411277 |
| ENSG00000 | gene | 0,92157635 | 0,02522056 | 0,09077569 | SUPT5H | -0,1178244 |
| ENSG00000 | gene | 0,92992853 | 0,02530859 | 0,0910041 | SLC35G2 | -0,1048083 |
| ENSG00000: | gene | 1,32464731 | 0,02531182 | 0,0910041 | PGM5P2 | 0,40560829 |
| ENSG00000 | gene | 1,19377707 | 0,02532857 | 0,0910041 | CRIM1 | 0,25553345 |
| ENSG00000 | gene | 0,92201678 | 0,02534472 | 0,0910041 | PIGG | -0,1171351 |
| ENSG00000 | gene | 0,93308736 | 0,02534527 | 0,0910041 | STX18 | -0,0999159 |
| ENSG00000 | gene | 1,36052701 | 0,02535359 | 0,0910041 | CHD9 | 0,4441656 |
| ENSG00000 | gene | 0,8885163 | 0,02536125 | 0,0910041 | SGSM2 | -0,1705299 |
| ENSG00000 | gene | 1,21390376 | 0,02536751 | 0,0910041 | LPAR1 | 0,27965405 |
| ENSG00000( | gene | 1,12448159 | 0,02537293 | 0,0910041 | TNPO1 | 0,16926004 |
| ENSG00000 | gene | 1,12431064 | 0,02538765 | 0,0910041 | MSRA | 0,1690407 |
| ENSG00000 | gene | 0,9126844 | 0,02540007 | 0,0910041 | MTMR14 | -0,131812 |
| ENSG00000 | gene | 1,25349456 | 0,02540125 | 0,0910041 | XRCC3 | 0,32595574 |
| ENSG00000 | gene | 0,85801515 | 0,02540332 | 0,0910041 | PIM1 | -0,220925 |
| ENSG00000: | gene | 1,10792696 | 0,02542664 | 0,09105473 | AC018638.4 | 0,14786278 |
| ENSG00000 | gene | 1,14673558 | 0,02544647 | 0,09109285 | GPD2 | 0,19753276 |
| ENSG00000: | gene | 1,61225115 | 0,02552714 | 0,09134866 | MIR3661 | 0,6890765 |
| ENSG00000( | gene | 0,89897747 | 0,02554191 | 0,09136854 | SMARCE1 | -0,1536431 |
| ENSG00000 | gene | 0,85599084 | 0,02555944 | 0,09139826 | NCLN | -0,2243327 |
| ENSG00000: | gene | 1,09890731 | 0,0255713 | 0,09140771 | NRAS | 0,13606971 |
| ENSG00000 | gene | 1,15901475 | 0,02560569 | 0,09148856 | ARL5A | 0,21289893 |
| ENSG00000: | gene | 0,76927926 | 0,02561237 | 0,09148856 | LINC01504 | -0,3784207 |
| ENSG00000( | gene | 0,85601768 | 0,02562331 | 0,09149467 | PDCD2 | -0,2242875 |
| ENSG00000 | gene | 1,11849267 | 0,02564249 | 0,09153022 | PRKACB | 0,16155581 |
| ENSG00000 | gene | 1,06995206 | 0,02566327 | 0,09157144 | GGCX | 0,09754616 |
| ENSG00000 | gene | 0,88642715 | 0,02568457 | 0,09161446 | SNN | -0,173926 |
| ENSG00000 | gene | 1,13685165 | 0,02576648 | 0,09187358 | GUF1 | 0,185044 |
| ENSG00000 | gene | 0,87530022 | 0,02581502 | 0,09201356 | NPIPB12 | -0,1921502 |
| ENSG00000 | gene | 0,94864597 | 0,0258932 | 0,09225907 | RAB35 | -0,0760583 |
| ENSG00000 | gene | 0,93505677 | 0,02591361 | 0,09226792 | ETFA | -0,0968741 |
| ENSG00000 | gene | 0,94484844 | 0,0259143 | 0,09226792 | CCT3 | -0,0818452 |
| ENSG00000 | gene | 1,06076284 | 0,02593947 | 0,09232441 | ATXN10 | 0,08510214 |
| ENSG00000 | gene | 0,80016875 | 0,02597765 | 0,09242711 | KDM6B | -0,3216238 |
| ENSG00000 | gene | 0,85760702 | 0,02603505 | 0,09259811 | GPR89A | -0,2216114 |
| ENSG00000: | gene | 1,26747311 | 0,02606693 | 0,09267827 | AP000763.3 | 0,34195514 |
| ENSG00000( | gene | 0,91914704 | 0,02614619 | 0,09292673 | ALKBH5 | -0,1216324 |
| ENSG00000 | gene | 0,88929185 | 0,0262334 | 0,09318425 | RPL10A | -0,1692711 |

| | | | | | | |
|---|---|---|---|---|---|---|
| ENSG00000( | gene | 0,92422355 | 0,02623744 | 0,09318425 | FKBP1A | -0,1136862 |
| ENSG00000' | gene | 0,93046887 | 0,02626312 | 0,09324207 | PEX19 | -0,1039702 |
| ENSG00000' | gene | 0,92156264 | 0,02631347 | 0,09338376 | SHKBP1 | -0,1178459 |
| ENSG00000; | gene | 1,33545348 | 0,02634219 | 0,09338376 | AF129075.4 | 0,41732972 |
| ENSG00000' | gene | 0,91910868 | 0,02634626 | 0,09338376 | OCRL | -0,1216926 |
| ENSG00000' | gene | 0,90400397 | 0,02634646 | 0,09338376 | NUMBL | -0,145599 |
| ENSG00000' | gene | 0,80059604 | 0,02635012 | 0,09338376 | QTRT2 | -0,3208536 |
| ENSG00000' | gene | 1,25603183 | 0,02642317 | 0,09359809 | KLC3 | 0,32887303 |
| ENSG00000' | gene | 0,94747344 | 0,02643434 | 0,09359809 | EMC3 | -0,0778426 |
| ENSG00000( | gene | 0,88254827 | 0,02643891 | 0,09359809 | TMEM161A | -0,1802529 |
| ENSG00000' | gene | 1,14182918 | 0,02649273 | 0,09375513 | TBX2 | 0,19134684 |
| ENSG00000( | gene | 1,43548761 | 0,02650987 | 0,09378234 | WDR7 | 0,52154087 |
| ENSG00000; | gene | 1,21522502 | 0,02655864 | 0,09389583 | AL138689.2 | 0,28122348 |
| ENSG00000' | gene | 1,32885038 | 0,02656089 | 0,09389583 | EPHA1 | 0,41017868 |
| ENSG00000( | gene | 1,07531878 | 0,02661617 | 0,09405773 | CRKL | 0,10476441 |
| ENSG00000( | gene | 0,84716671 | 0,02662909 | 0,09406791 | TAF11 | -0,2392822 |
| ENSG00000' | gene | 0,90730048 | 0,02664594 | 0,09406791 | POLR1D | -0,1403477 |
| ENSG00000' | gene | 0,89637536 | 0,02664751 | 0,09406791 | MRAS | -0,1578251 |
| ENSG00000' | gene | 1,09510528 | 0,02665771 | 0,09407042 | SLC38A2 | 0,13106957 |
| ENSG00000' | gene | 0,95731631 | 0,02666908 | 0,09407707 | CCT5 | -0,0629324 |
| ENSG00000' | gene | 0,87873764 | 0,0266939 | 0,09413111 | CCS | -0,1864956 |
| ENSG00000' | gene | 0,9061772 | 0,02673992 | 0,0942599 | CDR2 | -0,1421349 |
| ENSG00000; | gene | 1,23042673 | 0,02675148 | 0,09426711 | AL591684.1 | 0,29915875 |
| ENSG00000( | gene | 1,15530418 | 0,02682744 | 0,09450121 | TUT7 | 0,20827274 |
| ENSG00000' | gene | 0,8981763 | 0,02689289 | 0,09469811 | PPP1R14BP | -0,1549294 |
| ENSG00000' | gene | 1,108232 | 0,02692907 | 0,09479185 | PRRG1 | 0,14825992 |
| ENSG00000( | gene | 0,9229022 | 0,02700157 | 0,09501334 | METTL1 | -0,1157503 |
| ENSG00000' | gene | 0,83120591 | 0,0270309 | 0,0950828 | EIF1AD | -0,2667222 |
| ENSG00000' | gene | 0,77850159 | 0,02704897 | 0,09511263 | CEP131 | -0,3612281 |
| ENSG00000; | gene | 1,26090026 | 0,02707501 | 0,09517043 | OSTN-AS1 | 0,33445416 |
| ENSG00000' | gene | 1,34195106 | 0,02712084 | 0,09529775 | MAGOHB | 0,42433205 |
| ENSG00000( | gene | 0,87692539 | 0,02715262 | 0,09535701 | ALDH3B1 | -0,189474 |
| ENSG00000' | gene | 0,72273185 | 0,02716447 | 0,09535701 | GAL3ST4 | -0,4684676 |
| ENSG00000( | gene | 0,92978497 | 0,02716656 | 0,09535701 | SPPL2B | -0,105031 |
| ENSG00000; | gene | 1,1065297 | 0,02718293 | 0,09538073 | TMED7-TICA | 0,14604218 |
| ENSG00000' | gene | 1,06348785 | 0,02725452 | 0,09559808 | MINPP1 | 0,08880356 |
| ENSG00000' | gene | 0,9054397 | 0,02734715 | 0,09586136 | PNKD | -0,1433095 |
| ENSG00000; | gene | 1,2048015 | 0,02734891 | 0,09586136 | CALM2P2 | 0,26879547 |
| ENSG00000' | gene | 0,89596028 | 0,02741595 | 0,0960624 | DMAC2 | -0,1584933 |
| ENSG00000' | gene | 0,89322476 | 0,02743838 | 0,09610701 | FAM53C | -0,1629049 |
| ENSG00000' | gene | 0,88440818 | 0,02745491 | 0,09613096 | RUBCN | -0,1772157 |
| ENSG00000' | gene | 0,91740782 | 0,02746554 | 0,09613423 | EIF2B4 | -0,1243649 |
| ENSG00000' | gene | 0,9439986 | 0,0275211 | 0,09629471 | RBMX | -0,0831434 |
| ENSG00000' | gene | 1,51641157 | 0,02758108 | 0,09647053 | SRRM3 | 0,60066137 |
| ENSG00000' | gene | 0,92901499 | 0,02762527 | 0,09659102 | BOD1 | -0,1062262 |
| ENSG00000' | gene | 0,91458137 | 0,02763695 | 0,0965978 | PTK2 | -0,1288166 |
| ENSG00000( | gene | 0,92502634 | 0,02766317 | 0,09665539 | ATP6AP1 | -0,1124337 |
| ENSG00000' | gene | 1,05839676 | 0,0276791 | 0,096677 | VIRMA | 0,08188055 |
| ENSG00000' | gene | 1,13874388 | 0,02785625 | 0,09726149 | MAP7D3 | 0,18744331 |
| ENSG00000' | gene | 0,893326 | 0,02790533 | 0,09739853 | BECN1 | -0,1627413 |
| ENSG00000' | gene | 1,09545856 | 0,02797134 | 0,09754879 | TMEM167A | 0,13153491 |
| ENSG00000' | gene | 0,89556777 | 0,02797926 | 0,09754879 | EAF1 | -0,1591255 |
| ENSG00000' | gene | 1,19786442 | 0,02798439 | 0,09754879 | RBM24 | 0,26046462 |
| ENSG00000' | gene | 1,13490174 | 0,02798773 | 0,09754879 | PPP4R2 | 0,18256739 |
| ENSG00000' | gene | 0,91767196 | 0,02801214 | 0,09756443 | MTERF4 | -0,1239496 |
| ENSG00000' | gene | 0,92828165 | 0,02801425 | 0,09756443 | PRKRIP1 | -0,1073655 |
| ENSG00000' | gene | 1,09346816 | 0,02802173 | 0,09756443 | MBOAT2 | 0,12891121 |

| | | | | | | |
|---|---|---|---|---|---|---|
| ENSG00000 | gene | 0,89674637 | 0,02805724 | 0,09761245 | TMEM59L | -0,1572281 |
| ENSG00000 | gene | 0,72523142 | 0,02806414 | 0,09761245 | HADH | -0,4634867 |
| ENSG00000 | gene | 1,31008657 | 0,02807164 | 0,09761245 | AC245033.3 | 0,38966215 |
| ENSG00000 | gene | 0,92040215 | 0,0280749 | 0,09761245 | PLOD3 | -0,1196637 |
| ENSG00000 | gene | 0,8815501 | 0,02812312 | 0,09774583 | PRKAB1 | -0,1818855 |
| ENSG00000 | gene | 1,37046115 | 0,02815256 | 0,09781385 | ZMYM3 | 0,45466143 |
| ENSG00000 | gene | 1,07101022 | 0,02829178 | 0,09826315 | UBR5 | 0,09897224 |
| ENSG00000 | gene | 1,21779531 | 0,02831543 | 0,09831085 | SEMA3C | 0,28427166 |
| ENSG00000 | gene | 0,89074042 | 0,02833858 | 0,09835678 | AGAP3 | -0,166923 |
| ENSG00000 | gene | 0,9201272 | 0,02840747 | 0,09853334 | ATP10A | -0,1200948 |
| ENSG00000 | gene | 0,81114802 | 0,02840932 | 0,09853334 | STX1A | -0,3019629 |
| ENSG00000 | gene | 1,26019828 | 0,02842809 | 0,09856397 | AC009511.1 | 0,33365074 |
| ENSG00000 | gene | 0,77142163 | 0,02845525 | 0,09857815 | FTH1P7 | -0,3744085 |
| ENSG00000 | gene | 1,22480178 | 0,02845578 | 0,09857815 | AC108010.1 | 0,29254828 |
| ENSG00000 | gene | 0,91959678 | 0,028462 | 0,09857815 | MAP7D1 | -0,1209267 |
| ENSG00000 | gene | 0,94583504 | 0,02847813 | 0,09859957 | FBL | -0,0803395 |
| ENSG00000 | gene | 1,40137972 | 0,02850465 | 0,09862979 | Z94160.2 | 0,48684792 |
| ENSG00000 | gene | 1,10092392 | 0,02851091 | 0,09862979 | PDS5A | 0,13871478 |
| ENSG00000 | gene | 1,10036878 | 0,02853634 | 0,09862979 | EFR3A | 0,13798712 |
| ENSG00000 | gene | 0,92087947 | 0,02854083 | 0,09862979 | RPTOR | -0,1189158 |
| ENSG00000 | gene | 0,91305062 | 0,02854535 | 0,09862979 | GPS1 | -0,1312332 |
| ENSG00000 | gene | 1,11727934 | 0,02855828 | 0,09862979 | DPY19L4 | 0,15998993 |
| ENSG00000 | gene | 0,92859866 | 0,02855969 | 0,09862979 | CUEDC1 | -0,1068729 |
| ENSG00000 | gene | 1,12246968 | 0,02856643 | 0,09862979 | EBAG9 | 0,16667647 |
| ENSG00000 | gene | 0,94282471 | 0,02860927 | 0,0987433 | RNF123 | -0,0849385 |
| ENSG00000 | gene | 0,87474099 | 0,02866201 | 0,09889092 | G6PC3 | -0,1930722 |
| ENSG00000 | gene | 0,91027038 | 0,02868732 | 0,09894381 | SMNDC1 | -0,135633 |
| ENSG00000 | gene | 1,31205366 | 0,02871753 | 0,09901356 | AL645933.1 | 0,39182673 |
| ENSG00000 | gene | 0,83910819 | 0,02875687 | 0,09911474 | SLC52A2 | -0,2530713 |
| ENSG00000 | gene | 1,2457097 | 0,0287726 | 0,09913449 | SNORD58A | 0,31696791 |
| ENSG00000 | gene | 0,8957066 | 0,02879257 | 0,09915477 | FAM162A | -0,1589019 |
| ENSG00000 | gene | 1,1143705 | 0,02879848 | 0,09915477 | ARV1 | 0,15622897 |
| ENSG00000 | gene | 1,31006842 | 0,02882242 | 0,09920275 | MTBP | 0,38964216 |
| ENSG00000 | gene | 1,15180054 | 0,02885449 | 0,09927866 | EIF5AP4 | 0,2038909 |
| ENSG00000 | gene | 0,92533115 | 0,02887287 | 0,09929869 | SPRYD3 | -0,1119583 |
| ENSG00000 | gene | 0,70010265 | 0,02888034 | 0,09929869 | FAM13A | -0,5143616 |
| ENSG00000 | gene | 1,13228488 | 0,02896221 | 0,09951979 | POLE | 0,17923698 |
| ENSG00000 | gene | 1,17311699 | 0,02896471 | 0,09951979 | IL10RB-DT | 0,23034689 |
| ENSG00000 | gene | 1,10361307 | 0,02897476 | 0,09951984 | ECHDC1 | 0,14223445 |
| ENSG00000 | gene | 1,32396028 | 0,02901752 | 0,09963219 | S100A2 | 0,40485984 |
| ENSG00000 | gene | 0,90428171 | 0,02904904 | 0,09970587 | NDUFS2 | -0,1451558 |
| ENSG00000 | gene | 1,10131188 | 0,02909299 | 0,09980554 | MT-TE | 0,13922308 |
| ENSG00000 | gene | 0,79532357 | 0,02909821 | 0,09980554 | EFNA4 | -0,3303862 |
| ENSG00000 | gene | 0,76635917 | 0,0291334 | 0,09986034 | MSC-AS1 | -0,3839074 |
| ENSG00000 | gene | 1,20666541 | 0,02913433 | 0,09986034 | VPS8 | 0,27102569 |
| ENSG00000 | gene | 1,6899613 | 0,02920497 | 0,10006788 | APOLD1 | 0,75699021 |
| ENSG00000 | gene | 1,48652874 | 0,02927449 | 0,10027146 | LYRM9 | 0,57194736 |
| ENSG00000 | gene | 1,11101716 | 0,02931696 | 0,10038224 | THOC7 | 0,15188109 |
| ENSG00000 | gene | 0,91964789 | 0,0293329 | 0,10040214 | STYXL1 | -0,1208465 |
| ENSG00000 | gene | 1,15336587 | 0,02934579 | 0,10041161 | PI4K2B | 0,20585023 |
| ENSG00000 | gene | 1,2457237 | 0,02942181 | 0,10057307 | SHROOM3 | 0,31698411 |
| ENSG00000 | gene | 0,78481675 | 0,02942204 | 0,10057307 | JPT2 | -0,3495723 |
| ENSG00000 | gene | 0,88264652 | 0,0294234 | 0,10057307 | TRAPPC2B | -0,1800923 |
| ENSG00000 | gene | 0,79218983 | 0,02949025 | 0,10076681 | AC104066.5 | -0,3360819 |
| ENSG00000 | gene | 1,2508386 | 0,02952032 | 0,10083482 | BTBD19 | 0,32289564 |
| ENSG00000 | gene | 1,30647472 | 0,02953354 | 0,10084525 | POT1 | 0,38567921 |
| ENSG00000 | gene | 1,08831087 | 0,0295977 | 0,10102952 | WDR43 | 0,12209072 |

| | | | | | | |
|---|---|---|---|---|---|---|
| ENSG00000 | gene | 0,89505992 | 0,02961385 | 0,10104986 | RAB24 | -0,1599438 |
| ENSG00000 | gene | 1,21935531 | 0,02964349 | 0,1011162 | SERPINB7 | 0,28611857 |
| ENSG00000 | gene | 1,39909958 | 0,02966403 | 0,10115148 | FASTKD3 | 0,48449865 |
| ENSG00000 | gene | 1,26642399 | 0,0296934 | 0,10121683 | PDGFD | 0,34076049 |
| ENSG00000 | gene | 1,17915252 | 0,02974601 | 0,1013613 | DIPK1A | 0,23775033 |
| ENSG00000 | gene | 0,91584435 | 0,02976831 | 0,10140246 | LAMP1 | -0,1268257 |
| ENSG00000 | gene | 0,90731269 | 0,02981082 | 0,10151239 | PRCP | -0,1403283 |
| ENSG00000 | gene | 0,9141664 | 0,02983904 | 0,1015736 | DUSP12 | -0,1294713 |
| ENSG00000 | gene | 0,94568815 | 0,02987003 | 0,10163443 | SLC39A7 | -0,0805636 |
| ENSG00000 | gene | 0,86330255 | 0,02987741 | 0,10163443 | TOMM40 | -0,2120618 |
| ENSG00000 | gene | 0,88141389 | 0,02990088 | 0,10167938 | NSMCE1 | -0,1821085 |
| ENSG00000 | gene | 1,25014984 | 0,02992465 | 0,10171753 | EXOC6 | 0,32210102 |
| ENSG00000 | gene | 1,23898114 | 0,02993641 | 0,10171753 | AC010834.1 | 0,30915422 |
| ENSG00000 | gene | 0,94665691 | 0,02994287 | 0,10171753 | TCF25 | -0,0790864 |
| ENSG00000 | gene | 1,09587463 | 0,0299652 | 0,10175853 | INTS8 | 0,13208276 |
| ENSG00000 | gene | 1,24499991 | 0,02999126 | 0,10181218 | AL592146.1 | 0,31614563 |
| ENSG00000 | gene | 1,16175624 | 0,03007923 | 0,10207585 | ADAM9 | 0,21630739 |
| ENSG00000 | gene | 0,8850946 | 0,03009439 | 0,10208679 | CCNB1IP1 | -0,1760964 |
| ENSG00000 | gene | 0,7778433 | 0,03010304 | 0,10208679 | YDJC | -0,3624485 |
| ENSG00000 | gene | 0,91574066 | 0,03011831 | 0,10210364 | CABIN1 | -0,126989 |
| ENSG00000 | gene | 0,86384405 | 0,03016467 | 0,10222585 | CLK3 | -0,2111572 |
| ENSG00000 | gene | 0,93992589 | 0,03020029 | 0,10228572 | MORF4L1 | -0,0893811 |
| ENSG00000 | gene | 0,89310921 | 0,03020296 | 0,10228572 | PACS1 | -0,1630915 |
| ENSG00000 | gene | 1,09063612 | 0,03022522 | 0,10232614 | SSH2 | 0,12516985 |
| ENSG00000 | gene | 0,88575231 | 0,03027592 | 0,10246281 | VAC14 | -0,1750248 |
| ENSG00000 | gene | 0,88004469 | 0,03029288 | 0,10248449 | SREBF1 | -0,1843513 |
| ENSG00000 | gene | 0,92937451 | 0,030303 | 0,10248449 | LPCAT4 | -0,105668 |
| ENSG00000 | gene | 0,84769918 | 0,03031434 | 0,1024879 | DPYSL4 | -0,2383757 |
| ENSG00000 | gene | 1,41379399 | 0,03033662 | 0,10252827 | AC006116.8 | 0,49957192 |
| ENSG00000 | gene | 0,94064578 | 0,03035804 | 0,1025657 | YARS2 | -0,0882766 |
| ENSG00000 | gene | 1,05991837 | 0,03038493 | 0,10262159 | NCOR1 | 0,08395315 |
| ENSG00000 | gene | 0,87450877 | 0,03039529 | 0,1026216 | ZNF706 | -0,1934552 |
| ENSG00000 | gene | 0,96884413 | 0,03041863 | 0,10266547 | EXT1 | -0,0456635 |
| ENSG00000 | gene | 0,86271313 | 0,03043738 | 0,10269381 | TRIM52-AS1 | -0,2130472 |
| ENSG00000 | gene | 0,72073317 | 0,03044948 | 0,10269966 | NEAT1 | -0,4724629 |
| ENSG00000 | gene | 0,84906887 | 0,03046312 | 0,10271073 | C1S | -0,2360465 |
| ENSG00000 | gene | 0,88704808 | 0,03049056 | 0,10272045 | MBD3 | -0,1729158 |
| ENSG00000 | gene | 0,94589189 | 0,03049177 | 0,10272045 | IFT46 | -0,0802528 |
| ENSG00000 | gene | 1,21419004 | 0,03049708 | 0,10272045 | PLK4 | 0,27999425 |
| ENSG00000 | gene | 0,84796129 | 0,03058413 | 0,1029787 | KDM2B | -0,2379297 |
| ENSG00000 | gene | 0,81112007 | 0,0306282 | 0,10305441 | IFITM2 | -0,3020126 |
| ENSG00000 | gene | 1,19344134 | 0,0306354 | 0,10305441 | LXN | 0,25512765 |
| ENSG00000 | gene | 0,88511606 | 0,03064797 | 0,10305441 | WAC | -0,1760614 |
| ENSG00000 | gene | 1,26472609 | 0,03064819 | 0,10305441 | LYPD1 | 0,33882497 |
| ENSG00000 | gene | 0,8447709 | 0,03073974 | 0,1033272 | THOC6 | -0,243368 |
| ENSG00000 | gene | 0,90076461 | 0,03075538 | 0,10334473 | CD81 | -0,1507779 |
| ENSG00000 | gene | 0,92561709 | 0,0307697 | 0,10335784 | NPM1 | -0,1115126 |
| ENSG00000 | gene | 0,85762243 | 0,03079081 | 0,1033769 | TBC1D31 | -0,2215855 |
| ENSG00000 | gene | 0,93541501 | 0,0308027 | 0,1033769 | PRPF3 | -0,0963215 |
| ENSG00000 | gene | 0,76951078 | 0,03081566 | 0,1033769 | AC067751.1 | -0,3779866 |
| ENSG00000 | gene | 0,88125372 | 0,03081707 | 0,1033769 | SIRT6 | -0,1823707 |
| ENSG00000 | gene | 1,22444398 | 0,03088395 | 0,1035662 | SFRP1 | 0,29212677 |
| ENSG00000 | gene | 1,1879243 | 0,0309104 | 0,10361984 | SCOC | 0,24844291 |
| ENSG00000 | gene | 0,81661069 | 0,03100226 | 0,10389269 | RPL14P1 | -0,2922796 |
| ENSG00000 | gene | 0,88218249 | 0,03102792 | 0,10394355 | PLK3 | -0,180851 |
| ENSG00000 | gene | 1,12569926 | 0,03107049 | 0,10402661 | NUFIP2 | 0,17082146 |
| ENSG00000 | gene | 0,93499846 | 0,03107962 | 0,10402661 | CSNK1G2 | -0,0969641 |

| | | | | | | |
|---|---|---|---|---|---|---|
| ENSG00000 | gene | 1,11418072 | 0,03108419 | 0,10402661 | SMIM15 | 0,15598326 |
| ENSG00000 | gene | 1,45645029 | 0,03112592 | 0,10410166 | WWTR1-AS1 | 0,54245647 |
| ENSG00000 | gene | 1,11410303 | 0,03113784 | 0,10410166 | AL390728.4 | 0,15588265 |
| ENSG00000 | gene | 0,76652046 | 0,03114685 | 0,10410166 | RPL41P5 | -0,3836038 |
| ENSG00000 | gene | 1,16950895 | 0,03114861 | 0,10410166 | TSLP | 0,2259029 |
| ENSG00000 | gene | 0,89749565 | 0,03122992 | 0,1043315 | LSM1 | -0,1560232 |
| ENSG00000 | gene | 0,87868689 | 0,03124741 | 0,1043315 | SCAMP4 | -0,1865789 |
| ENSG00000 | gene | 1,1433984 | 0,03124895 | 0,1043315 | TUB | 0,19332817 |
| ENSG00000 | gene | 0,8906638 | 0,03130288 | 0,1044764 | PAIP2 | -0,1670471 |
| ENSG00000 | gene | 0,78111843 | 0,03134535 | 0,10458294 | AC073861.1 | -0,3563868 |
| ENSG00000 | gene | 0,68894045 | 0,03138419 | 0,10467732 | MT1X | -0,5375488 |
| ENSG00000 | gene | 0,84759958 | 0,03144033 | 0,1048293 | TINF2 | -0,2385452 |
| ENSG00000 | gene | 0,69842864 | 0,03147306 | 0,10487429 | MTAP | -0,5178154 |
| ENSG00000 | gene | 1,3028951 | 0,03147709 | 0,10487429 | AGAP6 | 0,38172094 |
| ENSG00000 | gene | 1,15098859 | 0,03148751 | 0,10487429 | SPCS3 | 0,20287354 |
| ENSG00000 | gene | 0,85026891 | 0,03149613 | 0,10487429 | AFAP1L1 | -0,2340089 |
| ENSG00000 | gene | 1,08838923 | 0,03153745 | 0,10497662 | ASPH | 0,12219459 |
| ENSG00000 | gene | 0,92847949 | 0,03159953 | 0,10514797 | GBA2 | -0,1070581 |
| ENSG00000 | gene | 0,87853982 | 0,03163266 | 0,10519158 | STEAP3 | -0,1868204 |
| ENSG00000 | gene | 1,11441914 | 0,03164292 | 0,10519158 | ZNF280D | 0,15629194 |
| ENSG00000 | gene | 1,36961658 | 0,03165326 | 0,10519158 | AC008149.1 | 0,45377207 |
| ENSG00000 | gene | 0,87538598 | 0,03165507 | 0,10519158 | OST4 | -0,1920088 |
| ENSG00000 | gene | 0,90728533 | 0,03173624 | 0,10542598 | SP110 | -0,1403718 |
| ENSG00000 | gene | 0,9316208 | 0,03176567 | 0,10548842 | HABP4 | -0,1021852 |
| ENSG00000 | gene | 0,91701751 | 0,03180497 | 0,10556126 | GATAD1 | -0,1249788 |
| ENSG00000 | gene | 0,92782264 | 0,03181497 | 0,10556126 | KIF7 | -0,108079 |
| ENSG00000 | gene | 1,14152374 | 0,03181955 | 0,10556126 | HAUS6 | 0,19096086 |
| ENSG00000 | gene | 1,42726765 | 0,03190442 | 0,10580744 | GPX1P1 | 0,5132559 |
| ENSG00000 | gene | 0,88319397 | 0,0319559 | 0,10594274 | CSNK2B | -0,1791978 |
| ENSG00000 | gene | 1,15366551 | 0,03199442 | 0,10603497 | ANP32E | 0,206225 |
| ENSG00000 | gene | 0,88379932 | 0,0320136 | 0,10606311 | DCAF11 | -0,1782093 |
| ENSG00000 | gene | 1,11532545 | 0,03202994 | 0,10608178 | CETN3 | 0,15746475 |
| ENSG00000 | gene | 0,94832028 | 0,03207708 | 0,10620245 | RBM10 | -0,0765537 |
| ENSG00000 | gene | 0,92066655 | 0,03209178 | 0,10621564 | PFKL | -0,1192494 |
| ENSG00000 | gene | 1,23476508 | 0,03213559 | 0,1063051 | TYMS | 0,30423659 |
| ENSG00000 | gene | 0,68604176 | 0,03214025 | 0,1063051 | IRF9 | -0,5436317 |
| ENSG00000 | gene | 0,69871682 | 0,03218218 | 0,1063607 | C11orf96 | -0,5172202 |
| ENSG00000 | gene | 0,81531938 | 0,03218851 | 0,1063607 | ZFAND2A | -0,2945628 |
| ENSG00000 | gene | 0,88432657 | 0,03218924 | 0,1063607 | DDX47 | -0,1773489 |
| ENSG00000 | gene | 0,94418109 | 0,03232276 | 0,10670603 | UQCRC1 | -0,0828645 |
| ENSG00000 | gene | 0,69776225 | 0,03232674 | 0,10670603 | PRR5 | -0,5191925 |
| ENSG00000 | gene | 1,38631068 | 0,03232719 | 0,10670603 | DOP1A | 0,47125061 |
| ENSG00000 | gene | 0,86703234 | 0,03233679 | 0,10670603 | CHCHD5 | -0,2058423 |
| ENSG00000 | gene | 0,92410608 | 0,03238792 | 0,10683919 | LSM14B | -0,1138696 |
| ENSG00000 | gene | 0,52876653 | 0,03241368 | 0,10688861 | RNASEK-C1 | -0,9192972 |
| ENSG00000 | gene | 1,16168717 | 0,03243416 | 0,1069206 | TBCAP1 | 0,21622161 |
| ENSG00000 | gene | 0,92260161 | 0,03244883 | 0,1069334 | AP3D1 | -0,1162203 |
| ENSG00000 | gene | 1,10691845 | 0,03248064 | 0,10700268 | FAM114A2 | 0,14654894 |
| ENSG00000 | gene | 1,13977779 | 0,03254123 | 0,10716668 | CTF1 | 0,18875259 |
| ENSG00000 | gene | 0,6458679 | 0,03255695 | 0,10718283 | AC027031.2 | -0,630689 |
| ENSG00000 | gene | 0,84754485 | 0,03257057 | 0,10719208 | TBC1D20 | -0,2386384 |
| ENSG00000 | gene | 0,9368111 | 0,03258711 | 0,10720933 | MAPK3 | -0,0941699 |
| ENSG00000 | gene | 1,12074185 | 0,0326017 | 0,10720933 | PFAS | 0,164454 |
| ENSG00000 | gene | 1,32912605 | 0,03260824 | 0,10720933 | NUTM2B-AS | 0,41047793 |
| ENSG00000 | gene | 1,21654845 | 0,03263353 | 0,10723878 | AL023584.1 | 0,28279378 |
| ENSG00000 | gene | 0,90875238 | 0,03263936 | 0,10723878 | RGL2 | -0,1380409 |
| ENSG00000 | gene | 1,19086367 | 0,03264964 | 0,10723878 | AL034380.1 | 0,25200826 |

| | | | | | | |
|---|---|---|---|---|---|---|
| ENSG00000 | gene | 0,91875719 | 0,03269545 | 0,10735367 | BBS2 | -0,1222445 |
| ENSG00000 | gene | 1,07776596 | 0,03273798 | 0,10745773 | PGM2 | 0,10804393 |
| ENSG00000 | gene | 0,90300681 | 0,03288539 | 0,10790587 | NRBP2 | -0,1471912 |
| ENSG00000 | gene | 0,94680585 | 0,03292061 | 0,10794124 | PFKM | -0,0788595 |
| ENSG00000; | gene | 1,21717232 | 0,03292214 | 0,10794124 | AC060766.5 | 0,28353343 |
| ENSG00000; | gene | 1,46336345 | 0,03293804 | 0,10794124 | AL162258.2 | 0,54928813 |
| ENSG00000; | gene | 0,64109802 | 0,03293971 | 0,10794124 | HSPA1A | -0,6413831 |
| ENSG00000 | gene | 0,91446846 | 0,03296373 | 0,10798427 | ABCF3 | -0,1289947 |
| ENSG00000 | gene | 1,16744216 | 0,0330054 | 0,10808505 | ANO4 | 0,22335108 |
| ENSG00000( | gene | 0,94366371 | 0,03303856 | 0,10815249 | BAK1 | -0,0836553 |
| ENSG00000; | gene | 1,33250575 | 0,03305821 | 0,10815249 | MIR3120 | 0,41414176 |
| ENSG00000 | gene | 0,92024018 | 0,03305871 | 0,10815249 | SH2B1 | -0,1199176 |
| ENSG00000 | gene | 1,0677371 | 0,03309414 | 0,10820154 | IARS1 | 0,09455646 |
| ENSG00000 | gene | 0,89512905 | 0,03309553 | 0,10820154 | VKORC1L1 | -0,1598324 |
| ENSG00000 | gene | 1,08654689 | 0,03314907 | 0,10834088 | SKI | 0,11975044 |
| ENSG00000 | gene | 0,9300345 | 0,03322161 | 0,1085339 | EIF2B2 | -0,1046439 |
| ENSG00000 | gene | 0,87680149 | 0,03323002 | 0,1085339 | ASCC1 | -0,1896778 |
| ENSG00000 | gene | 0,88445638 | 0,03333215 | 0,10883161 | NT5C | -0,1771371 |
| ENSG00000 | gene | 0,84189546 | 0,03342505 | 0,1090636 | STUB1 | -0,248287 |
| ENSG00000 | gene | 0,9155555 | 0,0334252 | 0,1090636 | TNFAIP2 | -0,1272807 |
| ENSG00000 | gene | 0,79689575 | 0,03344392 | 0,1090821 | IFT22 | -0,3275371 |
| ENSG00000; | gene | 1,16812403 | 0,03345302 | 0,1090821 | AC138028.4 | 0,22419347 |
| ENSG00000 | gene | 1,10258019 | 0,03346387 | 0,1090821 | SMAD3 | 0,14088359 |
| ENSG00000 | gene | 0,95394717 | 0,03360239 | 0,10946047 | GOT1 | -0,0680187 |
| ENSG00000 | gene | 0,92800806 | 0,03360817 | 0,10946047 | SPATS2L | -0,1077908 |
| ENSG00000 | gene | 0,74863722 | 0,03361306 | 0,10946047 | INSIG1 | -0,4176613 |
| ENSG00000 | gene | 1,17358137 | 0,03369705 | 0,10969793 | UFM1 | 0,23091788 |
| ENSG00000 | gene | 0,91547959 | 0,03375838 | 0,10986154 | ILVBL | -0,1274004 |
| ENSG00000; | gene | 0,93333429 | 0,03377354 | 0,10987481 | RCC1L | -0,0995342 |
| ENSG00000( | gene | 0,9323646 | 0,03389942 | 0,11024815 | CFAP20 | -0,1010339 |
| ENSG00000 | gene | 1,18957726 | 0,03391231 | 0,11025393 | PDCD1LG2 | 0,25044897 |
| ENSG00000 | gene | 0,90679716 | 0,03395028 | 0,11030075 | SLC25A11 | -0,1411482 |
| ENSG00000( | gene | 0,8797149 | 0,03395415 | 0,11030075 | ST3GAL1 | -0,184892 |
| ENSG00000 | gene | 0,79695932 | 0,03396008 | 0,11030075 | SLC4A3 | -0,327422 |
| ENSG00000 | gene | 0,91507713 | 0,03397192 | 0,11030306 | PRPSAP1 | -0,1280347 |
| ENSG00000 | gene | 0,81712165 | 0,03400942 | 0,11038867 | WARS1 | -0,2913772 |
| ENSG00000 | gene | 0,84145916 | 0,03403444 | 0,1104247 | IFI27L1 | -0,2490348 |
| ENSG00000 | gene | 0,9381509 | 0,03404279 | 0,1104247 | ATP8B2 | -0,0921081 |
| ENSG00000 | gene | 0,85485097 | 0,03412371 | 0,11064153 | TRNAU1AP | -0,2262552 |
| ENSG00000 | gene | 0,94324996 | 0,03413195 | 0,11064153 | PWWP3A | -0,084288 |
| ENSG00000; | gene | 1,34954334 | 0,03427386 | 0,11106523 | AC136469.2 | 0,43247131 |
| ENSG00000( | gene | 0,93126736 | 0,03430589 | 0,11111886 | CTSA | -0,1027327 |
| ENSG00000 | gene | 0,8938828 | 0,03431282 | 0,11111886 | MAP1S | -0,1618424 |
| ENSG00000 | gene | 0,87187686 | 0,03433898 | 0,11116728 | FZR1 | -0,1978037 |
| ENSG00000 | gene | 0,91202627 | 0,03440698 | 0,11135106 | NDUFAF2 | -0,1328527 |
| ENSG00000 | gene | 1,15072732 | 0,03444342 | 0,11142415 | SLC38A4 | 0,20254601 |
| ENSG00000 | gene | 0,93220994 | 0,03445204 | 0,11142415 | MDH2 | -0,1012732 |
| ENSG00000 | gene | 0,93199911 | 0,0344729 | 0,11145525 | MTCH1 | -0,1015995 |
| ENSG00000 | gene | 0,90376504 | 0,03451315 | 0,111526 | SIL1 | -0,1459803 |
| ENSG00000 | gene | 0,89008389 | 0,03451728 | 0,111526 | LPCAT3 | -0,1679868 |
| ENSG00000 | gene | 0,92526805 | 0,03456175 | 0,11163333 | KDM5C | -0,1120567 |
| ENSG00000 | gene | 0,89816223 | 0,03458032 | 0,11165695 | RPUSD4 | -0,154952 |
| ENSG00000 | gene | 0,80518446 | 0,03463053 | 0,11175332 | NGF | -0,3126088 |
| ENSG00000 | gene | 0,8656797 | 0,03463271 | 0,11175332 | KLF9 | -0,2080948 |
| ENSG00000; | gene | 1,30405878 | 0,03464968 | 0,11177171 | AC007292.1 | 0,3830089 |
| ENSG00000 | gene | 1,49486438 | 0,03468822 | 0,11182655 | EXO1 | 0,5800146 |
| ENSG00000 | gene | 0,8656471 | 0,0346959 | 0,11182655 | MDK | -0,2081491 |

| | | | | | | |
|---|---|---|---|---|---|---|
| ENSG00002 | gene | 1,28189759 | 0,03470051 | 0,11182655 | N4BP2L2 | 0,35828101 |
| ENSG00000 | gene | 0,88718055 | 0,0347227 | 0,11186168 | CFAP410 | -0,1727004 |
| ENSG00000 | gene | 0,79242979 | 0,03477856 | 0,11200526 | ZNF395 | -0,335645 |
| ENSG00000 | gene | 0,74141071 | 0,03481375 | 0,11204992 | CXCL5 | -0,4316551 |
| ENSG00000 | gene | 0,89506348 | 0,03481503 | 0,11204992 | UBALD2 | -0,1599381 |
| ENSG00000 | gene | 1,12373881 | 0,03484603 | 0,1121133 | PSIP1 | 0,16830675 |
| ENSG00000 | gene | 0,94585607 | 0,03485973 | 0,112121 | HNRNPUL1 | -0,0803074 |
| ENSG00000 | gene | 0,92426832 | 0,03491888 | 0,11227485 | PABPC4 | -0,1136164 |
| ENSG00000 | gene | 0,94616834 | 0,0349469 | 0,1123285 | GPKOW | -0,0798312 |
| ENSG00000 | gene | 0,94550142 | 0,03501933 | 0,11252354 | COPZ1 | -0,0808485 |
| ENSG00000 | gene | 0,91826544 | 0,03503985 | 0,11252354 | DLGAP4 | -0,1230168 |
| ENSG00000 | gene | 0,9111935 | 0,03504453 | 0,11252354 | TRADD | -0,1341706 |
| ENSG00000 | gene | 0,87691517 | 0,03505297 | 0,11252354 | NAGK | -0,1894908 |
| ENSG00000 | gene | 0,76440275 | 0,03508979 | 0,11260531 | SERF1A | -0,3875951 |
| ENSG00000 | gene | 1,38369269 | 0,03510608 | 0,11262113 | ZNF276 | 0,46852356 |
| ENSG00000 | gene | 1,11574806 | 0,03512169 | 0,11263474 | ENDOD1 | 0,1580113 |
| ENSG00000 | gene | 0,8920589 | 0,03518239 | 0,11276937 | TIMM13 | -0,1647891 |
| ENSG00002 | gene | 0,91573612 | 0,03518893 | 0,11276937 | MTATP6P1 | -0,1269962 |
| ENSG00002 | gene | 1,25543515 | 0,03519778 | 0,11276937 | AL354919.2 | 0,32818751 |
| ENSG00002 | gene | 1,26656212 | 0,03523634 | 0,11285645 | AC109587.1 | 0,34091783 |
| ENSG00000 | gene | 0,91729374 | 0,03528854 | 0,1129763 | TRIM56 | -0,1245443 |
| ENSG00000 | gene | 1,08226754 | 0,03529655 | 0,1129763 | CLIP1 | 0,11405718 |
| ENSG00000 | gene | 0,8690124 | 0,0353586 | 0,11310544 | RNF24 | -0,2025513 |
| ENSG00000 | gene | 0,84865955 | 0,03537671 | 0,11310544 | SH3D19 | -0,2367422 |
| ENSG00000 | gene | 0,86880846 | 0,035382 | 0,11310544 | CNP | -0,2028899 |
| ENSG00000 | gene | 0,91969928 | 0,03538252 | 0,11310544 | NDUFV2 | -0,1207659 |
| ENSG00000 | gene | 0,88183331 | 0,03545448 | 0,11328688 | RPS19 | -0,1814221 |
| ENSG00000 | gene | 1,11099329 | 0,03546213 | 0,11328688 | PPWD1 | 0,1518501 |
| ENSG00000 | gene | 1,11387095 | 0,03555565 | 0,113498 | SUZ12 | 0,1555821 |
| ENSG00002 | gene | 1,24693916 | 0,03555596 | 0,113498 | VGLL3 | 0,31839108 |
| ENSG00000 | gene | 0,81447656 | 0,03556256 | 0,113498 | SFSWAP | -0,2960549 |
| ENSG00000 | gene | 1,13149061 | 0,03563473 | 0,11369175 | NFE2L3 | 0,17822461 |
| ENSG00000 | gene | 0,85156694 | 0,03565729 | 0,11372716 | TOP3B | -0,2318082 |
| ENSG00000 | gene | 1,11063334 | 0,03569991 | 0,11382645 | POP1 | 0,1513826 |
| ENSG00000 | gene | 0,91411789 | 0,03571746 | 0,11384581 | GAK | -0,1295479 |
| ENSG00000 | gene | 0,93005656 | 0,03573893 | 0,11387766 | PLXND1 | -0,1046096 |
| ENSG00000 | gene | 1,05127226 | 0,03583588 | 0,11414989 | GLUD1 | 0,07213634 |
| ENSG00000 | gene | 0,8104877 | 0,0358558 | 0,11417664 | PPP1R15A | -0,3031378 |
| ENSG00000 | gene | 0,94632924 | 0,03590398 | 0,11428349 | THRA | -0,0795859 |
| ENSG00000 | gene | 1,14399071 | 0,0359124 | 0,11428349 | SMAP2 | 0,19407534 |
| ENSG00000 | gene | 0,75857967 | 0,03594569 | 0,11435272 | BTN3A2 | -0,3986274 |
| ENSG00002 | gene | 1,22815559 | 0,03596684 | 0,1143833 | AC087588.3 | 0,29649335 |
| ENSG00000 | gene | 0,94126225 | 0,03600866 | 0,11447961 | ATXN7L3 | -0,0873314 |
| ENSG00000 | gene | 1,3428416 | 0,03608757 | 0,1146937 | RELN | 0,42528914 |
| ENSG00002 | gene | 1,19447321 | 0,03611607 | 0,11474749 | AC025682.2 | 0,2563745 |
| ENSG00000 | gene | 0,89307301 | 0,03616007 | 0,1148505 | CTSD | -0,16315 |
| ENSG00002 | gene | 1,33193637 | 0,03626155 | 0,11513593 | AL121987.2 | 0,41352517 |
| ENSG00002 | gene | 0,63745355 | 0,03631707 | 0,11526776 | HSPA1B | -0,6496079 |
| ENSG00002 | gene | 0,75474773 | 0,03632632 | 0,11526776 | AL390719.1 | -0,4059336 |
| ENSG00000 | gene | 0,92943164 | 0,03634025 | 0,1152751 | BLVRA | -0,1055793 |
| ENSG00000 | gene | 1,04569285 | 0,03637784 | 0,11535742 | CNOT9 | 0,06445916 |
| ENSG00000 | gene | 1,21721334 | 0,03639709 | 0,11538157 | CHRNB1 | 0,28358205 |
| ENSG00000 | gene | 1,04316083 | 0,03650148 | 0,11567552 | DHX15 | 0,06096161 |
| ENSG00000 | gene | 0,88117132 | 0,0365239 | 0,11570958 | SVBP | -0,1825055 |
| ENSG00000 | gene | 1,40946662 | 0,0366288 | 0,11597578 | SLC8A1 | 0,49514931 |
| ENSG00000 | gene | 0,87198115 | 0,03664686 | 0,11597578 | RPL24P4 | -0,1976311 |
| ENSG00002 | gene | 0,85668104 | 0,03665456 | 0,11597578 | AC009093.8 | -0,2231699 |

| | | | | | | |
|---|---|---|---|---|---|---|
| ENSG00000( | gene | 1,29768069 | 0,03666369 | 0,11597578 | MCM10 | 0,37593544 |
| ENSG00000( | gene | 0,87046729 | 0,03666641 | 0,11597578 | SCD | -0,200138 |
| ENSG00000 | gene | 1,13139684 | 0,03672115 | 0,11601633 | CEP63 | 0,17810505 |
| ENSG00000: | gene | 2,14116551 | 0,03672905 | 0,11601633 | TGIF2-RAB5 | 1,09839632 |
| ENSG00000( | gene | 0,81408612 | 0,0367314 | 0,11601633 | CDV3 | -0,2967467 |
| ENSG00000 | gene | 0,93177232 | 0,0367376 | 0,11601633 | MEIS1 | -0,1019506 |
| ENSG00000 | gene | 0,94078332 | 0,03673919 | 0,11601633 | FAM234A | -0,0880656 |
| ENSG00000( | gene | 0,90815122 | 0,03676031 | 0,11601633 | SLC9A3R2 | -0,1389955 |
| ENSG00000 | gene | 1,04698665 | 0,03676112 | 0,11601633 | CBX6 | 0,06624304 |
| ENSG00000 | gene | 0,93594757 | 0,03686569 | 0,11630932 | FCHSD1 | -0,0955004 |
| ENSG00000 | gene | 0,85525043 | 0,03691868 | 0,11643945 | ZNF205 | -0,2255812 |
| ENSG00000 | gene | 0,93963338 | 0,03693208 | 0,11644467 | SNRPC | -0,0898301 |
| ENSG00000 | gene | 0,94243883 | 0,03698961 | 0,116589 | NOLC1 | -0,0855291 |
| ENSG00000( | gene | 0,92880093 | 0,03706717 | 0,11679634 | RRAGB | -0,1065587 |
| ENSG00000: | gene | 0,76254614 | 0,03720315 | 0,11718757 | PCDHGA12 | -0,3911035 |
| ENSG00000: | gene | 0,89343704 | 0,03736634 | 0,11765833 | AL117190.2 | -0,162562 |
| ENSG00000: | gene | 0,80778093 | 0,03737634 | 0,11765833 | AC068631.4 | -0,307964 |
| ENSG00000: | gene | 1,1473101 | 0,0374105 | 0,11772849 | AC060766.1 | 0,19825538 |
| ENSG00000 | gene | 0,80560542 | 0,03747038 | 0,11787952 | UHMK1 | -0,3118547 |
| ENSG00000 | gene | 0,80318969 | 0,0375025 | 0,11794315 | VKORC1 | -0,3161873 |
| ENSG00000 | gene | 0,86996956 | 0,03751855 | 0,11795213 | GRPEL1 | -0,2009632 |
| ENSG00000 | gene | 0,93431556 | 0,03752914 | 0,11795213 | PPP1R9B | -0,0980182 |
| ENSG00000: | gene | 1,2482031 | 0,0375428 | 0,11795767 | TMED2-DT | 0,3198527 |
| ENSG00000: | gene | 0,74543009 | 0,03763822 | 0,11818638 | AC018804.1 | -0,423855 |
| ENSG00000: | gene | 0,8872543 | 0,03764284 | 0,11818638 | AP000943.2 | -0,1725804 |
| ENSG00000: | gene | 0,8986801 | 0,03765135 | 0,11818638 | ATP5MF | -0,1541204 |
| ENSG00000 | gene | 0,91672526 | 0,0376805 | 0,11824044 | BAIAP2 | -0,1254387 |
| ENSG00000 | gene | 0,66002085 | 0,03769925 | 0,11826186 | CXCL3 | -0,5994165 |
| ENSG00000 | gene | 0,93037035 | 0,03778736 | 0,11850078 | CCT4 | -0,104123 |
| ENSG00000 | gene | 1,27854177 | 0,03781947 | 0,11856399 | TMSB4XP8 | 0,35449929 |
| ENSG00000( | gene | 0,76837842 | 0,03789525 | 0,11876401 | ICAM1 | -0,3801111 |
| ENSG00000: | gene | 1,29012272 | 0,03791295 | 0,11878193 | AC025257.1 | 0,3675083 |
| ENSG00000: | gene | 1,36103859 | 0,03792553 | 0,11878381 | MIR4292 | 0,44470797 |
| ENSG00000( | gene | 1,16115477 | 0,03797785 | 0,11891013 | MCM6 | 0,21556029 |
| ENSG00000 | gene | 0,70483424 | 0,03800099 | 0,11892236 | HNRNPA1L2 | -0,5046441 |
| ENSG00000 | gene | 0,80743162 | 0,03801224 | 0,11892236 | SNRK | -0,308588 |
| ENSG00000 | gene | 0,89636302 | 0,03801774 | 0,11892236 | RTN2 | -0,157845 |
| ENSG00000 | gene | 0,81182065 | 0,03804261 | 0,11892507 | QKI | -0,3007671 |
| ENSG00000: | gene | 0,75362345 | 0,03805087 | 0,11892507 | AC092691.1 | -0,4080842 |
| ENSG00000 | gene | 1,08429199 | 0,03805458 | 0,11892507 | TCAF1 | 0,11675332 |
| ENSG00000 | gene | 0,92379224 | 0,03809389 | 0,11901039 | RHEB | -0,1143597 |
| ENSG00000 | gene | 1,11797637 | 0,03812157 | 0,11905937 | PRIM2 | 0,16088969 |
| ENSG00000 | gene | 0,90441572 | 0,03817891 | 0,11920091 | ERF | -0,144942 |
| ENSG00000: | gene | 1,34452404 | 0,03830008 | 0,1195348 | AC244669.1 | 0,42709555 |
| ENSG00000 | gene | 0,83402419 | 0,03830996 | 0,1195348 | PFKFB4 | -0,2618389 |
| ENSG00000 | gene | 0,80164009 | 0,03833826 | 0,11958545 | ISYNA1 | -0,3189734 |
| ENSG00000 | gene | 0,88366564 | 0,03835306 | 0,11959151 | RNF41 | -0,1784275 |
| ENSG00000 | gene | 0,89427604 | 0,03836432 | 0,11959151 | NDUFB5 | -0,1612079 |
| ENSG00000 | gene | 0,80482893 | 0,03844253 | 0,11979765 | RPL7L1 | -0,3132459 |
| ENSG00000( | gene | 0,90615319 | 0,03854803 | 0,12008868 | OGFR | -0,1421731 |
| ENSG00000 | gene | 0,95659926 | 0,03856233 | 0,12009549 | GART | -0,0640134 |
| ENSG00000 | gene | 0,77168284 | 0,03858972 | 0,12012292 | RNASEH2B | -0,3739201 |
| ENSG00000( | gene | 1,11352233 | 0,03860129 | 0,12012292 | MAP4K3 | 0,15513049 |
| ENSG00000 | gene | 0,75474819 | 0,03861461 | 0,12012292 | KLF4 | -0,4059327 |
| ENSG00000 | gene | 0,79137892 | 0,03861959 | 0,12012292 | PRNP | -0,3375595 |
| ENSG00000 | gene | 0,90647205 | 0,03867876 | 0,12026923 | AEBP1 | -0,1416656 |
| ENSG00000: | gene | 0,60534207 | 0,03869906 | 0,12029463 | Z83844.3 | -0,7241775 |

| | | | | | | |
|---|---|---|---|---|---|---|
| ENSG00000 | gene | 0,90479737 | 0,03872509 | 0,12033783 | LEMD2 | -0,1443334 |
| ENSG00000( | gene | 1,17192495 | 0,03888959 | 0,12080318 | REX1BD | 0,22888018 |
| ENSG00000 | gene | 0,93919154 | 0,03889921 | 0,12080318 | AIFM1 | -0,0905087 |
| ENSG00000 | gene | 1,10785839 | 0,03897849 | 0,1210115 | DCUN1D4 | 0,14777348 |
| ENSG00000( | gene | 0,93993825 | 0,03904153 | 0,12111982 | SLC25A5 | -0,0893621 |
| ENSG00000: | gene | 0,92895747 | 0,03904238 | 0,12111982 | RHEBP2 | -0,1063155 |
| ENSG00000: | gene | 0,84728345 | 0,03905003 | 0,12111982 | CKS1BP3 | -0,2390834 |
| ENSG00000 | gene | 0,88767667 | 0,0390896 | 0,12120464 | RPL5 | -0,1718938 |
| ENSG00000 | gene | 1,12724143 | 0,03919326 | 0,12148808 | RHOF | 0,17279655 |
| ENSG00000 | gene | 0,67822528 | 0,03930989 | 0,12181153 | NR4A2 | -0,5601635 |
| ENSG00000 | gene | 0,6374103 | 0,0393346 | 0,12183514 | GPRASP2 | -0,6497058 |
| ENSG00000( | gene | 0,87749848 | 0,03934208 | 0,12183514 | FKBP4 | -0,1885315 |
| ENSG00000( | gene | 0,91342191 | 0,03937159 | 0,12188846 | LUC7L | -0,1306467 |
| ENSG00000: | gene | 1,11820129 | 0,03942233 | 0,12200745 | AC100793.3 | 0,16117992 |
| ENSG00000 | gene | 0,86617713 | 0,03946201 | 0,12209212 | COX16 | -0,207266 |
| ENSG00000 | gene | 0,79415634 | 0,03952028 | 0,12223429 | ZBTB46 | -0,332505 |
| ENSG00000 | gene | 1,08145138 | 0,03955211 | 0,12229457 | COX5A | 0,1129688 |
| ENSG00000 | gene | 0,93785611 | 0,03958282 | 0,12235139 | IRAK1 | -0,0925615 |
| ENSG00000 | gene | 0,81825148 | 0,0396825 | 0,12262127 | SETD5 | -0,2893838 |
| ENSG00000 | gene | 0,94886945 | 0,03971626 | 0,12264938 | MRPL32 | -0,0757185 |
| ENSG00000: | gene | 1,3086629 | 0,03971633 | 0,12264938 | IFNWP19 | 0,38809352 |
| ENSG00000: | gene | 0,87886809 | 0,03973305 | 0,12266281 | SNHG6 | -0,1862815 |
| ENSG00000 | gene | 1,16801102 | 0,03978886 | 0,12279308 | ARHGAP5 | 0,22405389 |
| ENSG00000( | gene | 0,87764184 | 0,03980002 | 0,12279308 | POLB | -0,1882958 |
| ENSG00000 | gene | 0,76408629 | 0,03984362 | 0,12288936 | LDAH | -0,3881925 |
| ENSG00000 | gene | 1,09149188 | 0,03996993 | 0,12321997 | TGFBR2 | 0,1263014 |
| ENSG00000 | gene | 1,27998165 | 0,03997566 | 0,12321997 | TRIM55 | 0,35612313 |
| ENSG00000 | gene | 0,92385727 | 0,03999081 | 0,12322837 | TLK2 | -0,1142581 |
| ENSG00000 | gene | 0,81200487 | 0,04005991 | 0,12338598 | DCTPP1 | -0,3004397 |
| ENSG00000( | gene | 0,93487646 | 0,04007611 | 0,12338598 | PAF1 | -0,0971524 |
| ENSG00000: | gene | 0,9206087 | 0,04009008 | 0,12338598 | LCMT1 | -0,11934 |
| ENSG00000 | gene | 0,93761689 | 0,040093 | 0,12338598 | ENTR1 | -0,0929295 |
| ENSG00000: | gene | 0,74315256 | 0,04011322 | 0,12338598 | AP003108.2 | -0,4282697 |
| ENSG00000 | gene | 1,12668313 | 0,04011662 | 0,12338598 | SCP2 | 0,17208182 |
| ENSG00000 | gene | 0,85824177 | 0,04013933 | 0,12341479 | RPL23A | -0,220544 |
| ENSG00000 | gene | 0,95309071 | 0,04015088 | 0,12341479 | TOB2 | -0,0693146 |
| ENSG00000 | gene | 1,11957004 | 0,04016573 | 0,12342217 | ZDHHC20 | 0,16294478 |
| ENSG00000 | gene | 0,89833656 | 0,04021796 | 0,12354439 | FLCN | -0,154672 |
| ENSG00000 | gene | 1,15211897 | 0,04025716 | 0,12362649 | GOLGA4 | 0,2042897 |
| ENSG00000 | gene | 0,92878542 | 0,04033866 | 0,12383842 | SURF1 | -0,1065828 |
| ENSG00000 | gene | 1,14884637 | 0,0403801 | 0,1239273 | GBP3 | 0,20018588 |
| ENSG00000( | gene | 1,20725875 | 0,04043029 | 0,12399719 | ZNF302 | 0,27173492 |
| ENSG00000: | gene | 1,22026728 | 0,04043186 | 0,12399719 | AC131238.1 | 0,28719718 |
| ENSG00000: | gene | 0,82814211 | 0,04044039 | 0,12399719 | SPCS2P4 | -0,2720497 |
| ENSG00000: | gene | 1,38340094 | 0,04046978 | 0,12403026 | AC125494.1 | 0,46821934 |
| ENSG00000( | gene | 1,08445872 | 0,04047619 | 0,12403026 | MTIF2 | 0,11697514 |
| ENSG00000 | gene | 0,92142426 | 0,04049595 | 0,12405247 | GSS | -0,1180625 |
| ENSG00000 | gene | 0,81646602 | 0,04060826 | 0,1243581 | DPH1 | -0,2925352 |
| ENSG00000 | gene | 0,92994175 | 0,04072631 | 0,12468112 | MRPL18 | -0,1047877 |
| ENSG00000 | gene | 0,90550846 | 0,04077572 | 0,12479385 | CCDC59 | -0,1432 |
| ENSG00000 | gene | 0,94264194 | 0,04080164 | 0,12483465 | CXorf40B | -0,0852182 |
| ENSG00000 | gene | 0,93642688 | 0,04086533 | 0,12499092 | JAGN1 | -0,0947617 |
| ENSG00000 | gene | 1,28962618 | 0,04093929 | 0,12517853 | IRGQ | 0,36695294 |
| ENSG00000( | gene | 0,93044484 | 0,04096978 | 0,12523314 | PSEN1 | -0,1040075 |
| ENSG00000 | gene | 0,89696696 | 0,0410019 | 0,12529271 | ARHGEF18 | -0,1568733 |
| ENSG00000 | gene | 0,85848365 | 0,04105281 | 0,12540963 | ATPAF2 | -0,2201374 |
| ENSG00000 | gene | 0,89727577 | 0,04108042 | 0,12545532 | SH3BP5 | -0,1563766 |

| | | | | | | |
|---|---|---|---|---|---|---|
| ENSG00000( | gene | 0,94863261 | 0,04111955 | 0,12553337 | LSG1 | -0,0760786 |
| ENSG00000' | gene | 0,89743018 | 0,04113129 | 0,12553337 | KIFC3 | -0,1561284 |
| ENSG00000' | gene | 0,85768231 | 0,0411941 | 0,12568636 | TMEM184B | -0,2214847 |
| ENSG00000' | gene | 0,92073846 | 0,04121052 | 0,12569779 | TRIM28 | -0,1191367 |
| ENSG00000( | gene | 0,87875301 | 0,04132839 | 0,12600722 | CPOX | -0,1864704 |
| ENSG00000' | gene | 1,17353203 | 0,04133738 | 0,12600722 | NEK7 | 0,23085722 |
| ENSG00000' | gene | 1,19050615 | 0,04138875 | 0,12612501 | MYOCD | 0,25157507 |
| ENSG00000' | gene | 1,21934213 | 0,04141615 | 0,12616976 | SHCBP1 | 0,28610298 |
| ENSG00000( | gene | 0,96094392 | 0,04147896 | 0,1263223 | TFE3 | -0,0574759 |
| ENSG00000: | gene | 0,82623144 | 0,04152444 | 0,12638562 | CHMP1B-AS | -0,2753821 |
| ENSG00000' | gene | 0,82121785 | 0,04152968 | 0,12638562 | IMPA2 | -0,2841631 |
| ENSG00000: | gene | 1,39897813 | 0,04153799 | 0,12638562 | AL645941.3 | 0,4843734 |
| ENSG00000( | gene | 0,86959203 | 0,04157684 | 0,12646502 | SLC35C2 | -0,2015894 |
| ENSG00000' | gene | 1,07273438 | 0,04165685 | 0,12666952 | COL6A3 | 0,10129289 |
| ENSG00000' | gene | 0,87943932 | 0,04167655 | 0,12669058 | OTUB1 | -0,1853441 |
| ENSG00000' | gene | 1,09148462 | 0,04172219 | 0,12673048 | FARSB | 0,12629181 |
| ENSG00000' | gene | 0,84558168 | 0,04172718 | 0,12673048 | CD55 | -0,241984 |
| ENSG00000: | gene | 1,15043383 | 0,04172802 | 0,12673048 | APOBEC3C | 0,202178 |
| ENSG00000' | gene | 0,9136317 | 0,04187135 | 0,12712686 | ATP5IF1 | -0,1303154 |
| ENSG00000: | gene | 1,33631398 | 0,04188479 | 0,12712874 | AC022167.1 | 0,41825902 |
| ENSG00000' | gene | 0,89097026 | 0,04197574 | 0,12736579 | B4GALT4 | -0,1665508 |
| ENSG00000' | gene | 0,84870313 | 0,04202302 | 0,12747024 | SPSB1 | -0,2366681 |
| ENSG00000: | gene | 1,5496548 | 0,04205797 | 0,12752916 | AC006441.3 | 0,63194688 |
| ENSG00000( | gene | 0,89997383 | 0,04206816 | 0,12752916 | HERPUD1 | -0,152045 |
| ENSG00000: | gene | 1,61081329 | 0,0420844 | 0,12753941 | AC015674.1 | 0,68778928 |
| ENSG00000' | gene | 0,95631372 | 0,0421333 | 0,12764859 | CDK16 | -0,0644441 |
| ENSG00000' | gene | 0,84497518 | 0,04234251 | 0,12824323 | SPG7 | -0,2430191 |
| ENSG00000' | gene | 0,91013606 | 0,04236235 | 0,12826414 | PRKCD | -0,1358459 |
| ENSG00000: | gene | 0,83109157 | 0,04241462 | 0,12838321 | AP000944.3 | -0,2669207 |
| ENSG00000' | gene | 1,08236026 | 0,04245196 | 0,12845701 | TNKS2 | 0,11418078 |
| ENSG00000' | gene | 0,94811886 | 0,04248519 | 0,12851834 | TMCO1 | -0,0768602 |
| ENSG00000' | gene | 0,81713557 | 0,04254165 | 0,1286499 | AFMID | -0,2913526 |
| ENSG00000' | gene | 0,88698194 | 0,04257031 | 0,12867333 | SRGAP1 | -0,1730234 |
| ENSG00000' | gene | 0,88979565 | 0,04259138 | 0,12867333 | MAGED4B | -0,168454 |
| ENSG00000: | gene | 0,94930486 | 0,04260963 | 0,12867333 | RING1 | -0,0750566 |
| ENSG00000( | gene | 0,94217663 | 0,04261149 | 0,12867333 | TSG101 | -0,0859306 |
| ENSG00000: | gene | 0,77835076 | 0,04261428 | 0,12867333 | AL049873.1 | -0,3615076 |
| ENSG00000' | gene | 1,0516408 | 0,04263775 | 0,12870499 | KIAA0232 | 0,07264202 |
| ENSG00000' | gene | 0,93900743 | 0,04269083 | 0,12874782 | SUMO2 | -0,0907915 |
| ENSG00000: | gene | 0,81400956 | 0,04270279 | 0,12874782 | AC091045.2 | -0,2968824 |
| ENSG00000( | gene | 0,66024527 | 0,04270519 | 0,12874782 | IL11 | -0,598926 |
| ENSG00000' | gene | 0,9209278 | 0,04270879 | 0,12874782 | PEX5 | -0,11884 |
| ENSG00000' | gene | 0,85674311 | 0,04271685 | 0,12874782 | RPSAP19 | -0,2230654 |
| ENSG00000' | gene | 1,10375276 | 0,04278962 | 0,12892794 | NIPSNAP3A | 0,14241705 |
| ENSG00000' | gene | 0,87831984 | 0,04285867 | 0,12909678 | MEF2D | -0,1871817 |
| ENSG00000: | gene | 0,71339771 | 0,04293984 | 0,129302 | AC114490.2 | -0,4872215 |
| ENSG00000: | gene | 1,21253223 | 0,04302688 | 0,12948678 | ELOA-AS1 | 0,27802309 |
| ENSG00000' | gene | 0,86450872 | 0,04302732 | 0,12948678 | SRSF2 | -0,2100476 |
| ENSG00000' | gene | 0,94708926 | 0,04310076 | 0,12964004 | TAOK2 | -0,0784277 |
| ENSG00000' | gene | 0,95889052 | 0,0431044 | 0,12964004 | APMAP | -0,060562 |
| ENSG00000' | gene | 1,29444531 | 0,04315325 | 0,1297476 | HARBI1 | 0,37233401 |
| ENSG00000' | gene | 0,8014385 | 0,0432002 | 0,1298494 | GOSR2 | -0,3193363 |
| ENSG00000' | gene | 0,89518569 | 0,04322764 | 0,1298925 | SLC6A8 | -0,1597411 |
| ENSG00000: | gene | 0,82070329 | 0,0432714 | 0,12998462 | AC098614.3 | -0,2850674 |
| ENSG00000' | gene | 1,42286675 | 0,04329351 | 0,13001164 | RIMKLB | 0,50880057 |
| ENSG00000( | gene | 1,17209251 | 0,04338761 | 0,13025478 | TRAM1 | 0,22908645 |
| ENSG00000: | gene | 0,89048342 | 0,04344022 | 0,13037325 | TCTEX1D2 | -0,1673394 |

| | | | | | | |
|---|---|---|---|---|---|---|
| ENSG00000 | gene | 0,90309486 | 0,04348947 | 0,13048157 | PRR13P5 | -0,1470506 |
| ENSG00000 | gene | 0,89268092 | 0,04359611 | 0,13074046 | CSNK2A1 | -0,1637835 |
| ENSG00000 | gene | 0,87287674 | 0,04360486 | 0,13074046 | LURAP1L | -0,1961501 |
| ENSG00000 | gene | 1,54814319 | 0,04361543 | 0,13074046 | AC010542.2 | 0,63053891 |
| ENSG00000 | gene | 1,11405071 | 0,04364236 | 0,13074046 | FLNA | 0,1558149 |
| ENSG00000 | gene | 1,10957182 | 0,04364511 | 0,13074046 | MEIS2 | 0,15000306 |
| ENSG00000 | gene | 0,88156303 | 0,04365487 | 0,13074046 | COPS9 | -0,1818644 |
| ENSG00000 | gene | 0,84169928 | 0,04370344 | 0,13084152 | TMEM132A | -0,2486232 |
| ENSG00000 | gene | 0,80474307 | 0,04371841 | 0,13084152 | DNAJC15 | -0,3133999 |
| ENSG00000 | gene | 0,93124804 | 0,04373335 | 0,13084152 | PELP1 | -0,1027626 |
| ENSG00000 | gene | 1,40275077 | 0,04374139 | 0,13084152 | PLEKHG5 | 0,48825871 |
| ENSG00000 | gene | 0,73955448 | 0,04386623 | 0,13114482 | RBM38 | -0,4352717 |
| ENSG00000 | gene | 0,86207773 | 0,04387342 | 0,13114482 | GRINA | -0,2141101 |
| ENSG00000 | gene | 0,8160343 | 0,04388986 | 0,13114482 | ZFYVE27 | -0,2932983 |
| ENSG00000 | gene | 1,12624709 | 0,04389569 | 0,13114482 | RWDD4 | 0,17152338 |
| ENSG00000 | gene | 1,20794608 | 0,04393933 | 0,13123565 | KIT | 0,27255606 |
| ENSG00000 | gene | 0,77606788 | 0,04404778 | 0,13151995 | CEP290 | -0,3657452 |
| ENSG00000 | gene | 1,04858399 | 0,04406942 | 0,13154496 | RARS1 | 0,06844242 |
| ENSG00000 | gene | 0,91130802 | 0,04414958 | 0,13170625 | EXOSC5 | -0,1339893 |
| ENSG00000 | gene | 1,19895464 | 0,04415002 | 0,13170625 | MAP3K20 | 0,26177708 |
| ENSG00000 | gene | 2,23305983 | 0,04419928 | 0,13181356 | AC023509.1 | 1,1590219 |
| ENSG00000 | gene | 1,41995787 | 0,04421744 | 0,13182806 | OXTR | 0,50584813 |
| ENSG00000 | gene | 1,17918088 | 0,04432636 | 0,13211307 | H2AX | 0,23778503 |
| ENSG00000 | gene | 0,92716553 | 0,04435542 | 0,13212655 | SLC35B1 | -0,1091012 |
| ENSG00000 | gene | 0,81423491 | 0,04435753 | 0,13212655 | TMEM184A | -0,296483 |
| ENSG00000 | gene | 0,74628016 | 0,04437539 | 0,13214007 | SGCE | -0,4222108 |
| ENSG00000 | gene | 0,92060013 | 0,04442976 | 0,13226223 | HOXB6 | -0,1193535 |
| ENSG00000 | gene | 1,11192475 | 0,04448087 | 0,13234061 | NDC1 | 0,15305915 |
| ENSG00000 | gene | 1,06410704 | 0,044495 | 0,13234061 | MTDH | 0,08964328 |
| ENSG00000 | gene | 0,90075799 | 0,04449613 | 0,13234061 | SRC | -0,1507886 |
| ENSG00000 | gene | 1,0523563 | 0,04453041 | 0,13238246 | UBE2J1 | 0,07362325 |
| ENSG00000 | gene | 0,92112075 | 0,04454584 | 0,13238246 | PPP1R7 | -0,1185378 |
| ENSG00000 | gene | 1,11499085 | 0,04455025 | 0,13238246 | WDR33 | 0,15703187 |
| ENSG00000 | gene | 0,89656895 | 0,04461195 | 0,13249993 | ADAR | -0,1575136 |
| ENSG00000 | gene | 0,76246115 | 0,0446165 | 0,13249993 | U2AF1L4 | -0,3912643 |
| ENSG00000 | gene | 0,91452195 | 0,04467149 | 0,13262349 | OSER1 | -0,1289103 |
| ENSG00000 | gene | 0,89943458 | 0,04470991 | 0,13269784 | SLC66A2 | -0,1529097 |
| ENSG00000 | gene | 1,24034993 | 0,04477238 | 0,13284348 | GCC2 | 0,3107472 |
| ENSG00000 | gene | 1,11888889 | 0,04485348 | 0,13300696 | GALNT6 | 0,16206677 |
| ENSG00000 | gene | 0,81671587 | 0,0448543 | 0,13300696 | CSRNP1 | -0,2920938 |
| ENSG00000 | gene | 0,83144145 | 0,04490512 | 0,13311783 | ZFHX2-AS1 | -0,2663134 |
| ENSG00000 | gene | 1,0697741 | 0,04505882 | 0,13353356 | DCTN6 | 0,09730618 |
| ENSG00000 | gene | 0,81439576 | 0,04514261 | 0,13368325 | TP53TG1 | -0,296198 |
| ENSG00000 | gene | 0,90742131 | 0,04514528 | 0,13368325 | FXR2 | -0,1401555 |
| ENSG00000 | gene | 1,07955553 | 0,04514978 | 0,13368325 | UBXN4 | 0,11043745 |
| ENSG00000 | gene | 0,86368532 | 0,04519792 | 0,133773 | C1orf109 | -0,2114223 |
| ENSG00000 | gene | 1,21576207 | 0,04520707 | 0,133773 | AC025043.1 | 0,28186091 |
| ENSG00000 | gene | 1,43311862 | 0,04528545 | 0,13396495 | MIR210HG | 0,51915802 |
| ENSG00000 | gene | 1,12440702 | 0,04552032 | 0,13459511 | RNF141 | 0,16916437 |
| ENSG00000 | gene | 1,0631271 | 0,04552562 | 0,13459511 | PRKD3 | 0,08831408 |
| ENSG00000 | gene | 0,88505926 | 0,04557927 | 0,13471358 | HSPA8 | -0,176154 |
| ENSG00000 | gene | 0,85036304 | 0,04560764 | 0,13473582 | JTB | -0,2338492 |
| ENSG00000 | gene | 0,81963309 | 0,04561397 | 0,13473582 | NIFK-AS1 | -0,2869499 |
| ENSG00000 | gene | 1,0787528 | 0,04564481 | 0,13478676 | CHD3 | 0,10936431 |
| ENSG00000 | gene | 0,87411366 | 0,04569132 | 0,13483122 | PSENEN | -0,1941072 |
| ENSG00000 | gene | 1,31398438 | 0,04569881 | 0,13483122 | AC135050.7 | 0,39394812 |
| ENSG00000 | gene | 1,03914876 | 0,04570066 | 0,13483122 | ACTR8 | 0,0554022 |

| | | | | | | |
|---|---|---|---|---|---|---|
| ENSG00000' | gene | 0,9277288 | 0,04574064 | 0,13488265 | DIP2A | -0,108225 |
| ENSG00000' | gene | 0,88303853 | 0,0457453 | 0,13488265 | RNPEPL1 | -0,1794517 |
| ENSG00000( | gene | 0,92957015 | 0,0460271 | 0,13567323 | TOLLIP | -0,1053644 |
| ENSG00000' | gene | 0,94672093 | 0,04613587 | 0,13595342 | ILF2 | -0,0789889 |
| ENSG00000' | gene | 1,22456689 | 0,04620286 | 0,1361104 | RPRD2 | 0,29227158 |
| ENSG00000; | gene | 1,21154985 | 0,04628987 | 0,1363107 | AF111169.3 | 0,27685377 |
| ENSG00000' | gene | 0,913454 | 0,04629835 | 0,1363107 | YKT6 | -0,130596 |
| ENSG00000; | gene | 0,88124594 | 0,04640106 | 0,13657255 | AC016394.4 | -0,1823834 |
| ENSG00000; | gene | 0,77125026 | 0,04656929 | 0,13702703 | AC011979.1 | -0,374729 |
| ENSG00000' | gene | 0,87736371 | 0,04659858 | 0,13702784 | KLHDC4 | -0,1887531 |
| ENSG00000' | gene | 0,85881048 | 0,04660231 | 0,13702784 | ATP6V0C | -0,2195883 |
| ENSG00000' | gene | 0,89718121 | 0,04661102 | 0,13702784 | SNRPG | -0,1565287 |
| ENSG00000( | gene | 1,23685598 | 0,04665445 | 0,13711312 | DIS3 | 0,30667753 |
| ENSG00000' | gene | 1,07696779 | 0,04667036 | 0,13711312 | CCM2 | 0,1069751 |
| ENSG00000' | gene | 0,4770728 | 0,04668923 | 0,13711312 | AL365205.1 | -1,0677187 |
| ENSG00000' | gene | 1,0822174 | 0,04669534 | 0,13711312 | HNRNPR | 0,11399034 |
| ENSG00000' | gene | 0,88989572 | 0,04678309 | 0,1373301 | ARFRP1 | -0,1682918 |
| ENSG00000' | gene | 0,84572732 | 0,04681849 | 0,13736457 | NIPSNAP1 | -0,2417355 |
| ENSG00000' | gene | 1,12531497 | 0,04682254 | 0,13736457 | RB1 | 0,17032887 |
| ENSG00000' | gene | 1,55996021 | 0,04686114 | 0,13740802 | UCHL5 | 0,64150923 |
| ENSG00000; | gene | 0,8029582 | 0,04686506 | 0,13740802 | SOCS7 | -0,3166032 |
| ENSG00000' | gene | 0,93242969 | 0,04690909 | 0,13749648 | S100A10 | -0,1009332 |
| ENSG00000( | gene | 0,94552319 | 0,04693615 | 0,13753513 | TMEM101 | -0,0808153 |
| ENSG00000( | gene | 0,9201321 | 0,0469819 | 0,13760533 | SF3A1 | -0,1200871 |
| ENSG00000' | gene | 1,07741088 | 0,04698786 | 0,13760533 | ECD | 0,10756854 |
| ENSG00000' | gene | 0,90116771 | 0,04700363 | 0,13761087 | TAF6L | -0,1501325 |
| ENSG00000' | gene | 1,32009029 | 0,04703616 | 0,13766544 | F3 | 0,40063661 |
| ENSG00000' | gene | 1,06583873 | 0,04708275 | 0,13776114 | SNX14 | 0,09198916 |
| ENSG00000' | gene | 0,81934164 | 0,047152 | 0,13792308 | PHKG2 | -0,287463 |
| ENSG00000' | gene | 0,89328617 | 0,04727856 | 0,13821631 | RPL34 | -0,1628057 |
| ENSG00000' | gene | 0,86815729 | 0,04728013 | 0,13821631 | DTX3 | -0,2039716 |
| ENSG00000( | gene | 1,35858795 | 0,04734177 | 0,13835572 | ERCC8 | 0,44210796 |
| ENSG00000' | gene | 0,92416503 | 0,04735593 | 0,13835632 | SNRPA1 | -0,1137776 |
| ENSG00000( | gene | 0,6810115 | 0,04743992 | 0,13854758 | CXCL2 | -0,5542489 |
| ENSG00000' | gene | 0,85569981 | 0,0474626 | 0,13854758 | UNC5B | -0,2248233 |
| ENSG00000( | gene | 1,13908352 | 0,04747146 | 0,13854758 | WWTR1 | 0,18787353 |
| ENSG00000' | gene | 0,92360862 | 0,04747728 | 0,13854758 | LOXL2 | -0,1146464 |
| ENSG00000' | gene | 1,21039696 | 0,0475189 | 0,13862826 | MYPN | 0,27548027 |
| ENSG00000' | gene | 0,86775974 | 0,04762968 | 0,13891055 | SYDE1 | -0,2046324 |
| ENSG00000' | gene | 0,90808163 | 0,04768301 | 0,1390067 | CYTH2 | -0,1391061 |
| ENSG00000( | gene | 1,13524985 | 0,04769068 | 0,1390067 | SLK | 0,18300984 |
| ENSG00000' | gene | 1,74762462 | 0,04771978 | 0,13905064 | UBXN11 | 0,80539533 |
| ENSG00000' | gene | 0,89702005 | 0,04775167 | 0,13910268 | NACC2 | -0,1567879 |
| ENSG00000; | gene | 1,14479514 | 0,04777844 | 0,1391398 | C4orf46 | 0,19508945 |
| ENSG00000; | gene | 0,88175291 | 0,04792451 | 0,13952421 | RPS18 | -0,1815537 |
| ENSG00000; | gene | 1,09131878 | 0,04794186 | 0,13953374 | AC084024.3 | 0,12607259 |
| ENSG00000' | gene | 1,09176907 | 0,04796696 | 0,13956585 | PTEN | 0,12666774 |
| ENSG00000' | gene | 0,48179974 | 0,04799867 | 0,13961714 | RAB4B-EGL | -1,0534945 |
| ENSG00000' | gene | 0,85283686 | 0,04807501 | 0,13979817 | DUSP22 | -0,2296583 |
| ENSG00000; | gene | 0,84707306 | 0,04810581 | 0,13982787 | AC010642.2 | -0,2394417 |
| ENSG00000( | gene | 1,17224777 | 0,04811342 | 0,13982787 | DHX8 | 0,22927753 |
| ENSG00000; | gene | 0,75239834 | 0,04817724 | 0,13994088 | RPS10-NUD | -0,4104314 |
| ENSG00000' | gene | 1,18539462 | 0,04818053 | 0,13994088 | PBK | 0,24536741 |
| ENSG00000' | gene | 0,88446807 | 0,0482366 | 0,14002193 | MIB2 | -0,177118 |
| ENSG00000' | gene | 0,77151665 | 0,04823668 | 0,14002193 | HSPA6 | -0,3742308 |
| ENSG00000; | gene | 1,20907713 | 0,04835171 | 0,14027785 | AC110597.3 | 0,27390629 |
| ENSG00000; | gene | 1,1959353 | 0,04839255 | 0,14027785 | AC022400.1 | 0,25813934 |

| | | | | | | |
|---|---|---|---|---|---|---|
| ENSG00000 | gene | 1,2880073 | 0,04839342 | 0,14027785 | GDF5OS | 0,36514077 |
| ENSG00000 | gene | 0,92511722 | 0,04839485 | 0,14027785 | RBM42 | -0,1122919 |
| ENSG00000 | gene | 1,13813428 | 0,04839558 | 0,14027785 | SCRN3 | 0,18667078 |
| ENSG00000 | gene | 1,41768559 | 0,04845985 | 0,1404231 | AC008738.7 | 0,50353761 |
| ENSG00000 | gene | 0,93919405 | 0,0484809 | 0,14044308 | KHDRBS1 | -0,0905048 |
| ENSG00000 | gene | 1,16338434 | 0,04850374 | 0,14045921 | SPTY2D1OS | 0,21832778 |
| ENSG00000 | gene | 0,8971035 | 0,0485148 | 0,14045921 | DGCR8 | -0,1566536 |
| ENSG00000 | gene | 1,0796249 | 0,04853455 | 0,14047537 | CPNE3 | 0,11053015 |
| ENSG00000 | gene | 1,10190467 | 0,04855399 | 0,14049063 | PLXNB3 | 0,13999942 |
| ENSG00000 | gene | 0,962333 | 0,04857269 | 0,14050373 | MCM3AP | -0,0553919 |
| ENSG00000 | gene | 0,92915664 | 0,04867868 | 0,14076926 | EIF2D | -0,1060063 |
| ENSG00000 | gene | 0,66763706 | 0,04869632 | 0,14077922 | AC020934.2 | -0,5828641 |
| ENSG00000 | gene | 0,94255896 | 0,04876733 | 0,14093621 | RAPGEF1 | -0,0853452 |
| ENSG00000 | gene | 0,942567 | 0,04881565 | 0,14093621 | EIF4A3 | -0,0853329 |
| ENSG00000 | gene | 1,17804023 | 0,04881668 | 0,14093621 | YES1 | 0,23638881 |
| ENSG00000 | gene | 1,23119651 | 0,04882023 | 0,14093621 | RBM45 | 0,30006105 |
| ENSG00000 | gene | 1,18539061 | 0,04882169 | 0,14093621 | AC074044.1 | 0,24536253 |
| ENSG00000 | gene | 0,75851906 | 0,04891048 | 0,14114545 | MYC | -0,3987427 |
| ENSG00000 | gene | 1,05726162 | 0,04892808 | 0,14114545 | EIF2AK4 | 0,08033242 |
| ENSG00000 | gene | 1,20749534 | 0,04893688 | 0,14114545 | PRR11 | 0,27201762 |
| ENSG00000 | gene | 0,95815118 | 0,04898758 | 0,1412506 | SRPRA | -0,0616748 |
| ENSG00000 | gene | 0,75059583 | 0,04900823 | 0,14126908 | AC073896.2 | -0,4138918 |
| ENSG00000 | gene | 1,06125982 | 0,04913574 | 0,14159546 | NUP50 | 0,0857779 |
| ENSG00000 | gene | 1,09930739 | 0,04915744 | 0,14161685 | PRKACA | 0,13659485 |
| ENSG00000 | gene | 0,90313635 | 0,04920529 | 0,14169845 | PANK4 | -0,1469843 |
| ENSG00000 | gene | 1,18014298 | 0,04921435 | 0,14169845 | DNAJB14 | 0,23896165 |
| ENSG00000 | gene | 1,13113646 | 0,04926075 | 0,14179087 | PON2 | 0,17777299 |
| ENSG00000 | gene | 1,12610893 | 0,04945732 | 0,14228791 | RAPGEF2 | 0,17134639 |
| ENSG00000 | gene | 1,05785348 | 0,04946212 | 0,14228791 | LSM12 | 0,08113981 |
| ENSG00000 | gene | 0,92164163 | 0,04954395 | 0,1424565 | SRXN1 | -0,1177222 |
| ENSG00000 | gene | 0,91685508 | 0,04954946 | 0,1424565 | ATP5MC3 | -0,1252344 |
| ENSG00000 | gene | 0,84824752 | 0,04959033 | 0,14252866 | GLIPR2 | -0,2374428 |
| ENSG00000 | gene | 0,94557973 | 0,04960331 | 0,14252866 | STAU1 | -0,080729 |
| ENSG00000 | gene | 1,22647294 | 0,04963329 | 0,14257349 | PRC1-AS1 | 0,2945154 |
| ENSG00000 | gene | 0,92375846 | 0,04967207 | 0,14264356 | POLR3K | -0,1144124 |
| ENSG00000 | gene | 0,88318681 | 0,04974429 | 0,14279774 | RPL18 | -0,1792095 |
| ENSG00000 | gene | 0,93089439 | 0,04975456 | 0,14279774 | RBBP7 | -0,1033106 |
| ENSG00000 | gene | 0,89917705 | 0,04978245 | 0,14282241 | SMURF1 | -0,1533229 |
| ENSG00000 | gene | 1,27360298 | 0,04981579 | 0,14282241 | MED22 | 0,34891562 |
| ENSG00000 | gene | 0,9040924 | 0,04981971 | 0,14282241 | TNFRSF1A | -0,1454579 |
| ENSG00000 | gene | 0,92300869 | 0,04982379 | 0,14282241 | MTX1P1 | -0,1155839 |
| ENSG00000 | gene | 0,95043187 | 0,04983517 | 0,14282241 | ARMCX1 | -0,0733449 |
| ENSG00000 | gene | 0,91836178 | 0,04985379 | 0,1428345 | EEF1A1 | -0,1228655 |
| ENSG00000 | gene | 1,18435048 | 0,04987237 | 0,14284644 | USP44 | 0,24409607 |
| ENSG00000 | gene | 0,93591849 | 0,04990456 | 0,14289737 | WDR13 | -0,0955452 |
| ENSG00000 | gene | 1,1739295 | 0,04995483 | 0,1429486 | BRF2 | 0,23134577 |
| ENSG00000 | gene | 1,19614814 | 0,04995781 | 0,1429486 | USP53 | 0,25839608 |
| ENSG00000 | gene | 1,19867897 | 0,04996745 | 0,1429486 | FARP1-AS1 | 0,26144533 |
| ENSG00000 | gene | 1,3204599 | 0,04998011 | 0,1429486 | STAMBPL1 | 0,40104049 |
| ENSG00000 | gene | 0,83242434 | 0,04999666 | 0,1429547 | SUPT4H1 | -0,2646089 |

**MERS-CoV_DEGs**

| id | feature | fc | pval | qval | gene_name | de |
|---|---|---|---|---|---|---|
| ENSG00000 | gene | 1,15218809 | 1,5965E-13 | 1,7761E-09 | LRP1 | 0,20437625 |
| ENSG00000 | gene | 1,34404063 | 9,9032E-13 | 5,5086E-09 | TRPA1 | 0,42657675 |
| ENSG00000 | gene | 0,72672779 | 1,96E-12 | 7,2683E-09 | FP236383.5 | -0,460513 |
| ENSG00000 | gene | 0,87855048 | 4,7025E-12 | 1,3079E-08 | RPS2P5 | -0,1868029 |
| ENSG00000 | gene | 1,17512669 | 1,135E-10 | 2,4881E-07 | LTBP1 | 0,2328163 |
| ENSG00000 | gene | 1,43314082 | 1,3419E-10 | 2,4881E-07 | GPCPD1 | 0,51918037 |
| ENSG00000 | gene | 1,23446859 | 2,5513E-10 | 4,0547E-07 | SEMA3A | 0,30389013 |
| ENSG00000 | gene | 1,22379638 | 3,391E-09 | 4,7156E-06 | MSMO1 | 0,29136353 |
| ENSG00000 | gene | 0,8095107 | 1,9096E-08 | 2,3604E-05 | NDUFB11 | -0,3048779 |
| ENSG00000 | gene | 0,85682538 | 2,5378E-08 | 2,8233E-05 | CHCHD2 | -0,2229269 |
| ENSG00000 | gene | 1,21663065 | 4,5597E-08 | 4,6115E-05 | TPR | 0,28289126 |
| ENSG00000 | gene | 0,88083944 | 6,6518E-08 | 5,8905E-05 | TUBB4B | -0,183049 |
| ENSG00000 | gene | 1,08791871 | 7,5156E-08 | 5,8905E-05 | HTRA1 | 0,12157076 |
| ENSG00000 | gene | 1,31019911 | 7,9063E-08 | 5,8905E-05 | CDH6 | 0,38978608 |
| ENSG00000 | gene | 1,13745923 | 7,9422E-08 | 5,8905E-05 | FN1 | 0,18581483 |
| ENSG00000 | gene | 1,14623319 | 1,3781E-07 | 9,4022E-05 | COL4A1 | 0,19690057 |
| ENSG00000 | gene | 1,24673853 | 1,4367E-07 | 9,4022E-05 | EPAS1 | 0,31815893 |
| ENSG00000 | gene | 1,20524299 | 2,0936E-07 | 0,00012939 | AP002490.1 | 0,26932403 |
| ENSG00000 | gene | 0,84363766 | 2,4753E-07 | 0,00014494 | EPHA2 | -0,2453046 |
| ENSG00000 | gene | 1,38096233 | 4,2081E-07 | 0,00023408 | CABLES1 | 0,46567396 |
| ENSG00000 | gene | 1,0897653 | 8,3382E-07 | 0,00044173 | STC2 | 0,12401746 |
| ENSG00000 | gene | 1,12856649 | 1,0321E-06 | 0,00052192 | ERGIC1 | 0,17449142 |
| ENSG00000 | gene | 0,77504253 | 1,3969E-06 | 0,00067569 | DUSP14 | -0,3676526 |
| ENSG00000 | gene | 0,70130419 | 1,5875E-06 | 0,00073588 | COX5B | -0,5118877 |
| ENSG00000 | gene | 1,18762336 | 1,7537E-06 | 0,00078039 | COL4A2 | 0,24807738 |
| ENSG00000 | gene | 4,69383094 | 2,0795E-06 | 0,00088885 | ZBTB38 | 2,23076588 |
| ENSG00000 | gene | 1,18073856 | 2,1572E-06 | 0,00088885 | COL6A3 | 0,23968956 |
| ENSG00000 | gene | 1,16950578 | 2,6992E-06 | 0,00107244 | PLK2 | 0,22589899 |
| ENSG00000 | gene | 1,12541369 | 3,9762E-06 | 0,0014848 | DYNC1H1 | 0,17045541 |
| ENSG00000 | gene | 0,90817912 | 4,004E-06 | 0,0014848 | MT-TS1 | -0,1389512 |
| ENSG00000 | gene | 0,90350763 | 5,0604E-06 | 0,00181601 | MT-ND6 | -0,1463913 |
| ENSG00000 | gene | 1,08924857 | 5,3363E-06 | 0,00185521 | POLR2A | 0,12333322 |
| ENSG00000 | gene | 1,48008707 | 6,5087E-06 | 0,00208025 | AC011731.1 | 0,56568205 |
| ENSG00000 | gene | 1,55878077 | 6,54E-06 | 0,00208025 | CSGALNAC | 0,64041804 |
| ENSG00000 | gene | 1,5475778 | 6,5446E-06 | 0,00208025 | DKK1 | 0,63001194 |
| ENSG00000 | gene | 1,18541469 | 6,8176E-06 | 0,00210684 | AGPS | 0,24539184 |
| ENSG00000 | gene | 0,84506571 | 1,0799E-05 | 0,00318382 | PTX3 | -0,2428646 |
| ENSG00000 | gene | 1,17274008 | 1,0875E-05 | 0,00318382 | GOPC | 0,2298833 |
| ENSG00000 | gene | 0,74961418 | 1,1647E-05 | 0,00332247 | AL353644.2 | -0,4157799 |
| ENSG00000 | gene | 1,22650437 | 1,238E-05 | 0,00344319 | GJA1 | 0,29455237 |
| ENSG00000 | gene | 0,86522514 | 1,4447E-05 | 0,00382964 | TIMM22 | -0,2088525 |
| ENSG00000 | gene | 0,74873442 | 1,4458E-05 | 0,00382964 | RNA5-8SN4 | -0,417474 |
| ENSG00000 | gene | 0,8733911 | 1,5159E-05 | 0,00392202 | RPL39P3 | -0,1953003 |
| ENSG00000 | gene | 0,7370877 | 1,5726E-05 | 0,00397618 | RNA5-8SN3 | -0,4400918 |
| ENSG00000 | gene | 2,66849774 | 1,7801E-05 | 0,00440076 | PDE4B | 1,41602779 |
| ENSG00000 | gene | 0,91889275 | 2,1278E-05 | 0,00514608 | TFE3 | -0,1220316 |
| ENSG00000 | gene | 1,14311782 | 2,3293E-05 | 0,00551354 | FAT1 | 0,19297411 |
| ENSG00000 | gene | 0,80655082 | 2,6702E-05 | 0,00612502 | ZNF703 | -0,3101626 |
| ENSG00000 | gene | 0,75240162 | 2,6978E-05 | 0,00612502 | RNA5-8SN2 | -0,4104251 |
| ENSG00000 | gene | 0,84550941 | 2,7607E-05 | 0,00614249 | AC027319.1 | -0,2421073 |
| ENSG00000 | gene | 0,87833037 | 2,9044E-05 | 0,00633554 | PRELID3B | -0,1871644 |
| ENSG00000 | gene | 0,65972515 | 3,2734E-05 | 0,00700317 | FAM216A | -0,600063 |
| ENSG00000 | gene | 0,85934003 | 3,8128E-05 | 0,00782604 | RBMS1 | -0,218699 |
| ENSG00000 | gene | 0,84705833 | 3,8611E-05 | 0,00782604 | AC011603.3 | -0,2394668 |
| ENSG00000 | gene | 0,77816811 | 3,8691E-05 | 0,00782604 | FP236383.4 | -0,3618462 |

| | | | | | |
|---|---|---|---|---|---|
| ENSG00000 gene | 1,21493866 | 4,495E-05 | 0,00892977 | BICC1 | 0,28088348 |
| ENSG00000 gene | 0,92590762 | 4,7288E-05 | 0,00894468 | UBE2H | -0,1110598 |
| ENSG00000 gene | 1,18160287 | 4,7465E-05 | 0,00894468 | SALL1 | 0,24074524 |
| ENSG00000 gene | 1,53235873 | 4,7686E-05 | 0,00894468 | CYP2S1 | 0,61575407 |
| ENSG00000 gene | 1,41584988 | 4,8315E-05 | 0,00894468 | AC113386.1 | 0,5016683 |
| ENSG00000 gene | 1,42192631 | 4,9045E-05 | 0,00894468 | DNER | 0,5078467 |
| ENSG00000 gene | 1,06741086 | 5,0731E-05 | 0,0091029 | GNS | 0,0941156 |
| ENSG00000 gene | 0,93353877 | 5,5772E-05 | 0,00984858 | RPS28 | -0,0992182 |
| ENSG00000 gene | 0,93005563 | 5,883E-05 | 0,01022635 | MT-ND3 | -0,1046111 |
| ENSG00000 gene | 1,2163005 | 7,0124E-05 | 0,012002 | USP11 | 0,28249971 |
| ENSG00000 gene | 1,26429606 | 8,1224E-05 | 0,01338438 | FAP | 0,33833434 |
| ENSG00000 gene | 0,87638572 | 8,1786E-05 | 0,01338438 | MIF | -0,1903621 |
| ENSG00000 gene | 1,13930672 | 8,181E-05 | 0,01338438 | RGP1 | 0,1881562 |
| ENSG00000 gene | 1,41401174 | 8,6149E-05 | 0,0138899 | ADAMTS5 | 0,4997941 |
| ENSG00000 gene | 0,95689904 | 9,0711E-05 | 0,01420959 | RPS9 | -0,0635614 |
| ENSG00000 gene | 0,8750387 | 9,1361E-05 | 0,01420959 | TUBA1B | -0,1925813 |
| ENSG00000 gene | 2,16221867 | 9,1963E-05 | 0,01420959 | IL33 | 1,11251243 |
| ENSG00000 gene | 1,26315584 | 9,5672E-05 | 0,01447876 | ANGPTL2 | 0,33703264 |
| ENSG00000 gene | 1,15886559 | 9,6308E-05 | 0,01447876 | DDX10 | 0,21271325 |
| ENSG00000 gene | 0,90829681 | 0,00010129 | 0,01502489 | RPS12 | -0,1387643 |
| ENSG00000 gene | 0,93173002 | 0,00010546 | 0,01536694 | SERBP1 | -0,1020161 |
| ENSG00000 gene | 0,91545849 | 0,00010636 | 0,01536694 | GINM1 | -0,1274336 |
| ENSG00000 gene | 1,09161149 | 0,00011519 | 0,01617867 | EXT1 | 0,12645948 |
| ENSG00000 gene | 0,68256723 | 0,00011632 | 0,01617867 | AC109322.1 | -0,550957 |
| ENSG00000 gene | 0,86370361 | 0,00011634 | 0,01617867 | SNX6 | -0,2113918 |
| ENSG00000 gene | 1,28255114 | 0,00011935 | 0,01639279 | SLC6A9 | 0,35901636 |
| ENSG00000 gene | 1,15802345 | 0,0001245 | 0,01681041 | SC5D | 0,21166447 |
| ENSG00000 gene | 0,83729426 | 0,00012542 | 0,01681041 | LSM7 | -0,2561934 |
| ENSG00000 gene | 1,24603548 | 0,00012748 | 0,01688391 | RNF213 | 0,31734514 |
| ENSG00000 gene | 2,35411757 | 0,00013024 | 0,01704655 | Z84488.2 | 1,23518637 |
| ENSG00000 gene | 0,74978326 | 0,0001335 | 0,01718736 | SELPLG | -0,4154545 |
| ENSG00000 gene | 0,90625283 | 0,00013441 | 0,01718736 | SNHG5 | -0,1420145 |
| ENSG00000 gene | 1,1451241 | 0,0001362 | 0,01721818 | PAM | 0,19550395 |
| ENSG00000 gene | 0,87793947 | 0,00014351 | 0,01793879 | TSPAN5 | -0,1878066 |
| ENSG00000 gene | 1,30778864 | 0,00014888 | 0,01840304 | FADS1 | 0,3871294 |
| ENSG00000 gene | 1,5202681 | 0,00015129 | 0,01849554 | SYNGAP1 | 0,60432576 |
| ENSG00000 gene | 0,93475581 | 0,0001606 | 0,01941999 | EEF1D | -0,0973386 |
| ENSG00000 gene | 1,10912518 | 0,00018027 | 0,02156506 | AC004477.2 | 0,1494222 |
| ENSG00000 gene | 0,89192185 | 0,00018306 | 0,0216659 | PLCB3 | -0,1650108 |
| ENSG00000 gene | 0,88518732 | 0,00018947 | 0,022188 | ITGB1BP1 | -0,1759453 |
| ENSG00000 gene | 1,10391784 | 0,00019687 | 0,02281441 | DST | 0,1426328 |
| ENSG00000 gene | 0,87707638 | 0,00019895 | 0,02281782 | AC015674.1 | -0,1892256 |
| ENSG00000 gene | 0,81808864 | 0,00021046 | 0,02389188 | AC079250.1 | -0,2896709 |
| ENSG00000 gene | 0,7599214 | 0,00022891 | 0,02572331 | STING1 | -0,3960779 |
| ENSG00000 gene | 0,87560781 | 0,00023883 | 0,0265696 | COX8A | -0,1916433 |
| ENSG00000 gene | 1,63035331 | 0,00025721 | 0,02833128 | KLC4 | 0,70518464 |
| ENSG00000 gene | 1,17764141 | 0,00026075 | 0,02844003 | RHOB | 0,23590031 |
| ENSG00000 gene | 0,89841172 | 0,00026684 | 0,02882094 | RPS21 | -0,1545513 |
| ENSG00000 gene | 0,87884352 | 0,00029008 | 0,03102992 | PTN | -0,1863218 |
| ENSG00000 gene | 0,76990465 | 0,00030399 | 0,03220854 | SH3GLB2 | -0,3772483 |
| ENSG00000 gene | 0,91106357 | 0,00033657 | 0,03532387 | IMMT | -0,1343764 |
| ENSG00000 gene | 1,27515508 | 0,00037324 | 0,03880646 | PTGS1 | 0,35067272 |
| ENSG00000 gene | 0,76868568 | 0,00038085 | 0,03923105 | TIMP3 | -0,3795343 |
| ENSG00000 gene | 1,45178392 | 0,0003943 | 0,04024401 | FGF7 | 0,53782674 |
| ENSG00000 gene | 1,42107717 | 0,0004024 | 0,04069755 | CLMP | 0,5069849 |
| ENSG00000 gene | 0,88182967 | 0,0004739 | 0,04749628 | H2AX | -0,1814281 |
| ENSG00000 gene | 0,85399163 | 0,0004954 | 0,04862994 | AC125611.3 | -0,2277062 |

| | | | | | |
|---|---|---|---|---|---|
| ENSG00000 gene | 1,2920331 | 0,00049558 | 0,04862994 | ATG13 | 0,36964303 |
| ENSG00000 gene | 0,84544847 | 0,00049832 | 0,04862994 | PDE6D | -0,2422113 |
| ENSG00000 gene | 1,82966426 | 0,00051276 | 0,04960353 | SLC16A6 | 0,87157894 |
| ENSG00000 gene | 1,08773332 | 0,00052272 | 0,05013136 | GTPBP2 | 0,12132489 |
| ENSG00000 gene | 0,84107918 | 0,00053267 | 0,05064954 | UAP1 | -0,2496865 |
| ENSG00000 gene | 0,86354012 | 0,00056843 | 0,05359144 | POMP | -0,2116649 |
| ENSG00000 gene | 1,41000274 | 0,00058757 | 0,05493027 | RSAD2 | 0,49569796 |
| ENSG00000 gene | 0,55993322 | 0,00060914 | 0,05647229 | MITF | -0,8366733 |
| ENSG00000 gene | 0,91342067 | 0,00063152 | 0,05769704 | RFLNB | -0,1306487 |
| ENSG00000 gene | 0,87152772 | 0,00063272 | 0,05769704 | NIFK | -0,1983815 |
| ENSG00000 gene | 0,85976034 | 0,00066379 | 0,06003833 | SIPA1 | -0,2179935 |
| ENSG00000 gene | 0,89472587 | 0,00069427 | 0,06166327 | SRP19 | -0,1604824 |
| ENSG00000 gene | 0,9118777 | 0,00069696 | 0,06166327 | APMAP | -0,1330877 |
| ENSG00000 gene | 0,8450581 | 0,000699 | 0,06166327 | TRNP1 | -0,2428776 |
| ENSG00000 gene | 0,93617944 | 0,00070393 | 0,06166327 | TMEM50A | -0,095143 |
| ENSG00000 gene | 0,83943535 | 0,00071629 | 0,06223646 | NDUFAB1 | -0,2525089 |
| ENSG00000 gene | 0,83103017 | 0,00072851 | 0,06223646 | TEAD4 | -0,2670272 |
| ENSG00000 gene | 0,68824719 | 0,00073214 | 0,06223646 | NUP35 | -0,5390013 |
| ENSG00000 gene | 0,94755897 | 0,00073285 | 0,06223646 | PSMB2 | -0,0777124 |
| ENSG00000 gene | 1,08303222 | 0,00076292 | 0,06429886 | ARHGEF11 | 0,11507616 |
| ENSG00000 gene | 0,74662186 | 0,00077983 | 0,06501955 | RPL22L1 | -0,4215503 |
| ENSG00000 gene | 0,81623351 | 0,00078316 | 0,06501955 | POLR2E | -0,2929462 |
| ENSG00000 gene | 0,84960358 | 0,00079587 | 0,06558532 | EVA1B | -0,2351383 |
| ENSG00000 gene | 0,92919842 | 0,00081507 | 0,06667389 | GNB2 | -0,1059414 |
| ENSG00000 gene | 1,19095543 | 0,0008861 | 0,07195529 | MOSPD1 | 0,25211942 |
| ENSG00000 gene | 0,8971792 | 0,00090055 | 0,07259866 | UBE2L3 | -0,1565319 |
| ENSG00000 gene | 0,8855714 | 0,00092384 | 0,07394033 | C8orf33 | -0,1753195 |
| ENSG00000 gene | 1,08294532 | 0,00094017 | 0,07470993 | MAVS | 0,1149604 |
| ENSG00000 gene | 1,143033 | 0,00097681 | 0,07661406 | MCC | 0,19286706 |
| ENSG00000 gene | 0,90337706 | 0,00097791 | 0,07661406 | CSTB | -0,1465998 |
| ENSG00000 gene | 0,70563362 | 0,00100565 | 0,07823652 | MYL12BP1 | -0,5030088 |
| ENSG00000 gene | 0,44492056 | 0,00103161 | 0,07894908 | VDR | -1,1683803 |
| ENSG00000 gene | 1,24508536 | 0,00103798 | 0,07894908 | CDCP1 | 0,31624465 |
| ENSG00000 gene | 0,89117943 | 0,00104176 | 0,07894908 | FOXL1 | -0,1662122 |
| ENSG00000 gene | 0,91632057 | 0,00104319 | 0,07894908 | AL583722.2 | -0,1260757 |
| ENSG00000 gene | 1,20524335 | 0,00106464 | 0,0800279 | IGF2R | 0,26932447 |
| ENSG00000 gene | 0,92797461 | 0,00109327 | 0,08162839 | EIF3E | -0,1078428 |
| ENSG00000 gene | 0,86044008 | 0,00111669 | 0,08282154 | H2AZ1 | -0,2168534 |
| ENSG00000 gene | 0,7707074 | 0,00112574 | 0,08293959 | SCAND1 | -0,3757448 |
| ENSG00000 gene | 0,78984498 | 0,00121001 | 0,08856153 | RN7SL1 | -0,3403586 |
| ENSG00000 gene | 1,73111732 | 0,00122755 | 0,08925818 | AC112496.1 | 0,7917035 |
| ENSG00000 gene | 0,93215963 | 0,00125005 | 0,09030427 | TUFM | -0,1013511 |
| ENSG00000 gene | 1,11922567 | 0,00126808 | 0,09101523 | DDX24 | 0,16250096 |
| ENSG00000 gene | 0,87132744 | 0,0013063 | 0,0931576 | CLIC1 | -0,1987131 |
| ENSG00000 gene | 1,30421078 | 0,00135542 | 0,09561637 | THBS2 | 0,38317705 |
| ENSG00000 gene | 0,93263708 | 0,00135797 | 0,09561637 | SNRPB2 | -0,1006123 |
| ENSG00000 gene | 0,86358989 | 0,00136924 | 0,09580341 | TBC1D2B | -0,2115817 |
| ENSG00000 gene | 1,2355209 | 0,00141548 | 0,09842038 | MSMP | 0,30511941 |
| ENSG00000 gene | 1,0454327 | 0,00145841 | 0,10077552 | PSAP | 0,0641002 |
| ENSG00000 gene | 0,85783475 | 0,00148721 | 0,10213104 | REEP3 | -0,2212283 |
| ENSG00000 gene | 0,86321171 | 0,00149695 | 0,10216935 | ARHGEF10L | -0,2122137 |
| ENSG00000 gene | 1,8997164 | 0,00154402 | 0,10457083 | YPEL2 | 0,92578406 |
| ENSG00000 gene | 1,08878054 | 0,00155094 | 0,10457083 | TOPBP1 | 0,12271319 |
| ENSG00000 gene | 0,89251039 | 0,00158776 | 0,10640861 | FCHSD2 | -0,1640591 |
| ENSG00000 gene | 0,79388902 | 0,00163578 | 0,10849391 | FJX1 | -0,3329908 |
| ENSG00000 gene | 0,85229339 | 0,00164045 | 0,10849391 | CORO1B | -0,230578 |
| ENSG00000 gene | 0,89413497 | 0,00165956 | 0,10849391 | AC087393.2 | -0,1614355 |

| | | | | | | |
|---|---|---|---|---|---|---|
| ENSG00000 gene | | 0,92006078 | 0,00166972 | 0,10849391 | HNRNPL | -0,1201989 |
| ENSG00000 gene | | 0,94113193 | 0,00167609 | 0,10849391 | PAPSS2 | -0,0875311 |
| ENSG00000 gene | | 0,89363153 | 0,00167739 | 0,10849391 | RALY | -0,162248 |
| ENSG00000 gene | | 0,86014715 | 0,00170619 | 0,10971895 | MRPS22 | -0,2173446 |
| ENSG00000 gene | | 0,91197859 | 0,00173597 | 0,11011201 | PPP1R12A | -0,1329281 |
| ENSG00000 gene | | 2,14831177 | 0,00174162 | 0,11011201 | RAPGEF6 | 1,10320337 |
| ENSG00000 gene | | 1,88632441 | 0,001742 | 0,11011201 | LINC01705 | 0,91557781 |
| ENSG00000 gene | | 0,97694677 | 0,00176167 | 0,11072618 | TUBB | -0,0336481 |
| ENSG00000 gene | | 1,09564271 | 0,00178361 | 0,11147549 | FDPS | 0,13177741 |
| ENSG00000 gene | | 0,79670335 | 0,00181402 | 0,11230503 | CKB | -0,3278855 |
| ENSG00000 gene | | 0,85593208 | 0,00181707 | 0,11230503 | TOMM22 | -0,2244318 |
| ENSG00000 gene | | 1,11058108 | 0,00189866 | 0,11669911 | MT-ND4L | 0,15131473 |
| ENSG00000 gene | | 0,92049787 | 0,00195467 | 0,1194816 | CYFIP1 | -0,1195137 |
| ENSG00000 gene | | 1,08008659 | 0,00199281 | 0,12083479 | DDX6 | 0,11114697 |
| ENSG00000 gene | | 0,88329639 | 0,00199853 | 0,12083479 | RPS28P7 | -0,1790305 |
| ENSG00000 gene | | 0,89498684 | 0,00203375 | 0,12229982 | BOP1 | -0,1600616 |
| ENSG00000 gene | | 0,81477976 | 0,00204583 | 0,12236463 | MYL12B | -0,295518 |
| ENSG00000 gene | | 1,87151176 | 0,00207462 | 0,12342297 | MME | 0,90420412 |
| ENSG00000 gene | | 1,23017064 | 0,00216189 | 0,1276547 | BX322234.1 | 0,29885845 |
| ENSG00000 gene | | 1,30160327 | 0,0021687 | 0,1276547 | HAS2 | 0,38028978 |
| ENSG00000 gene | | 0,90293365 | 0,00219825 | 0,12871328 | PPP4R1 | -0,1473081 |
| ENSG00000 gene | | 1,18556014 | 0,00222373 | 0,12932563 | ARRDC3 | 0,24556884 |
| ENSG00000 gene | | 1,0701788 | 0,00223196 | 0,12932563 | SMC1A | 0,09785185 |
| ENSG00000 gene | | 1,08092874 | 0,00224362 | 0,12932782 | ZKSCAN1 | 0,11227142 |
| ENSG00000 gene | | 0,77153063 | 0,00226034 | 0,12961992 | ZNRD1 | -0,3742047 |
| ENSG00000 gene | | 0,77246292 | 0,00234478 | 0,13377294 | IGFBP6 | -0,3724624 |
| ENSG00000 gene | | 0,94330246 | 0,00236618 | 0,13389917 | AP2A1 | -0,0842077 |
| ENSG00000 gene | | 0,74117915 | 0,00238155 | 0,13389917 | DAD1 | -0,4321058 |
| ENSG00000 gene | | 1,13239728 | 0,0023831 | 0,13389917 | SUPT6H | 0,17938019 |
| ENSG00000 gene | | 0,79922003 | 0,00244313 | 0,13456062 | PPDPF | -0,3233354 |
| ENSG00000 gene | | 0,81899489 | 0,00244433 | 0,13456062 | C9orf16 | -0,2880736 |
| ENSG00000 gene | | 1,51088289 | 0,00245152 | 0,13456062 | IL1R1 | 0,59539184 |
| ENSG00000 gene | | 0,58983847 | 0,00245741 | 0,13456062 | AC060766.5 | -0,7616082 |
| ENSG00000 gene | | 1,10595387 | 0,00248073 | 0,13456062 | PXDN | 0,14529121 |
| ENSG00000 gene | | 0,9287895 | 0,00248787 | 0,13456062 | RPLP0P6 | -0,1065764 |
| ENSG00000 gene | | 1,13616516 | 0,00248934 | 0,13456062 | RAB23 | 0,18417257 |
| ENSG00000 gene | | 0,77253334 | 0,00249164 | 0,13456062 | TIMM13 | -0,3723309 |
| ENSG00000 gene | | 0,88014723 | 0,00254351 | 0,13669828 | RPL35A | -0,1841832 |
| ENSG00000 gene | | 0,90189427 | 0,00258811 | 0,13842641 | COX6A1 | -0,1489698 |
| ENSG00000 gene | | 1,91736376 | 0,00262793 | 0,13988387 | MIR4746 | 0,93912407 |
| ENSG00000 gene | | 1,07391509 | 0,00274591 | 0,14546806 | PBXIP1 | 0,10287992 |
| ENSG00000 gene | | 1,12358779 | 0,00276024 | 0,14553402 | TSPYL2 | 0,16811285 |
| ENSG00000 gene | | 0,90274211 | 0,00281462 | 0,14770107 | SDHB | -0,1476142 |
| ENSG00000 gene | | 0,79555188 | 0,00284939 | 0,14882368 | TK1 | -0,3299721 |
| ENSG00000 gene | | 0,73355647 | 0,00287039 | 0,14921998 | KISS1 | -0,4470201 |
| ENSG00000 gene | | 1,35401204 | 0,00291175 | 0,15058541 | PRKG1-AS1 | 0,43724057 |
| ENSG00000 gene | | 0,77354578 | 0,00292373 | 0,15058541 | CYLD | -0,3704414 |
| ENSG00000 gene | | 1,11749354 | 0,00293762 | 0,15060394 | ANPEP | 0,16026649 |
| ENSG00000 gene | | 1,4106321 | 0,00298624 | 0,15203188 | MME-AS1 | 0,49634177 |
| ENSG00000 gene | | 0,75651149 | 0,00299281 | 0,15203188 | OST4 | -0,4025661 |
| ENSG00000 gene | | 0,5357723 | 0,00309483 | 0,15649983 | SLC25A14 | -0,9003081 |
| ENSG00000 gene | | 1,89391713 | 0,00314587 | 0,15836103 | ZNF584 | 0,92137321 |
| ENSG00000 gene | | 1,13581371 | 0,00316036 | 0,15837398 | TOP2A | 0,18372623 |
| ENSG00000 gene | | 0,8609407 | 0,00322422 | 0,1608154 | S100A11 | -0,2160142 |
| ENSG00000 gene | | 0,84389151 | 0,00323799 | 0,1608154 | PPT1 | -0,2448706 |
| ENSG00000 gene | | 1,21336578 | 0,00325388 | 0,16088652 | GPNMB | 0,27901453 |
| ENSG00000 gene | | 0,83986473 | 0,0032939 | 0,16214433 | RABAC1 | -0,2517711 |

| | | | | | |
|---|---|---|---|---|---|
| ENSG00000 gene | 0,64927829 | 0,00337597 | 0,16545241 | RPPH1 | -0,6230911 |
| ENSG00000 gene | 1,21102487 | 0,00341289 | 0,16630127 | ANTXR1 | 0,27622849 |
| ENSG00000 gene | 0,77885927 | 0,00342319 | 0,16630127 | MBIP | -0,3605654 |
| ENSG00000 gene | 0,91326898 | 0,00345605 | 0,16659867 | PDHA1 | -0,1308883 |
| ENSG00000 gene | 0,89042704 | 0,00345926 | 0,16659867 | C5orf30 | -0,1674307 |
| ENSG00000 gene | 0,67701363 | 0,00368202 | 0,17656239 | VKORC1 | -0,5627432 |
| ENSG00000 gene | 0,66244115 | 0,00370256 | 0,17678525 | RMRP | -0,5941358 |
| ENSG00000 gene | 0,87963861 | 0,00373687 | 0,17741828 | SMAGP | -0,1850172 |
| ENSG00000 gene | 0,67159372 | 0,00374771 | 0,17741828 | AC073641.1 | -0,5743394 |
| ENSG00000 gene | 1,38306632 | 0,00377451 | 0,17753507 | AC009511.1 | 0,46787033 |
| ENSG00000 gene | 1,03937207 | 0,00378715 | 0,17753507 | SEC62 | 0,05571219 |
| ENSG00000 gene | 0,78984605 | 0,00381154 | 0,17753507 | SMCO4 | -0,3403566 |
| ENSG00000 gene | 0,88752061 | 0,00382344 | 0,17753507 | AC243919.1 | -0,1721475 |
| ENSG00000 gene | 0,83383589 | 0,00382997 | 0,17753507 | RN7SL4P | -0,2621646 |
| ENSG00000 gene | 1,20615816 | 0,00386352 | 0,17834694 | IGF1R | 0,27041909 |
| ENSG00000 gene | 2,58927263 | 0,00395413 | 0,1817757 | NRG1 | 1,37254688 |
| ENSG00000 gene | 1,69386312 | 0,00403523 | 0,18474037 | PTGES | 0,7603173 |
| ENSG00000 gene | 1,92600752 | 0,00406851 | 0,18550087 | FOXO3B | 0,94561333 |
| ENSG00000 gene | 0,93812103 | 0,00411496 | 0,18685288 | CAPN2 | -0,092154 |
| ENSG00000 gene | 1,98020193 | 0,00414557 | 0,18747741 | RNU6-26P | 0,98564755 |
| ENSG00000 gene | 1,10850486 | 0,0041725 | 0,18793136 | ARFGEF2 | 0,14861509 |
| ENSG00000 gene | 0,91581158 | 0,00419029 | 0,18797175 | SLC25A3 | -0,1268773 |
| ENSG00000 gene | 0,65351223 | 0,00423781 | 0,18825223 | GADD45B | -0,6137139 |
| ENSG00000 gene | 0,93720028 | 0,00423859 | 0,18825223 | OPTN | -0,0935707 |
| ENSG00000 gene | 0,90111508 | 0,00425199 | 0,18825223 | RPS13 | -0,1502167 |
| ENSG00000 gene | 1,2262306 | 0,00426423 | 0,18825223 | SLC38A7 | 0,29423031 |
| ENSG00000 gene | 0,82994517 | 0,0043662 | 0,19199208 | MRPL41 | -0,2689121 |
| ENSG00000 gene | 0,95055421 | 0,00442709 | 0,19351923 | SEC24D | -0,0731592 |
| ENSG00000 gene | 0,48806897 | 0,00443572 | 0,19351923 | AC009065.1( | -1,0348431 |
| ENSG00000 gene | 0,6474718 | 0,0044988 | 0,19550444 | MIR3615 | -0,6271107 |
| ENSG00000 gene | 0,84748366 | 0,00463964 | 0,2008404 | SDF4 | -0,2387425 |
| ENSG00000 gene | 0,82935991 | 0,00468125 | 0,20185607 | STK11 | -0,2699298 |
| ENSG00000 gene | 1,0768307 | 0,00471618 | 0,2020054 | CASC3 | 0,10679144 |
| ENSG00000 gene | 0,85588406 | 0,00472103 | 0,2020054 | HAT1 | -0,2245127 |
| ENSG00000 gene | 0,88669024 | 0,00479054 | 0,20419444 | UBL5 | -0,1734979 |
| ENSG00000 gene | 1,85546858 | 0,0048356 | 0,20450969 | SNORD124 | 0,89178357 |
| ENSG00000 gene | 0,84759811 | 0,00484423 | 0,20450969 | GPR137 | -0,2385477 |
| ENSG00000 gene | 1,57409006 | 0,00485308 | 0,20450969 | EPB41L5 | 0,65451808 |
| ENSG00000 gene | 1,25167617 | 0,00492739 | 0,20685751 | SGIP1 | 0,32386136 |
| ENSG00000 gene | 0,89639076 | 0,0049516 | 0,20709246 | SEH1L | -0,1578003 |
| ENSG00000 gene | 1,11133489 | 0,00502636 | 0,20870941 | SRXN1 | 0,15229362 |
| ENSG00000 gene | 1,15773782 | 0,00502779 | 0,20870941 | WLS | 0,21130858 |
| ENSG00000 gene | 0,8799212 | 0,00510654 | 0,21119065 | RBCK1 | -0,1845538 |
| ENSG00000 gene | 0,89332281 | 0,00515344 | 0,21234084 | RANBP3 | -0,1627465 |
| ENSG00000 gene | 0,8792231 | 0,00521965 | 0,2136109 | GPAA1 | -0,1856988 |
| ENSG00000 gene | 1,48114544 | 0,00522267 | 0,2136109 | SHC2 | 0,56671332 |
| ENSG00000 gene | 0,79016798 | 0,00532675 | 0,21707002 | MT-TR | -0,3397687 |
| ENSG00000 gene | 0,91751576 | 0,00538624 | 0,21869308 | AC012640.6 | -0,1241952 |
| ENSG00000 gene | 0,81116354 | 0,00547333 | 0,22078767 | SFT2D1 | -0,3019353 |
| ENSG00000 gene | 0,91848472 | 0,00547752 | 0,22078767 | PPM1F | -0,1226724 |
| ENSG00000 gene | 0,84530566 | 0,00550036 | 0,22090801 | MRPS15 | -0,242455 |
| ENSG00000 gene | 2,12450656 | 0,00554236 | 0,22179398 | SPIN1 | 1,0871278 |
| ENSG00000 gene | 1,6934525 | 0,00559909 | 0,22326125 | AP000346.2 | 0,75996752 |
| ENSG00000 gene | 1,41584294 | 0,00567199 | 0,22536021 | PAPPA-AS1 | 0,50166123 |
| ENSG00000 gene | 1,32079528 | 0,00573145 | 0,22691242 | FADS2 | 0,40140686 |
| ENSG00000 gene | 0,86496911 | 0,00577817 | 0,22795076 | CDC42 | -0,2092795 |
| ENSG00000 gene | 0,88727753 | 0,00581778 | 0,22799583 | RPL22 | -0,1725427 |

| | | | | | |
|---|---|---|---|---|---|
| ENSG00000 gene | 1,72817217 | 0,0058203 | 0,22799583 | PLEKHA5 | 0,78924696 |
| ENSG00000 gene | 1,44768928 | 0,00600586 | 0,23443923 | PAPPA | 0,53375199 |
| ENSG00000 gene | 0,92019426 | 0,00603446 | 0,23473208 | SRRD | -0,1199896 |
| ENSG00000 gene | 0,86866088 | 0,00606766 | 0,2352011 | RN7SL2 | -0,203135 |
| ENSG00000 gene | 0,8220214 | 0,00613579 | 0,23550016 | TSPO | -0,2827522 |
| ENSG00000 gene | 0,96439984 | 0,00613807 | 0,23550016 | BRK1 | -0,0522967 |
| ENSG00000 gene | 1,2717708 | 0,00613888 | 0,23550016 | PRRC2B | 0,34683869 |
| ENSG00000 gene | 0,89315378 | 0,00617963 | 0,23624855 | CD320 | -0,1630195 |
| ENSG00000 gene | 0,78111516 | 0,00628741 | 0,23954589 | CTDNEP1 | -0,3563928 |
| ENSG00000 gene | 1,17405071 | 0,00638055 | 0,24151959 | PRPSAP1 | 0,23149472 |
| ENSG00000 gene | 2,09701074 | 0,00638263 | 0,24151959 | AC005740.3 | 1,06833425 |
| ENSG00000 gene | 0,90902043 | 0,00644348 | 0,2423193 | RASSF3 | -0,1376154 |
| ENSG00000 gene | 1,61234331 | 0,00646138 | 0,2423193 | MICAL3 | 0,68915896 |
| ENSG00000 gene | 0,76538829 | 0,00649739 | 0,2423193 | NDUFS6 | -0,3857363 |
| ENSG00000 gene | 0,79363729 | 0,00651501 | 0,2423193 | CDC42EP1 | -0,3334483 |
| ENSG00000 gene | 1,09037926 | 0,00653502 | 0,2423193 | SF3B1 | 0,12483002 |
| ENSG00000 gene | 0,93711098 | 0,00656925 | 0,2423193 | YWHAQ | -0,0937082 |
| ENSG00000 gene | 0,70292594 | 0,00657264 | 0,2423193 | UBE2V1 | -0,5085554 |
| ENSG00000 gene | 0,91095889 | 0,00657802 | 0,2423193 | P3H3 | -0,1345421 |
| ENSG00000 gene | 0,83383828 | 0,00665174 | 0,24275865 | XPNPEP1 | -0,2621605 |
| ENSG00000 gene | 0,94996939 | 0,00666438 | 0,24275865 | RAC1 | -0,0740471 |
| ENSG00000 gene | 1,11708439 | 0,00667014 | 0,24275865 | ZNF623 | 0,15973818 |
| ENSG00000 gene | 0,72281432 | 0,00667723 | 0,24275865 | USP10 | -0,468303 |
| ENSG00000 gene | 0,81251468 | 0,00672953 | 0,2438631 | RNF185-AS1 | -0,2995342 |
| ENSG00000 gene | 0,91235682 | 0,00676945 | 0,24451359 | VGLL4 | -0,1323299 |
| ENSG00000 gene | 1,30742846 | 0,00687225 | 0,24742315 | HIF1A | 0,386732 |
| ENSG00000 gene | 1,12268094 | 0,00691993 | 0,24833621 | ZNF217 | 0,16694798 |
| ENSG00000 gene | 0,88871957 | 0,00697526 | 0,24863651 | CEACAM19 | -0,1701998 |
| ENSG00000 gene | 1,9980268 | 0,00698952 | 0,24863651 | NPHP3-ACA | 0,99857593 |
| ENSG00000 gene | 1,16749551 | 0,00699535 | 0,24863651 | STARD13 | 0,223417 |
| ENSG00000 gene | 1,05789216 | 0,00709445 | 0,2505973 | BGN | 0,08119256 |
| ENSG00000 gene | 1,77085371 | 0,00709556 | 0,2505973 | CREB5 | 0,82444504 |
| ENSG00000 gene | 0,91974832 | 0,00716483 | 0,2519139 | HPS1 | -0,120689 |
| ENSG00000 gene | 1,33726522 | 0,00717813 | 0,2519139 | CHD2 | 0,41928563 |
| ENSG00000 gene | 1,14042002 | 0,00722963 | 0,25292327 | POSTN | 0,18956527 |
| ENSG00000 gene | 1,07220796 | 0,00730117 | 0,25414029 | NORAD | 0,10058474 |
| ENSG00000 gene | 0,90506836 | 0,00732499 | 0,25414029 | PHB | -0,1439013 |
| ENSG00000 gene | 0,87646169 | 0,00734033 | 0,25414029 | ARF3 | -0,1902371 |
| ENSG00000 gene | 0,75175832 | 0,00735579 | 0,25414029 | BRIX1 | -0,4116592 |
| ENSG00000 gene | 1,34875974 | 0,00744995 | 0,25596874 | FHOD3 | 0,43163338 |
| ENSG00000 gene | 0,91074036 | 0,00748866 | 0,25596874 | CAP1 | -0,1348883 |
| ENSG00000 gene | 1,26112077 | 0,00752403 | 0,25596874 | ROBO1 | 0,33470644 |
| ENSG00000 gene | 0,79366591 | 0,00752495 | 0,25596874 | PXK | -0,3333963 |
| ENSG00000 gene | 0,75815353 | 0,00753844 | 0,25596874 | CSF1 | -0,3994381 |
| ENSG00000 gene | 1,0778459 | 0,00758581 | 0,25596874 | AP2A2 | 0,10815093 |
| ENSG00000 gene | 0,82504471 | 0,00759844 | 0,25596874 | SUB1 | -0,2774558 |
| ENSG00000 gene | 1,34810149 | 0,00760339 | 0,25596874 | SLC6A15 | 0,43092911 |
| ENSG00000 gene | 0,80256229 | 0,00761579 | 0,25596874 | TMEM68 | -0,3173147 |
| ENSG00000 gene | 1,18308876 | 0,0076523 | 0,25614425 | GPC6 | 0,24255832 |
| ENSG00000 gene | 0,58086636 | 0,00766706 | 0,25614425 | SNORD54 | -0,7837218 |
| ENSG00000 gene | 1,58088424 | 0,00771276 | 0,25689955 | LINC00161 | 0,66073173 |
| ENSG00000 gene | 1,16254838 | 0,00778151 | 0,25841578 | VMP1 | 0,21729075 |
| ENSG00000 gene | 1,17196826 | 0,00794231 | 0,26236964 | PLAT | 0,2289335 |
| ENSG00000 gene | 1,25222747 | 0,00794774 | 0,26236964 | OLR1 | 0,32449665 |
| ENSG00000 gene | 0,39212241 | 0,00801995 | 0,26397031 | AC002985.1 | -1,350624 |
| ENSG00000 gene | 0,80402538 | 0,00810776 | 0,26607319 | PRPSAP2 | -0,314687 |
| ENSG00000 gene | 1,21719853 | 0,00815695 | 0,26690034 | EIF4BP6 | 0,2835645 |

| | | | | | |
|---|---|---|---|---|---|
| ENSG00000 gene | 0,85846423 | 0,00818188 | 0,26693083 | SURF2 | -0,2201701 |
| ENSG00000 gene | 0,67441174 | 0,00821165 | 0,26711871 | TMEM138 | -0,5682985 |
| ENSG00000 gene | 0,89973903 | 0,00823957 | 0,26724555 | YARS2 | -0,1524215 |
| ENSG00000 gene | 0,86087065 | 0,00831823 | 0,2689126 | METTL2A | -0,2161316 |
| ENSG00000 gene | 0,91570526 | 0,0083631 | 0,2689126 | EIF5B | -0,1270448 |
| ENSG00000 gene | 0,81991071 | 0,00836348 | 0,2689126 | BUD31 | -0,2864613 |
| ENSG00000 gene | 1,08646161 | 0,00848124 | 0,27191304 | DNAJC24 | 0,1196372 |
| ENSG00000 gene | 1,07175358 | 0,00854248 | 0,27308941 | WASHC2A | 0,09997324 |
| ENSG00000 gene | 1,91713202 | 0,00856858 | 0,2731388 | GTF2IRD2 | 0,93894969 |
| ENSG00000 gene | 0,94203452 | 0,00861978 | 0,27352313 | HNRNPAB | -0,0861482 |
| ENSG00000 gene | 1,05132317 | 0,00865461 | 0,27352313 | GANAB | 0,07220621 |
| ENSG00000 gene | 1,36891857 | 0,00867306 | 0,27352313 | DPP8 | 0,45303663 |
| ENSG00000 gene | 0,86614767 | 0,00867898 | 0,27352313 | RPS29 | -0,2073151 |
| ENSG00000 gene | 0,84349753 | 0,00872737 | 0,27414673 | AC115223.1 | -0,2455442 |
| ENSG00000 gene | 1,18671432 | 0,00875241 | 0,27414673 | CEMP1 | 0,24697267 |
| ENSG00000 gene | 0,77182391 | 0,00881997 | 0,27414673 | FARP1-AS1 | -0,3736564 |
| ENSG00000 gene | 1,23400459 | 0,00884236 | 0,27414673 | FTH1P11 | 0,30334777 |
| ENSG00000 gene | 1,57147599 | 0,00887184 | 0,27414673 | PABPC4-AS | 0,65212023 |
| ENSG00000 gene | 1,12183726 | 0,00888126 | 0,27414673 | MARF1 | 0,1658634 |
| ENSG00000 gene | 1,37059687 | 0,00889577 | 0,27414673 | AL732414.1 | 0,4548043 |
| ENSG00000 gene | 0,92329454 | 0,00889591 | 0,27414673 | SGSM3 | -0,1151371 |
| ENSG00000 gene | 0,77845615 | 0,00893903 | 0,27471464 | SGMS2 | -0,3613123 |
| ENSG00000 gene | 0,61943748 | 0,0090773 | 0,27819534 | DCTPP1 | -0,6909694 |
| ENSG00000 gene | 1,1912511 | 0,00914586 | 0,27952658 | DLGAP1-AS | 0,25247755 |
| ENSG00000 gene | 0,90960637 | 0,00921277 | 0,27985893 | CCDC12 | -0,1366857 |
| ENSG00000 gene | 1,28100612 | 0,00921674 | 0,27985893 | CYP51A1 | 0,35727736 |
| ENSG00000 gene | 1,11060405 | 0,0092322 | 0,27985893 | ATE1 | 0,15134456 |
| ENSG00000 gene | 2,17909773 | 0,0093536 | 0,28276837 | OSBPL6 | 1,1237309 |
| ENSG00000 gene | 0,57453264 | 0,00939992 | 0,28339871 | KCTD7 | -0,7995392 |
| ENSG00000 gene | 0,72458291 | 0,00958712 | 0,28748462 | DENND3 | -0,4647773 |
| ENSG00000 gene | 1,12361872 | 0,00958713 | 0,28748462 | TP53INP1 | 0,16815257 |
| ENSG00000 gene | 1,53368558 | 0,00963846 | 0,28824697 | COX7A2L | 0,61700275 |
| ENSG00000 gene | 0,83589306 | 0,00966561 | 0,28828396 | SCD5 | -0,2586097 |
| ENSG00000 gene | 1,0557168 | 0,009751 | 0,29005302 | AP3B1 | 0,07822288 |
| ENSG00000 gene | 1,07627315 | 0,00978144 | 0,29018269 | FKBP15 | 0,10604427 |
| ENSG00000 gene | 1,17058729 | 0,0099005 | 0,29293372 | ACSL3 | 0,22723252 |
| ENSG00000 gene | 1,20470048 | 0,00993037 | 0,29303815 | ACYP1 | 0,2686745 |
| ENSG00000 gene | 0,92507298 | 0,00995942 | 0,29311783 | RPL27A | -0,1123609 |
| ENSG00000 gene | 0,85951685 | 0,01006545 | 0,29545685 | SLC1A5 | -0,2184022 |
| ENSG00000 gene | 1,49589714 | 0,01012862 | 0,29652868 | AC092683.1 | 0,58101098 |
| ENSG00000 gene | 2,01908954 | 0,01016616 | 0,29684664 | BCL2L2-PAE | 1,01370489 |
| ENSG00000 gene | 0,83482807 | 0,0102104 | 0,29735793 | KHDRBS3 | -0,260449 |
| ENSG00000 gene | 0,90124744 | 0,01031046 | 0,29948798 | CASP3 | -0,1500048 |
| ENSG00000 gene | 0,81944251 | 0,01035838 | 0,29959494 | AC104758.1 | -0,2872853 |
| ENSG00000 gene | 1,08421302 | 0,010368 | 0,29959494 | TAF6 | 0,11664824 |
| ENSG00000 gene | 1,4647142 | 0,01047722 | 0,30196658 | KSR1 | 0,55061919 |
| ENSG00000 gene | 1,07970063 | 0,0105467 | 0,30274761 | CPD | 0,11063135 |
| ENSG00000 gene | 1,11085079 | 0,01055875 | 0,30274761 | RTL6 | 0,15166505 |
| ENSG00000 gene | 0,96448909 | 0,01073644 | 0,30510206 | TPD52L2 | -0,0521632 |
| ENSG00000 gene | 0,95564915 | 0,01073656 | 0,30510206 | ANXA1 | -0,065447 |
| ENSG00000 gene | 0,76667229 | 0,01074941 | 0,30510206 | NME2 | -0,3833181 |
| ENSG00000 gene | 0,87407712 | 0,01075056 | 0,30510206 | MT-TE | -0,1941675 |
| ENSG00000 gene | 0,91767979 | 0,01080654 | 0,30568634 | FHL2 | -0,1239373 |
| ENSG00000 gene | 1,14027406 | 0,0108261 | 0,30568634 | NCOR1 | 0,18938061 |
| ENSG00000 gene | 0,84185567 | 0,01091264 | 0,30692241 | DCTN3 | -0,2483552 |
| ENSG00000 gene | 1,08498379 | 0,01092506 | 0,30692241 | GLG1 | 0,11767349 |
| ENSG00000 gene | 1,08950632 | 0,01096226 | 0,30719189 | ATAD1 | 0,12367456 |

| | | | | | | |
|---|---|---|---|---|---|---|
| ENSG00000 | gene | 2,71666897 | 0,01108542 | 0,30904367 | MIR3658 | 1,44183878 |
| ENSG00000 | gene | 0,89852343 | 0,01113414 | 0,30904367 | ABHD17A | -0,154372 |
| ENSG00000 | gene | 1,89214052 | 0,01113444 | 0,30904367 | EREG | 0,92001924 |
| ENSG00000 | gene | 0,93459449 | 0,01116363 | 0,30904367 | CNOT7 | -0,0975876 |
| ENSG00000 | gene | 1,08907009 | 0,01116724 | 0,30904367 | ATP2C1 | 0,1230968 |
| ENSG00000 | gene | 0,8944357 | 0,01119505 | 0,30904458 | WBP1L | -0,1609503 |
| ENSG00000 | gene | 0,55549207 | 0,01135517 | 0,31198146 | AC010776.2 | -0,8481618 |
| ENSG00000 | gene | 1,3114031 | 0,01138426 | 0,31198146 | CRTC1 | 0,39111121 |
| ENSG00000 | gene | 0,8276296 | 0,01138557 | 0,31198146 | SGMS1 | -0,2729428 |
| ENSG00000 | gene | 0,83732137 | 0,01145059 | 0,3129922 | MICB | -0,2561466 |
| ENSG00000 | gene | 1,3090368 | 0,01150327 | 0,31366135 | FRG1HP | 0,38850565 |
| ENSG00000 | gene | 0,6707608 | 0,01162629 | 0,31510304 | YTHDF2 | -0,5761297 |
| ENSG00000 | gene | 0,76181078 | 0,0116473 | 0,31510304 | PIMREG | -0,3924954 |
| ENSG00000 | gene | 0,86405956 | 0,01166537 | 0,31510304 | LMAN2 | -0,2107973 |
| ENSG00000 | gene | 3,55453793 | 0,01166943 | 0,31510304 | TMEM189-U | 1,82966203 |
| ENSG00000 | gene | 0,94001041 | 0,01174392 | 0,31556078 | ZCCHC24 | -0,0892514 |
| ENSG00000 | gene | 0,91810406 | 0,01175933 | 0,31556078 | STK40 | -0,1232704 |
| ENSG00000 | gene | 0,79933956 | 0,01180636 | 0,31556078 | AL451085.3 | -0,3231196 |
| ENSG00000 | gene | 0,93702837 | 0,01181401 | 0,31556078 | RPS2P46 | -0,0938354 |
| ENSG00000 | gene | 1,59354886 | 0,01182821 | 0,31556078 | DIPK1B | 0,67224325 |
| ENSG00000 | gene | 0,88384044 | 0,01196126 | 0,31717158 | PFN1 | -0,1781421 |
| ENSG00000 | gene | 0,86778784 | 0,01196339 | 0,31717158 | FAM3C | -0,2045857 |
| ENSG00000 | gene | 0,66626229 | 0,01199408 | 0,31717158 | MAP4K3-DT | -0,5858379 |
| ENSG00000 | gene | 0,60083026 | 0,01200263 | 0,31717158 | ABHD11 | -0,7349706 |
| ENSG00000 | gene | 1,3478209 | 0,01213937 | 0,32002475 | TXNRD1 | 0,4306288 |
| ENSG00000 | gene | 0,9297601 | 0,01222963 | 0,32164219 | SCARA3 | -0,1050696 |
| ENSG00000 | gene | 1,62209565 | 0,01231316 | 0,32307531 | PIK3R1 | 0,6978589 |
| ENSG00000 | gene | 0,92392572 | 0,01239856 | 0,32455067 | RPSA | -0,1141512 |
| ENSG00000 | gene | 0,921307 | 0,01248837 | 0,32613395 | SLC20A1 | -0,1182461 |
| ENSG00000 | gene | 0,94243805 | 0,01253849 | 0,32667617 | PA2G4 | -0,0855303 |
| ENSG00000 | gene | 0,8791231 | 0,01257143 | 0,32676893 | SCCPDH | -0,1858629 |
| ENSG00000 | gene | 1,65348484 | 0,01262419 | 0,32737558 | GK-AS1 | 0,72550981 |
| ENSG00000 | gene | 0,8157091 | 0,01265413 | 0,32738891 | AC020909.3 | -0,2938733 |
| ENSG00000 | gene | 1,22878654 | 0,01271994 | 0,32832802 | UTP25 | 0,29723431 |
| ENSG00000 | gene | 1,51304534 | 0,01280989 | 0,32975858 | FGF7P4 | 0,59745522 |
| ENSG00000 | gene | 0,58859167 | 0,01286001 | 0,32975858 | CCDC189 | -0,764661 |
| ENSG00000 | gene | 0,92261887 | 0,01286429 | 0,32975858 | UQCRC1 | -0,1161933 |
| ENSG00000 | gene | 0,88184664 | 0,01303004 | 0,33106629 | PROCR | -0,1814003 |
| ENSG00000 | gene | 0,9318857 | 0,01303387 | 0,33106629 | LRRC75A | -0,1017751 |
| ENSG00000 | gene | 0,84647147 | 0,01303807 | 0,33106629 | DPAGT1 | -0,2404667 |
| ENSG00000 | gene | 1,0578491 | 0,01312049 | 0,33106629 | CLIP3 | 0,08113384 |
| ENSG00000 | gene | 1,1148567 | 0,01314822 | 0,33106629 | MEG3 | 0,15685829 |
| ENSG00000 | gene | 1,10022196 | 0,01315744 | 0,33106629 | UBR2 | 0,13779461 |
| ENSG00000 | gene | 0,85676061 | 0,0132158 | 0,33106629 | MICU2 | -0,2230359 |
| ENSG00000 | gene | 0,94190823 | 0,01327007 | 0,33106629 | TEAD3 | -0,0863416 |
| ENSG00000 | gene | 0,86087805 | 0,01330258 | 0,33106629 | KIRREL3 | -0,2161192 |
| ENSG00000 | gene | 1,14625518 | 0,01335024 | 0,33106629 | TBX5 | 0,19692826 |
| ENSG00000 | gene | 1,28758247 | 0,01344196 | 0,33106629 | WNT5A-AS1 | 0,36466485 |
| ENSG00000 | gene | 0,88402122 | 0,01345124 | 0,33106629 | FAM104A | -0,1778471 |
| ENSG00000 | gene | 0,68579845 | 0,01346905 | 0,33106629 | HIGD1A | -0,5441435 |
| ENSG00000 | gene | 0,55126398 | 0,01350306 | 0,33106629 | AL161787.1 | -0,8591847 |
| ENSG00000 | gene | 0,91109984 | 0,01350718 | 0,33106629 | PHLDA3 | -0,1343189 |
| ENSG00000 | gene | 1,19073418 | 0,01353171 | 0,33106629 | PCM1 | 0,25185138 |
| ENSG00000 | gene | 0,93765475 | 0,01354732 | 0,33106629 | MIF-AS1 | -0,0928713 |
| ENSG00000 | gene | 0,93356734 | 0,0135634 | 0,33106629 | RPL18A | -0,099174 |
| ENSG00000 | gene | 0,93926871 | 0,01358747 | 0,33106629 | TSR2 | -0,0903901 |
| ENSG00000 | gene | 0,80774799 | 0,01358898 | 0,33106629 | GADD45A | -0,3080228 |

| | | | | | |
|---|---|---|---|---|---|
| ENSG00000 gene | 0,72153522 | 0,01360526 | 0,33106629 | TJP2 | -0,4708583 |
| ENSG00000 gene | 1,37465344 | 0,01361815 | 0,33106629 | FNTA | 0,45906795 |
| ENSG00000 gene | 0,85471388 | 0,0136397 | 0,33106629 | MT-TG | -0,2264865 |
| ENSG00000 gene | 0,85388702 | 0,01365277 | 0,33106629 | RPL12P4 | -0,2278829 |
| ENSG00000 gene | 0,90875024 | 0,01366157 | 0,33106629 | RPS7 | -0,1380443 |
| ENSG00000 gene | 1,67973004 | 0,01373445 | 0,33106629 | FAM167A | 0,74822938 |
| ENSG00000 gene | 0,86831449 | 0,01373501 | 0,33106629 | ZFAND6 | -0,2037104 |
| ENSG00000 gene | 0,82957349 | 0,01374855 | 0,33106629 | NDUFS3 | -0,2695583 |
| ENSG00000 gene | 0,83494593 | 0,01380407 | 0,33127342 | ATP5MC3 | -0,2602453 |
| ENSG00000 gene | 1,19376322 | 0,01383905 | 0,33127342 | ZFYVE26 | 0,25551672 |
| ENSG00000 gene | 0,90240974 | 0,01384648 | 0,33127342 | RPS15A | -0,1481455 |
| ENSG00000 gene | 1,4340588 | 0,01388469 | 0,33147456 | LIAS | 0,52010418 |
| ENSG00000 gene | 0,90247816 | 0,01400435 | 0,33293534 | POP7 | -0,1480361 |
| ENSG00000 gene | 0,75704037 | 0,01403025 | 0,33293534 | CLTB | -0,4015579 |
| ENSG00000 gene | 0,89575285 | 0,01403566 | 0,33293534 | MRPL14 | -0,1588274 |
| ENSG00000 gene | 0,91993932 | 0,01409294 | 0,33358284 | HARS1 | -0,1203894 |
| ENSG00000 gene | 0,83334242 | 0,0141273 | 0,33366293 | SSR3 | -0,2630187 |
| ENSG00000 gene | 0,66065543 | 0,01415631 | 0,33366293 | C4orf48 | -0,5980301 |
| ENSG00000 gene | 0,83205834 | 0,0142526 | 0,33503613 | ALCAM | -0,2652434 |
| ENSG00000 gene | 1,40584831 | 0,0142748 | 0,33503613 | GNG11 | 0,49144094 |
| ENSG00000 gene | 1,7980317 | 0,01442249 | 0,33777337 | ZNF331 | 0,84641845 |
| ENSG00000 gene | 0,89502886 | 0,01447359 | 0,33777337 | RPS2 | -0,1599939 |
| ENSG00000 gene | 1,27551873 | 0,01449298 | 0,33777337 | PRICKLE2 | 0,35108408 |
| ENSG00000 gene | 1,83449652 | 0,01457067 | 0,33777337 | RAB31 | 0,87538417 |
| ENSG00000 gene | 0,81556515 | 0,01462165 | 0,33777337 | RPS15 | -0,294128 |
| ENSG00000 gene | 1,1510448 | 0,01465628 | 0,33777337 | CSNK2A3 | 0,20294398 |
| ENSG00000 gene | 1,12207447 | 0,01467151 | 0,33777337 | ANKRD17 | 0,16616842 |
| ENSG00000 gene | 0,86522119 | 0,01468367 | 0,33777337 | MBNL1 | -0,2088591 |
| ENSG00000 gene | 1,18674011 | 0,01470665 | 0,33777337 | EIF4BP7 | 0,24700403 |
| ENSG00000 gene | 0,93337631 | 0,01473241 | 0,33777337 | CCT5 | -0,0994693 |
| ENSG00000 gene | 0,89878653 | 0,01474515 | 0,33777337 | RBX1 | -0,1539496 |
| ENSG00000 gene | 0,88548329 | 0,01475576 | 0,33777337 | GPN3 | -0,175463 |
| ENSG00000 gene | 0,88294022 | 0,01481832 | 0,33796659 | SCARF2 | -0,1796123 |
| ENSG00000 gene | 0,92496395 | 0,01485112 | 0,33796659 | RPS11 | -0,112531 |
| ENSG00000 gene | 0,77999515 | 0,01485534 | 0,33796659 | MAEA | -0,3584629 |
| ENSG00000 gene | 1,13573762 | 0,01493316 | 0,33904361 | SLC23A2 | 0,18362958 |
| ENSG00000 gene | 0,43062807 | 0,01503842 | 0,34028328 | OAZ3 | -1,2154857 |
| ENSG00000 gene | 1,56152779 | 0,01504893 | 0,34028328 | CEP290 | 0,64295824 |
| ENSG00000 gene | 0,83925022 | 0,01510834 | 0,34070936 | CLEC11A | -0,2528271 |
| ENSG00000 gene | 0,87091959 | 0,01513165 | 0,34070936 | ZNF593 | -0,1993886 |
| ENSG00000 gene | 1,12206848 | 0,01516098 | 0,34070936 | AL158212.3 | 0,16616072 |
| ENSG00000 gene | 0,94768476 | 0,01519028 | 0,34070936 | XRCC6 | -0,0775208 |
| ENSG00000 gene | 0,85091239 | 0,01525998 | 0,34158403 | RTN4 | -0,2329175 |
| ENSG00000 gene | 1,08238782 | 0,01529804 | 0,34174829 | HNRNPU | 0,11421752 |
| ENSG00000 gene | 1,95173539 | 0,01550959 | 0,34535861 | TCIM | 0,96475747 |
| ENSG00000 gene | 0,69079613 | 0,0155271 | 0,34535861 | HMMR | -0,5336681 |
| ENSG00000 gene | 1,53444343 | 0,01557811 | 0,34535861 | ABHD17C | 0,61771546 |
| ENSG00000 gene | 0,82915382 | 0,01558382 | 0,34535861 | SNRPD1 | -0,2702883 |
| ENSG00000 gene | 0,69407586 | 0,01571921 | 0,34679631 | ZNF266 | -0,5268347 |
| ENSG00000 gene | 0,86261162 | 0,01576674 | 0,34679631 | SERPINE1 | -0,2132169 |
| ENSG00000 gene | 0,75171442 | 0,01581839 | 0,34679631 | AP001107.6 | -0,4117434 |
| ENSG00000 gene | 1,67156139 | 0,01587651 | 0,34679631 | LYST | 0,74119634 |
| ENSG00000 gene | 0,94107228 | 0,01587796 | 0,34679631 | NT5DC2 | -0,0876226 |
| ENSG00000 gene | 0,77802176 | 0,01587926 | 0,34679631 | AC002347.2 | -0,3621176 |
| ENSG00000 gene | 0,92771301 | 0,01590917 | 0,34679631 | PTCD1 | -0,1082495 |
| ENSG00000 gene | 0,89214657 | 0,0159169 | 0,34679631 | ORAI1 | -0,1646473 |
| ENSG00000 gene | 1,29543538 | 0,01593802 | 0,34679631 | KLF9 | 0,37343705 |

| | | | | | | |
|---|---|---|---|---|---|---|
| ENSG00000 | gene | 1,60581308 | 0,01596042 | 0,34679631 | MRRF | 0,68330397 |
| ENSG00000 | gene | 0,77671429 | 0,01603529 | 0,34749372 | MRPS28 | -0,3645441 |
| ENSG00000 | gene | 0,56952508 | 0,01606649 | 0,34749372 | ZNF286A | -0,8121687 |
| ENSG00000 | gene | 0,90471412 | 0,01608623 | 0,34749372 | PGM2 | -0,1444661 |
| ENSG00000 | gene | 0,89938418 | 0,01622677 | 0,34929777 | MAD2L2 | -0,1529906 |
| ENSG00000 | gene | 1,37626946 | 0,01623253 | 0,34929777 | HIF1A-AS1 | 0,46076297 |
| ENSG00000 | gene | 0,81203639 | 0,01639456 | 0,35207536 | SMARCD3 | -0,3003837 |
| ENSG00000 | gene | 0,8933569 | 0,01642491 | 0,35207536 | AC009119.1 | -0,1626914 |
| ENSG00000 | gene | 0,61821484 | 0,01647678 | 0,3525081 | NKAPD1 | -0,6938198 |
| ENSG00000 | gene | 0,60955656 | 0,01658908 | 0,3538689 | TTLL4 | -0,714168 |
| ENSG00000 | gene | 1,50167186 | 0,01664316 | 0,3538689 | ENC1 | 0,58656959 |
| ENSG00000 | gene | 1,44005777 | 0,01671187 | 0,3538689 | CD55 | 0,52612669 |
| ENSG00000 | gene | 2,01948699 | 0,01671332 | 0,3538689 | ATF3 | 1,01398885 |
| ENSG00000 | gene | 0,91662702 | 0,0167229 | 0,3538689 | WBP4 | -0,1255933 |
| ENSG00000 | gene | 0,79284886 | 0,01676136 | 0,3538689 | NDUFAF3 | -0,3348822 |
| ENSG00000 | gene | 1,86735855 | 0,01679187 | 0,3538689 | CRAMP1 | 0,90099897 |
| ENSG00000 | gene | 0,94740431 | 0,01679486 | 0,3538689 | PTP4A2 | -0,0779479 |
| ENSG00000 | gene | 1,12957128 | 0,01684028 | 0,35405252 | PURB | 0,17577531 |
| ENSG00000 | gene | 0,67135788 | 0,01686722 | 0,35405252 | CFL2 | -0,5748461 |
| ENSG00000 | gene | 0,89804238 | 0,01694452 | 0,3546422 | MRPS12 | -0,1551446 |
| ENSG00000 | gene | 1,18764911 | 0,01701866 | 0,3546422 | CLDN23 | 0,24810866 |
| ENSG00000 | gene | 0,92724203 | 0,01702032 | 0,3546422 | TMED9 | -0,1089821 |
| ENSG00000 | gene | 0,8510287 | 0,01704178 | 0,3546422 | ARF5 | -0,2327203 |
| ENSG00000 | gene | 0,78691679 | 0,0170547 | 0,3546422 | SHQ1 | -0,345717 |
| ENSG00000 | gene | 1,49584972 | 0,01709305 | 0,3547764 | RNF31 | 0,58096525 |
| ENSG00000 | gene | 0,9455049 | 0,0171602 | 0,35505605 | IKBIP | -0,0808432 |
| ENSG00000 | gene | 0,88198665 | 0,01717035 | 0,35505605 | SNHG6 | -0,1811713 |
| ENSG00000 | gene | 0,85137257 | 0,01723623 | 0,35575716 | AC016739.1 | -0,2321375 |
| ENSG00000 | gene | 0,89706016 | 0,01744409 | 0,35872046 | TMEM123 | -0,1567234 |
| ENSG00000 | gene | 0,71630755 | 0,01744429 | 0,35872046 | LNCOG | -0,4813489 |
| ENSG00000 | gene | 0,83324283 | 0,01753971 | 0,35921212 | AC091607.2 | -0,2631911 |
| ENSG00000 | gene | 2,17234377 | 0,01756212 | 0,35921212 | CSF3 | 1,11925242 |
| ENSG00000 | gene | 1,29594564 | 0,01756507 | 0,35921212 | ABHD2 | 0,3740052 |
| ENSG00000 | gene | 1,4756029 | 0,01762515 | 0,35977953 | BBS12 | 0,56130453 |
| ENSG00000 | gene | 0,8846383 | 0,01769913 | 0,36025307 | RPL13AP5 | -0,1768404 |
| ENSG00000 | gene | 0,95092837 | 0,0177653 | 0,36025307 | AFG3L2 | -0,0725914 |
| ENSG00000 | gene | 0,91453141 | 0,01783847 | 0,36025307 | RPL24 | -0,1288954 |
| ENSG00000 | gene | 0,90513455 | 0,01784262 | 0,36025307 | RHBDD3 | -0,1437958 |
| ENSG00000 | gene | 1,30512609 | 0,01784842 | 0,36025307 | AC007319.1 | 0,38418919 |
| ENSG00000 | gene | 0,9548884 | 0,01787044 | 0,36025307 | EIF2S1 | -0,066596 |
| ENSG00000 | gene | 0,48538622 | 0,01788312 | 0,36025307 | AC008522.1 | -1,042795 |
| ENSG00000 | gene | 0,81752435 | 0,01790741 | 0,36025307 | HACD2 | -0,2906664 |
| ENSG00000 | gene | 0,8551956 | 0,01794228 | 0,36030296 | MARCHF4 | -0,2256737 |
| ENSG00000 | gene | 0,59204929 | 0,01807271 | 0,36037185 | USP41 | -0,7562108 |
| ENSG00000 | gene | 0,64618919 | 0,01808832 | 0,36037185 | PRXL2C | -0,6299715 |
| ENSG00000 | gene | 0,9471825 | 0,01810112 | 0,36037185 | PRMT1 | -0,0782857 |
| ENSG00000 | gene | 1,52993869 | 0,01810454 | 0,36037185 | DHPS | 0,61347384 |
| ENSG00000 | gene | 0,86348754 | 0,01825206 | 0,36037185 | NDUFB8 | -0,2117527 |
| ENSG00000 | gene | 0,87885973 | 0,01825642 | 0,36037185 | KRT10 | -0,1862952 |
| ENSG00000 | gene | 0,72845723 | 0,01827274 | 0,36037185 | AL445183.3 | -0,4570838 |
| ENSG00000 | gene | 0,79806567 | 0,01827428 | 0,36037185 | AC092069.1 | -0,3254206 |
| ENSG00000 | gene | 1,05694304 | 0,0182774 | 0,36037185 | EIF4G3 | 0,07989764 |
| ENSG00000 | gene | 0,8943626 | 0,01833445 | 0,36037185 | MRPS24 | -0,1610682 |
| ENSG00000 | gene | 0,64368205 | 0,01834713 | 0,36037185 | FTX | -0,6355799 |
| ENSG00000 | gene | 1,09587322 | 0,01835748 | 0,36037185 | SMC5 | 0,13208091 |
| ENSG00000 | gene | 0,47203422 | 0,01836682 | 0,36037185 | AC087190.3 | -1,0830367 |
| ENSG00000 | gene | 1,08727886 | 0,01840893 | 0,36056225 | DDX42 | 0,120722 |

| | | | | | | |
|---|---|---|---|---|---|---|
| ENSG00000 | gene | 0,88924725 | 0,01850294 | 0,36106541 | RPL34 | -0,1693435 |
| ENSG00000 | gene | 0,8154222 | 0,01854061 | 0,36106541 | CD151 | -0,2943809 |
| ENSG00000 | gene | 1,14986448 | 0,01862771 | 0,36106541 | FAM189B | 0,20146384 |
| ENSG00000 | gene | 0,87859766 | 0,01867026 | 0,36106541 | RPSAP19 | -0,1867254 |
| ENSG00000 | gene | 1,12805508 | 0,01868032 | 0,36106541 | MVD | 0,17383751 |
| ENSG00000 | gene | 1,18457953 | 0,01868953 | 0,36106541 | YAP1 | 0,24437507 |
| ENSG00000 | gene | 1,08483036 | 0,01874554 | 0,36106541 | SLC26A2 | 0,11746945 |
| ENSG00000 | gene | 1,40795927 | 0,01875447 | 0,36106541 | GON4L | 0,4936056 |
| ENSG00000 | gene | 1,36813399 | 0,01879822 | 0,36106541 | ADAMTS6 | 0,45220953 |
| ENSG00000 | gene | 1,32833503 | 0,01880345 | 0,36106541 | SYNE1 | 0,40961907 |
| ENSG00000 | gene | 1,34582085 | 0,01880983 | 0,36106541 | FAM120A | 0,42848638 |
| ENSG00000 | gene | 1,39756681 | 0,01883984 | 0,36106541 | TOP3B | 0,48291726 |
| ENSG00000 | gene | 0,74512847 | 0,01885654 | 0,36106541 | TUBB2B | -0,4244389 |
| ENSG00000 | gene | 0,90851368 | 0,01893439 | 0,36153106 | TUBA1C | -0,1384199 |
| ENSG00000 | gene | 0,69052922 | 0,01894585 | 0,36153106 | ANAPC10 | -0,5342256 |
| ENSG00000 | gene | 0,93298641 | 0,01904123 | 0,36227733 | ZFP36L2 | -0,100072 |
| ENSG00000 | gene | 1,1029631 | 0,01905009 | 0,36227733 | GRSF1 | 0,14138452 |
| ENSG00000 | gene | 0,75762091 | 0,01908938 | 0,36240509 | ZFAND5 | -0,4004519 |
| ENSG00000 | gene | 0,88643392 | 0,01914627 | 0,36286592 | GALNT18 | -0,173915 |
| ENSG00000 | gene | 1,05639871 | 0,01923596 | 0,36394576 | DNAJC13 | 0,07915444 |
| ENSG00000 | gene | 1,44878945 | 0,01936882 | 0,36583717 | AL162274.2 | 0,53484795 |
| ENSG00000 | gene | 0,61531871 | 0,01943506 | 0,36611129 | ZNF414 | -0,7005942 |
| ENSG00000 | gene | 1,06990481 | 0,01944915 | 0,36611129 | GTF3C3 | 0,09748245 |
| ENSG00000 | gene | 0,89889193 | 0,01953937 | 0,36718834 | EMP3 | -0,1537804 |
| ENSG00000 | gene | 1,08688146 | 0,01957813 | 0,36729619 | NBAS | 0,1201946 |
| ENSG00000 | gene | 0,86704544 | 0,01970036 | 0,36844308 | S100A16 | -0,2058205 |
| ENSG00000 | gene | 0,91679216 | 0,01972814 | 0,36844308 | DIMT1 | -0,1253334 |
| ENSG00000 | gene | 1,11575487 | 0,01979861 | 0,36844308 | DHX32 | 0,15802011 |
| ENSG00000 | gene | 0,90119089 | 0,01980851 | 0,36844308 | RPLP2 | -0,1500954 |
| ENSG00000 | gene | 1,19803794 | 0,01983781 | 0,36844308 | AC007750.1 | 0,2606736 |
| ENSG00000 | gene | 1,23421779 | 0,01986188 | 0,36844308 | DNMBP | 0,30359699 |
| ENSG00000 | gene | 0,79905054 | 0,01990911 | 0,36844308 | GGCT | -0,3236413 |
| ENSG00000 | gene | 1,43052702 | 0,01990918 | 0,36844308 | PLEKHG5 | 0,51654674 |
| ENSG00000 | gene | 0,94171924 | 0,01993732 | 0,36844308 | STK17A | -0,0866311 |
| ENSG00000 | gene | 0,9005741 | 0,02000419 | 0,36906564 | CD81 | -0,1510831 |
| ENSG00000 | gene | 0,90888953 | 0,0200788 | 0,36982897 | AP3M1 | -0,1378231 |
| ENSG00000 | gene | 0,89189025 | 0,02017976 | 0,37107409 | DBN1 | -0,1650619 |
| ENSG00000 | gene | 2,36006176 | 0,02034209 | 0,37180823 | AC006001.4 | 1,23882462 |
| ENSG00000 | gene | 0,90454961 | 0,02034529 | 0,37180823 | PPP1R2 | -0,1447285 |
| ENSG00000 | gene | 1,7806983 | 0,02035732 | 0,37180823 | MIR1281 | 0,8324431 |
| ENSG00000 | gene | 0,85328888 | 0,02040003 | 0,37180823 | MFSD5 | -0,2288938 |
| ENSG00000 | gene | 0,94093241 | 0,0204092 | 0,37180823 | OLA1 | -0,087837 |
| ENSG00000 | gene | 0,35746099 | 0,02042143 | 0,37180823 | ATG10 | -1,4841423 |
| ENSG00000 | gene | 1,08106608 | 0,02050378 | 0,37180823 | SKAP2 | 0,11245471 |
| ENSG00000 | gene | 1,20497506 | 0,02054233 | 0,37180823 | RC3H2 | 0,26900329 |
| ENSG00000 | gene | 0,61208115 | 0,02056079 | 0,37180823 | CC2D1B | -0,7082052 |
| ENSG00000 | gene | 0,830698 | 0,02059809 | 0,37180823 | PPA1 | -0,267604 |
| ENSG00000 | gene | 0,73636633 | 0,02060103 | 0,37180823 | PVR | -0,4415044 |
| ENSG00000 | gene | 0,94684272 | 0,02064335 | 0,37180823 | ALDH18A1 | -0,0788033 |
| ENSG00000 | gene | 1,04306568 | 0,02065416 | 0,37180823 | GCN1 | 0,06083001 |
| ENSG00000 | gene | 1,59757467 | 0,02081659 | 0,37412689 | AC104411.1 | 0,67588337 |
| ENSG00000 | gene | 1,11736315 | 0,02098806 | 0,37581011 | FTH1P2 | 0,16009815 |
| ENSG00000 | gene | 1,6437701 | 0,02100625 | 0,37581011 | IFNAR2 | 0,71700854 |
| ENSG00000 | gene | 1,11203575 | 0,02101159 | 0,37581011 | ATXN1 | 0,15320317 |
| ENSG00000 | gene | 0,90814218 | 0,02119597 | 0,37849952 | RING1 | -0,1390099 |
| ENSG00000 | gene | 1,0518682 | 0,02124762 | 0,37881365 | PABPC1 | 0,07295395 |
| ENSG00000 | gene | 1,12371005 | 0,02134948 | 0,3800208 | PTPN23 | 0,16826982 |

| | | | | | |
|---|---|---|---|---|---|
| ENSG00000 gene | 0,71609914 | 0,02140617 | 0,38015323 | NUAK2 | -0,4817688 |
| ENSG00000 gene | 0,90579179 | 0,02142527 | 0,38015323 | UQCRC2 | -0,1427486 |
| ENSG00000 gene | 0,67519815 | 0,02148024 | 0,3805217 | L3MBTL2 | -0,5666171 |
| ENSG00000 gene | 0,8422008 | 0,02153346 | 0,38085804 | CUEDC1 | -0,2477638 |
| ENSG00000 gene | 0,89618623 | 0,02171619 | 0,3834804 | PAPSS1 | -0,1581295 |
| ENSG00000 gene | 0,95313414 | 0,02182163 | 0,38356089 | TMED7 | -0,0692488 |
| ENSG00000 gene | 0,79195141 | 0,02183893 | 0,38356089 | ARCN1 | -0,3365162 |
| ENSG00000 gene | 1,62116284 | 0,02183895 | 0,38356089 | PCYT2 | 0,69702901 |
| ENSG00000 gene | 0,87163768 | 0,02185866 | 0,38356089 | PSMA2 | -0,1981995 |
| ENSG00000 gene | 0,90250324 | 0,02200083 | 0,38544755 | CDC26 | -0,147996 |
| ENSG00000 gene | 1,14984357 | 0,02206376 | 0,38566197 | LURAP1L | 0,2014376 |
| ENSG00000 gene | 0,92572996 | 0,0220824 | 0,38566197 | RPL9P7 | -0,1113367 |
| ENSG00000 gene | 0,85044541 | 0,022127 | 0,38583514 | PTS | -0,2337095 |
| ENSG00000 gene | 0,9476993 | 0,02223849 | 0,38717244 | NDFIP1 | -0,0774987 |
| ENSG00000 gene | 1,32852351 | 0,0224358 | 0,3894412 | AL354811.1 | 0,40982376 |
| ENSG00000 gene | 0,91883153 | 0,02243881 | 0,3894412 | UBL4A | -0,1221277 |
| ENSG00000 gene | 1,13956735 | 0,02253374 | 0,38996838 | FTH1P5 | 0,1884862 |
| ENSG00000 gene | 0,91822218 | 0,02257809 | 0,38996838 | RPL13 | -0,1230848 |
| ENSG00000 gene | 0,91506123 | 0,0225873 | 0,38996838 | BAD | -0,1280598 |
| ENSG00000 gene | 1,25320937 | 0,0226094 | 0,38996838 | CEP164 | 0,32562746 |
| ENSG00000 gene | 1,18496037 | 0,02270968 | 0,39109163 | CDC25B | 0,24483881 |
| ENSG00000 gene | 0,88178232 | 0,02276632 | 0,39146098 | INPP5K | -0,1815055 |
| ENSG00000 gene | 0,90563957 | 0,02297659 | 0,39391319 | ELOF1 | -0,1429911 |
| ENSG00000 gene | 1,54446261 | 0,02297974 | 0,39391319 | PTPRN | 0,62710495 |
| ENSG00000 gene | 1,09667432 | 0,02309118 | 0,39424816 | XRCC1 | 0,13313516 |
| ENSG00000 gene | 0,77822917 | 0,02309989 | 0,39424816 | DMTF1 | -0,361733 |
| ENSG00000 gene | 0,80077676 | 0,0231056 | 0,39424816 | POLR3H | -0,320528 |
| ENSG00000 gene | 1,05286591 | 0,02314153 | 0,39425648 | PNMA1 | 0,07432171 |
| ENSG00000 gene | 1,44157513 | 0,02322401 | 0,3949306 | CDC42SE1 | 0,52764603 |
| ENSG00000 gene | 0,9150674 | 0,02325209 | 0,3949306 | ACAP3 | -0,1280501 |
| ENSG00000 gene | 1,55421857 | 0,02334689 | 0,39513194 | FBLIM1 | 0,63618941 |
| ENSG00000 gene | 0,93300107 | 0,02337077 | 0,39513194 | AC010883.3 | -0,1000494 |
| ENSG00000 gene | 1,14277247 | 0,0233846 | 0,39513194 | SART3 | 0,19253818 |
| ENSG00000 gene | 0,94626203 | 0,02340602 | 0,39513194 | FBXL3 | -0,0796884 |
| ENSG00000 gene | 1,11811348 | 0,0235027 | 0,39616293 | DHCR7 | 0,16106661 |
| ENSG00000 gene | 0,91733011 | 0,02364026 | 0,39787889 | RPL12 | -0,1244871 |
| ENSG00000 gene | 0,53552811 | 0,02380123 | 0,39807763 | IFT22 | -0,9009658 |
| ENSG00000 gene | 0,8595987 | 0,02380783 | 0,39807763 | CENPT | -0,2182648 |
| ENSG00000 gene | 0,92387801 | 0,02384442 | 0,39807763 | RPS7P1 | -0,1142257 |
| ENSG00000 gene | 0,54454242 | 0,02386023 | 0,39807763 | AP003068.3 | -0,8768836 |
| ENSG00000 gene | 1,06102266 | 0,02386483 | 0,39807763 | SHLD2 | 0,08545547 |
| ENSG00000 gene | 1,09417634 | 0,02388752 | 0,39807763 | CLSTN3 | 0,12984527 |
| ENSG00000 gene | 0,89473904 | 0,02395183 | 0,39807763 | EEF1G | -0,1604611 |
| ENSG00000 gene | 1,07035036 | 0,02399968 | 0,39807763 | USP46 | 0,09808311 |
| ENSG00000 gene | 0,89535879 | 0,02401996 | 0,39807763 | AC073861.1 | -0,1594622 |
| ENSG00000 gene | 0,90589298 | 0,02402175 | 0,39807763 | RPL23 | -0,1425875 |
| ENSG00000 gene | 0,85931196 | 0,02404568 | 0,39807763 | LINC01943 | -0,2187461 |
| ENSG00000 gene | 0,80585203 | 0,02412543 | 0,39880447 | ZFPM2-AS1 | -0,3114131 |
| ENSG00000 gene | 0,86760346 | 0,02422308 | 0,39982455 | SEC61B | -0,2048923 |
| ENSG00000 gene | 1,25944936 | 0,02438541 | 0,40190774 | H3-5 | 0,33279311 |
| ENSG00000 gene | 0,91674499 | 0,02464068 | 0,40551423 | RPS8 | -0,1254076 |
| ENSG00000 gene | 0,66608726 | 0,0246841 | 0,40562863 | MSRB3 | -0,5862169 |
| ENSG00000 gene | 1,37613821 | 0,02488277 | 0,40829035 | ZKSCAN8 | 0,46062537 |
| ENSG00000 gene | 1,08326272 | 0,02502125 | 0,40995781 | NLE1 | 0,11538317 |
| ENSG00000 gene | 1,06825784 | 0,02515633 | 0,41069652 | KIAA0232 | 0,09525991 |
| ENSG00000 gene | 0,93407146 | 0,02515739 | 0,41069652 | RSPRY1 | -0,0983952 |
| ENSG00000 gene | 1,3942074 | 0,02517708 | 0,41069652 | FRMD8 | 0,47944519 |

| | | | | | |
|---|---|---|---|---|---|
| ENSG00000 gene | 0,8249925 | 0,02525785 | 0,41141078 | MRPL16 | -0,2775471 |
| ENSG00000 gene | 0,62517387 | 0,02530251 | 0,41153571 | TES | -0,6776706 |
| ENSG00000 gene | 1,16041461 | 0,02539765 | 0,41167279 | CD109 | 0,21464037 |
| ENSG00000 gene | 1,12352102 | 0,02540983 | 0,41167279 | AC069234.2 | 0,16802712 |
| ENSG00000 gene | 0,91739076 | 0,02542195 | 0,41167279 | TXN | -0,1243917 |
| ENSG00000 gene | 1,63125581 | 0,02554225 | 0,41285525 | RASD1 | 0,70598304 |
| ENSG00000 gene | 0,89690579 | 0,02556919 | 0,41285525 | PSMA6 | -0,1569716 |
| ENSG00000 gene | 1,67119804 | 0,02563576 | 0,41303751 | TTC39B | 0,74088271 |
| ENSG00000 gene | 1,04719975 | 0,02566014 | 0,41303751 | LAMP2 | 0,06653666 |
| ENSG00000 gene | 0,81904431 | 0,02569186 | 0,41303751 | FP671120.7 | -0,2879866 |
| ENSG00000 gene | 0,87263329 | 0,02576935 | 0,41368543 | RPL35 | -0,1965526 |
| ENSG00000 gene | 1,75949723 | 0,02584388 | 0,41400347 | AC016629.3 | 0,81516324 |
| ENSG00000 gene | 0,7830537 | 0,02587122 | 0,41400347 | LMBR1 | -0,3528168 |
| ENSG00000 gene | 1,09633125 | 0,02590134 | 0,41400347 | UBAP2L | 0,13268377 |
| ENSG00000 gene | 0,86889292 | 0,02593802 | 0,41400347 | MPC2 | -0,2027497 |
| ENSG00000 gene | 0,68361158 | 0,02611072 | 0,41571683 | ASDURF | -0,5487513 |
| ENSG00000 gene | 0,89764096 | 0,02613431 | 0,41571683 | RPSAP58 | -0,1557896 |
| ENSG00000 gene | 1,41155685 | 0,02615746 | 0,41571683 | AC004264.1 | 0,49728724 |
| ENSG00000 gene | 1,24571152 | 0,02623298 | 0,41632229 | PACS1 | 0,31697001 |
| ENSG00000 gene | 0,4558919 | 0,02627478 | 0,41639161 | LINC00847 | -1,1332363 |
| ENSG00000 gene | 0,61196155 | 0,02634161 | 0,41685377 | SNORD14A | -0,7084871 |
| ENSG00000 gene | 0,86947754 | 0,02638158 | 0,41685377 | EXOSC4 | -0,2017793 |
| ENSG00000 gene | 1,80444179 | 0,02641635 | 0,41685377 | TPTEP2-CSI | 0,8515526 |
| ENSG00000 gene | 0,82668584 | 0,02671137 | 0,42091221 | OGFOD3 | -0,2745889 |
| ENSG00000 gene | 1,19664145 | 0,02685339 | 0,42183137 | STXBP1 | 0,25899095 |
| ENSG00000 gene | 0,63620645 | 0,02687117 | 0,42183137 | GRPEL2 | -0,6524331 |
| ENSG00000 gene | 0,94241202 | 0,0269526 | 0,42183137 | NAA20 | -0,0855701 |
| ENSG00000 gene | 1,3227502 | 0,02697117 | 0,42183137 | BMP6 | 0,40354064 |
| ENSG00000 gene | 1,7830111 | 0,02698618 | 0,42183137 | TBXAS1 | 0,83431568 |
| ENSG00000 gene | 0,68468941 | 0,02708727 | 0,42183137 | PCDHGA12 | -0,5464784 |
| ENSG00000 gene | 1,33611416 | 0,02709907 | 0,42183137 | SOX4 | 0,41804328 |
| ENSG00000 gene | 0,88095206 | 0,02712854 | 0,42183137 | CDC123 | -0,1828646 |
| ENSG00000 gene | 0,77826575 | 0,02717755 | 0,42183137 | FP671120.8 | -0,3616652 |
| ENSG00000 gene | 1,39649597 | 0,02719648 | 0,42183137 | BTBD6 | 0,48181141 |
| ENSG00000 gene | 0,94428306 | 0,02723565 | 0,42183137 | NOL7 | -0,0827087 |
| ENSG00000 gene | 1,25828782 | 0,02727363 | 0,42183137 | HOXA5 | 0,33146196 |
| ENSG00000 gene | 0,79154997 | 0,02729695 | 0,42183137 | C8orf58 | -0,3372477 |
| ENSG00000 gene | 0,89367592 | 0,02730055 | 0,42183137 | B3GAT3 | -0,1621763 |
| ENSG00000 gene | 0,91738148 | 0,02736796 | 0,42192142 | RFTN1 | -0,1244063 |
| ENSG00000 gene | 1,1493885 | 0,02738223 | 0,42192142 | OSBPL8 | 0,20086652 |
| ENSG00000 gene | 0,89109127 | 0,02761763 | 0,42496009 | BID | -0,1663549 |
| ENSG00000 gene | 1,28466001 | 0,02769581 | 0,42557436 | SLC3A2 | 0,3613866 |
| ENSG00000 gene | 0,90079978 | 0,0277833 | 0,42632998 | PARG | -0,1507216 |
| ENSG00000 gene | 0,8967807 | 0,02788695 | 0,42648808 | MAPK3 | -0,1571729 |
| ENSG00000 gene | 0,92365693 | 0,02793632 | 0,42648808 | ACOT9 | -0,114571 |
| ENSG00000 gene | 0,86459488 | 0,02794833 | 0,42648808 | PSMC3IP | -0,2099038 |
| ENSG00000 gene | 0,80881438 | 0,02797673 | 0,42648808 | AC011511.3 | -0,3061195 |
| ENSG00000 gene | 1,11849287 | 0,02802041 | 0,42648808 | ARID1A | 0,16155606 |
| ENSG00000 gene | 1,08671629 | 0,02805048 | 0,42648808 | RNF123 | 0,11997534 |
| ENSG00000 gene | 0,51079343 | 0,02807337 | 0,42648808 | EBPL | -0,9691881 |
| ENSG00000 gene | 0,90682374 | 0,02810029 | 0,42648808 | RNF130 | -0,1411059 |
| ENSG00000 gene | 0,78845888 | 0,02821462 | 0,42731857 | PPP2R3C | -0,3428926 |
| ENSG00000 gene | 0,94665501 | 0,02823183 | 0,42731857 | RAB11B | -0,0790893 |
| ENSG00000 gene | 0,56846946 | 0,02833505 | 0,42793323 | SYS1-DBND | -0,8148453 |
| ENSG00000 gene | 1,61321278 | 0,02837615 | 0,42793323 | CA9 | 0,68993674 |
| ENSG00000 gene | 0,82315398 | 0,02838784 | 0,42793323 | TUFT1 | -0,2807658 |
| ENSG00000 gene | 0,85233481 | 0,0284576 | 0,42840431 | PDCD2 | -0,2305078 |

| | | | | | | |
|---|---|---|---|---|---|---|
| ENSG00000 | gene | 0,89008905 | 0,02855088 | 0,42890117 | DOK4 | -0,1679784 |
| ENSG00000 | gene | 0,89487791 | 0,02857139 | 0,42890117 | AC116533.1 | -0,1602372 |
| ENSG00000 | gene | 0,96005913 | 0,02861994 | 0,42890117 | CHPF | -0,0588048 |
| ENSG00000 | gene | 1,10872708 | 0,02864482 | 0,42890117 | SECISBP2 | 0,14890428 |
| ENSG00000 | gene | 1,51467425 | 0,02888107 | 0,43166686 | TRIM56 | 0,59900756 |
| ENSG00000 | gene | 1,66895725 | 0,02890713 | 0,43166686 | TRABD2A | 0,738947 |
| ENSG00000 | gene | 0,32430866 | 0,0289485 | 0,43170517 | TOMM6 | -1,6245605 |
| ENSG00000 | gene | 0,88404321 | 0,02906935 | 0,43292712 | ALKBH4 | -0,1778112 |
| ENSG00000 | gene | 0,94965355 | 0,02915308 | 0,43310124 | IGF2BP2 | -0,0745268 |
| ENSG00000 | gene | 1,08068813 | 0,02917806 | 0,43310124 | PRPF8 | 0,11195025 |
| ENSG00000 | gene | 1,13363014 | 0,02921527 | 0,43310124 | PGM1 | 0,18095002 |
| ENSG00000 | gene | 0,75602409 | 0,02930459 | 0,43310124 | PAXX | -0,4034959 |
| ENSG00000 | gene | 0,94097765 | 0,02932807 | 0,43310124 | UQCRH | -0,0877676 |
| ENSG00000 | gene | 1,20765322 | 0,02933496 | 0,43310124 | UBBP4 | 0,27220625 |
| ENSG00000 | gene | 1,04963542 | 0,02935356 | 0,43310124 | FAF2 | 0,06988831 |
| ENSG00000 | gene | 1,26580209 | 0,02946011 | 0,43321883 | AC068888.1 | 0,34005185 |
| ENSG00000 | gene | 1,26416048 | 0,02949437 | 0,43321883 | MEIS1 | 0,33817962 |
| ENSG00000 | gene | 1,19888893 | 0,02949947 | 0,43321883 | ABCD4 | 0,261698 |
| ENSG00000 | gene | 0,91373427 | 0,02953269 | 0,43321883 | AHCY | -0,1301534 |
| ENSG00000 | gene | 0,86259572 | 0,02958749 | 0,43321883 | SPATC1L | -0,2132435 |
| ENSG00000 | gene | 1,07112168 | 0,02966804 | 0,43321883 | UNC119B | 0,09912238 |
| ENSG00000 | gene | 0,87771319 | 0,0296881 | 0,43321883 | CENPU | -0,1881785 |
| ENSG00000 | gene | 1,11490879 | 0,02970704 | 0,43321883 | COL4A2-AS | 0,1569257 |
| ENSG00000 | gene | 1,36556978 | 0,029712 | 0,43321883 | FAM13A | 0,44950304 |
| ENSG00000 | gene | 0,88565511 | 0,02993372 | 0,43588043 | E2F1 | -0,1751831 |
| ENSG00000 | gene | 0,67458389 | 0,03001462 | 0,43648711 | HSF4 | -0,5679302 |
| ENSG00000 | gene | 0,90537411 | 0,03017247 | 0,43767911 | GNG5 | -0,143414 |
| ENSG00000 | gene | 0,77441008 | 0,03017527 | 0,43767911 | CTNNAL1 | -0,3688304 |
| ENSG00000 | gene | 0,63547678 | 0,03033495 | 0,43857123 | SIMC1 | -0,6540887 |
| ENSG00000 | gene | 0,71429753 | 0,03034685 | 0,43857123 | SNAPIN | -0,485403 |
| ENSG00000 | gene | 0,87753549 | 0,03035504 | 0,43857123 | AC016596.3 | -0,1884706 |
| ENSG00000 | gene | 1,81822093 | 0,03043075 | 0,43909487 | AL031591.1 | 0,86252751 |
| ENSG00000 | gene | 0,66557933 | 0,03047468 | 0,43915912 | AC243964.4 | -0,5873175 |
| ENSG00000 | gene | 1,73600929 | 0,03053689 | 0,4394863 | NFATC1 | 0,79577467 |
| ENSG00000 | gene | 0,91289102 | 0,03061694 | 0,44006904 | MORF4L1P1 | -0,1314855 |
| ENSG00000 | gene | 1,09944435 | 0,03077687 | 0,44179703 | GNL1 | 0,13677458 |
| ENSG00000 | gene | 0,89620676 | 0,03103061 | 0,44396392 | BUD23 | -0,1580965 |
| ENSG00000 | gene | 1,80578621 | 0,03104196 | 0,44396392 | AP000926.2 | 0,8526271 |
| ENSG00000 | gene | 0,72820059 | 0,03105364 | 0,44396392 | C16orf74 | -0,4575922 |
| ENSG00000 | gene | 1,24931969 | 0,03108745 | 0,44396392 | SOX9 | 0,3211427 |
| ENSG00000 | gene | 1,11106698 | 0,03133159 | 0,44510053 | YBX1P1 | 0,1519458 |
| ENSG00000 | gene | 0,91957308 | 0,03136232 | 0,44510053 | RPS25 | -0,1209639 |
| ENSG00000 | gene | 1,25857497 | 0,03140193 | 0,44510053 | PTPRU | 0,33179116 |
| ENSG00000 | gene | 1,14893884 | 0,03144225 | 0,44510053 | HS1BP3 | 0,200302 |
| ENSG00000 | gene | 1,39710678 | 0,03145048 | 0,44510053 | BCAP29 | 0,48244229 |
| ENSG00000 | gene | 0,72370044 | 0,03145164 | 0,44510053 | RN7SK | -0,4665355 |
| ENSG00000 | gene | 1,76430886 | 0,03156902 | 0,44510053 | AC000041.1 | 0,81910314 |
| ENSG00000 | gene | 0,91156475 | 0,03158473 | 0,44510053 | PYGL | -0,133583 |
| ENSG00000 | gene | 0,89979413 | 0,03161281 | 0,44510053 | ROCK2 | -0,1523331 |
| ENSG00000 | gene | 1,51419284 | 0,03168984 | 0,44510053 | NUP155 | 0,59854895 |
| ENSG00000 | gene | 1,31334972 | 0,03170233 | 0,44510053 | OSGIN2 | 0,39325113 |
| ENSG00000 | gene | 0,75363728 | 0,03172544 | 0,44510053 | CHCHD2P9 | -0,4080578 |
| ENSG00000 | gene | 0,62358877 | 0,0317428 | 0,44510053 | AC025370.1 | -0,6813331 |
| ENSG00000 | gene | 0,80525651 | 0,0317513 | 0,44510053 | ITPA | -0,3124797 |
| ENSG00000 | gene | 0,73898271 | 0,03176717 | 0,44510053 | PKIG | -0,4363875 |
| ENSG00000 | gene | 0,66444558 | 0,03209111 | 0,44802619 | SNORD19 | -0,589777 |
| ENSG00000 | gene | 1,14695925 | 0,03209526 | 0,44802619 | HUWE1 | 0,19781413 |

| | | | | | | |
|---|---|---|---|---|---|---|
| ENSG00000 | gene | 0,82843116 | 0,03216318 | 0,44802619 | TRAPPC2 | -0,2715463 |
| ENSG00000 | gene | 0,91868638 | 0,03220378 | 0,44802619 | RPL9 | -0,1223556 |
| ENSG00000 | gene | 1,44832189 | 0,03223642 | 0,44802619 | IDUA | 0,53438227 |
| ENSG00000 | gene | 1,25670968 | 0,03224633 | 0,44802619 | ZXDC | 0,3296514 |
| ENSG00000 | gene | 0,54740929 | 0,03225789 | 0,44802619 | SNORD36B | -0,8693082 |
| ENSG00000 | gene | 0,8094348 | 0,03243502 | 0,44992467 | CYTH3 | -0,3050132 |
| ENSG00000 | gene | 0,52417016 | 0,03251677 | 0,4502545 | AL157935.3 | -0,9318929 |
| ENSG00000 | gene | 0,49291283 | 0,03256177 | 0,4502545 | SNORA6 | -1,0205955 |
| ENSG00000 | gene | 0,90354414 | 0,03258021 | 0,4502545 | LYSMD3 | -0,146333 |
| ENSG00000 | gene | 0,78770058 | 0,0326552 | 0,45073084 | FTH1P15 | -0,3442808 |
| ENSG00000 | gene | 0,9423246 | 0,03270982 | 0,45092533 | TUBG1 | -0,085704 |
| ENSG00000 | gene | 0,70164597 | 0,03275928 | 0,45104825 | MFSD11 | -0,5111848 |
| ENSG00000 | gene | 1,08721575 | 0,03283156 | 0,45148473 | SLC39A6 | 0,12063826 |
| ENSG00000 | gene | 0,94005799 | 0,0329715 | 0,45284936 | ALYREF | -0,0891783 |
| ENSG00000 | gene | 1,20532317 | 0,03304202 | 0,45325832 | AP000331.1 | 0,26942001 |
| ENSG00000 | gene | 0,87208116 | 0,03315013 | 0,45418132 | AC084024.3 | -0,1974657 |
| ENSG00000 | gene | 1,35074301 | 0,03320994 | 0,45427647 | CEMIP | 0,43375322 |
| ENSG00000 | gene | 0,89895566 | 0,03326798 | 0,45427647 | DNAJB11 | -0,1536781 |
| ENSG00000 | gene | 0,58923154 | 0,03328755 | 0,45427647 | MIR4690 | -0,7630934 |
| ENSG00000 | gene | 0,90229116 | 0,03332041 | 0,45427647 | SRA1 | -0,148335 |
| ENSG00000 | gene | 1,49197159 | 0,03349023 | 0,45600896 | DUSP10 | 0,57722006 |
| ENSG00000 | gene | 0,82700794 | 0,03356039 | 0,45600896 | GFM1 | -0,2740269 |
| ENSG00000 | gene | 1,14319408 | 0,03357046 | 0,45600896 | AC036108.2 | 0,19307035 |
| ENSG00000 | gene | 0,80763899 | 0,03362843 | 0,45621095 | SDC4 | -0,3082175 |
| ENSG00000 | gene | 0,50915046 | 0,03369457 | 0,45621095 | ASS1 | -0,973836 |
| ENSG00000 | gene | 0,81152924 | 0,03374105 | 0,45621095 | SBDS | -0,301285 |
| ENSG00000 | gene | 1,96684564 | 0,03377821 | 0,45621095 | BRWD1 | 0,97588374 |
| ENSG00000 | gene | 1,2353432 | 0,03386919 | 0,45621095 | VLDLR-AS1 | 0,3049119 |
| ENSG00000 | gene | 0,87414456 | 0,0339178 | 0,45621095 | AP002383.2 | -0,1940562 |
| ENSG00000 | gene | 2,24020781 | 0,03392711 | 0,45621095 | CXCL1 | 1,16363257 |
| ENSG00000 | gene | 0,79616896 | 0,03394527 | 0,45621095 | S100A6 | -0,3288535 |
| ENSG00000 | gene | 0,88572331 | 0,0339544 | 0,45621095 | RCN2 | -0,175072 |
| ENSG00000 | gene | 1,04574073 | 0,03403112 | 0,45631835 | SMARCA5 | 0,0645252 |
| ENSG00000 | gene | 1,46916997 | 0,03404443 | 0,45631835 | AL138785.2 | 0,55500131 |
| ENSG00000 | gene | 0,86610238 | 0,03414075 | 0,45695987 | AC022167.1 | -0,2073905 |
| ENSG00000 | gene | 1,95338424 | 0,03426316 | 0,45695987 | HEATR5A | 0,96597576 |
| ENSG00000 | gene | 1,73039002 | 0,03427208 | 0,45695987 | RMND5B | 0,79109725 |
| ENSG00000 | gene | 0,78034077 | 0,0342944 | 0,45695987 | GADD45G | -0,3578238 |
| ENSG00000 | gene | 1,41757274 | 0,03437091 | 0,45695987 | CLDN12 | 0,50342277 |
| ENSG00000 | gene | 0,85472626 | 0,03437357 | 0,45695987 | CNOT4 | -0,2264656 |
| ENSG00000 | gene | 0,73062234 | 0,0343849 | 0,45695987 | PRAF2 | -0,4528022 |
| ENSG00000 | gene | 1,10531104 | 0,03442089 | 0,45695987 | MTX1P1 | 0,14445241 |
| ENSG00000 | gene | 0,8751643 | 0,03452387 | 0,45778075 | TIMM23 | -0,1923742 |
| ENSG00000 | gene | 0,92133529 | 0,03457709 | 0,45793394 | RPL23A | -0,1182018 |
| ENSG00000 | gene | 0,83863524 | 0,03461775 | 0,45793394 | RPLP1 | -0,2538846 |
| ENSG00000 | gene | 1,46223946 | 0,03468133 | 0,45803717 | ZNF35 | 0,54817959 |
| ENSG00000 | gene | 1,84267717 | 0,0347079 | 0,45803717 | DNAJC16 | 0,88180334 |
| ENSG00000 | gene | 0,93603709 | 0,03475037 | 0,45805438 | TSPAN9 | -0,0953624 |
| ENSG00000 | gene | 1,28755882 | 0,0348553 | 0,45889381 | ANKRD10-IT | 0,36463834 |
| ENSG00000 | gene | 1,50065018 | 0,03500184 | 0,46027828 | MIR570 | 0,5855877 |
| ENSG00000 | gene | 0,8306024 | 0,03507255 | 0,46066365 | AL110504.1 | -0,2677701 |
| ENSG00000 | gene | 0,82021614 | 0,03529592 | 0,4630508 | PCCB | -0,285924 |
| ENSG00000 | gene | 0,89845709 | 0,03540716 | 0,46336874 | NECTIN2 | -0,1544785 |
| ENSG00000 | gene | 0,92966082 | 0,03551673 | 0,46336874 | CCDC124 | -0,1052236 |
| ENSG00000 | gene | 0,84674347 | 0,03557016 | 0,46336874 | SNRPF | -0,2400031 |
| ENSG00000 | gene | 0,82657777 | 0,03561606 | 0,46336874 | CNN3 | -0,2747775 |
| ENSG00000 | gene | 1,51191964 | 0,03562138 | 0,46336874 | FMR1 | 0,59638146 |

| | | | | | |
|---|---|---|---|---|---|
| ENSG00000 gene | 0,62268318 | 0,03562623 | 0,46336874 | HLA-DMA | -0,6834298 |
| ENSG00000 gene | 1,21728812 | 0,03573584 | 0,46336874 | BOD1L1 | 0,28367068 |
| ENSG00000 gene | 0,70906329 | 0,03574531 | 0,46336874 | HMGB1 | -0,4960137 |
| ENSG00000 gene | 0,79564893 | 0,03585632 | 0,46336874 | ANXA11 | -0,3297961 |
| ENSG00000 gene | 1,21067469 | 0,03587058 | 0,46336874 | SATB2 | 0,27581126 |
| ENSG00000 gene | 0,78085433 | 0,03592626 | 0,46336874 | SELENOM | -0,3568747 |
| ENSG00000 gene | 0,7436519 | 0,03592787 | 0,46336874 | ATPAF2 | -0,4273006 |
| ENSG00000 gene | 1,12731446 | 0,03593449 | 0,46336874 | NFYA | 0,17289001 |
| ENSG00000 gene | 0,86203251 | 0,03593462 | 0,46336874 | STMN3 | -0,2141858 |
| ENSG00000 gene | 0,74145186 | 0,03594492 | 0,46336874 | MB21D2 | -0,4315751 |
| ENSG00000 gene | 0,92526538 | 0,03614692 | 0,46501481 | HNRNPC | -0,1120609 |
| ENSG00000 gene | 0,81527419 | 0,03615621 | 0,46501481 | FDX2 | -0,2946428 |
| ENSG00000 gene | 1,14364939 | 0,03623375 | 0,46545929 | GATAD1 | 0,19364484 |
| ENSG00000 gene | 1,23080487 | 0,03629707 | 0,46545929 | AVPI1 | 0,29960206 |
| ENSG00000 gene | 1,48698141 | 0,03632279 | 0,46545929 | GPAT3 | 0,57238661 |
| ENSG00000 gene | 0,72372351 | 0,03641606 | 0,46545929 | H4C15 | -0,4664895 |
| ENSG00000 gene | 0,95972532 | 0,03642655 | 0,46545929 | P3H1 | -0,0593065 |
| ENSG00000 gene | 0,84562973 | 0,0364418 | 0,46545929 | FAM76B | -0,241902 |
| ENSG00000 gene | 1,14308315 | 0,03650321 | 0,4657034 | PPP1R3C | 0,19293036 |
| ENSG00000 gene | 0,92002666 | 0,03654464 | 0,4657034 | RPL38 | -0,1202524 |
| ENSG00000 gene | 0,68150832 | 0,03668293 | 0,46693095 | PDCD1LG2 | -0,5531968 |
| ENSG00000 gene | 1,11699978 | 0,03682806 | 0,46712928 | AURKA | 0,1596289 |
| ENSG00000 gene | 0,89693378 | 0,03683564 | 0,46712928 | NET1 | -0,1569266 |
| ENSG00000 gene | 0,92593389 | 0,03688058 | 0,46712928 | NSMCE3 | -0,1110189 |
| ENSG00000 gene | 0,90015233 | 0,03688367 | 0,46712928 | NADK | -0,1517589 |
| ENSG00000 gene | 0,93037799 | 0,03690846 | 0,46712928 | RPS4X | -0,1041111 |
| ENSG00000 gene | 0,88478515 | 0,03695698 | 0,46721185 | SIRT2 | -0,1766009 |
| ENSG00000 gene | 0,89019487 | 0,03711332 | 0,46798668 | GLYR1 | -0,1678069 |
| ENSG00000 gene | 0,93958706 | 0,03723766 | 0,46798668 | AFAP1-AS1 | -0,0899013 |
| ENSG00000 gene | 0,78674016 | 0,03724225 | 0,46798668 | ETFDH | -0,3460409 |
| ENSG00000 gene | 0,9449766 | 0,03728617 | 0,46798668 | AFAP1 | -0,0816495 |
| ENSG00000 gene | 0,88238099 | 0,03728722 | 0,46798668 | ARPC1A | -0,1805264 |
| ENSG00000 gene | 1,09979584 | 0,03734091 | 0,46798668 | DDX50 | 0,13723573 |
| ENSG00000 gene | 0,64656096 | 0,03735124 | 0,46798668 | DGAT1 | -0,6291417 |
| ENSG00000 gene | 1,30629076 | 0,03736582 | 0,46798668 | HSD11B1 | 0,38547605 |
| ENSG00000 gene | 0,79462243 | 0,03742989 | 0,46798668 | AC006441.3 | -0,3316586 |
| ENSG00000 gene | 0,92271317 | 0,03748932 | 0,46798668 | GTF2F2 | -0,1160459 |
| ENSG00000 gene | 0,85984305 | 0,03751946 | 0,46798668 | REX1BD | -0,2178548 |
| ENSG00000 gene | 1,32747388 | 0,03752307 | 0,46798668 | SOD2 | 0,40868348 |
| ENSG00000 gene | 1,40106255 | 0,0376131 | 0,46809087 | PCDHGB5 | 0,48652137 |
| ENSG00000 gene | 0,79759926 | 0,0376596 | 0,46809087 | ROMO1 | -0,326264 |
| ENSG00000 gene | 1,25920054 | 0,03770915 | 0,46809087 | KMT2D | 0,33250807 |
| ENSG00000 gene | 1,20248891 | 0,03772688 | 0,46809087 | CD58 | 0,26602359 |
| ENSG00000 gene | 0,86616987 | 0,0377418 | 0,46809087 | SLC25A11 | -0,2072781 |
| ENSG00000 gene | 1,13108152 | 0,03779472 | 0,46822528 | PLIN2 | 0,17770291 |
| ENSG00000 gene | 1,3590369 | 0,03785957 | 0,46850693 | DTX2 | 0,44258462 |
| ENSG00000 gene | 1,15000071 | 0,03794502 | 0,46862262 | ASAH1 | 0,20163476 |
| ENSG00000 gene | 0,87061808 | 0,03796338 | 0,46862262 | ERH | -0,1998881 |
| ENSG00000 gene | 1,16752814 | 0,03801814 | 0,46862262 | TWF1P1 | 0,22345732 |
| ENSG00000 gene | 1,16833443 | 0,03807537 | 0,46862262 | CEMIP2 | 0,2244533 |
| ENSG00000 gene | 3,50424492 | 0,03807954 | 0,46862262 | AREG | 1,80910361 |
| ENSG00000 gene | 0,86523777 | 0,03824687 | 0,47016175 | NUCB2 | -0,2088315 |
| ENSG00000 gene | 0,8616382 | 0,03829837 | 0,47027525 | HSD17B10 | -0,2148459 |
| ENSG00000 gene | 0,88907392 | 0,03841333 | 0,47105827 | PEAR1 | -0,1696247 |
| ENSG00000 gene | 0,61304191 | 0,03844682 | 0,47105827 | L3MBTL3 | -0,7059424 |
| ENSG00000 gene | 0,53303687 | 0,03854461 | 0,47117511 | BBS1 | -0,9076928 |
| ENSG00000 gene | 0,94159313 | 0,03854633 | 0,47117511 | RPL5 | -0,0868243 |

| | | | | | |
|---|---|---|---|---|---|
| ENSG00000 gene | 1,22125459 | 0,03858342 | 0,47117511 | TOR1A | 0,28836398 |
| ENSG00000 gene | 1,52689147 | 0,03871089 | 0,47221342 | MIR7161 | 0,61059752 |
| ENSG00000 gene | 0,64443848 | 0,03881117 | 0,47291809 | HNRNPKP4 | -0,6338855 |
| ENSG00000 gene | 0,82529806 | 0,0389167 | 0,47364307 | SNRPG | -0,2770129 |
| ENSG00000 gene | 0,93287851 | 0,03895581 | 0,47364307 | SNRNP35 | -0,1002389 |
| ENSG00000 gene | 0,92263217 | 0,03908441 | 0,47458635 | RPL3 | -0,1161725 |
| ENSG00000 gene | 0,88002781 | 0,03913198 | 0,47458635 | ISCU | -0,184379 |
| ENSG00000 gene | 1,38476034 | 0,03916499 | 0,47458635 | ACSL4 | 0,46963631 |
| ENSG00000 gene | 0,83434419 | 0,03925773 | 0,47458635 | FASTK | -0,2612854 |
| ENSG00000 gene | 1,10015058 | 0,03928138 | 0,47458635 | ACLY | 0,137701 |
| ENSG00000 gene | 0,89740668 | 0,03931412 | 0,47458635 | CHPF2 | -0,1561662 |
| ENSG00000 gene | 0,87647429 | 0,03934548 | 0,47458635 | MRPL4 | -0,1902163 |
| ENSG00000 gene | 0,48797431 | 0,0393759 | 0,47458635 | AC008750.8 | -1,0351229 |
| ENSG00000 gene | 1,05234195 | 0,03942866 | 0,47458635 | SLC7A11 | 0,07360357 |
| ENSG00000 gene | 0,90759071 | 0,03946836 | 0,47458635 | ABT1 | -0,1398863 |
| ENSG00000 gene | 0,76145812 | 0,03951994 | 0,47458635 | WDR77 | -0,3931634 |
| ENSG00000 gene | 0,85371356 | 0,03955158 | 0,47458635 | BRD9 | -0,228176 |
| ENSG00000 gene | 0,90314419 | 0,03969067 | 0,47458635 | SUMO2 | -0,1469718 |
| ENSG00000 gene | 0,91630633 | 0,03970399 | 0,47458635 | PSMG2 | -0,1260981 |
| ENSG00000 gene | 0,92103492 | 0,03977949 | 0,47458635 | GAR1 | -0,1186722 |
| ENSG00000 gene | 1,12233204 | 0,0398129 | 0,47458635 | KIF3B | 0,16649956 |
| ENSG00000 gene | 0,87780493 | 0,03982056 | 0,47458635 | TLE4 | -0,1880277 |
| ENSG00000 gene | 1,0504721 | 0,03983303 | 0,47458635 | SERINC1 | 0,07103785 |
| ENSG00000 gene | 1,52307309 | 0,03993912 | 0,47458635 | TM4SF1 | 0,60698518 |
| ENSG00000 gene | 1,04274502 | 0,03998679 | 0,47458635 | SND1 | 0,06038642 |
| ENSG00000 gene | 1,22897304 | 0,04000633 | 0,47458635 | MID1 | 0,29745327 |
| ENSG00000 gene | 0,89657918 | 0,04003836 | 0,47458635 | PI4K2B | -0,1574971 |
| ENSG00000 gene | 0,84993555 | 0,04007841 | 0,47458635 | ARPC2 | -0,2345746 |
| ENSG00000 gene | 1,32547251 | 0,0401211 | 0,47458635 | IKBKB | 0,40650675 |
| ENSG00000 gene | 1,14582878 | 0,04013142 | 0,47458635 | CTPS1 | 0,19639148 |
| ENSG00000 gene | 1,18934201 | 0,04014254 | 0,47458635 | SIN3B | 0,25016364 |
| ENSG00000 gene | 0,93242671 | 0,04026909 | 0,47556715 | JAGN1 | -0,1009378 |
| ENSG00000 gene | 1,25057111 | 0,04031099 | 0,47556715 | KCNMA1-AS | 0,3225871 |
| ENSG00000 gene | 1,1671361 | 0,04039165 | 0,47599972 | UBAP2 | 0,22297281 |
| ENSG00000 gene | 0,64713133 | 0,04043323 | 0,47599972 | H2BC4 | -0,6278696 |
| ENSG00000 gene | 0,91171837 | 0,04053232 | 0,47650859 | RPS27 | -0,1333398 |
| ENSG00000 gene | 0,92239265 | 0,04058823 | 0,47650859 | PUF60 | -0,1165471 |
| ENSG00000 gene | 0,9251102 | 0,04060496 | 0,47650859 | TST | -0,1123029 |
| ENSG00000 gene | 1,17275392 | 0,04072103 | 0,47736715 | FDFT1 | 0,22990032 |
| ENSG00000 gene | 0,88153366 | 0,04079748 | 0,47775993 | MXD4 | -0,1819124 |
| ENSG00000 gene | 0,61094363 | 0,04089021 | 0,47828904 | CU633904.2 | -0,7108888 |
| ENSG00000 gene | 1,56981757 | 0,04092864 | 0,47828904 | PACRGL | 0,65059691 |
| ENSG00000 gene | 1,52858916 | 0,04100276 | 0,47865242 | RNMT | 0,61220071 |
| ENSG00000 gene | 0,87778266 | 0,04123112 | 0,48058264 | ACTR1A | -0,1880643 |
| ENSG00000 gene | 1,34545129 | 0,04125451 | 0,48058264 | ZEB2 | 0,42809016 |
| ENSG00000 gene | 0,90774116 | 0,04147519 | 0,48247709 | ZNF512B | -0,1396471 |
| ENSG00000 gene | 0,82791648 | 0,04156191 | 0,48247709 | USP53 | -0,2724429 |
| ENSG00000 gene | 0,9212983 | 0,04160521 | 0,48247709 | USP16 | -0,1182597 |
| ENSG00000 gene | 0,79221965 | 0,0416523 | 0,48247709 | PSMG4 | -0,3360276 |
| ENSG00000 gene | 0,9254075 | 0,04167712 | 0,48247709 | TMEM183A | -0,1118393 |
| ENSG00000 gene | 0,88871067 | 0,04167735 | 0,48247709 | FADD | -0,1702143 |
| ENSG00000 gene | 0,68418265 | 0,04191794 | 0,48396459 | COPS7B | -0,5475466 |
| ENSG00000 gene | 1,70501367 | 0,04192461 | 0,48396459 | HAUS8 | 0,76978331 |
| ENSG00000 gene | 2,06046204 | 0,0419603 | 0,48396459 | MMP10 | 1,04296789 |
| ENSG00000 gene | 1,41933745 | 0,04197985 | 0,48396459 | ABHD6 | 0,50521764 |
| ENSG00000 gene | 1,07703147 | 0,04222486 | 0,48628528 | CCNB1 | 0,1070604 |
| ENSG00000 gene | 0,80009783 | 0,0422951 | 0,4864026 | CDK2AP2 | -0,3217517 |

| | | | | | |
|---|---|---|---|---|---|
| ENSG00000 gene | 1,1139923 | 0,04232249 | 0,4864026 | DDX60 | 0,15573926 |
| ENSG00000 gene | 0,66367956 | 0,0424131 | 0,48655053 | C17orf80 | -0,5914413 |
| ENSG00000 gene | 0,89883207 | 0,04245219 | 0,48655053 | STOML2 | -0,1538765 |
| ENSG00000 gene | 0,72886065 | 0,04246657 | 0,48655053 | DDT | -0,4562851 |
| ENSG00000 gene | 1,55206815 | 0,04264588 | 0,4871231 | CRLF1 | 0,6341919 |
| ENSG00000 gene | 0,90288356 | 0,04264757 | 0,4871231 | RPL27 | -0,1473881 |
| ENSG00000 gene | 0,83409993 | 0,04271067 | 0,4871231 | ANXA2P2 | -0,2617079 |
| ENSG00000 gene | 0,53242755 | 0,04274534 | 0,4871231 | APOLD1 | -0,9093429 |
| ENSG00000 gene | 0,7761447 | 0,04277225 | 0,4871231 | MGST1 | -0,3656025 |
| ENSG00000 gene | 0,84186411 | 0,04277926 | 0,4871231 | COTL1 | -0,2483407 |
| ENSG00000 gene | 0,63328415 | 0,04294583 | 0,48787441 | ARHGAP27I | -0,6590751 |
| ENSG00000 gene | 1,08458179 | 0,04295656 | 0,48787441 | CTNNB1 | 0,11713885 |
| ENSG00000 gene | 0,80063889 | 0,0429768 | 0,48787441 | RSRC1 | -0,3207764 |
| ENSG00000 gene | 1,19510114 | 0,04324118 | 0,48992921 | TRRAP | 0,25713271 |
| ENSG00000 gene | 0,72276041 | 0,04324589 | 0,48992921 | CALM2P2 | -0,4684106 |
| ENSG00000 gene | 0,84628633 | 0,04331676 | 0,49023287 | ANXA2 | -0,2407822 |
| ENSG00000 gene | 0,92791479 | 0,04337229 | 0,49036255 | RPL23AP42 | -0,1079358 |
| ENSG00000 gene | 1,69596607 | 0,04344152 | 0,49064664 | LRRC8E | 0,76210731 |
| ENSG00000 gene | 1,0847064 | 0,04358084 | 0,49079601 | PTAR1 | 0,11730459 |
| ENSG00000 gene | 0,90284598 | 0,0436241 | 0,49079601 | TIMM17A | -0,1474482 |
| ENSG00000 gene | 0,88647818 | 0,0436583 | 0,49079601 | NKIRAS1 | -0,173843 |
| ENSG00000 gene | 0,69637177 | 0,04366068 | 0,49079601 | ALDH4A1 | -0,5220704 |
| ENSG00000 gene | 0,57711061 | 0,04367966 | 0,49079601 | AC093525.2 | -0,7930803 |
| ENSG00000 gene | 1,58278312 | 0,04371945 | 0,49079601 | SNORD12B | 0,66246359 |
| ENSG00000 gene | 0,91289145 | 0,0437812 | 0,49099379 | RNF181 | -0,1314848 |
| ENSG00000 gene | 1,03447508 | 0,04396892 | 0,49245902 | CKAP5 | 0,04889889 |
| ENSG00000 gene | 0,92812819 | 0,04404331 | 0,49245902 | JSRP1 | -0,107604 |
| ENSG00000 gene | 0,90919203 | 0,04406091 | 0,49245902 | RPL6 | -0,1373431 |
| ENSG00000 gene | 1,24493399 | 0,04408892 | 0,49245902 | SH3PXD2A | 0,31606925 |
| ENSG00000 gene | 1,36053066 | 0,04430663 | 0,49352742 | AKR1C1 | 0,44416947 |
| ENSG00000 gene | 0,65207402 | 0,04440586 | 0,49352742 | GMPPB | -0,6168923 |
| ENSG00000 gene | 1,30374649 | 0,04441392 | 0,49352742 | FOXO3 | 0,38266337 |
| ENSG00000 gene | 1,17428025 | 0,04444155 | 0,49352742 | NUP214 | 0,23177676 |
| ENSG00000 gene | 0,86525785 | 0,04444359 | 0,49352742 | STK39 | -0,208798 |
| ENSG00000 gene | 0,21099686 | 0,04445074 | 0,49352742 | HCG20 | -2,2447066 |
| ENSG00000 gene | 0,87454421 | 0,04454401 | 0,4937598 | RABEP2 | -0,1933968 |
| ENSG00000 gene | 1,23535964 | 0,04456043 | 0,4937598 | LTBP2 | 0,3049311 |
| ENSG00000 gene | 0,80560772 | 0,04489921 | 0,4970186 | RGS10 | -0,3118506 |
| ENSG00000 gene | 0,78821798 | 0,04499406 | 0,49757351 | ACTR3 | -0,3433334 |
| ENSG00000 gene | 0,94360101 | 0,04519764 | 0,49932843 | KLHDC3 | -0,0837511 |
| ENSG00000 gene | 1,21450629 | 0,04527929 | 0,49963108 | NF1 | 0,28036997 |
| ENSG00000 gene | 1,05621959 | 0,04536239 | 0,49963108 | RANBP2 | 0,0789098 |
| ENSG00000 gene | 0,86763319 | 0,04536972 | 0,49963108 | NDUFB2 | -0,2048429 |
| ENSG00000 gene | 0,88747518 | 0,04540557 | 0,49963108 | LTO1 | -0,1722213 |
| ENSG00000 gene | 0,94970807 | 0,04544959 | 0,49963108 | UBA52 | -0,074444 |
| ENSG00000 gene | 1,23915788 | 0,04557936 | 0,50056301 | NDUFB9 | 0,30936001 |
| ENSG00000 gene | 1,08435236 | 0,04569785 | 0,5005865 | NCAPD2 | 0,11683363 |
| ENSG00000 gene | 0,76568353 | 0,04570663 | 0,5005865 | C12orf65 | -0,3851799 |
| ENSG00000 gene | 1,35798046 | 0,04571648 | 0,5005865 | COL7A1 | 0,44146272 |
| ENSG00000 gene | 0,85435872 | 0,04580717 | 0,50108627 | MTG1 | -0,2270861 |
| ENSG00000 gene | 0,70543361 | 0,04585275 | 0,50109222 | THSD1 | -0,5034178 |
| ENSG00000 gene | 0,88735072 | 0,04599124 | 0,50168709 | EMC6 | -0,1724237 |
| ENSG00000 gene | 0,58347767 | 0,04599738 | 0,50168709 | LINC00680 | -0,7772507 |
| ENSG00000 gene | 0,93673266 | 0,04607545 | 0,50200395 | STXBP3 | -0,0942907 |
| ENSG00000 gene | 0,91741338 | 0,04615112 | 0,50200395 | RPL37 | -0,1243562 |
| ENSG00000 gene | 1,1919995 | 0,0461618 | 0,50200395 | ABCA1 | 0,25338363 |
| ENSG00000 gene | 0,80441661 | 0,04630478 | 0,50269243 | CMTM3 | -0,3139852 |

| | | | | | |
|---|---|---|---|---|---|
| ENSG00000 gene | 1,08197335 | 0,04637665 | 0,50269243 | USP19 | 0,11366497 |
| ENSG00000 gene | 0,81817894 | 0,04646177 | 0,50269243 | SSR2 | -0,2895117 |
| ENSG00000 gene | 1,18682695 | 0,04646632 | 0,50269243 | HMBOX1 | 0,24710959 |
| ENSG00000 gene | 0,74845692 | 0,04646714 | 0,50269243 | SNORA4 | -0,4180088 |
| ENSG00000 gene | 0,74928032 | 0,04649623 | 0,50269243 | RPL39 | -0,4164225 |
| ENSG00000 gene | 0,74462989 | 0,04657936 | 0,50310235 | LHFPL6 | -0,4254046 |
| ENSG00000 gene | 0,42893654 | 0,04677892 | 0,50476773 | AC008878.1 | -1,2211639 |
| ENSG00000 gene | 1,12778871 | 0,04686444 | 0,50490085 | PALB2 | 0,17349681 |
| ENSG00000 gene | 0,90658426 | 0,04688203 | 0,50490085 | EIF5A | -0,141487 |
| ENSG00000 gene | 0,91169302 | 0,04697206 | 0,50513652 | HDAC5 | -0,13338 |
| ENSG00000 gene | 0,86738452 | 0,04702817 | 0,50513652 | C1orf122 | -0,2052564 |
| ENSG00000 gene | 0,94221187 | 0,04704013 | 0,50513652 | HEXIM1 | -0,0858766 |
| ENSG00000 gene | 0,83258731 | 0,04708642 | 0,50514599 | PRKCI | -0,2643265 |
| ENSG00000 gene | 1,14030478 | 0,04714799 | 0,5053193 | ZSCAN29 | 0,18941948 |
| ENSG00000 gene | 0,82975462 | 0,04721844 | 0,50558725 | GPR89A | -0,2692433 |
| ENSG00000 gene | 1,50809077 | 0,04729675 | 0,50593881 | AL139393.3 | 0,59272326 |
| ENSG00000 gene | 0,56884256 | 0,04735368 | 0,50601068 | HHEX | -0,8138987 |
| ENSG00000 gene | 1,22005096 | 0,04739444 | 0,50601068 | FAM193A | 0,28694141 |
| ENSG00000 gene | 1,15742115 | 0,04747832 | 0,50642025 | QPCTL | 0,21091392 |
| ENSG00000 gene | 1,16903858 | 0,04758894 | 0,50688514 | INCENP | 0,22532254 |
| ENSG00000 gene | 0,81585917 | 0,04761303 | 0,50688514 | RPL15 | -0,293608 |
| ENSG00000 gene | 1,39289362 | 0,04769012 | 0,50703436 | ZFYVE16 | 0,47808508 |
| ENSG00000 gene | 1,14720399 | 0,04771917 | 0,50703436 | IRF3 | 0,19812195 |
| ENSG00000 gene | 0,70646402 | 0,04778879 | 0,50703436 | ARPC4 | -0,501312 |
| ENSG00000 gene | 0,66010848 | 0,04783451 | 0,50703436 | CDKN1A | -0,599225 |
| ENSG00000 gene | 0,74002116 | 0,04785493 | 0,50703436 | SH2D5 | -0,4343616 |
| ENSG00000 gene | 1,64149476 | 0,04808692 | 0,50878252 | PDE10A | 0,71501014 |
| ENSG00000 gene | 0,98380562 | 0,04811139 | 0,50878252 | MT-ATP6 | -0,0235548 |
| ENSG00000 gene | 0,96574052 | 0,04837711 | 0,51033017 | EIF2S2 | -0,0502925 |
| ENSG00000 gene | 0,60939625 | 0,04840233 | 0,51033017 | SNORD26 | -0,7145475 |
| ENSG00000 gene | 1,05881887 | 0,04840741 | 0,51033017 | CCAR1 | 0,08245581 |
| ENSG00000 gene | 0,76937246 | 0,04844968 | 0,51033017 | IER2 | -0,3782459 |
| ENSG00000 gene | 1,34010927 | 0,0484871 | 0,51033017 | ZNF451-AS1 | 0,42235064 |
| ENSG00000 gene | 0,83389022 | 0,04861177 | 0,51115875 | AC112777.1 | -0,2620706 |
| ENSG00000 gene | 1,20437499 | 0,04874228 | 0,5120162 | RNF25 | 0,26828465 |
| ENSG00000 gene | 0,93821755 | 0,04880673 | 0,5120162 | RPL18AP3 | -0,0920056 |
| ENSG00000 gene | 0,65866596 | 0,04885036 | 0,5120162 | FGF5 | -0,6023811 |
| ENSG00000 gene | 1,1147475 | 0,04892414 | 0,5120162 | TMEM87A | 0,15671697 |
| ENSG00000 gene | 1,08551019 | 0,04893868 | 0,5120162 | HCFC1 | 0,11837327 |
| ENSG00000 gene | 0,85848858 | 0,04896946 | 0,5120162 | APBA3 | -0,2201291 |
| ENSG00000 gene | 1,34487744 | 0,04913304 | 0,51312768 | SLC2A13 | 0,4274747 |
| ENSG00000 gene | 0,94643331 | 0,04916801 | 0,51312768 | HMGN4 | -0,0794272 |
| ENSG00000 gene | 0,92319309 | 0,04946714 | 0,51576564 | C5orf15 | -0,1152957 |
| ENSG00000 gene | 1,30508652 | 0,04957075 | 0,51620413 | MT-TQ | 0,38414545 |
| ENSG00000 gene | 1,1498168 | 0,049602 | 0,51620413 | AL353689.2 | 0,20140402 |
| ENSG00000 gene | 0,92819856 | 0,04971126 | 0,51685776 | BTF3 | -0,1074946 |
| ENSG00000 gene | 2,04366248 | 0,04988942 | 0,51775082 | MEDAG | 1,03115695 |
| ENSG00000 gene | 2,14855804 | 0,04990515 | 0,51775082 | SNORD14C | 1,10336875 |
| ENSG00000 gene | 1,19745454 | 0,04993678 | 0,51775082 | METTL9 | 0,25997089 |

**PBMC_DEGs**

| id | feature | fc | pval | qval | gene_name | de |
|---|---|---|---|---|---|---|
| ENSG000001( | gene | 0,040277187 | 5,29794E-05 | 0,129441475 | CLC | -4,63389324 |
| ENSG000001: | gene | 0,044524105 | 0,001152044 | 0,224332495 | AHNAK | -4,48926957 |
| ENSG000001( | gene | 0,070881288 | 0,039341104 | 0,283183815 | CXCL8 | -3,81845137 |
| ENSG000002‹ | gene | 0,113876551 | 0,029200987 | 0,270832371 | DEFA1B | -3,13445739 |
| ENSG000001: | gene | 0,120958605 | 0,002218514 | 0,231249696 | GSTM1 | -3,04741469 |
| ENSG000001‹ | gene | 0,173317928 | 0,027958327 | 0,268594209 | HDC | -2,5285072 |
| ENSG000002: | gene | 0,1796499 | 0,035288247 | 0,2774496 | DEFA3 | -2,47673996 |
| ENSG000001' | gene | 0,194677729 | 0,010607073 | 0,24500668 | KLRB1 | -2,36084024 |
| ENSG000001: | gene | 0,200676557 | 0,03467302 | 0,27606048 | FGFBP2 | -2,317056 |
| ENSG000002: | gene | 0,202686275 | 0,001232698 | 0,224332495 | AC104809.2 | -2,30267969 |
| ENSG000002: | gene | 0,203135979 | 0,039192388 | 0,283069015 | TRBV29-1 | -2,29948231 |
| ENSG000000: | gene | 0,206035646 | 0,000157757 | 0,143227097 | IKZF2 | -2,27903413 |
| ENSG000001ε | gene | 0,208011005 | 0,006801118 | 0,239194577 | MYBL1 | -2,26526823 |
| ENSG000001( | gene | 0,210414893 | 0,013214473 | 0,252112392 | CPA3 | -2,24869127 |
| ENSG000000( | gene | 0,216406482 | 0,012641407 | 0,249534675 | CACNA2D2 | -2,20818438 |
| ENSG000002' | gene | 0,218479835 | 0,013566448 | 0,252327435 | TRDC | -2,19442797 |
| ENSG000001( | gene | 0,222174553 | 0,023572108 | 0,263452051 | GATA3 | -2,17023451 |
| ENSG000001( | gene | 0,226329232 | 0,012466222 | 0,249534675 | IER2 | -2,14350516 |
| ENSG000002: | gene | 0,229396026 | 0,015115637 | 0,252327435 | AC022413.1 | -2,1240877 |
| ENSG000001ε | gene | 0,229593248 | 0,013703095 | 0,252327435 | CRIP2 | -2,12284788 |
| ENSG000000( | gene | 0,23157367 | 0,00717918 | 0,239194577 | PDZD4 | -2,11045686 |
| ENSG000001: | gene | 0,238835608 | 0,001367553 | 0,224332495 | KLRG1 | -2,06591015 |
| ENSG000000( | gene | 0,241830504 | 0,022878276 | 0,260623396 | MATK | -2,04793186 |
| ENSG000001; | gene | 0,246100268 | 0,001920181 | 0,2305768 | ZNF266 | -2,02268187 |
| ENSG000001' | gene | 0,247153634 | 0,0068228 | 0,239194577 | IFNG | -2,01651997 |
| ENSG000001; | gene | 0,247504571 | 0,005023837 | 0,239194577 | SIGLEC17P | -2,01447292 |
| ENSG000002; | gene | 0,256555826 | 0,02669163 | 0,267343734 | AC008033.3 | -1,96265531 |
| ENSG000001! | gene | 0,26134565 | 0,011354789 | 0,249534675 | AUTS2 | -1,93596895 |
| ENSG000002! | gene | 0,26299726 | 0,006548935 | 0,239194577 | BTG1P1 | -1,92688032 |
| ENSG000002: | gene | 0,263145034 | 0,023954294 | 0,26347532 | AL359962.1 | -1,92606993 |
| ENSG000002; | gene | 0,264395215 | 0,023800506 | 0,263452051 | CCL4 | -1,91923203 |
| ENSG000001: | gene | 0,266434876 | 0,002012245 | 0,2305768 | CDKN1C | -1,90814515 |
| ENSG000000; | gene | 0,268929591 | 0,031231864 | 0,271804321 | ITM2A | -1,89469959 |
| ENSG000000; | gene | 0,270412766 | 0,007522833 | 0,239194577 | CST7 | -1,88676483 |
| ENSG000002( | gene | 0,272218025 | 0,046627975 | 0,294694579 | SNORD36B | -1,8771655 |
| ENSG000002! | gene | 0,272643887 | 0,01901971 | 0,254046692 | LINC02384 | -1,87491028 |
| ENSG000000( | gene | 0,272680759 | 0,011068355 | 0,249392265 | TGFBR3 | -1,87471519 |
| ENSG000001( | gene | 0,27404852 | 0,012400295 | 0,249534675 | PTGDS | -1,86749675 |
| ENSG000002' | gene | 0,275240743 | 0,034143112 | 0,275250624 | TRAV14DV4 | -1,86123405 |
| ENSG000002! | gene | 0,277763018 | 0,024600055 | 0,264837204 | KRT17P8 | -1,84807356 |
| ENSG000001( | gene | 0,279169483 | 0,007362699 | 0,239194577 | SLC1A7 | -1,84078685 |
| ENSG000000( | gene | 0,282827361 | 0,035525511 | 0,277679087 | TKTL1 | -1,8220064 |
| ENSG000002; | gene | 0,285070052 | 0,032230906 | 0,272856143 | AC044849.1 | -1,81061161 |
| ENSG000001ε | gene | 0,286678733 | 0,021980802 | 0,25813016 | COL18A1 | -1,80249321 |
| ENSG000001ε | gene | 0,288417113 | 0,005603419 | 0,239194577 | FCRL6 | -1,79377133 |
| ENSG000001ε | gene | 0,291872069 | 0,023396649 | 0,263452051 | ARL4C | -1,77659194 |
| ENSG000002! | gene | 0,292113703 | 0,012820466 | 0,249534675 | AC008750.2 | -1,77539806 |
| ENSG000002ε | gene | 0,292362802 | 0,0238751 | 0,263452051 | AL499604.2 | -1,77416833 |
| ENSG000002( | gene | 0,295417683 | 0,035200702 | 0,277446768 | AL451085.1 | -1,75917191 |
| ENSG000002ε | gene | 0,296502401 | 0,009813431 | 0,242573226 | SNORD98 | -1,75388431 |
| ENSG000000: | gene | 0,29664437 | 0,022570631 | 0,260020801 | RUNX3 | -1,75319369 |
| ENSG000001‹ | gene | 0,296892934 | 0,026675084 | 0,267343734 | COL6A2 | -1,75198534 |
| ENSG000001! | gene | 0,297749436 | 0,029600974 | 0,271738191 | SPON2 | -1,74782932 |
| ENSG000002; | gene | 0,298364868 | 0,03388793 | 0,275250624 | CCL5 | -1,74485043 |
| ENSG000001! | gene | 0,299017102 | 0,027494549 | 0,267488561 | DUSP2 | -1,7417001 |

| | | | | | | |
|---|---|---|---|---|---|---|
| ENSG0000019 | gene | 0,299345235 | 0,010609675 | 0,24500668 | SH2D1B | -1,74011779 |
| ENSG0000011 | gene | 0,299874636 | 0,042101021 | 0,288019516 | GNLY | -1,7375686 |
| ENSG0000010 | gene | 0,302249836 | 0,048705192 | 0,298041903 | NKG7 | -1,72618654 |
| ENSG0000018 | gene | 0,302344735 | 0,017090057 | 0,252327435 | S1PR5 | -1,72573364 |
| ENSG0000009 | gene | 0,303123887 | 0,013759102 | 0,252327435 | LTBP4 | -1,72202055 |
| ENSG0000014 | gene | 0,303355687 | 0,027115787 | 0,267343734 | SOX13 | -1,72091774 |
| ENSG0000023 | gene | 0,303377068 | 0,017229337 | 0,252327435 | RPL10P19 | -1,72081606 |
| ENSG0000019 | gene | 0,303495956 | 0,045204971 | 0,2922856 | TPM2 | -1,7202508 |
| ENSG0000021 | gene | 0,304071506 | 0,043837001 | 0,292071193 | TRGC1 | -1,71751747 |
| ENSG0000016 | gene | 0,30441536 | 0,016972587 | 0,252327435 | NSG1 | -1,71588694 |
| ENSG0000014 | gene | 0,305754742 | 0,009069357 | 0,242573226 | MRPL49 | -1,70955322 |
| ENSG0000027 | gene | 0,308169536 | 0,008211203 | 0,242573226 | AL359962.2 | -1,69820384 |
| ENSG0000017 | gene | 0,311803724 | 0,004107887 | 0,239194577 | AC020659.1 | -1,68128994 |
| ENSG0000014 | gene | 0,313249344 | 0,031131672 | 0,271804321 | PGGHG | -1,67461661 |
| ENSG0000010 | gene | 0,313717356 | 0,037480826 | 0,281299108 | GZMB | -1,67246275 |
| ENSG0000023 | gene | 0,314477609 | 0,031560755 | 0,272282875 | LINC00402 | -1,6689708 |
| ENSG0000017 | gene | 0,320630166 | 0,045926677 | 0,292991357 | RARG | -1,64101793 |
| ENSG0000015 | gene | 0,320666348 | 0,008714753 | 0,242573226 | GOLGA7B | -1,64085514 |
| ENSG0000010 | gene | 0,32157889 | 0,027643472 | 0,267527236 | IL2RB | -1,63675539 |
| ENSG0000019 | gene | 0,323961929 | 0,010052755 | 0,242573226 | CD247 | -1,62610381 |
| ENSG0000013 | gene | 0,325270818 | 0,034547213 | 0,27606048 | PRR5L | -1,6202867 |
| ENSG0000020 | gene | 0,325582643 | 0,03844449 | 0,28223387 | SNORA2B | -1,6189043 |
| ENSG0000021 | gene | 0,326459302 | 0,026571655 | 0,267343734 | KLRK1 | -1,61502494 |
| ENSG0000017 | gene | 0,326652395 | 0,022676481 | 0,260623396 | CARNS1 | -1,61417188 |
| ENSG0000016 | gene | 0,329013009 | 0,009623113 | 0,242573226 | KIR3DL1 | -1,60378347 |
| ENSG0000015 | gene | 0,329625186 | 0,01227591 | 0,249534675 | CD226 | -1,60110161 |
| ENSG0000022 | gene | 0,329926431 | 0,01835628 | 0,252510181 | KIR2DS4 | -1,59978373 |
| ENSG0000018 | gene | 0,330343637 | 0,049052714 | 0,299174578 | BCL9L | -1,59796054 |
| ENSG0000016 | gene | 0,332086162 | 0,010292835 | 0,244080246 | PTGDR | -1,59037049 |
| ENSG0000024 | gene | 0,333818387 | 0,013972192 | 0,252327435 | LINC002481 | -1,58286467 |
| ENSG0000015 | gene | 0,335599455 | 0,002577965 | 0,237725877 | ZFYVE28 | -1,57518772 |
| ENSG0000012 | gene | 0,337203778 | 0,02163342 | 0,2564603 | PLEKHG3 | -1,56830739 |
| ENSG0000025 | gene | 0,339982328 | 0,033899537 | 0,275250624 | AC008750.1 | -1,55646834 |
| ENSG0000028 | gene | 0,340247784 | 0,023873018 | 0,263452051 | AL645933.5 | -1,55534233 |
| ENSG0000018 | gene | 0,340712622 | 0,009169564 | 0,242573226 | PRSS57 | -1,5533727 |
| ENSG0000013 | gene | 0,341791212 | 0,007335509 | 0,239194577 | LGR6 | -1,54881279 |
| ENSG0000010 | gene | 0,346870299 | 0,048996613 | 0,29910791 | NFATC2 | -1,52753178 |
| ENSG0000013 | gene | 0,347641096 | 0,031957959 | 0,272282875 | STAT4 | -1,52432946 |
| ENSG0000027 | gene | 0,349085086 | 0,004759987 | 0,239194577 | LINC00869 | -1,51834938 |
| ENSG0000012 | gene | 0,349208681 | 0,014736994 | 0,252327435 | KLF2 | -1,51783867 |
| ENSG0000016 | gene | 0,349345039 | 0,023403698 | 0,263452051 | IL7R | -1,51727544 |
| ENSG0000007 | gene | 0,354608054 | 0,015656027 | 0,252327435 | LLGL2 | -1,49570279 |
| ENSG0000002 | gene | 0,355380019 | 0,038887304 | 0,282708484 | SH2D2A | -1,49256553 |
| ENSG0000016 | gene | 0,35607888 | 0,009258877 | 0,242573226 | CAMP | -1,48973123 |
| ENSG0000012 | gene | 0,357078149 | 0,001344338 | 0,224332495 | KIR2DL1 | -1,48568824 |
| ENSG0000021 | gene | 0,358090964 | 0,029706678 | 0,271738191 | MT-TH | -1,48160198 |
| ENSG0000014 | gene | 0,358532307 | 0,0262236 | 0,266237968 | PMAIP1 | -1,47982497 |
| ENSG0000009 | gene | 0,359347862 | 0,028069917 | 0,268594209 | HIVEP1 | -1,47654699 |
| ENSG0000019 | gene | 0,360072937 | 0,031921774 | 0,272282875 | LYPD2 | -1,47363892 |
| ENSG0000013 | gene | 0,361125527 | 0,014968182 | 0,252327435 | TBC1D4 | -1,46942769 |
| ENSG0000016 | gene | 0,36531535 | 0,021110411 | 0,256239299 | PBXIP1 | -1,45278572 |
| ENSG0000021 | gene | 0,365751891 | 0,005288625 | 0,239194577 | TRAV10 | -1,45106277 |
| ENSG0000014 | gene | 0,366205614 | 0,039530659 | 0,284051162 | MS4A2 | -1,44927419 |
| ENSG0000025 | gene | 0,366926881 | 0,000836915 | 0,208351114 | AC025884.1 | -1,44643549 |
| ENSG0000014 | gene | 0,367991774 | 0,001841306 | 0,226771947 | ASXL2 | -1,44225458 |
| ENSG0000013 | gene | 0,371554167 | 0,015272117 | 0,252327435 | SYTL2 | -1,42835554 |
| ENSG0000018 | gene | 0,372239568 | 0,004771137 | 0,239194577 | TSPYL1 | -1,42569668 |

| | | | | | | |
|---|---|---|---|---|---|---|
| ENSG000001[illegible] | gene | 0,373399566 | 0,033399522 | 0,274926551 | TOB1 | -1,42120785 |
| ENSG000001[illegible] | gene | 0,374458368 | 0,010851664 | 0,246139857 | CLDND2 | -1,41712277 |
| ENSG000001[illegible] | gene | 0,375106878 | 0,006509403 | 0,239194577 | C16orf54 | -1,41462638 |
| ENSG000001[illegible] | gene | 0,375545774 | 0,006090329 | 0,239194577 | NMUR1 | -1,41293933 |
| ENSG000001[illegible] | gene | 0,379676615 | 0,000728212 | 0,205762815 | MTERF2 | -1,39715695 |
| ENSG000000[illegible] | gene | 0,379748375 | 0,027170853 | 0,267343734 | CA11 | -1,3968843 |
| ENSG000001[illegible] | gene | 0,384051393 | 0,03369779 | 0,275250624 | SIGIRR | -1,38062871 |
| ENSG000001[illegible] | gene | 0,384071198 | 0,04972997 | 0,299174578 | TTC16 | -1,38055432 |
| ENSG000002[illegible] | gene | 0,384213902 | 0,003006575 | 0,237725877 | AP003733.3 | -1,38001837 |
| ENSG000002[illegible] | gene | 0,386292769 | 0,002795591 | 0,237725877 | TRDJ1 | -1,37223342 |
| ENSG000001[illegible] | gene | 0,386728562 | 0,011496771 | 0,249534675 | PYHIN1 | -1,37060677 |
| ENSG000001[illegible] | gene | 0,386999318 | 0,037938348 | 0,28223387 | KLF12 | -1,36959707 |
| ENSG000001[illegible] | gene | 0,387523547 | 0,004090849 | 0,239194577 | AMDHD2 | -1,36764412 |
| ENSG000001[illegible] | gene | 0,387530373 | 0,046562867 | 0,294465415 | CCDC65 | -1,36761871 |
| ENSG000001[illegible] | gene | 0,387655596 | 0,002780998 | 0,237725877 | EDRF1 | -1,3671526 |
| ENSG000002[illegible] | gene | 0,387871233 | 0,0166157 | 0,252327435 | AL365203.2 | -1,36635031 |
| ENSG000001[illegible] | gene | 0,388016942 | 0,046235701 | 0,294036015 | ZNF91 | -1,36580845 |
| ENSG000001[illegible] | gene | 0,388166016 | 0,044909852 | 0,2922856 | DHRS3 | -1,36525428 |
| ENSG000001[illegible] | gene | 0,388402901 | 0,003693006 | 0,239194577 | RPS17 | -1,36437412 |
| ENSG000002[illegible] | gene | 0,388824197 | 0,014526582 | 0,252327435 | KIR2DL3 | -1,36281009 |
| ENSG000001[illegible] | gene | 0,389150544 | 0,002110201 | 0,231249696 | HLA-DQA1 | -1,36159972 |
| ENSG000001[illegible] | gene | 0,390688806 | 0,049497697 | 0,299174578 | MARCHF9 | -1,35590817 |
| ENSG000002[illegible] | gene | 0,391317984 | 0,043447655 | 0,291373337 | LINC00987 | -1,35358668 |
| ENSG000001[illegible] | gene | 0,391795009 | 0,002852244 | 0,237725877 | TFDP2 | -1,35182907 |
| ENSG000002[illegible] | gene | 0,392676663 | 0,03961952 | 0,284051162 | MLLT6 | -1,34858623 |
| ENSG000001[illegible] | gene | 0,393976682 | 0,01983203 | 0,25449173 | ZNF365 | -1,34381785 |
| ENSG000001[illegible] | gene | 0,394756317 | 0,029018737 | 0,270832371 | MARCKSL1 | -1,34096574 |
| ENSG000001[illegible] | gene | 0,39713289 | 0,000166244 | 0,143227097 | ARHGEF9 | -1,33230625 |
| ENSG000002[illegible] | gene | 0,397387557 | 0,023791918 | 0,263452051 | AL365361.1 | -1,3313814 |
| ENSG000001[illegible] | gene | 0,397783436 | 0,009724038 | 0,242573226 | B3GAT1 | -1,32994489 |
| ENSG000001[illegible] | gene | 0,39784384 | 0,030861898 | 0,271804321 | TCN1 | -1,32972584 |
| ENSG000002[illegible] | gene | 0,398395255 | 0,004980711 | 0,239194577 | RPL34P20 | -1,32772763 |
| ENSG000002[illegible] | gene | 0,399341688 | 0,035293941 | 0,2774496 | Z94721.1 | -1,32430441 |
| ENSG000001[illegible] | gene | 0,399433034 | 0,01817747 | 0,252327435 | ZNF831 | -1,32397444 |
| ENSG000002[illegible] | gene | 0,400122045 | 0,018015914 | 0,252327435 | AC026124.2 | -1,32148798 |
| ENSG000002[illegible] | gene | 0,401766903 | 0,026941025 | 0,267343734 | FCGR3A | -1,31556937 |
| ENSG000002[illegible] | gene | 0,402608064 | 0,00497111 | 0,239194577 | LINC01619 | -1,31255203 |
| ENSG000002[illegible] | gene | 0,405238195 | 0,049261422 | 0,299174578 | RNU7-181P | -1,30315794 |
| ENSG000002[illegible] | gene | 0,405725707 | 0,00533353 | 0,239194577 | SNORA75 | -1,30142338 |
| ENSG000001[illegible] | gene | 0,407463917 | 0,027565925 | 0,267527236 | KIF21A | -1,29525579 |
| ENSG000001[illegible] | gene | 0,407573316 | 0,004666603 | 0,239194577 | ANKRD36C | -1,29486849 |
| ENSG000000[illegible] | gene | 0,407872456 | 0,038061258 | 0,28223387 | TBX21 | -1,29381001 |
| ENSG000001[illegible] | gene | 0,409165716 | 0,00308436 | 0,237725877 | SLC25A29 | -1,28924283 |
| ENSG000001[illegible] | gene | 0,411109582 | 0,011902524 | 0,249534675 | SAMD3 | -1,2824051 |
| ENSG000001[illegible] | gene | 0,411116089 | 0,009558292 | 0,242573226 | MAGEE1 | -1,28238226 |
| ENSG000000[illegible] | gene | 0,411137394 | 0,026110784 | 0,266237968 | ZNF264 | -1,2823075 |
| ENSG000001[illegible] | gene | 0,411632329 | 0,043421898 | 0,291373337 | MAF | -1,2805718 |
| ENSG000001[illegible] | gene | 0,412507849 | 0,01960903 | 0,25449173 | SMAD7 | -1,27750652 |
| ENSG000000[illegible] | gene | 0,415613364 | 0,00851907 | 0,242573226 | UNG | -1,26668605 |
| ENSG000001[illegible] | gene | 0,417189619 | 0,000111185 | 0,141858162 | YPEL1 | -1,26122483 |
| ENSG000002[illegible] | gene | 0,417395919 | 0,001476799 | 0,224332495 | AC026979.2 | -1,2605116 |
| ENSG000001[illegible] | gene | 0,419143582 | 0,009997162 | 0,242573226 | WNT10B | -1,25448355 |
| ENSG000002[illegible] | gene | 0,41925532 | 0,017462496 | 0,252327435 | AC016957.2 | -1,254099 |
| ENSG000001[illegible] | gene | 0,419869195 | 0,046550863 | 0,294465415 | TSEN54 | -1,25198815 |
| ENSG000000[illegible] | gene | 0,421181072 | 5,94664E-05 | 0,129441475 | TSPOAP1 | -1,24748749 |
| ENSG000001[illegible] | gene | 0,422281818 | 0,040487206 | 0,286069252 | CAMK2N1 | -1,24372196 |
| ENSG000002[illegible] | gene | 0,422291141 | 0,04096591 | 0,286069252 | PVRIG2P | -1,24369011 |

| | | | | | | |
|---|---|---|---|---|---|---|
| ENSG000002( | gene | 0,424734623 | 0,034137151 | 0,275250624 | AC008750.8 | -1,23536638 |
| ENSG000001( | gene | 0,425153515 | 0,002895858 | 0,237725877 | TOPORS | -1,23394423 |
| ENSG00000( | gene | 0,425736969 | 0,006889996 | 0,239194577 | FHL1 | -1,23196572 |
| ENSG000002( | gene | 0,426910653 | 0,014713681 | 0,252327435 | VDAC2P2 | -1,22799393 |
| ENSG000001( | gene | 0,427047552 | 0,022874273 | 0,260623396 | TOP1MT | -1,22753137 |
| ENSG000001( | gene | 0,428885235 | 0,033121874 | 0,274068881 | ZFPM1 | -1,22133644 |
| ENSG000002( | gene | 0,429078206 | 0,012584008 | 0,249534675 | LINC00299 | -1,22068747 |
| ENSG000000( | gene | 0,429304855 | 0,03829243 | 0,28223387 | HIVEP2 | -1,21992561 |
| ENSG000002( | gene | 0,429555388 | 0,030354091 | 0,271804321 | AC243829.1 | -1,21908393 |
| ENSG000002( | gene | 0,430977482 | 0,030123607 | 0,271804321 | RN7SKP80 | -1,2143156 |
| ENSG000001( | gene | 0,431052903 | 0,03965812 | 0,284051162 | KLRC4 | -1,21406315 |
| ENSG000002( | gene | 0,431170048 | 0,010398741 | 0,244080246 | AJM1 | -1,21367113 |
| ENSG000001( | gene | 0,43127517 | 0,035958272 | 0,279532435 | SNPH | -1,21331944 |
| ENSG000001( | gene | 0,43142475 | 0,042272721 | 0,288197095 | DMPK | -1,21281915 |
| ENSG000001( | gene | 0,432073797 | 0,0271763 | 0,267343734 | DHRS13 | -1,21065035 |
| ENSG000001( | gene | 0,436959687 | 0,009583881 | 0,242573226 | ZNF600 | -1,19442791 |
| ENSG000002( | gene | 0,437075316 | 0,036577662 | 0,280909481 | AC020910.5 | -1,19404619 |
| ENSG000000( | gene | 0,438951275 | 0,038333822 | 0,28223387 | TUBE1 | -1,18786729 |
| ENSG000001( | gene | 0,439282905 | 0,002637262 | 0,237725877 | YTHDF2 | -1,18677774 |
| ENSG000001( | gene | 0,439925032 | 0,00717589 | 0,239194577 | SYTL3 | -1,1846704 |
| ENSG000001( | gene | 0,441189648 | 0,020532056 | 0,256090678 | KIF21B | -1,18052915 |
| ENSG000001( | gene | 0,442605405 | 0,022429655 | 0,258981325 | RRAS2 | -1,17590703 |
| ENSG000001( | gene | 0,443372843 | 0,024497894 | 0,264837204 | PPP1R16B | -1,17340769 |
| ENSG000001( | gene | 0,444537803 | 0,023821974 | 0,263452051 | AGAP1 | -1,16962198 |
| ENSG000001( | gene | 0,444759402 | 0,005798535 | 0,239194577 | LRIG1 | -1,16890299 |
| ENSG000002( | gene | 0,444902662 | 0,033467047 | 0,275163475 | AC010175.1 | -1,16843836 |
| ENSG000000( | gene | 0,446952856 | 0,040897102 | 0,286069252 | JADE2 | -1,16180543 |
| ENSG000001( | gene | 0,447139977 | 0,018446089 | 0,253404085 | DMWD | -1,16120156 |
| ENSG000001( | gene | 0,447605933 | 0,033105733 | 0,274068881 | PHF1 | -1,15969894 |
| ENSG000002( | gene | 0,447955729 | 0,006484656 | 0,239194577 | C2CD4D-AS1 | -1,15857194 |
| ENSG000001( | gene | 0,450861284 | 0,045048448 | 0,2922856 | HPS4 | -1,14924446 |
| ENSG000001( | gene | 0,451486192 | 0,008115586 | 0,242573226 | C1orf21 | -1,14724623 |
| ENSG000001( | gene | 0,451968385 | 0,001543237 | 0,224332495 | AKR1C3 | -1,14570623 |
| ENSG000001( | gene | 0,45280984 | 0,037284629 | 0,2810952 | CMC1 | -1,14302279 |
| ENSG000001( | gene | 0,453731341 | 0,005289201 | 0,239194577 | AHDC1 | -1,14008978 |
| ENSG000001( | gene | 0,453880319 | 0,009490196 | 0,242573226 | PDGFD | -1,13961616 |
| ENSG000001( | gene | 0,454830333 | 0,029213916 | 0,270832371 | SLC9A3R1 | -1,13659962 |
| ENSG000002( | gene | 0,45545321 | 0,026177981 | 0,266237968 | SNORA77 | -1,13462524 |
| ENSG000001( | gene | 0,455514706 | 0,033836044 | 0,275250624 | RNF144A | -1,13443046 |
| ENSG000001( | gene | 0,455560641 | 0,036472279 | 0,280613979 | SYNE1 | -1,13428499 |
| ENSG000000( | gene | 0,456350043 | 0,034113851 | 0,275250624 | IDS | -1,13178723 |
| ENSG000001( | gene | 0,456490953 | 0,003562662 | 0,239194577 | CBX7 | -1,13134183 |
| ENSG000002( | gene | 0,456869638 | 0,015152817 | 0,252327435 | VAMP2 | -1,13014553 |
| ENSG000000( | gene | 0,457330395 | 0,019233147 | 0,254107998 | ICAM3 | -1,12869129 |
| ENSG000001( | gene | 0,458584955 | 0,035908706 | 0,279532435 | ENO2 | -1,12473907 |
| ENSG000002( | gene | 0,459602988 | 0,04975601 | 0,299174578 | AC010894.5 | -1,12153992 |
| ENSG000002( | gene | 0,460900277 | 0,040446625 | 0,286069252 | SNORA12 | -1,11747346 |
| ENSG000002( | gene | 0,461072687 | 0,024936887 | 0,265412724 | AC116348.6 | -1,11693389 |
| ENSG000001( | gene | 0,461487174 | 0,017763331 | 0,252327435 | PDLIM5 | -1,11563754 |
| ENSG000002( | gene | 0,461648037 | 0,005677163 | 0,239194577 | LINC00623 | -1,11513474 |
| ENSG000002( | gene | 0,461790098 | 0,018904924 | 0,254046692 | RNU6ATAC24 | -1,11469086 |
| ENSG000002( | gene | 0,463858972 | 0,045630552 | 0,292582729 | TRAJ19 | -1,10824185 |
| ENSG000000( | gene | 0,464307009 | 0,047914248 | 0,296879689 | NUDCD3 | -1,10684903 |
| ENSG000001( | gene | 0,464766419 | 0,020386545 | 0,256090678 | EPHA4 | -1,10542226 |
| ENSG000001( | gene | 0,467653046 | 0,04629517 | 0,294106514 | XCL2 | -1,09648951 |
| ENSG000002( | gene | 0,46801168 | 0,023060736 | 0,261281502 | AC004918.1 | -1,09538356 |
| ENSG000002( | gene | 0,469141536 | 0,020590067 | 0,256090678 | TMEM191A | -1,09190486 |

| | | | | | | |
|---|---|---|---|---|---|---|
| ENSG0000015 | gene | 0,469442752 | 0,016343375 | 0,252327435 | NADK2 | -1,09097886 |
| ENSG0000025 | gene | 0,469556681 | 0,040258128 | 0,286069252 | AC027559.1 | -1,09062878 |
| ENSG0000018 | gene | 0,469607509 | 0,033752829 | 0,275250624 | TSHZ2 | -1,09047262 |
| ENSG0000027 | gene | 0,46998419 | 0,003974663 | 0,239194577 | LINC02084 | -1,08931587 |
| ENSG0000014 | gene | 0,472158574 | 0,011590493 | 0,249534675 | PNRC1 | -1,08265663 |
| ENSG0000018 | gene | 0,472399942 | 0,009740996 | 0,242573226 | ZFP90 | -1,08191931 |
| ENSG0000020 | gene | 0,473017254 | 0,037928864 | 0,28223387 | PDCD4-AS1 | -1,08003529 |
| ENSG0000000 | gene | 0,474386615 | 0,013929844 | 0,252327435 | TTC22 | -1,07586479 |
| ENSG0000019 | gene | 0,475046112 | 0,038981049 | 0,282785762 | MIR103A2 | -1,07386053 |
| ENSG0000009 | gene | 0,47511415 | 0,031155047 | 0,271804321 | WHRN | -1,07365392 |
| ENSG0000013 | gene | 0,476003889 | 0,040731663 | 0,286069252 | RHOF | -1,07095473 |
| ENSG0000025 | gene | 0,476028076 | 0,011808751 | 0,249534675 | KDM2B-DT | -1,07088143 |
| ENSG0000014 | gene | 0,47613662 | 0,025839193 | 0,266059172 | NARF | -1,0705525 |
| ENSG0000000 | gene | 0,476317207 | 0,000352621 | 0,19726282 | CRY1 | -1,07000543 |
| ENSG0000027 | gene | 0,476324445 | 0,035510563 | 0,277679087 | AC133644.2 | -1,06998351 |
| ENSG0000025 | gene | 0,476369663 | 0,023571741 | 0,263452051 | DENND11 | -1,06984656 |
| ENSG0000013 | gene | 0,477165597 | 0,005081774 | 0,239194577 | CNOT6L | -1,06743806 |
| ENSG0000024 | gene | 0,478193285 | 0,048177328 | 0,296879689 | PDCL3P4 | -1,06433422 |
| ENSG0000015 | gene | 0,47834048 | 0,032288793 | 0,272856143 | IFNAR2 | -1,06389021 |
| ENSG0000026 | gene | 0,478483089 | 0,003199533 | 0,238752762 | AC004687.1 | -1,06346016 |
| ENSG0000028 | gene | 0,478515714 | 0,045281976 | 0,2922856 | AC079781.4 | -1,06336179 |
| ENSG0000010 | gene | 0,47980774 | 0,001506477 | 0,224332495 | ZNF574 | -1,05947166 |
| ENSG0000025 | gene | 0,480825725 | 0,040820495 | 0,286069252 | AC037459.2 | -1,05641401 |
| ENSG0000008 | gene | 0,481344773 | 0,024209952 | 0,264579509 | PPP1R15A | -1,05485747 |
| ENSG0000014 | gene | 0,481800803 | 0,013019489 | 0,250263503 | USP53 | -1,0534913 |
| ENSG0000016 | gene | 0,481869534 | 0,006776727 | 0,239194577 | SEMA4C | -1,0532855 |
| ENSG0000012 | gene | 0,482137944 | 0,021564569 | 0,256239299 | TBCC | -1,05248212 |
| ENSG0000021 | gene | 0,48261918 | 0,047600911 | 0,296879689 | TRAJ21 | -1,05104284 |
| ENSG0000005 | gene | 0,483272824 | 0,037023383 | 0,281022131 | TRAF1 | -1,04909022 |
| ENSG0000024 | gene | 0,483637314 | 0,007496605 | 0,239194577 | DNAJC25-GN | -1,04800254 |
| ENSG0000016 | gene | 0,484025405 | 0,009237008 | 0,242573226 | CNNM3 | -1,04684532 |
| ENSG0000019 | gene | 0,48620843 | 0,019766142 | 0,25449173 | ITPRIPL1 | -1,04035319 |
| ENSG0000021 | gene | 0,486587578 | 0,01877431 | 0,254046692 | TRAJ31 | -1,0392286 |
| ENSG0000010 | gene | 0,48747588 | 0,014099197 | 0,252327435 | AGFG2 | -1,03659726 |
| ENSG0000005 | gene | 0,487705721 | 0,013801068 | 0,252327435 | ARID4B | -1,0359172 |
| ENSG0000015 | gene | 0,489950924 | 0,020776473 | 0,256090678 | TTC39B | -1,02929085 |
| ENSG0000017 | gene | 0,490239178 | 0,028348615 | 0,269111214 | RESF1 | -1,02844231 |
| ENSG0000018 | gene | 0,490606749 | 0,005001267 | 0,239194577 | PTCH1 | -1,02736101 |
| ENSG0000008 | gene | 0,49113593 | 0,025039722 | 0,265635455 | ZBTB25 | -1,02580573 |
| ENSG0000017 | gene | 0,491835485 | 0,03120162 | 0,271804321 | ZBTB4 | -1,02375227 |
| ENSG0000017 | gene | 0,492120539 | 0,014063518 | 0,252327435 | GK5 | -1,02291637 |
| ENSG0000024 | gene | 0,492611046 | 0,0137698 | 0,252327435 | NAA80 | -1,02147912 |
| ENSG0000020 | gene | 0,492950882 | 0,015029567 | 0,252327435 | SNORD20 | -1,02048419 |
| ENSG0000012 | gene | 0,493430728 | 0,013582274 | 0,252327435 | PIGT | -1,01908053 |
| ENSG0000011 | gene | 0,493473409 | 0,013964278 | 0,252327435 | BRD8 | -1,01895575 |
| ENSG0000016 | gene | 0,493483489 | 0,027259256 | 0,267343734 | PLD4 | -1,01892628 |
| ENSG0000019 | gene | 0,494511446 | 0,014049913 | 0,252327435 | NOL4L | -1,01592418 |
| ENSG0000016 | gene | 0,494839064 | 0,00771306 | 0,239194577 | GFI1 | -1,0149687 |
| ENSG0000020 | gene | 0,495460589 | 0,006855255 | 0,239194577 | SNORA33 | -1,01315779 |
| ENSG0000005 | gene | 0,495513431 | 0,0456632 | 0,292582729 | FLYWCH1 | -1,01300393 |
| ENSG0000016 | gene | 0,496096721 | 0,019871648 | 0,25449173 | EOMES | -1,01130667 |
| ENSG0000014 | gene | 0,496287196 | 0,001332264 | 0,224332495 | CBX4 | -1,01075286 |
| ENSG0000011 | gene | 0,496903443 | 0,038788724 | 0,282581813 | FOXP1 | -1,00896256 |
| ENSG0000027 | gene | 0,497333247 | 0,022145769 | 0,25813016 | MIR6812 | -1,00771521 |
| ENSG0000014 | gene | 0,497586827 | 0,002280178 | 0,23273779 | TMEM181 | -1,0069798 |
| ENSG0000027 | gene | 0,497614419 | 0,013663442 | 0,252327435 | TRAJ37 | -1,0068998 |
| ENSG0000021 | gene | 0,498069107 | 0,00325136 | 0,238752762 | MMP23A | -1,00558216 |

| | | | | | | |
|---|---|---|---|---|---|---|
| ENSG000001. | gene | 0,498346286 | 0,045250176 | 0,2922856 | KBTBD4 | -1,00477952 |
| ENSG000002 | gene | 0,498648094 | 0,048430464 | 0,297073163 | AC093010.3 | -1,00390606 |
| ENSG000001. | gene | 0,499149094 | 0,026211518 | 0,266237968 | MIF4GD | -1,00245729 |
| ENSG000002 | gene | 0,500432244 | 0,044789801 | 0,2922856 | TRGV9 | -0,99875335 |
| ENSG000002 | gene | 0,500502402 | 0,018692654 | 0,254046692 | AL139246.5 | -0,9985511 |
| ENSG000000 | gene | 0,502544885 | 0,023971148 | 0,26347532 | PTPN4 | -0,99267564 |
| ENSG000001 | gene | 0,502641726 | 0,039159662 | 0,283069015 | NCR1 | -0,99239766 |
| ENSG000000 | gene | 0,503460606 | 0,026928526 | 0,267343734 | SYNE2 | -0,9900492 |
| ENSG000002 | gene | 0,504220432 | 0,027284684 | 0,267343734 | SNORA3B | -0,98787352 |
| ENSG000001 | gene | 0,504397463 | 0,009135318 | 0,242573226 | RPUSD2 | -0,98736708 |
| ENSG000001 | gene | 0,504401869 | 0,013718483 | 0,252327435 | TMUB1 | -0,98735447 |
| ENSG000001 | gene | 0,505333729 | 0,028284383 | 0,269057501 | TSHZ1 | -0,98469162 |
| ENSG000001 | gene | 0,50556142 | 0,015913387 | 0,252327435 | FUT11 | -0,98404172 |
| ENSG000001. | gene | 0,505695508 | 0,029373022 | 0,271124889 | RBM48 | -0,98365913 |
| ENSG000001 | gene | 0,505937849 | 0,040931184 | 0,286069252 | DCUN1D4 | -0,98296792 |
| ENSG000001 | gene | 0,506309782 | 0,008245631 | 0,242573226 | GPR183 | -0,98190774 |
| ENSG000001 | gene | 0,507280056 | 0,018254865 | 0,252327435 | PLEKHA1 | -0,97914565 |
| ENSG000002 | gene | 0,507310383 | 0,040905898 | 0,286069252 | AC005831.1 | -0,97905941 |
| ENSG000001 | gene | 0,507511033 | 0,014323997 | 0,252327435 | ZNF830 | -0,97848891 |
| ENSG000001 | gene | 0,507827364 | 0,001562524 | 0,224332495 | RCE1 | -0,97758996 |
| ENSG000002 | gene | 0,508004649 | 0,049611421 | 0,299174578 | MIR4518 | -0,97708639 |
| ENSG000000 | gene | 0,509475372 | 0,040891519 | 0,286069252 | RNF216 | -0,97291569 |
| ENSG000001 | gene | 0,509969695 | 0,044139238 | 0,2922856 | S100B | -0,97151658 |
| ENSG000001 | gene | 0,511855563 | 0,035132528 | 0,277400936 | PLXNA3 | -0,96619133 |
| ENSG000001 | gene | 0,511963611 | 0,019126963 | 0,254046692 | SKI | -0,96588682 |
| ENSG000001 | gene | 0,51231469 | 0,013506861 | 0,252327435 | ZXDB | -0,96489783 |
| ENSG000001 | gene | 0,513247139 | 0,032203341 | 0,272856143 | TPK1 | -0,96227442 |
| ENSG000001 | gene | 0,513559403 | 0,038420111 | 0,28223387 | FBXL14 | -0,96139693 |
| ENSG000001 | gene | 0,513831754 | 0,008970774 | 0,242573226 | ADGRG5 | -0,96063205 |
| ENSG000001 | gene | 0,51386971 | 0,006908498 | 0,239194577 | RAB11FIP4 | -0,96052548 |
| ENSG000000 | gene | 0,514254527 | 0,049612874 | 0,299174578 | KIF22 | -0,95944551 |
| ENSG000001 | gene | 0,51438815 | 0,012826658 | 0,249534675 | ZNF83 | -0,95907069 |
| ENSG000001 | gene | 0,514532452 | 0,047926527 | 0,296879689 | LUC7L3 | -0,95866602 |
| ENSG000002 | gene | 0,515229399 | 0,041317295 | 0,286109654 | Z94721.3 | -0,95671318 |
| ENSG000001 | gene | 0,516255363 | 0,013446957 | 0,252327435 | THAP1 | -0,95384323 |
| ENSG000000 | gene | 0,516398871 | 0,012743538 | 0,249534675 | OSBPL7 | -0,95344225 |
| ENSG000001. | gene | 0,516743039 | 0,040313963 | 0,286069252 | SBDS | -0,95248105 |
| ENSG000002 | gene | 0,51778438 | 0,032175677 | 0,272856143 | SNORD63 | -0,94957665 |
| ENSG000001 | gene | 0,518019075 | 0,010203802 | 0,244080246 | ABCA2 | -0,94892287 |
| ENSG000002 | gene | 0,518283481 | 0,037480486 | 0,281299108 | AC005674.2 | -0,94818668 |
| ENSG000001 | gene | 0,518798622 | 0,040922226 | 0,286069252 | RIOX1 | -0,94675345 |
| ENSG000001 | gene | 0,518941404 | 0,000530589 | 0,200582152 | ARHGAP25 | -0,94635645 |
| ENSG000000 | gene | 0,520350506 | 0,02560129 | 0,265635455 | KAT6A | -0,94244435 |
| ENSG000001 | gene | 0,52159992 | 0,000745883 | 0,205762815 | ANKRD49 | -0,93898445 |
| ENSG000001 | gene | 0,52176025 | 0,000699011 | 0,205762815 | YRDC | -0,93854106 |
| ENSG000001 | gene | 0,522356491 | 0,032292566 | 0,272856143 | MOAP1 | -0,93689336 |
| ENSG000000 | gene | 0,523237125 | 0,000300151 | 0,180214301 | TTC38 | -0,93446319 |
| ENSG000002 | gene | 0,523325823 | 0,026925125 | 0,267343734 | MIR6752 | -0,93421865 |
| ENSG000001 | gene | 0,523595549 | 0,011957439 | 0,249534675 | DNAAF2 | -0,93347526 |
| ENSG000002 | gene | 0,524298552 | 0,037148911 | 0,2810952 | MIR6731 | -0,93153953 |
| ENSG000001 | gene | 0,524381575 | 0,049668571 | 0,299174578 | CHST2 | -0,9313111 |
| ENSG000002 | gene | 0,524735004 | 0,03603303 | 0,27990041 | SNORD15B | -0,93033906 |
| ENSG000001 | gene | 0,524895333 | 0,034751528 | 0,27613317 | IQCN | -0,92989833 |
| ENSG000002 | gene | 0,525231808 | 0,026294225 | 0,266237968 | AC242842.2 | -0,92897381 |
| ENSG000001 | gene | 0,525918551 | 0,007234271 | 0,239194577 | KRR1 | -0,92708871 |
| ENSG000002 | gene | 0,527055939 | 0,014369445 | 0,252327435 | AC011477.1 | -0,923972 |
| ENSG000002 | gene | 0,527329708 | 0,044085104 | 0,2922856 | RNU6-48P | -0,92322282 |

| | | | | | | |
|---|---|---|---|---|---|---|
| ENSG000001( | gene | 0,527869613 | 0,016138742 | 0,252327435 | MRPL55 | -0,92174648 |
| ENSG000001; | gene | 0,528391028 | 0,016078828 | 0,252327435 | CREBZF | -0,92032212 |
| ENSG000001' | gene | 0,529022404 | 0,047912419 | 0,296879689 | DROSHA | -0,91859927 |
| ENSG000002, | gene | 0,529691844 | 0,030359801 | 0,271804321 | PRAF2 | -0,9167748 |
| ENSG000001; | gene | 0,53022662 | 0,015811252 | 0,252327435 | PARP15 | -0,91531899 |
| ENSG000002( | gene | 0,53088453 | 0,033546193 | 0,275245975 | TRBJ1-5 | -0,91352999 |
| ENSG000001; | gene | 0,531163498 | 0,02086188 | 0,256090678 | CERS5 | -0,91277209 |
| ENSG000002; | gene | 0,532882506 | 0,03423494 | 0,275250624 | AC146949.1 | -0,90811062 |
| ENSG00000( | gene | 0,533141339 | 0,04969301 | 0,299174578 | RGS1 | -0,90741004 |
| ENSG000002; | gene | 0,533311672 | 0,035441986 | 0,277679087 | AC099804.1 | -0,90694919 |
| ENSG000001; | gene | 0,534416663 | 0,01981114 | 0,25449173 | CLSTN3 | -0,9039631 |
| ENSG000002; | gene | 0,534973202 | 0,019126403 | 0,254046692 | LINC00852 | -0,90246147 |
| ENSG000001( | gene | 0,535447554 | 0,02620232 | 0,266237968 | LSS | -0,90118282 |
| ENSG000001; | gene | 0,535574896 | 0,042468426 | 0,288598684 | CLCF1 | -0,90083976 |
| ENSG000001; | gene | 0,537633249 | 0,007452914 | 0,239194577 | DCP2 | -0,89530573 |
| ENSG000000( | gene | 0,538148736 | 0,041392469 | 0,286241825 | ANKRD44 | -0,89392313 |
| ENSG000002; | gene | 0,538412259 | 0,000591361 | 0,205762815 | AL645933.3 | -0,89321684 |
| ENSG000000( | gene | 0,538494574 | 0,003591311 | 0,239194577 | ABCB1 | -0,89299629 |
| ENSG000002( | gene | 0,539660218 | 0,012379869 | 0,249534675 | AC092652.2 | -0,88987676 |
| ENSG000001( | gene | 0,53990995 | 0,01244162 | 0,249534675 | ZC3HAV1 | -0,88920929 |
| ENSG000001( | gene | 0,541114388 | 0,032758315 | 0,274068881 | DFFB | -0,88599449 |
| ENSG000000( | gene | 0,541240217 | 0,001943633 | 0,2305768 | RASSF1 | -0,88565905 |
| ENSG000001( | gene | 0,541896589 | 0,006378033 | 0,239194577 | CFAP410 | -0,88391053 |
| ENSG000001, | gene | 0,543185125 | 0,043868055 | 0,292071193 | PPIP5K2 | -0,88048412 |
| ENSG000001; | gene | 0,543592594 | 0,017068493 | 0,252327435 | ZHX2 | -0,8794023 |
| ENSG000000; | gene | 0,545901459 | 0,025348568 | 0,265635455 | ZNF839 | -0,87328754 |
| ENSG000001; | gene | 0,547258405 | 0,01945647 | 0,254107998 | ANKRD36 | -0,86970589 |
| ENSG000001( | gene | 0,547505809 | 0,021488207 | 0,256239299 | CAMK1D | -0,86905382 |
| ENSG000001( | gene | 0,547997483 | 0,024471885 | 0,264837204 | ARHGAP5 | -0,86775883 |
| ENSG000002; | gene | 0,549581408 | 0,006610905 | 0,239194577 | AL662844.4 | -0,8635949 |
| ENSG000000; | gene | 0,550035673 | 0,014513778 | 0,252327435 | ARID4A | -0,86240291 |
| ENSG000001( | gene | 0,55086902 | 0,01639441 | 0,252327435 | CYTH1 | -0,86021877 |
| ENSG000001; | gene | 0,551026174 | 0,023033484 | 0,261281502 | DNAJB9 | -0,85980725 |
| ENSG000001' | gene | 0,551886048 | 0,043852186 | 0,292071193 | CEP85L | -0,85755768 |
| ENSG000001! | gene | 0,553059158 | 0,040355931 | 0,286069252 | PPP2R2B | -0,85449429 |
| ENSG000001( | gene | 0,553287907 | 0,032745459 | 0,274068881 | TLE5 | -0,8538977 |
| ENSG000001; | gene | 0,554264615 | 0,013001712 | 0,250263503 | TRERF1 | -0,85135319 |
| ENSG000001; | gene | 0,556633385 | 0,037405758 | 0,281149169 | KLC4 | -0,84520066 |
| ENSG000002; | gene | 0,557065735 | 0,011693034 | 0,249534675 | MIR6819 | -0,84408052 |
| ENSG000002; | gene | 0,557249501 | 0,014960059 | 0,252327435 | ABHD17AP6 | -0,84360467 |
| ENSG000001! | gene | 0,557588906 | 0,038462838 | 0,28223387 | FOXO1 | -0,84272624 |
| ENSG000000, | gene | 0,557797416 | 0,025271537 | 0,265635455 | RSF1 | -0,84218684 |
| ENSG000001( | gene | 0,560096071 | 0,042331096 | 0,288197095 | TNFRSF18 | -0,83625379 |
| ENSG000001! | gene | 0,560612311 | 0,03255749 | 0,274068881 | ARMCX6 | -0,83492467 |
| ENSG000001( | gene | 0,561725283 | 0,016390949 | 0,252327435 | CNPY4 | -0,83206335 |
| ENSG000001; | gene | 0,562155385 | 0,007675094 | 0,239194577 | NAP1L5 | -0,83095914 |
| ENSG000002( | gene | 0,562764345 | 0,001623978 | 0,224332495 | FP565260.6 | -0,82939717 |
| ENSG000002; | gene | 0,563627799 | 0,017046478 | 0,252327435 | HOXB-AS1 | -0,82718533 |
| ENSG000001; | gene | 0,563898448 | 0,004965278 | 0,239194577 | FGD3 | -0,82649272 |
| ENSG000001; | gene | 0,564825108 | 0,005447641 | 0,239194577 | TAF7 | -0,82412387 |
| ENSG000002' | gene | 0,56497985 | 0,036685919 | 0,280909481 | TRDJ2 | -0,82372868 |
| ENSG000001, | gene | 0,565344547 | 0,044569525 | 0,2922856 | SCML4 | -0,82279771 |
| ENSG000001( | gene | 0,565677297 | 0,033040721 | 0,274068881 | ARHGAP12 | -0,82194882 |
| ENSG000001! | gene | 0,566656453 | 0,021150549 | 0,256239299 | TOX | -0,81945376 |
| ENSG000002; | gene | 0,567552626 | 0,039545959 | 0,284051162 | ORAI1 | -0,81717392 |
| ENSG000000! | gene | 0,567574971 | 0,000178726 | 0,143227097 | CDK17 | -0,81711712 |
| ENSG000001; | gene | 0,567631477 | 0,017051093 | 0,252327435 | EGLN1 | -0,8169735 |

| | | | | | | |
|---|---|---|---|---|---|---|
| ENSG000000 | gene | 0,567641757 | 0,00741721 | 0,239194577 | PPP2R5C | -0,81694737 |
| ENSG000001 | gene | 0,56948023 | 0,046091269 | 0,293382847 | AGAP2 | -0,81228234 |
| ENSG000000 | gene | 0,569912759 | 0,003088465 | 0,237725877 | BOD1L1 | -0,811187 |
| ENSG000002 | gene | 0,571388098 | 0,02098314 | 0,256239299 | AL021707.8 | -0,80745711 |
| ENSG000001 | gene | 0,571658008 | 0,024264157 | 0,264598559 | GLCCI1 | -0,80677578 |
| ENSG000002 | gene | 0,571794044 | 0,013901029 | 0,252327435 | AL691432.2 | -0,8064325 |
| ENSG000002 | gene | 0,572508715 | 0,004396267 | 0,239194577 | AC005332.3 | -0,80463044 |
| ENSG000002 | gene | 0,573679782 | 0,023350241 | 0,263452051 | N4BP2L2-IT2 | -0,80168242 |
| ENSG000001 | gene | 0,574305202 | 0,008493577 | 0,242573226 | ANKRD12 | -0,80011047 |
| ENSG000001 | gene | 0,575262165 | 0,035310597 | 0,2774496 | COQ10A | -0,79770851 |
| ENSG000001 | gene | 0,575715265 | 0,035489588 | 0,277679087 | IKZF3 | -0,79657263 |
| ENSG000001 | gene | 0,576174195 | 0,019652467 | 0,25449173 | PPP1R2 | -0,79542305 |
| ENSG000001 | gene | 0,576237258 | 0,042233425 | 0,288197095 | TAGLN2 | -0,79526515 |
| ENSG000001 | gene | 0,576621435 | 0,00445753 | 0,239194577 | EIF1AD | -0,79430363 |
| ENSG000002 | gene | 0,576709266 | 0,018123613 | 0,252327435 | AC116366.1 | -0,79408389 |
| ENSG000000 | gene | 0,577034914 | 0,008328267 | 0,242573226 | FYN | -0,79326948 |
| ENSG000001 | gene | 0,577687183 | 0,047284825 | 0,295914294 | NECTIN1 | -0,79163961 |
| ENSG000001 | gene | 0,577788608 | 0,02459337 | 0,264837204 | AAK1 | -0,79138634 |
| ENSG000001 | gene | 0,57790469 | 0,036482292 | 0,280613979 | CBLB | -0,79109652 |
| ENSG000001 | gene | 0,578754353 | 0,048044211 | 0,296879689 | KCNA3 | -0,78897695 |
| ENSG000000 | gene | 0,578920908 | 0,031705342 | 0,272282875 | SIDT1 | -0,78856183 |
| ENSG000001 | gene | 0,579225028 | 0,043798053 | 0,292071193 | CD244 | -0,78780415 |
| ENSG000001 | gene | 0,580360449 | 0,029147572 | 0,270832371 | DYNLT3 | -0,78497889 |
| ENSG000001 | gene | 0,580361755 | 0,035247786 | 0,2774496 | TBCCD1 | -0,78497564 |
| ENSG000001 | gene | 0,581126343 | 0,031214023 | 0,271804321 | USP20 | -0,78307624 |
| ENSG000002 | gene | 0,581619475 | 0,017765396 | 0,252327435 | PTOV1-AS2 | -0,78185252 |
| ENSG000001 | gene | 0,582595974 | 0,005326533 | 0,239194577 | PLCD1 | -0,77943236 |
| ENSG000001 | gene | 0,583665652 | 0,045779928 | 0,292595946 | ZSCAN25 | -0,77678592 |
| ENSG000002 | gene | 0,583945099 | 0,044042293 | 0,2922856 | AC021188.1 | -0,77609536 |
| ENSG000001 | gene | 0,584053994 | 0,005744361 | 0,239194577 | ENO3 | -0,77582635 |
| ENSG000002 | gene | 0,585629687 | 0,038555668 | 0,28223387 | AL356273.6 | -0,7719394 |
| ENSG000000 | gene | 0,585998353 | 0,020831019 | 0,256090678 | ING3 | -0,77103149 |
| ENSG000000 | gene | 0,586785462 | 0,040420178 | 0,286069252 | CBFB | -0,76909497 |
| ENSG000001 | gene | 0,588689381 | 0,006200871 | 0,239194577 | ARRDC3 | -0,76442149 |
| ENSG000001 | gene | 0,589081333 | 0,01913187 | 0,254046692 | TCEA2 | -0,76346126 |
| ENSG000002 | gene | 0,58947019 | 0,015885239 | 0,252327435 | AL356356.2 | -0,76250924 |
| ENSG000001 | gene | 0,589477833 | 0,016562783 | 0,252327435 | CD47 | -0,76249053 |
| ENSG000001 | gene | 0,590053043 | 0,002633522 | 0,237725877 | BMI1 | -0,76108344 |
| ENSG000001 | gene | 0,59084712 | 0,014866155 | 0,252327435 | STAG1 | -0,75914321 |
| ENSG000001 | gene | 0,59158611 | 0,020433007 | 0,256090678 | PNMA1 | -0,75733991 |
| ENSG000000 | gene | 0,591871267 | 0,030719556 | 0,271804321 | TBC1D10A | -0,75664467 |
| ENSG000000 | gene | 0,591959004 | 0,044380671 | 0,2922856 | FYB1 | -0,75643083 |
| ENSG000001 | gene | 0,593070724 | 0,008600234 | 0,242573226 | FBXO33 | -0,75372394 |
| ENSG000002 | gene | 0,593140248 | 0,026485411 | 0,267131022 | MT1XP1 | -0,75355482 |
| ENSG000002 | gene | 0,593451859 | 0,026819268 | 0,267343734 | HLA-F | -0,75279709 |
| ENSG000000 | gene | 0,594617437 | 0,007420761 | 0,239194577 | KMT2E | -0,74996632 |
| ENSG000002 | gene | 0,597013116 | 0,005654477 | 0,239194577 | SNORD100 | -0,74416547 |
| ENSG000002 | gene | 0,597549771 | 0,011663634 | 0,249534675 | FAM229A | -0,74286921 |
| ENSG000002 | gene | 0,598361593 | 0,023700211 | 0,263452051 | TRGV7 | -0,74091052 |
| ENSG000002 | gene | 0,59852888 | 0,016708966 | 0,252327435 | RN7SL809P | -0,74050723 |
| ENSG000002 | gene | 0,599275738 | 0,029129291 | 0,270832371 | AC069281.2 | -0,73870813 |
| ENSG000002 | gene | 0,599805375 | 0,036469096 | 0,280613979 | COA6-AS1 | -0,73743364 |
| ENSG000002 | gene | 0,600229723 | 0,044546033 | 0,2922856 | ZBTB10 | -0,73641333 |
| ENSG000001 | gene | 0,600251274 | 0,009999999 | 0,242573226 | MECP2 | -0,73636153 |
| ENSG000002 | gene | 0,601275264 | 0,029930769 | 0,271804321 | UBAP1L | -0,73390249 |
| ENSG000001 | gene | 0,601599849 | 0,013816214 | 0,252327435 | CAMK2G | -0,73312389 |
| ENSG000001 | gene | 0,601828682 | 0,037992704 | 0,28223387 | MVD | -0,73257523 |

| | | | | | | |
|---|---|---|---|---|---|---|
| ENSG000001( | gene | 0,602196226 | 0,011173005 | 0,249534675 | TNRC6B | -0,73169443 |
| ENSG000001( | gene | 0,602473914 | 0,024179744 | 0,264579509 | CPSF4 | -0,73102932 |
| ENSG000002‹ | gene | 0,605600916 | 0,040963023 | 0,286069252 | APOBEC3D | -0,72356071 |
| ENSG000002( | gene | 0,605808655 | 0,034703979 | 0,276090036 | MIR573 | -0,72306591 |
| ENSG000002( | gene | 0,606350086 | 0,019627626 | 0,25449173 | SRSF8 | -0,7217771 |
| ENSG000001( | gene | 0,608080791 | 0,003709305 | 0,239194577 | ZNF148 | -0,71766508 |
| ENSG000001{ | gene | 0,608593086 | 0,035461622 | 0,277679087 | SIVA1 | -0,71645015 |
| ENSG0000017 | gene | 0,609646346 | 0,03058279 | 0,271804321 | EIF1 | -0,71395551 |
| ENSG000001( | gene | 0,61013672 | 0,044561351 | 0,2922856 | SLAMF6 | -0,71279553 |
| ENSG000001( | gene | 0,611228387 | 0,042317341 | 0,288197095 | RUBCNL | -0,71021655 |
| ENSG000001' | gene | 0,611424638 | 0,027101526 | 0,267343734 | LNPEP | -0,70975341 |
| ENSG000001! | gene | 0,611838115 | 0,041253851 | 0,286069252 | SSBP3 | -0,70877811 |
| ENSG000000{ | gene | 0,611921306 | 0,028256159 | 0,269039756 | RSBN1 | -0,70858196 |
| ENSG000000{ | gene | 0,612119598 | 0,01439729 | 0,252327435 | ERO1B | -0,70811454 |
| ENSG0000007 | gene | 0,612556718 | 0,039800811 | 0,284088728 | ATP2A3 | -0,70708466 |
| ENSG000000! | gene | 0,613851104 | 0,009487712 | 0,242573226 | PUM2 | -0,70403934 |
| ENSG000001‹ | gene | 0,614277007 | 0,041308524 | 0,286109654 | DGKZ | -0,70303871 |
| ENSG000001( | gene | 0,614547132 | 0,015474749 | 0,252327435 | DYRK1B | -0,70240443 |
| ENSG0000027 | gene | 0,614729621 | 0,024525161 | 0,264837204 | AC004918.5 | -0,70197609 |
| ENSG000000( | gene | 0,614865714 | 0,019468253 | 0,254107998 | THUMPD1 | -0,70165673 |
| ENSG000001' | gene | 0,614960289 | 0,002458274 | 0,237725877 | DDX6 | -0,70143484 |
| ENSG000001: | gene | 0,615317554 | 0,015991301 | 0,252327435 | RFTN1 | -0,70059694 |
| ENSG000001{ | gene | 0,617226151 | 0,037894757 | 0,28223387 | RTL6 | -0,69612891 |
| ENSG0000027 | gene | 0,61738016 | 0,026815085 | 0,267343734 | LINC00653 | -0,69576897 |
| ENSG000001! | gene | 0,617460355 | 0,037345409 | 0,2810952 | CDC42SE1 | -0,69558159 |
| ENSG000002( | gene | 0,617650918 | 0,010786343 | 0,246139857 | MIR4639 | -0,6951364 |
| ENSG0000012 | gene | 0,617809385 | 0,017491308 | 0,252327435 | TARDBP | -0,69476631 |
| ENSG0000012 | gene | 0,618152735 | 0,006390672 | 0,239194577 | ZBTB1 | -0,69396475 |
| ENSG000002: | gene | 0,618155364 | 0,03092022 | 0,271804321 | USP32P2 | -0,69395861 |
| ENSG000001! | gene | 0,61918168 | 0,006092965 | 0,239194577 | RABGAP1L | -0,69156531 |
| ENSG0000017 | gene | 0,619275707 | 0,011782714 | 0,249534675 | RNF139 | -0,69134624 |
| ENSG000001: | gene | 0,61947902 | 0,00845043 | 0,242573226 | CSNK1G2 | -0,69087267 |
| ENSG0000012 | gene | 0,619597571 | 0,039777825 | 0,284088728 | SLC44A2 | -0,69059661 |
| ENSG0000007 | gene | 0,62049634 | 0,005534198 | 0,239194577 | STK10 | -0,68850539 |
| ENSG000001{ | gene | 0,620781902 | 0,038637275 | 0,28223387 | MRPL30 | -0,6878416 |
| ENSG000001( | gene | 0,620892659 | 0,024264092 | 0,264598559 | PCSK7 | -0,68758422 |
| ENSG000001! | gene | 0,621237558 | 0,024210567 | 0,264579509 | ARID5B | -0,68678304 |
| ENSG0000027 | gene | 0,621499657 | 0,015129728 | 0,252327435 | AC139256.3 | -0,6861745 |
| ENSG0000027 | gene | 0,622321928 | 0,01559018 | 0,252327435 | AC005332.5 | -0,68426701 |
| ENSG000001: | gene | 0,622562944 | 0,000967375 | 0,224332495 | PCNX2 | -0,68370839 |
| ENSG000001' | gene | 0,624358943 | 0,020608466 | 0,256090678 | EXOC8 | -0,67955242 |
| ENSG0000017 | gene | 0,624502049 | 0,028814041 | 0,270317021 | ERN1 | -0,67922179 |
| ENSG000000' | gene | 0,62642447 | 0,024746655 | 0,264837204 | CCDC88C | -0,67478752 |
| ENSG000001( | gene | 0,626720816 | 0,027651106 | 0,267527236 | AXIN1 | -0,67410518 |
| ENSG0000027 | gene | 0,627258043 | 0,010371325 | 0,244080246 | AL662884.1 | -0,67286903 |
| ENSG000001' | gene | 0,628319915 | 0,03054666 | 0,271804321 | CDKN1B | -0,67042879 |
| ENSG000002{ | gene | 0,629367709 | 0,003950658 | 0,239194577 | TRG-AS1 | -0,66802493 |
| ENSG000001: | gene | 0,629381242 | 0,016071655 | 0,252327435 | SETD1B | -0,66799391 |
| ENSG000002: | gene | 0,630623132 | 0,004635843 | 0,239194577 | AC108693.1 | -0,66515 |
| ENSG000001' | gene | 0,630988017 | 0,005401434 | 0,239194577 | NCBP2 | -0,66431549 |
| ENSG000001( | gene | 0,631772707 | 0,041194434 | 0,286069252 | APMAP | -0,66252248 |
| ENSG0000027 | gene | 0,632019486 | 0,028369126 | 0,269111214 | AC092747.4 | -0,66195906 |
| ENSG000001‹ | gene | 0,63316687 | 0,022778622 | 0,260623396 | FBXL17 | -0,65934232 |
| ENSG000001' | gene | 0,633611456 | 0,04199799 | 0,287700327 | CRBN | -0,65832968 |
| ENSG000001! | gene | 0,633769196 | 0,002033106 | 0,2305768 | FBXO32 | -0,65797055 |
| ENSG000001( | gene | 0,634238003 | 0,026254786 | 0,266237968 | CRTC2 | -0,65690377 |
| ENSG000001! | gene | 0,634385559 | 0,020748644 | 0,256090678 | RASA2 | -0,65656817 |

| | | | | | | |
|---|---|---|---|---|---|---|
| ENSG000001( | gene | 0,635978179 | 0,025975777 | 0,266237968 | CCSER2 | -0,65295083 |
| ENSG000001! | gene | 0,636623364 | 0,017246681 | 0,252327435 | IFIT5 | -0,65148799 |
| ENSG000000! | gene | 0,638357513 | 0,040058488 | 0,285528621 | ARHGAP4 | -0,64756346 |
| ENSG000002; | gene | 0,639076053 | 0,003097633 | 0,237725877 | MIR6802 | -0,64594047 |
| ENSG000002‹ | gene | 0,639665712 | 0,013456961 | 0,252327435 | STAG3L5P | -0,64460994 |
| ENSG000001; | gene | 0,642650218 | 0,005358625 | 0,239194577 | PIN1 | -0,63789437 |
| ENSG000001' | gene | 0,643549383 | 0,036301861 | 0,280613979 | BCAS2 | -0,63587724 |
| ENSG000002' | gene | 0,645613995 | 0,009432732 | 0,242573226 | DNASE1 | -0,63125624 |
| ENSG000001( | gene | 0,645873598 | 0,006607786 | 0,239194577 | JAK1 | -0,63067625 |
| ENSG000001( | gene | 0,646492115 | 0,02641688 | 0,266967423 | TSC22D4 | -0,62929532 |
| ENSG000001! | gene | 0,646494514 | 0,036240752 | 0,280613979 | ZXDA | -0,62928997 |
| ENSG000001‹ | gene | 0,646527309 | 0,007043076 | 0,239194577 | CHST7 | -0,62921679 |
| ENSG000001( | gene | 0,646901037 | 0,043717343 | 0,292071193 | BAG5 | -0,62838307 |
| ENSG000001; | gene | 0,646969417 | 0,030357888 | 0,271804321 | BTG1 | -0,62823058 |
| ENSG000001! | gene | 0,64717106 | 0,033098982 | 0,274068881 | ZNF770 | -0,627781 |
| ENSG000001! | gene | 0,64773624 | 0,002829729 | 0,237725877 | AKAP17A | -0,62652163 |
| ENSG000001( | gene | 0,650210482 | 0,042961113 | 0,290207865 | TCP11L2 | -0,62102128 |
| ENSG000002! | gene | 0,650634204 | 0,015127652 | 0,252327435 | AC010186.3 | -0,62008143 |
| ENSG000002‹ | gene | 0,652125025 | 0,019312197 | 0,254107998 | ACAD11 | -0,61677951 |
| ENSG000000! | gene | 0,652218585 | 0,023703677 | 0,263452051 | SRRT | -0,61657254 |
| ENSG000002! | gene | 0,652835107 | 0,048390697 | 0,297007722 | ATXN7L3B | -0,61520945 |
| ENSG000001; | gene | 0,653475612 | 0,029763266 | 0,271738191 | APPL2 | -0,6137947 |
| ENSG000001' | gene | 0,653974057 | 0,03674066 | 0,280909481 | BNIP1 | -0,61269469 |
| ENSG000001! | gene | 0,653988012 | 0,025594859 | 0,265635455 | IKZF1 | -0,6126639 |
| ENSG000001; | gene | 0,654377257 | 0,027328398 | 0,267343734 | ARGLU1 | -0,61180549 |
| ENSG000001! | gene | 0,65448732 | 0,041452834 | 0,286271363 | SLC25A42 | -0,61156285 |
| ENSG000001; | gene | 0,655364085 | 0,036232862 | 0,280613979 | ASXL1 | -0,60963148 |
| ENSG000001; | gene | 0,655687394 | 0,049436303 | 0,299174578 | TOM1L2 | -0,60891994 |
| ENSG000001! | gene | 0,655971311 | 0,031919363 | 0,272282875 | RSBN1L | -0,60829537 |
| ENSG000001( | gene | 0,656577813 | 0,026885824 | 0,267343734 | DNAJB14 | -0,6069621 |
| ENSG000001; | gene | 0,656801906 | 0,036472635 | 0,280613979 | GABARAPL1 | -0,60646978 |
| ENSG000001( | gene | 0,657150125 | 0,022864857 | 0,260623396 | ABHD15 | -0,60570511 |
| ENSG000001! | gene | 0,657249502 | 0,00496897 | 0,239194577 | ZNF292 | -0,60548695 |
| ENSG000001' | gene | 0,657523678 | 0,002691614 | 0,237725877 | WIPF1 | -0,60488525 |
| ENSG000001; | gene | 0,659569135 | 0,001938902 | 0,2305768 | KLHL28 | -0,60040421 |
| ENSG000001! | gene | 0,660266199 | 0,045208397 | 0,2922856 | BUB3 | -0,5988803 |
| ENSG000001; | gene | 0,660666803 | 0,002242943 | 0,231249696 | KANSL2 | -0,59800524 |
| ENSG000001; | gene | 0,661533318 | 0,016676142 | 0,252327435 | FAM199X | -0,59611427 |
| ENSG000001! | gene | 0,662519053 | 0,047718676 | 0,296879689 | PRXL2C | -0,59396615 |
| ENSG000000! | gene | 0,663101027 | 0,014752843 | 0,252327435 | KDM2B | -0,59269941 |
| ENSG000001( | gene | 0,663193696 | 0,01764775 | 0,252327435 | FCRL3 | -0,5924978 |
| ENSG000001! | gene | 0,663493898 | 0,024914433 | 0,265412724 | SFMBT2 | -0,5918449 |
| ENSG000001( | gene | 0,666333298 | 0,007248777 | 0,239194577 | ANKRD54 | -0,5856841 |
| ENSG000001( | gene | 0,666499344 | 0,020594287 | 0,256090678 | GLOD4 | -0,58532464 |
| ENSG000001! | gene | 0,667420481 | 0,043708882 | 0,292071193 | TWF1 | -0,58333214 |
| ENSG000001' | gene | 0,667668427 | 0,029712668 | 0,271738191 | ORC2 | -0,58279628 |
| ENSG000001! | gene | 0,668556259 | 0,000932648 | 0,224332495 | PAFAH2 | -0,58087913 |
| ENSG000001; | gene | 0,668674545 | 0,002225113 | 0,231249696 | SRPK2 | -0,5806239 |
| ENSG000001( | gene | 0,668705239 | 0,034321177 | 0,275250624 | PCF11 | -0,58055768 |
| ENSG000002; | gene | 0,66968856 | 0,014580266 | 0,252327435 | SYNRG | -0,57843777 |
| ENSG000002( | gene | 0,670444049 | 0,015860827 | 0,252327435 | AL031432.2 | -0,57681115 |
| ENSG000002; | gene | 0,670688694 | 0,005125125 | 0,239194577 | LINC01578 | -0,57628481 |
| ENSG000002; | gene | 0,670879037 | 0,008586172 | 0,242573226 | AL606834.2 | -0,57587543 |
| ENSG000001‹ | gene | 0,671011828 | 0,012702772 | 0,249534675 | GIGYF1 | -0,5755899 |
| ENSG000002( | gene | 0,671668201 | 0,003474095 | 0,239194577 | MIR4690 | -0,57417937 |
| ENSG000001; | gene | 0,672638825 | 0,004548726 | 0,239194577 | UBE2N | -0,57209604 |
| ENSG000001; | gene | 0,673776726 | 0,049282743 | 0,299174578 | SEPTIN7 | -0,5696575 |

| | | | | | | |
|---|---|---|---|---|---|---|
| ENSG000001: | gene | 0,674212476 | 0,044249135 | 0,2922856 | S1PR4 | -0,56872477 |
| ENSG000001 | gene | 0,674706134 | 0,026149165 | 0,266237968 | C16orf91 | -0,56766882 |
| ENSG000001: | gene | 0,67568821 | 0,038241772 | 0,28223387 | HUS1 | -0,56557041 |
| ENSG000000 | gene | 0,678058471 | 0,002723927 | 0,237725877 | FAM76B | -0,56051841 |
| ENSG000001 | gene | 0,678623455 | 0,03673887 | 0,280909481 | XPC | -0,5593168 |
| ENSG000001 | gene | 0,678631423 | 0,000125671 | 0,142524983 | TRIM33 | -0,55929986 |
| ENSG000000 | gene | 0,67923316 | 0,046949248 | 0,295089709 | ATG16L1 | -0,5580212 |
| ENSG000001 | gene | 0,680496306 | 0,019426326 | 0,254107998 | SDE2 | -0,55534076 |
| ENSG000001 | gene | 0,680795608 | 0,000473626 | 0,19726282 | TPRG1L | -0,55470637 |
| ENSG000001: | gene | 0,681971282 | 0,048638442 | 0,297990743 | WWP1 | -0,55221711 |
| ENSG000002 | gene | 0,682352626 | 0,012897001 | 0,24999756 | AC055811.4 | -0,55141061 |
| ENSG000001 | gene | 0,682881413 | 0,018034429 | 0,252327435 | SETBP1 | -0,55029303 |
| ENSG000001 | gene | 0,683713265 | 0,030850605 | 0,271804321 | H3P47 | -0,54853668 |
| ENSG000001 | gene | 0,684170999 | 0,016131491 | 0,252327435 | POFUT2 | -0,54757114 |
| ENSG000002: | gene | 0,684351141 | 0,030771206 | 0,271804321 | NPIPB14P | -0,54719133 |
| ENSG000001 | gene | 0,685275778 | 0,013401014 | 0,252327435 | RBM4 | -0,5452434 |
| ENSG000001( | gene | 0,685828959 | 0,009260155 | 0,242573226 | AKNA | -0,54407927 |
| ENSG000001 | gene | 0,687932508 | 0,011268975 | 0,249534675 | CCDC144B | -0,53966106 |
| ENSG000001 | gene | 0,688827728 | 0,020705359 | 0,256090678 | CBR4 | -0,53778488 |
| ENSG000000 | gene | 0,689243614 | 0,025573561 | 0,265635455 | OTUD5 | -0,5369141 |
| ENSG000001: | gene | 0,689767253 | 0,006849821 | 0,239194577 | CDK9 | -0,53581846 |
| ENSG000001 | gene | 0,68977732 | 0,006601329 | 0,239194577 | TRIM62 | -0,5357974 |
| ENSG000000 | gene | 0,690013876 | 0,03335426 | 0,274926551 | RC3H2 | -0,53530272 |
| ENSG000001 | gene | 0,690039608 | 0,041451408 | 0,286271363 | LANCL1 | -0,53524892 |
| ENSG000002: | gene | 0,690669185 | 0,046526224 | 0,294465415 | AC243919.1 | -0,53393324 |
| ENSG000000 | gene | 0,691634998 | 0,03262024 | 0,274068881 | KIZ | -0,53191722 |
| ENSG000001 | gene | 0,691817261 | 0,04432286 | 0,2922856 | ELK4 | -0,53153708 |
| ENSG000001 | gene | 0,691855356 | 0,020038187 | 0,2559822 | CCDC186 | -0,53145765 |
| ENSG000000 | gene | 0,692616921 | 0,024775403 | 0,264837204 | FAM214A | -0,52987046 |
| ENSG000001 | gene | 0,69318444 | 0,021394116 | 0,256239299 | LEMD3 | -0,52868883 |
| ENSG000001( | gene | 0,693499444 | 0,025633411 | 0,265635455 | TBC1D17 | -0,52803337 |
| ENSG000001: | gene | 0,69512166 | 0,047888223 | 0,296879689 | CHST12 | -0,52466259 |
| ENSG000001: | gene | 0,695605952 | 0,005063947 | 0,239194577 | C2orf49 | -0,52365782 |
| ENSG000001: | gene | 0,696256307 | 0,040924007 | 0,286069252 | RBBP6 | -0,5223096 |
| ENSG000001 | gene | 0,697538406 | 0,0220615 | 0,25813016 | DIP2A | -0,51965544 |
| ENSG000001( | gene | 0,697630932 | 0,013468097 | 0,252327435 | SMURF2 | -0,51946409 |
| ENSG000001 | gene | 0,69804544 | 0,025913889 | 0,266099662 | E4F1 | -0,51860714 |
| ENSG000002 | gene | 0,698216091 | 0,010042639 | 0,242573226 | PCBP2-OT1 | -0,51825449 |
| ENSG000002( | gene | 0,698739267 | 0,008936678 | 0,242573226 | HLA-E | -0,51717388 |
| ENSG000001 | gene | 0,699308517 | 0,025857813 | 0,266059172 | RPRD2 | -0,51599902 |
| ENSG000002 | gene | 0,699707378 | 0,009055376 | 0,242573226 | TRIM59 | -0,51517639 |
| ENSG000001 | gene | 0,69979515 | 0,000182419 | 0,143227097 | TOB2 | -0,51499543 |
| ENSG000001 | gene | 0,701098022 | 0,010212735 | 0,244080246 | SMAD3 | -0,51231193 |
| ENSG000002: | gene | 0,701105949 | 0,007123116 | 0,239194577 | DFFBP1 | -0,51229562 |
| ENSG000001 | gene | 0,702045354 | 0,013047567 | 0,250331798 | CRIPT | -0,51036386 |
| ENSG000001 | gene | 0,702109302 | 0,008406134 | 0,242573226 | SF1 | -0,51023245 |
| ENSG000001 | gene | 0,702461654 | 0,043895093 | 0,292071193 | VPS13A | -0,50950862 |
| ENSG000002 | gene | 0,702959736 | 0,00210726 | 0,231249696 | AL121753.2 | -0,50848604 |
| ENSG000000 | gene | 0,702998931 | 0,017656282 | 0,252327435 | REST | -0,5084056 |
| ENSG000001 | gene | 0,703178929 | 0,020191837 | 0,256090678 | CFL2 | -0,50803625 |
| ENSG000001 | gene | 0,703900728 | 0,049696427 | 0,299174578 | ZBTB40 | -0,50655612 |
| ENSG000002 | gene | 0,704437595 | 0,004225798 | 0,239194577 | PDXDC2P | -0,50545619 |
| ENSG000000 | gene | 0,705681365 | 0,026779106 | 0,267343734 | ZNF324 | -0,50291118 |
| ENSG000001: | gene | 0,70647588 | 8,66159E-05 | 0,129441475 | PRKAA1 | -0,50128779 |
| ENSG000002 | gene | 0,706651027 | 0,036417668 | 0,280613979 | AL049840.6 | -0,50093016 |
| ENSG000001( | gene | 0,708131184 | 0,012652917 | 0,249534675 | CREBRF | -0,49791144 |
| ENSG000002: | gene | 0,708835681 | 0,02916995 | 0,270832371 | ASH1L-AS1 | -0,49647687 |

| | | | | | | |
|---|---|---|---|---|---|---|
| ENSG000001( | gene | 0,709437597 | 0,042254127 | 0,288197095 | MED30 | -0,49525231 |
| ENSG000001! | gene | 0,709482019 | 0,010728566 | 0,246139857 | GXYLT1 | -0,49516197 |
| ENSG000001{ | gene | 0,710026793 | 0,020411772 | 0,256090678 | TMEM222 | -0,49405463 |
| ENSG000001; | gene | 0,710525647 | 0,019750705 | 0,25449173 | PMS2 | -0,49304137 |
| ENSG000001‹ | gene | 0,711268651 | 0,038654625 | 0,28223387 | ESCO1 | -0,49153352 |
| ENSG000002; | gene | 0,711333346 | 0,033762331 | 0,275250624 | PHF20 | -0,4914023 |
| ENSG000001' | gene | 0,7113846 | 0,038624553 | 0,28223387 | KMT2A | -0,49129835 |
| ENSG000001; | gene | 0,711849653 | 0,010463396 | 0,244080246 | PHC3 | -0,49035553 |
| ENSG000001; | gene | 0,712416747 | 0,007653582 | 0,239194577 | UBL3 | -0,48920666 |
| ENSG000001' | gene | 0,712616555 | 0,030664554 | 0,271804321 | HECA | -0,48880209 |
| ENSG000001! | gene | 0,713082003 | 0,030007894 | 0,271804321 | RNF166 | -0,4878601 |
| ENSG000001' | gene | 0,713922407 | 0,021694136 | 0,256881727 | RPF1 | -0,48616081 |
| ENSG000001' | gene | 0,71480311 | 0,022027699 | 0,25813016 | GNPTAB | -0,48438218 |
| ENSG000001( | gene | 0,716073884 | 0,014915277 | 0,252327435 | KIAA1143 | -0,48181964 |
| ENSG000001; | gene | 0,716087916 | 0,044828311 | 0,2922856 | HNRNPA0 | -0,48179137 |
| ENSG000000' | gene | 0,722056667 | 0,04518491 | 0,2922856 | MBTD1 | -0,46981603 |
| ENSG000000( | gene | 0,722146386 | 0,030930454 | 0,271804321 | PKD1 | -0,46963678 |
| ENSG000001' | gene | 0,722958947 | 0,033115029 | 0,274068881 | PAPOLG | -0,46801437 |
| ENSG000001{ | gene | 0,723442372 | 0,033073929 | 0,274068881 | TTC3 | -0,46705 |
| ENSG000001‹ | gene | 0,724407394 | 0,028104106 | 0,268594209 | RETREG3 | -0,46512682 |
| ENSG000001( | gene | 0,725275977 | 0,039214224 | 0,283069015 | UBE2S | -0,46339803 |
| ENSG000001( | gene | 0,725544086 | 0,041510487 | 0,286440934 | OTUD3 | -0,46286482 |
| ENSG000001( | gene | 0,72836382 | 0,010585642 | 0,24500668 | ACTR5 | -0,45726883 |
| ENSG000000! | gene | 0,729157401 | 0,038887174 | 0,282708484 | RFFL | -0,45569782 |
| ENSG000001; | gene | 0,729382414 | 0,003302909 | 0,239097848 | NR2C2 | -0,45525268 |
| ENSG000001; | gene | 0,729562531 | 0,026676717 | 0,267343734 | ASB8 | -0,45489646 |
| ENSG000001; | gene | 0,729701752 | 0,027082998 | 0,267343734 | CHD6 | -0,45462118 |
| ENSG000000( | gene | 0,732650888 | 0,025061731 | 0,265635455 | KIF2A | -0,44880218 |
| ENSG000001! | gene | 0,732943648 | 0,03872968 | 0,282486259 | NEK7 | -0,44822581 |
| ENSG000002{ | gene | 0,734561784 | 0,025356184 | 0,265635455 | AC125232.2 | -0,44504425 |
| ENSG000001; | gene | 0,734638048 | 0,005895984 | 0,239194577 | GCC1 | -0,44489448 |
| ENSG000001; | gene | 0,734756907 | 0,00774523 | 0,239194577 | AMD1 | -0,44466108 |
| ENSG000001( | gene | 0,735138508 | 0,013344923 | 0,252327435 | ACIN1 | -0,443912 |
| ENSG000001‹ | gene | 0,736571739 | 0,02483092 | 0,264837204 | SMAD4 | -0,44110205 |
| ENSG000002‹ | gene | 0,736652482 | 0,04262724 | 0,28929271 | N4BP2L2 | -0,44094391 |
| ENSG000001; | gene | 0,7369467 | 0,026590576 | 0,267343734 | RELA | -0,44036782 |
| ENSG000001( | gene | 0,737366989 | 0,02142565 | 0,256239299 | MON1B | -0,43954527 |
| ENSG000001; | gene | 0,738965921 | 0,039785909 | 0,284088728 | ZNF767P | -0,43642026 |
| ENSG000001( | gene | 0,739581716 | 0,041239317 | 0,286069252 | SNX20 | -0,43521854 |
| ENSG000002( | gene | 0,741604893 | 0,006043821 | 0,239194577 | DYNLL2 | -0,43127733 |
| ENSG000002‹ | gene | 0,743379991 | 0,044148506 | 0,2922856 | CDK11B | -0,42782824 |
| ENSG000001! | gene | 0,743772013 | 0,039437097 | 0,283674733 | STK39 | -0,42706763 |
| ENSG000002{ | gene | 0,744474897 | 0,022516338 | 0,259688433 | AL590064.1 | -0,42570489 |
| ENSG000000( | gene | 0,745206944 | 0,038618638 | 0,28223387 | IP6K2 | -0,42428698 |
| ENSG000001; | gene | 0,745993588 | 0,018559043 | 0,254046692 | KLF16 | -0,42276486 |
| ENSG000001( | gene | 0,749601719 | 0,020509134 | 0,256090678 | SNTB2 | -0,41580383 |
| ENSG000001; | gene | 0,750662459 | 0,009876265 | 0,242573226 | MXD4 | -0,41376376 |
| ENSG000001' | gene | 0,750749334 | 0,045949574 | 0,292991357 | IK | -0,4135968 |
| ENSG000001( | gene | 0,751190345 | 0,049409894 | 0,299174578 | RAMAC | -0,41274957 |
| ENSG000001‹ | gene | 0,752834648 | 0,03462839 | 0,27606048 | MINK1 | -0,40959507 |
| ENSG000001' | gene | 0,753474485 | 0,01249553 | 0,249534675 | FAM172A | -0,40836944 |
| ENSG000001{ | gene | 0,754891242 | 0,021096731 | 0,256239299 | PTAR1 | -0,40565929 |
| ENSG000002; | gene | 0,756952458 | 0,045522323 | 0,292582729 | AP000347.1 | -0,4017254 |
| ENSG000001; | gene | 0,757618793 | 0,031279515 | 0,271804321 | MARCHF7 | -0,40045598 |
| ENSG000001; | gene | 0,758640157 | 0,043204077 | 0,291078555 | DGKZP1 | -0,39851235 |
| ENSG000000( | gene | 0,759059494 | 0,026577942 | 0,267343734 | CDK13 | -0,39771513 |
| ENSG000000! | gene | 0,759349602 | 0,005463777 | 0,239194577 | ATP11B | -0,39716384 |

| | | | | | | |
|---|---|---|---|---|---|---|
| ENSG000001( | gene | 0,759388791 | 0,025300748 | 0,265635455 | CCDC88B | -0,39708939 |
| ENSG000001{ | gene | 0,760020749 | 0,00733501 | 0,239194577 | PRPF39 | -0,39588929 |
| ENSG000001( | gene | 0,760191265 | 0,031780595 | 0,272282875 | DCAF16 | -0,39556565 |
| ENSG000000: | gene | 0,761958423 | 0,031729713 | 0,272282875 | PNKP | -0,39221582 |
| ENSG00002! | gene | 0,763731998 | 0,017945535 | 0,252327435 | GLUD1P3 | -0,38886163 |
| ENSG000000 | gene | 0,764691782 | 0,041189115 | 0,286069252 | CIC | -0,38704972 |
| ENSG000000( | gene | 0,765616106 | 0,018670194 | 0,254046692 | CNN2 | -0,38530691 |
| ENSG000001: | gene | 0,765707076 | 0,012153724 | 0,249534675 | STARD3 | -0,3851355 |
| ENSG000000! | gene | 0,765862529 | 0,016825735 | 0,252327435 | PPP1R12A | -0,38484264 |
| ENSG000001: | gene | 0,766065399 | 0,041255407 | 0,286069252 | PRPF38A | -0,38446053 |
| ENSG000001; | gene | 0,76672219 | 0,017780881 | 0,252327435 | CCNT1 | -0,38322416 |
| ENSG000001 | gene | 0,766779845 | 0,018552318 | 0,254046692 | MEPCE | -0,38311568 |
| ENSG000001( | gene | 0,767210887 | 0,034653822 | 0,27606048 | YY1 | -0,3823049 |
| ENSG000001: | gene | 0,767218381 | 0,025374728 | 0,265635455 | GPS2 | -0,38229081 |
| ENSG000001: | gene | 0,767511896 | 0,006305312 | 0,239194577 | PRPF38B | -0,38173898 |
| ENSG000001: | gene | 0,767922038 | 0,009310195 | 0,242573226 | RNF38 | -0,38096824 |
| ENSG000001: | gene | 0,768371141 | 0,000265201 | 0,180214301 | SP110 | -0,38012476 |
| ENSG000001 | gene | 0,769807097 | 0,013607474 | 0,252327435 | MAN1B1 | -0,37743112 |
| ENSG000001: | gene | 0,770120492 | 0,045188948 | 0,2922856 | DCAF15 | -0,37684391 |
| ENSG000001; | gene | 0,770294978 | 0,027332929 | 0,267343734 | MAU2 | -0,37651708 |
| ENSG000001 | gene | 0,770352403 | 0,031632141 | 0,272282875 | EMB | -0,37640953 |
| ENSG000000 | gene | 0,771102747 | 0,023253395 | 0,262843195 | GTF3C1 | -0,37500499 |
| ENSG000001 | gene | 0,771171567 | 0,036919174 | 0,281022131 | FBXO34 | -0,37487623 |
| ENSG000001! | gene | 0,771644554 | 0,028036951 | 0,268594209 | VPS26B | -0,37399165 |
| ENSG000002 | gene | 0,77326254 | 0,018919447 | 0,254046692 | AL445363.3 | -0,37096977 |
| ENSG000001 | gene | 0,773794953 | 0,021762714 | 0,257387001 | GATAD2B | -0,36997678 |
| ENSG000001! | gene | 0,773958533 | 0,033971613 | 0,275250624 | L3MBTL3 | -0,36967182 |
| ENSG000001 | gene | 0,777467285 | 0,037069304 | 0,2810952 | DCAF1 | -0,36314613 |
| ENSG000001! | gene | 0,777609418 | 0,016862094 | 0,252327435 | PCNX3 | -0,3628824 |
| ENSG000000( | gene | 0,777681661 | 0,019233173 | 0,254107998 | SLK | -0,36274838 |
| ENSG000001 | gene | 0,77906643 | 0,038234946 | 0,28223387 | RALGAPA1 | -0,36018174 |
| ENSG000001: | gene | 0,78094629 | 0,006269694 | 0,239194577 | RBM39 | -0,35670477 |
| ENSG000000( | gene | 0,781208537 | 0,001140855 | 0,224332495 | FBXL3 | -0,35622038 |
| ENSG000001 | gene | 0,781687619 | 0,016882596 | 0,252327435 | PRDM4 | -0,35533591 |
| ENSG000001 | gene | 0,782748288 | 0,038336914 | 0,28223387 | RBM25 | -0,35337965 |
| ENSG000002( | gene | 0,784705351 | 0,008046921 | 0,242573226 | SYS1 | -0,34977706 |
| ENSG000001 | gene | 0,785700004 | 0,032961586 | 0,274068881 | EHD1 | -0,34794953 |
| ENSG000001( | gene | 0,787158239 | 0,016673545 | 0,252327435 | CLASRP | -0,34527441 |
| ENSG000001! | gene | 0,787671826 | 0,004652446 | 0,239194577 | MBIP | -0,34433342 |
| ENSG000001: | gene | 0,790460763 | 0,044318473 | 0,2922856 | TIMM10B | -0,33923424 |
| ENSG000000 | gene | 0,791465127 | 0,010150579 | 0,244080246 | KDM5A | -0,33740231 |
| ENSG000001: | gene | 0,791588662 | 0,011675735 | 0,249534675 | G3BP2 | -0,33717715 |
| ENSG000001 | gene | 0,791856749 | 0,006812082 | 0,239194577 | RPS6KA3 | -0,33668863 |
| ENSG000002 | gene | 0,792703458 | 0,04441621 | 0,2922856 | STAG3L2 | -0,33514682 |
| ENSG000001 | gene | 0,792924595 | 0,005149806 | 0,239194577 | TOMM5 | -0,33474442 |
| ENSG000000( | gene | 0,794275492 | 0,031347809 | 0,272027301 | SFSWAP | -0,33228861 |
| ENSG000001 | gene | 0,794937375 | 0,004150144 | 0,239194577 | VGLL4 | -0,33108689 |
| ENSG000001 | gene | 0,795479773 | 0,018935866 | 0,254046692 | ASB6 | -0,33010285 |
| ENSG000001 | gene | 0,795740942 | 0,018252741 | 0,252327435 | SLC39A6 | -0,32962926 |
| ENSG000001 | gene | 0,797247054 | 0,014785546 | 0,252327435 | C2orf69 | -0,32690123 |
| ENSG000002 | gene | 0,799756681 | 0,012575809 | 0,249534675 | ARFGAP3 | -0,32236696 |
| ENSG000001 | gene | 0,801395173 | 0,049752426 | 0,299174578 | RIPOR2 | -0,31941427 |
| ENSG000001 | gene | 0,80263098 | 0,021040519 | 0,256239299 | PPWD1 | -0,31719125 |
| ENSG000001: | gene | 0,804246395 | 0,039690573 | 0,284051162 | SMARCC2 | -0,31429053 |
| ENSG000001 | gene | 0,804341979 | 0,005928915 | 0,239194577 | CEP104 | -0,31411908 |
| ENSG000002( | gene | 0,805378883 | 0,044735807 | 0,2922856 | GNL1 | -0,31226045 |
| ENSG000000{ | gene | 0,805822197 | 0,027679997 | 0,267527236 | GMCL1 | -0,31146655 |

| | | | | | | |
|---|---|---|---|---|---|---|
| ENSG0000017 | gene | 0,805848221 | 0,022971515 | 0,261238821 | PLEKHF2 | -0,31141996 |
| ENSG0000010 | gene | 0,805986015 | 0,041051572 | 0,286069252 | ACBD5 | -0,31117329 |
| ENSG0000011 | gene | 0,806386896 | 0,02150818 | 0,256239299 | ZNF451 | -0,3104559 |
| ENSG0000016 | gene | 0,807074363 | 0,03291894 | 0,274068881 | ZBTB39 | -0,30922649 |
| ENSG0000014 | gene | 0,809235312 | 0,00993457 | 0,242573226 | BDP1 | -0,30536882 |
| ENSG0000007 | gene | 0,810423658 | 0,016011252 | 0,252327435 | MKRN2 | -0,3032518 |
| ENSG0000001 | gene | 0,810643539 | 0,016028232 | 0,252327435 | RALBP1 | -0,30286043 |
| ENSG0000015 | gene | 0,811183335 | 0,046104251 | 0,293382847 | CWC15 | -0,30190008 |
| ENSG0000010 | gene | 0,811559924 | 0,027812121 | 0,26780973 | TFAM | -0,30123047 |
| ENSG0000014 | gene | 0,81170503 | 0,046779804 | 0,295089709 | POLE3 | -0,30097254 |
| ENSG0000013 | gene | 0,812301554 | 0,009524776 | 0,242573226 | KRAS | -0,29991269 |
| ENSG0000011 | gene | 0,813230471 | 0,002675108 | 0,237725877 | ATF2 | -0,29826382 |
| ENSG0000014 | gene | 0,813394838 | 0,002180013 | 0,231249696 | TMF1 | -0,29797226 |
| ENSG0000008 | gene | 0,813549791 | 0,04981237 | 0,299174578 | WBP11 | -0,29769745 |
| ENSG0000001 | gene | 0,814621349 | 0,012830705 | 0,249534675 | ZNF207 | -0,29579847 |
| ENSG0000009 | gene | 0,816038664 | 0,04691179 | 0,295089709 | WAC | -0,29329059 |
| ENSG0000005 | gene | 0,818473036 | 0,047686239 | 0,296879689 | THRAP3 | -0,28899321 |
| ENSG0000027 | gene | 0,819118458 | 0,002470853 | 0,237725877 | POM121C | -0,28785599 |
| ENSG0000014 | gene | 0,820371497 | 0,033864957 | 0,275250624 | NSD3 | -0,28565073 |
| ENSG0000010 | gene | 0,822081317 | 0,031431454 | 0,272112682 | TBCB | -0,28264699 |
| ENSG0000018 | gene | 0,82232538 | 0,017060366 | 0,252327435 | ATP6V0A2 | -0,28221874 |
| ENSG0000019 | gene | 0,823193619 | 0,034157925 | 0,275250624 | ZNF335 | -0,2806963 |
| ENSG0000012 | gene | 0,823387211 | 0,020742775 | 0,256090678 | HERC2 | -0,28035706 |
| ENSG0000012 | gene | 0,82926129 | 0,04847363 | 0,297159366 | PIK3CA | -0,27010135 |
| ENSG0000017 | gene | 0,82995604 | 0,012492129 | 0,249534675 | EPM2AIP1 | -0,26889317 |
| ENSG0000010 | gene | 0,833866888 | 0,007102082 | 0,239194577 | POLR2F | -0,26211099 |
| ENSG0000016 | gene | 0,83441943 | 0,044333658 | 0,2922856 | TRA2A | -0,26115534 |
| ENSG0000013 | gene | 0,834593255 | 0,02099379 | 0,256239299 | SH3BP5 | -0,26085483 |
| ENSG0000011 | gene | 0,835179041 | 0,022851701 | 0,260623396 | UBE4A | -0,25984259 |
| ENSG0000003 | gene | 0,835750866 | 0,038013044 | 0,28223387 | EDC4 | -0,25885515 |
| ENSG0000016 | gene | 0,836128998 | 0,049597351 | 0,299174578 | RAD21 | -0,25820256 |
| ENSG0000011 | gene | 0,838526905 | 0,010421331 | 0,244080246 | SUB1 | -0,25407102 |
| ENSG0000003 | gene | 0,84221418 | 0,030093498 | 0,271804321 | FAM13B | -0,24774093 |
| ENSG0000011 | gene | 0,843236003 | 0,014402399 | 0,252327435 | SENP6 | -0,24599163 |
| ENSG0000007 | gene | 0,843795728 | 0,041970472 | 0,287700327 | MAP2K7 | -0,24503431 |
| ENSG0000018 | gene | 0,844147786 | 0,042937395 | 0,290207865 | SECISBP2 | -0,2444325 |
| ENSG0000011 | gene | 0,845118279 | 0,004401561 | 0,239194577 | RAB14 | -0,24277483 |
| ENSG0000016 | gene | 0,845949467 | 0,048058277 | 0,296879689 | PRKCB | -0,24135661 |
| ENSG0000016 | gene | 0,848128208 | 0,008401637 | 0,242573226 | BRD7 | -0,23764573 |
| ENSG0000025 | gene | 0,849055273 | 0,02945797 | 0,271124889 | INAFM1 | -0,23606962 |
| ENSG0000012 | gene | 0,849790627 | 0,002734442 | 0,237725877 | MMP24OS | -0,23482066 |
| ENSG0000017 | gene | 0,85437441 | 0,019870806 | 0,25449173 | GRK2 | -0,22705966 |
| ENSG0000011 | gene | 0,860312887 | 0,010299975 | 0,244080246 | STK38 | -0,21706665 |
| ENSG0000018 | gene | 0,863689395 | 0,031002091 | 0,271804321 | DGAT1 | -0,21141552 |
| ENSG0000021 | gene | 0,866621627 | 0,02342976 | 0,263452051 | SCAF8 | -0,20652585 |
| ENSG0000013 | gene | 0,868456446 | 0,016431794 | 0,252327435 | DSTYK | -0,2034746 |
| ENSG0000016 | gene | 0,870737738 | 0,031389341 | 0,272027301 | GOLGA2 | -0,19968984 |
| ENSG0000011 | gene | 0,872446784 | 0,025241026 | 0,265635455 | PPP6R3 | -0,19686096 |
| ENSG0000016 | gene | 0,87287375 | 0,03411288 | 0,275250624 | MAML1 | -0,19615509 |
| ENSG0000011 | gene | 0,87378478 | 0,047960437 | 0,296879689 | MADD | -0,19465012 |
| ENSG0000018 | gene | 0,876101421 | 0,035940038 | 0,279532435 | KPNA4 | -0,1908302 |
| ENSG0000019 | gene | 0,88298104 | 0,017349582 | 0,252327435 | PHF2 | -0,17954564 |
| ENSG0000012 | gene | 0,883600487 | 0,045174858 | 0,2922856 | ARPP19 | -0,17853388 |
| ENSG0000015 | gene | 0,88434535 | 0,046459262 | 0,294465415 | UBR1 | -0,17731822 |
| ENSG0000020 | gene | 0,886813073 | 0,004275221 | 0,239194577 | PRRC2A | -0,17329806 |
| ENSG0000011 | gene | 0,88684728 | 0,023584636 | 0,263452051 | EXOC2 | -0,17324241 |
| ENSG0000007 | gene | 0,892693357 | 0,009176123 | 0,242573226 | CHFR | -0,1637634 |

| | | | | | | |
|---|---|---|---|---|---|---|
| ENSG000001 | gene | 0,893003967 | 0,001257482 | 0,224332495 | SCAF11 | -0,16326151 |
| ENSG000001 | gene | 0,894268573 | 0,011718782 | 0,249534675 | EPC2 | -0,16121992 |
| ENSG000000 | gene | 0,899715233 | 0,016816388 | 0,252327435 | TNPO1 | -0,15245965 |
| ENSG000001 | gene | 0,951921401 | 0,045399504 | 0,2922856 | CNOT6 | -0,07108564 |
| ENSG000001 | gene | 1,049051048 | 0,028598437 | 0,269541526 | ZBTB2 | 0,069084882 |
| ENSG000002 | gene | 1,059756592 | 0,042951886 | 0,290207865 | LSM14A | 0,083732941 |
| ENSG000001 | gene | 1,06345326 | 0,005575731 | 0,239194577 | NR1H2 | 0,088756627 |
| ENSG000001 | gene | 1,087020161 | 0,021475761 | 0,256239299 | MAN2A1 | 0,120378699 |
| ENSG000001 | gene | 1,115660806 | 0,025201319 | 0,265635455 | EWSR1 | 0,157898471 |
| ENSG000002 | gene | 1,1250967 | 0,005768439 | 0,239194577 | ATF6B | 0,170049004 |
| ENSG000001 | gene | 1,126282602 | 0,04509354 | 0,2922856 | APH1A | 0,171568867 |
| ENSG000001 | gene | 1,127177587 | 0,008837066 | 0,242573226 | STK11IP | 0,17271483 |
| ENSG000002 | gene | 1,132939597 | 0,036622633 | 0,280909481 | DDOST | 0,180070946 |
| ENSG000001 | gene | 1,137959231 | 0,04100051 | 0,286069252 | VPS29 | 0,186448872 |
| ENSG000001 | gene | 1,142220479 | 0,022047887 | 0,25813016 | USP25 | 0,191841156 |
| ENSG000001 | gene | 1,156925384 | 0,007351588 | 0,239194577 | PDIA3 | 0,21029582 |
| ENSG000001 | gene | 1,157379782 | 0,00291942 | 0,237725877 | ELMSAN1 | 0,210862348 |
| ENSG000001 | gene | 1,158091726 | 0,000651697 | 0,205762815 | GLE1 | 0,211749526 |
| ENSG000001 | gene | 1,165670317 | 0,006715862 | 0,239194577 | PPP1CA | 0,221159813 |
| ENSG000001 | gene | 1,168911849 | 0,038963104 | 0,282785762 | DUS1L | 0,225166136 |
| ENSG000001 | gene | 1,170507369 | 0,025316627 | 0,265635455 | EIF4H | 0,227134017 |
| ENSG000001 | gene | 1,181419327 | 0,045390185 | 0,2922856 | SERPINB6 | 0,240521119 |
| ENSG000001 | gene | 1,188116159 | 0,046807124 | 0,295089709 | GNPAT | 0,248675892 |
| ENSG000001 | gene | 1,189280465 | 0,038453876 | 0,28223387 | LIMD1 | 0,250088982 |
| ENSG000001 | gene | 1,193794366 | 0,019468018 | 0,254107998 | SRPRA | 0,25555435 |
| ENSG000000 | gene | 1,204323268 | 0,018607591 | 0,254046692 | SPAG7 | 0,268222696 |
| ENSG000001 | gene | 1,206387129 | 0,034244123 | 0,275250624 | PARVG | 0,270692941 |
| ENSG000001 | gene | 1,209034689 | 0,035051694 | 0,277400936 | GLRX3 | 0,273855639 |
| ENSG000000 | gene | 1,210630269 | 0,015360318 | 0,252327435 | UBE3C | 0,275758328 |
| ENSG000000 | gene | 1,211002613 | 0,021822735 | 0,257387001 | GNPTG | 0,276201978 |
| ENSG000001 | gene | 1,211312982 | 0,028702609 | 0,270016158 | RNF7 | 0,27657168 |
| ENSG000001 | gene | 1,212637064 | 0,031276539 | 0,271804321 | TANGO2 | 0,278147824 |
| ENSG000001 | gene | 1,214808126 | 0,030364413 | 0,271804321 | SLC4A2 | 0,280728464 |
| ENSG000001 | gene | 1,217419518 | 0,035693217 | 0,278745727 | INTS4 | 0,283826401 |
| ENSG000001 | gene | 1,220507462 | 0,021293843 | 0,256239299 | NOP9 | 0,287481116 |
| ENSG000001 | gene | 1,22419078 | 0,015009776 | 0,252327435 | RRAGA | 0,291828408 |
| ENSG000001 | gene | 1,225801315 | 0,006488745 | 0,239194577 | YARS1 | 0,293725158 |
| ENSG000001 | gene | 1,225870447 | 0,014705409 | 0,252327435 | DDI2 | 0,293806519 |
| ENSG000000 | gene | 1,22622036 | 0,043679397 | 0,292071193 | ATRN | 0,294218264 |
| ENSG000001 | gene | 1,232790907 | 0,014168995 | 0,252327435 | SLC39A7 | 0,301928125 |
| ENSG000001 | gene | 1,236081519 | 0,024704688 | 0,264837204 | PPFIA1 | 0,305773892 |
| ENSG000001 | gene | 1,23739447 | 0,039306195 | 0,283132205 | UBA1 | 0,307305492 |
| ENSG000002 | gene | 1,2399234 | 0,00081603 | 0,208351114 | AC090589.3 | 0,310250997 |
| ENSG000001 | gene | 1,243186793 | 0,045348021 | 0,2922856 | CLP1 | 0,314043083 |
| ENSG000001 | gene | 1,243724896 | 0,033482222 | 0,275163475 | HTATIP2 | 0,314667405 |
| ENSG000000 | gene | 1,246982607 | 0,016968136 | 0,252327435 | M6PR | 0,318441342 |
| ENSG000001 | gene | 1,249173869 | 0,007083229 | 0,239194577 | SURF1 | 0,320974295 |
| ENSG000001 | gene | 1,249666674 | 0,004524975 | 0,239194577 | POLR2G | 0,321543333 |
| ENSG000001 | gene | 1,25184377 | 0,035741632 | 0,278910423 | DDX21 | 0,324054525 |
| ENSG000000 | gene | 1,252385665 | 0,031475078 | 0,272259424 | TTC17 | 0,324678901 |
| ENSG000001 | gene | 1,253420798 | 0,009681519 | 0,242573226 | SLC25A11 | 0,325870837 |
| ENSG000001 | gene | 1,254735158 | 0,029067527 | 0,270832371 | RAN | 0,327382881 |
| ENSG000001 | gene | 1,261743708 | 0,006664526 | 0,239194577 | WASHC3 | 0,335418893 |
| ENSG000002 | gene | 1,263947828 | 0,047095242 | 0,295089709 | TRIM26 | 0,337936915 |
| ENSG000001 | gene | 1,264565593 | 0,013135471 | 0,251545499 | SMU1 | 0,338641872 |
| ENSG000001 | gene | 1,26513043 | 0,016375686 | 0,252327435 | MAGT1 | 0,339286128 |
| ENSG000001 | gene | 1,266150826 | 0,020276975 | 0,256090678 | DHX9 | 0,340449271 |

| | | | | | | |
|---|---|---|---|---|---|---|
| ENSG000001: | gene | 1,266713855 | 0,049231086 | 0,299174578 | EMC7 | 0,341090663 |
| ENSG000001: | gene | 1,268635242 | 0,029355207 | 0,271124889 | CEPT1 | 0,343277325 |
| ENSG000000 | gene | 1,268663839 | 0,009915542 | 0,242573226 | DNAJA2 | 0,343309846 |
| ENSG000001 | gene | 1,269087935 | 0,034908699 | 0,276640596 | SH3GL1 | 0,343792037 |
| ENSG000001: | gene | 1,269974233 | 0,027811717 | 0,26780973 | BLOC1S1 | 0,344799226 |
| ENSG000001 | gene | 1,274169124 | 0,009206057 | 0,242573226 | HECTD3 | 0,349556783 |
| ENSG000001 | gene | 1,275297194 | 0,040675443 | 0,286069252 | PTDSS1 | 0,350833491 |
| ENSG000002 | gene | 1,276696701 | 0,038922927 | 0,282766059 | MORF4L1P1 | 0,352415831 |
| ENSG000001: | gene | 1,27676786 | 0,045416377 | 0,2922856 | YIPF3 | 0,35249624 |
| ENSG000001 | gene | 1,276939897 | 0,023438674 | 0,263452051 | GFM1 | 0,352690622 |
| ENSG000001 | gene | 1,27717507 | 0,001679905 | 0,225615654 | TMEM115 | 0,352956297 |
| ENSG000002 | gene | 1,279287853 | 0,039082151 | 0,283069015 | IPO7 | 0,355340922 |
| ENSG000000 | gene | 1,282186878 | 0,049952664 | 0,299174578 | GRAMD4 | 0,358606549 |
| ENSG000001 | gene | 1,283299807 | 0,020874639 | 0,256090678 | SRP72 | 0,359858256 |
| ENSG000001 | gene | 1,28368772 | 0,038470801 | 0,28223387 | RCC2 | 0,360294284 |
| ENSG000001 | gene | 1,283898308 | 0,01348619 | 0,252327435 | PEAK1 | 0,360530937 |
| ENSG000001 | gene | 1,284848979 | 8,87715E-05 | 0,129441475 | ATP13A1 | 0,361598795 |
| ENSG000000 | gene | 1,286787052 | 0,00853624 | 0,242573226 | IDH3G | 0,363773325 |
| ENSG000001 | gene | 1,289316383 | 0,047831569 | 0,296879689 | IMPAD1 | 0,366606327 |
| ENSG000001 | gene | 1,293209145 | 0,009574816 | 0,242573226 | TRAK2 | 0,370955615 |
| ENSG000001 | gene | 1,297540788 | 0,018299399 | 0,252408062 | NDUFA6 | 0,37577989 |
| ENSG000000 | gene | 1,302490998 | 0,040605041 | 0,286069252 | CWF19L1 | 0,381273402 |
| ENSG000001 | gene | 1,304399466 | 0,018620648 | 0,254046692 | PLEKHM2 | 0,383385756 |
| ENSG000001 | gene | 1,305447657 | 0,028928312 | 0,270642788 | RSPRY1 | 0,384544613 |
| ENSG000001 | gene | 1,307086982 | 0,040122608 | 0,285786083 | SDHC | 0,38635515 |
| ENSG000001: | gene | 1,307240106 | 0,00775128 | 0,239194577 | ADRM1 | 0,386524151 |
| ENSG000000 | gene | 1,310131368 | 0,042068156 | 0,287987702 | KCNK6 | 0,389711479 |
| ENSG000001 | gene | 1,311812701 | 0,018011266 | 0,252327435 | USP3 | 0,391561748 |
| ENSG000001 | gene | 1,312925123 | 0,003052849 | 0,237725877 | DCTN6 | 0,392784641 |
| ENSG000001 | gene | 1,312953579 | 0,012980024 | 0,250263503 | RMDN1 | 0,392815909 |
| ENSG000000 | gene | 1,313688574 | 0,021229943 | 0,256239299 | ZFR | 0,393623307 |
| ENSG000002 | gene | 1,314594498 | 0,048019245 | 0,296879689 | C6orf47 | 0,394617852 |
| ENSG000001 | gene | 1,315758613 | 0,041126848 | 0,286069252 | TMEM147 | 0,395894839 |
| ENSG000001: | gene | 1,316386843 | 0,010348764 | 0,244080246 | TRAF7 | 0,396583512 |
| ENSG000001 | gene | 1,316475283 | 0,021947303 | 0,258083089 | DMXL1 | 0,396680435 |
| ENSG000001: | gene | 1,317486472 | 0,047772606 | 0,296879689 | TPMT | 0,397788148 |
| ENSG000001 | gene | 1,326581747 | 0,034218747 | 0,275250624 | UNC50 | 0,407713581 |
| ENSG000000 | gene | 1,328770619 | 0,014892328 | 0,252327435 | POLR2E | 0,410092079 |
| ENSG000002 | gene | 1,32906746 | 0,01617232 | 0,252327435 | IKBKE | 0,410414334 |
| ENSG000000 | gene | 1,332475965 | 0,048342751 | 0,296993939 | SRI | 0,41410951 |
| ENSG000001 | gene | 1,338913228 | 0,038562706 | 0,28223387 | AP2A1 | 0,421062466 |
| ENSG000001 | gene | 1,33923785 | 0,015617672 | 0,252327435 | SCAMP2 | 0,421412208 |
| ENSG000001 | gene | 1,341351545 | 0,007271075 | 0,239194577 | PTRHD1 | 0,423687393 |
| ENSG000001: | gene | 1,342211465 | 0,005526963 | 0,239194577 | IAH1 | 0,424611986 |
| ENSG000001 | gene | 1,343161228 | 0,027370844 | 0,267343734 | ARMH3 | 0,425632491 |
| ENSG000001 | gene | 1,343922909 | 0,020294923 | 0,256090678 | LHPP | 0,426450383 |
| ENSG000001: | gene | 1,346962495 | 0,019769415 | 0,25449173 | UBL7 | 0,429709681 |
| ENSG000001 | gene | 1,347102921 | 0,032121601 | 0,272856143 | GUSB | 0,42986008 |
| ENSG000001: | gene | 1,347244337 | 0,001050221 | 0,224332495 | TUBGCP2 | 0,430011523 |
| ENSG000001: | gene | 1,349396022 | 0,002238941 | 0,231249696 | AAR2 | 0,432313814 |
| ENSG000000 | gene | 1,349544617 | 0,011820161 | 0,249534675 | SOAT1 | 0,432472674 |
| ENSG000000 | gene | 1,35074464 | 0,019114054 | 0,254046692 | ZFYVE26 | 0,433754957 |
| ENSG000001 | gene | 1,35244668 | 0,004576263 | 0,239194577 | DYM | 0,435571717 |
| ENSG000000 | gene | 1,352508614 | 0,026304242 | 0,266237968 | CS | 0,435637783 |
| ENSG000001 | gene | 1,35344879 | 0,031817609 | 0,272282875 | MICOS10 | 0,436640302 |
| ENSG000000 | gene | 1,353636195 | 0,001789693 | 0,225855677 | PACC1 | 0,43684005 |
| ENSG000001 | gene | 1,354467955 | 0,038656114 | 0,28223387 | DOK2 | 0,437726261 |

| | | | | | | |
|---|---|---|---|---|---|---|
| ENSG000001 | gene | 1,356276787 | 0,014995616 | 0,252327435 | DDX41 | 0,439651631 |
| ENSG000001 | gene | 1,356650696 | 0,031956378 | 0,272282875 | NDUFV2 | 0,44004931 |
| ENSG000001 | gene | 1,356752064 | 0,034055564 | 0,275250624 | SSNA1 | 0,440157104 |
| ENSG000001 | gene | 1,356895635 | 0,046888011 | 0,295089709 | DESI1 | 0,440309761 |
| ENSG000001 | gene | 1,358284763 | 0,015229713 | 0,252327435 | SELENON | 0,441785971 |
| ENSG000001 | gene | 1,359503347 | 0,041980796 | 0,287700327 | ATF4 | 0,443079704 |
| ENSG000001 | gene | 1,367869984 | 0,025490696 | 0,265635455 | POLK | 0,451931109 |
| ENSG000000 | gene | 1,368350291 | 0,008265121 | 0,242573226 | BAX | 0,4524376 |
| ENSG000001 | gene | 1,370649048 | 0,036940511 | 0,281022131 | GMPPA | 0,45485922 |
| ENSG000001 | gene | 1,371882117 | 0,020708736 | 0,256090678 | FOXN2 | 0,456156519 |
| ENSG000001 | gene | 1,374479513 | 0,012704931 | 0,249534675 | MAP1S | 0,458885403 |
| ENSG000001 | gene | 1,37650452 | 0,027356986 | 0,267343734 | ATPAF2 | 0,461009347 |
| ENSG000001 | gene | 1,37735069 | 0,001530649 | 0,224332495 | ZDHHC5 | 0,461895933 |
| ENSG000001 | gene | 1,379531015 | 0,006206623 | 0,239194577 | SUMO3 | 0,464177892 |
| ENSG000000 | gene | 1,379748392 | 0,002627599 | 0,237725877 | PSMB1 | 0,464405204 |
| ENSG000001 | gene | 1,382334574 | 0,046304415 | 0,294106514 | SNX12 | 0,467106841 |
| ENSG000001 | gene | 1,384081287 | 0,000448344 | 0,19726282 | CALCOCO2 | 0,468928675 |
| ENSG000002 | gene | 1,384116855 | 0,041636329 | 0,286762487 | MRPL33 | 0,468965749 |
| ENSG000001 | gene | 1,384956542 | 0,017178308 | 0,252327435 | UQCRC2 | 0,469840708 |
| ENSG000000 | gene | 1,385605405 | 0,045889256 | 0,292991357 | KIAA0100 | 0,470516463 |
| ENSG000000 | gene | 1,393391612 | 0,04181762 | 0,28750291 | ME2 | 0,478600783 |
| ENSG000001 | gene | 1,396060022 | 0,008618134 | 0,242573226 | STAU1 | 0,48136097 |
| ENSG000001 | gene | 1,398969227 | 0,048693819 | 0,298041903 | AP3M1 | 0,484364228 |
| ENSG000001 | gene | 1,39920646 | 0,01879843 | 0,254046692 | AKR1A1 | 0,484608855 |
| ENSG000001 | gene | 1,399641826 | 0,040637563 | 0,286069252 | PCBD2 | 0,485057683 |
| ENSG000001 | gene | 1,402464619 | 0,005193994 | 0,239194577 | SERPINB9 | 0,487964376 |
| ENSG000001 | gene | 1,40264533 | 0,018261853 | 0,252327435 | PLD2 | 0,488150258 |
| ENSG000000 | gene | 1,403346434 | 0,009992153 | 0,242573226 | MRPS10 | 0,488871201 |
| ENSG000001 | gene | 1,404311902 | 0,006357902 | 0,239194577 | TRAFD1 | 0,489863398 |
| ENSG000001 | gene | 1,405622035 | 0,04817144 | 0,296879689 | IMMT | 0,491208713 |
| ENSG000000 | gene | 1,406596066 | 0,035182522 | 0,277446768 | ARSD | 0,492208088 |
| ENSG000001 | gene | 1,406663716 | 0,012706544 | 0,249534675 | VPS33A | 0,492277472 |
| ENSG000000 | gene | 1,408139379 | 0,018155475 | 0,252327435 | KARS1 | 0,493790141 |
| ENSG000001 | gene | 1,409903919 | 0,031216685 | 0,271804321 | SNX17 | 0,49559685 |
| ENSG000001 | gene | 1,410837708 | 0,019186653 | 0,254107998 | PSMG2 | 0,496552041 |
| ENSG000001 | gene | 1,412732198 | 0,043131822 | 0,290783689 | UBQLN1 | 0,49848801 |
| ENSG000001 | gene | 1,412788094 | 0,025010171 | 0,265635455 | PSMD13 | 0,49854509 |
| ENSG000000 | gene | 1,413657708 | 0,008126526 | 0,242573226 | GDE1 | 0,499432839 |
| ENSG000001 | gene | 1,414469387 | 0,010894781 | 0,246314595 | GTF2F2 | 0,500260953 |
| ENSG000001 | gene | 1,415489403 | 0,016914793 | 0,252327435 | MIEN1 | 0,501300949 |
| ENSG000001 | gene | 1,417702597 | 0,045175385 | 0,2922856 | KLHL18 | 0,503554919 |
| ENSG000001 | gene | 1,419260309 | 0,009076458 | 0,242573226 | TMEM126A | 0,505139221 |
| ENSG000000 | gene | 1,420343478 | 0,003632269 | 0,239194577 | PIGS | 0,506239855 |
| ENSG000001 | gene | 1,421025538 | 0,037264238 | 0,2810952 | GYS1 | 0,506932483 |
| ENSG000001 | gene | 1,421498248 | 0,002980796 | 0,237725877 | NAA38 | 0,50741232 |
| ENSG000001 | gene | 1,423279902 | 0,001844036 | 0,226771947 | ERH | 0,50921941 |
| ENSG000000 | gene | 1,425700439 | 0,045128046 | 0,2922856 | RHOA | 0,511670882 |
| ENSG000001 | gene | 1,427248 | 0,022138542 | 0,25813016 | PPP2CB | 0,513236041 |
| ENSG000001 | gene | 1,428667446 | 0,019360444 | 0,254107998 | MRPS12 | 0,514670137 |
| ENSG000001 | gene | 1,428952083 | 0,001706949 | 0,225855677 | ATXN10 | 0,514957539 |
| ENSG000000 | gene | 1,428977025 | 0,040524212 | 0,286069252 | TBC1D22B | 0,514982721 |
| ENSG000001 | gene | 1,429682768 | 0,017274036 | 0,252327435 | YY1AP1 | 0,515695063 |
| ENSG000001 | gene | 1,429905055 | 0,006142779 | 0,239194577 | CCNJ | 0,515919356 |
| ENSG000001 | gene | 1,429961339 | 0,043413669 | 0,291373337 | CLCN7 | 0,515976142 |
| ENSG000001 | gene | 1,431119254 | 0,018060051 | 0,252327435 | COX6A1 | 0,517143896 |
| ENSG000001 | gene | 1,432745186 | 0,02983995 | 0,271804321 | RABGGTA | 0,518782049 |
| ENSG000001 | gene | 1,435223753 | 0,045753529 | 0,292595946 | CCDC159 | 0,521275672 |

| | | | | | | |
|---|---|---|---|---|---|---|
| ENSG000001[illegible] | gene | 1,435339816 | 0,03393724 | 0,275250624 | ATP5F1A | 0,521392335 |
| ENSG000000[illegible] | gene | 1,435981069 | 0,034555014 | 0,27606048 | NSUN2 | 0,522036729 |
| ENSG000000[illegible] | gene | 1,439123578 | 0,029852508 | 0,271804321 | MRPS18A | 0,525190482 |
| ENSG000000[illegible] | gene | 1,439609964 | 0,023080848 | 0,261281502 | RUFY3 | 0,525677992 |
| ENSG000000[illegible] | gene | 1,441575816 | 0,022269184 | 0,258543709 | RETSAT | 0,527646713 |
| ENSG000002[illegible] | gene | 1,446290215 | 0,005063471 | 0,239194577 | AL022312.1 | 0,532357075 |
| ENSG000001[illegible] | gene | 1,44636077 | 0,042178481 | 0,288197095 | ASPSCR1 | 0,532427453 |
| ENSG000001[illegible] | gene | 1,446381758 | 0,013835443 | 0,252327435 | NOC4L | 0,532448388 |
| ENSG000001[illegible] | gene | 1,446577853 | 0,03961712 | 0,284051162 | ABRACL | 0,532643969 |
| ENSG000002[illegible] | gene | 1,447132377 | 0,046488176 | 0,294465415 | BAG6 | 0,533196899 |
| ENSG000001[illegible] | gene | 1,448802297 | 0,014447125 | 0,252327435 | OGDH | 0,534860739 |
| ENSG000002[illegible] | gene | 1,449290074 | 0,045333683 | 0,2922856 | DPP3 | 0,535346378 |
| ENSG000001[illegible] | gene | 1,449656146 | 0,009128174 | 0,242573226 | ERLIN2 | 0,535710738 |
| ENSG000001[illegible] | gene | 1,451108667 | 0,049973613 | 0,299174578 | STX12 | 0,53715556 |
| ENSG000001[illegible] | gene | 1,45409369 | 0,034673218 | 0,27606048 | RPN2 | 0,540120228 |
| ENSG000001[illegible] | gene | 1,454367193 | 0,030069763 | 0,271804321 | GET3 | 0,540391561 |
| ENSG000001[illegible] | gene | 1,454815896 | 0,008821621 | 0,242573226 | THBS3 | 0,540836595 |
| ENSG000001[illegible] | gene | 1,455387903 | 0,022208847 | 0,258184169 | RPAP1 | 0,541403725 |
| ENSG000001[illegible] | gene | 1,456329501 | 0,004590511 | 0,239194577 | VAC14 | 0,542336809 |
| ENSG000001[illegible] | gene | 1,456810822 | 0,042407486 | 0,288376553 | LAIR1 | 0,542813545 |
| ENSG000001[illegible] | gene | 1,457742276 | 0,030093951 | 0,271804321 | NAA20 | 0,543735679 |
| ENSG000002[illegible] | gene | 1,459150675 | 0,011854126 | 0,249534675 | TWF2 | 0,545128866 |
| ENSG000001[illegible] | gene | 1,460181063 | 0,031939943 | 0,272282875 | EIF4E2 | 0,546147275 |
| ENSG000001[illegible] | gene | 1,463012098 | 0,011595384 | 0,249534675 | MRPS15 | 0,5489417 |
| ENSG000001[illegible] | gene | 1,463389173 | 0,02273769 | 0,260623396 | GMDS | 0,54931349 |
| ENSG000001[illegible] | gene | 1,463447513 | 0,036846043 | 0,281022131 | MRPL47 | 0,549371004 |
| ENSG000000[illegible] | gene | 1,463965231 | 0,008614712 | 0,242573226 | OGFR | 0,54988129 |
| ENSG000001[illegible] | gene | 1,464990584 | 0,002935088 | 0,237725877 | PSMA1 | 0,550891392 |
| ENSG000001[illegible] | gene | 1,465005868 | 0,003995323 | 0,239194577 | TOR3A | 0,550906444 |
| ENSG000001[illegible] | gene | 1,472255777 | 0,038325102 | 0,28223387 | COPS2 | 0,558028334 |
| ENSG000001[illegible] | gene | 1,473093645 | 0,025811728 | 0,266059172 | NDUFS2 | 0,558849145 |
| ENSG000001[illegible] | gene | 1,473601162 | 0,007613983 | 0,239194577 | NDUFB3 | 0,559346105 |
| ENSG000001[illegible] | gene | 1,474277548 | 0,029112946 | 0,270832371 | CD99L2 | 0,560008152 |
| ENSG000001[illegible] | gene | 1,474575815 | 0,022983488 | 0,261238821 | TNFRSF10D | 0,5603 |
| ENSG000001[illegible] | gene | 1,474757869 | 0,037970222 | 0,28223387 | SEC13 | 0,560478108 |
| ENSG000002[illegible] | gene | 1,475238073 | 0,037009574 | 0,281022131 | NME2 | 0,560947795 |
| ENSG000001[illegible] | gene | 1,476041768 | 0,001608152 | 0,224332495 | VKORC1 | 0,561733546 |
| ENSG000001[illegible] | gene | 1,476153742 | 0,004102581 | 0,239194577 | KCTD5 | 0,561842986 |
| ENSG000001[illegible] | gene | 1,477209027 | 0,039168153 | 0,283069015 | SMS | 0,562873984 |
| ENSG000001[illegible] | gene | 1,477862466 | 0,002573862 | 0,237725877 | GHITM | 0,563512015 |
| ENSG000001[illegible] | gene | 1,477877874 | 0,002986586 | 0,237725877 | POP7 | 0,563527055 |
| ENSG000001[illegible] | gene | 1,478898904 | 0,015329832 | 0,252327435 | MRPL16 | 0,564523434 |
| ENSG000001[illegible] | gene | 1,480082176 | 0,023684822 | 0,263452051 | AHCYL1 | 0,565677278 |
| ENSG000001[illegible] | gene | 1,484757459 | 0,002633691 | 0,237725877 | PCCB | 0,57022728 |
| ENSG000001[illegible] | gene | 1,48504582 | 0,048359354 | 0,296993939 | LRPAP1 | 0,570507445 |
| ENSG000001[illegible] | gene | 1,486894069 | 0,000483603 | 0,19726282 | TMEM179B | 0,572301869 |
| ENSG000001[illegible] | gene | 1,486993894 | 0,00778021 | 0,239194577 | UBAP1 | 0,572398724 |
| ENSG000001[illegible] | gene | 1,487704797 | 0,005680203 | 0,239194577 | AP1S1 | 0,573088283 |
| ENSG000001[illegible] | gene | 1,48803424 | 0,014127633 | 0,252327435 | CHID1 | 0,573407723 |
| ENSG000002[illegible] | gene | 1,48849099 | 0,022153622 | 0,25813016 | LTB4R | 0,573850489 |
| ENSG000002[illegible] | gene | 1,488937736 | 0,009741646 | 0,242573226 | HEXA | 0,574283425 |
| ENSG000001[illegible] | gene | 1,489611628 | 0,030010718 | 0,271804321 | PTPA | 0,57493624 |
| ENSG000001[illegible] | gene | 1,491488544 | 0,042214307 | 0,288197095 | FKBP1C | 0,576752897 |
| ENSG000000[illegible] | gene | 1,491490398 | 0,003425862 | 0,239194577 | COASY | 0,57675469 |
| ENSG000001[illegible] | gene | 1,49336619 | 0,000218698 | 0,159446371 | MRPL21 | 0,578567974 |
| ENSG000001[illegible] | gene | 1,494746714 | 0,031278084 | 0,271804321 | SLC9A9 | 0,57990104 |
| ENSG000001[illegible] | gene | 1,494998838 | 0,041924376 | 0,287700327 | PMVK | 0,580144363 |

| | | | | | | |
|---|---|---|---|---|---|---|
| ENSG000001! | gene | 1,495179218 | 0,015718203 | 0,252327435 | CMTM7 | 0,580318422 |
| ENSG000001 | gene | 1,495624911 | 0,026009972 | 0,266237968 | DOP1B | 0,580748405 |
| ENSG000001 | gene | 1,495947702 | 0,007434816 | 0,239194577 | ATG101 | 0,58105974 |
| ENSG000001( | gene | 1,496943969 | 0,020760335 | 0,256090678 | SEC61B | 0,582020222 |
| ENSG000001 | gene | 1,49736278 | 0,012065117 | 0,249534675 | SNX19 | 0,582423799 |
| ENSG000001( | gene | 1,499913448 | 0,049650462 | 0,299174578 | NIPAL2 | 0,584879253 |
| ENSG000001 | gene | 1,499918123 | 0,01806631 | 0,252327435 | THG1L | 0,58488375 |
| ENSG000000 | gene | 1,500548003 | 0,004883406 | 0,239194577 | CLPTM1L | 0,585489472 |
| ENSG000001 | gene | 1,502412521 | 0,003556761 | 0,239194577 | RFX5 | 0,587280992 |
| ENSG000000 | gene | 1,502517988 | 0,01873496 | 0,254046692 | TMEM131 | 0,587382263 |
| ENSG000001 | gene | 1,502759864 | 0,042662147 | 0,289337232 | SIL1 | 0,587614489 |
| ENSG000001 | gene | 1,50289323 | 0,004561369 | 0,239194577 | PSMD2 | 0,58774252 |
| ENSG000000! | gene | 1,503676718 | 0,009622432 | 0,242573226 | SEC61A1 | 0,588494429 |
| ENSG000002( | gene | 1,504584185 | 0,024304254 | 0,264629928 | AC022167.3 | 0,589364831 |
| ENSG000001( | gene | 1,504751821 | 0,031745398 | 0,272282875 | AHCY | 0,589525562 |
| ENSG000001( | gene | 1,505137228 | 0,013192838 | 0,252112392 | PSMA7 | 0,589895028 |
| ENSG000001( | gene | 1,505773286 | 0,017556387 | 0,252327435 | PSMD11 | 0,59050457 |
| ENSG000001 | gene | 1,507722224 | 0,000831965 | 0,208351114 | NDUFS6 | 0,592370657 |
| ENSG000001( | gene | 1,507989945 | 0,028576909 | 0,269541526 | CAPN2 | 0,592626809 |
| ENSG000001 | gene | 1,50882775 | 0,019353013 | 0,254107998 | PSMA5 | 0,593428115 |
| ENSG000001 | gene | 1,509998498 | 0,002928633 | 0,237725877 | S100Z | 0,594547114 |
| ENSG000001 | gene | 1,510564274 | 0,040468209 | 0,286069252 | JAGN1 | 0,595087572 |
| ENSG000001 | gene | 1,510772558 | 0,016251678 | 0,252327435 | HCFC1 | 0,595286484 |
| ENSG000002 | gene | 1,511583554 | 0,017284457 | 0,252327435 | UQCRFS1P1 | 0,596060727 |
| ENSG000001 | gene | 1,514046137 | 0,015268451 | 0,252327435 | GNL2 | 0,598409169 |
| ENSG000001( | gene | 1,517761011 | 0,022380516 | 0,258763174 | ECH1 | 0,601944639 |
| ENSG000001( | gene | 1,517969863 | 0,024808299 | 0,264837204 | KDELR1 | 0,602143149 |
| ENSG000001! | gene | 1,521489273 | 0,03353996 | 0,275245975 | GTF3C6 | 0,605484163 |
| ENSG000001( | gene | 1,521733405 | 0,023977904 | 0,26347532 | ZMAT5 | 0,605715633 |
| ENSG000001( | gene | 1,52229647 | 0,027288595 | 0,267343734 | WASHC5 | 0,606249354 |
| ENSG000000 | gene | 1,522725575 | 0,025302712 | 0,265635455 | SNX1 | 0,606655963 |
| ENSG000001! | gene | 1,523066691 | 0,004417543 | 0,239194577 | GART | 0,606979115 |
| ENSG000001( | gene | 1,524542353 | 0,00176494 | 0,225855677 | HPRT1 | 0,60837623 |
| ENSG000001( | gene | 1,52651476 | 0,034309137 | 0,275250624 | PHKB | 0,610241539 |
| ENSG000001 | gene | 1,527295322 | 0,008136381 | 0,242573226 | TMEM60 | 0,610979052 |
| ENSG000000( | gene | 1,528135896 | 0,001786592 | 0,225855677 | SLC25A5 | 0,611772847 |
| ENSG000001 | gene | 1,528231514 | 0,027659061 | 0,267527236 | UCHL3 | 0,611863116 |
| ENSG000001( | gene | 1,528741273 | 0,008120348 | 0,242573226 | SUCLG1 | 0,612344263 |
| ENSG000001( | gene | 1,529652518 | 0,025297874 | 0,265635455 | DDB1 | 0,613203962 |
| ENSG000001 | gene | 1,530913748 | 0,041131497 | 0,286069252 | TMEM11 | 0,614393003 |
| ENSG000001! | gene | 1,532457355 | 0,03332517 | 0,274926551 | ATP5PF | 0,615846928 |
| ENSG000000! | gene | 1,533092611 | 0,018705763 | 0,254046692 | PRDM1 | 0,61644485 |
| ENSG000000( | gene | 1,533098667 | 0,009801336 | 0,242573226 | RALA | 0,616450548 |
| ENSG000001 | gene | 1,533521823 | 0,031394941 | 0,272027301 | YIF1A | 0,616848697 |
| ENSG000001( | gene | 1,533622999 | 0,02025841 | 0,256090678 | UBE2L3 | 0,616943878 |
| ENSG000001( | gene | 1,53381992 | 0,038523128 | 0,28223387 | TNFSF8 | 0,617129112 |
| ENSG000001 | gene | 1,533937442 | 0,00179233 | 0,225855677 | ANXA7 | 0,617239647 |
| ENSG000000 | gene | 1,5350299 | 0,004951489 | 0,239194577 | MRPL28 | 0,618266757 |
| ENSG000001 | gene | 1,535637722 | 0,018753258 | 0,254046692 | COMMD1 | 0,618837904 |
| ENSG000001 | gene | 1,536026541 | 0,032923928 | 0,274068881 | OS9 | 0,619203145 |
| ENSG000000 | gene | 1,536176114 | 0,003471203 | 0,239194577 | SNX29 | 0,619343623 |
| ENSG000001! | gene | 1,536434729 | 0,010829276 | 0,246139857 | COPG2 | 0,619586479 |
| ENSG000000 | gene | 1,537214162 | 0,019545806 | 0,25449173 | COX15 | 0,620318173 |
| ENSG000001( | gene | 1,538970807 | 0,035953878 | 0,279532435 | STING1 | 0,621965865 |
| ENSG000001 | gene | 1,541054776 | 0,049488463 | 0,299174578 | PSMD14 | 0,623918142 |
| ENSG000001( | gene | 1,541370769 | 0,00504667 | 0,239194577 | MCFD2 | 0,624213936 |
| ENSG000001( | gene | 1,542416395 | 0,030570426 | 0,271804321 | RPS19BP1 | 0,625192292 |

| | | | | | | |
|---|---|---|---|---|---|---|
| ENSG000001 | gene | 1,544052558 | 0,036688335 | 0,280909481 | S100A4 | 0,626721862 |
| ENSG000001 | gene | 1,544065922 | 0,033589935 | 0,275250624 | CHCHD1 | 0,626734348 |
| ENSG000001 | gene | 1,544829659 | 0,003521746 | 0,239194577 | GARS1 | 0,627447768 |
| ENSG000001 | gene | 1,544898869 | 0,005938212 | 0,239194577 | NDUFB10 | 0,627512401 |
| ENSG000001 | gene | 1,547643226 | 0,024608619 | 0,264837204 | TFCP2 | 0,630072929 |
| ENSG000000 | gene | 1,548060536 | 0,015553734 | 0,252327435 | EDEM2 | 0,630461888 |
| ENSG000001 | gene | 1,548790707 | 0,034321108 | 0,275250624 | PIGF | 0,631142202 |
| ENSG000001 | gene | 1,550680116 | 0,043533771 | 0,291567719 | ACP5 | 0,632901109 |
| ENSG000001 | gene | 1,55077345 | 0,019120883 | 0,254046692 | PRDX6 | 0,632987941 |
| ENSG000000 | gene | 1,551408395 | 0,03175942 | 0,272282875 | GDI2 | 0,633578514 |
| ENSG000001 | gene | 1,551496196 | 0,03821509 | 0,28223387 | THOC6 | 0,633660159 |
| ENSG000002 | gene | 1,553629986 | 0,015254881 | 0,252327435 | ARPC1A | 0,635642951 |
| ENSG000001 | gene | 1,55667072 | 0,006370616 | 0,239194577 | TMED9 | 0,638463806 |
| ENSG000001 | gene | 1,561668922 | 0,033142429 | 0,274068881 | DUSP7 | 0,643088631 |
| ENSG000001 | gene | 1,563312514 | 0,001591567 | 0,224332495 | PSMD4 | 0,644606209 |
| ENSG000001 | gene | 1,565348982 | 0,017323471 | 0,252327435 | TMEM258 | 0,64648433 |
| ENSG000001 | gene | 1,565460608 | 0,020461759 | 0,256090678 | ALAD | 0,646587206 |
| ENSG000000 | gene | 1,568759934 | 0,011487019 | 0,249534675 | PSMD8 | 0,649624595 |
| ENSG000001 | gene | 1,569670789 | 0,001460283 | 0,224332495 | SYNE3 | 0,65046201 |
| ENSG000001 | gene | 1,569861801 | 0,037293868 | 0,2810952 | ZNF330 | 0,65063756 |
| ENSG000001 | gene | 1,571385629 | 0,006357428 | 0,239194577 | TOMM22 | 0,652037272 |
| ENSG000001 | gene | 1,572335865 | 0,008770682 | 0,242573226 | PLCB3 | 0,652909423 |
| ENSG000001 | gene | 1,57276669 | 0,047462849 | 0,296846382 | OXR1 | 0,653304672 |
| ENSG000001 | gene | 1,573873391 | 0,033019424 | 0,274068881 | UQCRH | 0,654319489 |
| ENSG000001 | gene | 1,577215089 | 0,035076824 | 0,277400936 | SLX1A | 0,657379417 |
| ENSG000002 | gene | 1,578386038 | 0,011335694 | 0,249534675 | PSMB3 | 0,658450099 |
| ENSG000001 | gene | 1,578565911 | 0,036745683 | 0,280909481 | BECN1 | 0,6586145 |
| ENSG000001 | gene | 1,578690127 | 0,000728154 | 0,205762815 | ETHE1 | 0,65872802 |
| ENSG000001 | gene | 1,580005197 | 0,012150413 | 0,249534675 | SDHB | 0,659929303 |
| ENSG000001 | gene | 1,581053698 | 0,049500544 | 0,299174578 | MRPL40 | 0,660886368 |
| ENSG000000 | gene | 1,583337729 | 0,041173137 | 0,286069252 | ERP44 | 0,662969018 |
| ENSG000001 | gene | 1,58413692 | 0,016251459 | 0,252327435 | COPG1 | 0,663697035 |
| ENSG000001 | gene | 1,585077955 | 0,006840644 | 0,239194577 | TMEM69 | 0,664553795 |
| ENSG000001 | gene | 1,587762296 | 0,012804829 | 0,249534675 | SMIM15 | 0,666994942 |
| ENSG000001 | gene | 1,587825628 | 0,009135566 | 0,242573226 | VAMP8 | 0,667052487 |
| ENSG000001 | gene | 1,588499927 | 0,02559304 | 0,265635455 | LAMTOR2 | 0,667665024 |
| ENSG000002 | gene | 1,588840079 | 0,037371038 | 0,2810952 | GANC | 0,667973921 |
| ENSG000001 | gene | 1,590705972 | 0,044736648 | 0,2922856 | GIMAP8 | 0,669667191 |
| ENSG000001 | gene | 1,590712739 | 0,034162558 | 0,275250624 | SRRD | 0,669673329 |
| ENSG000001 | gene | 1,592398565 | 0,048081366 | 0,296879689 | ATP5F1E | 0,671201476 |
| ENSG000001 | gene | 1,592643546 | 0,028469978 | 0,269245724 | PA2G4 | 0,671423409 |
| ENSG000001 | gene | 1,592885053 | 0,032401985 | 0,273554229 | ACLY | 0,671642162 |
| ENSG000001 | gene | 1,593718706 | 0,040978437 | 0,286069252 | SLC27A3 | 0,672397013 |
| ENSG000001 | gene | 1,594671806 | 0,006934029 | 0,239194577 | SLC2A6 | 0,673259538 |
| ENSG000002 | gene | 1,595461328 | 0,035336973 | 0,2774496 | AC004556.3 | 0,67397364 |
| ENSG000001 | gene | 1,598688805 | 0,021148998 | 0,256239299 | RPN1 | 0,676889136 |
| ENSG000001 | gene | 1,601739441 | 0,017467778 | 0,252327435 | TIMMDC1 | 0,67963948 |
| ENSG000000 | gene | 1,604601225 | 0,006805906 | 0,239194577 | UQCRC1 | 0,682214803 |
| ENSG000000 | gene | 1,605097231 | 0,015281007 | 0,252327435 | ISOC2 | 0,682660693 |
| ENSG000002 | gene | 1,605711459 | 0,015866343 | 0,252327435 | EIF6 | 0,683212669 |
| ENSG000001 | gene | 1,607674297 | 0,01385349 | 0,252327435 | MDH2 | 0,684975157 |
| ENSG000001 | gene | 1,608077942 | 0,026825214 | 0,267343734 | BST2 | 0,685337334 |
| ENSG000000 | gene | 1,613064832 | 0,012386031 | 0,249534675 | PRR11 | 0,689804424 |
| ENSG000001 | gene | 1,613465796 | 0,006576749 | 0,239194577 | UBE2E1 | 0,690162994 |
| ENSG000001 | gene | 1,613511387 | 0,026318616 | 0,266237968 | BABAM2 | 0,690203759 |
| ENSG000001 | gene | 1,614397328 | 0,009123658 | 0,242573226 | PDIA6 | 0,690995692 |
| ENSG000001 | gene | 1,615406045 | 0,00684313 | 0,239194577 | MPV17 | 0,691896844 |

| | | | | | | |
|---|---|---|---|---|---|---|
| ENSG000001 | gene | 1,616179191 | 0,021837674 | 0,257387001 | MRPL36 | 0,692587163 |
| ENSG000000 | gene | 1,616495217 | 0,001292602 | 0,224332495 | HSD17B10 | 0,692869239 |
| ENSG000001 | gene | 1,617825113 | 0,046931991 | 0,295089709 | HM13 | 0,694055661 |
| ENSG000001 | gene | 1,618482809 | 0,001239295 | 0,224332495 | ACAA2 | 0,694642042 |
| ENSG000001 | gene | 1,622897933 | 0,014929967 | 0,252327435 | SNF8 | 0,698572269 |
| ENSG000001 | gene | 1,623383463 | 0,003246323 | 0,238752762 | DHRS4 | 0,699003822 |
| ENSG000001 | gene | 1,624424391 | 0,04235286 | 0,288197095 | KCTD10 | 0,699928595 |
| ENSG000001 | gene | 1,625562189 | 0,039046705 | 0,283060878 | PGAM1 | 0,70093875 |
| ENSG000001 | gene | 1,625758234 | 0,029343346 | 0,271124889 | ETFA | 0,70111273 |
| ENSG000000 | gene | 1,628310787 | 0,02238541 | 0,258763174 | PDK2 | 0,703376086 |
| ENSG000001 | gene | 1,628468812 | 0,045956614 | 0,292991357 | RIC1 | 0,70351609 |
| ENSG000001 | gene | 1,631175333 | 0,002756401 | 0,237725877 | MRPL37 | 0,705911864 |
| ENSG000000 | gene | 1,633099081 | 0,045652488 | 0,292582729 | MPND | 0,707612323 |
| ENSG000002 | gene | 1,633155439 | 0,022315764 | 0,258543709 | CYTOR | 0,707662109 |
| ENSG000001 | gene | 1,635374978 | 0,041903853 | 0,287700327 | HAUS1 | 0,709621472 |
| ENSG000002 | gene | 1,637189494 | 0,000713858 | 0,205762815 | TIMM23 | 0,711221314 |
| ENSG000001 | gene | 1,637596358 | 0,016589023 | 0,252327435 | ATP5MC3 | 0,711579799 |
| ENSG000001 | gene | 1,637902008 | 0,024753026 | 0,264837204 | ACTG1 | 0,711849046 |
| ENSG000001 | gene | 1,641531934 | 0,001159282 | 0,224332495 | COX8A | 0,715042816 |
| ENSG000001 | gene | 1,641565319 | 0,028023222 | 0,268594209 | RHBDF2 | 0,715072157 |
| ENSG000001 | gene | 1,64251483 | 0,005589675 | 0,239194577 | NACC2 | 0,715906396 |
| ENSG000001 | gene | 1,642591912 | 0,028207529 | 0,269039756 | PGK1 | 0,715974099 |
| ENSG000001 | gene | 1,643695637 | 0,004762376 | 0,239194577 | ATP5MPL | 0,71694318 |
| ENSG000001 | gene | 1,644642785 | 0,049967551 | 0,299174578 | STX10 | 0,717774266 |
| ENSG000001 | gene | 1,646978684 | 0,044098189 | 0,2922856 | ARPC3 | 0,719821883 |
| ENSG000001 | gene | 1,647981336 | 0,020852747 | 0,256090678 | SNX3 | 0,720699904 |
| ENSG000001 | gene | 1,648272013 | 0,021795414 | 0,257387001 | TMEM165 | 0,720954349 |
| ENSG000001 | gene | 1,650209258 | 0,01810596 | 0,252327435 | OLA1 | 0,72264898 |
| ENSG000000 | gene | 1,654725165 | 0,007659475 | 0,239194577 | SNX11 | 0,726591619 |
| ENSG000001 | gene | 1,656473915 | 0,032198926 | 0,272856143 | MRPL18 | 0,728115485 |
| ENSG000001 | gene | 1,658130241 | 0,000488701 | 0,19726282 | CYC1 | 0,72955733 |
| ENSG000001 | gene | 1,660917997 | 0,042535285 | 0,288860717 | STYXL1 | 0,731980846 |
| ENSG000001 | gene | 1,661854448 | 0,049918463 | 0,299174578 | ACSS2 | 0,73279403 |
| ENSG000001 | gene | 1,661970682 | 0,010473905 | 0,244080246 | PSMD3 | 0,732894932 |
| ENSG000001 | gene | 1,663719279 | 0,023980462 | 0,26347532 | CCT8 | 0,734412027 |
| ENSG000001 | gene | 1,666578345 | 0,047064385 | 0,295089709 | ELOB | 0,736889139 |
| ENSG000001 | gene | 1,667014321 | 0,020363813 | 0,256090678 | CZIB | 0,737266498 |
| ENSG000001 | gene | 1,667065672 | 0,008904809 | 0,242573226 | MGST3 | 0,737310938 |
| ENSG000002 | gene | 1,671210968 | 0,015785568 | 0,252327435 | EIF4HP1 | 0,740893866 |
| ENSG000000 | gene | 1,672106806 | 0,008453832 | 0,242573226 | TSG101 | 0,741667003 |
| ENSG000000 | gene | 1,673679883 | 0,024352811 | 0,264716875 | COMT | 0,743023617 |
| ENSG000001 | gene | 1,676839103 | 0,018081216 | 0,252327435 | FIBP | 0,745744265 |
| ENSG000002 | gene | 1,677171968 | 0,012733532 | 0,249534675 | TMEM256 | 0,746030622 |
| ENSG000002 | gene | 1,677645671 | 0,032506121 | 0,273980159 | BRK1 | 0,746438042 |
| ENSG000001 | gene | 1,677721037 | 0,018197209 | 0,252327435 | NDUFV3 | 0,746502852 |
| ENSG000001 | gene | 1,680610898 | 0,016470028 | 0,252327435 | RIOK3 | 0,748985744 |
| ENSG000001 | gene | 1,680649391 | 0,001385567 | 0,224332495 | ATP5F1B | 0,749018788 |
| ENSG000001 | gene | 1,68066857 | 0,027516703 | 0,267488561 | PSMA2 | 0,749035251 |
| ENSG000000 | gene | 1,680890163 | 0,007760337 | 0,239194577 | ATP1B3 | 0,749225455 |
| ENSG000001 | gene | 1,683350735 | 0,013006614 | 0,250263503 | COA4 | 0,751335801 |
| ENSG000002 | gene | 1,683529223 | 0,032961623 | 0,274068881 | MPV17L2 | 0,751488764 |
| ENSG000001 | gene | 1,683809083 | 0,007744485 | 0,239194577 | FBP1 | 0,751728569 |
| ENSG000001 | gene | 1,684060654 | 0,02885546 | 0,270349417 | DCAF6 | 0,7519441 |
| ENSG000001 | gene | 1,685862575 | 0,012773338 | 0,249534675 | RBBP8 | 0,753486938 |
| ENSG000000 | gene | 1,686482372 | 0,012676676 | 0,249534675 | POLDIP2 | 0,754017238 |
| ENSG000001 | gene | 1,687618009 | 0,008351597 | 0,242573226 | SUMF1 | 0,754988388 |
| ENSG000001 | gene | 1,689581823 | 0,021403561 | 0,256239299 | ATP6V1D | 0,756666219 |

| | | | | | | |
|---|---|---|---|---|---|---|
| ENSG000001 | gene | 1,691935608 | 0,032073741 | 0,272839834 | AP1S2 | 0,758674663 |
| ENSG000001 | gene | 1,692511531 | 0,025634459 | 0,265635455 | POLR1D | 0,759165663 |
| ENSG000001 | gene | 1,694244381 | 0,017411139 | 0,252327435 | HSF1 | 0,760641987 |
| ENSG000001 | gene | 1,695042911 | 0,004318283 | 0,239194577 | ATP5PD | 0,761321797 |
| ENSG000001 | gene | 1,697135967 | 0,004008806 | 0,239194577 | ORMDL2 | 0,763102152 |
| ENSG000000 | gene | 1,697158083 | 0,016895094 | 0,252327435 | HADHA | 0,763120952 |
| ENSG000001 | gene | 1,698848314 | 0,011778211 | 0,249534675 | HDGF | 0,764557043 |
| ENSG000001 | gene | 1,699878735 | 0,003694418 | 0,239194577 | C14orf119 | 0,765431832 |
| ENSG000000 | gene | 1,70011975 | 0,032940228 | 0,274068881 | GADD45B | 0,765636368 |
| ENSG000001 | gene | 1,701127365 | 0,033139031 | 0,274068881 | COX6B1 | 0,766491161 |
| ENSG000001 | gene | 1,702519052 | 0,019110711 | 0,254046692 | WDR61 | 0,767670943 |
| ENSG000001 | gene | 1,703569655 | 0,015180368 | 0,252327435 | XXYLT1 | 0,768560937 |
| ENSG000001 | gene | 1,704524939 | 0,038575622 | 0,28223387 | NRROS | 0,769369707 |
| ENSG000001 | gene | 1,705210395 | 0,005585845 | 0,239194577 | PREB | 0,769949755 |
| ENSG000002 | gene | 1,706335368 | 0,010049224 | 0,242573226 | AC135048.1 | 0,770901226 |
| ENSG000001 | gene | 1,708577197 | 0,004432577 | 0,239194577 | COMMD5 | 0,772795433 |
| ENSG000001 | gene | 1,710794409 | 0,030868935 | 0,271804321 | ATP5F1C | 0,774666397 |
| ENSG000001 | gene | 1,713769213 | 0,043481906 | 0,291411567 | GABARAP | 0,77717284 |
| ENSG000001 | gene | 1,713777557 | 0,026126995 | 0,266237968 | PALM2AKAP2 | 0,777179864 |
| ENSG000002 | gene | 1,717066403 | 0,019780368 | 0,25449173 | PSMC1P1 | 0,779945833 |
| ENSG000001 | gene | 1,717087182 | 0,034790561 | 0,27613317 | SQLE | 0,779963291 |
| ENSG000001 | gene | 1,717533744 | 0,032434855 | 0,273605422 | ALDH9A1 | 0,780338444 |
| ENSG000001 | gene | 1,720461962 | 0,037236242 | 0,2810952 | CPSF3 | 0,782795995 |
| ENSG000000 | gene | 1,721965984 | 0,019649709 | 0,25449173 | MTHFD2 | 0,784056644 |
| ENSG000001 | gene | 1,722521072 | 0,039712061 | 0,284051162 | MFSD14A | 0,784521632 |
| ENSG000001 | gene | 1,722641924 | 0,012069329 | 0,249534675 | ATP5PB | 0,784622848 |
| ENSG000001 | gene | 1,72803725 | 0,011839061 | 0,249534675 | NDOR1 | 0,789134317 |
| ENSG000001 | gene | 1,728897934 | 0,014779103 | 0,252327435 | HNRNPAB | 0,789852702 |
| ENSG000001 | gene | 1,729116259 | 0,010472596 | 0,244080246 | EMC8 | 0,790034874 |
| ENSG000000 | gene | 1,731392335 | 0,04404164 | 0,2922856 | SNX10 | 0,791932677 |
| ENSG000001 | gene | 1,732269605 | 0,020157414 | 0,256090678 | DECR1 | 0,792663484 |
| ENSG000001 | gene | 1,732394779 | 0,005571465 | 0,239194577 | EIF4G1 | 0,792767729 |
| ENSG000002 | gene | 1,733452888 | 0,007377385 | 0,239194577 | MIR5010 | 0,793648627 |
| ENSG000000 | gene | 1,734068812 | 0,012815778 | 0,249534675 | RNH1 | 0,794161149 |
| ENSG000001 | gene | 1,734900166 | 0,018162029 | 0,252327435 | MGAT1 | 0,794852646 |
| ENSG000000 | gene | 1,736419063 | 0,04426094 | 0,2922856 | MYDGF | 0,796115166 |
| ENSG000001 | gene | 1,736501145 | 0,036996079 | 0,281022131 | CMAS | 0,796183362 |
| ENSG000001 | gene | 1,73705713 | 0,016573254 | 0,252327435 | ALDOA | 0,796645204 |
| ENSG000001 | gene | 1,738208262 | 0,026474425 | 0,267131022 | DBI | 0,797600948 |
| ENSG000001 | gene | 1,739263292 | 0,027769571 | 0,26780973 | MSRB2 | 0,798476346 |
| ENSG000001 | gene | 1,743882512 | 0,035140532 | 0,277400936 | COG5 | 0,802302847 |
| ENSG000001 | gene | 1,744084139 | 0,047274968 | 0,295914294 | RAD23B | 0,802469641 |
| ENSG000001 | gene | 1,744886564 | 0,041148292 | 0,286069252 | TPI1 | 0,803133249 |
| ENSG000000 | gene | 1,749133737 | 0,027063514 | 0,267343734 | EIF2AK1 | 0,806640601 |
| ENSG000001 | gene | 1,749565706 | 0,037190937 | 0,2810952 | CYB5R1 | 0,806996847 |
| ENSG000001 | gene | 1,750990772 | 0,008316185 | 0,242573226 | ACO2 | 0,808171481 |
| ENSG000001 | gene | 1,753223663 | 0,016768601 | 0,252327435 | HIGD2A | 0,810010056 |
| ENSG000000 | gene | 1,754929059 | 0,047021959 | 0,295089709 | PLOD1 | 0,811412712 |
| ENSG000001 | gene | 1,759594884 | 0,02147941 | 0,256239299 | CDC42BPB | 0,815243312 |
| ENSG000001 | gene | 1,76159564 | 0,047960966 | 0,296879689 | RCBTB2 | 0,816882803 |
| ENSG000001 | gene | 1,762264294 | 0,004962714 | 0,239194577 | NDUFAF3 | 0,817430307 |
| ENSG000001 | gene | 1,76696251 | 0,022696547 | 0,260623396 | PEPD | 0,82127143 |
| ENSG000000 | gene | 1,768206526 | 0,02767881 | 0,267527236 | HCCS | 0,822286791 |
| ENSG000001 | gene | 1,770327726 | 0,019439765 | 0,254107998 | CCT5 | 0,824016459 |
| ENSG000001 | gene | 1,771502767 | 0,003245891 | 0,238752762 | CAPRIN1 | 0,824973719 |
| ENSG000002 | gene | 1,771976834 | 0,0016357 | 0,224332495 | AC018521.5 | 0,825359743 |
| ENSG000001 | gene | 1,77472736 | 0,035089301 | 0,277400936 | C16orf70 | 0,82759741 |

| | | | | | | |
|---|---|---|---|---|---|---|
| ENSG00000019 | gene | 1,77530005 | 0,015484872 | 0,252327435 | LTN1 | 0,82806288 |
| ENSG0000013 | gene | 1,777716005 | 0,016383374 | 0,252327435 | STT3A | 0,830024868 |
| ENSG0000014 | gene | 1,780981186 | 0,020781378 | 0,256090678 | C1orf43 | 0,832672276 |
| ENSG0000012 | gene | 1,788906789 | 0,009506316 | 0,242573226 | SEM1 | 0,839078217 |
| ENSG0000017 | gene | 1,790003054 | 0,033693238 | 0,275250624 | GRINA | 0,839962049 |
| ENSG0000010 | gene | 1,791585672 | 0,02614716 | 0,266237968 | RHEB | 0,841237034 |
| ENSG0000014 | gene | 1,79214903 | 0,036768401 | 0,280909481 | DNAJB12 | 0,841690613 |
| ENSG0000026 | gene | 1,792479994 | 0,030753431 | 0,271804321 | AC067852.3 | 0,841957017 |
| ENSG0000010 | gene | 1,792808376 | 0,040369439 | 0,286069252 | SNX8 | 0,842221294 |
| ENSG0000010 | gene | 1,79684934 | 0,005070353 | 0,239194577 | CTSC | 0,845469448 |
| ENSG0000011 | gene | 1,798768744 | 0,028078788 | 0,268594209 | DENND1A | 0,84700972 |
| ENSG0000018 | gene | 1,805070069 | 0,025168543 | 0,265635455 | UQCR10 | 0,852054841 |
| ENSG0000016 | gene | 1,806840123 | 0,044690144 | 0,2922856 | NDUFS4 | 0,853468856 |
| ENSG0000016 | gene | 1,809643223 | 0,030242451 | 0,271804321 | SPTSSA | 0,855705293 |
| ENSG0000004 | gene | 1,81704698 | 0,001645092 | 0,224332495 | PSMA4 | 0,861595721 |
| ENSG0000019 | gene | 1,817140015 | 0,027441263 | 0,267488561 | ANXA4 | 0,861669587 |
| ENSG0000016 | gene | 1,81850356 | 0,015564928 | 0,252327435 | FAM234A | 0,86275175 |
| ENSG0000015 | gene | 1,818697221 | 0,030518673 | 0,271804321 | PEX10 | 0,862905382 |
| ENSG0000011 | gene | 1,827278012 | 0,027278469 | 0,267343734 | IFI27L2 | 0,86969615 |
| ENSG0000003 | gene | 1,832845356 | 0,011411676 | 0,249534675 | BAK1 | 0,874085065 |
| ENSG0000013 | gene | 1,833073271 | 0,023609548 | 0,263452051 | PSMB7 | 0,874264454 |
| ENSG0000012 | gene | 1,833293972 | 0,006342132 | 0,239194577 | ATP6V1F | 0,874438143 |
| ENSG0000015 | gene | 1,833502148 | 0,012907682 | 0,24999756 | ATP5MC1 | 0,874601956 |
| ENSG0000018 | gene | 1,833686751 | 0,019056539 | 0,254046692 | TMEM121B | 0,874747204 |
| ENSG0000026 | gene | 1,83386239 | 0,016998642 | 0,252327435 | AC016876.3 | 0,874885386 |
| ENSG0000016 | gene | 1,834001765 | 0,032076791 | 0,272839834 | RNF181 | 0,874995027 |
| ENSG0000014 | gene | 1,837458788 | 0,040593747 | 0,286069252 | SNRPG | 0,877711892 |
| ENSG0000010 | gene | 1,838328915 | 0,00849844 | 0,242573226 | NUBP1 | 0,878394917 |
| ENSG0000022 | gene | 1,838668359 | 0,001988836 | 0,2305768 | TPI1P1 | 0,878661284 |
| ENSG0000016 | gene | 1,842423495 | 0,044924495 | 0,2922856 | PPFIBP2 | 0,881604714 |
| ENSG0000009 | gene | 1,842731987 | 0,035898499 | 0,279532435 | FH | 0,881846256 |
| ENSG0000017 | gene | 1,843210813 | 0,021277158 | 0,256239299 | SLFN11 | 0,882221085 |
| ENSG0000001 | gene | 1,845067331 | 0,019121842 | 0,254046692 | RNF14 | 0,883673465 |
| ENSG0000011 | gene | 1,847123312 | 0,039670749 | 0,284051162 | ID2 | 0,885280182 |
| ENSG0000027 | gene | 1,853841644 | 0,008310769 | 0,242573226 | HERC2P2 | 0,890518014 |
| ENSG0000021 | gene | 1,855537872 | 0,01573492 | 0,252327435 | NUP62 | 0,891837447 |
| ENSG0000006 | gene | 1,855654946 | 0,01558033 | 0,252327435 | YBX3 | 0,89192847 |
| ENSG0000014 | gene | 1,857318934 | 0,036800322 | 0,280943071 | SLC43A1 | 0,893221572 |
| ENSG0000011 | gene | 1,86146395 | 0,028479247 | 0,269245724 | GNPDA1 | 0,896437677 |
| ENSG0000007 | gene | 1,862259884 | 0,000286599 | 0,180214301 | SCARB1 | 0,897054419 |
| ENSG0000012 | gene | 1,864960342 | 0,003274988 | 0,23877005 | RRAS | 0,899144952 |
| ENSG0000012 | gene | 1,870438154 | 0,01534125 | 0,252327435 | TNFSF10 | 0,903376264 |
| ENSG0000007 | gene | 1,871820383 | 0,022314117 | 0,258543709 | ACTB | 0,904442003 |
| ENSG0000017 | gene | 1,874070442 | 0,030344829 | 0,271804321 | CRTAP | 0,906175182 |
| ENSG0000010 | gene | 1,877140373 | 0,024048553 | 0,263812153 | SRP54 | 0,908536539 |
| ENSG0000018 | gene | 1,878436737 | 0,031199571 | 0,271804321 | FGD6 | 0,909532529 |
| ENSG0000012 | gene | 1,880521945 | 0,04537543 | 0,2922856 | PRDX5 | 0,911133142 |
| ENSG0000010 | gene | 1,8809375 | 0,013532515 | 0,252327435 | EZH2 | 0,911451912 |
| ENSG0000013 | gene | 1,881680441 | 0,040609693 | 0,286069252 | COX7B | 0,912021641 |
| ENSG0000013 | gene | 1,882453798 | 0,032657557 | 0,274068881 | NDUFB5 | 0,912614457 |
| ENSG0000014 | gene | 1,888718821 | 0,048332459 | 0,296993939 | PDSS1 | 0,917407939 |
| ENSG0000027 | gene | 1,889823327 | 0,031762799 | 0,272282875 | C11orf98 | 0,918251368 |
| ENSG0000016 | gene | 1,890111147 | 0,038589417 | 0,28223387 | CSTB | 0,918471074 |
| ENSG0000010 | gene | 1,891139046 | 0,017754841 | 0,252327435 | GALK1 | 0,919255441 |
| ENSG0000013 | gene | 1,892125108 | 0,04324538 | 0,291164637 | CASP1 | 0,920007483 |
| ENSG0000021 | gene | 1,893443019 | 0,012575118 | 0,249534675 | AC078819.1 | 0,921012006 |
| ENSG0000012 | gene | 1,895543659 | 0,040411477 | 0,286069252 | ITFG1 | 0,922611685 |

| | | | | | | |
|---|---|---|---|---|---|---|
| ENSG000002 | gene | 1,897386613 | 0,037759889 | 0,28223387 | IGHA2 | 0,924013673 |
| ENSG000001 | gene | 1,898447422 | 0,022180042 | 0,258143322 | EXOSC4 | 0,924820044 |
| ENSG000001 | gene | 1,898767543 | 0,040373551 | 0,286069252 | CHKA | 0,925063294 |
| ENSG000001 | gene | 1,900031452 | 0,021102553 | 0,256239299 | CIAO2A | 0,9260233 |
| ENSG000001 | gene | 1,902280088 | 0,048837229 | 0,298670817 | AP1B1 | 0,927729681 |
| ENSG000001 | gene | 1,902520502 | 0,013851765 | 0,252327435 | PRMT5 | 0,927912001 |
| ENSG000002 | gene | 1,90450298 | 0,017992408 | 0,252327435 | LINC01637 | 0,929414545 |
| ENSG000001 | gene | 1,90607783 | 0,020296293 | 0,256090678 | PLP2 | 0,930607029 |
| ENSG000001 | gene | 1,908507042 | 0,043104438 | 0,290783689 | OTULINL | 0,932444509 |
| ENSG000002 | gene | 1,910869428 | 0,037580755 | 0,281841856 | LINC02705 | 0,934229201 |
| ENSG000000 | gene | 1,913214629 | 0,044798689 | 0,2922856 | HMGB3 | 0,935998728 |
| ENSG000001 | gene | 1,91354875 | 0,008773509 | 0,242573226 | ABCD1 | 0,936250656 |
| ENSG000001 | gene | 1,917203493 | 0,00746671 | 0,239194577 | DOK1 | 0,939003473 |
| ENSG000002 | gene | 1,920791173 | 0,000489272 | 0,19726282 | ASMTL-AS1 | 0,941700679 |
| ENSG000001 | gene | 1,920844184 | 0,01943327 | 0,254107998 | TNFAIP8L2 | 0,941740495 |
| ENSG000001 | gene | 1,927581673 | 0,00158782 | 0,224332495 | ELOC | 0,94679199 |
| ENSG000001 | gene | 1,933666143 | 0,047067608 | 0,295089709 | SRD5A1 | 0,951338728 |
| ENSG000001 | gene | 1,935299783 | 0,032938725 | 0,274068881 | FKBP15 | 0,952557061 |
| ENSG000001 | gene | 1,936658186 | 0,009360368 | 0,242573226 | NDUFB9 | 0,953569345 |
| ENSG000001 | gene | 1,936740552 | 0,03776278 | 0,28223387 | DUSP3 | 0,953630702 |
| ENSG000001 | gene | 1,94162872 | 0,029790833 | 0,271738191 | NKIRAS2 | 0,957267354 |
| ENSG000001 | gene | 1,942883769 | 0,044576448 | 0,2922856 | DPM2 | 0,958199595 |
| ENSG000001 | gene | 1,944687405 | 0,01264218 | 0,249534675 | MAN2B1 | 0,95953827 |
| ENSG000001 | gene | 1,94505481 | 0,049585719 | 0,299174578 | HDLBP | 0,95981081 |
| ENSG000001 | gene | 1,947489392 | 0,031246624 | 0,271804321 | AC130689.1 | 0,96161547 |
| ENSG000001 | gene | 1,948693425 | 0,014605826 | 0,252327435 | COPS3 | 0,962507139 |
| ENSG000001 | gene | 1,953344444 | 0,010402768 | 0,244080246 | NDUFB1 | 0,96594637 |
| ENSG000001 | gene | 1,953384772 | 0,023872325 | 0,263452051 | TPP1 | 0,965976155 |
| ENSG000001 | gene | 1,956532155 | 0,041828336 | 0,28750291 | SNAPIN | 0,968298821 |
| ENSG000002 | gene | 1,957978983 | 0,028401051 | 0,269163906 | TSPAN4 | 0,969365279 |
| ENSG000002 | gene | 1,959560278 | 0,006056439 | 0,239194577 | PFDN6 | 0,970529952 |
| ENSG000001 | gene | 1,964542936 | 0,012376687 | 0,249534675 | PARL | 0,974193699 |
| ENSG000002 | gene | 1,966974914 | 0,045490226 | 0,292576391 | ATP5MC2P4 | 0,975978558 |
| ENSG000001 | gene | 1,967070764 | 0,042879255 | 0,290207865 | PYCARD | 0,976048859 |
| ENSG000002 | gene | 1,970902853 | 0,020807405 | 0,256090678 | RNA5SP450 | 0,978856667 |
| ENSG000001 | gene | 1,97367022 | 0,021176217 | 0,256239299 | PSMB6 | 0,98088095 |
| ENSG000000 | gene | 1,977430891 | 0,028050513 | 0,268594209 | MAPK6 | 0,983627275 |
| ENSG000001 | gene | 1,978556969 | 0,002900813 | 0,237725877 | MGME1 | 0,984448606 |
| ENSG000002 | gene | 1,979263597 | 0,011987015 | 0,249534675 | AGAP9 | 0,984963763 |
| ENSG000001 | gene | 1,980906091 | 0,039111222 | 0,283069015 | NAPSB | 0,986160488 |
| ENSG000001 | gene | 1,981636615 | 0,010181561 | 0,244080246 | NFE2 | 0,986692431 |
| ENSG000002 | gene | 1,982008555 | 0,036419966 | 0,280613979 | CENPW | 0,98696319 |
| ENSG000001 | gene | 1,984606889 | 0,027214418 | 0,267343734 | RPP30 | 0,988853267 |
| ENSG000001 | gene | 1,985122508 | 0,001122482 | 0,224332495 | COX5A | 0,989228043 |
| ENSG000001 | gene | 1,986258995 | 0,021562111 | 0,256239299 | PDHB | 0,990053753 |
| ENSG000001 | gene | 1,986302634 | 0,000619265 | 0,205762815 | PAPSS1 | 0,99008545 |
| ENSG000002 | gene | 1,998431501 | 0,006706816 | 0,239194577 | CBWD6 | 0,998868123 |
| ENSG000001 | gene | 2,000237484 | 0,017390381 | 0,252327435 | HVCN1 | 1,000171298 |
| ENSG000000 | gene | 2,002157756 | 0,012053897 | 0,249534675 | KCNQ1 | 1,001555653 |
| ENSG000000 | gene | 2,004302885 | 0,027487587 | 0,267488561 | ALAS1 | 1,003100541 |
| ENSG000002 | gene | 2,004343023 | 0,013932739 | 0,252327435 | COA8 | 1,003129433 |
| ENSG000001 | gene | 2,005451344 | 0,015129306 | 0,252327435 | CD300LB | 1,003926964 |
| ENSG000000 | gene | 2,006491173 | 0,028164733 | 0,26892182 | SCD | 1,004674809 |
| ENSG000001 | gene | 2,008625483 | 0,009476696 | 0,242573226 | TM9SF4 | 1,006208592 |
| ENSG000001 | gene | 2,009157996 | 0,029581419 | 0,271738191 | HAVCR2 | 1,006591019 |
| ENSG000001 | gene | 2,011350485 | 0,00544256 | 0,239194577 | COPB2 | 1,008164498 |
| ENSG000001 | gene | 2,02489541 | 0,004661017 | 0,239194577 | DHCR24 | 1,017847392 |

| | | | | | | |
|---|---|---|---|---|---|---|
| ENSG00000( | gene | 2,025643344 | 0,012741136 | 0,249534675 | ADA2 | 1,018380181 |
| ENSG00001( | gene | 2,025749939 | 0,001290145 | 0,224332495 | MTCH2 | 1,018456097 |
| ENSG00001z | gene | 2,026104147 | 0,016314169 | 0,252327435 | UROD | 1,018708334 |
| ENSG00001' | gene | 2,033437718 | 0,017983872 | 0,252327435 | COMMD9 | 1,023920803 |
| ENSG00001{ | gene | 2,034708078 | 0,04825908 | 0,296914065 | UPP1 | 1,024821824 |
| ENSG00001{ | gene | 2,038379424 | 0,015223839 | 0,252327435 | EIF4EBP1 | 1,02742262 |
| ENSG00001! | gene | 2,042630449 | 0,031104569 | 0,271804321 | S100A6 | 1,030428217 |
| ENSG00002( | gene | 2,043878073 | 0,018155009 | 0,252327435 | NDUFA7 | 1,031309136 |
| ENSG00001z | gene | 2,044146239 | 0,033379497 | 0,274926551 | SLC35B1 | 1,031498411 |
| ENSG00001‹ | gene | 2,049694319 | 0,028742374 | 0,27014126 | ATP10D | 1,035408769 |
| ENSG00001( | gene | 2,051605823 | 0,027488988 | 0,267488561 | PGM2 | 1,036753571 |
| ENSG00002( | gene | 2,062046814 | 0,02576811 | 0,266004869 | AC005632.3 | 1,044077086 |
| ENSG00001( | gene | 2,072020609 | 0,017538124 | 0,252327435 | MRPL27 | 1,051038352 |
| ENSG00001‹ | gene | 2,074314208 | 0,005634803 | 0,239194577 | G6PC3 | 1,052634444 |
| ENSG00000( | gene | 2,075146445 | 0,01773846 | 0,252327435 | PKM | 1,053213152 |
| ENSG00001! | gene | 2,075577744 | 5,39566E-05 | 0,129441475 | PPARGC1B | 1,053512972 |
| ENSG00001; | gene | 2,076955871 | 0,038279902 | 0,28223387 | HADHB | 1,054470563 |
| ENSG00002‹ | gene | 2,077736168 | 0,045196781 | 0,2922856 | DBNDD2 | 1,055012472 |
| ENSG00001; | gene | 2,081986411 | 0,002832377 | 0,237725877 | CBWD2 | 1,057960652 |
| ENSG00001; | gene | 2,082321734 | 0,001018428 | 0,224332495 | VPS25 | 1,058192993 |
| ENSG00001( | gene | 2,086696912 | 0,020442579 | 0,256090678 | POLL | 1,061221068 |
| ENSG00001' | gene | 2,093072254 | 0,021142673 | 0,256239299 | COQ5 | 1,065622115 |
| ENSG00001¨ | gene | 2,094541429 | 0,030301883 | 0,271804321 | NUDT4 | 1,06663442 |
| ENSG00002‹ | gene | 2,099020352 | 0,009308864 | 0,242573226 | ATP5MF | 1,069716155 |
| ENSG00001' | gene | 2,104570294 | 0,001285085 | 0,224332495 | MRPL51 | 1,073525698 |
| ENSG00000{ | gene | 2,107535563 | 0,029643473 | 0,271738191 | MEF2C | 1,075556976 |
| ENSG00001z | gene | 2,115998327 | 0,014946966 | 0,252327435 | MRM2 | 1,081338487 |
| ENSG00001( | gene | 2,118407388 | 0,004928197 | 0,239194577 | CDK5 | 1,082980059 |
| ENSG00001‹ | gene | 2,123763805 | 0,042694422 | 0,289363855 | COMMD3 | 1,086623326 |
| ENSG00002( | gene | 2,124559151 | 0,014181014 | 0,252327435 | RNVU1-15 | 1,087163511 |
| ENSG00001; | gene | 2,125807327 | 0,00364375 | 0,239194577 | POMP | 1,088010844 |
| ENSG00001( | gene | 2,126177733 | 0,045164223 | 0,2922856 | AGPAT2 | 1,088262201 |
| ENSG00001( | gene | 2,137948091 | 0,019348284 | 0,254107998 | VPS35L | 1,096226825 |
| ENSG00000‹ | gene | 2,143834835 | 0,01438985 | 0,252327435 | AP2S1 | 1,100193762 |
| ENSG00002( | gene | 2,146538823 | 0,017287774 | 0,252327435 | MFSD2B | 1,102012266 |
| ENSG00001{ | gene | 2,14762661 | 0,042793636 | 0,289843826 | RGS6 | 1,102743186 |
| ENSG00001( | gene | 2,15560654 | 0,020757229 | 0,256090678 | TSPO | 1,108093869 |
| ENSG00001{ | gene | 2,15662088 | 0,038503257 | 0,28223387 | OAF | 1,108772582 |
| ENSG00002( | gene | 2,157752125 | 0,004922373 | 0,239194577 | RNU5D-1 | 1,109529142 |
| ENSG00001‹ | gene | 2,159584592 | 0,006679651 | 0,239194577 | SEC11A | 1,110753829 |
| ENSG00001( | gene | 2,161983548 | 0,019967261 | 0,255395777 | AIMP2 | 1,112355545 |
| ENSG00001! | gene | 2,162854472 | 0,010740545 | 0,246139857 | CBR1 | 1,112936596 |
| ENSG00001z | gene | 2,165262526 | 0,015845504 | 0,252327435 | CLTA | 1,114541954 |
| ENSG00002‹ | gene | 2,166236213 | 0,025774352 | 0,266004869 | AC139495.1 | 1,115190567 |
| ENSG00001¨ | gene | 2,166682769 | 0,008685866 | 0,242573226 | CNP | 1,115487939 |
| ENSG00001( | gene | 2,171595507 | 0,008686023 | 0,242573226 | NFAT5 | 1,118755404 |
| ENSG00001¨ | gene | 2,174669114 | 0,019799803 | 0,25449173 | GPBAR1 | 1,120795905 |
| ENSG00001; | gene | 2,180894501 | 0,030996627 | 0,271804321 | NRM | 1,124919983 |
| ENSG00002( | gene | 2,186028502 | 0,007988938 | 0,242505758 | AC138207.4 | 1,128312212 |
| ENSG00001( | gene | 2,188503048 | 0,00076911 | 0,206587025 | CPED1 | 1,129944394 |
| ENSG00002; | gene | 2,189116576 | 0,017391298 | 0,252327435 | KIFC1 | 1,130348784 |
| ENSG00001‹ | gene | 2,191957952 | 0,028335289 | 0,269111214 | TRIM7 | 1,132220124 |
| ENSG00001; | gene | 2,193651477 | 0,016076451 | 0,252327435 | KCTD3 | 1,133334331 |
| ENSG00001‹ | gene | 2,200241664 | 0,036278007 | 0,280613979 | NOL3 | 1,137661991 |
| ENSG00001( | gene | 2,203401819 | 0,001307797 | 0,224332495 | RCHY1 | 1,139732613 |
| ENSG00001( | gene | 2,203970272 | 0,031731425 | 0,272282875 | E2F1 | 1,140104765 |
| ENSG00001( | gene | 2,207087154 | 0,008744023 | 0,242573226 | URM1 | 1,1421436 |

| | | | | | | |
|---|---|---|---|---|---|---|
| ENSG000001 | gene | 2,215051572 | 0,003168408 | 0,238752762 | SRP19 | 1,147340289 |
| ENSG000002 | gene | 2,215459123 | 0,01224451 | 0,249534675 | VDAC1 | 1,147605708 |
| ENSG000001 | gene | 2,22097818 | 0,014856894 | 0,252327435 | ACVR1B | 1,151195219 |
| ENSG000001 | gene | 2,229128029 | 0,012614491 | 0,249534675 | PRDX1 | 1,15647948 |
| ENSG000001 | gene | 2,230560987 | 0,007776051 | 0,239194577 | CKS2 | 1,157406594 |
| ENSG000001 | gene | 2,231918233 | 0,016228359 | 0,252327435 | LPCAT3 | 1,158284175 |
| ENSG000001 | gene | 2,245501564 | 0,018179724 | 0,252327435 | AC012085.1 | 1,167037727 |
| ENSG000002 | gene | 2,24633529 | 0,047596935 | 0,296879689 | SDHD | 1,167573282 |
| ENSG000001 | gene | 2,249599305 | 0,00291418 | 0,237725877 | TMEM208 | 1,169668054 |
| ENSG000001 | gene | 2,251133829 | 0,025312846 | 0,265635455 | BMF | 1,170651827 |
| ENSG000001 | gene | 2,25174004 | 0,03834404 | 0,28223387 | LGALS1 | 1,17104028 |
| ENSG000000 | gene | 2,252513105 | 0,003642136 | 0,239194577 | CYBRD1 | 1,1715355 |
| ENSG000001 | gene | 2,252992495 | 0,012323421 | 0,249534675 | FADS1 | 1,171842508 |
| ENSG000000 | gene | 2,253225741 | 0,04339299 | 0,291373337 | PDIA5 | 1,171991858 |
| ENSG000001 | gene | 2,253300891 | 0,030973816 | 0,271804321 | BCKDK | 1,172039974 |
| ENSG000001 | gene | 2,257236087 | 0,004772609 | 0,239194577 | LDHA | 1,174557319 |
| ENSG000001 | gene | 2,258799582 | 0,007103596 | 0,239194577 | SLC37A2 | 1,17555627 |
| ENSG000002 | gene | 2,258846838 | 0,046870058 | 0,295089709 | AC090181.1 | 1,175586452 |
| ENSG000001 | gene | 2,259738293 | 0,029097138 | 0,270832371 | RASGEF1B | 1,1761557 |
| ENSG000001 | gene | 2,262398644 | 0,003878805 | 0,239194577 | PSMA6 | 1,177853161 |
| ENSG000001 | gene | 2,263385795 | 0,010719141 | 0,246139857 | MICU1 | 1,178482514 |
| ENSG000001 | gene | 2,270393712 | 0,029766139 | 0,271738191 | QTRT2 | 1,182942499 |
| ENSG000001 | gene | 2,274357589 | 0,049478659 | 0,299174578 | PSAP | 1,185459102 |
| ENSG000002 | gene | 2,275674083 | 0,019126039 | 0,254046692 | AC011195.1 | 1,186293953 |
| ENSG000001 | gene | 2,27775881 | 0,021617512 | 0,2564603 | TMEM106C | 1,187614989 |
| ENSG000001 | gene | 2,281928627 | 0,028983381 | 0,270832371 | FAM89A | 1,190253669 |
| ENSG000002 | gene | 2,285152213 | 0,020855773 | 0,256090678 | SENP3-EIF4A | 1,192290266 |
| ENSG000001 | gene | 2,295809863 | 0,012042737 | 0,249534675 | TMEM106A | 1,199003164 |
| ENSG000000 | gene | 2,297076731 | 0,027028012 | 0,267343734 | ME1 | 1,199799049 |
| ENSG000001 | gene | 2,297467341 | 0,036911193 | 0,281022131 | PCTP | 1,200044354 |
| ENSG000001 | gene | 2,298390914 | 0,011207795 | 0,249534675 | RENBP | 1,200624195 |
| ENSG000000 | gene | 2,308284955 | 0,039304453 | 0,283132205 | ROGDI | 1,206821334 |
| ENSG000001 | gene | 2,309373176 | 0,007421577 | 0,239194577 | HK1 | 1,20750132 |
| ENSG000000 | gene | 2,31092186 | 0,014539503 | 0,252327435 | PHTF2 | 1,208468478 |
| ENSG000000 | gene | 2,314581766 | 0,008006705 | 0,242505758 | MGST2 | 1,210751529 |
| ENSG000001 | gene | 2,321729863 | 0,025906799 | 0,266099662 | SLC46A2 | 1,215200122 |
| ENSG000001 | gene | 2,32226366 | 0,015249724 | 0,252327435 | RPS27L | 1,215531779 |
| ENSG000000 | gene | 2,325033041 | 0,019642326 | 0,25449173 | NANS | 1,217251218 |
| ENSG000001 | gene | 2,32642243 | 0,029740567 | 0,271738191 | GLB1 | 1,218113084 |
| ENSG000000 | gene | 2,327121655 | 0,026008946 | 0,266237968 | RPL26L1 | 1,218546633 |
| ENSG000001 | gene | 2,333948865 | 0,002199954 | 0,231249696 | RUSC2 | 1,222772953 |
| ENSG000001 | gene | 2,336337538 | 0,009215758 | 0,242573226 | LTA4H | 1,22424872 |
| ENSG000001 | gene | 2,340290398 | 0,00780473 | 0,239227871 | TXNRD2 | 1,22668756 |
| ENSG000001 | gene | 2,340416976 | 0,024318886 | 0,264629928 | S100A10 | 1,226765588 |
| ENSG000001 | gene | 2,346656628 | 0,011671492 | 0,249534675 | SLC1A5 | 1,230606756 |
| ENSG000001 | gene | 2,351270729 | 0,02693858 | 0,267343734 | NBR2 | 1,233440663 |
| ENSG000001 | gene | 2,358829853 | 0,016684033 | 0,252327435 | CDCA5 | 1,238071358 |
| ENSG000002 | gene | 2,368152093 | 0,005072414 | 0,239194577 | TAX1BP3 | 1,24376174 |
| ENSG000002 | gene | 2,369933323 | 0,012120616 | 0,249534675 | NEU1 | 1,24484647 |
| ENSG000001 | gene | 2,374474488 | 0,041561578 | 0,286440934 | SLC39A11 | 1,247608256 |
| ENSG000001 | gene | 2,375477016 | 0,009600423 | 0,242573226 | SLC16A1 | 1,248217248 |
| ENSG000002 | gene | 2,376632999 | 0,012624102 | 0,249534675 | RNVU1-6 | 1,248919139 |
| ENSG000001 | gene | 2,378155781 | 0,005749467 | 0,239194577 | UAP1L1 | 1,249843222 |
| ENSG000002 | gene | 2,380151354 | 0,023728811 | 0,263452051 | AL034430.2 | 1,251053317 |
| ENSG000001 | gene | 2,384111311 | 0,012187833 | 0,249534675 | GTF2H1 | 1,253451595 |
| ENSG000001 | gene | 2,384186813 | 0,021360414 | 0,256239299 | ASL | 1,253497283 |
| ENSG000002 | gene | 2,38443685 | 0,010585398 | 0,24500668 | Z93241.1 | 1,253648575 |

| | | | | | | |
|---|---|---|---|---|---|---|
| ENSG000007 | gene | 2,384464653 | 0,009726212 | 0,242573226 | ENO1 | 1,253665397 |
| ENSG000019 | gene | 2,386351758 | 0,007002814 | 0,239194577 | SULT1A1 | 1,254806718 |
| ENSG000023 | gene | 2,389807824 | 0,000502482 | 0,19726282 | LINC02723 | 1,256894609 |
| ENSG000010 | gene | 2,393671319 | 0,043789779 | 0,292071193 | TFEC | 1,259225066 |
| ENSG000012 | gene | 2,395161045 | 0,018355011 | 0,252510181 | CAT | 1,260122663 |
| ENSG000011 | gene | 2,397227049 | 0,043330572 | 0,29135385 | NPC2 | 1,261366558 |
| ENSG000008 | gene | 2,412580755 | 0,023614482 | 0,263452051 | DYNLL1 | 1,270577234 |
| ENSG000001 | gene | 2,414735584 | 0,027704153 | 0,267527236 | TTC27 | 1,271865221 |
| ENSG000010 | gene | 2,419375282 | 0,02548517 | 0,265635455 | DHX35 | 1,274634571 |
| ENSG000014 | gene | 2,42139912 | 0,009862386 | 0,242573226 | NCEH1 | 1,275840899 |
| ENSG000013 | gene | 2,430445432 | 0,045360513 | 0,2922856 | HSD17B4 | 1,281220743 |
| ENSG000016 | gene | 2,432969592 | 0,029418537 | 0,271124889 | TK1 | 1,282718289 |
| ENSG000019 | gene | 2,440958428 | 0,048115252 | 0,296879689 | NUDT16 | 1,287447725 |
| ENSG000013 | gene | 2,443698712 | 0,024549518 | 0,264837204 | SLC43A3 | 1,289066424 |
| ENSG000023 | gene | 2,447935522 | 0,046957789 | 0,295089709 | GPX1 | 1,291565558 |
| ENSG000019 | gene | 2,461312286 | 0,029724595 | 0,271738191 | NAGA | 1,299427715 |
| ENSG000011 | gene | 2,464455246 | 0,03048864 | 0,271804321 | PLA2G4A | 1,301268782 |
| ENSG000013 | gene | 2,46770763 | 0,01504592 | 0,252327435 | MPP1 | 1,303171476 |
| ENSG000028 | gene | 2,471799585 | 0,02942086 | 0,271124889 | AC138207.9 | 1,305561773 |
| ENSG000024 | gene | 2,472679497 | 0,048088293 | 0,296879689 | CEBPA | 1,306075253 |
| ENSG000003 | gene | 2,475992614 | 0,036103774 | 0,280023723 | ATP6V0A1 | 1,308007011 |
| ENSG000010 | gene | 2,490850517 | 0,004268154 | 0,239194577 | LIPA | 1,316638444 |
| ENSG000010 | gene | 2,49113312 | 0,046992036 | 0,295089709 | TIMP1 | 1,316802118 |
| ENSG000014 | gene | 2,493546697 | 0,024666103 | 0,264837204 | GSTO1 | 1,318199221 |
| ENSG000013 | gene | 2,50752931 | 0,000551048 | 0,200876676 | PPT1 | 1,326266565 |
| ENSG000012 | gene | 2,508319865 | 0,006284369 | 0,239194577 | HAT1 | 1,326721335 |
| ENSG000010 | gene | 2,510200778 | 0,044847019 | 0,2922856 | PCK2 | 1,327802762 |
| ENSG000013 | gene | 2,512333258 | 0,034898551 | 0,276640596 | SQOR | 1,329027849 |
| ENSG000023 | gene | 2,512948256 | 0,003735961 | 0,239194577 | LINC00106 | 1,329380965 |
| ENSG000010 | gene | 2,515014465 | 0,005495408 | 0,239194577 | GATA1 | 1,330566698 |
| ENSG000009 | gene | 2,516574927 | 0,031590639 | 0,272282875 | JAK2 | 1,331461552 |
| ENSG000013 | gene | 2,521800413 | 0,03281744 | 0,274068881 | GNS | 1,334454098 |
| ENSG000014 | gene | 2,529469423 | 0,043301619 | 0,291351106 | SLC24A4 | 1,3388348 |
| ENSG000004 | gene | 2,533320793 | 0,03127969 | 0,271804321 | HEXB | 1,341029776 |
| ENSG000026 | gene | 2,533782753 | 0,018120109 | 0,252327435 | MIR4767 | 1,341292833 |
| ENSG000014 | gene | 2,542676928 | 0,021542518 | 0,256239299 | IRF8 | 1,346348165 |
| ENSG000021 | gene | 2,545022124 | 0,030016509 | 0,271804321 | S1PR3 | 1,347678198 |
| ENSG000010 | gene | 2,554557964 | 0,005614624 | 0,239194577 | TMEM205 | 1,353073671 |
| ENSG000017 | gene | 2,55686435 | 0,006127126 | 0,239194577 | NETO2 | 1,354375623 |
| ENSG000010 | gene | 2,558569617 | 0,024522031 | 0,264837204 | KIAA0930 | 1,355337489 |
| ENSG000017 | gene | 2,575821558 | 0,034317251 | 0,275250624 | YWHAG | 1,365032653 |
| ENSG000010 | gene | 2,581908275 | 0,049776381 | 0,299174578 | BLVRA | 1,368437748 |
| ENSG000013 | gene | 2,583097766 | 0,008998657 | 0,242573226 | MRPL15 | 1,369102248 |
| ENSG000000 | gene | 2,583932404 | 0,021426766 | 0,256239299 | FUCA2 | 1,36956833 |
| ENSG000002 | gene | 2,586235794 | 0,016197948 | 0,252327435 | IFNGR1 | 1,370853816 |
| ENSG000009 | gene | 2,597835653 | 0,045562899 | 0,292582729 | MFSD11 | 1,377310164 |
| ENSG000016 | gene | 2,603431234 | 0,012258655 | 0,249534675 | SIGLEC11 | 1,380414301 |
| ENSG000026 | gene | 2,606948397 | 0,018268833 | 0,252327435 | AC025857.2 | 1,382362026 |
| ENSG000018 | gene | 2,610959615 | 0,045693718 | 0,292594592 | UBE2E2 | 1,384580143 |
| ENSG000016 | gene | 2,620597563 | 0,001577522 | 0,224332495 | MT1E | 1,389895821 |
| ENSG000016 | gene | 2,621377677 | 0,032277099 | 0,272856143 | YIF1B | 1,390325226 |
| ENSG000013 | gene | 2,621548347 | 0,012775311 | 0,249534675 | LGALS3 | 1,390419152 |
| ENSG000013 | gene | 2,632139041 | 0,015618228 | 0,252327435 | ST6GALNAC4 | 1,3962357 |
| ENSG000017 | gene | 2,641840313 | 0,012726255 | 0,249534675 | ATOX1 | 1,401543265 |
| ENSG000010 | gene | 2,647641508 | 0,046450046 | 0,294465415 | ERLIN1 | 1,404707794 |
| ENSG000016 | gene | 2,647759442 | 0,005515635 | 0,239194577 | PLK1 | 1,404772054 |
| ENSG000013 | gene | 2,649652909 | 0,010907632 | 0,246314595 | TOP2A | 1,405803386 |

| | | | | | | |
|---|---|---|---|---|---|---|
| ENSG000000 | gene | 2,651267293 | 0,034765226 | 0,27613317 | SMARCD3 | 1,406682126 |
| ENSG000001 | gene | 2,653428743 | 0,03454364 | 0,27606048 | SPINT1 | 1,407857806 |
| ENSG000002 | gene | 2,65478292 | 0,031289319 | 0,271804321 | AP002358.2 | 1,408593897 |
| ENSG000001 | gene | 2,666691989 | 0,024793186 | 0,264837204 | SAPCD2 | 1,415051199 |
| ENSG000001 | gene | 2,701945371 | 0,007554588 | 0,239194577 | SLIRP | 1,433998506 |
| ENSG000001 | gene | 2,708675667 | 0,007908065 | 0,240948122 | PLD3 | 1,437587658 |
| ENSG000002 | gene | 2,714438653 | 0,0074898 | 0,239194577 | RNU5E-4P | 1,440653879 |
| ENSG000002 | gene | 2,723831563 | 0,045580751 | 0,292582729 | AL031284.1 | 1,445637492 |
| ENSG000001 | gene | 2,75631769 | 0,048951593 | 0,299059804 | CFP | 1,462742181 |
| ENSG000001 | gene | 2,774300056 | 0,023089636 | 0,261281502 | FABP5 | 1,472123832 |
| ENSG000001 | gene | 2,785981118 | 0,031884389 | 0,272282875 | P2RX4 | 1,47818548 |
| ENSG000001 | gene | 2,793014235 | 0,01361851 | 0,252327435 | SCIMP | 1,481822926 |
| ENSG000000 | gene | 2,826081301 | 0,002110891 | 0,231249696 | E2F2 | 1,49880297 |
| ENSG000001 | gene | 2,837417297 | 0,029403516 | 0,271124889 | LONRF3 | 1,504578343 |
| ENSG000001 | gene | 2,842627932 | 0,036585144 | 0,280909481 | ANXA2 | 1,507225279 |
| ENSG000001 | gene | 2,850864225 | 0,022821916 | 0,260623396 | TP53I3 | 1,511399331 |
| ENSG000001 | gene | 2,86225439 | 0,044383904 | 0,2922856 | ATP6V1B2 | 1,517151901 |
| ENSG000001 | gene | 2,871836155 | 0,005603162 | 0,239194577 | ADAM15 | 1,521973442 |
| ENSG000001 | gene | 2,878352647 | 0,038746033 | 0,282486259 | SMCO4 | 1,525243357 |
| ENSG000001 | gene | 2,886236869 | 0,03377884 | 0,275250624 | NRG1 | 1,529189704 |
| ENSG000001 | gene | 2,895153531 | 0,014515328 | 0,252327435 | GAPDH | 1,533639857 |
| ENSG000001 | gene | 2,908960379 | 0,0168947 | 0,252327435 | TMTC2 | 1,540503646 |
| ENSG000001 | gene | 2,92094027 | 0,039837998 | 0,284155449 | ASGR1 | 1,546432857 |
| ENSG000001 | gene | 2,921985693 | 0,031223392 | 0,271804321 | GAA | 1,546949114 |
| ENSG000001 | gene | 2,926136812 | 0,007867291 | 0,240423462 | ALDH1A1 | 1,548997224 |
| ENSG000001 | gene | 2,929711111 | 0,049464003 | 0,299174578 | SPTLC2 | 1,550758412 |
| ENSG000000 | gene | 2,944508911 | 0,030711462 | 0,271804321 | LAP3 | 1,558027039 |
| ENSG000001 | gene | 2,950845012 | 0,030207068 | 0,271804321 | MLST8 | 1,561128148 |
| ENSG000001 | gene | 2,954459565 | 0,000496079 | 0,19726282 | UBE2C | 1,562894254 |
| ENSG000002 | gene | 2,967846281 | 0,033161073 | 0,274068881 | CYFIP1 | 1,56941637 |
| ENSG000001 | gene | 3,001688367 | 0,002388009 | 0,237725877 | CCNA2 | 1,585774205 |
| ENSG000002 | gene | 3,007081544 | 0,025628458 | 0,265635455 | MIR6774 | 1,58836399 |
| ENSG000001 | gene | 3,014718047 | 0,004928216 | 0,239194577 | SLITRK4 | 1,59202308 |
| ENSG000001 | gene | 3,016626714 | 0,00750332 | 0,239194577 | ANXA5 | 1,592936184 |
| ENSG000001 | gene | 3,02534096 | 0,033731967 | 0,275250624 | SCPEP1 | 1,597097745 |
| ENSG000001 | gene | 3,03744663 | 0,006717261 | 0,239194577 | CCNB2 | 1,60285906 |
| ENSG000001 | gene | 3,039345649 | 0,047497169 | 0,296879119 | ALCAM | 1,603760754 |
| ENSG000001 | gene | 3,041933285 | 0,036389452 | 0,280613979 | NIBAN2 | 1,604988513 |
| ENSG000001 | gene | 3,052257281 | 0,005585495 | 0,239194577 | CHST13 | 1,609876575 |
| ENSG000002 | gene | 3,056369902 | 0,041559563 | 0,286440934 | AC139495.2 | 1,611819158 |
| ENSG000001 | gene | 3,063677485 | 0,018019836 | 0,252327435 | ARMC10 | 1,615264432 |
| ENSG000001 | gene | 3,103544259 | 0,049763408 | 0,299174578 | DPYD | 1,63391672 |
| ENSG000001 | gene | 3,114247184 | 0,026681132 | 0,267343734 | RPE | 1,638883459 |
| ENSG000001 | gene | 3,15167815 | 0,010838245 | 0,246139857 | ACCS | 1,656120214 |
| ENSG000001 | gene | 3,158504449 | 0,030201103 | 0,271804321 | CD86 | 1,659241605 |
| ENSG000001 | gene | 3,159236516 | 0,009744813 | 0,242573226 | CDC20 | 1,659575948 |
| ENSG000001 | gene | 3,167205132 | 0,02848882 | 0,269245724 | TMEM144 | 1,66321031 |
| ENSG000001 | gene | 3,169802534 | 0,032685748 | 0,274068881 | FGD2 | 1,664392969 |
| ENSG000001 | gene | 3,173346482 | 0,003610878 | 0,239194577 | CREG1 | 1,66600505 |
| ENSG000002 | gene | 3,187411335 | 0,046859093 | 0,295089709 | AC245014.3 | 1,672385211 |
| ENSG000000 | gene | 3,197546834 | 0,030069224 | 0,271804321 | MSR1 | 1,67696549 |
| ENSG000001 | gene | 3,198726215 | 0,007141405 | 0,239194577 | NECTIN2 | 1,677497515 |
| ENSG000002 | gene | 3,220058003 | 0,010319909 | 0,244080246 | MILR1 | 1,687086676 |
| ENSG000000 | gene | 3,234477631 | 0,033927984 | 0,275250624 | OAS1 | 1,693532735 |
| ENSG000002 | gene | 3,296326925 | 0,014609806 | 0,252327435 | FAM198B-AS1 | 1,720859334 |
| ENSG000001 | gene | 3,304063865 | 0,02209204 | 0,25813016 | FXYD6 | 1,724241573 |
| ENSG000001 | gene | 3,306526395 | 0,03068155 | 0,271804321 | CBWD1 | 1,725316418 |

| | | | | | | |
|---|---|---|---|---|---|---|
| ENSG0000019 | gene | 3,313423236 | 7,13696E-05 | 0,129441475 | ZNF808 | 1,728322496 |
| ENSG0000010 | gene | 3,380464495 | 0,028779184 | 0,270238389 | CTSZ | 1,757221495 |
| ENSG0000016 | gene | 3,39090732 | 0,00363739 | 0,239194577 | CTSB | 1,761671353 |
| ENSG0000020 | gene | 3,404066505 | 0,047975434 | 0,296879689 | MRPL53 | 1,767259223 |
| ENSG0000013 | gene | 3,409578854 | 0,028599341 | 0,269541526 | TNS3 | 1,769593551 |
| ENSG0000017 | gene | 3,413124272 | 0,041667018 | 0,286780347 | KCNE3 | 1,771092944 |
| ENSG0000013 | gene | 3,44015707 | 0,015307549 | 0,252327435 | ECHDC3 | 1,782474437 |
| ENSG0000015 | gene | 3,44141632 | 0,049407346 | 0,299174578 | IL18 | 1,78300243 |
| ENSG0000016 | gene | 3,499396296 | 0,005541334 | 0,239194577 | NT5DC2 | 1,807106055 |
| ENSG0000017 | gene | 3,514819116 | 0,021915293 | 0,258003915 | MYD88 | 1,813450445 |
| ENSG0000028 | gene | 3,531661922 | 0,024700665 | 0,264837204 | AL121933.2 | 1,820347244 |
| ENSG0000026 | gene | 3,549330182 | 0,012834888 | 0,249534675 | SULT1A3 | 1,82754679 |
| ENSG0000025 | gene | 3,555620361 | 0,041372521 | 0,286241825 | TMED7-TICAM | 1,830101294 |
| ENSG0000025 | gene | 3,557158621 | 0,024062811 | 0,263812153 | CLEC12B | 1,830725309 |
| ENSG0000004 | gene | 3,565961329 | 0,012790442 | 0,249534675 | HDAC9 | 1,834291058 |
| ENSG0000017 | gene | 3,597613729 | 0,028246171 | 0,269039756 | TYMS | 1,847040295 |
| ENSG0000008 | gene | 3,64704361 | 0,001648372 | 0,224332495 | TPX2 | 1,866727451 |
| ENSG0000013 | gene | 3,650439086 | 0,048219074 | 0,296879689 | LMO2 | 1,868070006 |
| ENSG0000025 | gene | 3,675871247 | 0,043094767 | 0,290783689 | IGKV2-26 | 1,878086235 |
| ENSG0000013 | gene | 3,676382458 | 0,03552943 | 0,277679087 | TUBG1 | 1,87828686 |
| ENSG0000015 | gene | 3,685043972 | 0,002895542 | 0,237725877 | TIRAP | 1,881681834 |
| ENSG0000023 | gene | 3,694345526 | 0,008623717 | 0,242573226 | TXNDC5 | 1,885318806 |
| ENSG0000013 | gene | 3,698580746 | 0,028870468 | 0,270349417 | IDH1 | 1,886971773 |
| ENSG0000009 | gene | 3,712357635 | 0,031215957 | 0,271804321 | TBC1D2 | 1,892335701 |
| ENSG0000014 | gene | 3,721147394 | 0,00039017 | 0,19726282 | TRPM2 | 1,895747537 |
| ENSG0000026 | gene | 3,734630079 | 0,007634468 | 0,239194577 | RN7SL138P | 1,900965349 |
| ENSG0000025 | gene | 3,849214472 | 0,037764001 | 0,28223387 | NAIPP1 | 1,944564058 |
| ENSG0000017 | gene | 3,850162809 | 0,003628298 | 0,239194577 | FAM20C | 1,944919453 |
| ENSG0000013 | gene | 3,864265632 | 0,015040819 | 0,252327435 | TMOD1 | 1,950194269 |
| ENSG0000013 | gene | 3,866936828 | 0,029432883 | 0,271124889 | SORT1 | 1,951191196 |
| ENSG0000021 | gene | 3,867900209 | 0,017909216 | 0,252327435 | IGHA1 | 1,951550574 |
| ENSG0000014 | gene | 3,902896594 | 0,01605466 | 0,252327435 | MKI67 | 1,96454524 |
| ENSG0000008 | gene | 3,928936657 | 0,041013785 | 0,286069252 | FCN1 | 1,974138909 |
| ENSG0000021 | gene | 3,933319011 | 0,048216745 | 0,296879689 | SULT1A4 | 1,9757472 |
| ENSG0000019 | gene | 3,954268993 | 0,00200111 | 0,2305768 | CTNND1 | 1,983411015 |
| ENSG0000016 | gene | 4,006806794 | 0,008912242 | 0,242573226 | C2 | 2,002452946 |
| ENSG0000013 | gene | 4,061699991 | 0,018596281 | 0,254046692 | RAB20 | 2,022083682 |
| ENSG0000016 | gene | 4,070150621 | 0,01436317 | 0,252327435 | PRDX3 | 2,025082184 |
| ENSG0000026 | gene | 4,111801425 | 0,006132937 | 0,239194577 | SNORD3C | 2,039770593 |
| ENSG0000013 | gene | 4,198911207 | 0,001168916 | 0,224332495 | MYOF | 2,07001528 |
| ENSG0000016 | gene | 4,22066104 | 0,038073192 | 0,28223387 | ADAM9 | 2,077468972 |
| ENSG0000007 | gene | 4,239671287 | 0,003977519 | 0,239194577 | ACER3 | 2,083952413 |
| ENSG0000014 | gene | 4,240303317 | 0,02013394 | 0,256090678 | LHFPL2 | 2,084167467 |
| ENSG0000016 | gene | 4,257635166 | 0,017825357 | 0,252327435 | SIGLEC16 | 2,090052331 |
| ENSG0000010 | gene | 4,273045936 | 0,033246655 | 0,27455389 | NTSR1 | 2,095264826 |
| ENSG0000021 | gene | 4,346756913 | 0,034328798 | 0,275250624 | IGKV3D-11 | 2,119939417 |
| ENSG0000011 | gene | 4,355707276 | 0,021179129 | 0,256239299 | FLVCR2 | 2,122907001 |
| ENSG0000018 | gene | 4,356315594 | 0,033088913 | 0,274068881 | LILRB4 | 2,123108474 |
| ENSG0000018 | gene | 4,41925733 | 0,027265969 | 0,267343734 | TMEM150B | 2,143803941 |
| ENSG0000021 | gene | 4,478261248 | 0,027896977 | 0,268373647 | IGLV7-43 | 2,162938693 |
| ENSG0000018 | gene | 4,48251988 | 0,025555736 | 0,265635455 | ADAP2 | 2,164309981 |
| ENSG0000015 | gene | 4,625861542 | 0,012391136 | 0,249534675 | CD1D | 2,209722085 |
| ENSG0000014 | gene | 4,722978547 | 0,046035243 | 0,293309443 | ST14 | 2,239696982 |
| ENSG0000017 | gene | 4,748607388 | 0,044553533 | 0,2922856 | KCTD12 | 2,24750448 |
| ENSG0000008 | gene | 4,971440033 | 0,003250296 | 0,238752762 | CRLS1 | 2,313663805 |
| ENSG0000024 | gene | 5,119617048 | 0,037329399 | 0,2810952 | IGKV1-27 | 2,356035899 |
| ENSG0000023 | gene | 5,28008444 | 0,025424511 | 0,265635455 | IGKV1-6 | 2,400561002 |

| | | | | | | |
|---|---|---|---|---|---|---|
| ENSG000002 | gene | 5,309340385 | 0,034583269 | 0,27606048 | NAIPP3 | 2,408532636 |
| ENSG000001 | gene | 5,312631279 | 0,031049489 | 0,271804321 | GASK1B | 2,409426587 |
| ENSG000001 | gene | 5,331478124 | 0,017125048 | 0,252327435 | MYBL2 | 2,414535568 |
| ENSG000002 | gene | 5,565374221 | 0,046432856 | 0,294465415 | IGKV1D-12 | 2,476478699 |
| ENSG000001 | gene | 5,580072581 | 0,021313407 | 0,256239299 | RAB13 | 2,480283888 |
| ENSG000002 | gene | 5,586499406 | 0,006637337 | 0,239194577 | AC024051.10 | 2,481944549 |
| ENSG000002 | gene | 5,623200062 | 0,031898163 | 0,272282875 | IGHV3-74 | 2,491391376 |
| ENSG000001 | gene | 5,777654624 | 0,019139993 | 0,254046692 | C1QA | 2,530483965 |
| ENSG000002 | gene | 5,829207192 | 0,037181482 | 0,2810952 | IGLV3-19 | 2,543299681 |
| ENSG000001 | gene | 5,833507284 | 0,02472624 | 0,264837204 | C1QC | 2,544363537 |
| ENSG000000 | gene | 6,109819764 | 0,006307556 | 0,239194577 | MGST1 | 2,611129822 |
| ENSG000001 | gene | 6,112461002 | 0,000731059 | 0,205762815 | SLC50A1 | 2,611753356 |
| ENSG000002 | gene | 6,236982014 | 0,037645404 | 0,282119411 | IGKV3-15 | 2,640848099 |
| ENSG000002 | gene | 6,314047591 | 0,049082154 | 0,299174578 | IGHV6-1 | 2,658565135 |
| ENSG000001 | gene | 6,360481152 | 4,25189E-05 | 0,129441475 | WDR13 | 2,669135905 |
| ENSG000001 | gene | 6,393005145 | 0,012795374 | 0,249534675 | CLEC12A | 2,676494255 |
| ENSG000002 | gene | 6,473745363 | 0,016527224 | 0,252327435 | IGHV4-61 | 2,69460062 |
| ENSG000002 | gene | 6,571092253 | 0,044272172 | 0,2922856 | IGHV1-18 | 2,716133197 |
| ENSG000001 | gene | 6,686331387 | 0,043851047 | 0,292071193 | ASGR2 | 2,741214859 |
| ENSG000002 | gene | 6,802078122 | 0,014718792 | 0,252327435 | IGKV2D-30 | 2,765975575 |
| ENSG000002 | gene | 6,81233756 | 0,045290377 | 0,2922856 | IGKC | 2,768149924 |
| ENSG000001 | gene | 7,041377033 | 0,034622557 | 0,27606048 | C1QB | 2,815857594 |
| ENSG000002 | gene | 7,160828093 | 0,02381507 | 0,263452051 | IGKV2-29 | 2,840126433 |
| ENSG000002 | gene | 7,465990662 | 0,047852647 | 0,296879689 | IGLV2-11 | 2,900333704 |
| ENSG000002 | gene | 7,720917492 | 0,014822239 | 0,252327435 | IGKV1-16 | 2,948772296 |
| ENSG000002 | gene | 7,811606269 | 0,037030936 | 0,281022131 | IGKV1-5 | 2,965619234 |
| ENSG000001 | gene | 7,954208725 | 0,020491662 | 0,256090678 | FN1 | 2,99171842 |
| ENSG000002 | gene | 8,026182437 | 0,045406924 | 0,2922856 | IGHV3-21 | 3,004713949 |
| ENSG000002 | gene | 8,075340929 | 0,022810369 | 0,260623396 | IGKV1-12 | 3,01352317 |
| ENSG000002 | gene | 9,10928726 | 0,00670578 | 0,239194577 | IGKV2-30 | 3,187338177 |
| ENSG000002 | gene | 9,395880744 | 0,03881451 | 0,282581813 | IGLV1-47 | 3,232028402 |
| ENSG000002 | gene | 9,426098268 | 0,016007023 | 0,252327435 | IGLV7-46 | 3,236660722 |
| ENSG000002 | gene | 9,705127077 | 0,015888898 | 0,252327435 | CES1P1 | 3,278747103 |
| ENSG000001 | gene | 9,79875092 | 0,016484243 | 0,252327435 | TCN2 | 3,292597856 |
| ENSG000002 | gene | 10,30399873 | 0,048224407 | 0,296879689 | IGHV3-72 | 3,365132415 |
| ENSG000002 | gene | 12,61432267 | 0,048959433 | 0,299059804 | IGHV3-66 | 3,656990837 |
| ENSG000002 | gene | 13,28898443 | 0,037296931 | 0,2810952 | IGHV3-48 | 3,73215895 |
| ENSG000002 | gene | 13,74189797 | 0,014180148 | 0,252327435 | IGHG4 | 3,780509371 |
| ENSG000002 | gene | 14,32402021 | 0,036075865 | 0,280020038 | IGKV1-39 | 3,840364554 |
| ENSG000002 | gene | 14,76818569 | 0,009838199 | 0,242573226 | IGKV3D-20 | 3,884420693 |
| ENSG000002 | gene | 15,18417138 | 0,044837617 | 0,2922856 | IGKV2-28 | 3,924496275 |
| ENSG000002 | gene | 15,34365752 | 0,010778794 | 0,246139857 | IGKV3-20 | 3,939570519 |
| ENSG000001 | gene | 15,43044425 | 0,039266235 | 0,283132205 | IFI27 | 3,947707693 |
| ENSG000002 | gene | 15,78978938 | 0,025760398 | 0,266004869 | IGHV3-53 | 3,980920022 |
| ENSG000002 | gene | 17,0842345 | 0,025747633 | 0,266004869 | IGHG3 | 4,094593701 |
| ENSG000002 | gene | 18,47792834 | 0,016121859 | 0,252327435 | IGHV3-20 | 4,207731112 |
| ENSG000002 | gene | 19,20657311 | 0,04574256 | 0,292595946 | IGHV3-33 | 4,263528227 |
| ENSG000001 | gene | 20,19791275 | 0,028638603 | 0,269662565 | VSIG4 | 4,336134307 |
| ENSG000002 | gene | 27,70333857 | 0,011386143 | 0,249534675 | IGHV4-39 | 4,791987943 |
| ENSG000002 | gene | 29,14549278 | 0,00481664 | 0,239194577 | IGHV1-2 | 4,865200889 |
| ENSG000002 | gene | 34,90653626 | 0,023800092 | 0,263452051 | IGHG1 | 5,125425302 |
| ENSG000001 | gene | 39,11478014 | 0,005367078 | 0,239194577 | CES1 | 5,289641951 |

**SARS-CoV2_unique**

| gene_name | id | feature | fc | pval | qval | de |
|---|---|---|---|---|---|---|
| RPS27AP16 | ENSG0000022 | gene | 0,033606837 | 0,04003026 | 0,765722912 | -4,89510141 |
| AC007383.4 | ENSG0000027 | gene | 0,164101059 | 0,003671247 | 0,493619508 | -2,60734355 |
| METTL21A | ENSG0000014 | gene | 0,169918461 | 0,028136883 | 0,727417021 | -2,55708549 |
| GNB3 | ENSG0000011 | gene | 0,18802193 | 0,006432253 | 0,560473839 | -2,41102715 |
| SURF4 | ENSG0000014 | gene | 0,254017884 | 2,10045E-05 | 0,067991613 | -1,97699802 |
| LRRC28 | ENSG0000016 | gene | 0,278264188 | 0,0452138 | 0,776242869 | -1,84547285 |
| SENP2 | ENSG0000016 | gene | 0,284212508 | 0,018535967 | 0,669155317 | -1,81495805 |
| PGPEP1 | ENSG0000013 | gene | 0,325010802 | 0,000282542 | 0,17148505 | -1,62144043 |
| ARL6 | ENSG0000011 | gene | 0,340469676 | 0,010885541 | 0,575745644 | -1,55440178 |
| ANG | ENSG0000021 | gene | 0,342405416 | 0,000892769 | 0,298954595 | -1,54622257 |
| NIPA1 | ENSG0000017 | gene | 0,345682561 | 0,034856571 | 0,761794317 | -1,53248027 |
| YTHDC2 | ENSG0000004 | gene | 0,361423715 | 0,012856328 | 0,605697652 | -1,46823692 |
| GNL3L | ENSG0000013 | gene | 0,404750714 | 0,006610906 | 0,560473839 | -1,30489447 |
| RPL29 | ENSG0000016 | gene | 0,426391216 | 0,0453228 | 0,776242869 | -1,22975038 |
| SGK2 | ENSG0000010 | gene | 0,440564497 | 0,002459398 | 0,493619508 | -1,18257485 |
| MFSD8 | ENSG0000016 | gene | 0,451875231 | 0,000548567 | 0,253673251 | -1,14600361 |
| SLC25A20 | ENSG0000017 | gene | 0,455426867 | 0,036498789 | 0,765722912 | -1,13470869 |
| PIK3R4 | ENSG0000019 | gene | 0,458482338 | 0,029867873 | 0,745621897 | -1,12506194 |
| MIR6784 | ENSG0000027 | gene | 0,462073106 | 0,029952549 | 0,745818463 | -1,11380697 |
| WRN | ENSG0000016 | gene | 0,46798039 | 0,032344129 | 0,751763165 | -1,09548002 |
| PPT2-EGFL8 | ENSG0000025 | gene | 0,480723547 | 0,029019106 | 0,735782087 | -1,05672062 |
| SERHL | ENSG0000017 | gene | 0,482675233 | 0,015975282 | 0,65735579 | -1,05087529 |
| TMEM164 | ENSG0000015 | gene | 0,489895041 | 0,035129134 | 0,761794317 | -1,02945541 |
| NDUFAF4 | ENSG0000012 | gene | 0,494640006 | 0,010845106 | 0,575745644 | -1,01554917 |
| VEGFA | ENSG0000011 | gene | 0,507545057 | 0,034762362 | 0,761794317 | -0,97839219 |
| TSC22D3 | ENSG0000015 | gene | 0,533656844 | 0,023108 | 0,700392548 | -0,90601575 |
| AC027796.4 | ENSG0000026 | gene | 0,54781914 | 0,029778635 | 0,745621897 | -0,86822842 |
| C1DP1 | ENSG0000023 | gene | 0,549782549 | 0,031322669 | 0,749219164 | -0,86306698 |
| STOML1 | ENSG0000006 | gene | 0,558273145 | 0,01526966 | 0,636410574 | -0,84095694 |
| RREB1 | ENSG0000012 | gene | 0,582712827 | 0,040537296 | 0,765722912 | -0,77914303 |
| GGT8P | ENSG0000023 | gene | 0,589520314 | 0,023633523 | 0,700392548 | -0,76238657 |
| AL513175.2 | ENSG0000023 | gene | 0,59030577 | 0,046367675 | 0,783814537 | -0,76046565 |
| MYCBP2 | ENSG0000000 | gene | 0,601435744 | 0,034666964 | 0,761794317 | -0,73351748 |
| SLC38A6 | ENSG0000013 | gene | 0,610976183 | 0,023953087 | 0,700392548 | -0,71081195 |
| AC048341.2 | ENSG0000027 | gene | 0,614761005 | 0,008114167 | 0,560473839 | -0,70190244 |
| PSRC1 | ENSG0000013 | gene | 0,62020154 | 0,032367476 | 0,751763165 | -0,68919099 |
| DANCR | ENSG0000022 | gene | 0,631052698 | 0,006832487 | 0,560473839 | -0,66416761 |
| RAP1GAP | ENSG0000007 | gene | 0,637704426 | 0,020084804 | 0,682146839 | -0,6490402 |
| LINC02313 | ENSG0000025 | gene | 0,639221808 | 0,049918014 | 0,788217613 | -0,64561147 |
| RAD54L | ENSG0000008 | gene | 0,644852533 | 0,043370404 | 0,767591009 | -0,63295882 |
| AP006621.3 | ENSG0000025 | gene | 0,650167628 | 0,006667442 | 0,560473839 | -0,62111637 |
| ACOT8 | ENSG0000010 | gene | 0,651588763 | 0,008814582 | 0,566479818 | -0,61796637 |
| KRT4 | ENSG0000017 | gene | 0,657524406 | 0,009136845 | 0,570481883 | -0,60488365 |
| TMEM97 | ENSG0000010 | gene | 0,667933669 | 9,0942E-05 | 0,092192886 | -0,58222326 |
| CDC42BPA | ENSG0000014 | gene | 0,67459005 | 0,00049806 | 0,2418332 | -0,56791706 |
| FTLP3 | ENSG0000022 | gene | 0,690578195 | 0,015550455 | 0,642597757 | -0,53412331 |
| POLE4 | ENSG0000011 | gene | 0,694194546 | 0,023294364 | 0,700392548 | -0,52658806 |
| AC130343.1 | ENSG0000026 | gene | 0,696187329 | 0,043025625 | 0,767591009 | -0,52245254 |
| MTX1 | ENSG0000017 | gene | 0,702108567 | 0,040795944 | 0,765722912 | -0,51023396 |
| RTN4R | ENSG0000004 | gene | 0,715363866 | 0,018703191 | 0,669657331 | -0,48325085 |
| GACAT2 | ENSG0000026 | gene | 0,716144366 | 0,024566171 | 0,702826025 | -0,48167765 |
| THOP1 | ENSG0000017 | gene | 0,71894428 | 0,005222027 | 0,524190612 | -0,47604813 |
| ATP6V0E2-AS | ENSG0000020 | gene | 0,723775759 | 0,029316716 | 0,739466574 | -0,46638531 |
| NMRAL2P | ENSG0000017 | gene | 0,724434734 | 0,025583031 | 0,702826025 | -0,46507237 |
| IFT172 | ENSG0000013 | gene | 0,732088874 | 0,00736261 | 0,560473839 | -0,44990929 |

| | | | | | |
|---|---|---|---|---|---|
| RPL32 | ENSG0000014 gene | 0,737940456 | 0,039154888 | 0,765722912 | -0,43842369 |
| JADE1 | ENSG0000007 gene | 0,74041954 | 0,047325219 | 0,785311848 | -0,43358513 |
| MED15 | ENSG0000009 gene | 0,743468076 | 0,023599552 | 0,700392548 | -0,4276573 |
| AL355312.3 | ENSG0000026 gene | 0,748444155 | 0,008205971 | 0,560473839 | -0,41803342 |
| CBS | ENSG0000016 gene | 0,755250701 | 0,034186264 | 0,761428461 | -0,40497248 |
| FMC1 | ENSG0000016 gene | 0,758906545 | 0,02323448 | 0,700392548 | -0,39800586 |
| GPANK1 | ENSG0000020 gene | 0,769484792 | 0,021697767 | 0,700392548 | -0,37803528 |
| FIZ1 | ENSG0000017 gene | 0,772974861 | 0,043053547 | 0,767591009 | -0,3715066 |
| NFYC | ENSG0000006 gene | 0,775610049 | 0,024190623 | 0,702826025 | -0,3665966 |
| HEXD | ENSG0000016 gene | 0,779077354 | 0,007090774 | 0,560473839 | -0,36016151 |
| CDKN2C | ENSG0000012 gene | 0,782139514 | 0,010755251 | 0,575745644 | -0,35450212 |
| NSMF | ENSG0000016 gene | 0,783722537 | 0,047026611 | 0,784665672 | -0,35158511 |
| FA2H | ENSG0000010 gene | 0,784086378 | 0,016276452 | 0,658924168 | -0,3509155 |
| RBM19 | ENSG0000012 gene | 0,786743818 | 0,025155263 | 0,702826025 | -0,34603416 |
| AC005586.1 | ENSG0000024 gene | 0,78830373 | 0,042157357 | 0,767591009 | -0,34317649 |
| C19orf71 | ENSG0000018 gene | 0,791426563 | 0,034316718 | 0,761794317 | -0,33747261 |
| PPP1R8 | ENSG0000011 gene | 0,79524243 | 0,012475324 | 0,605697652 | -0,33053336 |
| SEC31A | ENSG0000013 gene | 0,795455041 | 0,009659896 | 0,57333908 | -0,3301477 |
| SNORC | ENSG0000018 gene | 0,796868328 | 0,007488691 | 0,560473839 | -0,32758674 |
| AMH | ENSG0000010 gene | 0,797756623 | 0,009995316 | 0,57333908 | -0,32597941 |
| GALNT11 | ENSG0000017 gene | 0,802437554 | 0,045482656 | 0,777609275 | -0,31753897 |
| AC018638.5 | ENSG0000024 gene | 0,802701602 | 0,008270893 | 0,560473839 | -0,31706432 |
| ARMCX7P | ENSG0000020 gene | 0,802871166 | 0,022192992 | 0,700392548 | -0,31675959 |
| TIMP4 | ENSG0000015 gene | 0,805270411 | 0,045189923 | 0,776242869 | -0,31245477 |
| EEF1A1P11 | ENSG0000022 gene | 0,80964405 | 0,039246429 | 0,765722912 | -0,30464031 |
| TMEM79 | ENSG0000016 gene | 0,810246533 | 0,016615791 | 0,663077244 | -0,30356715 |
| RPL10P16 | ENSG0000017 gene | 0,810924443 | 0,033400227 | 0,754653699 | -0,3023606 |
| QSOX1 | ENSG0000011 gene | 0,813466392 | 0,024848811 | 0,702826025 | -0,29784535 |
| AC034102.6 | ENSG0000025 gene | 0,81369951 | 0,017715632 | 0,663077244 | -0,29743197 |
| EEF1A1P13 | ENSG0000025 gene | 0,816581702 | 0,019225677 | 0,674862552 | -0,29233085 |
| FDXR | ENSG0000016 gene | 0,817304749 | 0,008764492 | 0,566479818 | -0,29105398 |
| ADCK5 | ENSG0000017 gene | 0,817888645 | 0,002257158 | 0,487094782 | -0,29002366 |
| LPIN1 | ENSG0000013 gene | 0,824156927 | 0,046017319 | 0,782616789 | -0,27900903 |
| DPP7 | ENSG0000017 gene | 0,824316633 | 0,040724219 | 0,765722912 | -0,27872949 |
| ATP9A | ENSG0000005 gene | 0,824907198 | 0,048494986 | 0,785311848 | -0,27769627 |
| AC023355.2 | ENSG0000025 gene | 0,825346715 | 0,044539593 | 0,776242869 | -0,2769278 |
| EXD2 | ENSG0000008 gene | 0,826242075 | 0,003245229 | 0,493619508 | -0,27536357 |
| RPL3P4 | ENSG0000023 gene | 0,827896718 | 0,013800713 | 0,620457054 | -0,27247729 |
| RPS23P8 | ENSG0000023 gene | 0,828712064 | 0,036754672 | 0,765722912 | -0,27105717 |
| COPRS | ENSG0000017 gene | 0,829500249 | 0,014929858 | 0,633117253 | -0,26968568 |
| CERK | ENSG0000010 gene | 0,830820655 | 0,022158578 | 0,700392548 | -0,26739101 |
| LRFN4 | ENSG0000017 gene | 0,831416489 | 0,008039437 | 0,560473839 | -0,26635673 |
| STK16 | ENSG0000011 gene | 0,834968831 | 0,043194253 | 0,767591009 | -0,26020575 |
| RPL7 | ENSG0000014 gene | 0,838384786 | 0,003611306 | 0,493619508 | -0,25431556 |
| MARVELD1 | ENSG0000015 gene | 0,839045904 | 0,029307717 | 0,739466574 | -0,25317835 |
| MFSD10 | ENSG0000010 gene | 0,840327692 | 0,011249792 | 0,583167084 | -0,25097607 |
| MZF1 | ENSG0000009 gene | 0,841473284 | 0,048038479 | 0,785311848 | -0,24901063 |
| RPL7A | ENSG0000014 gene | 0,842916756 | 0,030982302 | 0,749158036 | -0,24653793 |
| PLEKHM1 | ENSG0000022 gene | 0,844425722 | 0,047874711 | 0,785311848 | -0,24395757 |
| FRAT2 | ENSG0000018 gene | 0,846372437 | 0,007933193 | 0,560473839 | -0,24063545 |
| CAMKK1 | ENSG0000000 gene | 0,847464014 | 0,003803245 | 0,497082126 | -0,23877599 |
| AC024293.1 | ENSG0000024 gene | 0,849077865 | 0,026162185 | 0,702826025 | -0,23603123 |
| NCKIPSD | ENSG0000021 gene | 0,849214554 | 0,022819958 | 0,700392548 | -0,235799 |
| AL589743.1 | ENSG0000024 gene | 0,849743515 | 0,009485725 | 0,57333908 | -0,23490065 |
| RNF185 | ENSG0000013 gene | 0,850439348 | 0,00797639 | 0,560473839 | -0,23371975 |
| H2AJ | ENSG0000024 gene | 0,851463312 | 0,021909991 | 0,700392548 | -0,23198373 |
| MCOLN1 | ENSG0000009 gene | 0,85510689 | 0,027148516 | 0,722299273 | -0,22582332 |

| | | | | | | |
|---|---|---|---|---|---|---|
| RPS3AP26 | ENSG0000021 | gene | 0,855589287 | 0,035767818 | 0,765722912 | -0,22500968 |
| RHOBTB2 | ENSG0000000 | gene | 0,85899502 | 0,001165677 | 0,365157756 | -0,21927833 |
| HOMER3 | ENSG0000005 | gene | 0,860478799 | 0,043186703 | 0,767591009 | -0,21678845 |
| TRIP6 | ENSG0000008 | gene | 0,861121663 | 0,013155379 | 0,61125306 | -0,21571101 |
| FAM160A2 | ENSG0000005 | gene | 0,862060566 | 0,038918803 | 0,765722912 | -0,21413886 |
| TGFB1I1 | ENSG0000014 | gene | 0,862700606 | 0,02601525 | 0,702826025 | -0,21306813 |
| CIT | ENSG0000012 | gene | 0,864250254 | 0,007966404 | 0,560473839 | -0,21047897 |
| DNAJC9-AS1 | ENSG0000023 | gene | 0,864618881 | 0,021499124 | 0,700392548 | -0,20986375 |
| NR4A1AS | ENSG0000025 | gene | 0,865492307 | 0,039312832 | 0,765722912 | -0,2084071 |
| LRRC20 | ENSG0000017 | gene | 0,866246482 | 0,000170382 | 0,137881717 | -0,20715051 |
| AL358781.1 | ENSG0000017 | gene | 0,870448709 | 0,033907738 | 0,756961016 | -0,20016881 |
| NTRK3 | ENSG0000014 | gene | 0,872087677 | 0,038206643 | 0,765722912 | -0,19745491 |
| ADI1 | ENSG0000018 | gene | 0,876232853 | 0,028877699 | 0,735782087 | -0,19061379 |
| INPP5E | ENSG0000014 | gene | 0,87635289 | 0,028103864 | 0,727417021 | -0,19041616 |
| ABCB8 | ENSG0000019 | gene | 0,87981523 | 0,04119009 | 0,765859559 | -0,18472752 |
| C19orf33 | ENSG0000016 | gene | 0,879996647 | 0,017310537 | 0,663077244 | -0,18443007 |
| NPIPA9 | ENSG0000023 | gene | 0,882857451 | 0,014435641 | 0,628093865 | -0,17974758 |
| LBH | ENSG0000021 | gene | 0,883281402 | 0,046010577 | 0,782616789 | -0,17905496 |
| BX255925.3 | ENSG0000028 | gene | 0,883721878 | 0,049758096 | 0,788217613 | -0,17833569 |
| MAP3K11 | ENSG0000017 | gene | 0,885751132 | 0,015234301 | 0,636410574 | -0,17502669 |
| ATP9B | ENSG0000016 | gene | 0,886564704 | 0,012660848 | 0,605697652 | -0,17370217 |
| COX14 | ENSG0000017 | gene | 0,887899615 | 0,03957076 | 0,765722912 | -0,17153152 |
| PACSIN3 | ENSG0000016 | gene | 0,891304401 | 0,042199912 | 0,767591009 | -0,16600987 |
| EIF3L | ENSG0000010 | gene | 0,891600191 | 0,029694438 | 0,745621897 | -0,16553117 |
| LAPTM4B | ENSG0000010 | gene | 0,894232367 | 0,011976032 | 0,599480655 | -0,16127833 |
| ACACA | ENSG0000027 | gene | 0,896176855 | 0,047122513 | 0,784917185 | -0,15814463 |
| DOCK10 | ENSG0000013 | gene | 0,89832196 | 0,007536909 | 0,560473839 | -0,15469549 |
| MRPL57 | ENSG0000017 | gene | 0,898750564 | 0,009145454 | 0,570481883 | -0,15400732 |
| ZNF219 | ENSG0000016 | gene | 0,899198667 | 0,049768101 | 0,788217613 | -0,1532882 |
| AC141586.3 | ENSG0000026 | gene | 0,902634571 | 0,030056659 | 0,746496713 | -0,14778606 |
| ZZEF1 | ENSG0000007 | gene | 0,902678678 | 0,024722804 | 0,702826025 | -0,14771557 |
| RPS10 | ENSG0000012 | gene | 0,912873177 | 0,019877673 | 0,682146839 | -0,13151365 |
| PDCD11 | ENSG0000014 | gene | 0,916438114 | 0,009897985 | 0,57333908 | -0,12589063 |
| NPC1 | ENSG0000014 | gene | 0,916891354 | 0,040170304 | 0,765722912 | -0,1251773 |
| HCN2 | ENSG0000009 | gene | 0,918418219 | 0,036144184 | 0,765722912 | -0,12277683 |
| NNT | ENSG0000011 | gene | 0,923813383 | 0,041697921 | 0,767591009 | -0,11432665 |
| ALG1 | ENSG0000003 | gene | 0,926933457 | 0,005788646 | 0,546307419 | -0,10946232 |
| CLSTN1 | ENSG0000017 | gene | 0,927071091 | 0,032985975 | 0,754653699 | -0,10924812 |
| MRPS18B | ENSG0000020 | gene | 0,936171925 | 0,010330278 | 0,575745644 | -0,09515459 |
| SLC6A6 | ENSG0000013 | gene | 0,938160363 | 0,033567281 | 0,754653699 | -0,09209355 |
| PEX6 | ENSG0000012 | gene | 0,941148679 | 0,03920324 | 0,765722912 | -0,08750544 |
| NAAA | ENSG0000013 | gene | 0,944829358 | 0,013881801 | 0,621226599 | -0,0818743 |
| NCAPG2 | ENSG0000014 | gene | 0,947591454 | 0,018952512 | 0,673063319 | -0,07766291 |
| FRG1CP | ENSG0000028 | gene | 0,966748422 | 0,028411772 | 0,729502277 | -0,04878759 |
| HMGN1 | ENSG0000020 | gene | 1,031677694 | 0,023143441 | 0,700392548 | 0,044992329 |
| MYO1E | ENSG0000015 | gene | 1,066107005 | 0,021953421 | 0,700392548 | 0,092352248 |
| JARID2 | ENSG0000000 | gene | 1,069991807 | 0,048383091 | 0,785311848 | 0,09759975 |
| LAMB1 | ENSG0000009 | gene | 1,073710116 | 0,02348464 | 0,700392548 | 0,102604542 |
| AMIGO2 | ENSG0000013 | gene | 1,076610668 | 0,007767082 | 0,560473839 | 0,106496626 |
| ARL3 | ENSG0000013 | gene | 1,08463965 | 0,042138938 | 0,767591009 | 0,117215816 |
| CA12 | ENSG0000007 | gene | 1,09183806 | 0,025305368 | 0,702826025 | 0,126758893 |
| CAV2 | ENSG0000010 | gene | 1,097765819 | 0,035077304 | 0,761794317 | 0,134570324 |
| DUSP4 | ENSG0000012 | gene | 1,105952172 | 0,044208034 | 0,773521117 | 0,145288996 |
| TP53BP2 | ENSG0000014 | gene | 1,111916565 | 0,031976947 | 0,750135666 | 0,153048537 |
| GLP2R | ENSG0000006 | gene | 1,113452779 | 0,021366408 | 0,700392548 | 0,155040375 |
| LAMC2 | ENSG0000005 | gene | 1,1171933 | 0,040933716 | 0,765722912 | 0,159878826 |
| KDM1A | ENSG0000000 | gene | 1,121092249 | 0,011854425 | 0,597348252 | 0,164904995 |

| | | | | | | |
|---|---|---|---|---|---|---|
| USP40 | ENSG0000008 | gene | 1,121186625 | 0,040337659 | 0,765722912 | 0,16502644 |
| HR | ENSG0000016 | gene | 1,12361015 | 0,041014892 | 0,765722912 | 0,168141562 |
| SAFB | ENSG0000016 | gene | 1,123659507 | 0,048435557 | 0,785311848 | 0,168204934 |
| MTA3 | ENSG0000005 | gene | 1,127997883 | 0,038301966 | 0,765722912 | 0,17376436 |
| MORC4 | ENSG0000013 | gene | 1,128016265 | 0,030827645 | 0,749158036 | 0,17378787 |
| MT-TA | ENSG0000021 | gene | 1,130271977 | 0,01797104 | 0,665808244 | 0,17666997 |
| AGR2 | ENSG0000010 | gene | 1,13131841 | 0,041325343 | 0,765859559 | 0,178005034 |
| GTF2E1 | ENSG0000015 | gene | 1,133118032 | 0,046939148 | 0,784665672 | 0,180298149 |
| ARF6 | ENSG0000016 | gene | 1,134531368 | 0,010556763 | 0,575745644 | 0,182096498 |
| CYP24A1 | ENSG0000001 | gene | 1,136712515 | 0,021449794 | 0,700392548 | 0,184867429 |
| TRIM4 | ENSG0000014 | gene | 1,136881547 | 0,014495373 | 0,628093865 | 0,185081946 |
| RANBP1 | ENSG0000009 | gene | 1,139768804 | 0,022048885 | 0,700392548 | 0,188741211 |
| ATMIN | ENSG0000016 | gene | 1,141347339 | 0,003587934 | 0,493619508 | 0,190737904 |
| HMCES | ENSG0000018 | gene | 1,141392637 | 0,02107691 | 0,700392548 | 0,190795162 |
| PDIA4 | ENSG0000015 | gene | 1,14405219 | 0,039440591 | 0,765722912 | 0,194152868 |
| MAPKAP1 | ENSG0000011 | gene | 1,145876762 | 0,024776748 | 0,702826025 | 0,196451892 |
| FBXO41 | ENSG0000016 | gene | 1,147791013 | 0,009164368 | 0,570481883 | 0,198859984 |
| MSH6 | ENSG0000011 | gene | 1,15223069 | 0,046934456 | 0,784665672 | 0,20442959 |
| PTPRM | ENSG0000017 | gene | 1,15608379 | 0,004627113 | 0,510612383 | 0,209245965 |
| DIS3L | ENSG0000016 | gene | 1,159895738 | 0,046491316 | 0,783814537 | 0,213995129 |
| DNAJC14 | ENSG0000013 | gene | 1,160642244 | 0,048409991 | 0,785311848 | 0,214923344 |
| BEX3 | ENSG0000016 | gene | 1,160671561 | 0,04030979 | 0,765722912 | 0,214959786 |
| KLC1 | ENSG0000012 | gene | 1,162713641 | 0,037180808 | 0,765722912 | 0,217495827 |
| CKAP4 | ENSG0000013 | gene | 1,163332134 | 0,034634263 | 0,761794317 | 0,218263048 |
| TNFSF12 | ENSG0000023 | gene | 1,164321576 | 0,043928744 | 0,771136021 | 0,219489574 |
| AKAP12 | ENSG0000013 | gene | 1,165477869 | 0,014296415 | 0,626651758 | 0,22092161 |
| COL5A2 | ENSG0000020 | gene | 1,166063893 | 0,030732392 | 0,749158036 | 0,221646841 |
| IGFBP3 | ENSG0000014 | gene | 1,166526417 | 0,009599271 | 0,57333908 | 0,222218979 |
| RARRES1 | ENSG0000011 | gene | 1,16747368 | 0,023153935 | 0,700392548 | 0,223390025 |
| TERF1 | ENSG0000014 | gene | 1,167806814 | 0,033150326 | 0,754653699 | 0,223801634 |
| CYB5A | ENSG0000016 | gene | 1,168777787 | 0,044665447 | 0,776242869 | 0,225000665 |
| CEBPD | ENSG0000022 | gene | 1,168898273 | 0,035331841 | 0,764159259 | 0,22514938 |
| ATP8B4 | ENSG0000010 | gene | 1,171830709 | 0,019651432 | 0,682146839 | 0,228764163 |
| PAQR5 | ENSG0000013 | gene | 1,175246543 | 0,014135068 | 0,626651758 | 0,232963437 |
| RRBP1 | ENSG0000012 | gene | 1,175491347 | 0,007640386 | 0,560473839 | 0,233263919 |
| HSPA5 | ENSG0000004 | gene | 1,176746543 | 0,016284811 | 0,658924168 | 0,234803615 |
| C2orf74 | ENSG0000023 | gene | 1,177196797 | 0,020090001 | 0,682146839 | 0,235355522 |
| PDHX | ENSG0000011 | gene | 1,179887008 | 0,025105087 | 0,702826025 | 0,238648706 |
| KRT18 | ENSG0000011 | gene | 1,18052875 | 0,005794425 | 0,546307419 | 0,239433177 |
| PLEKHG2 | ENSG0000009 | gene | 1,181876005 | 0,007343017 | 0,560473839 | 0,241078685 |
| PIGH | ENSG0000010 | gene | 1,182946979 | 0,009246541 | 0,571930932 | 0,242385411 |
| SH2D3A | ENSG0000012 | gene | 1,18296156 | 0,017818348 | 0,663077244 | 0,242403194 |
| PNMA2 | ENSG0000024 | gene | 1,186633183 | 0,019903583 | 0,682146839 | 0,246874031 |
| TCP1 | ENSG0000012 | gene | 1,187925422 | 0,047761519 | 0,785311848 | 0,248444267 |
| PSME3 | ENSG0000013 | gene | 1,188701867 | 0,016489312 | 0,66168473 | 0,249386925 |
| CD40 | ENSG0000010 | gene | 1,189318609 | 0,040684578 | 0,765722912 | 0,250135253 |
| PDCD6 | ENSG0000024 | gene | 1,193640393 | 0,004243918 | 0,505536224 | 0,255368263 |
| PRPF40B | ENSG0000011 | gene | 1,193922631 | 0,025470505 | 0,702826025 | 0,25570935 |
| PCED1B | ENSG0000017 | gene | 1,19685971 | 0,024841432 | 0,702826025 | 0,259254056 |
| MT-RNR2 | ENSG0000021 | gene | 1,200926339 | 0,036648516 | 0,765722912 | 0,264147663 |
| NIF3L1 | ENSG0000019 | gene | 1,202714265 | 0,035662712 | 0,765722912 | 0,266293935 |
| AC005082.2 | ENSG0000028 | gene | 1,211478636 | 0,02546473 | 0,702826025 | 0,276768963 |
| HMGB1P10 | ENSG0000021 | gene | 1,2214739 | 0,00679011 | 0,560473839 | 0,288623037 |
| FANCI | ENSG0000014 | gene | 1,223583294 | 0,048841347 | 0,787085197 | 0,291112315 |
| WDR92 | ENSG0000024 | gene | 1,224389525 | 0,010534006 | 0,575745644 | 0,292062607 |
| GHDC | ENSG0000016 | gene | 1,22610484 | 0,022145523 | 0,700392548 | 0,294082344 |
| SSB | ENSG0000013 | gene | 1,235441315 | 0,037305592 | 0,765722912 | 0,305026482 |

| | | | | | | |
|---|---|---|---|---|---|---|
| SPA17 | ENSG0000006 | gene | 1,236104825 | 0,000627211 | 0,273995771 | 0,305801093 |
| UQCRB | ENSG0000015 | gene | 1,240740507 | 0,041081417 | 0,765722912 | 0,311201417 |
| HSP90B1 | ENSG0000016 | gene | 1,241950497 | 0,028245909 | 0,727575645 | 0,312607671 |
| HSPA2 | ENSG0000012 | gene | 1,242355552 | 0,007977434 | 0,560473839 | 0,31307812 |
| EEF2KMT | ENSG0000011 | gene | 1,250688901 | 0,04625539 | 0,783814537 | 0,322722976 |
| PRAME | ENSG0000018 | gene | 1,256472898 | 0,004320822 | 0,505536224 | 0,329379553 |
| GPRC5A | ENSG0000001 | gene | 1,258239206 | 0,049614812 | 0,788217613 | 0,331406221 |
| HSP90AA2P | ENSG0000022 | gene | 1,260681645 | 0,0233899 | 0,700392548 | 0,334204004 |
| NEIL2 | ENSG0000015 | gene | 1,267783662 | 0,036158963 | 0,765722912 | 0,342308581 |
| SLTM | ENSG0000013 | gene | 1,273171118 | 0,026344215 | 0,702826025 | 0,348426335 |
| BBIP1 | ENSG0000021 | gene | 1,285436888 | 0,02853282 | 0,729502277 | 0,362258779 |
| SARNP | ENSG0000020 | gene | 1,285485409 | 0,028164844 | 0,727417021 | 0,362313235 |
| COCH | ENSG0000010 | gene | 1,28760916 | 0,008253275 | 0,560473839 | 0,364694745 |
| MB | ENSG0000019 | gene | 1,287680047 | 0,025632656 | 0,702826025 | 0,364774168 |
| SETP14 | ENSG0000024 | gene | 1,289720792 | 0,003125175 | 0,493619508 | 0,367058774 |
| DGCR6L | ENSG0000012 | gene | 1,303194445 | 0,045203615 | 0,776242869 | 0,382052359 |
| PITPNA-AS1 | ENSG0000023 | gene | 1,30353313 | 0,010036829 | 0,57333908 | 0,38242725 |
| AJUBA | ENSG0000012 | gene | 1,305663068 | 0,022420792 | 0,700392548 | 0,384782651 |
| THBD | ENSG0000017 | gene | 1,308231858 | 0,034799965 | 0,761794317 | 0,387618252 |
| SLC37A3 | ENSG0000015 | gene | 1,314444868 | 0,022990345 | 0,700392548 | 0,394453632 |
| FAM104B | ENSG0000018 | gene | 1,317416922 | 0,028005106 | 0,727417021 | 0,397711987 |
| KRT8 | ENSG0000017 | gene | 1,324810742 | 0,042441796 | 0,767591009 | 0,405786275 |
| ZNF274 | ENSG0000017 | gene | 1,325823971 | 0,002963873 | 0,493619508 | 0,406889242 |
| EDN1 | ENSG0000007 | gene | 1,326998114 | 0,005917436 | 0,550470362 | 0,40816632 |
| FGB | ENSG0000017 | gene | 1,327007095 | 0,026015137 | 0,702826025 | 0,408176084 |
| SEC11B | ENSG0000022 | gene | 1,337912676 | 0,048959812 | 0,787166786 | 0,419983956 |
| RNF121 | ENSG0000013 | gene | 1,338419447 | 0,023651921 | 0,700392548 | 0,420530313 |
| BCAM | ENSG0000018 | gene | 1,339854429 | 0,00321004 | 0,493619508 | 0,422076265 |
| C6orf226 | ENSG0000022 | gene | 1,342522307 | 0,004583157 | 0,510612383 | 0,42494606 |
| AD000091.1 | ENSG0000026 | gene | 1,348681891 | 0,012240547 | 0,60034321 | 0,431550105 |
| SKP2 | ENSG0000014 | gene | 1,353043435 | 0,012821293 | 0,605697652 | 0,436208153 |
| SULT2B1 | ENSG0000008 | gene | 1,353481197 | 0,042015951 | 0,767591009 | 0,436674846 |
| MED20 | ENSG0000012 | gene | 1,358072297 | 0,006342211 | 0,560473839 | 0,441560283 |
| SMIM11A | ENSG0000020 | gene | 1,361057936 | 0,017821353 | 0,663077244 | 0,44472848 |
| RBP4 | ENSG0000013 | gene | 1,363069904 | 0,044651463 | 0,776242869 | 0,446859552 |
| CP | ENSG0000004 | gene | 1,364003624 | 0,024351265 | 0,702826025 | 0,447847478 |
| LARS2 | ENSG0000001 | gene | 1,368216335 | 0,027512742 | 0,727417021 | 0,452296359 |
| DDX60L | ENSG0000018 | gene | 1,368517481 | 0,015346739 | 0,636889676 | 0,452613863 |
| TDRD7 | ENSG0000019 | gene | 1,37796219 | 0,019812582 | 0,682146839 | 0,462536303 |
| ZNFX1 | ENSG0000012 | gene | 1,37896036 | 0,012827486 | 0,605697652 | 0,463580985 |
| SNORD116-4 | ENSG0000027 | gene | 1,382450231 | 0,041986253 | 0,767591009 | 0,467227543 |
| AC015912.1 | ENSG0000026 | gene | 1,394201159 | 0,027659548 | 0,727417021 | 0,479438732 |
| ID1 | ENSG0000012 | gene | 1,395074841 | 0,005735389 | 0,546307419 | 0,48034252 |
| GSDME | ENSG0000010 | gene | 1,398886732 | 0,013657924 | 0,620457054 | 0,484279153 |
| DAZAP2 | ENSG0000018 | gene | 1,427535313 | 0,031306127 | 0,749219164 | 0,513526434 |
| HMGN1P36 | ENSG0000023 | gene | 1,430286995 | 0,03775359 | 0,765722912 | 0,51630466 |
| SAT1 | ENSG0000013 | gene | 1,433373862 | 0,010233417 | 0,575745644 | 0,519414952 |
| CDH1 | ENSG0000003 | gene | 1,447941036 | 0,000759653 | 0,295079662 | 0,534002853 |
| PDK4 | ENSG0000000 | gene | 1,451505046 | 0,040040794 | 0,765722912 | 0,537549588 |
| SERBP1P1 | ENSG0000021 | gene | 1,464343387 | 0,000201246 | 0,139592754 | 0,550253904 |
| TEX41 | ENSG0000022 | gene | 1,470342483 | 0,048873687 | 0,787085197 | 0,556152237 |
| CKS1BP2 | ENSG0000022 | gene | 1,478534462 | 0,022882364 | 0,700392548 | 0,564167871 |
| FGA | ENSG0000017 | gene | 1,485658294 | 0,021797648 | 0,700392548 | 0,571102329 |
| FNBP4 | ENSG0000010 | gene | 1,489136812 | 0,0137666 | 0,620457054 | 0,574476305 |
| SUSD2 | ENSG0000009 | gene | 1,50731792 | 0,004739665 | 0,516859327 | 0,591983739 |
| MAP3K9 | ENSG0000000 | gene | 1,508650947 | 0,035144048 | 0,761794317 | 0,593259051 |
| C3 | ENSG0000012 | gene | 1,570240462 | 0,033259839 | 0,754653699 | 0,650985506 |

| | | | | | | |
|---|---|---|---|---|---|---|
| SERPINA3 | ENSG0000019 | gene | 1,570756179 | 0,01176859 | 0,597348252 | 0,651459255 |
| TYMP | ENSG0000002 | gene | 1,577459369 | 0,003839065 | 0,497082126 | 0,657602846 |
| WASH6P | ENSG0000018 | gene | 1,590185655 | 0,013013039 | 0,607546277 | 0,669195211 |
| HERC5 | ENSG0000013 | gene | 1,60626865 | 0,008037056 | 0,560473839 | 0,683713205 |
| SEPTIN7P6 | ENSG0000024 | gene | 1,64053329 | 0,00361341 | 0,493619508 | 0,714164869 |
| SAMD9 | ENSG0000020 | gene | 1,64346026 | 0,029797748 | 0,745621897 | 0,716736572 |
| AC097709.1 | ENSG0000028 | gene | 1,714446231 | 0,000800706 | 0,297935015 | 0,777742659 |
| WDR60 | ENSG0000012 | gene | 1,781246251 | 0,044817761 | 0,776242869 | 0,832886978 |
| ARPC4-TTLL3 | ENSG0000025 | gene | 1,783510023 | 0,001692193 | 0,444131997 | 0,834719324 |
| MT-TV | ENSG0000021 | gene | 1,788345405 | 0,005235968 | 0,524190612 | 0,838625409 |
| TRIM16L | ENSG0000010 | gene | 1,8127658 | 0,006934586 | 0,560473839 | 0,858192549 |
| NPIPA1 | ENSG0000018 | gene | 1,836750153 | 0,024017168 | 0,700392548 | 0,877155395 |
| SPIRE2 | ENSG0000020 | gene | 1,84250797 | 0,025346057 | 0,702826025 | 0,88167086 |
| EIF2AK2 | ENSG0000005 | gene | 1,852429829 | 0,010684132 | 0,575745644 | 0,889418894 |
| MT-TF | ENSG0000021 | gene | 1,853024861 | 0,044900061 | 0,776242869 | 0,889882237 |
| TRIM14 | ENSG0000010 | gene | 1,968438831 | 0,001040527 | 0,336818531 | 0,977051882 |
| FUOM | ENSG0000014 | gene | 2,067204244 | 0,012184913 | 0,60034321 | 1,047680937 |
| PPP2R2A | ENSG0000022 | gene | 2,077111395 | 0,004003031 | 0,498377381 | 1,05457859 |
| TRIM34 | ENSG0000025 | gene | 2,092848592 | 0,025920022 | 0,702826025 | 1,065467943 |
| RTEL1 | ENSG0000025 | gene | 2,101149405 | 0,038817718 | 0,765722912 | 1,071178751 |
| AC004551.1 | ENSG0000025 | gene | 2,186659775 | 0,002205197 | 0,487094782 | 1,128728767 |
| TNK2 | ENSG0000006 | gene | 2,219487327 | 0,030721587 | 0,749158036 | 1,150226471 |
| GATAD2A | ENSG0000016 | gene | 2,223412679 | 0,032432901 | 0,751763165 | 1,152775746 |
| MED17 | ENSG0000004 | gene | 2,300980042 | 0,048482997 | 0,785311848 | 1,20224847 |
| BX890604.2 | ENSG0000028 | gene | 2,494387259 | 0,049526363 | 0,788217613 | 1,318685464 |
| MFSD14C | ENSG0000019 | gene | 2,923486382 | 0,049231457 | 0,788217613 | 1,547689871 |
| MMP17 | ENSG0000019 | gene | 2,940647796 | 0,049604428 | 0,788217613 | 1,556134002 |
| ZNF250 | ENSG0000019 | gene | 2,949093513 | 0,025449586 | 0,702826025 | 1,56027157 |
| HSD11B1L | ENSG0000016 | gene | 3,013928058 | 0,031900387 | 0,750135666 | 1,59164498 |
| POLR2J3 | ENSG0000028 | gene | 3,414944736 | 0,002592649 | 0,493619508 | 1,771862232 |
| ZNF566 | ENSG0000018 | gene | 3,434065909 | 0,025884119 | 0,702826025 | 1,779917729 |
| CCDC150 | ENSG0000014 | gene | 3,442209366 | 0,00595195 | 0,550470362 | 1,78333485 |
| AL392172.1 | ENSG0000022 | gene | 3,477577367 | 0,017324422 | 0,663077244 | 1,798082611 |
| EXOG | ENSG0000015 | gene | 3,543260864 | 0,001879868 | 0,468087114 | 1,825077684 |
| DPY19L3 | ENSG0000017 | gene | 3,659752756 | 0,005235647 | 0,524190612 | 1,871746187 |
| LINC01106 | ENSG0000017 | gene | 3,87233386 | 0,018990768 | 0,673063319 | 1,953203343 |
| SNORD14D | ENSG0000020 | gene | 4,151336543 | 0,00442352 | 0,510612383 | 2,053575894 |
| HERC6 | ENSG0000013 | gene | 4,2951874 | 0,005135044 | 0,524190612 | 2,102721078 |
| AL163636.3 | ENSG0000025 | gene | 4,577130303 | 0,004873672 | 0,52009038 | 2,194443364 |
| FAM71E1 | ENSG0000014 | gene | 5,127134404 | 0,00415356 | 0,504190259 | 2,358152717 |
| FXN | ENSG0000016 | gene | 6,017196653 | 0,041030368 | 0,765722912 | 2,589091507 |
| IRF7 | ENSG0000018 | gene | 6,299655065 | 0,009976856 | 0,57333908 | 2,655272837 |
| SNORD32A | ENSG0000020 | gene | 8,693482056 | 7,56732E-05 | 0,092192886 | 3,119934145 |
| FAM228B | ENSG0000021 | gene | 8,829039613 | 0,003442486 | 0,493619508 | 3,142256516 |
| AC004542.4 | ENSG0000027 | gene | 9,032413854 | 0,004311124 | 0,505536224 | 3,17511159 |

#### All Intersection genes

| MERS_COV_vs_SARS_COV | MERS_COV_vs_SARS_COV2 | MERS_COV_vs_PBMC | SARS_COV_vs_SARS_COV2 | SARS_COV_vs_PBMC | SARS_COV2_vs_PBMC | MERS_COV_vs_PBMC_vs_SARS_COV2 | MERS_COV_vs_SARS_COV_vs_PBMC | MERS_COV_vs_SARS_COV_vs_SARSC | PBMC_vs_SARS_COV_vs_SARSCOV2 | MERS_COV_vs_PBMC_vs_SARS_COV2 |
|---|---|---|---|---|---|---|---|---|---|---|
| FP236383.5 | GPCPD1 | FN1 | DNAJC13 | ATP6V1F | IFI27 | FN1 | FN1 | FN1 | AHNAK | FN1 |
| RPS2P5 | FN1 | GNS | IFITM1 | POLR2F | OAS1 | BRK1 | FADS1 | AC087393.2 | CXCL8 | BRK1 |
| SEMA3A | PLK2 | FADS1 | DDX27 | PFDN6 | LAP3 | HNRNPAB | PLCB3 | RPLP0P6 | DMPK | RSPRY1 |
| TPR | CSGALNACT1 | PLCB3 | RBM4 | RBM4 | SP110 | NME2 | KLC4 | GPR137 | LSS | NA |
| CDH6 | DKK1 | STING1 | SMARCA1 | ZNF83 | HEXA | RSPRY1 | APMAP | BRK1 | EIF1AD | NA |
| FN1 | RHOB | COX8A | PSME2 | DDX6 | TPRG1L | MRPL16 | POLR2E | CTDNEP1 | GLOD4 | NA |
| AP002490.1 | NIFK | KLC4 | ISG15 | PSMD13 | BCL9L | PCCB | CSTB | ARF3 | RBM4 | NA |
| STC2 | AC087393.2 | IMMT | MT-TS2 | BAX | ARL4C | NA | PSAP | RPSA | HLA-E | NA |
| COL6A3 | RALY | POMP | IFI35 | TWF2 | AMD1 | NA | PPP1R12A | RPL12P4 | STARD3 | NA |
| DYNC1H1 | RPLP0P6 | SRP19 | UBE2L6 | PPP1CA | RNH1 | NA | DDX6 | RPS7 | SP110 | NA |
| MT-ND6 | PRKG1-AS1 | APMAP | CIB1 | IFI27 | CDC42BPB | NA | COX6A1 | MBNL1 | SENP6 | NA |
| GOPC | ZNF584 | POLR2E | HLA-C | C1orf43 | MIR4767 | NA | TOP2A | EIF4G3 | BRD7 | NA |
| AL353644.2 | IGF1R | UBE2L3 | PHAX | SCAMP2 | POLR2G | NA | MBIP | DNAJC13 | CHFR | NA |
| RNA5-8SN4 | GPR137 | CSTB | SHFL | GET3 | SULT1A1 | NA | VKORC1 | AC073861.1 | USP25 | NA |
| RPL39P3 | PAPPA | PSAP | SAMHD1 | POLR2E | RSPRY1 | NA | GADD45B | RSPRY1 | POLR2G | NA |
| RNA5-8SN3 | BRK1 | PPP1R12A | IFI27 | CRTAP | STARD3 | NA | HAT1 | CXCL1 | RSPRY1 | NA |
| TFE3 | CTDNEP1 | TOMM22 | IFITM3 | PREB | MRPL53 | NA | SRRD | SMARCA5 | STAU1 | NA |
| RNA5-8SN2 | P3H3 | CYFIP1 | PUM3 | EIF1 | EIF2AK1 | NA | BRK1 | RPLP1 | TNFRSF10D | NA |
| AC027319.1 | CEACAM19 | DDX6 | ECSIT | SH3GL1 | DMPK | NA | FKBP15 | AL110504.1 | HEXA | NA |
| FP236383.4 | ARF3 | ARRDC3 | LAP3 | PRMT5 | ABHD15 | NA | UQCRC1 | AREG | CYTOR | NA |
| BICC1 | HNRNPAB | AP2A1 | EMD | BST2 | RHEB | NA | ATP5MC3 | TST | NACC2 | NA |
| RPS28 | NME2 | COX6A1 | MX1 | STX10 | HNRNPAB | NA | POP7 | NDUFB2 | BRK1 | NA |
| USP11 | CRTC1 | PBXIP1 | RIC8A | LAP3 | MEPCE | NA | ALCAM | AL353689.2 | RNH1 | NA |
| FAP | ABHD11 | SDHB | IFIT1 | PSMD4 | CHFR | NA | CCT5 | NA | EIF2AK1 | NA |
| RPS9 | RPSA | TK1 | PLSCR1 | HSF1 | CYTOR | NA | PGM2 | NA | CDC42BPB | NA |
| EXT1 | AC020909.3 | TOP2A | IFI6 | MYD88 | FABP5 | NA | SMARCD3 | NA | RHEB | NA |
| SLC6A9 | RPL12P4 | PPT1 | SMARCA5 | SEPTIN7 | HERC2P2 | NA | TMED9 | NA | HERC2P2 | NA |
| LSM7 | RPS7 | MBIP | STAT2 | COX6A1 | HERC2 | NA | MVD | NA | COQ5 | NA |
| SELPLG | MBNL1 | VKORC1 | STAT1 | PEPD | UROD | NA | AP3M1 | NA | MRM2 | NA |
| SNHG5 | TCIM | SMCO4 | TAP1 | ZMAT5 | PDIA3 | NA | PSMA2 | NA | LDHA | NA |
| FADS1 | ARF5 | AC243919.1 | USP18 | TMEM258 | STAU1 | NA | CDC42SE1 | NA | AL034430.2 | NA |
| EEF1D | ABHD2 | NRG1 | NTMT1 | PRDX1 | MRPL16 | NA | SEC61B | NA | NPC2 | NA |
| AC004477.2 | USP41 | CAPN2 | HERC2P2 | WASHC5 | HLA-E | NA | RSPRY1 | NA | TP53I3 | NA |
| PLCB3 | EIF4G3 | GADD45B | BRK1 | PKM | ELOC | NA | PSMA6 | NA | LAP3 | NA |
| DST | DNAJC13 | HAT1 | SENP6 | POFUT2 | SLFN11 | NA | UQCRH | NA | MRPL53 | NA |
| AC015674.1 | PVR | MRPS15 | HLA-E | MRPL51 | TXNRD2 | NA | AHCY | NA | FN1 | NA |
| KLC4 | NUAK2 | SRRD | SMARCD1 | SCPEP1 | GLOD4 | NA | SNAPIN | NA | IFI27 | NA |
| RPS21 | RPS7P1 | TSPO | MRPL2 | LHPP | MCFD2 | NA | GNL1 | NA | NA | NA |
| SH3GLB2 | AC073861.1 | BRK1 | RNH1 | ERH | VAMP2 | NA | TUBG1 | NA | NA | NA |
| H2AX | RSPRY1 | NDUFS6 | MBNL1 | APH1A | NACC2 | NA | GFM1 | NA | NA | NA |
| ATG13 | MRPL16 | VGLL4 | CUL5 | ATP6V1B2 | CXCL8 | NA | SBDS | NA | NA | NA |
| PDE6D | PSMC3IP | SUB1 | PSME1 | RAMAC | PCCB | NA | ATPAF2 | NA | NA | NA |
| GTPBP2 | CA9 | HNRNPAB | AHNAK | PSMA7 | TP53I3 | NA | GTF2F2 | NA | NA | NA |
| UAP1 | CXCL1 | FKBP15 | IL4R | AKR1A1 | LSS | NA | SLC25A11 | NA | NA | NA |
| APMAP | SMARCA5 | SLC1A5 | GLOD4 | PSMA5 | NPC2 | NA | ERH | NA | NA | NA |
| TRNP1 | RPLP1 | RTL6 | STARD3 | IER2 | COQ5 | NA | HSD17B10 | NA | NA | NA |
| TMEM50A | DNAJC16 | NME2 | TRIM25 | EIF4E2 | SNX8 | NA | SNRPG | NA | NA | NA |
| NDUFAB1 | AL110504.1 | YTHDF2 | CHTOP | GLB1 | IDS | NA | JAGN1 | NA | NA | NA |
| ARHGEF11 | PCCB | PA2G4 | OAS3 | MRPL21 | CDK5 | NA | USP53 | NA | NA | NA |
| POLR2E | PLIN2 | UQCRC1 | C16orf58 | GNPTAB | TCN2 | NA | PTAR1 | NA | NA | NA |
| GNB2 | AREG | ATP5MC3 | LSM6 | NANS | IFIT5 | NA | RNF181 | NA | NA | NA |
| MAVS | TST | ZFYVE26 | SPSB3 | CSTB | SENP6 | NA | STK39 | NA | NA | NA |
| CSTB | DDX60 | POP7 | CTDNEP1 | TSC22D4 | CDCA5 | NA | NDUFB9 | NA | NA | NA |
| VDR | ANXA2P2 | ALCAM | MRPL53 | DNAJB12 | BRK1 | NA | IER2 | NA | NA | NA |
| H2AZ1 | NDUFB2 | CCT5 | AREG | HERC2P2 | MRM2 | NA | NA | NA | NA | NA |
| TUFM | C12orf65 | ZNF266 | HEXA | COPS2 | AHNAK | NA | NA | NA | NA | NA |
| CLIC1 | RPL39 | NT5DC2 | ESF1 | BRK1 | ATP13A1 | NA | NA | NA | NA | NA |
| PSAP | HDAC5 | ORAI1 | RSPRY1 | GAPDH | LTA4H | NA | NA | NA | NA | NA |
| TOPBP1 | ARPC4 | PGM2 | IFIT3 | MT-TH | CZIB | NA | NA | NA | NA | NA |
| FJX1 | AL353689.2 | SMARCD3 | CHMP4B | SENP6 | EDEM2 | NA | NA | NA | NA | NA |
| CORO1B | NA | NDUFAF3 | STIM1 | DDX41 | TNFRSF10D | NA | NA | NA | NA | NA |
| AC087393.2 | NA | CFL2 | GUCD1 | HLA-E | TOMM5 | NA | NA | NA | NA | NA |
| MRPS22 | NA | MRPS12 | AC096772.2 | OTUD5 | STAG3L5P | NA | NA | NA | NA | NA |

| | | | | | | | | | | |
|---|---|---|---|---|---|---|---|---|---|---|
| PPP1R12A | NA | TMED9 | BTBD10 | NUBP1 | ACTG1 | NA | NA | NA | NA | NA |
| FDPS | NA | PRXL2C | AL353689.2 | HDGF | NME2 | NA | NA | NA | NA | NA |
| DDX6 | NA | MVD | IFIH1 | RHOA | PRDX6 | NA | NA | NA | NA | NA |
| RPS28P7 | NA | SYNE1 | AL110504.1 | DUS1L | RBM4 | NA | NA | NA | NA | NA |
| BX322234.1 | NA | AP3M1 | LGALS3BP | TRIM26 | DOK1 | NA | NA | NA | NA | NA |
| HAS2 | NA | PPP1R2 | G2E3 | SLC4A2 | NKIRAS2 | NA | NA | NA | NA | NA |
| DAD1 | NA | OLA1 | NPC2 | POP7 | CWC15 | NA | NA | NA | NA | NA |
| AC060766.5 | NA | RC3H2 | PHF21A | OS9 | BABAM2 | NA | NA | NA | NA | NA |
| RPLP0P6 | NA | IFNAR2 | CHTF8 | AC044849.1 | LDHA | NA | NA | NA | NA | NA |
| TIMM13 | NA | UQCRC2 | CFAP97 | GTF2F2 | NDUFB3 | NA | NA | NA | NA | NA |
| COX6A1 | NA | PAPSS1 | FN1 | SSNA1 | TPX2 | NA | NA | NA | NA | NA |
| KISS1 | NA | PSMA2 | NDUFB2 | ENO1 | GRAMD4 | NA | NA | NA | NA | NA |
| OST4 | NA | PNMA1 | EIF4G3 | DMXL1 | USP25 | NA | NA | NA | NA | NA |
| TOP2A | NA | CDC42SE1 | LDHA | DHRS4 | FN1 | NA | NA | NA | NA | NA |
| RABAC1 | NA | FBXL3 | STIP1 | KIAA0930 | EIF1AD | NA | NA | NA | NA | NA |
| RPPH1 | NA | CLSTN3 | PITPNA | YIPF3 | UAP1L1 | NA | NA | NA | NA | NA |
| MBIP | NA | SEC61B | CERS2 | UPP1 | AL034430.2 | NA | NA | NA | NA | NA |
| VKORC1 | NA | RSPRY1 | HELZ2 | PTDSS1 | ZBTB4 | NA | NA | NA | NA | NA |
| AC073641.1 | NA | MRPL16 | RPL12P4 | RNH1 | BRD7 | NA | NA | NA | NA | NA |
| AC009511.1 | NA | PSMA6 | TIGD2 | FADS1 | YARS1 | NA | NA | NA | NA | NA |
| SEC62 | NA | TTC39B | LSS | ARPP19 | NA | NA | NA | NA | NA | NA |
| PTGES | NA | EXOSC4 | CASP7 | SNX11 | NA | NA | NA | NA | NA | NA |
| RNU6-26P | NA | NAA20 | APLP2 | ATP5MC1 | NA | NA | NA | NA | NA | NA |
| GADD45B | NA | RFTN1 | NOP53 | GNL1 | NA | NA | NA | NA | NA | NA |
| SEC24D | NA | SECISBP2 | PTTG1IP | AHNAK | NA | NA | NA | NA | NA | NA |
| SDF4 | NA | UQCRH | CMTR1 | GLOD4 | NA | NA | NA | NA | NA | NA |
| CASC3 | NA | AHCY | PARP9 | LGALS1 | NA | NA | NA | NA | NA | NA |
| HAT1 | NA | E2F1 | HNRNPH1 | STARD3 | NA | NA | NA | NA | NA | NA |
| UBL5 | NA | SNAPIN | TST | ALDOA | NA | NA | NA | NA | NA | NA |
| GPR137 | NA | MORF4L1P1 | NDUFA2 | CYC1 | NA | NA | NA | NA | NA | NA |
| SRXN1 | NA | GNL1 | EIF2AK1 | WWP1 | NA | NA | NA | NA | NA | NA |
| RBCK1 | NA | SNORD36B | SHC1 | AP3M1 | NA | NA | NA | NA | NA | NA |
| RANBP3 | NA | TUBG1 | PDRG1 | HAT1 | NA | NA | NA | NA | NA | NA |
| MT-TR | NA | MFSD11 | SP100 | SEC11A | NA | NA | NA | NA | NA | NA |
| AC012640.6 | NA | SLC39A6 | POLR2G | SPAG7 | NA | NA | NA | NA | NA | NA |
| SFT2D1 | NA | MIR4690 | LTV1 | PTPA | NA | NA | NA | NA | NA | NA |
| PAPPA-AS1 | NA | GFM1 | OPA1 | GADD45B | NA | NA | NA | NA | NA | NA |
| FADS2 | NA | SBDS | ARF3 | CAPRIN1 | NA | NA | NA | NA | NA | NA |
| SRRD | NA | S100A6 | VRK3 | MVD | NA | NA | NA | NA | NA | NA |
| BRK1 | NA | PRAF2 | ASCC2 | STAG1 | NA | NA | NA | NA | NA | NA |
| PRRC2B | NA | TIMM23 | RAD17 | GNPTG | NA | NA | NA | NA | NA | NA |
| CTDNEP1 | NA | PCCB | TRIM21 | GUSB | NA | NA | NA | NA | NA | NA |
| PRPSAP1 | NA | NECTIN2 | AC007620.2 | IK | NA | NA | NA | NA | NA | NA |
| MICAL3 | NA | BOD1L1 | RPLP1 | CD47 | NA | NA | NA | NA | NA | NA |
| SF3B1 | NA | ATPAF2 | DTX3L | EIF4H | NA | NA | NA | NA | NA | NA |
| YWHAQ | NA | FAM76B | CHST3 | TRAF7 | NA | NA | NA | NA | NA | NA |
| USP10 | NA | ARPC1A | COQ5 | CYB5R1 | NA | NA | NA | NA | NA | NA |
| RNF185-AS1 | NA | DGAT1 | USP25 | TMEM11 | NA | NA | NA | NA | NA | NA |
| POSTN | NA | GTF2F2 | AC010978.1 | SEC13 | NA | NA | NA | NA | NA | NA |
| NORAD | NA | SLC25A11 | TRAF5 | NUP62 | NA | NA | NA | NA | NA | NA |
| PHB | NA | ERH | POMT2 | TMEM205 | NA | NA | NA | NA | NA | NA |
| ARF3 | NA | HSD17B10 | RSRP1 | ADRM1 | NA | NA | NA | NA | NA | NA |
| CAP1 | NA | L3MBTL3 | TNIP2 | ATG101 | NA | NA | NA | NA | NA | NA |
| AC002985.1 | NA | SNRPG | MT-RNR1 | COASY | NA | NA | NA | NA | NA | NA |
| SURF2 | NA | ACLY | SQSTM1 | ST6GALNAC4 | NA | NA | NA | NA | NA | NA |
| YARS2 | NA | AC008750.8 | PARP14 | MRPL53 | NA | NA | NA | NA | NA | NA |
| BUD31 | NA | PSMG2 | RPLP0P6 | CPED1 | NA | NA | NA | NA | NA | NA |
| RPS29 | NA | JAGN1 | NFKBIA | EIF6 | NA | NA | NA | NA | NA | NA |
| AC115223.1 | NA | MXD4 | SNAPC1 | CDC42SE1 | NA | NA | NA | NA | NA | NA |
| FARP1-AS1 | NA | USP53 | NPTX1 | PSMB7 | NA | NA | NA | NA | NA | NA |
| SGSM3 | NA | MGST1 | CDC42BPB | CMAS | NA | NA | NA | NA | NA | NA |
| DCTPP1 | NA | ANXA2 | NOP58 | HEXA | NA | NA | NA | NA | NA | NA |
| CCDC12 | NA | PTAR1 | AC087393.2 | CAMK2G | NA | NA | NA | NA | NA | NA |
| CYP51A1 | NA | RNF181 | CARHSP1 | BAG6 | NA | NA | NA | NA | NA | NA |
| COX7A2L | NA | STK39 | BCL2L1 | UNC50 | NA | NA | NA | NA | NA | NA |
| AP3B1 | NA | NDUFB9 | C1R | SNF8 | NA | NA | NA | NA | NA | NA |

| | | | | | | | | | | |
|---|---|---|---|---|---|---|---|---|---|---|
| FKBP15 | NA | IER2 | OSGIN1 | KARS1 | NA | NA | NA | NA | NA | NA |
| RPL27A | NA | HCFC1 | COQ4 | GSTO1 | NA | NA | NA | NA | NA | NA |
| BCL2L2-PABPI | NA | NA | AC012615.3 | RSPRY1 | NA | NA | NA | NA | NA | NA |
| CASP3 | NA | NA | FAU | LANCL1 | NA | NA | NA | NA | NA | NA |
| AC104758.1 | NA | NA | RPSA | MIEN1 | NA | NA | NA | NA | NA | NA |
| TAF6 | NA | NA | AC005306.1 | TBC1D17 | NA | NA | NA | NA | NA | NA |
| KSR1 | NA | NA | AL034430.2 | TMEM208 | NA | NA | NA | NA | NA | NA |
| CPD | NA | NA | GPR137 | FYN | NA | NA | NA | NA | NA | NA |
| TPD52L2 | NA | NA | TNFRSF10D | DPM2 | NA | NA | NA | NA | NA | NA |
| ANXA1 | NA | NA | DMPK | SNTB2 | NA | NA | NA | NA | NA | NA |
| MT-TE | NA | NA | BCCIP | UBE3C | NA | NA | NA | NA | NA | NA |
| NCOR1 | NA | NA | FAM102A | MKI67 | NA | NA | NA | NA | NA | NA |
| CNOT7 | NA | NA | CXCL1 | TPP1 | NA | NA | NA | NA | NA | NA |
| AC010776.2 | NA | NA | DDX58 | TMED9 | NA | NA | NA | NA | NA | NA |
| LMAN2 | NA | NA | IER3 | KANSL2 | NA | NA | NA | NA | NA | NA |
| AL451085.3 | NA | NA | BANF1 | SEM1 | NA | NA | NA | NA | NA | NA |
| PFN1 | NA | NA | RRS1 | ARID5B | NA | NA | NA | NA | NA | NA |
| TXNRD1 | NA | NA | TP53I3 | SNAPIN | NA | NA | NA | NA | NA | NA |
| RPSA | NA | NA | STAT3 | TWF1 | NA | NA | NA | NA | NA | NA |
| SLC20A1 | NA | NA | CHFR | NPC2 | NA | NA | NA | NA | NA | NA |
| SCCPDH | NA | NA | BRD7 | PLEKHM2 | NA | NA | NA | NA | NA | NA |
| UQCRC1 | NA | NA | MAPK8IP3 | MAN2A1 | NA | NA | NA | NA | NA | NA |
| PROCR | NA | NA | RPS7 | TLE5 | NA | NA | NA | NA | NA | NA |
| DPAGT1 | NA | NA | AC013489.3 | PSAP | NA | NA | NA | NA | NA | NA |
| MEG3 | NA | NA | CXCL8 | HDLBP | NA | NA | NA | NA | NA | NA |
| UBR2 | NA | NA | CYTOR | UBAP1 | NA | NA | NA | NA | NA | NA |
| MICU2 | NA | NA | CNBP | MRPS18A | NA | NA | NA | NA | NA | NA |
| KIRREL3 | NA | NA | SRSF1 | GFM1 | NA | NA | NA | NA | NA | NA |
| WNT5A-AS1 | NA | NA | TSR1 | IDH1 | NA | NA | NA | NA | NA | NA |
| PCM1 | NA | NA | PTBP2 | USP3 | NA | NA | NA | NA | NA | NA |
| RPL18A | NA | NA | RNF10 | RAB13 | NA | NA | NA | NA | NA | NA |
| FNTA | NA | NA | PPP1R16A | NARF | NA | NA | NA | NA | NA | NA |
| MT-TG | NA | NA | MRM2 | FN1 | NA | NA | NA | NA | NA | NA |
| RPL12P4 | NA | NA | OCRL | IP6K2 | NA | NA | NA | NA | NA | NA |
| RPS7 | NA | NA | EIF1AD | PGK1 | NA | NA | NA | NA | NA | NA |
| ZFAND6 | NA | NA | RBMX | LDHA | NA | NA | NA | NA | NA | NA |
| NDUFS3 | NA | NA | TMEM59L | ANXA5 | NA | NA | NA | NA | NA | NA |
| ATP5MC3 | NA | NA | EFR3A | PEAK1 | NA | NA | NA | NA | NA | NA |
| POP7 | NA | NA | NSMCE1 | YBX3 | NA | NA | NA | NA | NA | NA |
| MRPL14 | NA | NA | AC073861.1 | AGPAT2 | NA | NA | NA | NA | NA | NA |
| HARS1 | NA | NA | SP110 | TOP1MT | NA | NA | NA | NA | NA | NA |
| SSR3 | NA | NA | CETN3 | GABARAPL1 | NA | NA | NA | NA | NA | NA |
| ALCAM | NA | NA | IRF9 | RAD21 | NA | NA | NA | NA | NA | NA |
| RPS15 | NA | NA | IFI27L1 | RPS17 | NA | NA | NA | NA | NA | NA |
| ANKRD17 | NA | NA | CTSA | C11orf98 | NA | NA | NA | NA | NA | NA |
| MBNL1 | NA | NA | EXO1 | FAM214A | NA | NA | NA | NA | NA | NA |
| CCT5 | NA | NA | MDK | PSMA6 | NA | NA | NA | NA | NA | NA |
| SCARF2 | NA | NA | CXCL5 | CBR4 | NA | NA | NA | NA | NA | NA |
| RPS11 | NA | NA | PSIP1 | NSD3 | NA | NA | NA | NA | NA | NA |
| CEP290 | NA | NA | NAGK | CPSF3 | NA | NA | NA | NA | NA | NA |
| HMMR | NA | NA | SNRPC | TPI1 | NA | NA | NA | NA | NA | NA |
| ABHD17C | NA | NA | AL117190.2 | USP20 | NA | NA | NA | NA | NA | NA |
| SERPINE1 | NA | NA | CXCL3 | RBBP8 | NA | NA | NA | NA | NA | NA |
| AP001107.6 | NA | NA | ICAM1 | LSS | NA | NA | NA | NA | NA | NA |
| KLF9 | NA | NA | RHEB | PLD3 | NA | NA | NA | NA | NA | NA |
| MRPS28 | NA | NA | MRPL32 | PLOD1 | NA | NA | NA | NA | NA | NA |
| PGM2 | NA | NA | TGFBR2 | CLP1 | NA | NA | NA | NA | NA | NA |
| MAD2L2 | NA | NA | AC098614.3 | DGKZ | NA | NA | NA | NA | NA | NA |
| HIF1A-AS1 | NA | NA | TCTEX1D2 | PPP1R12A | NA | NA | NA | NA | NA | NA |
| SMARCD3 | NA | NA | CXCL2 | CLPTM1L | NA | NA | NA | NA | NA | NA |
| NKAPD1 | NA | NA | NACC2 | COX6B1 | NA | NA | NA | NA | NA | NA |
| ENC1 | NA | NA | STAU1 | SF1 | NA | NA | NA | NA | NA | NA |
| CD55 | NA | NA | NA | TMEM115 | NA | NA | NA | NA | NA | NA |
| WBP4 | NA | NA | NA | NR1H2 | NA | NA | NA | NA | NA | NA |
| CRAMP1 | NA | NA | NA | NDUFB9 | NA | NA | NA | NA | NA | NA |
| TMED9 | NA | NA | NA | EIF2AK1 | NA | NA | NA | NA | NA | NA |

| | | | | | | | | | | |
|---|---|---|---|---|---|---|---|---|---|---|
| IKBIP | NA | NA | NA | TOP2A | NA | NA | NA | NA | NA | NA |
| SNHG6 | NA | NA | NA | MICOS10 | NA | NA | NA | NA | NA | NA |
| AC016739.1 | NA | NA | NA | CHKA | NA | NA | NA | NA | NA | NA |
| TMEM123 | NA | NA | NA | POLR2G | NA | NA | NA | NA | NA | NA |
| RPL24 | NA | NA | NA | AC016876.3 | NA | NA | NA | NA | NA | NA |
| PRMT1 | NA | NA | NA | PTAR1 | NA | NA | NA | NA | NA | NA |
| NDUFB8 | NA | NA | NA | PSMB6 | NA | NA | NA | NA | NA | NA |
| KRT10 | NA | NA | NA | TRAK2 | NA | NA | NA | NA | NA | NA |
| AC092069.1 | NA | NA | NA | FKBP15 | NA | NA | NA | NA | NA | NA |
| EIF4G3 | NA | NA | NA | TARDBP | NA | NA | NA | NA | NA | NA |
| MRPS24 | NA | NA | NA | ALCAM | NA | NA | NA | NA | NA | NA |
| SMC5 | NA | NA | NA | MLLT6 | NA | NA | NA | NA | NA | NA |
| DDX42 | NA | NA | NA | SRRD | NA | NA | NA | NA | NA | NA |
| RPL34 | NA | NA | NA | RABGGTA | NA | NA | NA | NA | NA | NA |
| RPSAP19 | NA | NA | NA | MAP2K7 | NA | NA | NA | NA | NA | NA |
| MVD | NA | NA | NA | GDE1 | NA | NA | NA | NA | NA | NA |
| TOP3B | NA | NA | NA | ASXL1 | NA | NA | NA | NA | NA | NA |
| DNAJC13 | NA | NA | NA | TUBG1 | NA | NA | NA | NA | NA | NA |
| DHX32 | NA | NA | NA | AL022312.1 | NA | NA | NA | NA | NA | NA |
| RPLP2 | NA | NA | NA | SDE2 | NA | NA | NA | NA | NA | NA |
| PLEKHG5 | NA | NA | NA | ITFG1 | NA | NA | NA | NA | NA | NA |
| STK17A | NA | NA | NA | PEX10 | NA | NA | NA | NA | NA | NA |
| CD81 | NA | NA | NA | COQ5 | NA | NA | NA | NA | NA | NA |
| AP3M1 | NA | NA | NA | USP25 | NA | NA | NA | NA | NA | NA |
| DBN1 | NA | NA | NA | CDK11B | NA | NA | NA | NA | NA | NA |
| AC006001.4 | NA | NA | NA | ALAS1 | NA | NA | NA | NA | NA | NA |
| ATG10 | NA | NA | NA | YY1AP1 | NA | NA | NA | NA | NA | NA |
| AC104411.1 | NA | NA | NA | BRD8 | NA | NA | NA | NA | NA | NA |
| RING1 | NA | NA | NA | MPV17 | NA | NA | NA | NA | NA | NA |
| PABPC1 | NA | NA | NA | SNX3 | NA | NA | NA | NA | NA | NA |
| PTPN23 | NA | NA | NA | RNF181 | NA | NA | NA | NA | NA | NA |
| CUEDC1 | NA | NA | NA | LRPAP1 | NA | NA | NA | NA | NA | NA |
| PSMA2 | NA | NA | NA | AAR2 | NA | NA | NA | NA | NA | NA |
| CDC26 | NA | NA | NA | GNPDA1 | NA | NA | NA | NA | NA | NA |
| LURAP1L | NA | NA | NA | TTC27 | NA | NA | NA | NA | NA | NA |
| AL354811.1 | NA | NA | NA | ASB6 | NA | NA | NA | NA | NA | NA |
| UBL4A | NA | NA | NA | GMPPA | NA | NA | NA | NA | NA | NA |
| BAD | NA | NA | NA | ATF4 | NA | NA | NA | NA | NA | NA |
| CEP164 | NA | NA | NA | ACO2 | NA | NA | NA | NA | NA | NA |
| INPP5K | NA | NA | NA | MAPK6 | NA | NA | NA | NA | NA | NA |
| ELOF1 | NA | NA | NA | EMC8 | NA | NA | NA | NA | NA | NA |
| XRCC1 | NA | NA | NA | FUT11 | NA | NA | NA | NA | NA | NA |
| DMTF1 | NA | NA | NA | PHF1 | NA | NA | NA | NA | NA | NA |
| CDC42SE1 | NA | NA | NA | CNOT6 | NA | NA | NA | NA | NA | NA |
| ACAP3 | NA | NA | NA | PRKAA1 | NA | NA | NA | NA | NA | NA |
| FBLIM1 | NA | NA | NA | HTATIP2 | NA | NA | NA | NA | NA | NA |
| DHCR7 | NA | NA | NA | FBXL17 | NA | NA | NA | NA | NA | NA |
| IFT22 | NA | NA | NA | SBDS | NA | NA | NA | NA | NA | NA |
| SHLD2 | NA | NA | NA | UQCRH | NA | NA | NA | NA | NA | NA |
| AC073861.1 | NA | NA | NA | PLD2 | NA | NA | NA | NA | NA | NA |
| SEC61B | NA | NA | NA | PCTP | NA | NA | NA | NA | NA | NA |
| NLE1 | NA | NA | NA | PRDX5 | NA | NA | NA | NA | NA | NA |
| KIAA0232 | NA | NA | NA | PIGT | NA | NA | NA | NA | NA | NA |
| RSPRY1 | NA | NA | NA | CLCN7 | NA | NA | NA | NA | NA | NA |
| FRMD8 | NA | NA | NA | MYBL1 | NA | NA | NA | NA | NA | NA |
| CD109 | NA | NA | NA | AL365203.2 | NA | NA | NA | NA | NA | NA |
| PSMA6 | NA | NA | NA | OAF | NA | NA | NA | NA | NA | NA |
| PACS1 | NA | NA | NA | SMURF2 | NA | NA | NA | NA | NA | NA |
| PCDHGA12 | NA | NA | NA | SRI | NA | NA | NA | NA | NA | NA |
| CDC123 | NA | NA | NA | DMWD | NA | NA | NA | NA | NA | NA |
| BTBD6 | NA | NA | NA | PHKB | NA | NA | NA | NA | NA | NA |
| SLC3A2 | NA | NA | NA | PLCB3 | NA | NA | NA | NA | NA | NA |
| MAPK3 | NA | NA | NA | NOL3 | NA | NA | NA | NA | NA | NA |
| RNF123 | NA | NA | NA | MAN1B1 | NA | NA | NA | NA | NA | NA |
| RAB11B | NA | NA | NA | CDC42BPB | NA | NA | NA | NA | NA | NA |
| PDCD2 | NA | NA | NA | RELA | NA | NA | NA | NA | NA | NA |

| | | | | | | | | | | |
|---|---|---|---|---|---|---|---|---|---|---|
| TRIM56 | NA | NA | NA | CNPY4 | NA | NA | NA | NA | NA | NA |
| TRABD2A | NA | NA | NA | MYOF | NA | NA | NA | NA | NA | NA |
| PGM1 | NA | NA | NA | TCEA2 | NA | NA | NA | NA | NA | NA |
| UQCRH | NA | NA | NA | LINC00106 | NA | NA | NA | NA | NA | NA |
| AC068888.1 | NA | NA | NA | HPRT1 | NA | NA | NA | NA | NA | NA |
| MEIS1 | NA | NA | NA | E4F1 | NA | NA | NA | NA | NA | NA |
| ABCD4 | NA | NA | NA | NETO2 | NA | NA | NA | NA | NA | NA |
| AHCY | NA | NA | NA | HSD17B10 | NA | NA | NA | NA | NA | NA |
| CENPU | NA | NA | NA | LEMD3 | NA | NA | NA | NA | NA | NA |
| COL4A2-AS1 | NA | NA | NA | PSMB3 | NA | NA | NA | NA | NA | NA |
| FAM13A | NA | NA | NA | TSPYL1 | NA | NA | NA | NA | NA | NA |
| GNG5 | NA | NA | NA | UBL3 | NA | NA | NA | NA | NA | NA |
| CTNNAL1 | NA | NA | NA | YIF1B | NA | NA | NA | NA | NA | NA |
| SNAPIN | NA | NA | NA | STK39 | NA | NA | NA | NA | NA | NA |
| AC016596.3 | NA | NA | NA | COPS3 | NA | NA | NA | NA | NA | NA |
| AC243964.4 | NA | NA | NA | ARGLU1 | NA | NA | NA | NA | NA | NA |
| GNL1 | NA | NA | NA | DHX9 | NA | NA | NA | NA | NA | NA |
| BUD23 | NA | NA | NA | COPB2 | NA | NA | NA | NA | NA | NA |
| AP000926.2 | NA | NA | NA | ATP5MPL | NA | NA | NA | NA | NA | NA |
| SOX9 | NA | NA | NA | PSMA2 | NA | NA | NA | NA | NA | NA |
| PTPRU | NA | NA | NA | AHCY | NA | NA | NA | NA | NA | NA |
| HS1BP3 | NA | NA | NA | MBIP | NA | NA | NA | NA | NA | NA |
| ROCK2 | NA | NA | NA | KLC4 | NA | NA | NA | NA | NA | NA |
| AC025370.1 | NA | NA | NA | AL034430.2 | NA | NA | NA | NA | NA | NA |
| ITPA | NA | NA | NA | WHRN | NA | NA | NA | NA | NA | NA |
| TUBG1 | NA | NA | NA | ANXA7 | NA | NA | NA | NA | NA | NA |
| ALYREF | NA | NA | NA | ANXA4 | NA | NA | NA | NA | NA | NA |
| AC084024.3 | NA | NA | NA | CRTC2 | NA | NA | NA | NA | NA | NA |
| SRA1 | NA | NA | NA | KRR1 | NA | NA | NA | NA | NA | NA |
| GFM1 | NA | NA | NA | TNFRSF10D | NA | NA | NA | NA | NA | NA |
| SBDS | NA | NA | NA | TRIM59 | NA | NA | NA | NA | NA | NA |
| AP002383.2 | NA | NA | NA | URM1 | NA | NA | NA | NA | NA | NA |
| CXCL1 | NA | NA | NA | DMPK | NA | NA | NA | NA | NA | NA |
| RCN2 | NA | NA | NA | SLC9A3R1 | NA | NA | NA | NA | NA | NA |
| SMARCA5 | NA | NA | NA | MAU2 | NA | NA | NA | NA | NA | NA |
| AC022167.1 | NA | NA | NA | ETHE1 | NA | NA | NA | NA | NA | NA |
| MTX1P1 | NA | NA | NA | PGAM1 | NA | NA | NA | NA | NA | NA |
| RPL23A | NA | NA | NA | PSMD8 | NA | NA | NA | NA | NA | NA |
| RPLP1 | NA | NA | NA | OXR1 | NA | NA | NA | NA | NA | NA |
| TSPAN9 | NA | NA | NA | TBC1D10A | NA | NA | NA | NA | NA | NA |
| MIR570 | NA | NA | NA | NDUFA6 | NA | NA | NA | NA | NA | NA |
| AL110504.1 | NA | NA | NA | TP53I3 | NA | NA | NA | NA | NA | NA |
| CCDC124 | NA | NA | NA | SNX1 | NA | NA | NA | NA | NA | NA |
| FMR1 | NA | NA | NA | GABARAP | NA | NA | NA | NA | NA | NA |
| ATPAF2 | NA | NA | NA | GRK2 | NA | NA | NA | NA | NA | NA |
| FDX2 | NA | NA | NA | LINC00623 | NA | NA | NA | NA | NA | NA |
| GATAD1 | NA | NA | NA | LTBP4 | NA | NA | NA | NA | NA | NA |
| AVPI1 | NA | NA | NA | AL031284.1 | NA | NA | NA | NA | NA | NA |
| P3H1 | NA | NA | NA | PIN1 | NA | NA | NA | NA | NA | NA |
| PDCD1LG2 | NA | NA | NA | CBX4 | NA | NA | NA | NA | NA | NA |
| SIRT2 | NA | NA | NA | CHFR | NA | NA | NA | NA | NA | NA |
| GLYR1 | NA | NA | NA | NEU1 | NA | NA | NA | NA | NA | NA |
| AFAP1-AS1 | NA | NA | NA | BRD7 | NA | NA | NA | NA | NA | NA |
| DDX50 | NA | NA | NA | CKS2 | NA | NA | NA | NA | NA | NA |
| AC006441.3 | NA | NA | NA | SMARCD3 | NA | NA | NA | NA | NA | NA |
| GTF2F2 | NA | NA | NA | DYNLT3 | NA | NA | NA | NA | NA | NA |
| REX1BD | NA | NA | NA | CXCL8 | NA | NA | NA | NA | NA | NA |
| SOD2 | NA | NA | NA | SRRT | NA | NA | NA | NA | NA | NA |
| SLC25A11 | NA | NA | NA | TAGLN2 | NA | NA | NA | NA | NA | NA |
| ERH | NA | NA | NA | MRPS10 | NA | NA | NA | NA | NA | NA |
| TWF1P1 | NA | NA | NA | COA6-AS1 | NA | NA | NA | NA | NA | NA |
| AREG | NA | NA | NA | CYTOR | NA | NA | NA | NA | NA | NA |
| NUCB2 | NA | NA | NA | POLE3 | NA | NA | NA | NA | NA | NA |
| HSD17B10 | NA | NA | NA | TMEM181 | NA | NA | NA | NA | NA | NA |
| RPL5 | NA | NA | NA | CALCOCO2 | NA | NA | NA | NA | NA | NA |
| SNRPG | NA | NA | NA | PNRC1 | NA | NA | NA | NA | NA | NA |

| | | | | | | | | | | |
|---|---|---|---|---|---|---|---|---|---|---|
| RPL3 | NA | NA | NA | PRPF39 | NA | NA | NA | NA | NA | NA |
| FASTK | NA | NA | NA | SEC61B | NA | NA | NA | NA | NA | NA |
| SUMO2 | NA | NA | NA | XPC | NA | NA | NA | NA | NA | NA |
| PI4K2B | NA | NA | NA | PRPF38B | NA | NA | NA | NA | NA | NA |
| JAGN1 | NA | NA | NA | CHID1 | NA | NA | NA | NA | NA | NA |
| TST | NA | NA | NA | DDX21 | NA | NA | NA | NA | NA | NA |
| FDFT1 | NA | NA | NA | SLC43A3 | NA | NA | NA | NA | NA | NA |
| ZEB2 | NA | NA | NA | IDH3G | NA | NA | NA | NA | NA | NA |
| USP53 | NA | NA | NA | MRM2 | NA | NA | NA | NA | NA | NA |
| USP16 | NA | NA | NA | TNPO1 | NA | NA | NA | NA | NA | NA |
| TMEM183A | NA | NA | NA | ETFA | NA | NA | NA | NA | NA | NA |
| FADD | NA | NA | NA | ATXN10 | NA | NA | NA | NA | NA | NA |
| CDK2AP2 | NA | NA | NA | QTRT2 | NA | NA | NA | NA | NA | NA |
| STOML2 | NA | NA | NA | POLR1D | NA | NA | NA | NA | NA | NA |
| CRLF1 | NA | NA | NA | CCT5 | NA | NA | NA | NA | NA | NA |
| RPL27 | NA | NA | NA | EIF1AD | NA | NA | NA | NA | NA | NA |
| APOLD1 | NA | NA | NA | TMED7-TICAM | NA | NA | NA | NA | NA | NA |
| CTNNB1 | NA | NA | NA | BECN1 | NA | NA | NA | NA | NA | NA |
| CALM2P2 | NA | NA | NA | G6PC3 | NA | NA | NA | NA | NA | NA |
| RPL23AP42 | NA | NA | NA | NDUFS2 | NA | NA | NA | NA | NA | NA |
| PTAR1 | NA | NA | NA | STYXL1 | NA | NA | NA | NA | NA | NA |
| RNF181 | NA | NA | NA | PDGFD | NA | NA | NA | NA | NA | NA |
| CKAP5 | NA | NA | NA | SLC39A7 | NA | NA | NA | NA | NA | NA |
| JSRP1 | NA | NA | NA | ADAM9 | NA | NA | NA | NA | NA | NA |
| AKR1C1 | NA | NA | NA | VAC14 | NA | NA | NA | NA | NA | NA |
| STK39 | NA | NA | NA | KDM2B | NA | NA | NA | NA | NA | NA |
| KLHDC3 | NA | NA | NA | WAC | NA | NA | NA | NA | NA | NA |
| NF1 | NA | NA | NA | THOC6 | NA | NA | NA | NA | NA | NA |
| RANBP2 | NA | NA | NA | CSNK1G2 | NA | NA | NA | NA | NA | NA |
| NDUFB2 | NA | NA | NA | SMIM15 | NA | NA | NA | NA | NA | NA |
| UBA52 | NA | NA | NA | SP110 | NA | NA | NA | NA | NA | NA |
| NDUFB9 | NA | NA | NA | TYMS | NA | NA | NA | NA | NA | NA |
| STXBP3 | NA | NA | NA | UQCRC1 | NA | NA | NA | NA | NA | NA |
| SSR2 | NA | NA | NA | PGM2 | NA | NA | NA | NA | NA | NA |
| HMBOX1 | NA | NA | NA | BAK1 | NA | NA | NA | NA | NA | NA |
| SNORA4 | NA | NA | NA | SKI | NA | NA | NA | NA | NA | NA |
| EIF5A | NA | NA | NA | SMAD3 | NA | NA | NA | NA | NA | NA |
| GPR89A | NA | NA | NA | SLC25A11 | NA | NA | NA | NA | NA | NA |
| IRF3 | NA | NA | NA | MAP1S | NA | NA | NA | NA | NA | NA |
| MT-ATP6 | NA | NA | NA | MDH2 | NA | NA | NA | NA | NA | NA |
| CCAR1 | NA | NA | NA | SIL1 | NA | NA | NA | NA | NA | NA |
| IER2 | NA | NA | NA | LPCAT3 | NA | NA | NA | NA | NA | NA |
| RNF25 | NA | NA | NA | N4BP2L2 | NA | NA | NA | NA | NA | NA |
| FGF5 | NA | NA | NA | CFAP410 | NA | NA | NA | NA | NA | NA |
| TMEM87A | NA | NA | NA | CNP | NA | NA | NA | NA | NA | NA |
| MT-TQ | NA | NA | NA | NDUFV2 | NA | NA | NA | NA | NA | NA |
| AL353689.2 | NA | NA | NA | PPWD1 | NA | NA | NA | NA | NA | NA |
| SNORD14C | NA | NA | NA | SFSWAP | NA | NA | NA | NA | NA | NA |
| NA | NA | NA | NA | PPP1R15A | NA | NA | NA | NA | NA | NA |
| NA | NA | NA | NA | BLVRA | NA | NA | NA | NA | NA | NA |
| NA | NA | NA | NA | SCD | NA | NA | NA | NA | NA | NA |
| NA | NA | NA | NA | FAM234A | NA | NA | NA | NA | NA | NA |
| NA | NA | NA | NA | VKORC1 | NA | NA | NA | NA | NA | NA |
| NA | NA | NA | NA | ATP5MF | NA | NA | NA | NA | NA | NA |
| NA | NA | NA | NA | RHEB | NA | NA | NA | NA | NA | NA |
| NA | NA | NA | NA | NDUFB5 | NA | NA | NA | NA | NA | NA |
| NA | NA | NA | NA | OGFR | NA | NA | NA | NA | NA | NA |
| NA | NA | NA | NA | GART | NA | NA | NA | NA | NA | NA |
| NA | NA | NA | NA | DCUN1D4 | NA | NA | NA | NA | NA | NA |
| NA | NA | NA | NA | SLC25A5 | NA | NA | NA | NA | NA | NA |
| NA | NA | NA | NA | RHOF | NA | NA | NA | NA | NA | NA |
| NA | NA | NA | NA | COX5A | NA | NA | NA | NA | NA | NA |
| NA | NA | NA | NA | ARHGAP5 | NA | NA | NA | NA | NA | NA |
| NA | NA | NA | NA | TOB2 | NA | NA | NA | NA | NA | NA |
| NA | NA | NA | NA | SURF1 | NA | NA | NA | NA | NA | NA |
| NA | NA | NA | NA | MRPL18 | NA | NA | NA | NA | NA | NA |

| | | | | | | | | | | |
|---|---|---|---|---|---|---|---|---|---|---|
| NA | NA | NA | NA | JAGN1 | NA | NA | NA | NA | NA | NA |
| NA | NA | NA | NA | ATPAF2 | NA | NA | NA | NA | NA | NA |
| NA | NA | NA | NA | SH3BP5 | NA | NA | NA | NA | NA | NA |
| NA | NA | NA | NA | NEK7 | NA | NA | NA | NA | NA | NA |
| NA | NA | NA | NA | TSG101 | NA | NA | NA | NA | NA | NA |
| NA | NA | NA | NA | APMAP | NA | NA | NA | NA | NA | NA |
| NA | NA | NA | NA | GRINA | NA | NA | NA | NA | NA | NA |
| NA | NA | NA | NA | SLC35B1 | NA | NA | NA | NA | NA | NA |
| NA | NA | NA | NA | DCTN6 | NA | NA | NA | NA | NA | NA |
| NA | NA | NA | NA | DIP2A | NA | NA | NA | NA | NA | NA |
| NA | NA | NA | NA | RPRD2 | NA | NA | NA | NA | NA | NA |
| NA | NA | NA | NA | SNRPG | NA | NA | NA | NA | NA | NA |
| NA | NA | NA | NA | S100A10 | NA | NA | NA | NA | NA | NA |
| NA | NA | NA | NA | SLK | NA | NA | NA | NA | NA | NA |
| NA | NA | NA | NA | NACC2 | NA | NA | NA | NA | NA | NA |
| NA | NA | NA | NA | PRR11 | NA | NA | NA | NA | NA | NA |
| NA | NA | NA | NA | SRPRA | NA | NA | NA | NA | NA | NA |
| NA | NA | NA | NA | DNAJB14 | NA | NA | NA | NA | NA | NA |
| NA | NA | NA | NA | ATP5MC3 | NA | NA | NA | NA | NA | NA |
| NA | NA | NA | NA | STAU1 | NA | NA | NA | NA | NA | NA |
| NA | NA | NA | NA | WDR13 | NA | NA | NA | NA | NA | NA |
| NA | NA | NA | NA | USP53 | NA | NA | NA | NA | NA | NA |

**GEO Accessions of the Samples**

| GSE56192_MERS-INFECTED | GSE56192_MERS-MOCK | GSE56192_SARS-INFECTED | GSE56192_SARS-MOCK | PBMC-INFECTED |
|---|---|---|---|---|
| SRR1192413 | SRR1190365 | SRR1193006 | SRR1192354 | CRR119890 |
| SRR1192414 | SRR1190366 | SRR1193007 | SRR1192370 | CRR125445 |
| SRR1192415 | SRR1190367 | SRR1193008 | SRR1192371 | CRR125446 |
| SRR1192416 | SRR1190368 | SRR1193009 | SRR1192398 | |
| SRR1192417 | SRR1190369 | SRR1193010 | SRR1192399 | |
| SRR1192418 | SRR1190370 | SRR1193011 | SRR1192419 | |
| SRR1192419 | SRR1190371 | SRR1193012 | SRR1192464 | |
| SRR1192420 | SRR1190372 | SRR1193013 | SRR1192465 | |
| SRR1192464 | SRR1191342 | SRR1193014 | SRR1192471 | |
| SRR1192465 | SRR1191343 | SRR1193015 | SRR1192496 | |
| SRR1192466 | SRR1191344 | SRR1193016 | SRR1192497 | |
| SRR1192467 | SRR1191345 | SRR1193017 | SRR1192498 | |
| SRR1192468 | SRR1191346 | SRR1193018 | SRR1192499 | |
| SRR1192469 | SRR1191347 | SRR1193100 | | |
| SRR1192470 | SRR1191348 | SRR1193101 | | |
| SRR1192471 | SRR1191349 | SRR1193102 | | |
| | | SRR1193103 | | |
| | | SRR1193104 | | |
| | | SRR1193105 | | |
| | | SRR1193106 | | |
| | | SRR1193197 | | |
| | | SRR1193198 | | |
| | | SRR1193199 | | |
| | | SRR1193403 | | |
| | | SRR1193404 | | |
| | | SRR1193405 | | |
| | | SRR1193522 | | |
| | | SRR1193523 | | |
| | | SRR1193524 | | |
| | | SRR1193638 | | |
| | | SRR1195619 | | |
| | | SRR1195620 | | |
| | | SRR1195621 | | |

| PBMC-MOCK | GSE147507_SARS-CoV-2-INFECTED | GSE147507_SARS-CoV-2-MOCK |
|---|---|---|
| CRR119891 | SRR11412251 | SRR11412239 |
| CRR119892 | SRR11412252 | SRR11412240 |
| CRR119893 | SRR11412253 | SRR11412241 |
| | SRR11412254 | SRR11412242 |
| | SRR11412255 | SRR11412243 |
| | SRR11412256 | SRR11412244 |
| | SRR11412257 | SRR11412245 |
| | SRR11412258 | SRR11412246 |
| | SRR11412259 | SRR11412247 |
| | SRR11412260 | SRR11412248 |
| | SRR11412261 | SRR11412249 |
| | SRR11412262 | SRR11412250 |
